# Non-target Analysis of Wastewater Treatment Plant Effluents: Chemical Fingerprinting as a Monitoring Tool

**DOI:** 10.1101/2023.08.03.551870

**Authors:** Marie Rønne Aggerbeck, Emil Egede Frøkjær, Anders Johansen, Lea Ellegaard-Jensen, Lars Hestbjerg Hansen, Martin Hansen

## Abstract

This study aims at discovering and characterizing the plethora of xenobiotic substances released into the environment with wastewater effluents. We present a novel non-targeted screening methodology based on ultra-high resolution Orbitrap mass spectrometry and nanoflow ultra-high performance liquid chromatography together with a new data-processing pipeline. This approach was applied to effluent samples from two state-of-the-art urban, and one small rural wastewater treatment facility. In total, 785 structures were obtained, of these 38 were identified as single compounds, while 480 structures were identified at a putative level. The vast majority of these were therapeutics and drugs, present as parent compounds and metabolites. Using the R packages Phyloseq and MetacodeR, we here present a novel way of visualizing LCMS data while showing significant difference in xenobiotic presence in the wastewater effluents between the three sites.

**Significance:** We characterized a wide spectrum of xenobiotic substances using ultra-high performance liquid chromatography, and analysed the data with a new data-processing pipeline using microbial ecological tools to visualize and perform statistical testing of the chemical data to reveal trends in compound composition at the three WWTPs. This approach was applied to obtain and analyse data from effluent samples collected at three wastewater treatment facilities. In total, 785 chemical structures were achieved, with a majority identified as therapeutics and drugs. Several of the compounds are suspected endocrine disruptors. The data reveal a significant difference in compound diversity persisting in the wastewater effluents at the three sites. Our findings reveal the presence of undesirable compounds in effluent released into waterways, and address the greatest challenge in environmental chemistry – pinpointing single compounds of interest from masses of data produced.

## 2. Introduction

Wastewater treatment plants (WWTPs) are essential for removing pollutants from wastewater before it is discharged into waterways. However, current WWTPs are not capable of completely removing all types of xenobiotics^1^, which are synthetic or naturally occurring compounds that can pose risks to the environment and public health. Xenobiotics that are partially or fully degraded can still be released into waterways^2–4^, potentially causing adverse effects such as endocrine disruption^5,6^or antimicrobial resistance^7^. Recent studies have also found recalcitrant xenobiotics in groundwater and drinking water systems^8, 9^, indicating that leaching is a problem in the drinking-water cycle.

To improve the removal of xenobiotics from wastewater, the first step is to assess a vast range of compounds present in the wastewater effluent. However, using a targeted chemical approach can be expensive and challenging due to the sheer number of compounds typically present in wastewater samples^10^.

Non-targeted analysis (NTA) is a promising approach requiring minimal sample preparation (such as dilute-and-shoot^11^ or solid-phase extraction (SPE) capable of capturing a broad chemical space^12–14^) that utilizes high-resolution mass spectrometry (HRMS) and ultra-high performance liquid chromatography (UHPLC) to comprehensively discover xenobiotics in environmental samples^15, 16^. NTA avoids the limitations of a pre-defined list of compounds to search for, but results in large, complex datasets containing both known and unknown compounds that need to be curated^17^ and filtered for statistical analysis.

Despite the increased use of high-resolution mass spectrometry and data-processing software, the computational challenges of analysing large datasets containing thousands of compounds remain. Multivariate non-parametric analyses are needed to unravel relevant trends in the datasets.

While data-treatment tools have been developed to explore complex ecological datasets in biological fields, they remain underexploited in environmental chemistry. Basic statistics tools have become integrated in applications such as Compound Discoverer (Thermo Fischer Scientific), but comprehensive tool packages for extensive exploratory and comparative data analysis remain underdeveloped. Ecological community and non-targeted chemical datasets both consist of matrices of discrete values with object identification and related variables, and can be analyzed using the same statistical methods, allowing the use of already validated, peer-reviewed methods from the field of microbial ecology.

We conducted this study on effluent samples from three WWTPs in the Greater Copenhagen area, ranging in population equivalent (PE) from 12,000 to 400,000 and with different proximity to city, hospitals, industry, and rural areas. By sampling fully treated effluent water immediately before its release, we aimed to identify both parent substances and their degradation metabolites of pharmaceuticals, household chemicals, and a variety of human-related metabolites. We hypothesize recovery a wide range of known environmental pollutants.

The main aim of this case study was to employ state-of-the-art NTA on effluents from three Danish WWTPs, and to develop a robust versatile pipeline for processing non-targeted data in a pollution research context. We also aim to adapt and apply validated statistical approaches developed for microbial community analysis to non-targeted chemical analysis and develop a robust, reproducible data analysis pipeline for screening WWTP effluent samples for both known and unknown xenobiotics. To achieve this, we employed HRMS, developed a data processing pipeline for multivariate chemical compound analysis, and annotated substances using tandem MS spectral library matches. We then used an R-based array of ecological multivariate statistical analysis tools to explore and visualize the between-site dynamics of a vast number of xenobiotics.

Together, these tools will enable a more precise characterization of xenobiotic diversity in wastewater effluent and explore the potential to identify novel and unknown pollutants not targeted by classical analytical chemistry. This paper presents a novel approach to non-targeted sensitive environmental analysis of wastewater effluent, with the aim of investigating the extent of the release of potentially harmful compounds from a treatment plant into the surrounding waterways.

## 3. Results

Following data analysis, we observed 4094 unique substances across the dataset. Of these 1482 were filtered out as background. We applied the tiered identification system proposed by Schymanski et al.^18^, where a level 1 identification represents a confirmed substance with in-house experimental data on retention time and fragmentation spectra, level 2 is obtained using fragmentation data from other libraries, level 3 when no reference fragmentation data is available, while level 4 and 5 are used when only MS1 data was available. Using online tandem MS spectral databases, we confirmed 785 compounds above level 2 (a table of all confirmed compounds are available as Supplementary Table I). Of these, we were able to assign compound classes to 451 compounds. Using in-house libraries, 38 compounds were confirmed to level 1, as listed in table 1. Metabolites of confirmed level 1 substances are listed in Supplementary Metabolite Catalogue 1.

**Table 1.** All compounds confirmed at level 1 annotation. Class, subclass and compound name are stated, along with CAS numbers where possible. Compound presence in samples are given as the relative abundance of group areas for each site, and the number of putative identified metabolites is stated for all compounds significantly differing in abundance between sites (grey background on compound name). Each site is described by population equivalent and community size. Heatmap colouring denotes relative abundance of each compound between sites. Dark blue = high relative abundance; white = low relative abundance. * denote steroids with several unidentified molecules with a steroid-like structure.

| Class/Subclass /Compound Name | CAS | Relative abundance,<br>Group Area, site A<br>*Suburban, PE = 400k* | Relative abundance,<br>Group Area, site D<br>*Urban, PE = 350k* | Relative abundance,<br>Group Area, site S<br>*Rural, PE = 12k* | Metabolites<br>putative<br>identified in<br>dataset* |
| --- | --- | --- | --- | --- | --- |
| **Industrial Chemicals** |  |  |  |  |  |
| **Flame retardant** |  |  |  |  |  |
| Tris(2-butoxyethyl) phosphate | 78-51-3 | 27.6% | 45.3% | 27.0% | N/A |
| **Photoinitiator** |  |  |  |  |  |
| Photoinitiator TPO |  | 50.4% | 41.2% | 8.4% | N/A |
| **Plasticizer** |  |  |  |  |  |
| Tributyl Phosphate | 126-73-8 | 29.8% | 53.9% | 16.3% | 1 |
| **Metabolite** |  |  |  |  |  |
| **Human Metabolites** |  |  |  |  |  |
| Pregnenolone | 145-13-1 | 99.3% | 0.4% | 0.3% | 1* |
| Aldosterone | 52-39-1 | 93.6% | 4.6% | 1.8% | 0* |
| **Personal Care Product** |  |  |  |  |  |
| **Sunscreen** |  |  |  |  |  |
| Octisalate |  | 53.1% | 23.4% | 23.5% | 0 |
| **Pesticides** |  |  |  |  |  |
| **Fungicide** |  |  |  |  |  |
| DEET | 134-62-3 | 46.9% | 32.4% | 20.7% | N/A |
| Propiconazole | 60207-90-1 | 2.5% | 95.2% | 2.2% | 0 |
| Tebuconazole | 107534-96-3 | 36.9% | 29.4% | 33.7% | N/A |
| **Herbicide** |  |  |  |  |  |
| Prometryn | 7287-19-6 | 3.3% | 89.4% | 7.3% | 0 |
| Prosulfocarb | 52888-80-9 | 84.7% | 5.6% | 9.7% | 1 |
| **Reagents and standards** |  |  |  |  |  |
| Dibutyl phosphate | 107-66-4 | 29.6% | 54.2% | 16.2% | N/A |
| **Therapeutics/Drugs** |  |  |  |  |  |
| **Analgesic** |  |  |  |  |  |
| Diclofenac | 15307-86-5 | 40.0% | 34.1% | 25.9% | N/A |
| Indomethacin | 53-86-1 | 94.6% | 4.2% | 1.1% | 0 |
| Naproxen | 22204-53-1 | 39.0% | 39.2% | 21.8% | N/A |
| **Antibiotic** |  |  |  |  |  |
| Azithromycin | 83905-01-5 | 57.7% | 28.3% | 14.0% | N/A |
| Clarithromycin | 81103-11-9 | 49.1% | 45.2% | 5.7% | 1 |
| Roxithromycin | 80214-83-1 | 39.1% | 27.5% | 33.4% | N/A |
| Trimethoprim | 738-70-5 | 36.2% | 34.5% | 29.3% | N/A |
| **Anticonvulsant** |  |  |  |  |  |
| Carbamazepine | 298-46-4 | 33.8% | 32.2% | 34.0% | N/A |
| Oxcarbazepine | 28721-07-5 | 46.9% | 36.4% | 16.6% | 2 |
| **Antihistamine** |  |  |  |  |  |
| Cetirizine | 83881-51-0 | 35.4% | 37.7% | 26.8% | N/A |
| Fexofenadine | 83799-24-0 | 29.6% | 29.4% | 41.0% | N/A |
| **Cancer treatment** |  |  |  |  |  |
| Bicalutamide | 90357-06-5 | 30.8% | 35.3% | 33.8% | N/A |
| **Cardiovascular regulation** |  |  |  |  |  |
| Bisoprolol | 66722-44-9 | 28.1% | 25.2% | 46.7% | N/A |
| Candesartan | 139481-59-7 | 42.5% | 48.7% | 8.8% | 0 |
| Diltiazem | 42399-41-7 | 55.0% | 33.4% | 11.6% | 2 |
| Losartan | 114798-26-4 | 41.3% | 30.8% | 27.9% | N/A |
| Metoprolol | 51384-51-1 | 38.4% | 37.1% | 24.5% | N/A |
| Propranolol | 525-66-6 | 53.8% | 37.0% | 9.2% | 0 |
| Valsartan | 137862-53-4 | 36.3% | 48.1% | 15.6% | 0 |
| **Diuretic** |  |  |  |  |  |
| Furosemide | 54-31-9 | 43.1% | 43.3% | 13.6% | N/A |
| **Psychopharmaca** |  |  |  |  |  |
| Citalopram | 59729-33-8 | 32.6% | 41.1% | 26.4% | N/A |
| Sertraline | 79617-96-2 | 35.3% | 40.4% | 24.3% | N/A |
| Venlafaxine | 93413-69-5 | 44.8% | 27.1% | 28.0% | N/A |
| **Steroid medication** |  |  |  |  |  |
| Prednisone | 53-03-2 | 88.6% | 3.9% | 7.4% | 1* |
| **Wormicide** |  |  |  |  |  |
| Mebendazole | 31431-39-7 | 30.2% | 40.9% | 28.8% | N/A |

Compound Discoverer identified 16 compounds as differing significantly in presence between sites (L2fC > 1; < 0.05). Of these, 10 belonged to the therapeutics and drugs class (e.g. the antibiotic clarithromycin and the anticonvulsant oxcarbazepine), 4 to pesticides (e.g. propiconazole and prosulfocarb), one was a flame retardant (tributyl phosphate), and one was a personal care product, specifically a sunscreen component (2-ethylhexyl salicylate).

The *in silico* fragmentation chemical fingerprint assisted in confirming and finding suspects not otherwise found in the dataset. Without this, the study would have been limited to only annotating compounds found in house or in the used spectral libraries (mzCloud, MassList, ChemSpider). Broadening the search beyond these few sources improves the detection field and improves the likelihood of identifying and annotating compounds, which might otherwise have gone unknown. Using this approach, we succeeded in identifying 12 further metabolites in the dataset. All the compounds with newly identified metabolites were in the therapeutics and drugs class, except for the herbicide prosulfocarb. All metabolites were found in all samples, except for the metabolite prosulfocarb sulfoxide, a metabolite of prosulfocarb, which was found only in site A.

In samples from site A, metabolites and unidentified natural products make up approximately 50% of the compounds in all samples, a higher proportion than in site D and S, where they make up 25-30%. We observed a good agreement across the three replicates.

The largest class, therapeutics and drugs, make up 37 - 45% of the annotated compounds released from each site. Industrial chemicals make up approximately 10% at site D, more than twice as much as site A and S. Ubiquitous amino acids, personal care products, reagents and standards, and textile chemicals/auxiliary dyes make up 5-15% of the compounds across the dataset. However, these fractions of annotations may be skewed by the used databases.

The PERMANOVA comparing the three sites yielded P=0.002, suggesting a significant difference in variance between sites. However, when a pairwise PERMANOVA with p-value correction is applied, the p-values rise to 0.1.

The bray-curtis Principal Coordinates Analysis (PCoA, figure 2A) reveals a distinct clustering of same-site samples, with the pooled (QC) samples clustering in the middle, explaining 91.3% of the variance between axis 1 and 2. This suggests that there is distinct difference between sites, and that samples from the same sites are highly similar. The QC samples clustering around the point of origin suggests corrections for systematic errors, such as instrument signal drift, were achieved with a good performance. Fig 2B shows the clustering patterns of each class. Amino acids and textile chemicals/auxiliary dyes cluster around the point of origin, suggesting an equal distribution and abundance between sites. Metabolites (known and unknown), reagents and standards, and therapeutics and drugs are all spread across the Euclidean space, suggesting no immediate affinity for one or more sites for most of the compounds in the classes. The industrial chemicals and pesticides cluster towards sites A and D, with a larger number of industrial chemicals closer to site A.

**Figure 1.**
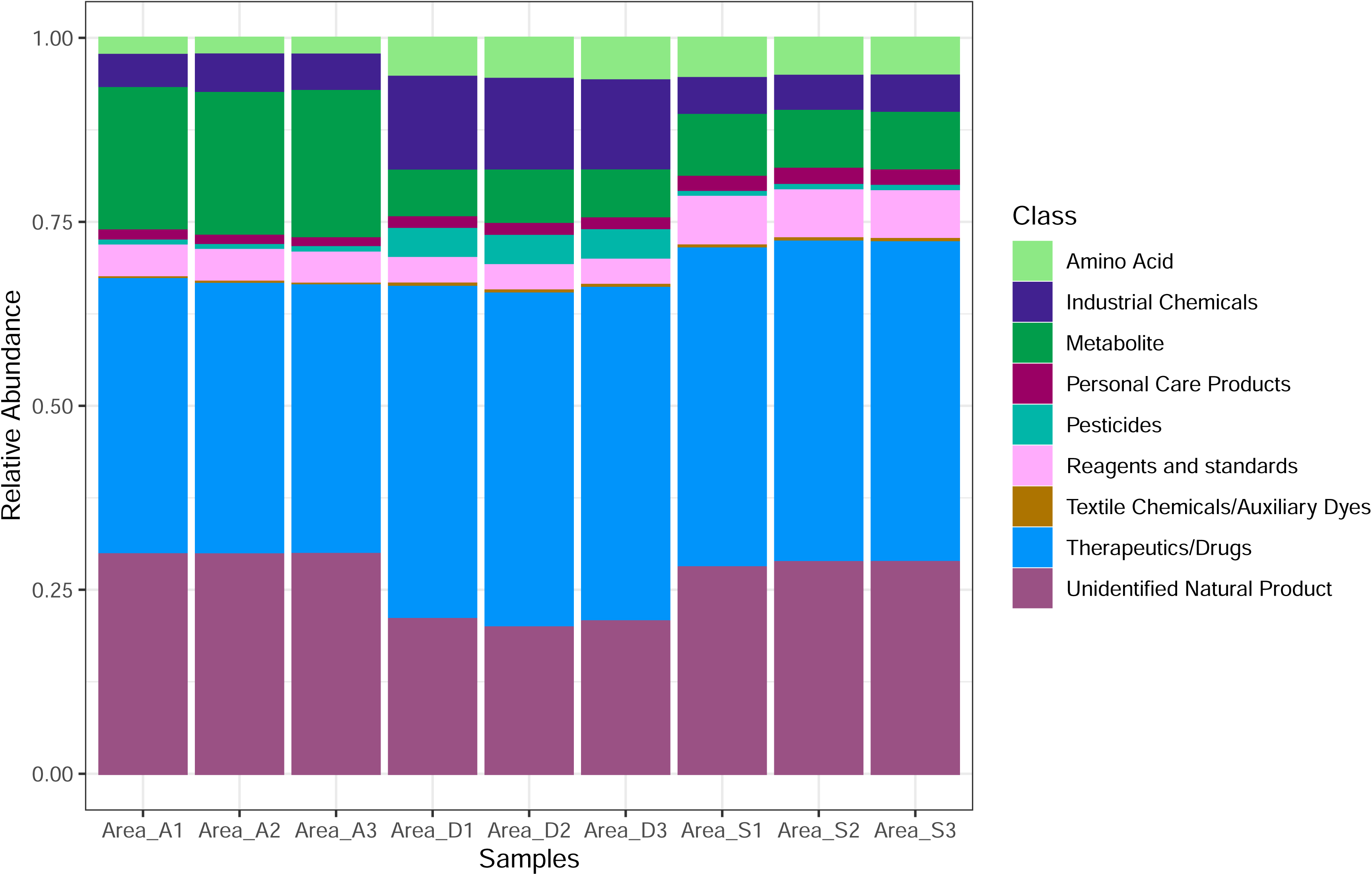
Figure 1 shows the composition of compound area of particular classes, relative to the total number of compounds in the sample. Each bar depicts one single sample, totalling nine samples across three sites. Areas of all identified molecules in each sample are stacked and coloured by compound class.

**Figure 2.**
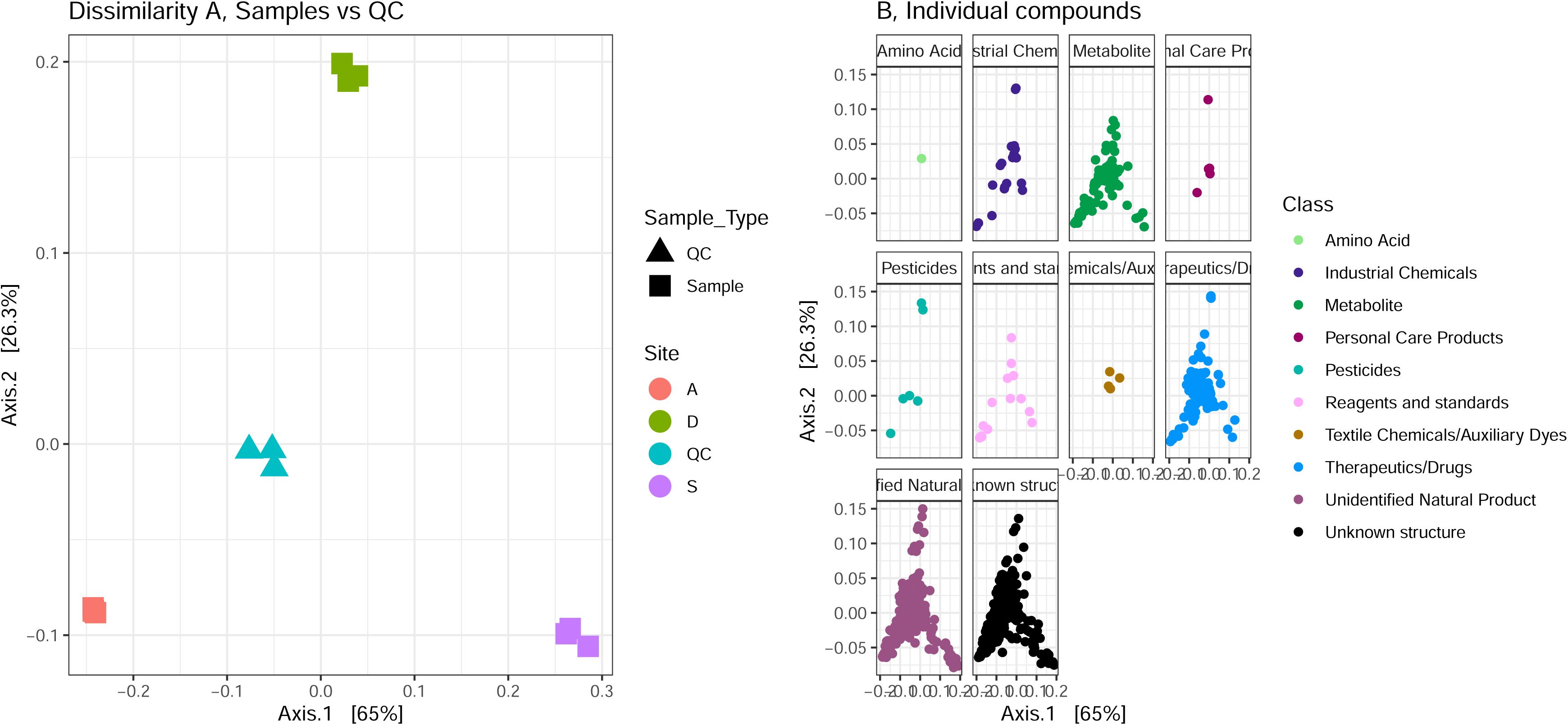
A) shows the principal coordinates plot of all 785 known compounds identified to either level 1 or 2. The square icons show the three sampling sites. The large blue triangles show the placement of the QC pools. B) shows the corresponding loadings plots for each compound class. The number of dots in each class plot corresponds to each individual compound belonging to that class (e.g. 1 dot in “Amino Acid” corresponds to the single compound identified as an amino acid in the overall dataset.)

The heat trees (figure 3) created to visualize single compound variance depict any compounds with a log2 fold-change (LfC) of more than 1 in any pairwise comparison. As both natural and synthetic substances were identified, we divide the following description into these categories to improve legibility:

**Figure 3.**
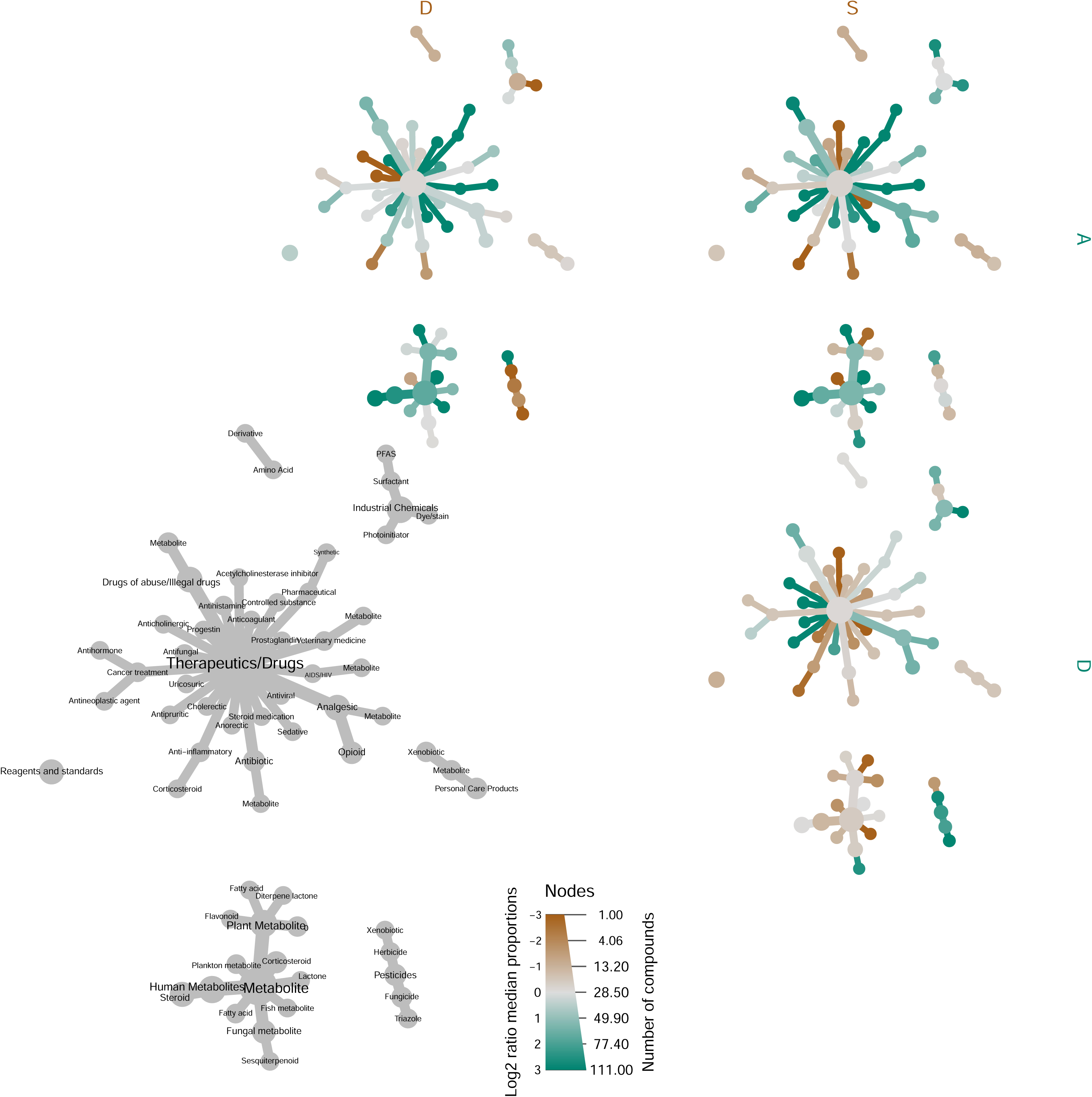
Figure 3 shows a differential heat tree displaying any compound with an LfC >1. Compounds are grouped by class. The grey tree relays compound class names. The coloured trees represent pairwise comparisons between sites. The size of nodes denotes the number of compounds belonging to that specific class or subclass. Coloured nodes represent an upregulation in the site with the same colour label. (e.g. a green node means a larger presence in the site noted in green.)

Natural substances:

We find only marginal difference in amino acid presence between sites, with a higher proportion at site D and site S.

Plant metabolites are more abundant at site S, human metabolites are more abundant at site A. Curiously, we find a number of human endogenous metabolites which remain undegraded at site A, but are less present at the larger site D.

Synthetic substances:

Industrial chemicals are generally more prevalent in sites A and D, which is in line with the site uplands. The dye Fluorescein is more prevalent at site D.

Personal care products are present in waste water effluent across all sites, and mostly in equal abundances at all sites (Table 1), with a few exceptions such as Galaxolidone, a fragrance ingredient most abundant in site D, and the sunscreen compound Enzacamene, which was found to be most abundant in site D. Likewise, pesticides escape degradation in all three plants, but with the largest overall proportion found in site D.

Reagents and standards were ubiquitously found across the dataset, with a few compounds (2-Ethyl-5-methylhexyl benzoate, 9-Azabicyclo[6.2.0]decan-10-one, and Cyclohexyl[(3S)-3-(5-methyl-1H-benzimidazol-2-yl)-1-pyrrolidinyl]methanone) classified as molecular building blocks being more abundant at site S.

Therapeutics and drugs were by far the most abundant group of compounds found in wastewater treatment plant effluent. 114 individual compounds were identified, of which 27 were identified to level 1. Acetylcholinesterase inhibitor was found to be most abundant in site S, along with antivirals, corticosteroids and cancer treatment compounds, as well as metabolites of antibiotics. Oppositely, analgesics are found in larger proportions at site A and D, which is in accordance with the size of the plants. Opioids make up almost half of the compounds found in this subclass. Along this result, a large number of controlled substances were identified in the dataset, as well as metabolites of these. The highest abundance of controlled substances were found in site A.

Antibiotics are found in highest abundances at site S, while antifungal drugs are most abundant in the effluent at site D.

A single anticholerectic drug was found in high abundances at site A and S.

## 4. Discussion

This study aimed to recover and identify the extent of the release of potentially harmful compounds from three wastewater treatment plants (PE 12.000 – 400.000) into the surrounding waterways. We employed a non-targeted analysis (NTA) approach utilizing high-resolution mass spectrometry (HRMS) and ultra-high performance liquid chromatography (UHPLC) to discover xenobiotics in effluent samples.

We observed 4094 unique substances across the dataset. Using the tiered identification system proposed by Schymanski et al., we putatively identified 785 compounds above level 2. Of these, we were able to assign compound classes to 451 compounds. Using in-house libraries, we further identified 38 compounds to level 1 (as listed in Table 1). 16 compounds differed significantly in presence between sites, with 10 af these belonging to the therapeutics and drugs class.

*In silico* fragmentation assisted in confirming and finding suspects not otherwise found in the dataset. The study identified 12 further metabolites in the dataset using this approach, and mostly from the therapeutics and drugs class. As the *in silico* fragmentation prediction relies on the investigated molecular structure, the approach is not hampered by observational bias as seen from mass spectral databases.

Interestingly, the only non-drug metabolite annotated *in silico* (a metabolite of prosulfocarb, prosulfocarb sulfoxide) was also the only metabolite not found across all samples. Given the presented differences between compounds (or chemical fingerprints) occurring in all three treatment plants, there seems to be no consistent compound metabolisation pattern between the observed plants, suggesting compound metabolisation is highly contextual and dependent on the individual plant, as well as sewage source and any infiltration processes.

Our study further tailored a robust, versatile analysis pipeline for processing non-targeted data in a pollution research context, aiming to adapt and apply validated statistical approaches developed for microbial community analysis to non-targeted chemical analysis.

The Non-Targeted Analysis as a descriptive tool relies on presenting sample differences in an understandable way. As such, the challenge lies in distinguishing compounds of interest from the vast amount of available data. To determine significant differences in single compounds between samples, Wilcoxon, Mann-Whitney-U, or similar non-parametric statistics are commonly used, but these tests can be less accurate with large datasets, as the presence of numerous ubiquitous compounds can mask or obscure statistical signals arising from the datapoints of interest, potentially leading to their disappearance. In such a situation, relying solely on p-values is not advisable, so we also report compounds with high log2 fold change (LfC) to address this issue.

Presenting the dataset as heat trees (figure 3) combines the convenient overview of a class-based dendrogram with the log fold scale, highlighting differences in per-compound area between sites that might otherwise have gone unnoticed.

To provide a concise summary of the findings, we here discuss each compound class separately. A number of substances of natural origin were found in the effluent samples.

### Amino acids

We found only a slight difference in the single amino acid presence between sites, with a higher relative proportion of dibenzyl L-aspartate at site D and site S.

Amino acids are the main intermediates from protein degradation^19^, and as higher protein concentrations in wastewater has been linked to eutrophication of waterways receiving effluent^20^, it might be a cause for concern to find excessive amounts of protein, amino acids or nitrogen-rich compounds in effluent. While outside the scope of this paper, measuring the extent of nitrogen release from WWTPs may provide further insight into mechanisms behind nitrification of Danish oceans and waterways.

### Endogenous Metabolites

Metabolites of various origin are to be expected in a wastewater treatment plant, especially with intake sources in closely populated areas. Most of the identified metabolites show some difference in relative abundance between sites.

Plant metabolites are more abundant at site S and D, which may in part be due to algae and fallen plant material being visibly present in the open effluent basin at the sites. The fatty acid Diisobutyl adipate, however, is more abundant at site A. While this compound is of ubiquitous algal origin^21^, its prevalence may also be due to its use in personal care products as an emollient, particularly in sunscreen lotions^22^. Whether this compound should be classified as a plant metabolite or a personal care product remains a subject of debate. However, due to the inability to ascertain its precise origin, we have chosen to assign it to the category that incurs the least ambiguity.

Metabolites found in fish are most prevalent at site A, while fungal sesquiterpenoids are equally common in sites A and D. Sesquiterpeneoids are the largest natural products from fungi^23^, and the metabolite nanangenine B in particular has been shown to exhibit strong cytotoxicity^24^, which might explain its resilience to treatment.

Endogenous human metabolites are more abundant at site A – particularly pregnenolone and aldosterone, both present in significantly higher relative abundance. Site A and D are of similar sizes, and both uplands contain domestic housing, making this result surprising. Using *in silico* fragmentation, a metabolite of pregnenolone, pregn-5-en-20-on-3b-yl sulfurate, was identified at all sites, suggesting partial degradation on site or by *in vivo* metabolisation of the parent compound.

We find a number of endogenous metabolites remaining undegraded at site A, but less at the larger site D. This could suggest more efficient degradation at site D, but is mostly likely an inflated result brought on by the sampling time: Endogenous steroids degrade at vastly different speeds depending on their structure^25^, but sampling was done in the morning, and site A with its chiefly suburbian upland would as such provide ample amounts of steroids of both endogenous and synthetic origin from the morning routines of the community.

### Synthetic substances

#### Industrial chemicals

Industrial chemicals are generally more prevalent in sites A and D, which matches the density of industrial sites in the corresponding WWTP uplands. Tributyl phosphate (TBP), a commonly used plasticizer, also applied as flame retardant to e.g. cotton fabric, was found in significantly higher relative abundances at site D. This compound is a known endocrine disruptor^26, 27^ and thus releasing it into the Danish seas poses an immediate ecological risk. Tributyl phosphate has shown effective removal efficiency in water resource recovery facilities^28^, so further investigation of the efficacy of site D might be advisable.

The dye Fluorescein is also most prevalent at site D, suggesting a recent application at the treatment plant itself, as it is used as a flow rate marker and the compound is known to photodegrade fast^29^.

Other industrial chemicals such as photoinitiators and surfactants are more prevalent at the larger sites A and D – especially worrying is the compound (2R,3R,4R,5S)-1-{5-[(4’-Fluoro-4-biphenylyl)methoxy]pentyl}-2-(hydroxymethyl)-3,4,5-piperidinetriol, which is classified by Pubchem^30^ as a PFAS. PFAS accumulation is increasingly recognized as a global concern, with Denmark also experiencing its impact and insufficient degradation at treatment plants may contribute to this issue^31^

#### Personal care products

We expected to find personal care products across all sites, and most of the compounds found in this class are present in equal abundances at all sites (Table 1). However, a few compounds differ. For instance the compound galaxolidone, which is the degradation product of a commonly used fragrance ingredient known to persist through wastewater treatment^32^, was most abundant in site D. Multiple fragrance production facilities are located in the upland of site D, which may explain the abundance of this particular compound at that site, while most other commonly used household compounds are found in higher or similar abundances at site A.

The sunscreen compound 2-ethylhexyl salicylate was found in all three sites but in significantly higher abundances in site A. This compound has recently been banned in several countries due to its endocrine disrupting effects^33, 34^. Another sunscreen compound, enzacamene (4-MBC), was found to be most abundant in site D. This compound has been investigated for its capacity to induce coral bleaching and its potential marine toxicity, as well as its documented endocrine-disrupting effects^35, 36^.

The uplands of both site A and D contain large residential areas made up of apartment buildings and detached houses, as well as parks and allotments, which see heightened use in the summer months. This dataset was collected in August, after a period of sunny weather coinciding with the Danish summer holiday. Sunscreen is likely used liberally during such a period and through personal hygiene practices is transported to the wastewater treatment system. Finding these two compounds undegraded in effluent piped directly into the sea raises further environmental concern. Such a result alone illustrates the use of the NTA as a monitoring tool and its capability to provide a snapshot of pollutants present in waterways.

#### Pesticides

We were surprised to find only a small number of pesticides in the wastewater effluent, as pesticide persistence through wastewater treatment has been reported often^37, 38^. Triazole fungicides were found at all three sites. Propiconazole, a carcinogenic endocrine disruptor outlawed as a pesticide in EU in 2019^39^ and currently under further legal scrutiny as an additive to wood preservatives^40^, was significantly more prevalent at site D. The herbicide prometryn, a broadleaf weedkiller, was also significantly more abundant at site D, while the herbicide prosulfocarb, the second-most used herbicide in Denmark^41^, was significantly more abundant at site A, suggesting a more common use in this area. As prosulfocarb is an agricultural herbicide, it is consistent with expectations to find it present only in a suburban setting. Concerns have been raised that pesticides droplets sprayed on fields may become airborne and be transported to adjacent neighborhoods, ultimately leading to their presence in wastewater^42^. This spray drift might account for a higher presence of pesticides in suburban site A, despite the distance of the upland to agricultural fields. A degradation product, prosulfocarb sulfoxide, was identified using the *in silico* fragmentation method.

Prosulfocarb sulfoxide was only recovered from site A, which is concordant with the parent compound abundance. This suggests partial on-site degradation of the pesticide. Pesticides were found in surprisingly low abundances at rural site S – given its location and minute size, we hypothesized a large number of pesticides escaping treatment plant degradation, in abundances greater than the urban plants. This could suggest efficient treatment at the site, but it is unfortunately more likely an effect of low intake - pesticides used in the upland likely escape the treatment plant entirely and leach into soil surrounding farms and houses, or wash directly into waterways via overflow systems.

#### Reagents and standards

Molecular building blocks (reagents and standards) were identified at all three sites, with a slightly overall relative abundance at site S, which is surprising, given that no biotech industry is apparently located near site S.

Dibutyl phosphate (DBP) was the only compound in this class identified to level 1. It was found at all three sites, and in highest relative abundances at site D. While DBP is used in its own right as a reagent and additive in several processes, it is also a degradation product of TBP^43^. Its relative abundances in each plant matches that of TBP, so its presence in our effluent samples is possibly the result of partial degradation of TBP. Whether this compound should have been classified as a metabolite or a reagent is another matter of debate. We have again chosen to assign the category incurring least ambiguity. Further studies of influent vs effluent composition would clarify the origin of DBP.

#### Therapeutics and drugs

Therapeutics and drugs were by far the most abundant group of compounds found in wastewater treatment plant effluent. 114 individual compounds were identified, of which 27 were identified to level 1.

This is consistent with our expectations, given at-home consumption of prescription drugs, and the tendency for administered drugs to pass partially unabsorbed through the digestive system^44^, thus introducing both partially metabolised and untransformed drugs to the wastewater system. [insert something about bias in over-focus on medical drugs.]. The currently used mass spectral libraries are over-represented by medicine. Moreover, it is plausible that substances with high abundance are recovered to a greater extent with the used sample preparation procedure, or yields a better mass spectrometer signal (e.g. from more efficient ionization processes).

The **anticholerectic** drug dehydrocholic acid was found in high abundances at site A and S, but not at site D, suggesting either lower consumption or a more efficient degradation at site D.

We expected to find a number of **analgesics** in the effluent, and especially the drug indomethacin – a powerful NSAID used to treat arthritis - was significantly more prevalent at site A.

Among the identified opioids were a number of controlled substances such as Tramadol, the most used opioid in Denmark^45^, and Methadone, the cause of most drug-related deaths in Denmark^46^. The prescription of both these opioids outside hospitals is restricted in Denmark. While in-home treatment of patients is a possibility, these findings may be evidence of opioid drug abuse, which is a rising problem worldwide^47^.

**Antibiotics** are found at all three sites. Antimicrobial resistance is of growing concern globally, and wastewater treatment plants play a large role in the release of antimicrobial resistance genes into surrounding waterways and soil^48^. We found a significant difference in the prevalence of clarithromycin, with higher relative abundances in sites A and D compared to site S. Clarithromycin, along with other recovered macrolides, are among the most common antibiotics found in wastewater effluent^49^, contributing to the ongoing resistance crisis. Further studies quantifying the extent of antimicrobial resistance in the three sites is advisable, to assess the impact of antimicrobial resistance genes being released from the plants. We did, however, find a metabolite of clarithromycin, 14-hydroxyclarithromycin, at all sites with the help of the *in silico* fragmentation. While not significant, the relative abundance of this metabolite was higher at site S. Based on this finding, we hypothesise that the small plant might not be affected as harshly by antimicrobial resistance as the two large plants, and further studies would clarify this.

**Anticonvulsants** (antiepileptics) were found in all three sites, with oxcarbazepine being present in significantly higher relative abundances at site A compared to site S. We obtained two metabolites of oxcarbazepine from the *in silico* fragmentation. Oxcarbazepine is a suspected carcinogenic, while its analogue carbamazepine, which was found in similar relative abundances across sites, is a confirmed carcinogenic compound^50^. Both compounds are known to persist through wastewater treatment. Two degradation products of oxcarbazepine was found with the *in silico* fragmentation approach, licarbazepine and dihydroxycarbazepine.

**Antifungal** drugs are found in highest abundances in the effluent at site D. Climbazole, an anti-dandruff additive to shampoo known to be ecotoxic, particularly to primary producers^51, 52^, and terbinafine, an over- the-counter remedy for athlete’s foot and other minor fungal infections, are both commonly used compounds.

**Antihistamines** were found ubiquitously across all three sites, and the antihistamines fexofenadine and cetirizine were among the compounds recovered with the highest areas. While we are unable to provide quantitative data on the concentration of antihistamines in our samples, this finding is consistent with the prevalence of pollen allergy in Denmark, particularly during the summer season

**Anti-inflammatory drugs** were also recovered across three sites. Corticosteroids, in particular, were most prevalent at site S. It is possible that this particular result may be an artifact, as many compounds with a steroid-like structure may be wrongfully identified as the steroids present in the spectral libraries.

**Cancer treatment drugs** were also present in the wastewater effluent. The anti-androgen bicalutamide, used as an antihormone to treat prostate cancer were present in similar abundances across all three sites, while the antineoplastic agent megestrol acetate, a chemotherapy agent used to treat breast cancer, was found in higher relative abundances at site A. Chemotherapy agents are known to have adverse environmental impact^53^, and megestrol acetate has indeed been shown to be a potent endocrine disruptor^54^.

A number of **cardiovascular regulating drugs** were recovered in our dataset. The compounds bisoprolol, losartan and metoprolol were found ubiquitously throughout all three sites, while candesartan, diltiazem, propranolol and valsartan were present in significantly higher relative abundances at sites A and D compared to site S. Of these, bisoprolol and diltiazem has displayed negative effects on aquatic life^55^. Two metabolites of diltiazem, N-monodesmethyl diltilazem and deacetyl diltiazem, were found in all three sites, suggesting at least partial degradation on site, or *in vivo*.

The effect of losartan on aquatic environments falls within acceptable limits, while metoprolol and propranolol has been shown to have adverse effects^56, 57^.

Valsartan has been shown to have adverse effects on soil microbial communities and annelids^58^.

Two **anti-cholesterol drugs** were found in our samples: Fenofibric acid was primarily recovered from sites A and D. This compound has been found to impact fish development even in low concentrations^59^. Another drug, gemfibrozil, was found in all three sites, and has been shown to inhibit both aquatic life^60^ and plant life^61^.

A number of **illegal drugs** were recovered from the dataset. In particular, cannabis-related products and metabolites of these were recovered, with compounds such as HU-331 and 11-Hydroxy-d(9)-THC being present in far higher relative abundances at site A.

**Psycopharmaca** was found across all three sites, in similar relative abundances. Citalopram has been shown to affect the behavior of aquatic invertebrates^62^, while sertraline and venlafaxine exhibits similar adverse effects of marine vertebrates^63, 64^.

Prednisone, a synthetic anti-inflammatory **corticosteroid** used to treat a wide range of conditions, was found in significantly higher relative abundances at site A. Together with many other steroids, glucocortocoids are known to be potent endocrine disruptors, even in low concentrations^65^. One metabolite, 20α-dihydro-prednisolone, was found via *in silico* fragmentation, suggesting partial degradation of the compound.

A single **wormicide**, mebendazole, was found in similar abundances across sites. A common treatment for hookworm, this neurotoxic biocide targets annelids and will as such pose a risk to native annelid populations if released into waterways. It is frequently detected in wastewater effluent, and is considered a major driver of WWTP effluent toxicity^52^.

A number of single compounds, tentatively identified at level 2, were found at varying abundances across the three sites:

The anticoagulant rivaroxaban was found ubiquitously across sites; the anticholinergic compound benztropine was found in higher relative abundances at site D; dehydrocholic acid, a cholerectic, was found in higher relative abundances at site A; benzphetamine, a controlled substance used primarily for weight loss, was found almost exclusively at site A.

Xylometazoline, a commonly used nasal decongestant, was found in similar abundance across sites; the diuretic furosemide was found in higher relative abundances at sites A and D; the multifunctional drug 2-benzylideneindan-1-one, found to inhibit cancer cell growth^66^ and treat alzheimer’s^67^, was found mainly at site A; the immunosuppressant mycophenolic acid, used for autoimmune conditions and to prevent organ rejections after transplantation, was found ubiquitously across sites.

Norgestrel, a progestin; the prostaglandin 13-(2-Fluorophenoxy)-12-hydroxy-8-(1-hydroxyethyl)tridecanoic acid; the sedative etaqualone;the synthetic steroid 1,4-Androstadiene-3,17-dione; the uricosuric probenecid; veterinary drugs.

Although a comparison of influent compound abundances is necessary to comment further on between-site difference in abundance of these compounds without having compared influent compound abundances, we can hypothesize that the difference in effluent compound contents may be due to differences in treatment plant structure and treatment approach, as well as the microbial diversity in flotation and sludge tanks.

Notably, we found an interesting combination of compounds at site S. The presence of acetylcholinesterase inhibitor, antivirals, corticosteroids and cancer treatment compounds at such a small site, without the concurrent presence of commonly used compounds such as over-the-counter analgesics suggests that the population at site S is receiving intensive, high-dose treatment - likely for cancer. This is another example illustrating the efficiency of the non-targeted analysis as a monitoring tool – a small plant receiving high doses of cytotoxic agents might experience adverse effects on their microbial communities.

Naturally, the diversity of compounds found in effluent is directly dependent on site intake sources, which is not taken into account by this analysis, but given the similar sizes of site A and D, and the similar proportion of dwellings at site D, seeing such a diversity of compounds escape degradation is surprising. This leads us to hypothesize a more efficient degradation process at site D, but without knowing the contents of the influent, further studies comparing influent and effluent at each site are needed to elucidate this hypothesis.

#### Overall discussion

Studies reliant on pre-assembled libraries for compound identification are generally limited in their predisposition toward known entries in the databases. As such, these approaches can introduce bias into the results. This is arguably evident in the proportion of therapeutics and drugs found in our study. As compounds are continually investigated and added to spectral libraries, the degree of this bias is expected to decrease.

A study like ours, generating large numbers of unknown or poorly characterized compounds, have the potential to further advance the field on environmental chemistry and metabolomics by bringing attention to the need for further investigations into specific compounds. A number of the identified compounds exhibit various degrees of cytotoxicity, raising concern for their impact on the microbial communities of the treatment plants, and those of the outlet sites. Chemotherapy agents are among classes of compounds known to directly impact bioreactors, but the cytotoxicity of metabolites of different origin and their impact on WWTP microbial communities warrants further investigation.

Due to the limited scope of our study, examining only a small number of samples from three wastewater treatment plants, it is challenging to draw conclusions regarding the state of the wastewater treatment processes at these facilities. The ultrasensitive analytical methods employed in this study enables the detection of minute quantities of compounds, potentially including single molecules. It is important to note that the mere presence of a compound in this dataset does not necessarily imply presence in concentrations that warrant any concern. Further quantitative analyses of similar samples would be necessary to elucidate problematic compounds and their concentrations in the samples

Our findings suggest that improving the removal of xenobiotics from wastewater benefits from assessing a vast range of compounds present in the wastewater effluent using NTA. This approach is particularly promising because it requires minimal sample preparation and can capture a broad chemical space. However, the computational challenges of analyzing large datasets containing thousands of compounds remain, and sound statistical approaches are needed to unravel relevant trends in the datasets. We here present a novel approach to non-targeted sensitive environmental analysis of wastewater effluent, which could be applied to other WWTPs and environmental samples. The study’s pipeline and statistical tools could be used to screen WWTP effluent samples for both known and unknown xenobiotics, enabling a more precise characterization of xenobiotic diversity in wastewater effluent. Additionally, the study’s findings highlight the need for continued research and development of wastewater treatment technologies that can effectively remove xenobiotics from wastewater.

#### Conclusion

In this paper, we present a comparison of wastewater treatment plant effluent from three sites, applying a non-targeted analysis to samples of wastewater. This provides a comprehensive approach to discovering xenobiotics and identifying novel pollutants not targeted by classical analytical chemistry. While this approach only provides a snapshot of the compounds escaping each treatment plant, it has proven effective in identifying compounds of concern and revealing significant differences in the relative abundances of compounds escaping from different sites. We identified a number of compounds of emerging concern in the effluent, among them endocrine disruptors and cytotoxic compounds detrimental to aquatic ecosystems. These findings further highlight the need for investigation into cytotoxic compound effect on the diversity and specialization of microbial communities at each plant.

Our comparison of wastewater treatment plant effluent from three sites using a Phyloseq+Metacoder wrapper for chemical analyses allowed us to discover a large amount of undegraded compounds escaping to waterways in Denmark. Therefore, we propose using the non-targeted analysis as a monitoring tool for detecting pollutants in wastewater effluent, and provide a robust analysis pipeline to obtain graphical overviews of extremely complex datasets, comprehensible to a broad audience.

## 5. Materials and Methods

### 5.1 Chemicals

Chemicals and solvents used were: methanol from Sigma-Aldrich (MeOH, HPLC-grade, Buchs, Switzerland), and acetonitrile from Fischer Scientific (MeCN, HPLC-grade, Finland). Ultrapure water was tapped daily from a Dionex™ IC Pure™ Water Purification System (Thermo Scientific), 18.2 MΩ cm. Supplier mixed solvent in LC-MS grade from Fischer Scientific (Vantaa, Finland) included, acetonitrile with formic acid (0.1% v/v), water with formic acid (0.1% v/v), and water with trifluoroacetic acid (0.1% v/v). MS calibration was performed with Pierce™ LTQ Velos ESI Positive Ion Calibration Solution (Thermo Scientific). Chemical standards used for compound identification (level 1) were all purchased from NEOCHEMA GmbH (Germany). All glassware was cleaned and heated to 450°C prior to use.

### 5.2 Sampling

500 ml grab samples were taken in August 2018 between 09.00 and 12.00, in triplicates, at three different locations: site A (PE 350 000, located near suburbs. Wastewater input issues mainly from industry and households), site D (PE 400 000, located in city. Input from households, hospitals and - to a lesser degree - industry) and site S (PE 12 000, rural village location. Input from households and farms). We directly submerged glass bottles (site D and S) or used on-site, basin-specific sampling equipment (site A), and sampled inside the plant, immediately before the effluent release pipe. Samples were contained in glass bottles, covered, and put on ice for immediate transfer to the laboratory for solid phase extraction (SPE), to minimise adsorption and degradation.

### 5.3 SPE

Using cartridges with a hydrophilic-lipophilic balanced material (Oasis HLB, 200 mg), SPE was performed on triplicate samples from each site, along with one procedural blank per site. Cartridges were pre-conditioned with 5 ml methanol and 5 ml ultrapure water. 100 ml homogenized sample was loaded through PTFE tubes via vacuum at 3 ml/min., while ultrapure water was loaded into the procedural blanks. After loading, the cartridges were washed with 5 ml ultrapure water, followed by 5 ml of aq. methanol (10% v/v ultrapure water). Cartridges dried on the vacuum manifold for 30 minutes, and subsequently stored at −20 °C overnight. After acclimatising in a fume hood for 15 minutes, samples were eluted into glass test tubes using 5 ml methanol at 3 ml/min.

Solvent was evaporated under a gentle flow of nitrogen. Samples were reconstituted in 100 µl acetonitrile, before addition of 900 µl of loading solvent, an aqueous solution of trifluoroacetic acid (0.05% v/v, TFA) and MeCN (2% v/v) resulting in a sample enrichment factor of 100.

#### 5.3.1 Quality control

All samples were processed immediately after sampling, on the same designated vacuum manifold, using the same glassware and disposable glass pipettes. Each site was processed as a single batch, with one procedural blank created per batch. Prior to analysis, pooled samples were made as a mix of all samples, representing as quality control (QC) for normalizing for any systematic errors during data acquisition^68^. 100 µl were taken out of each HPLC vial, combined and the vials were topped up to 1 ml with loading buffer. All samples were stored at 4 °C before analysis.

### 5.4 Instrumentation

Nanoflow liquid chromatography (Dionex Ultimate 3000 NCS-3500RS Proflow, Thermo Scientific) was performed utilizing online sample clean-up by solid-phase extraction upon a C18 pre-column (Acclaim PepMap, 100Å, C18, 0.3 mm × 5 mm, 5 µm, Thermo Scientific). During injection, 5 µL sample was loaded upon the pre-column for 2 minutes, injected using full-loop-pickup mode. The loading pump delivered a flow of 2% acetonitrile and 0.1% formic acid in water at 30 µL/min, after which a valve switch eluted trapped analytes onto a C18 (nanoflow-)column (PepMap RSLC, C18, 2 µm, 100Å, 75 µm x 15 cm, Thermo Scientific) using a flowrate of 300 nL/min. Mobile phases consisted of (A) 2% acetonitrile and 0.1% formic acid in water and (B) 98% acetonitrile and 0.1% formic acid in water. A runtime of 30 minutes was achieved using the following gradient: 10% B was held for 2 minutes followed by a linear increase to 95% B over 20 minutes. This plateau was maintained for 3 minutes, after which initial conditions were restored over 30 seconds. The column was then re-equilibrated for 4.5 minutes. The column temperature was held at 40°C, and samples were kept at 8 °C.

High-resolution accurate-mass spectrometric analysis (Q Exactive HF Orbitrap, Thermo Scientific) was performed using a combination of both full-scan and data-dependent acquisition mode. Positive polarity electrospray ionization was achieved using a spray voltage of 2.0 kV and capillary temperature of 250 °C. Full scan spectra were recorded in profile mode at scan range at a resolution of 240,000 (at m/z 200) using a mass range of 100 to 1500 m/z with the automated gain control (AGC) target set to 3e6. MS^2^ spectra were recorded at a resolution of 15,000 (at m/z 200), and were acquired using a Top10-approach with stepped collision energies of 15, 30, and 45 NCE. An AGC target of 1e5, maximum injection time of 50 ms, and isolation width of 4.0 m/z was used during data-dependent acquisition. 20 seconds of dynamic exclusion was enabled to prevent consecutive fragmentation of the same ion(s). Calibration of the mass spectrometer was performed weekly using a Pierce™ LTQ Velos ESI Positive Ion Calibration Solution (Thermo-Fischer Scientific). During acquisition, mass accuracy was adjusted towards a lock mass of m/z 371.10124^69^.

### 5.5 Data processing pipeline

Data acquisition was controlled with Thermo Scientific Xcalibur (v 4.1.31.9), and data was processed using Compound Discoverer (Thermo Scientific, v3.3). Retention time (RT) was set with a lower limit of 8 and an upper limit of 28 minutes, and RT was aligned using a linear model, with tolerance set to 0.5 minutes. Mass accuracy tolerance was set to 3 ppm. Any compounds with a sample-to-blank ratio of less than 100 were set to be ignored, as well as any compounds found in the QC blanks. Compounds without MS2 data were also omitted.

All level 2^18^ compound annotations were based on online spectral library searches (e.g. mzCloud), secondary mass list searching the ChemSpider database or in-house mass lists, and Compound Discoverer’s internal compound prediction software. Mass List and ChemSpider hits were run against mzLogic.

All level 1^18^ compound annotations were based on in-house spectral library searches (e.g. mzVault), secondary mass list searching the mzCloud and ChemSpider databases, and Compound Discoverer’s internal compound prediction software. Mass List and ChemSpider hits were run against mzLogic.

Post-processing descriptive statistics and differential analyses nodes were enabled in Compound Discoverer (See Supplementary Figure 1 for workflow).

Following data processing, any unidentified compounds were removed by manual data curation, and any compound not present in all three replicates from at least one sampling site was deemed a false positive and filtered out.

The obtained compounds were tentatively identified by either internal standard libraries (mzVault) or cloud databases; mzCloud, as well as ChemSpider, MassList, and Compound Discoverer’s predicted compositions. Compound classes and subclasses were assigned to each compound.

Using Compound Discoverer’s inbuilt differential analysis, a number of compounds identified to level 1 were found to significantly differ in output between treatment plants (LfC > 1 and p < 0.05). These were chosen for *in silico* fragmentation in an attempt to identify further possible metabolites. The most abundant compound in the dataset was also similarly investigated.

These significantly differing compounds were run separately on expected compounds workflows (environmental expected with transformations) in Compound Discoverer, resulting in a separate file for each of the compounds, containing all detected metabolites. Any known metabolites were identified by EnviPath, Pubchem, Drugbank or relevant literature (see references in [TABLE metabolites]), and any metabolites matching the molecular formula in each Compound Discoverer file were in silico fragmented using MetFrag, and cross referenced with ChemSpider.

Any unidentified molecules were dropped from the dataset, and any compounds not identifiable beyond molecular composition were classified as “unknown natural product”. All other compounds were researched and classified based on their most common application as stated by credible sources [SI X]. (e.g., the compound identified as Fexofenadine was classified as “Therapeutics and Drugs” and sub-classified as “Antihistamine”, as its most common application is in over-the-counter allergy medicine). In some less obvious cases (e.g. 1,4-Androstadiene-3,17-dione, a steroid precursor used in industry, also misused as a precursor for drugs of abuse) the final classification was agreed upon by the authors after careful evaluation of available sources. Metabolites of endogenous hormones were classified as “Metabolites”, and sub-classified by their organism of origin (e.g. fungal, human, etc.). References used as the base for these classifications can be found in [TABLE metabolites], along with simple encyclopaedia entry links for well-studied compounds such as cortisol.

### 5.6 Statistical analysis

All statistical analysis and data visualization was done in R 3.6.3^70^

For initial data visualisation, we constructed barplots of observed and relative abundances of compounds illustrated by compound classes.

To compare within-site and between-site variance, a PERMANOVA using a Bray-Curtis dissimilarity matrix with 999 permutations was applied to the filtered dataset, using a pairwise Adonis test for post hoc testing. We then performed a Principal Coordinates Analysis on QC vs site samples, applying a Bray-Curtis dissimilarity matrix using Phyloseq^71^.

Similar chemometrics were carried out by Sogin et al.^72^, using Bray-Curtis dissimilarity on an NMDS plot, and by Botticelli et al.^73^, who used a Mann-Whitney-U test on multivariate metabolomics data. We, however, chose to build a wrapper for an already established data analysis package, Phyloseq. This R package is originally intended for microbial ecological analysis, and integrates inputs in the form of abundance data, phylogenetic (identity and classification) information and covariates. This enables exploratory data transformation and plotting, as well as application of a suite of non-parametric, multivariate statistical analysis. While built for ecological analysis of microbial sequence data, basic Phyloseq input data (a Phyloseq Object) is composed from an array of matrices – tables of numerical abundance data, taxonomic/compound classification, and metadata such as sites or sample dates. We can apply this analysis framework to any structurally similar data, in this case for cheminformatics of NTA data. The advantage of constructing an environmental cheminformatics wrapper lies in Phyloseq being an established, peer-reviewed and widely used platform with an array of statistical tools for both parametric and non-parametric multivariate data – the data classes environmental NTA data generally belongs to.

To visualize single compound variance, we created differential heat trees using MetacodeR – a package visualizing log fold changes on cladograms. The compositionally transformed dataset was converted to a tibble using tidyverse packages, and the per-compound proportions and occurrences were calculated as log2 fold changes between wastewater sites. These were then drawn on to a cladogram constructed from compound classes and subclasses using the heat_tree function in MetacodeR. We used the Fructerman-Reingold algorithm for primary cladogram layout due to the number of subclasses. The standard Reingold-Tilford algorithm was applied for initial node layout.

An identical set of analyses was performed on a subset of the data, including only compounds in the “Therapeutics and Drugs” class.

## Supporting information

Supplementary Metabolite Catalogue 1

## Abbreviations

WWTP: Wastewater treatment plant
NTA: non-targeted analysis
HRMS: High-resolution mass spectrometry
UHPLC: ultra-high pressure liquid chromatography
SPE: solid-phase extraction
PE: population equivalent
CD: Compound Discoverer
RT: Retention Time
TFA: trifluoroacetic acid.

## Acknowledgements

The authors acknowledge the Danish EPA for partly funding of (HITLIST project) study, the Carlsberg Foundation (grant number CF20-0422), the Aarhus University Research Foundation (grant numbers AUFF-T-2017-FLS-7-4 and AUFF-E-2017-7-21) and the Aarhus University Centre for Water Technology (WATEC). Dr. Rune Græsbøll for insights and sparring on results.

## Supplementary materials

Supplementary Figure 1 shows the workflow setup in Compound Discoverer. Parameter setup for each node is stated after the figure.

Supplementary table 1

All collected data used for all 785 compounds.

Supplementary Metabolite Catalogue 1

All putative identified compound with detailed information about known metabolites and their prevalence in the dataset.

