## Supplementary Figure 1 for "Non-target Analysis of Wastewater Treatment Plant Effluents: Chemical Fingerprinting as a Monitoring Tool"

*Supplementary Figure 1: Compound Discoverer workflow*

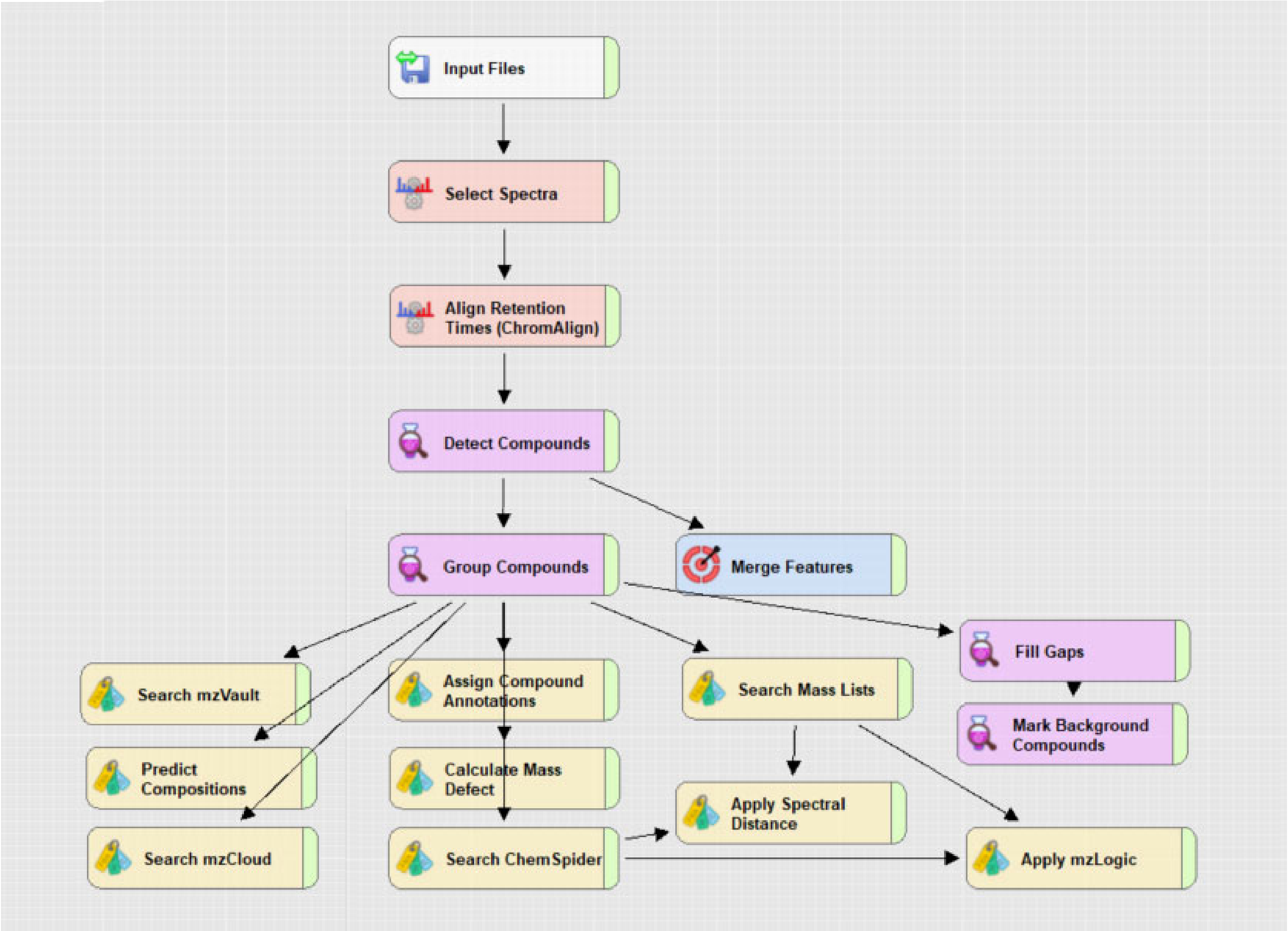


**Input files:**
12 samples (3 biological replicates and 1 procedural blank per site, for 3 sites), 3 QC pools and 3 MS2 identification pools.

**Selected spectra**
Lower and upper RT limit both set at 0.

**Align Retention times (ChromAlign)**
Reference file: QC pool 1.

**Detect Compounds**
Standard settings used, except for:
Mass tolerance set to 3 ppm. Min peak intensity set to 1.000.000 (1 million)

**Group compounds**
Standard settings used, except for:
Mass tolerance set to 3 ppm. Peak rating set to 2.

**Merge Features**
Mass tolerance set to 3 ppm

**Search mzVault**
mzVault library file used:  LC-pos – In-house xenobiotics v. 1.0

**Predict Compositions**
Mass tolerances set to 3 ppm

**Search mzCloud**
Standard settings used.

**Assign Compound Annotations**
Mass tolerance set to 3 ppm. Data sources set in order: Predicted compositions, mzCloud search, mzVault search, MassList search, ChemSpider search.

**Calculate Mass Defect**
Standard settings used.

**Search ChemSpider**
Databases used: BioCyc; ChEMBL, KEGG, Mass tolerance set to 3 ppm.

**Search Mass Lists**
Mass Lists used: EFS HRAM Compound Database, Extractables and Leachables HRAM Compound Databases. Mass tolerance set to 3 ppm.

**Apply Spectral Distance**
Mass tolerance set to 3 ppm.

**Fill Gaps**
Mass tolerance set to 3 ppm.

**Mark Background Compounds**
Standard settings used.

**Apply mzLogic**
Standard settings used.
