## Supplementary table 1 for "Non-target Analysis of Wastewater Treatment Plant Effluents: Chemical Fingerprinting as a Monitoring Tool"

| Compounds_ID | Tag_mzCloud_hit | Tag_mzCloud_best_conf_10plus | Tag_mzVault_hit | Name | Class | Subclass_I | Subclass_II | CAS_number |
|---|---|---|---|---|---|---|---|---|
| 6 | A | | | (S)-Dibenzy | Amino Acid | Derivative | | 2791-79-9 |
| 643 | A | B | | 4-Methylb | Industrial ( | Corrosion inhibitor | | 29878-31-[illegible] |
| 1790 | A | B | | 4-Methylb | Industrial ( | Corrosion inhibitor | | 29878-31-[illegible] |
| 9 | A | | | Fluoresceir | Industrial ( | Dye/stain | | 2321-07-5 |
| 99 | A | | | Tripropyl p | Industrial ( | Flame retardant | | 513-08-6 |
| 399 | A | B | | Tripropyl p | Industrial ( | Flame retardant | | 513-08-6 |
| 12 | A | B | C | Tris(2-butc | Industrial ( | Flame retardant | | 78-51-3 |
| 5276 | A | B | | Tris(2-chlo | Industrial ( | Flame retardant | | 115-96-8 |
| 5946 | | | C | Photoinitia | Industrial ( | Photoinitiator | | |
| 5118 | A | B | | Benzyl but | Industrial ( | Plasticizer | | 85-68-7 |
| 4412 | A | B | | beta-Buto | Industrial ( | Plasticizer | | 117-83-9 |
| 3690 | A | B | | Bisphenol | Industrial ( | Plasticizer | | 10192-62-[illegible] |
| 2461 | A | B | | Centralite | Industrial ( | Plasticizer | | 85-98-3 |
| 706 | A | | | Oxybisprop | Industrial ( | Plasticizer | | 27138-31-[illegible] |
| 781 | A | | | Oxybisprop | Industrial ( | Plasticizer | | 27138-31-[illegible] |
| 2000 | A | | | Oxybisprop | Industrial ( | Plasticizer | | 27138-31-[illegible] |
| 17 | A | B | C | Tributyl Ph | Industrial ( | Plasticizer | | 126-73-8 |
| 425 | A | | | (2R,3R,4R, | Industrial ( | Surfactant | PFAS | |
| 5574 | A | B | | Monolauri | Industrial ( | Surfactant | | 27215-38-[illegible] |
| 5710 | A | B | | 4-Aminodi | Industrial Chemicals | | | 101-54-2 |
| 2542 | A | B | | PPG Acryla | Industrial Chemicals | | | |
| 51 | A | B | | Corticoste | Metabolite | Corticosteroid | | 50-22-6 |
| 65 | A | B | | Corticoste | Metabolite | Corticosteroid | | 50-22-6 |
| 1745 | A | B | | Corticoste | Metabolite | Corticosteroid | | 50-22-6 |
| 4660 | A | B | | Corticoste | Metabolite | Corticosteroid | | 50-22-6 |
| 570 | A | B | | Cortisol | Metabolite | Corticosteroid | | 50-23-7 |
| 1446 | A | B | | Cortisol | Metabolite | Corticosteroid | | 50-23-7 |
| 5767 | A | B | | Eicosapent | Metabolite | Fatty acid | | 10417-94-[illegible] |
| 3503 | A | B | | Eicosapent | Metabolite | Fatty acid | | 2734-47-6 |
| 716 | A | | | 1,3,8-trime | Metabolite | Fish metabolite | | |
| 4639 | A | B | | Nanangeni | Metabolite | Fungal me | Sesquiterp | 1609469-7 |
| 728 | A | | | 6-[(3E,5E,7 | Metabolite | Fungal metabolite | | |
| 3946 | A | B | | 6-[(3E,5E,7 | Metabolite | Fungal metabolite | | |
| 4107 | A | | | 6-[(3E,5E,7 | Metabolite | Fungal metabolite | | |
| 1777 | A | B | | preussin E | Metabolite | Fungal metabolite | | |
| 3219 | A | B | | Verrucarol | Metabolite | Fungal metabolite | | |
| 5141 | A | B | | Verrucarol | Metabolite | Fungal metabolite | | |
| 5361 | A | B | | Verrucarol | Metabolite | Fungal metabolite | | |
| 5567 | A | B | | Verrucarol | Metabolite | Fungal metabolite | | |
| 6320 | A | B | | Verrucarol | Metabolite | Fungal metabolite | | |

| | | | | | | | | | |
|---|---|---|---|---|---|---|---|---|---|
| 1206 | A | B | | 16alpha-H | Metabolite | Human Me | Steroid | | 566-76-7 |
| 1322 | A | | | 17a-Hydro | Metabolite | Human Me | Steroid | | 68-96-2 |
| 997 | A | B | | 2-Methoxy | Metabolite | Human Me | Steroid | | 362-08-3 |
| 4204 | A | B | | 6beta-Hyd | Metabolite | Human Me | Steroid | | 62-99-7 |
| 704 | A | B | | 7a-Hydrox | Metabolite | Human Me | Steroid | | 62-83-9 |
| 8 | A | B | | 7alpha-Hy | Metabolite | Human Me | Steroid | | 30626-96- |
| 193 | A | B | | 7alpha-Hy | Metabolite | Human Me | Steroid | | 30626-96- |
| 2866 | A | B | | 7alpha-Hy | Metabolite | Human Me | Steroid | | 30626-96- |
| 717 | A | B | C | Aldosteror | Metabolite | Human Me | Steroid | | 52-39-1 |
| 3935 | A | B | | Estriol | Metabolite | Human Me | Steroid | | 50-27-1 |
| 5261 | A | B | | Estriol | Metabolite | Human Me | Steroid | | 50-27-1 |
| 1284 | A | B | | Pregnenol | Metabolite | Human Me | Steroid | | 145-13-1 |
| 1493 | A | B | C | Pregnenol | Metabolite | Human Me | Steroid | | 145-13-1 |
| 1699 | A | B | | Tetrahydr | Metabolite | Human Me | Steroid | | 53-05-4 |
| 5370 | A | | | 2-(1H-ben | Metabolite | Human Metabolites | | | |
| 3841 | A | B | | 5-OxoETE | Metabolite | Human Metabolites | | | 106154-18 |
| 4491 | A | B | | 5-OxoETE | Metabolite | Human Metabolites | | | 106154-18 |
| 603 | A | B | | Coenzyme | Metabolite | Human Metabolites | | | 606-06-4 |
| 909 | A | B | | Coenzyme | Metabolite | Human Metabolites | | | 606-06-4 |
| 1061 | A | B | | Coenzyme | Metabolite | Human Metabolites | | | 606-06-4 |
| 3218 | A | B | | Coenzyme | Metabolite | Human Metabolites | | | 606-06-4 |
| 683 | A | B | | (+/-)-C75 | Metabolite | Lactone | | | 218137-86 |
| 691 | A | B | | 13(S)-HOT | Metabolite | Plankton metabolite | | | |
| 4698 | A | B | | 13(S)-HOT | Metabolite | Plankton metabolite | | | |
| 168 | A | B | | Sedanolide | Metabolite | Plant Meta | | 0 | 6415-59-4 |
| 6196 | A | B | | Senkyunoli | Metabolite | Plant Meta | | 0 | 94596-27- |
| 3229 | A | | | Tanshinon | Metabolite | Plant Meta | | 0 | 568-72-9 |
| 4254 | A | | | daphmanic | Metabolite | Plant Meta | Alkaloid | | |
| 3005 | A | B | | Piperine | Metabolite | Plant Meta | Alkaloid | | 94-62-2 |
| 3177 | A | B | | Pinusolide | Metabolite | Plant Meta | Diterpene | | 31685-80- |
| 25 | A | B | | **Diisobutyl** | Metabolite | Plant Meta | Fatty acid | | 141-04-8 |
| 1914 | A | | | 7,9-di-tert- | Metabolite | Plant Meta | Flavonoid | | 82304-66- |
| 2731 | A | B | | (2E)-N-(4-a | Metabolite | Plant meta | Sesquiterpenoid | | |
| 5626 | A | B | | (3aR,4aS,5 | Metabolite | Plant meta | Sesquiterpenoid | | |
| 2318 | A | B | | (3aR,7aS,8 | Metabolite | Plant meta | Sesquiterpenoid | | |
| 6101 | A | B | | (3aR,7aS,8 | Metabolite | Plant meta | Sesquiterpenoid | | |
| 4486 | A | B | | Graveolide | Metabolite | Plant meta | Sesquiterpenoid | | |
| 4911 | A | | | Microhele | Metabolite | Plant meta | Sesquiterpenoid | | |
| 1519 | A | B | | Nootkaton | Metabolite | Plant Meta | Sesquiterpenoid | | |
| 4405 | A | B | | Psilostach | Metabolite | Plant meta | Sesquiterpenoid | | |
| 5036 | A | B | | Psilostach | Metabolite | Plant meta | Sesquiterpenoid | | |
| 5825 | A | B | | Psilostach | Metabolite | Plant meta | Sesquiterpenoid | | |
| 2096 | A | B | | Zedoarond | Metabolite | Plant meta | Sesquiterpenoid | | |
| 2119 | A | B | | Zedoarond | Metabolite | Plant meta | Sesquiterpenoid | | |
| 4479 | A | B | | (3beta,20S | Metabolite | Plant meta | Steroid | | |
| 3845 | A | B | | (5?)-12,13- | Metabolite | Plant meta | Trinorditerpene | | |
| 700 | A | B | | (11E,15Z)- | Metabolite | Plant metabolite | | | 185147-97 |
| 1650 | A | B | | 1,4a-Dime | Metabolite | Plant metabolite | | | |
| 4991 | A | B | | 1,4a-Dime | Metabolite | Plant metabolite | | | |
| 24 | A | | | 2-Methyl-S | Metabolite | Plant Metabolite | | | 615-22-5 |
| 143 | A | | | 6'-Methox | Metabolite | Plant Metabolite | | | |

| | | | | | | | | |
|---|---|---|---|---|---|---|---|---|
| 3460 | A | B | | Abietic aci | Metabolite | Plant Metabolite | | 514-10-3 |
| 3053 | A | B | | Arnicolide | Metabolite | Plant Metabolite | | 36505-53-( |
| 6296 | A | B | | Arnicolide | Metabolite | Plant Metabolite | | 34532-68-8 |
| 1000 | A | B | | Broussone | Metabolite | Plant metabolite | | 173327-05 |
| 1253 | A | B | | Broussone | Metabolite | Plant metabolite | | 173327-05 |
| 2373 | A | | | Celastrol | Metabolite | Plant Metabolite | | 34157-83-( |
| 5744 | A | B | | Dipropyl pl | Metabolite | Plant Metabolite | | 131-16-8 |
| 1462 | A | B | | Gibberellic | Metabolite | Plant Metabolite | | 77-06-5 |
| 3950 | A | B | | Hecogenin | Metabolite | Plant Metabolite | | 915-35-5 |
| 2927 | A | B | | Kahweol | Metabolite | Plant Metabolite | | 6894-43-5 |
| 928 | A | | | Norisoguai | Metabolite | Plant metabolite | | 50376-42-( |
| 128 | A | B | C | Octisalate | Personal C | Sunscreen | | |
| 3884 | A | | | Ostruthin | Metabolite | Plant Metabolite | | 148-83-4 |
| 5061 | A | B | | 11-Ketotes | Metabolite | Steroid | | 53187-98-7 |
| 5581 | A | B | | 11-Ketotes | Metabolite | Steroid | | 53187-98-7 |
| 50 | A | | | Galaxolido | Personal C | Metabolite | Xenobiotic | 507442-49 |
| 323 | A | | | Galaxolido | Personal C | Metabolite | Xenobiotic | 507442-49 |
| 460 | A | | | Galaxolido | Personal C | Metabolite | Xenobiotic | 507442-49 |
| 4430 | A | B | | Enzacamer | Personal C | Sunscreen | | 36861-47-9 |
| 57 | A | B | C | Propiconaz | Pesticides | Fungicide | Triazole | 60207-90-1 |
| 4544 | A | B | C | Tebuconaz | Pesticides | Fungicide | Triazole | 107534-96 |
| 248 | A | B | C | DEET | Pesticides | Fungicide | | 134-62-3 |
| 1048 | A | B | | Furmecycl | Pesticides | Fungicide | | 60568-05-( |
| 631 | A | B | C | Prosulfoca | Pesticides | Herbicide | Xenobiotic | 52888-80-9 |
| 118 | A | B | C | Prometryn | Pesticides | Herbicide | | 7287-19-6 |
| 1713 | A | | | QNK | Reagents a | Enzyme | | |
| 927 | A | B | | N-[(S)-(+)-1 | Reagents a | Peptide synthesis | | 82717-96-2 |
| 158 | A | B | | 1-Hydroxy- | Reagents and standards | | | 5408-46-8 |
| 1270 | A | B | | 1-Hydroxy- | Reagents and standards | | | 5408-46-8 |
| 61 | A | | | 2,7-Diacety | Reagents and standards | | | 961-27-3 |
| 23 | A | B | | 2-Ethyl-5-r | Reagents and standards | | | 131298-44 |
| 5484 | A | | | 2-Hydroxyl | Reagents and standards | | | 934-34-9 |
| 342 | A | B | | 9-Azabicyc | Reagents and standards | | | 4946-36-5 |
| 1755 | A | B | | Bis(2-ethyl | Reagents and standards | | | 106-20-7 |
| 2774 | A | B | | Bis(2-ethyl | Reagents and standards | | | 106-20-7 |
| 414 | A | | | Cyclohexyl | Reagents and standards | | | |
| 999 | A | | C | Dibutyl ph | Reagents and standards | | | 107-66-4 |
| 5057 | A | | | Dodecylam | Reagents and standards | | | 124-22-1 |
| 2479 | A | | | Octadecan | Reagents and standards | | | 124-30-1 |
| 3164 | A | B | | 3,3'-Dimet | Textile Che | Intermediate | | 119-90-4 |
| 1898 | A | B | | 4,4'-Bis(die | Textile Che | Intermediate | | 90-94-8 |
| 1385 | A | B | | OPEO | Textile Chemicals/Auxiliary Dyes | | | |
| 3509 | A | B | | OPEO | Textile Chemicals/Auxiliary Dyes | | | |
| 187 | A | | | Neostigmi | Therapeut | Acetylcholinesterase i | | 59-99-4 |
| 4572 | A | | | 4-(4-{[(3R)- | Therapeut | AIDS/HIV | Metabolite | |
| 3979 | A | B | | (6aS,7aS,8 | Therapeut | Analgesic | Anaesthetic | |
| 4504 | A | B | | Tramadol I | Therapeut | Analgesic | Metabolite | 27203-92-5 |
| 163 | A | B | C | Diclofenac | Therapeut | Analgesic | NSAID | 15307-86-5 |
| 1161 | A | | | Etodolac | Therapeut | Analgesic | NSAID | 41340-25-4 |
| 1611 | A | B | C | Indometha | Therapeut | Analgesic | NSAID | 53-86-1 |
| 1736 | A | B | | Ketoprofer | Therapeut | Analgesic | NSAID | 22071-15-4 |

| | | | | | | | | |
|---|---|---|---|---|---|---|---|---|
| 4502 | A | B | | Ketoprofer | Therapeut | Analgesic | NSAID | 22071-15-4 |
| 1511 | A | B | C | Naproxen | Therapeut | Analgesic | NSAID | 22204-53-1 |
| 2381 | A | | | Phenazone | Therapeut | Analgesic | NSAID | 60-80-0 |
| 549 | A | B | | Alphaprod | Therapeut | Analgesic | Opioid | 77-20-3 |
| 1950 | A | | | Meperidine | Therapeut | Analgesic | Opioid | 57-42-1 |
| 394 | A | B | | Methadon | Therapeut | Analgesic | Opioid | 76-99-3 |
| 1379 | A | | | Normorph | Therapeut | Analgesic | Opioid | 466-97-7 |
| 1 | A | | | Tapentado | Therapeut | Analgesic | Opioid | 175591-23 |
| 188 | A | | | Tramadol | Therapeut | Analgesic | Opioid | 27203-92-5 |
| 6114 | A | | | Tramadol | Therapeut | Analgesic | Opioid | 27203-92-5 |
| 1424 | A | | | Phendimet | Therapeut | Anorectic | | 634-03-7 |
| 1564 | A | B | | Clopidogre | Therapeut | Antiaggregant | | 113665-84 |
| 723 | A | B | | 2-Amino-5 | Therapeut | Antibiotic | Metabolite | 107752-89 |
| 410 | A | | C | Azithromy | Therapeut | Antibiotic | | 83905-01-5 |
| 2077 | A | B | C | Clarithrom | Therapeut | Antibiotic | | 81103-11-9 |
| 1454 | A | B | C | Roxithrom | Therapeut | Antibiotic | | 80214-83-1 |
| 3159 | A | | | Sutezolid | Therapeut | Antibiotic | | 168828-58 |
| 4191 | A | B | C | Trimethop | Therapeut | Antibiotic | | 738-70-5 |
| 292 | A | | | Benztropin | Therapeut | Anticholinergic | | 86-13-5 |
| 3997 | A | B | | Rivaroxaba | Therapeut | Anticoagulant | | 366789-02 |
| 496 | A | B | | 10,11-Dihy | Therapeut | Anticonvul | Metabolite | |
| 221 | A | | | 10-Hydrox | Therapeut | Anticonvul | Metabolite | |
| 658 | A | | | 10-Hydrox | Therapeut | Anticonvul | Metabolite | |
| 2692 | A | B | | Carbamaze | Therapeut | Anticonvul | Metabolite | 36507-30-9 |
| 5187 | A | B | | Carbamaze | Therapeut | Anticonvul | Metabolite | 36507-30-9 |
| 172 | A | B | C | Carbamaze | Therapeut | Anticonvulsant | | 298-46-4 |
| 268 | A | | | Lamotrigin | Therapeut | Anticonvulsant | | 84057-84-1 |
| 498 | A | B | C | Oxcarbaze | Therapeut | Anticonvulsant | | 28721-07-5 |
| 3981 | A | B | | Topiramate | Therapeut | Anticonvulsant | | |
| 361 | A | B | | Climbazole | Therapeut | Antifungal | | 38083-17-9 |
| 20 | A | B | | Terbinafine | Therapeut | Antifungal | | 91161-71-6 |
| 149 | A | | C | Cetirizine | Therapeut | Antihistamine | | 83881-51-0 |
| 5 | A | B | C | Fexofenad | Therapeut | Antihistamine | | 83799-24-0 |
| 5151 | A | B | | Hydrocorti | Therapeut | Anti-inflam | Corticoster | 13609-67-1 |
| 5498 | A | B | | L6H21 | Therapeut | Anti-inflammatory | | 18778-38-0 |
| 5654 | A | B | | Crotamitor | Therapeut | Antipruritic | | 483-63-6 |
| 1769 | A | B | | 1,3-Adama | Therapeut | Antiviral | | 17768-28-4 |
| 232 | A | B | | 15-deoxy-d | Metabolite | Human Metabolites | | 85235-11-0 |
| 2269 | A | B | | 4beta-Phor | Metabolite | Plant Metabolite | | 17673-25-5 |
| 2052 | A | B | C | Bicalutami | Therapeut | Cancer trea | Antihormo | 90357-06-5 |
| 5172 | A | | | Megestrol | Therapeut | Cancer trea | Antineopla | 595-33-5 |
| 3436 | A | B | | Alfuzosin | Therapeut | Cardiovasc | Alpha bloc | 81403-80-7 |
| 5625 | A | | | Flusoxolol | Therapeut | Cardiovasc | Beta block | 84057-96-5 |
| 59 | A | B | C | Metoprolo | Therapeut | Cardiovasc | Beta block | 51384-51-1 |
| 1438 | A | B | | Metoprolo | Therapeut | Cardiovasc | Beta block | 51384-51-1 |
| 979 | A | B | C | Propranolo | Therapeut | Cardiovasc | Beta block | 525-66-6 |
| 422 | A | B | | 3,4-Dimeth | Therapeut | Cardiovasc | Metabolite | 67775-97-7 |
| 2787 | A | B | C | Bisoprolol | Therapeut | Cardiovascular regula | | 66722-44-9 |
| 254 | A | B | C | Candesarta | Therapeut | Cardiovascular regula | | 139481-59 |
| 3542 | A | B | C | Diltiazem | Therapeut | Cardiovascular regula | | 42399-41-7 |
| 266 | A | B | | Flecainide | Therapeut | Cardiovascular regula | | 54143-55-4 |

| | | | | | | | | |
|---|---|---|---|---|---|---|---|---|
| 5353 | A | B | | Labetalol | Therapeut | Cardiovascular regula | | 36894-69-( |
| 14 | A | B | C | Losartan | Therapeut | Cardiovascular regula | | 114798-26 |
| 117 | A | B | | N,N'-Dicycl | Therapeut | Cardiovascular regula | | 2387-23-7 |
| 171 | A | B | C | Valsartan | Therapeut | Cardiovascular regula | | 137862-53 |
| 4691 | A | | | Dehydroch | Therapeut | Cholerectic | | 81-23-2 |
| 5842 | A | B | | Fenofibric | Therapeut | Cholesterol medicatio | | 42017-89-( |
| 90 | A | B | | Gemfibroz | Therapeut | Cholesterol medicatio | | 25812-30-( |
| 790 | A | B | | Benzpheta | Therapeut | Controlled substance | | 156-08-1 |
| 5106 | A | | | Xylometaz | Therapeut | Decongestant | | 526-36-3 |
| 1016 | | | C | Furosemid | Therapeut | Diuretic | | 54-31-9 |
| 45 | A | B | | 11-Hydrox | Therapeut | Drugs of al | Metabolite | |
| 448 | A | | | 2-Ethylider | Therapeut | Drugs of al | Metabolite | 30223-73-! |
| 2054 | A | B | | 6-Acetylm | Therapeut | Drugs of al | Metabolite | 2784-73-8 |
| 4075 | A | B | | Nor-9-carb | Therapeut | Drugs of al | Metabolite | 56354-06-4 |
| 2662 | A | B | | Methyldie | Therapeut | Drugs of al | Steroid | 14531-89-( |
| 864 | A | | | RCS-4 | Therapeut | Drugs of al | Synthetic c | 1345966-7 |
| 5823 | A | B | | (+/-)-Cann | Therapeut | Drugs of abuse/Illegal | | drugs |
| 1390 | A | B | | 5a-Andros | Therapeut | Drugs of abuse/Illegal | | 2243-05-2 |
| 1474 | A | B | | 5a-Andros | Therapeut | Drugs of abuse/Illegal | | 2243-05-2 |
| 6395 | A | B | | 5a-Andros | Therapeut | Drugs of abuse/Illegal | | 2243-05-2 |
| 6660 | A | B | | AB-CHMIN | Therapeut | Drugs of abuse/Illegal | | drugs |
| 3096 | A | | | Acetyl nor | Therapeut | Drugs of abuse/Illegal | | 22352-82-! |
| 302 | A | B | | DOAM | Therapeut | Drugs of abuse/Illegal | | 63779-90-8 |
| 465 | A | B | | HU-331 | Therapeut | Drugs of abuse/Illegal | | 137252-25 |
| 1233 | A | B | | HU-331 | Therapeut | Drugs of abuse/Illegal | | 137252-25 |
| 4253 | A | B | | HU-331 | Therapeut | Drugs of abuse/Illegal | | 137252-25 |
| 650 | A | B | | 2-benzylid | Therapeut | Immunosuppressant | | 17598-66-2 |
| 440 | A | B | | Mycophen | Therapeut | Immunosuppressant | | 24280-93-1 |
| 3873 | A | B | | (5R,7S,8R)- | Therapeut | Pharmace | Synthetic | |
| 4882 | A | B | | Norgestrel | Therapeut | Progestin | | 6533-00-2 |
| 4428 | A | | | 13-(2-Fluo | Therapeut | Prostaglandin | | |
| 918 | A | B | | Desmethyl | Therapeut | Psychopha | Metabolite | 62498-67-3 |
| 1093 | A | B | | Desmethyl | Therapeut | Psychopha | Metabolite | 58534-46-( |
| 3775 | A | B | | Venlafaxin | Therapeut | Psychopha | Metabolite | 93413-69-8 |
| 524 | A | B | | Amitriptyli | Therapeut | Psychopharmaca | | 50-48-6 |
| 1172 | A | B | | Amitriptyli | Therapeut | Psychopharmaca | | 50-48-6 |
| 1269 | A | B | | Chlorproth | Therapeut | Psychopharmaca | | 113-59-7 |
| 2500 | A | B | | Chlorproth | Therapeut | Psychopharmaca | | 113-59-7 |
| 198 | A | B | C | Citalopram | Therapeut | Psychopharmaca | | 59729-33-8 |
| 3826 | A | B | | Nortriptyli | Therapeut | Psychopharmaca | | 72-69-5 |
| 504 | A | B | | O-Desmetl | Therapeut | Psychopharmaca | | 93413-62-8 |
| 3813 | A | B | | O-Desmetl | Therapeut | Psychopharmaca | | 93413-62-8 |
| 427 | A | B | | Oxazepam | Therapeut | Psychopharmaca | | 604-75-1 |
| 1522 | A | B | C | Sertraline | Therapeut | Psychopharmaca | | 79617-96-2 |
| 104 | A | B | C | Venlafaxin | Therapeut | Psychopharmaca | | 93413-69-5 |
| 400 | A | B | C | Venlafaxin | Therapeut | Psychopharmaca | | 93413-69-5 |
| 2934 | A | | | Venlafaxin | Therapeut | Psychopharmaca | | 93413-69-5 |
| 1627 | A | | | Etaqualon | Therapeut | Sedative | | 7432-25-9 |
| 6555 | A | B | | 1,4-Andros | Therapeut | Steroid | Prohormone | |
| 2245 | A | B | C | Prednison | Therapeut | Steroid medication | | 53-03-2 |
| 2034 | A | B | | Probeneci | Therapeut | Uricosuric | | 57-66-9 |

| | | | | | | | |
|---|---|---|---|---|---|---|---|
| 3346 | A | B | | 20-Dihydrc | Therapeut | Veterinary Metabolite | 6923-42-8 |
| 1620 | A | B | | Cinchophe | Therapeut | Veterinary medicine | 132-60-5 |
| 4262 | A | B | C | Mebendaz | Therapeut | Wormicide | 31431-39-[illegible] |
| 2885 | A | B | | (17(18)-Ep | Metabolite | Plant Metabolite | 131339-23 |
| 1279 | A | B | | NP-004038 | Unidentified Natural Product | | |
| 2758 | A | B | | NP-004038 | Unidentified Natural Product | | |
| 4769 | A | B | | NP-004038 | Unidentified Natural Product | | |
| 5622 | A | B | | NP-004038 | Unidentified Natural Product | | |
| 5791 | A | B | | NP-005519 | Unidentified Natural Product | | |
| 1290 | A | B | | NP-008563 | Unidentified Natural Product | | |
| 2218 | A | B | | NP-008563 | Unidentified Natural Product | | |
| 1133 | A | B | | NP-013210 | Unidentified Natural Product | | |
| 4710 | A | B | | NP-013210 | Unidentified Natural Product | | |
| 3320 | A | B | | NP-014175 | Unidentified Natural Product | | |
| 391 | A | B | | NP-015468 | Unidentified Natural Product | | |
| 951 | A | B | | NP-016354 | Unidentified Natural Product | | |
| 318 | A | B | | NP-016928 | Unidentified Natural Product | | |
| 3155 | A | B | | NP-016928 | Unidentified Natural Product | | |
| 4337 | A | B | | NP-016928 | Unidentified Natural Product | | |
| 6292 | A | B | | NP-016928 | Unidentified Natural Product | | |
| 1857 | A | B | | NP-018362 | Unidentified Natural Product | | |
| 4006 | A | B | | NP-018817 | Unidentified Natural Product | | |
| 710 | A | B | | NP-020214 | Unidentified Natural Product | | |
| 4802 | A | B | | NP-020403 | Unidentified Natural Product | | |
| 2149 | A | B | | NP-020634 | Unidentified Natural Product | | |
| 2211 | A | B | | NP-020634 | Unidentified Natural Product | | |
| 3086 | A | B | | NP-020634 | Unidentified Natural Product | | |
| 4533 | A | B | | NP-020634 | Unidentified Natural Product | | |
| 2425 | A | B | | NP-022368 | Unidentified Natural Product | | |
| 4176 | A | B | | NP-022371 | Unidentified Natural Product | | |
| 605 | A | B | | NP-022405 | Unidentified Natural Product | | |
| 5263 | A | B | | NP-022473 | Unidentified Natural Product | | |
| 15 | A | B | | | Unknown structure | | |
| 100 | A | B | | | Unknown structure | | |
| 116 | A | | | | Unknown structure | | |
| 130 | A | | | | Unknown structure | | |
| 131 | A | | | | Unknown structure | | |
| 186 | A | | | | Unknown structure | | |
| 207 | A | B | | | Unknown structure | | |
| 215 | A | | | | Unknown structure | | |
| 218 | A | | | | Unknown structure | | |
| 238 | A | | | | Unknown structure | | |
| 243 | A | B | | | Unknown structure | | |
| 260 | A | | | | Unknown structure | | |
| 275 | A | B | | | Unknown structure | | |
| 310 | A | B | | | Unknown structure | | |
| 344 | A | B | | | Unknown structure | | |
| 359 | A | B | | | Unknown structure | | |
| 417 | A | B | | | Unknown structure | | |
| 424 | A | | | | Unknown structure | | |
| 433 | A | | | | Unknown structure | | |

| | | | |
|---|---|---|---|
| 441 | A | B | Unknown structure |
| 484 | A | | Unknown structure |
| 487 | A | B | Unknown structure |
| 508 | A | B | Unknown structure |
| 536 | A | | Unknown structure |
| 580 | A | B | Unknown structure |
| 585 | A | B | Unknown structure |
| 599 | A | B | Unknown structure |
| 604 | A | B | Unknown structure |
| 606 | A | B | Unknown structure |
| 620 | A | | Unknown structure |
| 625 | A | | Unknown structure |
| 626 | A | B | Unknown structure |
| 635 | A | B | Unknown structure |
| 636 | A | B | Unknown structure |
| 653 | A | | Unknown structure |
| 660 | A | | Unknown structure |
| 662 | A | | Unknown structure |
| 686 | A | | Unknown structure |
| 688 | A | B | Unknown structure |
| 722 | A | | Unknown structure |
| 724 | A | B | Unknown structure |
| 729 | A | B | Unknown structure |
| 735 | A | B | Unknown structure |
| 738 | A | | Unknown structure |
| 756 | A | | Unknown structure |
| 778 | A | B | Unknown structure |
| 795 | A | | Unknown structure |
| 807 | A | B | Unknown structure |
| 809 | A | B | Unknown structure |
| 813 | A | | Unknown structure |
| 820 | A | B | Unknown structure |
| 868 | A | | Unknown structure |
| 872 | A | B | Unknown structure |
| 873 | A | B | Unknown structure |
| 886 | A | | Unknown structure |
| 896 | A | B | Unknown structure |
| 915 | A | | Unknown structure |
| 922 | A | | Unknown structure |
| 926 | A | B | Unknown structure |
| 940 | A | | Unknown structure |
| 956 | A | B | Unknown structure |
| 963 | A | B | Unknown structure |
| 965 | A | B | Unknown structure |
| 967 | A | B | Unknown structure |
| 995 | A | B | Unknown structure |
| 998 | A | B | Unknown structure |
| 1023 | A | B | Unknown structure |
| 1056 | A | | Unknown structure |
| 1062 | A | B | Unknown structure |
| 1108 | A | B | Unknown structure |

| | | | |
|---|---|---|---|
| 1122 | A | | Unknown structure |
| 1134 | A | B | Unknown structure |
| 1135 | A | | Unknown structure |
| 1163 | A | | Unknown structure |
| 1164 | A | B | Unknown structure |
| 1183 | A | B | Unknown structure |
| 1200 | A | | Unknown structure |
| 1229 | A | B | Unknown structure |
| 1242 | A | | Unknown structure |
| 1313 | A | | Unknown structure |
| 1316 | A | | Unknown structure |
| 1356 | A | B | Unknown structure |
| 1364 | A | B | Unknown structure |
| 1377 | A | B | Unknown structure |
| 1395 | A | | Unknown structure |
| 1437 | A | B | Unknown structure |
| 1441 | A | | Unknown structure |
| 1490 | A | B | Unknown structure |
| 1530 | A | B | Unknown structure |
| 1533 | A | | Unknown structure |
| 1591 | A | B | Unknown structure |
| 1622 | A | B | Unknown structure |
| 1625 | A | B | Unknown structure |
| 1638 | A | B | Unknown structure |
| 1685 | A | B | Unknown structure |
| 1700 | A | B | Unknown structure |
| 1705 | A | B | Unknown structure |
| 1717 | A | B | Unknown structure |
| 1720 | A | B | Unknown structure |
| 1729 | A | B | Unknown structure |
| 1754 | A | B | Unknown structure |
| 1775 | A | B | Unknown structure |
| 1832 | A | B | Unknown structure |
| 1892 | A | | Unknown structure |
| 1925 | A | B | Unknown structure |
| 1965 | A | B | Unknown structure |
| 1968 | A | B | Unknown structure |
| 1993 | A | B | Unknown structure |
| 2035 | A | B | Unknown structure |
| 2036 | A | B | Unknown structure |
| 2046 | A | B | Unknown structure |
| 2060 | A | B | Unknown structure |
| 2145 | A | B | Unknown structure |
| 2154 | A | | Unknown structure |
| 2176 | A | B | Unknown structure |
| 2193 | A | B | Unknown structure |
| 2209 | A | | Unknown structure |
| 2210 | A | B | Unknown structure |
| 2248 | A | B | Unknown structure |
| 2255 | A | B | Unknown structure |
| 2271 | A | B | Unknown structure |

| | | | |
|---|---|---|---|
| 2278 | A | | Unknown structure |
| 2284 | A | B | Unknown structure |
| 2300 | A | B | Unknown structure |
| 2312 | A | B | Unknown structure |
| 2313 | A | | Unknown structure |
| 2316 | A | B | Unknown structure |
| 2366 | A | B | Unknown structure |
| 2370 | A | B | Unknown structure |
| 2379 | A | B | Unknown structure |
| 2439 | A | B | Unknown structure |
| 2446 | A | B | Unknown structure |
| 2459 | A | B | Unknown structure |
| 2492 | A | | Unknown structure |
| 2494 | A | B | Unknown structure |
| 2497 | A | B | Unknown structure |
| 2528 | A | B | Unknown structure |
| 2547 | A | | Unknown structure |
| 2625 | A | B | Unknown structure |
| 2628 | A | B | Unknown structure |
| 2630 | A | B | Unknown structure |
| 2632 | A | B | Unknown structure |
| 2657 | A | B | Unknown structure |
| 2665 | A | B | Unknown structure |
| 2669 | A | B | Unknown structure |
| 2707 | A | | Unknown structure |
| 2712 | A | | Unknown structure |
| 2725 | A | B | Unknown structure |
| 2772 | A | B | Unknown structure |
| 2807 | A | B | Unknown structure |
| 2848 | A | | Unknown structure |
| 2895 | A | B | Unknown structure |
| 2924 | A | B | Unknown structure |
| 2929 | A | B | Unknown structure |
| 2964 | A | B | Unknown structure |
| 2976 | A | B | Unknown structure |
| 3067 | A | | Unknown structure |
| 3069 | A | B | Unknown structure |
| 3082 | A | B | Unknown structure |
| 3089 | A | B | Unknown structure |
| 3094 | A | B | Unknown structure |
| 3126 | A | B | Unknown structure |
| 3129 | A | B | Unknown structure |
| 3146 | A | B | Unknown structure |
| 3186 | A | | Unknown structure |
| 3205 | A | B | Unknown structure |
| 3208 | A | | Unknown structure |
| 3210 | A | B | Unknown structure |
| 3211 | A | B | Unknown structure |
| 3248 | A | | Unknown structure |
| 3253 | A | B | Unknown structure |
| 3257 | A | B | Unknown structure |

| | | | |
|---|---|---|---|
| 3284 | A | B | Unknown structure |
| 3339 | A | B | Unknown structure |
| 3354 | A | | Unknown structure |
| 3361 | A | | Unknown structure |
| 3382 | A | | Unknown structure |
| 3396 | A | B | Unknown structure |
| 3423 | A | B | Unknown structure |
| 3441 | A | | Unknown structure |
| 3499 | A | B | Unknown structure |
| 3504 | A | B | Unknown structure |
| 3508 | A | B | Unknown structure |
| 3522 | A | B | Unknown structure |
| 3527 | A | B | Unknown structure |
| 3559 | A | B | Unknown structure |
| 3573 | A | B | Unknown structure |
| 3576 | A | B | Unknown structure |
| 3639 | A | B | Unknown structure |
| 3657 | A | B | Unknown structure |
| 3662 | A | | Unknown structure |
| 3685 | A | | Unknown structure |
| 3714 | A | B | Unknown structure |
| 3719 | A | B | Unknown structure |
| 3736 | A | B | Unknown structure |
| 3744 | A | | Unknown structure |
| 3766 | A | B | Unknown structure |
| 3791 | A | B | Unknown structure |
| 3815 | A | B | Unknown structure |
| 3830 | A | | Unknown structure |
| 3834 | A | B | Unknown structure |
| 3859 | A | B | Unknown structure |
| 3885 | A | B | Unknown structure |
| 3945 | A | | Unknown structure |
| 3958 | A | | Unknown structure |
| 3969 | A | B | Unknown structure |
| 4016 | A | B | Unknown structure |
| 4020 | A | B | Unknown structure |
| 4027 | A | B | Unknown structure |
| 4033 | A | B | Unknown structure |
| 4121 | A | B | Unknown structure |
| 4157 | A | B | Unknown structure |
| 4158 | A | B | Unknown structure |
| 4168 | A | | Unknown structure |
| 4199 | A | B | Unknown structure |
| 4211 | A | B | Unknown structure |
| 4217 | A | B | Unknown structure |
| 4222 | A | B | Unknown structure |
| 4235 | A | B | Unknown structure |
| 4237 | A | B | Unknown structure |
| 4263 | A | | Unknown structure |
| 4319 | A | B | Unknown structure |
| 4332 | A | B | Unknown structure |

| | | | |
|---|---|---|---|
| 4343 | A | B | Unknown structure |
| 4364 | A | B | Unknown structure |
| 4390 | A | B | Unknown structure |
| 4394 | A | B | Unknown structure |
| 4426 | A | B | Unknown structure |
| 4436 | A | B | Unknown structure |
| 4453 | A | B | Unknown structure |
| 4460 | A | B | Unknown structure |
| 4489 | A | B | Unknown structure |
| 4500 | A | B | Unknown structure |
| 4530 | A | B | Unknown structure |
| 4563 | A | B | Unknown structure |
| 4594 | A | B | Unknown structure |
| 4621 | A | B | Unknown structure |
| 4625 | A | B | Unknown structure |
| 4645 | A | B | Unknown structure |
| 4647 | A | B | Unknown structure |
| 4665 | A | B | Unknown structure |
| 4704 | A | B | Unknown structure |
| 4720 | A | B | Unknown structure |
| 4723 | A | B | Unknown structure |
| 4763 | A | | Unknown structure |
| 4771 | A | B | Unknown structure |
| 4782 | A | B | Unknown structure |
| 4789 | A | B | Unknown structure |
| 4806 | A | | Unknown structure |
| 4825 | A | | Unknown structure |
| 4831 | A | B | Unknown structure |
| 4839 | A | B | Unknown structure |
| 4845 | A | | Unknown structure |
| 4889 | A | B | Unknown structure |
| 4905 | A | B | Unknown structure |
| 4930 | A | B | Unknown structure |
| 5005 | A | B | Unknown structure |
| 5022 | A | B | Unknown structure |
| 5030 | A | B | Unknown structure |
| 5031 | A | B | Unknown structure |
| 5083 | A | B | Unknown structure |
| 5099 | A | | Unknown structure |
| 5205 | A | B | Unknown structure |
| 5230 | A | B | Unknown structure |
| 5237 | A | | Unknown structure |
| 5252 | A | B | Unknown structure |
| 5256 | A | | Unknown structure |
| 5289 | A | | Unknown structure |
| 5291 | A | B | Unknown structure |
| 5321 | A | | Unknown structure |
| 5416 | A | B | Unknown structure |
| 5439 | A | B | Unknown structure |
| 5494 | A | B | Unknown structure |
| 5524 | A | B | Unknown structure |

| | | | | |
|---|---|---|---|---|
| 5528 | A | B | | Unknown structure |
| 5539 | A | B | | Unknown structure |
| 5566 | A | B | | Unknown structure |
| 5649 | A | | | Unknown structure |
| 5678 | A | B | | Unknown structure |
| 5729 | A | B | | Unknown structure |
| 5739 | A | B | | Unknown structure |
| 5752 | A | B | | Unknown structure |
| 5798 | A | | | Unknown structure |
| 5824 | A | B | | Unknown structure |
| 5854 | A | B | | Unknown structure |
| 5870 | A | B | | Unknown structure |
| 5872 | A | | | Unknown structure |
| 5891 | A | B | | Unknown structure |
| 5897 | A | B | | Unknown structure |
| 5949 | A | B | | Unknown structure |
| 5958 | A | | | Unknown structure |
| 5987 | A | B | | Unknown structure |
| 5992 | A | | | Unknown structure |
| 6009 | A | B | | Unknown structure |
| 6044 | A | | | Unknown structure |
| 6050 | A | B | | Unknown structure |
| 6074 | A | B | | Unknown structure |
| 6089 | A | | | Unknown structure |
| 6136 | A | B | | Unknown structure |
| 6303 | A | B | | Unknown structure |
| 6347 | A | | | Unknown structure |
| 6384 | A | | | Unknown structure |
| 6607 | A | | | Unknown structure |
| 6665 | A | B | | Unknown structure |
| 6667 | A | B | | Unknown structure |
| 1123 | A | B | (10E)-10-P | Unidentified Natural Product |
| 2808 | A | B | (10E)-10-P | Unidentified Natural Product |
| 4025 | A | B | (10E)-10-P | Unidentified Natural Product |
| 3231 | A | B | (10Z)-2-Hy | Unidentified Natural Product |
| 3525 | A | B | (10Z)-2-Hy | Unidentified Natural Product |
| 4380 | A | B | (10Z)-2-Hy | Unidentified Natural Product |
| 595 | A | B | (11Z)-11-H | Unidentified Natural Product |
| 6348 | A | B | (11Z)-11-H | Unidentified Natural Product |
| 1151 | A | B | (1aR,3Z,5R | Unidentified Natural Product |
| 2389 | A | B | (1aR,3Z,5R | Unidentified Natural Product |
| 3516 | A | B | (1aR,3Z,5R | Unidentified Natural Product |
| 4044 | A | B | (1aR,3Z,5R | Unidentified Natural Product |
| 2456 | A | B | (1aR,6S,7a | Unidentified Natural Product |
| 3 | A | B | (1R,3aS,5a | Unidentified Natural Product |
| 88 | A | B | (1R,3aS,5a | Unidentified Natural Product |
| 1094 | A | B | (1R,3R,4S,5 | Unidentified Natural Product |
| 952 | A | B | (1R,5S,6R)- | Unidentified Natural Product |
| 1787 | A | B | (1S)-1-[(1S | Unidentified Natural Product |
| 4256 | A | B | (1S,2S,4R,7 | Unidentified Natural Product |
| 5115 | A | B | (1S,2S,4R,7 | Unidentified Natural Product |

| | | | | | |
|---|---|---|---|---|---|
| 1480 | A | B | (1S,4S,5R,8 | Unidentified Natural Product | |
| 26 | A | B | (1S,5S)-1,5 | Unidentified Natural Product | |
| 3928 | A | B | (1S,6R,11a | Unidentified Natural Product | |
| 5235 | A | B | (1S,6R,11a | Unidentified Natural Product | |
| 3943 | A | | (2,6-dimet | Unidentified Natural Product | |
| 4573 | A | B | (2E)-1-(2-H | Unidentified Natural Product | |
| 5801 | A | B | (2E)-3-[(1R | Unidentified Natural Product | |
| 2477 | A | | (2E)-3-{2-B | Unidentified Natural Product | |
| 4241 | A | B | (2E)-N-[2-( | Unidentified Natural Product | |
| 4714 | A | B | (2E,4E)-5-[ | Unidentified Natural Product | |
| 3457 | A | B | (2E,4E,6Z)- | Unidentified Natural Product | |
| 368 | A | B | (2R)-5-Met | Unidentified Natural Product | |
| 5942 | A | | (2R,3R,4R,! | Unidentified Natural Product | |
| 2935 | A | | (2S,3S,4S,5 | Unidentified Natural Product | |
| 4328 | A | B | (3aR,4aS,5 | Unidentified Natural Product | |
| 5496 | A | B | (3aR,4aS,5 | Unidentified Natural Product | |
| 2903 | A | B | (3aR,5aR,9 | Unidentified Natural Product | |
| 1393 | A | B | (3aR,8aR,9 | Unidentified Natural Product | |
| 4162 | A | B | (3aS,11aR) | Unidentified Natural Product | |
| 1192 | A | B | (3aS,6aR)-: | Unidentified Natural Product | |
| 1483 | A | B | (3aS,6aR)-: | Unidentified Natural Product | |
| 2454 | A | B | (3R)-12-Hy | Unidentified Natural Product | |
| 2959 | A | B | (3R,3aR,4E | Unidentified Natural Product | |
| 3889 | A | B | (3R,3aR,4E | Unidentified Natural Product | |
| 1837 | A | B | (3R,3aR,4R | Unidentified Natural Product | |
| 726 | A | B | (3R,4R)-1-[ | Unidentified Natural Product | |
| 2298 | A | B | (3S)-8,8-Di | Unidentified Natural Product | |
| 936 | A | B | (3S,6R,6aS | Unidentified Natural Product | |
| 2111 | A | B | (4aS,5R,6S | Unidentified Natural Product | |
| 2563 | A | B | (4R)-6-{2-[ | Unidentified Natural Product | |
| 4203 | A | B | (4S)-3-(Cyc | Unidentified Natural Product | |
| 3049 | A | B | (4S)-4-[(1R | Unidentified Natural Product | |
| 3510 | A | | (4S)-4-[(1R | Unidentified Natural Product | |
| 3431 | A | | (4S)-4-Hyd | Unidentified Natural Product | |
| 788 | A | B | (5R)-5-(4-{ | Unidentified Natural Product | |
| 2852 | A | B | (5R)-5-(4-{ | Unidentified Natural Product | |
| 878 | A | | (6a)-11,12- | Unidentified Natural Product | |
| 1815 | A | B | (6S,16R)-6- | Unidentified Natural Product | |
| 2527 | A | B | (6S,6aS,12 | Unidentified Natural Product | |
| 2126 | A | B | [(1R,5S,6r) | Unidentified Natural Product | |
| 4790 | A | B | [(2R,6R)-6- | Unidentified Natural Product | |
| 6639 | A | B | [(4aS,5aS,6 | Unidentified Natural Product | |
| 4351 | A | B | {(1R,3R,4R | Unidentified Natural Product | |
| 2793 | A | B | 1-(2-Metho | Unidentified Natural Product | |
| 134 | A | B | 1-(3,5-Dite | Unidentified Natural Product | 14035-33-7 |
| 235 | A | B | 1-(3,5-Dite | Unidentified Natural Product | 14035-33-7 |
| 737 | A | B | 1-(3,5-Dite | Unidentified Natural Product | 14035-33-7 |
| 3911 | A | | 1-(4-Fluoro | Unidentified Natural Product | |
| 4249 | A | B | 1-(Butylsul | Unidentified Natural Product | |
| 719 | A | B | 1,1'-[4-(All | Unidentified Natural Product | |
| 78 | A | | 1-[4-Fluoro | Unidentified Natural Product | |

| | | | | | |
|---|---|---|---|---|---|
| 2153 | A | | 1-{4-[(2R,3 | Unidentified Natural Product | |
| 2759 | A | | 1-{4-[(2R,3 | Unidentified Natural Product | |
| 4507 | A | B | 1-{4-Chlor( | Unidentified Natural Product | |
| 2212 | A | B | 11-[4-(Ben | Unidentified Natural Product | |
| 2854 | A | B | 1-amino-9- | Unidentified Natural Product | |
| 4810 | A | B | 2-(11,17-D | Unidentified Natural Product | |
| 164 | A | B | 2-(2-Meth( | Unidentified Natural Product | |
| 782 | A | | 2-(5-{(Z)-[1 | Unidentified Natural Product | |
| 385 | A | B | 2-(5-{1-[3-( | Unidentified Natural Product | |
| 5931 | A | B | 2-(Phenyls | Unidentified Natural Product | 4273-98-7 |
| 146 | A | B | 2,3-dihydr( | Unidentified Natural Product | |
| 147 | A | B | 2,3-dihydr( | Unidentified Natural Product | |
| 211 | A | B | 2,3-dihydr( | Unidentified Natural Product | |
| 1487 | A | B | 2,3-dihydr( | Unidentified Natural Product | |
| 1667 | A | B | 2,3-dihydr( | Unidentified Natural Product | |
| 1581 | A | B | 2,6-Difluor | Unidentified Natural Product | |
| 4080 | A | | 2,6-Dimetl | Unidentified Natural Product | |
| 4591 | A | B | 2-[(2R)-5-A | Unidentified Natural Product | |
| 4074 | A | B | 2-[(5R,8S,8 | Unidentified Natural Product | |
| 3507 | A | B | 2-[(Dimeth | Unidentified Natural Product | |
| 5311 | A | B | 2-[(Dimeth | Unidentified Natural Product | |
| 4440 | A | | 2-[2-({4-Ar | Unidentified Natural Product | |
| 5408 | A | B | 2-[2-(Dieth | Unidentified Natural Product | |
| 4236 | A | B | 2-Ethylhex | Unidentified Natural Product | |
| 137 | A | | 2-Fluoro-N | Unidentified Natural Product | |
| 4855 | A | B | 2-Isopropy | Unidentified Natural Product | |
| 2948 | A | B | 2-Methoxy | Unidentified Natural Product | |
| 6120 | A | | 2-Methyl-2 | Unidentified Natural Product | |
| 1460 | A | | 2-Phenyl-1 | Unidentified Natural Product | |
| 5624 | A | B | 3-(1-Hydro | Unidentified Natural Product | |
| 421 | A | | 3-(beta-D-( | Unidentified Natural Product | |
| 3748 | A | B | 3-(Dimethy | Unidentified Natural Product | 104753-60 |
| 2042 | A | B | 3-{3-[4-(4-I | Unidentified Natural Product | |
| 2538 | A | | 3-Acetyl-6- | Unidentified Natural Product | |
| 1932 | A | B | 3-amino-2- | Unidentified Natural Product | 74037-32-4 |
| 423 | A | B | 3-benzyl-4- | Unidentified Natural Product | |
| 800 | A | B | 3-Hexyl-4-i | Unidentified Natural Product | |
| 775 | A | B | 3-hydroxy- | Unidentified Natural Product | |
| 2051 | A | B | 3-hydroxy- | Unidentified Natural Product | |
| 2513 | A | B | 3-hydroxy- | Unidentified Natural Product | |
| 5167 | A | B | 3-Hydroxy- | Unidentified Natural Product | |
| 546 | A | | 3-Isoprope | Unidentified Natural Product | |
| 3240 | A | B | 3-Methyle | Unidentified Natural Product | |
| 1835 | A | B | 4-(tert-but | Unidentified Natural Product | |
| 3179 | A | B | 4-(tert-but | Unidentified Natural Product | |
| 4785 | A | B | 4,4'-Sulfan | Unidentified Natural Product | |
| 1607 | A | | 4-[(3S)-7-h | Unidentified Natural Product | |
| 1041 | A | B | 4-[(Adama | Unidentified Natural Product | |
| 4627 | A | B | 4-[2-(3,5-d | Unidentified Natural Product | |
| 1735 | A | | 4-[3-(2-Nit | Unidentified Natural Product | |
| 5045 | A | B | 4-[Hexade( | Unidentified Natural Product | |

| | | | | | |
|---|---|---|---|---|---|
| 2116 | A | B | 4-Biphenyl | Unidentified Natural Product | |
| 1654 | A | B | 4-carbamo | Unidentified Natural Product | |
| 514 | A | | 4-Ethyl-2-[ | Unidentified Natural Product | |
| 296 | A | | 4-heptyl-3- | Unidentified Natural Product | |
| 3148 | A | B | 4-Isopropo | Unidentified Natural Product | |
| 4371 | A | B | 4-Isopropy | Unidentified Natural Product | |
| 4401 | A | B | 4-Isopropy | Unidentified Natural Product | |
| 4577 | A | B | 4-Isopropy | Unidentified Natural Product | |
| 2853 | A | | 4-Methoxy | Unidentified Natural Product | 174680-14 |
| 2674 | A | B | 5-(2-{[3-(D | Unidentified Natural Product | |
| 5406 | A | B | 5-(4,5-Dihy | Unidentified Natural Product | |
| 1376 | A | B | 5-(Cyclooc | Unidentified Natural Product | |
| 1513 | A | B | 5-(Cyclooc | Unidentified Natural Product | |
| 5676 | A | B | 5,7-Dihydr | Unidentified Natural Product | |
| 3417 | A | B | 5-[2-(3-Fur | Unidentified Natural Product | |
| 4264 | A | B | 5-[3-(3-He | Unidentified Natural Product | |
| 76 | A | B | 5-[5-(meth | Unidentified Natural Product | |
| 2283 | A | | 5-Amino-1 | Unidentified Natural Product | |
| 2095 | A | | 5-Fluoro-N | Unidentified Natural Product | |
| 5573 | A | B | 5-Hydroxy- | Unidentified Natural Product | |
| 4998 | A | B | 5-Propyl-9 | Unidentified Natural Product | |
| 6327 | A | B | 6-(4-Fluoro | Unidentified Natural Product | |
| 2148 | A | B | 6-(4-pheny | Unidentified Natural Product | |
| 3246 | A | B | 6-(4-pheny | Unidentified Natural Product | |
| 3962 | A | B | 6-(4-pheny | Unidentified Natural Product | |
| 2936 | A | B | 6-Methoxy | Unidentified Natural Product | 58176-56-( |
| 1241 | A | B | 6-phenyl-3 | Unidentified Natural Product | |
| 5692 | A | B | 8-(Ethoxyn | Unidentified Natural Product | |
| 1990 | A | B | 8-[(4-Hydr | Unidentified Natural Product | |
| 1565 | A | B | 8-Isobutyl- | Unidentified Natural Product | |
| 5280 | A | | 9-(2-Chlor | Unidentified Natural Product | |
| 750 | A | B | 9,16-dihyd | Unidentified Natural Product | |
| 732 | A | B | 9a-Ethynyl | Unidentified Natural Product | |
| 2027 | A | B | 9-hydroxy- | Unidentified Natural Product | |
| 2357 | A | B | 9-hydroxy- | Unidentified Natural Product | |
| 2977 | A | B | 9-hydroxy- | Unidentified Natural Product | |
| 3537 | A | B | 9-hydroxy- | Unidentified Natural Product | |
| 3450 | A | | Benzo[kl]x | Unidentified Natural Product | 200-23-7 |
| 2418 | A | B | Benzyl oct | Unidentified Natural Product | |
| 6015 | A | B | D-Alanyl-N | Unidentified Natural Product | |
| 2141 | A | B | deoxoscala | Unidentified Natural Product | |
| 3939 | A | B | deoxoscala | Unidentified Natural Product | |
| 1962 | A | B | Diethyl (3- | Unidentified Natural Product | |
| 3446 | A | B | Diisopropy | Unidentified Natural Product | |
| 4391 | A | B | Diisopropy | Unidentified Natural Product | |
| 1331 | A | B | Dodecyl 2- | Unidentified Natural Product | |
| 1684 | A | B | Dodecyl 2- | Unidentified Natural Product | |
| 2159 | A | B | Ethyl 1-{2- | Unidentified Natural Product | |
| 4302 | A | B | Ethyl 1-oxo | Unidentified Natural Product | 16266-88-9 |
| 1351 | A | | ethyl 2-{[5- | Unidentified Natural Product | |
| 1138 | A | | Ethyl 3-hyd | Unidentified Natural Product | |

| | | | | | |
|---|---|---|---|---|---|
| 1228 | A | B | Ethyl 5-ber | Unidentified Natural Product | 134575-05 |
| 1257 | A | | Isobutyl [(4 | Unidentified Natural Product | |
| 3689 | A | B | mesityl (4- | Unidentified Natural Product | 5184-64-5 |
| 639 | A | | Methyl (1R | Unidentified Natural Product | |
| 528 | A | | Methyl (2,! | Unidentified Natural Product | |
| 53 | A | B | Methyl (4a | Unidentified Natural Product | |
| 473 | A | | Methyl 2,7 | Unidentified Natural Product | |
| 6226 | A | B | Methyl 2-[ | Unidentified Natural Product | |
| 4226 | A | B | Methyl N-( | Unidentified Natural Product | |
| 1937 | A | B | Methyl N-[ | Unidentified Natural Product | |
| 5719 | A | B | Methyl N-[ | Unidentified Natural Product | |
| 2641 | A | | N-(2-{[(2S) | Unidentified Natural Product | |
| 4690 | A | B | N-(3-Amin | Unidentified Natural Product | |
| 2561 | A | B | N,N-Diethy | Unidentified Natural Product | |
| 2898 | A | B | N,N-Diethy | Unidentified Natural Product | |
| 2243 | A | | N,N-Diethy | Unidentified Natural Product | |
| 3152 | A | | N,N-Diethy | Unidentified Natural Product | |
| 751 | A | B | N-[(1-Acet | Unidentified Natural Product | |
| 1184 | A | B | N-[4-(4-me | Unidentified Natural Product | |
| 4695 | A | B | N-[4-(4-me | Unidentified Natural Product | |
| 3385 | A | | N-{[(2R,4S, | Unidentified Natural Product | |
| 887 | A | B | N-{[4-(3-Fl | Unidentified Natural Product | |
| 87 | A | B | N-{4-[(2R,3 | Unidentified Natural Product | |
| 2160 | A | B | N-{4-[(2R,3 | Unidentified Natural Product | |
| 5560 | A | B | N-{4-[4-(3, | Unidentified Natural Product | |
| 401 | A | B | N1-[4-(ace | Unidentified Natural Product | |
| 395 | A | | N2-(2,2-Di | Unidentified Natural Product | |
| 3918 | A | | N2-Benzoy | Unidentified Natural Product | |
| 1823 | A | | N-Benzyl-N | Unidentified Natural Product | |
| 3091 | A | B | N-Cyclohex | Unidentified Natural Product | |
| 3707 | A | B | N-Hydroxy | Unidentified Natural Product | |

| Reference_classification |
| --- |
| https://pubchem.ncbi.nlm.nih.gov/compound/S_-Dibenzyl-2-aminosuccinate |
| https://www.scbt.com/p/4-methylbenzotriazole-29878-31-7 |
| https://www.scbt.com/p/4-methylbenzotriazole-29878-31-7 |
| https://en.wikipedia.org/wiki/Fluorescein |
| https://www.sigmaaldrich.com/DK/en/product/aldrich/255327 |
| https://www.sigmaaldrich.com/DK/en/product/aldrich/255327 |
| https://pubchem.ncbi.nlm.nih.gov/compound/6540 |
| https://www.sigmaaldrich.com/DK/en/product/aldrich/119660 |
| https://www.sigmaaldrich.com/DK/en/product/aldrich/906808 |
| https://en.wikipedia.org/wiki/Benzyl_butyl_phthalate |
| https://pubchem.ncbi.nlm.nih.gov/compound/Bis_2-butoxyethyl_-phthalate |
| https://en.wikipedia.org/wiki/Bisphenol_A |
| https://en.wikipedia.org/wiki/Centralite |
| https://www.eastman.com/Pages/ProductHome.aspx?product=71071434 |
| https://www.eastman.com/Pages/ProductHome.aspx?product=71071434 |
| https://www.eastman.com/Pages/ProductHome.aspx?product=71071434 |
| https://en.wikipedia.org/wiki/Tributyl_phosphate |
| https://pubchem.ncbi.nlm.nih.gov/compound/118719907#section=PFAS-and-Fl |
| https://en.wikipedia.org/wiki/Monolaurin |
| https://pubchem.ncbi.nlm.nih.gov/compound/7564 |
| https://pubchem.ncbi.nlm.nih.gov/compound/59018797 |
| https://en.wikipedia.org/wiki/Corticosterone |
| https://en.wikipedia.org/wiki/Corticosterone |
| https://en.wikipedia.org/wiki/Corticosterone |
| https://en.wikipedia.org/wiki/Corticosterone |
| https://en.wikipedia.org/wiki/Cortisol |
| https://en.wikipedia.org/wiki/Cortisol |
| https://en.wikipedia.org/wiki/Eicosapentaenoic_acid |
| https://www.caymanchem.com/product/9000295/eicosapentaenoic-acid-meth |
| https://www.lampreydb.com/material/769/view |
| https://pubchem.ncbi.nlm.nih.gov/compound/45359480; https://medkoo.com/ |
| https://pubchem.ncbi.nlm.nih.gov/compound/16745396 |
| https://pubchem.ncbi.nlm.nih.gov/compound/16745396 |
| https://pubchem.ncbi.nlm.nih.gov/compound/16745396 |
| https://pubchem.ncbi.nlm.nih.gov/compound/145975830 |
| https://pubchem.ncbi.nlm.nih.gov/compound/Verrucarol |
| https://pubchem.ncbi.nlm.nih.gov/compound/Verrucarol |
| https://pubchem.ncbi.nlm.nih.gov/compound/Verrucarol |
| https://pubchem.ncbi.nlm.nih.gov/compound/Verrucarol |
| https://pubchem.ncbi.nlm.nih.gov/compound/Verrucarol |

https://pubchem.ncbi.nlm.nih.gov/compound/115116#section=Molecular-Form
https://pubchem.ncbi.nlm.nih.gov/compound/Hydroxyprogesterone#section=M
https://pubchem.ncbi.nlm.nih.gov/compound/2-Methoxyestrone
https://pubchem.ncbi.nlm.nih.gov/compound/6beta-Hydroxytestosterone
https://pubchem.ncbi.nlm.nih.gov/compound/7alpha-Hydroxytestosterone
https://doi.org/10.1523/JNEUROSCI.3562-07.2008
https://doi.org/10.1523/JNEUROSCI.3562-07.2008
https://doi.org/10.1523/JNEUROSCI.3562-07.2008
https://en.wikipedia.org/wiki/Aldosterone
https://en.wikipedia.org/wiki/Estriol
https://en.wikipedia.org/wiki/Estriol
https://en.wikipedia.org/wiki/Pregnenolone
https://en.wikipedia.org/wiki/Pregnenolone
https://en.wikipedia.org/wiki/Tetrahydrocortisone
https://www.salivaryproteome.org/public/index.php/Special:Ontology_Term/Cl
https://en.wikipedia.org/wiki/5-Oxo-eicosatetraenoic_acid
https://en.wikipedia.org/wiki/5-Oxo-eicosatetraenoic_acid
https://www.caymanchem.com/product/17327/coenzyme-q2
https://www.caymanchem.com/product/17327/coenzyme-q2
https://www.caymanchem.com/product/17327/coenzyme-q2
https://www.caymanchem.com/product/17327/coenzyme-q2
https://pubchem.ncbi.nlm.nih.gov/compound/4248455
https://pubchem.ncbi.nlm.nih.gov/compound/10469728#:~:text=13%2DHOTrE%
https://pubchem.ncbi.nlm.nih.gov/compound/10469728#:~:text=13%2DHOTrE%
https://en.wikipedia.org/wiki/Sedanolide
https://pubchem.ncbi.nlm.nih.gov/compound/Senkyunolide-H
https://pubchem.ncbi.nlm.nih.gov/compound/Tanshinone-IIA
https://doi.org/10.1021/jo0258204
https://en.wikipedia.org/wiki/Piperine
https://pubchem.ncbi.nlm.nih.gov/compound/Pinusolide
https://journals.sagepub.com/doi/10.3109/10915818409010517 ; https://pubcl
https://www.chemicalbook.com/Article/What-is-7-9-ditert-butyl-1-oxaspiro-4-5
https://doi.org/10.1016/j.phyplu.2021.100183
https://jglobal.jst.go.jp/en/detail?JGLOBAL_ID=200907081939703640
https://jglobal.jst.go.jp/en/detail?JGLOBAL_ID=200907030666579816
https://jglobal.jst.go.jp/en/detail?JGLOBAL_ID=200907030666579816
https://pubchem.ncbi.nlm.nih.gov/compound/11043090
https://jglobal.jst.go.jp/en/detail?JGLOBAL_ID=200907088143107765#%7B%22
https://en.wikipedia.org/wiki/Nootkatone
https://pubchem.ncbi.nlm.nih.gov/compound/Psilostachyin-B#section=Synonyr
https://pubchem.ncbi.nlm.nih.gov/compound/Psilostachyin-B#section=Synonyr
https://pubchem.ncbi.nlm.nih.gov/compound/Psilostachyin-B#section=Synonyr
https://pubchem.ncbi.nlm.nih.gov/compound/Zedoarondiol
https://pubchem.ncbi.nlm.nih.gov/compound/Zedoarondiol
https://pubchem.ncbi.nlm.nih.gov/compound/3beta-Hydroxypregn-5-en-20-one
DOI:10.1002/hlca.200790209
https://www.sigmaaldrich.com/DK/en/product/sigma/smb00244
https://pubchem.ncbi.nlm.nih.gov/compound/14806212
https://pubchem.ncbi.nlm.nih.gov/compound/14806212
https://pubchem.ncbi.nlm.nih.gov/compound/2-_Methylthio_benzothiazole#se
https://pubchem.ncbi.nlm.nih.gov/compound/73832277

https://en.wikipedia.org/wiki/Abietic_acid
https://pubchem.ncbi.nlm.nih.gov/compound/Arnicolide-A
https://www.mdpi.com/1420-3049/24/10/1908
https://pubchem.ncbi.nlm.nih.gov/compound/177267
https://pubchem.ncbi.nlm.nih.gov/compound/177267
https://en.wikipedia.org/wiki/Celastrol
https://pubchem.ncbi.nlm.nih.gov/compound/Dipropyl-phthalate#::text=Dipro
https://pubchem.ncbi.nlm.nih.gov/compound/6466
https://drugs.ncats.io/drug/5K0WKD513R
https://pubchem.ncbi.nlm.nih.gov/compound/114778
https://pubchem.ncbi.nlm.nih.gov/compound/Norisoguaiacin
https://pubchem.ncbi.nlm.nih.gov/compound/8364
https://pubchem.ncbi.nlm.nih.gov/compound/Ostruthin
https://pubchem.ncbi.nlm.nih.gov/compound/5282365
https://pubchem.ncbi.nlm.nih.gov/compound/5282365
https://www.trc-canada.com/product-detail/?G189005 ; https://pubchem.ncbi.
https://www.trc-canada.com/product-detail/?G189005 ; https://pubchem.ncbi.
https://www.trc-canada.com/product-detail/?G189005 ; https://pubchem.ncbi.
https://en.wikipedia.org/wiki/Enzacamene
https://en.wikipedia.org/wiki/Propiconazole
https://en.wikipedia.org/wiki/Tebuconazole
https://en.wikipedia.org/wiki/DEET
https://pubchem.ncbi.nlm.nih.gov/compound/43359
https://pubchem.ncbi.nlm.nih.gov/compound/62020
https://pubchem.ncbi.nlm.nih.gov/compound/4929
https://doi.org/10.1093/nar/27.2.674
https://pubchem.ncbi.nlm.nih.gov/compound/n-_1-Ethoxycarbonyl-3-phenylpr
https://www.bldpharm.com/products/5408-46-8.html
https://www.bldpharm.com/products/5408-46-8.html
https://www.scbt.com/p/2-7-diacetyl-fluorene-961-27-3
https://www.vulcanchem.com/product/main-products/131298-44-7
https://www.trc-canada.com/product-detail/?B206660
https://www.trc-canada.com/product-detail/?B408375
https://pubchem.ncbi.nlm.nih.gov/compound/Bis_2-ethylhexyl_amine
https://pubchem.ncbi.nlm.nih.gov/compound/Bis_2-ethylhexyl_amine
https://chem-space.com/CSSS00159283679-D4FFF9
https://www.sigmaaldrich.com/DK/en/product/aldrich/68572
https://pubchem.ncbi.nlm.nih.gov/compound/Dodecylamine
https://pubchem.ncbi.nlm.nih.gov/compound/15793
https://pubchem.ncbi.nlm.nih.gov/compound/3_3_-Dimethoxybenzidine
https://en.wikipedia.org/wiki/Michler%27s_ketone
https://pubchem.ncbi.nlm.nih.gov/compound/Octyl-phenol-ethoxylate
https://pubchem.ncbi.nlm.nih.gov/compound/Octyl-phenol-ethoxylate
https://en.wikipedia.org/wiki/Neostigmine
https://doi.org/10.1016/j.bmc.2011.05.022
https://patents.google.com/patent/US9365502B2/en
https://en.wikipedia.org/wiki/Tramadol
https://en.wikipedia.org/wiki/Diclofenac
https://en.wikipedia.org/wiki/Etodolac
https://en.wikipedia.org/wiki/Indometacin
https://pubchem.ncbi.nlm.nih.gov/compound/3825

https://pubchem.ncbi.nlm.nih.gov/compound/3825
https://en.wikipedia.org/wiki/Naproxen
https://en.wikipedia.org/wiki/Phenazone
https://en.wikipedia.org/wiki/Prodine
https://en.wikipedia.org/wiki/Pethidine
https://en.wikipedia.org/wiki/Methadone
https://en.wikipedia.org/wiki/Normorphine
https://en.wikipedia.org/wiki/Tapentadol
https://en.wikipedia.org/wiki/Tramadol
https://en.wikipedia.org/wiki/Tramadol
https://en.wikipedia.org/wiki/Phendimetrazine
https://en.wikipedia.org/wiki/Clopidogrel
https://proceedings.science/brazmedchem-2019/papers/analysis-of-the-bacteri
https://pubchem.ncbi.nlm.nih.gov/compound/447043
https://en.wikipedia.org/wiki/Clarithromycin
https://en.wikipedia.org/wiki/Roxithromycin
https://en.wikipedia.org/wiki/Sutezolid
https://en.wikipedia.org/wiki/Trimethoprim
https://www.drugs.com/mtm/benztropine.html
https://en.wikipedia.org/wiki/Rivaroxaban
https://go.drugbank.com/metabolites/DBMET02803
https://pubchem.ncbi.nlm.nih.gov/compound/Licarbazepine
https://pubchem.ncbi.nlm.nih.gov/compound/Licarbazepine
DOI: 10.1097/00007691-199812000-00012
DOI: 10.1097/00007691-199812000-00012
https://www.drugs.com/carbamazepine.html
https://pubchem.ncbi.nlm.nih.gov/compound/3878
https://en.wikipedia.org/wiki/Oxcarbazepine
https://doi.org/10.1021/jm060807n
https://en.wikipedia.org/wiki/Climbazole
https://en.wikipedia.org/wiki/Terbinafine
https://www.drugs.com/cetirizine-hcl.html
https://en.wikipedia.org/wiki/Fexofenadine
https://pubchem.ncbi.nlm.nih.gov/compound/26133
https://bpspubs.onlinelibrary.wiley.com/doi/full/10.1111/bph.15741
https://en.wikipedia.org/wiki/Crotamiton
https://doi.org/10.1021/jm00241a015
https://doi.org/10.1124/jpet.104.079079
https://doi.org/10.1046/j.1432-1327.2001.02198.x
https://www.sciencedirect.com/science/article/pii/S0166445X14003956
https://en.wikipedia.org/wiki/Megestrol_acetate
https://en.wikipedia.org/wiki/Alfuzosin
https://en.wikipedia.org/wiki/Flusoxolol
https://en.wikipedia.org/wiki/Metoprolol
https://en.wikipedia.org/wiki/Metoprolol
https://en.wikipedia.org/wiki/Propranolol
https://www.sigmaaldrich.com/DK/en/substance/verapamilrelatedcompounda
https://en.wikipedia.org/wiki/Bisoprolol
https://en.wikipedia.org/wiki/Candesartan
https://en.wikipedia.org/wiki/Diltiazem
https://en.wikipedia.org/wiki/Flecainide

https://en.wikipedia.org/wiki/Labetalol
https://en.wikipedia.org/wiki/Losartan
https://en.wikipedia.org/wiki/Dicyclohexylurea
https://en.wikipedia.org/wiki/Valsartan
https://www.ndrugs.com/?s=decholin
https://www.webmd.com/drugs/2/drug-152860/fenofibric-acid-oral/details
https://en.wikipedia.org/wiki/Gemfibrozil
https://en.wikipedia.org/wiki/Benzphetamine
https://en.wikipedia.org/wiki/Xylometazoline
https://pubchem.ncbi.nlm.nih.gov/compound/3440
https://doi.org/10.1373/clinchem.2008.122119
https://doi.org/10.1093/jat/23.2.81
https://pubmed.ncbi.nlm.nih.gov/2046334/
https://en.wikipedia.org/wiki/11-Nor-9-carboxy-THC
https://en.wikipedia.org/wiki/Methyldienolone
https://en.wikipedia.org/wiki/RCS-4
https://www.caymanchem.com/product/21742/(%C2%B1)-cannabichromeorcir
https://www.caymanchem.com/product/17945/5%CE%B1-androstan-3%2C6%2
https://www.caymanchem.com/product/17945/5%CE%B1-androstan-3%2C6%2
https://www.caymanchem.com/product/17945/5%CE%B1-androstan-3%2C6%2
https://www.caymanchem.com/product/17084
https://www.caymanchem.com/product/ISO00130/acetyl-norfentanyl-(hydroch
https://en.wikipedia.org/wiki/2,5-Dimethoxy-4-amylamphetamine
https://en.wikipedia.org/wiki/HU-331
https://en.wikipedia.org/wiki/HU-331
https://en.wikipedia.org/wiki/HU-331
https://doi.org/10.1007/s00706-015-1426-7
https://en.wikipedia.org/wiki/Mycophenolic_acid
https://patents.google.com/patent/US6962938B2/en
https://en.wikipedia.org/wiki/Norgestrel
https://patents.google.com/patent/US4020177A/en
https://en.wikipedia.org/wiki/Desmethylcitalopram
https://en.wikipedia.org/wiki/Nordoxepin
https://en.wikipedia.org/wiki/Venlafaxine
https://en.wikipedia.org/wiki/Amitriptyline
https://en.wikipedia.org/wiki/Amitriptyline
https://en.wikipedia.org/wiki/Chlorprothixene
https://en.wikipedia.org/wiki/Chlorprothixene
https://en.wikipedia.org/wiki/Citalopram
https://en.wikipedia.org/wiki/Nortriptyline
https://en.wikipedia.org/wiki/Desvenlafaxine
https://en.wikipedia.org/wiki/Desvenlafaxine
https://en.wikipedia.org/wiki/Oxazepam
https://en.wikipedia.org/wiki/Sertraline
https://en.wikipedia.org/wiki/Venlafaxine
https://en.wikipedia.org/wiki/Venlafaxine
https://en.wikipedia.org/wiki/Venlafaxine
https://en.wikipedia.org/wiki/Etaqualone
https://pubmed.ncbi.nlm.nih.gov/16308875/
https://en.wikipedia.org/wiki/Prednisone
https://en.wikipedia.org/wiki/Probenecid

https://doi.org/10.1002/dta.1599
https://en.wikipedia.org/wiki/Cinchophen
https://en.wikipedia.org/wiki/Mebendazole
https://pubchem.ncbi.nlm.nih.gov/compound/16061089

7
7
7

https://pubchem.ncbi.nlm.nih.gov/compound/363819

-8

4

https://www.chemsrc.com/en/cas/174680-14-9_1641306.html#ebiemingDiv

)

9

| Recovered_as | Formula | Annot_Source_Predicted_Composi | Annot_Source_mzCloud_Search | Annot_Source_mzVault_Search | Annot_Source_ChemSpider_Search |
|---|---|---|---|---|---|
| | C18 H19 N | Full match | Not the to | No results | Partial mat |
| | C7 H7 N3 | Full match | Full match | No results | Not the to |
| | C7 H7 N3 | Full match | Full match | No results | Not the to |
| | C20 H12 O | Full match | Invalid ma: | No results | Partial mat |
| | C9 H21 O4 | Full match | Full match | No results | Partial mat |
| | C9 H21 O4 | Full match | Full match | No results | Partial mat |
| | C18 H39 O | Not the to | Full match | Full match | Full match |
| | C6 H12 Cl3 | Full match | Full match | No results | Full match |
| Diphenyl(2,4,6-trimethylbenzoyl)phosphine o: | C22 H21 O | Not the to | No results | Full match | Not the to |
| | C19 H20 O | Full match | Invalid ma: | No results | Partial mat |
| | C20 H30 O | Full match | Invalid ma: | No results | Partial mat |
| | C19 H20 O | Full match | Not the to | No results | Partial mat |
| | C17 H20 N | Full match | Full match | No results | Full match |
| | C20 H22 O | Full match | Invalid ma: | No results | Partial mat |
| | C20 H22 O | Full match | Invalid ma: | No results | Partial mat |
| | C20 H22 O | Full match | Invalid ma: | No results | Partial mat |
| | C12 H27 O | Full match | Full match | Not the to | Full match |
| uorinated-Organic-Compounds-in-PubChem | C24 H32 F | Full match | Invalid ma: | No results | Not the to |
| | C15 H30 O | Full match | Full match | No results | Full match |
| N-Phenyl-1,4-benzenediamine | C12 H12 N | Full match | Full match | No results | Partial mat |
| | C24 H46 O | Full match | Full match | No results | No results |
| | C21 H30 O | Full match | Full match | No results | Full match |
| | C21 H30 O | Full match | Full match | No results | Full match |
| | C21 H30 O | Full match | Full match | No results | Full match |
| | C21 H30 O | Full match | Full match | No results | Full match |
| | C21 H30 O | Full match | Full match | No results | Full match |
| | C21 H30 O | Full match | Full match | No results | Full match |
| | C20 H30 O | Full match | Not the to | No results | Partial mat |
| yl-ester#::text=Product%20Description&text=I | C21 H32 O | Full match | Full match | No results | Partial mat |
| | C14 H18 N | Full match | Not the to | No results | Partial mat |
| (5R,5aS,9R,9aR,9bR)-9,9b-Dihydroxy-6,6,9a-tr | C21 H32 O | Full match | Invalid ma: | No results | Full match |
| | C19 H24 O | Full match | Not the to | No results | Full match |
| | C19 H24 O | Full match | Full match | No results | Full match |
| | C19 H24 O | Full match | Not the to | No results | Full match |
| | C18 H27 N | Full match | Invalid ma: | No results | Full match |
| | C15 H22 O | Full match | Not the to | No results | Not the to |
| | C15 H22 O | Full match | Not the to | No results | Full match |
| | C15 H22 O | Full match | Not the to | No results | Not the to |
| | C15 H22 O | Full match | Not the to | No results | Full match |
| | C15 H22 O | Full match | Not the to | No results | Full match |

| | | | | | |
|---|---|---|---|---|---|
| ıula | C18 H22 O | Full match | Invalid ma: | No results | Full match |
| ⁄lolecular-Formula | C21 H30 O | Full match | Full match | No results | Full match |
| | C19 H24 O | Full match | Not the to| | No results | Full match |
| | C19 H28 O | Full match | Full match | No results | Not the to| |
| | C19 H28 O | Full match | Full match | No results | Not the to| |
| | C21 H32 O | Full match | Invalid ma: | No results | Full match |
| | C21 H32 O | Full match | Invalid ma: | No results | Full match |
| | C21 H32 O | Full match | Invalid ma: | No results | Full match |
| | C21 H28 O | Full match | Not the to| | Partial mat | Full match |
| | C18 H24 O | Full match | Full match | No results | Full match |
| | C18 H24 O | Full match | Full match | No results | Full match |
| | C21 H32 O | Full match | Full match | No results | Full match |
| | C21 H32 O | Not the to| | Partial mat | Full match | Full match |
| | C21 H32 O | Full match | Full match | No results | Full match |
| HEBI:121492 | C19 H17 N | Invalid ma: | Invalid ma: | No results | Invalid ma: |
| | C20 H30 O | Full match | Full match | No results | Not the to| |
| | C20 H30 O | Not the to| | Not the to| | No results | Partial mat |
| | C19 H26 O | Full match | Full match | No results | Full match |
| | C19 H26 O | Full match | Full match | No results | Full match |
| | C19 H26 O | Full match | Full match | No results | Full match |
| | C19 H26 O | Full match | Full match | No results | Full match |
| | C14 H22 O | Full match | Full match | No results | Full match |
| 20is%20a%20hydroxyoctadecatrienoic,fatty%: | C18 H30 O | Full match | Full match | No results | Not the to| |
| 20is%20a%20hydroxyoctadecatrienoic,fatty%: | C18 H30 O | Full match | Full match | No results | Not the to| |
| | C12 H18 O | Full match | Full match | No results | Full match |
| | C12 H16 O | Full match | Not the to| | No results | Full match |
| | C19 H18 O | Full match | Full match | No results | Full match |
| | C25 H33 N | Full match | Invalid ma: | No results | Full match |
| | C17 H19 N | Full match | Full match | No results | Partial mat |
| Methyl 1,4a-dimethyl-6-methylene-5-[2-(2-ox | C21 H30 O | Full match | Full match | No results | Full match |
| hem.ncbi.nlm.nih.gov/compound/Diisobutyl-a | C14 H26 O | Full match | Full match | No results | Not the to| |
| ;-deca-6-9-diene-2-8-dione-.htm | C17 H24 O | Full match | Invalid ma: | No results | Partial mat |
| | C16 H22 N | Full match | Full match | No results | Partial mat |
| | C15 H22 O | Full match | Not the to| | No results | Partial mat |
| | C15 H18 O | Full match | Not the to| | No results | Full match |
| | C15 H18 O | Full match | Not the to| | No results | Not the to| |
| | C15 H20 O | Full match | Not the to| | No results | Not the to| |
| (3S,3aR,4S,4aR,7aR,8R,9aR)-3,4a,8-trimethyl-: | C20 H26 O | Full match | Not the to| | No results | Partial mat |
| | C15 H22 O | Full match | Not the to| | No results | Partial mat |
| (3aS,10aR,10bR)-6,10a-Dimethyl-3-methylene | C15 H18 O | Full match | Not the to| | No results | Partial mat |
| (3aS,10aR,10bR)-6,10a-Dimethyl-3-methylene | C15 H18 O | Full match | Full match | No results | Partial mat |
| (3aS,10aR,10bR)-6,10a-Dimethyl-3-methylene | C15 H18 O | Full match | Full match | No results | Partial mat |
| 1,4-dihydroxy-1,4-dimethyl-7-(propan-2-ylide | C15 H24 O | Full match | Full match | No results | Not the to| |
| 1,4-dihydroxy-1,4-dimethyl-7-(propan-2-ylide | C15 H24 O | Full match | Full match | No results | Not the to| |
| e | C22 H34 O | Full match | Invalid ma: | No results | Full match |
| | C17 H22 O | Full match | Not the to| | No results | Full match |
| | C18 H32 O | Full match | Full match | No results | Partial mat |
| | C20 H28 O | Full match | Not the to| | No results | Not the to| |
| | C20 H28 O | Full match | Not the to| | No results | Not the to| |
| :ction=Canonical-SMILES | C8 H7 N S2 | Full match | Full match | No results | Full match |
| | C20 H21 N | Full match | Not the to| | No results | Partial mat |

| | | | | | |
|---|---|---|---|---|---|
| | C20 H30 O | Full match | Not the to| | No results | Not the to|
| | C17 H22 O | Full match | Invalid ma: | No results | Full match |
| | C19 H24 O | Full match | Invalid ma: | No results | Full match |
| 13-[(2R,3R,4R,5R)-3,4-Dihydroxy-5-(hydroxym | C18 H35 N | Full match | Invalid ma: | No results | Full match |
| | C18 H35 N | Full match | Invalid ma: | No results | Full match |
| | C29 H38 O | Full match | Not the to| | No results | Full match |
| ɔyl%20phthalate%20is%20a%20phthalate,Poly | C14 H18 O | Full match | Invalid ma: | No results | Partial mat |
| | C19 H22 O | Full match | Invalid ma: | No results | Partial mat |
| | C29 H44 O | Full match | Invalid ma: | No results | Full match |
| | C20 H26 O | Full match | Invalid ma: | No results | Full match |
| (5R,6R,7R)-5-(4-Hydroxy-3-methoxyphenyl)-6, | C19 H22 O | Full match | Invalid ma: | No results | Partial mat |
| 2-ethylhexyl salicylate | C15 H22 O | Full match | Partial mat | Full match | Partial mat |
| | C19 H22 O | Full match | Not the to| | No results | Full match |
| | C19 H26 O | Full match | Full match | No results | Full match |
| | C19 H26 O | Full match | Full match | No results | Full match |
| nlm.nih.gov/compound/69131857 | C18 H24 O | Full match | Full match | No results | Partial mat |
| nlm.nih.gov/compound/69131858 | C18 H24 O | Full match | Full match | No results | Partial mat |
| nlm.nih.gov/compound/69131859 | C18 H24 O | Full match | Full match | No results | Partial mat |
| | C18 H22 O | Full match | Invalid ma: | No results | Not the to| |
| | C15 H17 Cl | Full match | Full match | Full match | Full match |
| | C16 H22 Cl | Full match | Full match | Full match | Full match |
| | C12 H17 N | Full match | Full match | Full match | Partial mat |
| | C14 H21 N | Full match | Full match | No results | Full match |
| | C14 H21 N | Full match | Full match | Full match | Full match |
| | C10 H19 N | Full match | Not the to| | Full match | Full match |
| | C15 H28 N | Full match | Not the to| | No results | No match |
| ɔpyl_alanine | C15 H21 N | Full match | Full match | No results | Partial mat |
| | C17 H14 O | Full match | Invalid ma: | No results | Full match |
| | C17 H14 O | Full match | Invalid ma: | No results | Full match |
| | C17 H14 O | Full match | Invalid ma: | No results | Full match |
| | C16 H24 O | Full match | Invalid ma: | No results | Partial mat |
| | C7 H5 N O | Full match | Full match | No results | Full match |
| | C9 H15 N ( | Full match | Invalid ma: | No results | Full match |
| | C16 H35 N | Full match | Full match | No results | Full match |
| | C16 H35 N | Full match | Full match | No results | Full match |
| | C19 H25 N | Full match | Full match | No results | Partial mat |
| | C8 H19 O4 | Full match | Full match | Full match | Full match |
| | C12 H27 N | Full match | Full match | No results | Full match |
| | C18 H39 N | Full match | Full match | No results | Full match |
| | C14 H16 N | Full match | Full match | No results | Full match |
| | C21 H28 N | Full match | Full match | No results | Partial mat |
| | C16 H26 O | Full match | Full match | No results | Partial mat |
| | C16 H26 O | Full match | Full match | No results | Full match |
| | C12 H18 N | Full match | Full match | No results | Partial mat |
| | C33 H44 N | Full match | Invalid ma: | No results | Full match |
| | C18 H26 O | Full match | Invalid ma: | No results | Full match |
| | C16 H25 N | Full match | Full match | No results | Partial mat |
| | C14 H11 Cl | Full match | Full match | Full match | Full match |
| | C17 H21 N | Full match | Full match | No results | Full match |
| | C19 H16 Cl | Not the to| | Full match | Full match | Full match |
| | C16 H14 O | Full match | Not the to| | No results | Full match |

| | | | | | |
|---|---|---|---|---|---|
| | C16 H14 O | Not the to| | Not the to| | No results | Full match |
| | C14 H14 O | Full match | Full match | Full match | Full match |
| | C11 H12 N | Full match | Full match | No results | Full match |
| | C16 H23 N | Full match | Full match | No results | Not the to| |
| | C15 H21 N | Full match | Not the to| | No results | Partial mat |
| | C21 H27 N | Full match | Full match | No results | Full match |
| | C16 H17 N | Full match | Full match | No results | Full match |
| | C14 H23 N | Full match | Full match | No results | Full match |
| | C16 H25 N | Full match | Full match | No results | Full match |
| | C16 H25 N | Full match | Full match | No results | Full match |
| | C12 H17 N | Full match | Full match | No results | Full match |
| | C16 H16 Cl | Full match | Full match | No results | Full match |
| icidal-activity-of-2-amino-5-6-7-8-tetrahydro--1 | C16 H14 N | Full match | Invalid ma: | No results | Full match |
| | C38 H72 N | Not the to| | No match | Full match | Full match |
| | C38 H69 N | No match | Full match | Full match | Full match |
| | C41 H76 N | No match | Full match | Full match | Full match |
| | C16 H20 F | Invalid ma: | Invalid ma: | No results | Unused |
| | C14 H18 N | Not the to| | Partial mat | Full match | Partial mat |
| | C21 H25 N | Full match | Full match | No results | Full match |
| | C19 H18 Cl | Full match | Full match | No results | Full match |
| | C15 H14 N | Full match | Full match | No results | Full match |
| | C15 H14 N | Full match | Full match | No results | Full match |
| | C15 H14 N | Full match | Full match | No results | Full match |
| | C15 H12 N | Full match | Full match | No results | Partial mat |
| | C15 H12 N | Full match | Full match | No results | Partial mat |
| | C15 H12 N | Full match | Full match | Full match | Full match |
| | C9 H7 Cl2 I | Full match | Full match | No results | Full match |
| | C15 H12 N | Full match | Full match | Full match | Full match |
| N-{[(3as,5ar,8ar,8bs)-2,2,7,7-Tetramethyltetra | C12 H22 N | Full match | Invalid ma: | No results | Not the to| |
| | C15 H17 Cl | Full match | Full match | No results | Full match |
| | C21 H25 N | Full match | Not the to| | No results | Partial mat |
| | C21 H25 Cl | Full match | Full match | Full match | Full match |
| | C32 H39 N | Full match | Full match | Full match | Full match |
| | C25 H36 O | Full match | Full match | No results | Full match |
| | C18 H18 O | Full match | Invalid ma: | No results | Full match |
| | C13 H17 N | Full match | Full match | No results | Partial mat |
| | C14 H20 O | Full match | Invalid ma: | No results | Partial mat |
| | C20 H30 O | Full match | Not the to| | No results | Not the to| |
| Tumor-promoting agent used in research | C20 H28 O | Full match | Invalid ma: | No results | Partial mat |
| | C18 H14 F4 | Not the to| | Full match | Full match | Full match |
| | C24 H32 O | Full match | Not the to| | No results | Not the to| |
| | C19 H27 N | Full match | Full match | No results | Full match |
| | C22 H30 F | Full match | Unused | No results | Not the to| |
| | C15 H25 N | Full match | Full match | Full match | Full match |
| | C15 H25 N | Full match | Invalid ma: | No results | Partial mat |
| | C16 H21 N | Full match | Full match | Full match | Full match |
| 2-(3,4-Dimethoxyphenyl)-5-methylamino-2-iso | C17 H26 N | Full match | Full match | No results | Full match |
| | C18 H31 N | Not the to| | Full match | Full match | Full match |
| | C24 H20 N | Not the to| | Full match | Full match | Full match |
| | C22 H26 N | Not the to| | Full match | Full match | Full match |
| | C17 H20 F6 | Full match | Full match | No results | Full match |

| | | | | | |
|---|---|---|---|---|---|
| | C19 H24 N | Full match | Full match | No results | Full match |
| | C22 H23 Cl | Not the to| | Full match | Full match | Full match |
| | C13 H24 N | Full match | Full match | No results | Full match |
| | C24 H29 N | Full match | Full match | Full match | Full match |
| | C24 H34 O | Full match | Full match | No results | Full match |
| | C17 H15 Cl | Full match | Full match | No results | Full match |
| | C15 H22 O | Full match | Full match | No results | Full match |
| | C17 H21 N | Full match | Full match | No results | Full match |
| | C16 H24 N | Full match | Full match | No results | Partial mat |
| | C12 H11 Cl | Full match | No results | Full match | Partial mat |
| | C21 H30 O | Full match | Full match | No results | Not the to| |
| | C20 H23 N | Full match | Full match | No results | Partial mat |
| | C19 H21 N | Full match | Full match | No results | Not the to| |
| | C21 H28 O | Full match | Full match | No results | Full match |
| | C19 H26 O | Full match | Not the to| | No results | Not the to|
| | C21 H23 N | Full match | Full match | No results | Partial mat |
| ı-(crm) | C17 H22 O | Full match | Not the to| | No results | Partial mat |
| 2C17-trione#::text=5%CE%B1%2DAndrostan%2 | C19 H26 O | Full match | Full match | No results | Partial mat |
| 2C17-trione#::text=5%CE%B1%2DAndrostan%2 | C19 H26 O | Full match | Full match | No results | Partial mat |
| 2C17-trione#::text=5%CE%B1%2DAndrostan%2 | C19 H26 O | Full match | Full match | No results | Partial mat |
| | C20 H25 N | Full match | Full match | No results | Partial mat |
| ıloride) | C13 H18 N | Full match | Full match | No results | Partial mat |
| | C16 H27 N | Full match | Full match | No results | Full match |
| | C21 H28 O | Full match | Full match | No results | Partial mat |
| | C21 H28 O | Full match | Full match | No results | Partial mat |
| | C21 H28 O | Full match | Full match | No results | Partial mat |
| | C16 H12 O | Full match | Full match | No results | Not the to| |
| | C17 H20 O | Full match | Full match | No results | Full match |
| | C27 H30 N | Full match | Unused | No results | Not the to| |
| | C21 H28 O | Full match | Full match | No results | Full match |
| | C21 H33 F | Invalid ma: | Invalid ma: | No results | Invalid ma: |
| | C19 H19 F | Full match | Full match | No results | Full match |
| | C18 H19 N | Full match | Full match | No results | Partial mat |
| | C17 H27 N | Full match | Full match | No results | Partial mat |
| | C20 H23 N | Full match | Full match | No results | Partial mat |
| | C20 H23 N | Full match | Full match | No results | Partial mat |
| | C18 H18 Cl | Full match | Full match | No results | Full match |
| | C18 H18 Cl | Full match | Full match | No results | Full match |
| | C20 H21 F | Full match | Full match | Full match | Full match |
| | C19 H21 N | Full match | Full match | No results | Partial mat |
| | C16 H25 N | Full match | Full match | No results | Not the to| |
| | C16 H25 N | Full match | Full match | No results | Not the to| |
| | C15 H11 Cl | Full match | Full match | No results | Full match |
| | C17 H17 Cl | Full match | Full match | Full match | Full match |
| | C17 H27 N | Full match | Full match | Full match | Full match |
| | C17 H27 N | Full match | Full match | Full match | Full match |
| | C17 H27 N | Full match | Full match | No results | Full match |
| | C17 H16 N | Full match | Not the to| | No results | Partial mat |
| | C19 H24 O | Full match | Full match | No results | Full match |
| | C21 H26 O | Not the to| | Full match | Full match | Full match |
| | C13 H19 N | Full match | Full match | No results | Full match |

| | | | | |
|---|---|---|---|---|
| C22 H30 O | Full match | Full match | No results | Partial mat |
| C16 H11 N | Full match | Full match | No results | Partial mat |
| C16 H13 N | Not the to | Full match | Full match | Full match |
| C20 H30 O | Full match | Not the to | No results | Not the to |
| C16 H24 O | Full match | Full match | No results | Partial mat |
| C16 H24 O | Full match | Full match | No results | Partial mat |
| C16 H24 O | Full match | Not the to | No results | Partial mat |
| C16 H24 O | Full match | Full match | No results | Partial mat |
| C14 H18 O | Full match | Not the to | No results | Partial mat |
| C16 H28 O | Full match | Not the to | No results | Not the to |
| C16 H28 O | Full match | Not the to | No results | Not the to |
| C12 H14 O | Full match | Full match | No results | Partial mat |
| C12 H14 O | Full match | Full match | No results | Partial mat |
| C20 H28 O | Full match | Not the to | No results | Partial mat |
| C20 H16 O | Full match | Full match | No results | Partial mat |
| C22 H30 O | Full match | Not the to | No results | Partial mat |
| C20 H28 O | Full match | Not the to | No results | Partial mat |
| C20 H28 O | Full match | Not the to | No results | Partial mat |
| C20 H28 O | Full match | Not the to | No results | Partial mat |
| C20 H28 O | Full match | Not the to | No results | Partial mat |
| C20 H34 O | Full match | Not the to | No results | Partial mat |
| C15 H24 O | Full match | Not the to | No results | Partial mat |
| C16 H32 O | Full match | Full match | No results | Partial mat |
| C17 H20 O | Full match | Not the to | No results | Partial mat |
| C21 H30 O | Full match | Full match | No results | Partial mat |
| C21 H30 O | Full match | Full match | No results | Partial mat |
| C21 H30 O | Full match | Full match | No results | Partial mat |
| C21 H30 O | Full match | Full match | No results | Partial mat |
| C23 H25 N | Full match | Full match | No results | Partial mat |
| C19 H19 N | Full match | Full match | No results | Not the to |
| C27 H36 O | Full match | Full match | No results | Partial mat |
| C18 H26 O | Full match | Not the to | No results | Partial mat |
| C24 H37 F | Full match | No match | No results | No match |
| C26 H43 F | Full match | Invalid ma | No results | No match |
| C30 H59 N | Full match | Invalid ma | No results | No match |
| C15 H20 Cl | Invalid ma | Invalid ma | No results | No match |
| C27 H60 F | Full match | No match | No results | No match |
| C31 H51 F3 | Full match | No match | No results | No match |
| C38 H61 N | Full match | No match | No results | Not the to |
| C18 H34 F | Full match | Invalid ma | No results | No match |
| C13 H27 F | Full match | Invalid ma | No results | No match |
| C37 H72 N | Full match | Invalid ma | No results | No results |
| C26 H43 F | Full match | No match | No results | No match |
| C26 H28 F2 | Full match | Invalid ma | No results | No match |
| C26 H43 F | Full match | Invalid ma | No results | No match |
| C20 H29 F | Full match | No match | No results | No match |
| C23 H37 F | Full match | No match | No results | No match |
| C16 H27 F | Full match | Invalid ma | No results | No match |
| C26 H45 F | Full match | Invalid ma | No results | No match |
| C36 H76 F2 | Full match | Invalid ma | No results | No match |
| C16 H26 F | Invalid ma | No match | No results | No match |

| | | | | |
|---|---|---|---|---|
| C25 H40 F2 | Full match | No match | No results | No match |
| C21 H31 F | Full match | No match | No results | Invalid ma: |
| C32 H56 F2 | Invalid ma: | No match | No results | No match |
| C25 H59 N | Full match | No match | No results | No match |
| C20 H38 F3 | Invalid ma: | Invalid ma: | No results | No match |
| C30 H60 N | Full match | No match | No results | No results |
| C28 H47 F3 | Full match | Invalid ma: | No results | No match |
| C28 H40 N | Full match | No match | No results | No match |
| C16 H29 F3 | Full match | Invalid ma: | No results | No results |
| C26 H43 F | Full match | Invalid ma: | No results | No match |
| C15 H20 Cl | Full match | Invalid ma: | No results | No match |
| C30 H48 F | Full match | Invalid ma: | No results | No match |
| C19 H31 F | Full match | Invalid ma: | No results | No match |
| C29 H60 F3 | Full match | Invalid ma: | No results | No results |
| C29 H69 F | Full match | Invalid ma: | No results | No results |
| C23 H32 N | Full match | Invalid ma: | No results | No match |
| C28 H32 F2 | Full match | Invalid ma: | No results | No match |
| C26 H40 F | Full match | Invalid ma: | No results | No match |
| C42 H74 F2 | Full match | Invalid ma: | No results | No results |
| C35 H62 F2 | Full match | No match | No results | No results |
| C20 H27 Cl | Full match | Invalid ma: | No results | No match |
| C32 H66 F3 | Full match | Invalid ma: | No results | No results |
| C16 H21 Cl | Full match | Invalid ma: | No results | No match |
| C15 H24 F | Full match | No match | No results | No match |
| C13 H25 N | Invalid ma: | Invalid ma: | No results | No match |
| C17 H33 N | Full match | No match | No results | No match |
| C16 H27 F | Invalid ma: | No match | No results | No match |
| C21 H41 F | Full match | No match | No results | No match |
| C16 H33 N | Full match | Invalid ma: | No results | No match |
| C5 H13 F3 | Full match | Invalid ma: | No results | No match |
| C16 H28 N | Full match | Invalid ma: | No results | No match |
| C22 H32 F4 | Full match | Invalid ma: | No results | No match |
| C88 H145 I | Full match | Invalid ma: | No results | No results |
| C37 H58 F | Full match | No match | No results | No match |
| C22 H56 N | Full match | No match | No results | No match |
| C15 H25 F3 | Invalid ma: | Invalid ma: | No results | No match |
| C19 H24 F2 | Full match | Unused | No results | No match |
| C23 H52 F2 | Full match | Invalid ma: | No results | No match |
| C32 H76 N | Full match | Invalid ma: | No results | No results |
| C14 H29 N | Full match | Invalid ma: | No results | No results |
| C26 H32 F2 | Full match | Unused | No results | No match |
| C17 H29 F | Invalid ma: | Invalid ma: | No results | No match |
| C23 H51 F2 | Full match | Invalid ma: | No results | No results |
| C23 H45 F3 | Full match | No match | No results | No results |
| C26 H39 F | Full match | Invalid ma: | No results | No match |
| C12 H30 F2 | Full match | No match | No results | No match |
| C23 H33 N | Invalid ma: | Invalid ma: | No results | No match |
| C23 H41 F3 | Full match | No match | No results | No match |
| C19 H33 F3 | Full match | No match | No results | No match |
| C18 H32 F4 | Full match | No match | No results | No match |
| C20 H34 F2 | Full match | No match | No results | No match |

| | | | | |
|---|---|---|---|---|
| C82 H102 I | Full match | Invalid ma: | No results | No results |
| C20 H35 N | Full match | Invalid ma: | No results | No match |
| C25 H42 F2 | Full match | Invalid ma: | No results | No match |
| C41 H72 F4 | Full match | Invalid ma: | No results | No results |
| C6 H11 N5 | Full match | Invalid ma: | No results | No results |
| C37 H71 N | Invalid ma: | Invalid ma: | No results | No results |
| C28 H56 N | Full match | Invalid ma: | No results | No match |
| C22 H42 N | Full match | Invalid ma: | No results | No match |
| C16 H22 F2 | Full match | Invalid ma: | No results | No match |
| C17 H38 F2 | Full match | Invalid ma: | No results | No match |
| C20 H26 F4 | Full match | Invalid ma: | No results | No match |
| C23 H45 N | Full match | Invalid ma: | No results | No match |
| C16 H27 F | Invalid ma: | No match | No results | No match |
| C19 H33 F5 | Full match | Invalid ma: | No results | No match |
| C23 H35 F | Full match | Invalid ma: | No results | No match |
| C20 H42 F3 | Full match | Invalid ma: | No results | No results |
| C19 H35 F3 | Full match | Invalid ma: | No results | No match |
| C18 H20 Cl | Full match | Invalid ma: | No results | No match |
| C13 H21 F | Full match | No match | No results | No match |
| C27 H43 F2 | Full match | Invalid ma: | No results | No match |
| C21 H45 F | Full match | No match | No results | No results |
| C19 H31 F | Invalid ma: | No match | No results | No match |
| C12 H22 F2 | Full match | Invalid ma: | No results | No match |
| C13 H26 F | Full match | Invalid ma: | No results | No match |
| C19 H47 N | Full match | No match | No results | No match |
| C24 H47 F3 | Full match | Invalid ma: | No results | No match |
| C23 H34 F4 | Full match | Invalid ma: | No results | No match |
| C19 H43 N | Full match | No match | No results | No results |
| C18 H33 F | Invalid ma: | Unused | No results | No match |
| C10 H19 F3 | Full match | Invalid ma: | No results | No results |
| C22 H44 F | Full match | Invalid ma: | No results | No match |
| C20 H40 N | Full match | Invalid ma: | No results | No match |
| C14 H28 F2 | Full match | No match | No results | No match |
| C25 H50 N | Full match | Invalid ma: | No results | No match |
| C9 H21 N5 | Full match | Invalid ma: | No results | No results |
| C20 H38 F | Full match | No match | No results | No match |
| C10 H19 F2 | Full match | Invalid ma: | No results | No match |
| C19 H34 F | Full match | Unused | No results | No match |
| C19 H37 F | Invalid ma: | Invalid ma: | No results | No match |
| C16 H36 F | Full match | Invalid ma: | No results | No match |
| C32 H64 F6 | Full match | Invalid ma: | No results | No match |
| C26 H35 F3 | Invalid ma: | Invalid ma: | No results | No match |
| C18 H30 F4 | Full match | Invalid ma: | No results | No match |
| C26 H44 F3 | Full match | Invalid ma: | No results | No match |
| C18 H39 Cl | Full match | Invalid ma: | No results | No match |
| C19 H42 N | Full match | No match | No results | No match |
| C16 H36 F2 | Full match | Invalid ma: | No results | No match |
| C12 H21 F | Invalid ma: | Invalid ma: | No results | No match |
| C26 H50 F2 | Full match | No match | No results | No results |
| C19 H32 F3 | Full match | No match | No results | No match |
| C22 H39 F3 | Full match | No match | No results | No match |

| | | | | |
|---|---|---|---|---|
| C18 H32 F2 | Full match | Invalid ma: | No results | No match |
| C27 H54 F | Full match | Unused | No results | No match |
| C19 H46 N | Full match | Invalid ma: | No results | No results |
| C12 H23 F | Invalid ma: | No match | No results | No match |
| C23 H35 F | Full match | No match | No results | No match |
| C8 H17 N3 | Full match | No match | No results | No match |
| C10 H24 F | Full match | Invalid ma: | No results | No match |
| C32 H52 F2 | Invalid ma: | No match | No results | No results |
| C22 H44 F | Full match | Invalid ma: | No results | No match |
| C19 H46 N | Full match | Invalid ma: | No results | No results |
| C13 H18 F2 | Full match | No match | No results | No match |
| C22 H52 N | Full match | Invalid ma: | No results | No results |
| C10 H24 F | Full match | Invalid ma: | No results | No match |
| C29 H33 F3 | Full match | Invalid ma: | No results | No match |
| C22 H39 F5 | Full match | Invalid ma: | No results | No match |
| C8 H20 F2 | Full match | Invalid ma: | No results | No match |
| C25 H32 F | Full match | Invalid ma: | No results | No match |
| C10 H30 N | Full match | Invalid ma: | No results | No results |
| C15 H30 F2 | Full match | Invalid ma: | No results | No match |
| C10 H21 F | Full match | No match | No results | Invalid ma: |
| C18 H32 F3 | Full match | Invalid ma: | No results | No match |
| C18 H31 F | Full match | Invalid ma: | No results | No match |
| C20 H38 N | Full match | Invalid ma: | No results | No match |
| C22 H32 F4 | Full match | Invalid ma: | No results | No match |
| C17 H24 F2 | Full match | No match | No results | No match |
| C10 H27 N | Full match | No match | No results | No match |
| C17 H30 F | Invalid ma: | No match | No results | No match |
| C8 H24 N1 | Full match | Invalid ma: | No results | No match |
| C25 H38 F2 | Full match | No match | No results | No match |
| C22 H36 F2 | Full match | Invalid ma: | No results | No match |
| C9 H16 F N | Full match | Invalid ma: | No results | No match |
| C11 H28 N | Full match | Invalid ma: | No results | No match |
| C16 H27 N | Full match | Invalid ma: | No results | No match |
| C17 H29 F | Invalid ma: | Invalid ma: | No results | No match |
| C22 H36 F2 | Full match | No match | No results | No match |
| C17 H20 F2 | Full match | Unused | No results | No match |
| C18 H28 F2 | Full match | Unused | No results | No match |
| C13 H28 N | Full match | Invalid ma: | No results | No match |
| C16 H32 F2 | Full match | Invalid ma: | No results | No match |
| C15 H29 F3 | Full match | No match | No results | No match |
| C17 H29 F | Full match | Invalid ma: | No results | No match |
| C14 H21 F2 | Invalid ma: | No match | No results | No match |
| C17 H28 F2 | Full match | Invalid ma: | No results | No match |
| C19 H40 Cl | Full match | Invalid ma: | No results | No match |
| C16 H24 F4 | Full match | Invalid ma: | No results | No match |
| C22 H37 F | Full match | Invalid ma: | No results | No match |
| C13 H17 Cl | Full match | Invalid ma: | No results | No match |
| C9 H24 N6 | Full match | Unused | No results | No match |
| C23 H48 N | Full match | Invalid ma: | No results | No match |
| C15 H32 N | Full match | Invalid ma: | No results | No match |
| C22 H33 F | Full match | Invalid ma: | No results | No match |

| | | | | |
|---|---|---|---|---|
| C15 H30 N | Full match | Invalid ma: | No results | No match |
| C23 H45 F2 | Full match | Invalid ma: | No results | No match |
| C19 H35 F3 | Full match | Invalid ma: | No results | No match |
| C20 H31 F2 | Full match | Invalid ma: | No results | No match |
| C14 H22 F2 | Full match | Invalid ma: | No results | No match |
| C19 H42 F2 | Full match | Invalid ma: | No results | No match |
| C15 H29 N | Full match | Invalid ma: | No results | No match |
| C40 H75 F2 | Full match | Invalid ma: | No results | No results |
| C22 H33 F2 | Full match | Invalid ma: | No results | No match |
| C37 H56 N | Full match | Invalid ma: | No results | No match |
| C10 H30 N | Full match | Invalid ma: | No results | No results |
| C9 H21 N5 | Full match | Invalid ma: | No results | No results |
| C17 H34 F | Full match | Invalid ma: | No results | No match |
| C22 H39 F | Full match | No match | No results | No match |
| C30 H67 N | Full match | Invalid ma: | No results | No match |
| C17 H29 F | Full match | Invalid ma: | No results | No match |
| C11 H21 F | Full match | Invalid ma: | No results | No match |
| C12 H18 F6 | Full match | Invalid ma: | No results | No match |
| C8 H19 F N | Full match | Invalid ma: | No results | No match |
| C18 H24 F2 | Full match | No match | No results | No match |
| C20 H32 F2 | Full match | Invalid ma: | No results | No match |
| C20 H32 F2 | Full match | No match | No results | No match |
| C33 H65 F2 | Full match | Invalid ma: | No results | No results |
| C19 H34 F | Full match | Invalid ma: | No results | No match |
| C22 H32 F4 | Full match | Invalid ma: | No results | No match |
| C34 H60 F5 | Full match | Invalid ma: | No results | No match |
| C31 H59 F | Full match | No match | No results | No results |
| C12 H17 Cl | Full match | Invalid ma: | No results | No match |
| C32 H58 F | Full match | Invalid ma: | No results | No results |
| C22 H32 F2 | Full match | No match | No results | No match |
| C14 H22 F2 | Full match | Invalid ma: | No results | No match |
| C11 H20 F3 | Full match | No match | No results | No match |
| C20 H41 N | Full match | Invalid ma: | No results | No match |
| C19 H41 N | Full match | Invalid ma: | No results | Invalid ma: |
| C26 H42 F3 | Full match | Invalid ma: | No results | No match |
| C13 H27 N | Full match | Invalid ma: | No results | No match |
| C8 H18 F N | Full match | Invalid ma: | No results | No match |
| C15 H34 N | Full match | No match | No results | No match |
| C23 H37 F | Full match | No match | No results | No match |
| C12 H26 N | Full match | Invalid ma: | No results | No match |
| C23 H38 F5 | Full match | Invalid ma: | No results | No match |
| C12 H19 N | Full match | Invalid ma: | No results | No match |
| C17 H28 F2 | Full match | Invalid ma: | No results | No match |
| C15 H25 N | Full match | Invalid ma: | No results | No match |
| C35 H64 F | Full match | Invalid ma: | No results | No results |
| C26 H49 N | Full match | Invalid ma: | No results | No match |
| C15 H27 Cl | Full match | No match | No results | No match |
| C22 H42 F3 | Full match | Invalid ma: | No results | No match |
| C11 H19 F3 | Full match | Invalid ma: | No results | No match |
| C15 H35 N | Full match | Invalid ma: | No results | No match |
| C19 H38 Cl | Full match | Unused | No results | No match |

| | | | | |
|---|---|---|---|---|
| C15 H38 N | Full match | Invalid ma: | No results | No match |
| C24 H34 F | Full match | Invalid ma: | No results | No match |
| C12 H25 N | Full match | Invalid ma: | No results | No results |
| C18 H38 Cl | Full match | Invalid ma: | No results | No match |
| C11 H24 N | Full match | Invalid ma: | No results | No match |
| C12 H30 N | Full match | Invalid ma: | No results | No results |
| C18 H34 F2 | Full match | Invalid ma: | No results | No match |
| C18 H35 F2 | Full match | Invalid ma: | No results | No match |
| C19 H42 F2 | Full match | Invalid ma: | No results | No match |
| C15 H20 F2 | Full match | Invalid ma: | No results | No match |
| C10 H27 F3 | Full match | Invalid ma: | No results | No match |
| C16 H44 N | Full match | Invalid ma: | No results | No match |
| C18 H27 F2 | Full match | No match | No results | No match |
| C11 H26 F | Full match | Invalid ma: | No results | No match |
| C19 H37 N | Full match | Invalid ma: | No results | No match |
| C9 H24 N6 | Full match | Unused | No results | No match |
| C9 H19 N4 | Full match | Invalid ma: | No results | No match |
| C28 H51 N | Full match | Invalid ma: | No results | No results |
| C26 H43 F | Full match | Invalid ma: | No results | Unused |
| C28 H48 F2 | Full match | Invalid ma: | No results | No match |
| C14 H30 N | Full match | No match | No results | No match |
| C14 H23 F | Full match | Invalid ma: | No results | No match |
| C16 H32 F2 | Full match | Invalid ma: | No results | No match |
| C14 H26 F2 | Full match | No match | No results | No match |
| C13 H27 Cl | Full match | Invalid ma: | No results | No match |
| C13 H23 N | Invalid ma: | No match | No results | No match |
| C9 H20 F N | Full match | Invalid ma: | No results | No match |
| C16 H32 F2 | Full match | Invalid ma: | No results | No match |
| C15 H31 N | Full match | No match | No results | No match |
| C29 H65 F | Full match | No match | No results | No match |
| C11 H28 F2 | Full match | No match | No results | No match |
| C7 H10 Cl2 | Invalid ma: | Invalid ma: | No results | No match |
| C9 H18 N4 | Full match | Invalid ma: | No results | No match |
| C14 H18 F2 | Full match | No match | No results | No match |
| C11 H26 F | Full match | No match | No results | Invalid ma: |
| C28 H30 F2 | Full match | Invalid ma: | No results | Invalid ma: |
| C22 H38 F2 | Full match | Invalid ma: | No results | No match |
| C10 H22 N | Full match | Invalid ma: | No results | No match |
| C25 H25 N | Full match | Invalid ma: | No results | No match |
| C15 H20 F2 | Full match | Invalid ma: | No results | No match |
| C17 H32 N | Full match | Invalid ma: | No results | No match |
| C25 H42 F2 | Full match | Invalid ma: | No results | No match |
| C18 H28 Cl | Full match | Invalid ma: | No results | No match |
| C45 H91 F5 | Full match | Invalid ma: | No results | No results |
| C19 H37 Cl | Full match | Unused | No results | No match |
| C15 H21 Cl | Full match | Invalid ma: | No results | No match |
| C31 H57 F2 | Full match | Invalid ma: | No results | No match |
| C12 H28 N | Full match | No match | No results | No match |
| C27 H40 F2 | Full match | Invalid ma: | No results | No match |
| C21 H36 N | Full match | Invalid ma: | No results | No match |
| C11 H22 F | Full match | Invalid ma: | No results | No match |

| | | | | |
|---|---|---|---|---|
| C17 H32 F | Invalid ma: | Invalid ma: | No results | No match |
| C11 H27 N | Full match | No match | No results | No match |
| C31 H65 F! | Full match | Invalid ma: | No results | No match |
| C8 H23 N7 | Full match | Invalid ma: | No results | No match |
| C12 H27 N | Full match | No match | No results | No match |
| C11 H22 N | Full match | Invalid ma: | No results | No match |
| C15 H33 N | Full match | Invalid ma: | No results | No results |
| C20 H36 N | Full match | No match | No results | No match |
| C15 H36 N | Full match | Invalid ma: | No results | No match |
| C12 H26 Cl | Full match | Invalid ma: | No results | No match |
| C18 H42 F | Full match | No match | No results | No match |
| C37 H43 F | Full match | Invalid ma: | No results | No match |
| C8 H12 F N | Full match | Invalid ma: | No results | No match |
| C27 H43 Cl | Full match | Invalid ma: | No results | No match |
| C17 H35 Cl | Full match | Invalid ma: | No results | No match |
| C15 H20 Cl | Invalid ma: | No match | No results | No match |
| C14 H24 F4 | Full match | Invalid ma: | No results | No match |
| C11 H28 F2 | Full match | No match | No results | No match |
| C13 H19 Cl | Full match | Invalid ma: | No results | No match |
| C15 H31 N | Full match | Invalid ma: | No results | No match |
| C13 H29 F | Full match | Invalid ma: | No results | No match |
| C23 H48 Cl | Full match | Invalid ma: | No results | No match |
| C15 H26 F2 | Full match | No match | No results | No match |
| C17 H24 F2 | Full match | Invalid ma: | No results | No match |
| C23 H32 F4 | Full match | No match | No results | No match |
| C7 H16 F N | Full match | No match | No results | No match |
| C27 H39 F | Full match | Invalid ma: | No results | No match |
| C19 H30 F | Full match | Invalid ma: | No results | No match |
| C14 H31 F2 | Full match | Invalid ma: | No results | No match |
| C14 H25 F3 | Full match | Invalid ma: | No results | No match |
| C18 H32 F3 | Full match | Invalid ma: | No results | No match |
| C15 H20 O | Full match | Invalid ma: | No results | Full match |
| C15 H20 O | Full match | Invalid ma: | No results | Full match |
| C15 H20 O | Full match | Invalid ma: | No results | Full match |
| C16 H22 O | Full match | Invalid ma: | No results | Full match |
| C16 H22 O | Full match | Invalid ma: | No results | Full match |
| C16 H22 O | Full match | Invalid ma: | No results | Full match |
| C16 H22 O | Full match | Invalid ma: | No results | Full match |
| C16 H22 O | Full match | Invalid ma: | No results | Full match |
| C20 H28 O | Full match | Invalid ma: | No results | Full match |
| C20 H28 O | Full match | Invalid ma: | No results | Full match |
| C20 H28 O | Full match | Invalid ma: | No results | Full match |
| C20 H28 O | Full match | Invalid ma: | No results | Full match |
| C20 H31 N | Full match | Invalid ma: | No results | Full match |
| C29 H44 O | Full match | Invalid ma: | No results | Full match |
| C29 H44 O | Full match | Invalid ma: | No results | Full match |
| C15 H29 N | Full match | Invalid ma: | No results | Full match |
| C13 H18 O | Full match | Invalid ma: | No results | Full match |
| C17 H30 N | Full match | Invalid ma: | No results | Full match |
| C15 H20 O | Full match | Not the to | No results | Not the to |
| C15 H20 O | Full match | Full match | No results | Not the to |

| | | | | | |
|---|---|---|---|---|---|
| | C16 H23 N | Full match | Invalid ma: | No results | Full match |
| | C29 H44 O | Full match | Invalid ma: | No results | Full match |
| | C16 H24 O | Full match | Full match | No results | Not the to |
| | C16 H24 O | Full match | Full match | No results | Not the to |
| | C17 H25 N | Full match | Full match | No results | Partial mat |
| | C21 H22 O | Full match | Invalid ma: | No results | Full match |
| | C17 H26 O | Full match | Invalid ma: | No results | Full match |
| | C23 H23 F | Full match | Invalid ma: | No results | Full match |
| | C30 H38 N | Full match | Invalid ma: | No results | Full match |
| | C26 H30 F | Full match | Invalid ma: | No results | Full match |
| | C24 H32 O | Full match | Invalid ma: | No results | Full match |
| | C13 H14 O | Full match | Not the to | No results | Partial mat |
| | C18 H24 F | Full match | Invalid ma: | No results | Full match |
| | C15 H22 F | Invalid ma: | No match | No results | Unused |
| | C15 H20 O | Full match | Not the to | No results | Partial mat |
| | C15 H20 O | Full match | Not the to | No results | Partial mat |
| | C15 H20 O | Full match | Not the to | No results | Partial mat |
| | C27 H39 N | Full match | Invalid ma: | No results | Full match |
| | C17 H20 O | Full match | Invalid ma: | No results | Full match |
| | C15 H20 N | Full match | Invalid ma: | No results | Full match |
| | C15 H20 N | Full match | Invalid ma: | No results | Full match |
| | C17 H22 O | Full match | Invalid ma: | No results | Full match |
| | C15 H27 N | Full match | Invalid ma: | No results | Full match |
| | C15 H27 N | Full match | Invalid ma: | No results | Full match |
| | C17 H22 O | Full match | Not the to | No results | Full match |
| | C12 H15 Cl | Invalid ma: | No match | No results | No match |
| | C14 H23 N | Full match | Invalid ma: | No results | Full match |
| | C19 H20 O | Full match | Full match | No results | Not the to |
| | C21 H32 O | Full match | Full match | No results | Partial mat |
| | C17 H28 O | Full match | Invalid ma: | No results | Full match |
| | C22 H37 N | Full match | Invalid ma: | No results | Full match |
| | C22 H39 N | Full match | Invalid ma: | No results | Full match |
| | C22 H39 N | Full match | Invalid ma: | No results | Full match |
| | C29 H38 O | Full match | Invalid ma: | No results | Full match |
| | C18 H24 O | Full match | Invalid ma: | No results | Full match |
| | C18 H24 O | Full match | Invalid ma: | No results | Full match |
| | C20 H24 O | Full match | Not the to | No results | Full match |
| | C17 H30 O | Full match | Invalid ma: | No results | Full match |
| | C37 H44 N | Full match | Invalid ma: | No results | Full match |
| | C28 H32 N | Full match | No match | No results | Not the to |
| | C17 H24 O | Full match | Invalid ma: | No results | Full match |
| | C17 H20 O | Full match | Invalid ma: | No results | Full match |
| | C14 H29 N | Full match | Invalid ma: | No results | Not the to |
| | C16 H28 N | Full match | Invalid ma: | No results | Full match |
| 3,5-DI-TERT-BUTYL-4-HYDROXYACETOPHENOI | C16 H24 O | Full match | Invalid ma: | No results | Partial mat |
| 3,5-DI-TERT-BUTYL-4-HYDROXYACETOPHENOI | C16 H24 O | Full match | Invalid ma: | No results | Partial mat |
| 3,5-DI-TERT-BUTYL-4-HYDROXYACETOPHENOI | C16 H24 O | Full match | Invalid ma: | No results | Partial mat |
| | C20 H21 F | Full match | Invalid ma: | No results | Full match |
| | C14 H20 N | Full match | Invalid ma: | No results | Full match |
| | C19 H26 O | Full match | Invalid ma: | No results | Full match |
| | C15 H13 F | Full match | Invalid ma: | No results | Full match |

| | | | | | |
|---|---|---|---|---|---|
| | C16 H23 N | Full match | Not the to| | No results | Partial mat |
| | C16 H23 N | Full match | Not the to| | No results | Partial mat |
| | C18 H19 Cl | Full match | No match | No results | Full match |
| | C32 H33 F | Full match | Invalid ma: | No results | Full match |
| 1-AMINO-9-OXO-4-ACRIDNECARBOXYLIC ACIL | C14 H10 N | Full match | Invalid ma: | No results | Full match |
| | C24 H34 O | Full match | Full match | No results | Partial mat |
| | C21 H23 N | Full match | Full match | No results | Partial mat |
| | C27 H17 F | Full match | Invalid ma: | No results | Full match |
| | C26 H28 F2 | Full match | Invalid ma: | No results | Full match |
| | C12 H11 N | Full match | Full match | No results | Partial mat |
| | C17 H34 O | Full match | Full match | No results | Partial mat |
| | C17 H34 O | Full match | Full match | No results | Partial mat |
| | C17 H34 O | Full match | Full match | No results | Partial mat |
| | C17 H34 O | Full match | Full match | No results | Partial mat |
| | C17 H34 O | Full match | Full match | No results | Partial mat |
| | C21 H24 F2 | Full match | Invalid ma: | No results | Full match |
| | C13 H14 N | Full match | Not the to| | No results | Partial mat |
| | C18 H22 O | Full match | Invalid ma: | No results | Full match |
| | C15 H20 O | Full match | Not the to| | No results | Not the to|
| | C12 H13 N | Full match | Full match | No results | Partial mat |
| | C12 H13 N | Full match | Full match | No results | Partial mat |
| | C13 H22 N | Full match | Invalid ma: | No results | Full match |
| | C23 H23 F | Full match | Invalid ma: | No results | Full match |
| | C17 H24 O | Full match | Invalid ma: | No results | Partial mat |
| | C22 H29 F | Full match | Unused | No results | Not the to| |
| | C19 H20 O | Full match | Invalid ma: | No results | Full match |
| | C25 H30 N | Invalid ma: | Unused | No results | Invalid ma: |
| | C21 H30 N | Full match | Invalid ma: | No results | Full match |
| | C14 H10 N | Full match | Invalid ma: | No results | Full match |
| | C15 H21 N | Full match | Invalid ma: | No results | Full match |
| | C21 H40 O | Full match | Invalid ma: | No results | Full match |
| | C14 H21 N | Full match | Invalid ma: | No results | Partial mat |
| | C19 H19 N | Full match | Invalid ma: | No results | Full match |
| | C17 H12 N | Full match | Invalid ma: | No results | Full match |
| | C12 H10 N | Full match | Full match | No results | Partial mat |
| | C17 H14 O | Full match | Full match | No results | Partial mat |
| | C17 H28 O | Full match | Invalid ma: | No results | Full match |
| | C13 H24 O | Full match | Full match | No results | Partial mat |
| | C13 H24 O | Full match | Full match | No results | Partial mat |
| | C13 H24 O | Full match | Full match | No results | Partial mat |
| | C19 H19 N | Full match | Invalid ma: | No results | Full match |
| | C18 H26 | Full match | Invalid ma: | No results | Full match |
| | C15 H20 O | Full match | Full match | No results | Partial mat |
| | C16 H19 N | Full match | Full match | No results | Partial mat |
| | C16 H19 N | Full match | Full match | No results | Partial mat |
| | C14 H14 O | Full match | Invalid ma: | No results | Full match |
| | C20 H22 O | Full match | Not the to| | No results | Partial mat |
| | C19 H24 Cl | Full match | Invalid ma: | No results | Not the to| |
| | C15 H23 N | Full match | Not the to| | No results | Full match |
| | C13 H9 N5 | Full match | Invalid ma: | No results | Full match |
| | C25 H39 F3 | Full match | Invalid ma: | No results | Full match |

| | | | | |
|---|---|---|---|---|
| C16 H17 N | Full match | Invalid ma: | No results | Not the to| |
| C15 H23 N | Full match | Not the to| | No results | Partial mat |
| C17 H18 O | Full match | Invalid ma: | No results | Full match |
| C11 H20 N | Full match | Full match | No results | Partial mat |
| C20 H24 N | Full match | Invalid ma: | No results | Full match |
| C15 H18 O | Full match | Invalid ma: | No results | Partial mat |
| C15 H18 O | Full match | Invalid ma: | No results | Not the to| |
| C20 H24 O | Full match | Invalid ma: | No results | Full match |
| C17 H17 N | Full match | Unused | No results | Full match |
| C22 H40 N | Full match | No match | No results | Full match |
| C13 H16 N | Full match | Invalid ma: | No results | Partial mat |
| C13 H27 N | Full match | Invalid ma: | No results | Full match |
| C13 H27 N | Full match | Invalid ma: | No results | Full match |
| C21 H22 O | Full match | Invalid ma: | No results | Full match |
| C20 H28 O | Full match | Full match | No results | Not the to| |
| C16 H27 N | Full match | Invalid ma: | No results | Full match |
| C21 H34 O | Full match | Not the to| | No results | Partial mat |
| C20 H14 N | Full match | Invalid ma: | No results | Not the to| |
| C18 H22 F | Full match | Invalid ma: | No results | Not the to| |
| C20 H20 O | Full match | Invalid ma: | No results | Full match |
| C13 H26 N | Full match | No match | No results | Not the to| |
| C13 H11 F | Full match | Unused | No results | Not the to| |
| C16 H24 N | Full match | Full match | No results | Partial mat |
| C16 H24 N | Full match | Full match | No results | Partial mat |
| C16 H24 N | Full match | Full match | No results | Partial mat |
| C11 H13 N | Full match | Invalid ma: | No results | Partial mat |
| C16 H13 N | Full match | Not the to| | No results | Partial mat |
| C17 H20 O | Full match | Invalid ma: | No results | Full match |
| C23 H23 N | Full match | Invalid ma: | No results | Full match |
| C27 H34 N | Full match | Invalid ma: | No results | Not the to| |
| C21 H19 Cl | Full match | Invalid ma: | No results | Not the to| |
| C20 H30 O | Full match | Full match | No results | Partial mat |
| C16 H9 N ( | Full match | Invalid ma: | No results | Full match |
| C15 H22 O | Full match | Full match | No results | Partial mat |
| C15 H22 O | Full match | Full match | No results | Partial mat |
| C15 H22 O | Full match | Full match | No results | Partial mat |
| C15 H22 O | Full match | Full match | No results | Partial mat |
| C16 H10 O | Full match | Invalid ma: | No results | Full match |
| C15 H24 O | Full match | Invalid ma: | No results | Full match |
| C14 H27 Cl | Full match | Invalid ma: | No results | Not the to| |
| C29 H44 O | Full match | Invalid ma: | No results | Full match |
| C29 H44 O | Full match | Invalid ma: | No results | Full match |
| C20 H27 N | Full match | Invalid ma: | No results | Full match |
| C9 H21 O4 | Full match | Invalid ma: | No results | Full match |
| C13 H27 N | Invalid ma: | No match | No results | No match |
| C20 H39 N | Full match | Invalid ma: | No results | Full match |
| C20 H39 N | Full match | Invalid ma: | No results | Full match |
| C15 H19 N | Full match | Invalid ma: | No results | Full match |
| C12 H12 O | Full match | Invalid ma: | No results | Full match |
| C19 H19 N | Full match | Not the to| | No results | Partial mat |
| C16 H14 N | Full match | Full match | No results | Partial mat |

| | | | | |
|---|---|---|---|---|
| C15 H15 N | Full match | Invalid ma: | No results | Full match |
| C33 H47 N | Full match | Invalid ma: | No results | Full match |
| C16 H18 O | Full match | Full match | No results | Partial mat |
| C29 H27 F | Full match | Invalid ma: | No results | Full match |
| C15 H13 F: | Full match | Invalid ma: | No results | Full match |
| C14 H20 O | Full match | Invalid ma: | No results | Full match |
| C21 H23 N | Full match | Not the to| | No results | Partial mat |
| C21 H32 O | Full match | Invalid ma: | No results | Full match |
| C18 H20 N | Full match | Not the to| | No results | Partial mat |
| C20 H37 N | Full match | Invalid ma: | No results | Full match |
| C20 H37 N | Full match | Invalid ma: | No results | Full match |
| C18 H20 F | Full match | Invalid ma: | No results | Full match |
| C20 H34 N | Invalid ma: | Invalid ma: | No results | Invalid ma: |
| C16 H31 N | Full match | Invalid ma: | No results | Full match |
| C16 H31 N | Full match | Invalid ma: | No results | Full match |
| C14 H22 F | Invalid ma: | No match | No results | No match |
| C14 H22 F | Full match | No match | No results | Not the to| |
| C28 H33 N | Full match | No match | No results | Not the to| |
| C19 H23 N | Full match | Full match | No results | Partial mat |
| C19 H23 N | Full match | Full match | No results | Partial mat |
| C15 H18 N | Full match | Not the to| | No results | Partial mat |
| C21 H30 F | Full match | Invalid ma: | No results | Not the to| |
| C17 H24 N | Full match | Full match | No results | Partial mat |
| C17 H24 N | Full match | Full match | No results | Partial mat |
| C23 H23 N | Full match | Invalid ma: | No results | Full match |
| C14 H18 N | Full match | Full match | No results | Partial mat |
| C22 H44 N | Full match | Invalid ma: | No results | Full match |
| C24 H38 N | Full match | Invalid ma: | No results | Full match |
| C27 H30 F: | Full match | Invalid ma: | No results | Full match |
| C18 H22 N | Full match | Invalid ma: | No results | Full match |
| C19 H26 N | Full match | Invalid ma: | No results | Full match |

| Annot_Source_MassList_Search | Calc_MW | m_z | RT_min_ | Area_Max_ | _ChemSpider_Results | _mzCloud_Results | _mzVault_Results | mzCloud_Best_Match |
|---|---|---|---|---|---|---|---|---|
| Full match | 313.1314 | 314.1387 | 22.134 | 1.08E+09 | 286 | 6 | 0 | 80.9 |
| Full match | 133.0641 | 134.0714 | 12.817 | 1.79E+08 | 23 | 4 | 0 | 88.2 |
| Full match | 133.0642 | 134.0715 | 13.015 | 92808888 | 23 | 4 | 0 | 67.5 |
| Full match | 332.0684 | 333.0757 | 19.103 | 1.26E+09 | 15 | 4 | 0 | 99.2 |
| No results | 224.118 | 225.1252 | 19.355 | 1.91E+08 | 1 | 1 | 0 | 95.1 |
| No results | 224.118 | 225.1252 | 19.732 | 63376112 | 1 | 1 | 0 | 89.3 |
| Full match | 398.2432 | 399.2505 | 24.462 | 5.19E+08 | 29 | 6 | 1 | 99.3 |
| Full match | 283.9539 | 284.9612 | 18.197 | 6500166 | 1 | 1 | 0 | 38.9 |
| Full match | 348.1278 | 349.135 | 23.914 | 3196964 | 41 | 0 | 1 | |
| Full match | 312.1361 | 313.1434 | 22.052 | 8275782 | 132 | 2 | 0 | 43.4 |
| Full match | 366.2043 | 367.2116 | 18.724 | 28060308 | 234 | 1 | 0 | 30.8 |
| Partial mat | 312.1363 | 313.1435 | 19.403 | 14454056 | 132 | 3 | 0 | 44.3 |
| Partial mat | 268.1578 | 269.1651 | 23.322 | 10134925 | 213 | 4 | 0 | 71.7 |
| Full match | 342.1467 | 343.154 | 17.314 | 41474773 | 166 | 3 | 0 | 85.8 |
| Full match | 342.1467 | 343.154 | 17.166 | 30239134 | 166 | 2 | 0 | 84.8 |
| Full match | 342.1468 | 343.154 | 15.424 | 17140682 | 165 | 2 | 0 | 84.1 |
| Full match | 266.1649 | 267.1722 | 23.809 | 4.38E+08 | 3 | 3 | 2 | 99.5 |
| No results | 433.2254 | 434.2327 | 14.04 | 81170744 | 78 | 2 | 0 | 67.9 |
| No results | 274.2145 | 275.2218 | 21.357 | 55846242 | 8 | 1 | 0 | 44.6 |
| Partial mat | 184.1002 | 185.1075 | 9.059 | 8760804 | 46 | 2 | 0 | 31.9 |
| No results | 478.3146 | 479.3218 | 23.229 | 18168115 | 0 | 1 | 0 | 33.8 |
| No results | 346.2143 | 347.2216 | 18.217 | 3.27E+08 | 103 | 4 | 0 | 55.9 |
| No results | 346.2144 | 347.2216 | 17.25 | 5.25E+08 | 103 | 3 | 0 | 64.4 |
| No results | 346.2144 | 347.2217 | 19.082 | 32419417 | 103 | 4 | 0 | 58.4 |
| No results | 346.2144 | 347.2217 | 18.787 | 10423166 | 103 | 4 | 0 | 59 |
| Full match | 362.2093 | 363.2166 | 17.087 | 45182411 | 92 | 2 | 0 | 46.4 |
| Full match | 362.2093 | 363.2166 | 21.373 | 26738363 | 92 | 1 | 0 | 33.2 |
| Partial mat | 302.2246 | 303.2319 | 23.256 | 19272474 | 253 | 15 | 0 | 68.8 |
| No results | 316.2402 | 317.2475 | 23.975 | 8025877 | 106 | 3 | 0 | 46.8 |
| No results | 306.1329 | 307.1402 | 10.047 | 36929107 | 43 | 5 | 0 | 94.7 |
| No results | 380.2199 | 381.2272 | 14.877 | 8862753 | 34 | 1 | 0 | 33.9 |
| No results | 316.1675 | 317.1748 | 22.484 | 39198401 | 91 | 2 | 0 | 48.5 |
| No results | 316.1674 | 317.1747 | 21.55 | 24265628 | 89 | 1 | 0 | 31.3 |
| No results | 316.1675 | 317.1747 | 18.84 | 47762514 | 89 | 2 | 0 | 44.3 |
| No results | 305.1991 | 306.2064 | 20.001 | 26413218 | 98 | 1 | 0 | 38.6 |
| Full match | 266.152 | 267.1593 | 21.195 | 23008919 | 144 | 10 | 0 | 46.4 |
| Full match | 266.1522 | 267.1594 | 20.21 | 33690232 | 144 | 11 | 0 | 62 |
| Full match | 266.1521 | 267.1594 | 19.502 | 18966819 | 144 | 12 | 0 | 61.6 |
| Full match | 266.1521 | 267.1594 | 19.784 | 24801892 | 144 | 11 | 0 | 60.5 |
| Full match | 266.152 | 267.1593 | 15.67 | 10316713 | 144 | 7 | 0 | 54.2 |

| | | | | | | | | |
|---|---|---|---|---|---|---|---|---|
| No results | 286.1569 | 287.1644 | 19.578 | 15334105 | 73 | 2 | 0 | 47 |
| Full match | 330.2195 | 331.2268 | 23.01 | 24060418 | 170 | 7 | 0 | 80.7 |
| No results | 300.1726 | 301.1799 | 23.254 | 35843717 | 119 | 3 | 0 | 34.9 |
| No results | 304.2039 | 305.2112 | 22.714 | 57706418 | 97 | 2 | 0 | 54.9 |
| No results | 304.204 | 305.2112 | 24.353 | 26229530 | 97 | 2 | 0 | 53.7 |
| No results | 332.2351 | 333.2423 | 21.102 | 1.27E+09 | 132 | 3 | 0 | 69.7 |
| No results | 332.2352 | 333.2425 | 19.724 | 1.11E+08 | 132 | 3 | 0 | 66.2 |
| No results | 332.2351 | 333.2424 | 23.849 | 12654044 | 132 | 3 | 0 | 65.6 |
| Full match | 360.1936 | 361.201 | 17.279 | 26843860 | 84 | 11 | 1 | 58 |
| No results | 288.1727 | 289.18 | 17.426 | 16073264 | 71 | 3 | 0 | 44.8 |
| No results | 288.1726 | 289.1799 | 20.461 | 13824952 | 71 | 3 | 0 | 37 |
| No results | 316.2402 | 317.2475 | 21.53 | 18480598 | 106 | 5 | 0 | 57.4 |
| No results | 316.2403 | 317.2476 | 19.54 | 58792420 | 106 | 6 | 1 | 71.8 |
| Full match | 364.225 | 365.2322 | 15.146 | 17820640 | 64 | 2 | 0 | 60.1 |
| No results | 335.1082 | 336.1155 | 11.624 | 9607074 | 104 | 7 | 0 | 83.4 |
| No results | 318.2195 | 319.2267 | 23.08 | 42774961 | 296 | 16 | 0 | 55.7 |
| No results | 318.2196 | 319.2269 | 24.115 | 17659283 | 296 | 16 | 0 | 44.7 |
| No results | 318.1831 | 319.1904 | 21.447 | 86861092 | 61 | 1 | 0 | 62.6 |
| No results | 318.1832 | 319.1905 | 20.505 | 97882920 | 61 | 1 | 0 | 45.4 |
| No results | 318.1831 | 319.1904 | 19.578 | 45607268 | 61 | 1 | 0 | 70.6 |
| No results | 318.1831 | 319.1904 | 18.267 | 32680338 | 61 | 1 | 0 | 58.9 |
| No results | 254.152 | 255.1593 | 21.092 | 38649598 | 50 | 2 | 0 | 42.2 |
| No results | 294.2196 | 295.2269 | 23.034 | 47500802 | 35 | 6 | 0 | 56.3 |
| No results | 294.2197 | 295.227 | 23.835 | 23363953 | 35 | 6 | 0 | 61.1 |
| No results | 194.1308 | 195.1382 | 22.758 | 19352939 | 48 | 1 | 0 | 32.2 |
| Partial mat | 224.1052 | 225.1125 | 17.265 | 7590690 | 76 | 5 | 0 | 41.5 |
| No results | 294.1256 | 295.1329 | 20.328 | 13320041 | 134 | 6 | 0 | 38.3 |
| No results | 427.236 | 428.2433 | 13.604 | 7001117 | 54 | 1 | 0 | 79.9 |
| Full match | 285.1366 | 286.1439 | 21.934 | 14397056 | 276 | 4 | 0 | 63.8 |
| No results | 346.2145 | 347.2218 | 23.276 | 14337256 | 103 | 3 | 0 | 49.1 |
| Not the to | 258.1833 | 259.1906 | 24.687 | 2.87E+08 | 12 | 3 | 0 | 94.1 |
| Full match | 276.1727 | 277.18 | 21.142 | 64024035 | 56 | 2 | 0 | 42.2 |
| No results | 306.1581 | 307.1654 | 14.988 | 8424092 | 145 | 2 | 0 | 32.5 |
| Full match | 266.1521 | 267.1594 | 19.353 | 10110125 | 144 | 12 | 0 | 66 |
| No results | 246.1258 | 247.1331 | 20.715 | 39529559 | 176 | 9 | 0 | 52.5 |
| No results | 246.1258 | 247.1331 | 19.03 | 32422991 | 176 | 12 | 0 | 57.7 |
| Partial mat | 248.1415 | 249.1488 | 18.258 | 13497276 | 245 | 15 | 0 | 73.9 |
| No results | 346.1779 | 347.1852 | 16.809 | 21743374 | 157 | 3 | 0 | 52.5 |
| No results | 218.1672 | 219.1745 | 24.196 | 86325104 | 75 | 5 | 0 | 89.4 |
| No results | 262.1208 | 263.128 | 18.86 | 23688709 | 171 | 8 | 0 | 50.7 |
| No results | 262.1207 | 263.128 | 15.713 | 8591524 | 171 | 3 | 0 | 41.7 |
| No results | 262.1207 | 263.128 | 20.933 | 11444267 | 171 | 8 | 0 | 53.9 |
| No results | 252.1727 | 253.18 | 20.671 | 74342849 | 116 | 5 | 0 | 46.9 |
| No results | 252.1727 | 253.18 | 21.098 | 1.3E+08 | 116 | 5 | 0 | 59.1 |
| No results | 346.2507 | 347.258 | 23.975 | 16602295 | 78 | 3 | 0 | 53.2 |
| No results | 274.1571 | 275.1644 | 23.191 | 12336569 | 66 | 7 | 0 | 75.4 |
| No results | 328.2249 | 346.2587 | 22.731 | 31412279 | 16 | 1 | 0 | 43.2 |
| No results | 332.1988 | 333.2061 | 22.56 | 56942912 | 229 | 15 | 0 | 51.9 |
| No results | 332.1988 | 333.206 | 22.033 | 32528691 | 229 | 15 | 0 | 42.4 |
| Full match | 181.0021 | 182.0094 | 21.639 | 9.03E+08 | 6 | 2 | 0 | 98.1 |
| No results | 371.1367 | 372.144 | 22.542 | 1.21E+08 | 109 | 6 | 0 | 44.8 |

| | | | | | | | | |
|---|---|---|---|---|---|---|---|---|
| Not the to| | 302.2247 | 303.2319 | 24.698 | 8377783 | 253 | 15 | 0 | 62.9 |
| No results | 306.1468 | 307.1541 | 20.195 | 17733735 | 93 | 2 | 0 | 39.3 |
| No results | 332.1625 | 333.1698 | 15.85 | 7983169 | 80 | 5 | 0 | 48.1 |
| No results | 345.2515 | 346.2587 | 22.603 | 21514343 | 8 | 1 | 0 | 39.7 |
| No results | 345.2515 | 346.2588 | 22.729 | 31412279 | 8 | 1 | 0 | 46.3 |
| No results | 450.2772 | 451.2844 | 21.94 | 15624874 | 27 | 2 | 0 | 55.3 |
| Full match | 250.1207 | 251.128 | 20.016 | 7227313 | 80 | 3 | 0 | 45.6 |
| Full match | 346.1417 | 347.149 | 18.602 | 23646297 | 130 | 2 | 0 | 33.6 |
| No results | 472.3191 | 473.3263 | 21.227 | 7121566 | 47 | 1 | 0 | 35.3 |
| No results | 314.1882 | 315.1955 | 23.47 | 17045877 | 140 | 3 | 0 | 37 |
| Full match | 314.1518 | 315.1591 | 20.891 | 41038091 | 124 | 3 | 0 | 50.6 |
| Partial mat | 250.1571 | 251.1644 | 22.691 | 1.31E+08 | 228 | 8 | 1 | 76.3 |
| No results | 298.157 | 299.1643 | 20.9 | 24442143 | 115 | 3 | 0 | 45.3 |
| Full match | 302.1883 | 303.1956 | 22.519 | 27122901 | 123 | 8 | 0 | 54.4 |
| Full match | 302.1882 | 303.1955 | 22.775 | 12991367 | 123 | 8 | 0 | 54.4 |
| Full match | 272.1779 | 273.1851 | 24.84 | 1.51E+08 | 63 | 4 | 0 | 95.3 |
| Full match | 272.1779 | 273.1852 | 24.92 | 75285471 | 63 | 4 | 0 | 95.3 |
| Full match | 272.1778 | 273.1851 | 24.659 | 39919118 | 63 | 4 | 0 | 97.3 |
| Full match | 254.1672 | 255.1745 | 21.001 | 8755859 | 11 | 1 | 0 | 32.9 |
| Full match | 341.0698 | 342.0771 | 23.27 | 3.64E+08 | 25 | 2 | 1 | 87.4 |
| Full match | 307.1452 | 308.1525 | 22.629 | 5450557 | 28 | 2 | 1 | 81.5 |
| Full match | 191.1312 | 192.1384 | 18.932 | 1.04E+08 | 92 | 10 | 1 | 95.7 |
| Partial mat | 251.1524 | 252.1597 | 16.004 | 28095760 | 120 | 1 | 0 | 30.1 |
| Full match | 251.1345 | 252.1418 | 24.815 | 28485327 | 29 | 3 | 1 | 96.7 |
| Full match | 241.1363 | 242.1436 | 17.318 | 3.26E+08 | 3 | 3 | 2 | 93.5 |
| No results | 388.2072 | 389.2145 | 15.649 | 58616322 | 18 | 4 | 0 | 79.4 |
| Full match | 279.1472 | 280.1545 | 13.282 | 30114865 | 115 | 4 | 0 | 89.2 |
| No results | 266.0945 | 267.1018 | 21.461 | 1.2E+08 | 156 | 1 | 0 | 34.9 |
| No results | 266.0945 | 267.1017 | 21.23 | 22426268 | 156 | 1 | 0 | 39.4 |
| No results | 250.0996 | 251.1068 | 22.732 | 2.65E+08 | 82 | 1 | 0 | 39 |
| Full match | 248.1778 | 249.1851 | 21.209 | 6.31E+08 | 52 | 3 | 0 | 74.7 |
| Full match | 151.0094 | 152.0166 | 14.57 | 18158760 | 10 | 2 | 0 | 30.7 |
| No results | 153.1155 | 154.1228 | 14.394 | 1.06E+08 | 27 | 1 | 0 | 81.3 |
| Partial mat | 241.2772 | 242.2844 | 17.706 | 35609557 | 4 | 1 | 0 | 74 |
| Partial mat | 241.2772 | 242.2845 | 14.07 | 24364949 | 4 | 1 | 0 | 69.2 |
| No results | 311.1999 | 312.2072 | 21.119 | 46697618 | 141 | 4 | 0 | 37.3 |
| Full match | 210.1023 | 211.1096 | 23.809 | 20566226 | 3 | 1 | 1 | 84.2 |
| Partial mat | 185.2146 | 186.2219 | 14.283 | 74267350 | 3 | 1 | 0 | 31 |
| Partial mat | 269.3085 | 270.3158 | 19.183 | 10216219 | 2 | 2 | 0 | 74.7 |
| Full match | 244.1214 | 245.1287 | 17.332 | 12532400 | 295 | 2 | 0 | 39.1 |
| Full match | 324.2203 | 325.2275 | 13.463 | 37927779 | 106 | 1 | 0 | 32.5 |
| Not the to| | 250.1935 | 251.2007 | 21.25 | 28123751 | 43 | 1 | 0 | 59.3 |
| Full match | 250.1935 | 251.2007 | 22.875 | 30019510 | 43 | 1 | 0 | 34.3 |
| Full match | 222.1371 | 223.1444 | 15.338 | 1.24E+08 | 121 | 4 | 0 | 82.9 |
| No results | 592.3265 | 593.3338 | 23.442 | 5593656 | 14 | 1 | 0 | 49.1 |
| No results | 274.1934 | 275.2007 | 22.534 | 44092900 | 54 | 3 | 0 | 35.9 |
| No results | 279.1836 | 280.1909 | 10.233 | 9162629 | 113 | 1 | 0 | 36.9 |
| Full match | 295.0168 | 296.024 | 22.72 | 1.04E+08 | 37 | 2 | 1 | 92.6 |
| Full match | 287.1523 | 288.1596 | 22.205 | 20067823 | 218 | 1 | 0 | 86.2 |
| No results | 357.0768 | 358.084 | 22.766 | 13000428 | 55 | 1 | 1 | 86.2 |
| Partial mat | 254.0946 | 255.1018 | 20.294 | 22723761 | 158 | 3 | 0 | 50.4 |

| | | | | | | | | |
|---|---|---|---|---|---|---|---|---|
| Partial mat | 254.0944 | 255.1017 | 20.643 | 9682655 | 158 | 3 | 0 | 40.9 |
| Full match | 230.0946 | 231.1019 | 20.408 | 38281852 | 112 | 6 | 1 | 84.4 |
| Full match | 188.0952 | 189.1025 | 15.444 | 14767139 | 119 | 4 | 0 | 92.1 |
| No results | 261.1732 | 262.1805 | 10.752 | 63592442 | 192 | 5 | 0 | 74.7 |
| Full match | 247.1575 | 248.1648 | 9.635 | 19373389 | 246 | 5 | 0 | 48.4 |
| No results | 309.2094 | 310.2166 | 15.028 | 1.32E+08 | 130 | 3 | 0 | 92.3 |
| No results | 271.1211 | 272.1284 | 14.013 | 23601981 | 242 | 2 | 0 | 85.4 |
| No results | 221.1781 | 222.1854 | 21.429 | 2.17E+09 | 42 | 1 | 0 | 90.2 |
| Partial mat | 263.1888 | 264.196 | 9.666 | 1.31E+08 | 163 | 6 | 0 | 99.3 |
| Partial mat | 263.1888 | 264.1961 | 10.297 | 13036760 | 163 | 6 | 0 | 62.8 |
| Full match | 191.1312 | 192.1384 | 17.885 | 23157964 | 92 | 11 | 0 | 85.6 |
| No results | 321.059 | 322.0663 | 20.776 | 27957949 | 26 | 2 | 0 | 54.3 |
| No results | 266.1058 | 250.0865 | 21.669 | 39858508 | 358 | 1 | 0 | 37.6 |
| No results | 748.5084 | 375.2615 | 10.893 | 2.12E+08 | 13 | 1 | 1 | 39.6 |
| No results | 747.4757 | 748.483 | 15.829 | 20619508 | 9 | 1 | 1 | 92.8 |
| Full match | 836.5244 | 419.2696 | 16.119 | 29397158 | 9 | 2 | 1 | 93.5 |
| No results | 353.1198 | 354.1271 | 14.065 | 11828466 | 257 | 8 | 0 | 58.4 |
| Full match | 290.1381 | 291.1453 | 8.371 | 7276425 | 99 | 2 | 1 | 96.6 |
| No results | 307.1937 | 308.201 | 16.376 | 1.45E+08 | 134 | 1 | 0 | 31.4 |
| Full match | 435.0657 | 436.073 | 17.746 | 8273502 | 21 | 3 | 0 | 89.5 |
| No results | 270.1007 | 271.108 | 13.104 | 41957486 | 254 | 1 | 0 | 82 |
| Full match | 254.1058 | 255.1131 | 14.002 | 90137100 | 231 | 1 | 0 | 91.5 |
| Full match | 254.1057 | 255.113 | 15.63 | 31725486 | 242 | 1 | 0 | 64.1 |
| Full match | 252.0902 | 253.0974 | 15.566 | 10083647 | 239 | 9 | 0 | 67.8 |
| Full match | 252.0901 | 253.0974 | 14.746 | 10499134 | 239 | 9 | 0 | 63.2 |
| Full match | 236.0952 | 237.1025 | 17.482 | 1.25E+08 | 185 | 6 | 1 | 99 |
| Full match | 255.0081 | 256.0153 | 9.422 | 1.12E+08 | 9 | 1 | 0 | 99.3 |
| Full match | 252.0901 | 253.0974 | 15.844 | 51265175 | 239 | 6 | 1 | 94.3 |
| No match | 338.1153 | 339.1226 | 21.904 | 8200992 | 13 | 2 | 0 | 35 |
| No results | 292.098 | 293.1053 | 15.6 | 1.55E+08 | 44 | 6 | 0 | 87.7 |
| Full match | 291.1988 | 292.2061 | 15.894 | 1.11E+09 | 65 | 3 | 0 | 78.1 |
| Full match | 388.1554 | 389.1626 | 16.705 | 1.71E+08 | 46 | 1 | 1 | 93.8 |
| Full match | 501.2882 | 502.2955 | 15.663 | 2.43E+09 | 34 | 2 | 1 | 99.5 |
| Full match | 432.2515 | 433.2588 | 23.063 | 6325113 | 50 | 1 | 0 | 39.6 |
| No results | 298.1205 | 299.1278 | 20.047 | 7789807 | 175 | 1 | 0 | 54.1 |
| No results | 203.1312 | 204.1385 | 20.481 | 8579584 | 145 | 5 | 0 | 47.5 |
| Full match | 252.1365 | 253.1437 | 21.892 | 24367042 | 73 | 3 | 0 | 88.3 |
| Partial mat | 334.2145 | 335.2218 | 22.831 | 99612586 | 237 | 20 | 0 | 59.6 |
| Full match | 364.1887 | 365.1959 | 22.43 | 33889530 | 99 | 2 | 0 | 41.4 |
| No results | 430.061 | 431.0683 | 21.603 | 11426080 | 5 | 1 | 1 | 73.3 |
| Full match | 384.23 | 385.2373 | 22.001 | 7797389 | 83 | 3 | 0 | 66 |
| Full match | 389.2064 | 390.2137 | 11.162 | 13094727 | 25 | 4 | 0 | 92.9 |
| No results | 391.2148 | 392.222 | 12.813 | 12623377 | 64 | 2 | 0 | 87.8 |
| Full match | 267.1837 | 268.1909 | 9.761 | 3.05E+08 | 49 | 2 | 1 | 99.7 |
| Full match | 267.1838 | 268.191 | 19.872 | 39242712 | 49 | 10 | 0 | 54.1 |
| No results | 259.1575 | 260.1648 | 12.368 | 60220305 | 134 | 2 | 1 | 95.3 |
| No results | 290.1996 | 291.2069 | 12.371 | 94265363 | 173 | 2 | 0 | 95.9 |
| Full match | 325.2254 | 326.2327 | 11.55 | 15671337 | 22 | 1 | 1 | 73.4 |
| No results | 440.1598 | 441.1671 | 19.786 | 92563425 | 50 | 2 | 1 | 89 |
| Full match | 414.1615 | 415.1688 | 14.085 | 19350041 | 131 | 8 | 1 | 92.9 |
| Not the to | 414.138 | 415.1453 | 13.563 | 85057151 | 13 | 3 | 0 | 95.7 |

| | | | | | | | | |
|---|---|---|---|---|---|---|---|---|
| Full match | 328.1788 | 329.186 | 11.969 | 12816046 | 320 | 2 | 0 | 55.1 |
| No results | 422.1622 | 423.1695 | 19.089 | 8.68E+08 | 26 | 2 | 1 | 97.1 |
| Full match | 224.189 | 225.1963 | 20.369 | 1.92E+08 | 31 | 1 | 0 | 94.4 |
| Full match | 435.2271 | 436.2343 | 21.692 | 1.05E+08 | 146 | 1 | 1 | 95.4 |
| No results | 402.2406 | 403.2479 | 22.154 | 8955331 | 179 | 1 | 0 | 33.8 |
| No results | 318.0658 | 319.0731 | 22.908 | 3703364 | 27 | 1 | 0 | 49.1 |
| Partial mat | 250.1571 | 251.1644 | 23.747 | 1.43E+08 | 228 | 2 | 0 | 94.9 |
| Full match | 239.1677 | 240.175 | 13.144 | 86742246 | 49 | 1 | 0 | 33 |
| Full match | 244.1943 | 245.2015 | 13.658 | 15003994 | 40 | 3 | 0 | 87 |
| No results | 330.0077 | 331.015 | 17.745 | 32187524 | 6 | 0 | 1 | |
| Full match | 330.2194 | 331.2267 | 20.478 | 3.67E+08 | 170 | 7 | 0 | 85 |
| Full match | 277.1833 | 278.1906 | 13.626 | 1.35E+08 | 63 | 5 | 0 | 87.5 |
| Full match | 327.1471 | 328.1544 | 23.845 | 10408980 | 300 | 5 | 0 | 62.3 |
| Full match | 344.1988 | 345.2061 | 22.37 | 13226442 | 129 | 2 | 0 | 43.2 |
| Full match | 286.1934 | 287.2007 | 20.71 | 18403532 | 78 | 2 | 0 | 36.4 |
| No results | 321.173 | 322.1803 | 13.698 | 70551384 | 151 | 2 | 0 | 37.4 |
| No results | 258.1623 | 259.1695 | 18.01 | 7711226 | 59 | 3 | 0 | 47.7 |
| Full match | 302.1883 | 303.1956 | 22.921 | 36239034 | 123 | 8 | 0 | 53.6 |
| Full match | 302.1882 | 303.1955 | 20.62 | 34747304 | 123 | 8 | 0 | 62.6 |
| Full match | 302.1882 | 303.1955 | 22.252 | 15433510 | 123 | 6 | 0 | 44.5 |
| No results | 387.1795 | 388.1867 | 11.95 | 7609638 | 89 | 1 | 0 | 34.6 |
| No results | 218.1421 | 219.1494 | 20.082 | 12315484 | 145 | 1 | 0 | 32.4 |
| No results | 265.2044 | 266.2117 | 21.611 | 35014450 | 38 | 1 | 0 | 49.6 |
| Full match | 328.2038 | 329.2111 | 17.261 | 48767776 | 125 | 1 | 0 | 56.1 |
| Full match | 328.2038 | 329.2111 | 20.121 | 29674054 | 125 | 1 | 0 | 66.4 |
| Full match | 328.2038 | 329.2111 | 22.534 | 11292713 | 125 | 1 | 0 | 51.8 |
| No results | 220.089 | 221.0963 | 20.527 | 58314221 | 16 | 2 | 0 | 51.6 |
| Full match | 320.126 | 321.1333 | 19.856 | 58727587 | 54 | 10 | 0 | 90.2 |
| No results | 474.2256 | 475.2329 | 17.378 | 8322670 | 175 | 3 | 0 | 48.3 |
| No results | 312.2089 | 313.2162 | 23.261 | 8219907 | 112 | 2 | 0 | 51.1 |
| No match | 384.23 | 385.2373 | 22.427 | 12783351 | 1 | 3 | 0 | 49.2 |
| Full match | 310.1483 | 311.1556 | 13.503 | 79700616 | 61 | 5 | 0 | 64.8 |
| Full match | 265.1469 | 266.1541 | 22.328 | 27500987 | 136 | 4 | 0 | 66.4 |
| Full match | 293.1992 | 294.2065 | 12.34 | 15675738 | 83 | 3 | 0 | 63 |
| Full match | 277.1833 | 278.1906 | 15.11 | 1.07E+08 | 63 | 5 | 0 | 82.2 |
| Full match | 277.1832 | 278.1904 | 15.757 | 54850519 | 63 | 5 | 0 | 46.6 |
| Full match | 315.0849 | 316.0922 | 16.039 | 42103635 | 11 | 3 | 0 | 85.7 |
| Full match | 315.0849 | 316.0921 | 16.289 | 32258334 | 11 | 3 | 0 | 62.4 |
| Full match | 324.1639 | 325.1712 | 13.66 | 2.42E+08 | 54 | 6 | 1 | 95 |
| Full match | 263.1677 | 264.1749 | 14.956 | 19314102 | 71 | 5 | 0 | 49.7 |
| Partial mat | 263.1888 | 264.196 | 9.046 | 58588407 | 163 | 6 | 0 | 96.6 |
| Partial mat | 263.1889 | 264.1961 | 11.234 | 15892142 | 163 | 6 | 0 | 49.3 |
| Full match | 286.051 | 287.0583 | 18.2 | 57850171 | 78 | 13 | 0 | 93.7 |
| Full match | 305.0739 | 306.0812 | 15.825 | 42596886 | 28 | 1 | 1 | 87.2 |
| Partial mat | 277.2044 | 278.2116 | 11.412 | 2.76E+08 | 104 | 1 | 1 | 98.5 |
| Partial mat | 277.2046 | 278.2118 | 11.573 | 64738967 | 104 | 1 | 1 | 96.5 |
| Partial mat | 277.2044 | 278.2117 | 11.224 | 12792244 | 104 | 1 | 0 | 34.6 |
| Partial mat | 264.1265 | 265.1337 | 20.01 | 23473402 | 221 | 3 | 0 | 50.4 |
| Partial mat | 284.1777 | 285.185 | 21.216 | 21104286 | 72 | 1 | 0 | 31.7 |
| Partial mat | 358.178 | 359.1853 | 17.903 | 14980024 | 120 | 2 | 1 | 36.8 |
| Full match | 285.1036 | 286.1108 | 21.325 | 13453458 | 37 | 1 | 0 | 66.7 |

| | | | | | | | | |
|---|---|---|---|---|---|---|---|---|
| Full match | 374.2093 | 375.2166 | 19.473 | 27303723 | 109 | 2 | 0 | 40.5 |
| No results | 249.0792 | 250.0865 | 22.879 | 13147292 | 66 | 1 | 0 | 30.4 |
| Full match | 295.0958 | 296.1031 | 16.53 | 13696897 | 196 | 4 | 1 | 86.8 |
| No results | 318.2195 | 319.2267 | 24.851 | 8422488 | 296 | 16 | 0 | 57.9 |
| Partial mat | 264.1728 | 265.1801 | 21.527 | 37369834 | 60 | 3 | 0 | 47.1 |
| Partial mat | 264.1728 | 265.1801 | 20.813 | 45199486 | 60 | 3 | 0 | 44.4 |
| Partial mat | 264.1728 | 265.1801 | 19.977 | 16644056 | 60 | 4 | 0 | 56.8 |
| Partial mat | 264.1728 | 265.1801 | 20.379 | 19302991 | 60 | 1 | 0 | 35.1 |
| No results | 266.1157 | 267.1229 | 17.163 | 9442618 | 52 | 3 | 0 | 35.2 |
| Partial mat | 284.1988 | 317.2322 | 22.12 | 30262725 | 31 | 3 | 0 | 55.1 |
| Partial mat | 284.1989 | 302.2327 | 22.246 | 14757279 | 31 | 3 | 0 | 55 |
| No results | 206.0945 | 207.1018 | 20.539 | 39521302 | 86 | 2 | 0 | 34.7 |
| No results | 206.0945 | 207.1017 | 19.55 | 13193978 | 86 | 1 | 0 | 31.6 |
| No results | 316.2038 | 317.2111 | 22.656 | 54075306 | 272 | 20 | 0 | 73.7 |
| No results | 320.105 | 321.1123 | 20.871 | 85706270 | 49 | 2 | 0 | 38.7 |
| No results | 390.2043 | 391.2116 | 19.983 | 68917771 | 95 | 2 | 0 | 75.6 |
| No results | 316.2039 | 317.2111 | 24.003 | 75014218 | 272 | 19 | 0 | 74.4 |
| No results | 316.2039 | 317.2111 | 22.836 | 98211219 | 272 | 19 | 0 | 69.7 |
| No results | 316.2038 | 317.2111 | 22.947 | 15682416 | 272 | 19 | 0 | 69.7 |
| No results | 316.2039 | 317.2112 | 17.719 | 9038587 | 272 | 8 | 0 | 60.5 |
| Partial mat | 338.2459 | 356.2797 | 15.289 | 38793014 | 102 | 2 | 0 | 36.6 |
| No results | 268.1676 | 269.1749 | 20.834 | 70219417 | 81 | 4 | 0 | 59 |
| Partial mat | 288.2302 | 289.2375 | 24.988 | 51057528 | 10 | 1 | 0 | 31.2 |
| No results | 272.1415 | 273.1487 | 19.008 | 13171872 | 79 | 8 | 0 | 48.7 |
| Full match | 378.2043 | 379.2115 | 22.504 | 15510492 | 47 | 3 | 0 | 44 |
| Full match | 378.2043 | 379.2116 | 23.025 | 20693901 | 47 | 4 | 0 | 43.5 |
| Full match | 378.2043 | 379.2116 | 23.302 | 18180347 | 47 | 1 | 0 | 43.5 |
| Full match | 378.2043 | 379.2116 | 20.622 | 23419472 | 47 | 3 | 0 | 38 |
| No results | 363.1834 | 364.1907 | 22.463 | 13530315 | 184 | 1 | 0 | 37 |
| No results | 293.1417 | 294.149 | 20.501 | 8563114 | 177 | 2 | 0 | 36.2 |
| No results | 456.2514 | 457.2587 | 22.17 | 86830091 | 117 | 1 | 0 | 61.3 |
| Full match | 322.1781 | 323.1854 | 18.712 | 15302978 | 57 | 3 | 0 | 43.7 |
| No results | 456.2513 | 457.2586 | 21.822 | 6.29E+08 | 42 | 1 | 0 | 58.6 |
| No results | 486.2984 | 487.3056 | 20.834 | 1.82E+08 | 44 | 1 | 0 | 34.1 |
| No results | 648.3693 | 649.3766 | 22.31 | 2.74E+08 | 9 | 1 | 0 | 82.6 |
| No results | 421.0929 | 422.1002 | 20.475 | 1.35E+08 | 71 | 6 | 0 | 46.2 |
| No results | 648.3697 | 649.3769 | 22.417 | 4.31E+08 | 4 | 1 | 0 | 82.6 |
| No results | 648.3694 | 649.3767 | 21.682 | 3.02E+08 | 9 | 1 | 0 | 79.9 |
| No results | 715.4505 | 716.4578 | 15.245 | 1.58E+08 | 6 | 1 | 0 | 88.5 |
| No results | 423.2386 | 424.2458 | 21.539 | 70612002 | 1 | 1 | 0 | 32.1 |
| No results | 342.1807 | 343.188 | 19.834 | 88086369 | 126 | 7 | 0 | 47.5 |
| No results | 808.5156 | 405.2652 | 23.778 | 29953707 | 0 | 1 | 0 | 33.4 |
| No results | 486.2985 | 487.3058 | 22.165 | 1E+08 | 83 | 1 | 0 | 51.2 |
| No results | 486.1957 | 487.203 | 21.454 | 66023573 | 8 | 1 | 0 | 49.9 |
| No results | 486.2983 | 487.3057 | 21.933 | 1.07E+08 | 83 | 1 | 0 | 32.3 |
| No results | 444.2512 | 445.2585 | 21.035 | 1.7E+08 | 143 | 1 | 0 | 36 |
| No results | 444.2513 | 445.2585 | 20.04 | 67721811 | 35 | 1 | 0 | 39 |
| No results | 370.2119 | 186.1133 | 17.791 | 73252392 | 47 | 5 | 0 | 68.6 |
| No results | 472.319 | 473.3263 | 20.729 | 69207794 | 47 | 1 | 0 | 48.7 |
| No results | 748.5084 | 375.2615 | 11.006 | 1.91E+08 | 13 | 1 | 0 | 34.2 |
| No results | 311.1999 | 312.2072 | 21.203 | 67254833 | 141 | 4 | 0 | 37.2 |

| | | | | | | | | |
|---|---|---|---|---|---|---|---|---|
| No results | 458.267 | 459.2743 | 20.479 | 56503889 | 25 | 1 | 0 | 37.4 |
| No results | 442.2118 | 443.2191 | 19.553 | 50893879 | 55 | 1 | 0 | 41.1 |
| No results | 598.429 | 599.4368 | 21.592 | 1.35E+08 | 1 | 1 | 0 | 91.4 |
| No results | 598.4291 | 599.4369 | 21.896 | 1.17E+08 | 1 | 1 | 0 | 91.4 |
| No results | 453.2941 | 454.3015 | 18.649 | 2.1E+08 | 2 | 2 | 0 | 31.6 |
| No results | 656.4705 | 657.4783 | 22.447 | 1.01E+08 | 0 | 1 | 0 | 90 |
| No results | 576.3486 | 577.3559 | 21.089 | 88411352 | 5 | 1 | 0 | 45.3 |
| No match | 558.253 | 559.2603 | 22.689 | 29304974 | 5 | 2 | 0 | 91.9 |
| No results | 326.2068 | 327.2141 | 24.179 | 52192769 | 0 | 4 | 0 | 67 |
| No results | 486.2984 | 487.3057 | 22.066 | 34496080 | 83 | 1 | 0 | 51.2 |
| No results | 443.0475 | 444.0548 | 18.438 | 44816095 | 6 | 1 | 0 | 47.5 |
| No results | 645.3262 | 646.3335 | 18.581 | 80907356 | 5 | 1 | 0 | 63.7 |
| No results | 390.2044 | 391.2116 | 19.188 | 1.1E+08 | 95 | 1 | 0 | 61.9 |
| No results | 615.4556 | 616.4626 | 21.551 | 1.68E+08 | 0 | 1 | 0 | 91.4 |
| No results | 673.497 | 674.5039 | 22.436 | 1.54E+08 | 0 | 1 | 0 | 90.8 |
| No results | 500.2501 | 501.2574 | 15.37 | 45699202 | 8 | 2 | 0 | 67.3 |
| No results | 514.2273 | 515.2345 | 23.144 | 25490295 | 9 | 4 | 0 | 43.5 |
| No results | 497.2782 | 498.2855 | 17.16 | 45958867 | 38 | 1 | 0 | 60.9 |
| No results | 808.5155 | 809.5228 | 23.966 | 77245765 | 0 | 1 | 0 | 33.4 |
| No results | 656.4706 | 657.4782 | 22.726 | 71959446 | 0 | 1 | 0 | 90 |
| No results | 448.1403 | 449.1476 | 19.892 | 32167045 | 96 | 3 | 0 | 56.5 |
| No results | 673.4971 | 674.504 | 22.749 | 92356540 | 0 | 1 | 0 | 90 |
| No results | 443.0475 | 444.0548 | 18.785 | 51780942 | 6 | 1 | 0 | 43 |
| No results | 407.1182 | 408.1255 | 10.832 | 43927473 | 7 | 1 | 0 | 86 |
| No results | 316.1674 | 317.1747 | 22.172 | 89342952 | 89 | 2 | 0 | 31.8 |
| No results | 481.1687 | 482.176 | 21.124 | 29889094 | 65 | 1 | 0 | 81.3 |
| No match | 302.1884 | 303.1957 | 23.41 | 36096549 | 123 | 7 | 0 | 49.6 |
| No match | 504.3179 | 505.3252 | 14.313 | 36638498 | 5 | 3 | 0 | 92.5 |
| No results | 360.23 | 361.2372 | 21.231 | 30761432 | 102 | 2 | 0 | 43 |
| No results | 354.068 | 178.0413 | 14.651 | 14790065 | 85 | 7 | 0 | 55.8 |
| No results | 400.2073 | 401.2146 | 17.038 | 1.22E+08 | 36 | 3 | 0 | 66.6 |
| No match | 460.2463 | 461.2536 | 19.128 | 35070010 | 152 | 3 | 0 | 48.7 |
| No results | 1704.986 | 569.336 | 15.956 | 45697278 | 0 | 2 | 0 | 34 |
| No match | 704.3894 | 353.202 | 19.854 | 30648042 | 7 | 1 | 0 | 94.7 |
| No results | 557.4143 | 558.4211 | 20.882 | 87133945 | 2 | 1 | 0 | 94.5 |
| No results | 402.2229 | 403.2302 | 16.665 | 1.81E+08 | 4 | 1 | 0 | 61.5 |
| No results | 402.1314 | 403.1387 | 21.017 | 31138776 | 99 | 8 | 0 | 93.6 |
| No results | 576.3486 | 577.3559 | 20.745 | 60447949 | 5 | 1 | 0 | 46.2 |
| No results | 823.4984 | 824.5057 | 24.165 | 62268170 | 0 | 1 | 0 | 30.3 |
| No results | 299.2146 | 317.2476 | 21.561 | 18480598 | 0 | 5 | 0 | 57.4 |
| No results | 442.2258 | 443.2331 | 18.494 | 28139269 | 9 | 2 | 0 | 44.9 |
| No results | 348.1937 | 349.201 | 20.397 | 69289837 | 132 | 3 | 0 | 45.6 |
| No results | 499.3725 | 500.3798 | 19.655 | 70054024 | 0 | 1 | 0 | 86.6 |
| No results | 482.3458 | 483.353 | 19.647 | 98127043 | 0 | 1 | 0 | 97.3 |
| No results | 466.2721 | 467.2794 | 22.802 | 26105136 | 53 | 1 | 0 | 51.7 |
| No match | 344.2348 | 345.2423 | 23.509 | 17218891 | 86 | 6 | 0 | 63.8 |
| No results | 444.2276 | 445.2349 | 15.256 | 32638380 | 8 | 2 | 0 | 43.4 |
| No results | 478.3144 | 479.3217 | 22.53 | 23078234 | 1 | 1 | 0 | 34.5 |
| No results | 442.2118 | 443.2191 | 16.738 | 31260660 | 125 | 1 | 0 | 43.6 |
| No results | 481.1687 | 482.176 | 20.057 | 24458256 | 65 | 1 | 0 | 90.9 |
| No results | 408.2147 | 391.2114 | 16.528 | 25880868 | 41 | 1 | 0 | 55.5 |

| | | | | | | | | |
|---|---|---|---|---|---|---|---|---|
| No results | 1308.751 | 655.3826 | 15.101 | 43263875 | 0 | 2 | 0 | 30.5 |
| No results | 458.2307 | 459.238 | 19.635 | 25639593 | 181 | 2 | 0 | 55.1 |
| No results | 460.2827 | 461.2899 | 17.924 | 25362113 | 75 | 2 | 0 | 73.2 |
| No results | 808.5156 | 809.5229 | 24.105 | 46288682 | 0 | 1 | 0 | 32.2 |
| No results | 217.0633 | 435.1333 | 21.02 | 22298729 | 0 | 2 | 0 | 31.7 |
| No results | 673.4972 | 674.5042 | 22.22 | 39244239 | 0 | 1 | 0 | 86.8 |
| No results | 620.4108 | 621.418 | 21.565 | 64675013 | 3 | 1 | 0 | 91.8 |
| No results | 486.2986 | 487.3058 | 19.598 | 20312928 | 83 | 1 | 0 | 41.6 |
| No results | 332.126 | 333.1333 | 17.865 | 23872163 | 74 | 3 | 0 | 53.5 |
| No results | 460.2651 | 461.2723 | 18.733 | 42499058 | 15 | 1 | 0 | 56.9 |
| No results | 446.1942 | 447.2015 | 15.841 | 18733831 | 229 | 2 | 0 | 34 |
| No results | 472.319 | 473.3263 | 21.934 | 28406240 | 47 | 1 | 0 | 52.5 |
| No results | 334.1781 | 335.1854 | 19.558 | 31739508 | 45 | 2 | 0 | 46.6 |
| No match | 460.2464 | 461.2537 | 19.698 | 40491654 | 151 | 3 | 0 | 54.7 |
| No results | 426.2407 | 427.2479 | 19.437 | 23749897 | 35 | 3 | 0 | 39.2 |
| No results | 441.3302 | 442.3375 | 17.606 | 24066274 | 0 | 1 | 0 | 56.2 |
| No results | 368.2537 | 369.261 | 25.477 | 41620809 | 2 | 2 | 0 | 55.7 |
| No results | 438.1209 | 439.1281 | 15.75 | 17680351 | 56 | 1 | 0 | 67.5 |
| No results | 411.05 | 412.0573 | 20.678 | 15769126 | 19 | 1 | 0 | 32.5 |
| No results | 499.2939 | 500.3012 | 18.402 | 22388670 | 22 | 2 | 0 | 31.5 |
| No results | 536.3561 | 554.3898 | 23.319 | 30342568 | 0 | 1 | 0 | 41 |
| No match | 374.2093 | 375.2166 | 16.59 | 17027468 | 2 | 2 | 0 | 48.1 |
| No results | 236.1414 | 237.1487 | 17.672 | 36479405 | 57 | 3 | 0 | 52.5 |
| No results | 432.142 | 433.1493 | 21.578 | 17925267 | 109 | 6 | 0 | 63.7 |
| No results | 482.3457 | 483.353 | 19.015 | 34481422 | 1 | 1 | 0 | 89.1 |
| No results | 472.3192 | 473.3264 | 20.294 | 16660898 | 47 | 1 | 0 | 59.1 |
| No results | 490.257 | 508.2909 | 24.529 | 9299936 | 101 | 2 | 0 | 37.9 |
| No results | 478.3146 | 496.3485 | 23.338 | 26069390 | 0 | 1 | 0 | 33.8 |
| No match | 348.2302 | 349.2374 | 23.074 | 18229908 | 79 | 5 | 0 | 61 |
| No results | 228.134 | 229.1413 | 29.312 | 31679107 | 0 | 1 | 0 | 33.6 |
| No results | 518.3071 | 519.3144 | 19.576 | 1.06E+08 | 10 | 2 | 0 | 44.8 |
| No results | 460.2652 | 461.2725 | 18.627 | 72288721 | 15 | 1 | 0 | 53.8 |
| No results | 298.1783 | 281.175 | 19.434 | 24611311 | 62 | 2 | 0 | 36.4 |
| No results | 562.3692 | 563.3764 | 20.529 | 25892987 | 1 | 1 | 0 | 91.5 |
| No results | 247.1462 | 265.1801 | 22.937 | 18985961 | 0 | 1 | 0 | 33 |
| No results | 391.2722 | 392.2795 | 21.103 | 17864398 | 19 | 1 | 0 | 33.1 |
| No results | 269.0916 | 270.0989 | 13.921 | 32319912 | 50 | 3 | 0 | 49.7 |
| No results | 419.2422 | 420.2495 | 14.892 | 12101442 | 29 | 3 | 0 | 32.7 |
| No results | 396.2511 | 379.2478 | 23.695 | 10605703 | 19 | 1 | 0 | 30.2 |
| No results | 398.2279 | 399.2352 | 21.986 | 79720290 | 1 | 5 | 0 | 79.9 |
| No results | 715.4499 | 748.4831 | 15.815 | 20619508 | 4 | 1 | 0 | 81 |
| No results | 528.243 | 529.2503 | 21.745 | 10231830 | 9 | 1 | 0 | 31.1 |
| No results | 463.1581 | 464.1654 | 21.992 | 10454311 | 63 | 2 | 0 | 79 |
| No results | 504.3091 | 505.3164 | 20.833 | 16174539 | 69 | 1 | 0 | 35.6 |
| No results | 458.2497 | 459.257 | 18.586 | 48989635 | 10 | 1 | 0 | 48.2 |
| No match | 454.2835 | 455.2908 | 15.248 | 22471265 | 36 | 2 | 0 | 61.7 |
| No results | 446.2493 | 447.2566 | 17.246 | 29017151 | 18 | 1 | 0 | 60.3 |
| No results | 264.1365 | 265.1438 | 19.431 | 38085233 | 231 | 15 | 0 | 45.6 |
| No results | 536.3561 | 554.3898 | 23.954 | 24426284 | 0 | 1 | 0 | 39.6 |
| No results | 408.2147 | 391.2114 | 16.869 | 15703787 | 41 | 1 | 0 | 34.6 |
| No results | 456.2513 | 457.2586 | 20.989 | 13803296 | 120 | 1 | 0 | 60.3 |

| | | | | | | | | |
|---|---|---|---|---|---|---|---|---|
| No results | 486.1957 | 487.203 | 21.829 | 9747039 | 42 | 1 | 0 | 40.4 |
| No results | 551.3944 | 552.4017 | 15.147 | 19650290 | 4 | 2 | 0 | 81.5 |
| No results | 495.341 | 496.3484 | 23.256 | 35193420 | 0 | 1 | 0 | 33.8 |
| No match | 250.1571 | 251.1644 | 20.199 | 38256757 | 228 | 11 | 0 | 64 |
| No match | 426.2407 | 427.248 | 20.535 | 20827587 | 35 | 3 | 0 | 35.8 |
| No results | 249.0792 | 250.0865 | 21.747 | 37037138 | 68 | 1 | 0 | 31.2 |
| No results | 282.1621 | 283.1694 | 22.087 | 17478315 | 55 | 1 | 0 | 49.5 |
| No results | 594.3976 | 612.4312 | 23.924 | 19263127 | 0 | 1 | 0 | 43.6 |
| No results | 544.2403 | 545.2475 | 14.126 | 10024215 | 28 | 3 | 0 | 70.3 |
| No results | 495.3411 | 496.3484 | 22.514 | 45788309 | 0 | 1 | 0 | 34.5 |
| No results | 292.0948 | 293.102 | 17.159 | 25507871 | 51 | 1 | 0 | 33.3 |
| No results | 553.3829 | 554.3899 | 23.301 | 30342568 | 0 | 1 | 0 | 41 |
| No results | 298.157 | 299.1643 | 19.102 | 18490679 | 119 | 3 | 0 | 60.9 |
| No results | 546.2174 | 547.2247 | 22.657 | 8953596 | 1 | 3 | 0 | 39.7 |
| No results | 502.2935 | 503.3008 | 20.244 | 24904955 | 49 | 1 | 0 | 35.1 |
| No results | 302.1518 | 303.1591 | 20.364 | 37089032 | 93 | 5 | 0 | 63.6 |
| No results | 514.2272 | 515.2345 | 23.662 | 6356271 | 24 | 5 | 0 | 71.5 |
| No results | 322.2549 | 323.2621 | 24.337 | 8086663 | 0 | 5 | 0 | 38.5 |
| No results | 356.1947 | 357.202 | 18.301 | 10173170 | 3 | 2 | 0 | 35.1 |
| No results | 277.1469 | 278.1542 | 12.749 | 23624848 | 20 | 1 | 0 | 31 |
| No results | 504.1797 | 505.187 | 11.962 | 16318076 | 4 | 3 | 0 | 64.6 |
| No results | 458.2391 | 459.2464 | 15.723 | 15294328 | 17 | 3 | 0 | 62 |
| No results | 474.262 | 475.2693 | 19.617 | 15109990 | 123 | 1 | 0 | 39.8 |
| No results | 476.2412 | 477.2485 | 17.874 | 13043777 | 171 | 2 | 0 | 33.2 |
| No match | 314.1519 | 315.1592 | 23.022 | 12473569 | 124 | 6 | 0 | 38.4 |
| No match | 277.2044 | 278.2117 | 10.539 | 16667558 | 104 | 1 | 0 | 31.7 |
| No match | 331.2149 | 332.2221 | 16.916 | 11890703 | 77 | 2 | 0 | 58.7 |
| No results | 308.2029 | 309.2102 | 17.818 | 12337599 | 10 | 4 | 0 | 42.7 |
| No results | 456.2515 | 457.2587 | 20.234 | 15279224 | 120 | 1 | 0 | 57.3 |
| No results | 530.2609 | 531.2682 | 13.815 | 11803560 | 7 | 2 | 0 | 80.4 |
| No results | 346.0227 | 347.03 | 23.357 | 7309310 | 6 | 2 | 0 | 55.1 |
| No results | 400.1861 | 401.1933 | 20.429 | 8559950 | 40 | 2 | 0 | 32.5 |
| No match | 382.1892 | 366.1699 | 22.783 | 8901907 | 268 | 1 | 0 | 35.8 |
| No results | 332.1987 | 333.206 | 22.177 | 96188247 | 229 | 15 | 0 | 48.5 |
| No results | 418.2355 | 419.2428 | 22.093 | 12283383 | 80 | 1 | 0 | 45.3 |
| No results | 342.1103 | 343.1175 | 21.168 | 11036678 | 144 | 6 | 0 | 78.9 |
| No match | 362.1729 | 363.1802 | 19.01 | 18882411 | 1 | 8 | 0 | 54.6 |
| No results | 332.1988 | 333.2061 | 17.978 | 18595317 | 229 | 14 | 0 | 54.2 |
| No results | 350.2497 | 351.257 | 21.448 | 45678367 | 1 | 3 | 0 | 45 |
| No match | 330.1831 | 331.1903 | 22.484 | 68734030 | 1 | 2 | 0 | 35.8 |
| No results | 444.2236 | 445.2309 | 14.245 | 9460598 | 24 | 1 | 0 | 30.8 |
| No results | 349.126 | 350.1333 | 24.809 | 5022583 | 41 | 2 | 0 | 42.1 |
| No results | 334.1781 | 335.1854 | 15.027 | 17798693 | 45 | 2 | 0 | 52.1 |
| No results | 442.2543 | 443.2615 | 21.506 | 24471261 | 14 | 1 | 0 | 76.1 |
| No match | 364.1886 | 365.1959 | 21.487 | 12879531 | 100 | 1 | 0 | 52.2 |
| No results | 416.2563 | 417.2636 | 25.643 | 4425162 | 56 | 2 | 0 | 30.7 |
| No results | 429.0244 | 430.0316 | 9.98 | 11541497 | 9 | 1 | 0 | 32 |
| No results | 280.1677 | 281.175 | 22.418 | 12877981 | 95 | 4 | 0 | 47.9 |
| No results | 504.3279 | 505.3352 | 19.282 | 38090534 | 1 | 1 | 0 | 87.9 |
| No results | 360.2301 | 361.2374 | 19.501 | 13701765 | 100 | 4 | 0 | 61.1 |
| No results | 468.2213 | 469.2286 | 24.53 | 5149102 | 2 | 2 | 0 | 58.6 |

| | | | | | | | | |
|---|---|---|---|---|---|---|---|---|
| No results | 390.2043 | 391.2116 | 16.871 | 15703787 | 1 | 1 | 0 | 35 |
| No results | 501.2944 | 485.275 | 21.77 | 6658611 | 14 | 2 | 0 | 44 |
| No results | 368.2539 | 369.2612 | 25.757 | 12477974 | 2 | 1 | 0 | 39.2 |
| No results | 415.2109 | 416.2182 | 10.843 | 20602148 | 62 | 1 | 0 | 39.5 |
| No results | 392.1445 | 393.1518 | 9.952 | 10254806 | 38 | 5 | 0 | 55.9 |
| No results | 504.2916 | 505.2989 | 18.724 | 15220873 | 10 | 1 | 0 | 31.9 |
| No results | 421.1474 | 422.1546 | 23.062 | 6993395 | 60 | 2 | 0 | 48 |
| No results | 864.5413 | 865.5486 | 24.461 | 19204724 | 0 | 1 | 0 | 57.5 |
| No results | 473.2164 | 474.2237 | 15.748 | 16212445 | 52 | 1 | 0 | 37.2 |
| No results | 720.3127 | 361.1636 | 23.071 | 6387276 | 12 | 1 | 0 | 30.1 |
| No results | 322.2548 | 323.2621 | 24.142 | 16134333 | 0 | 2 | 0 | 32.5 |
| No results | 247.1462 | 289.18 | 22.21 | 45082465 | 0 | 4 | 0 | 59.1 |
| No results | 351.2411 | 352.2484 | 13.619 | 20866487 | 18 | 3 | 0 | 44.7 |
| No results | 418.2719 | 419.2792 | 22.136 | 10830675 | 50 | 1 | 0 | 30.7 |
| No results | 678.5043 | 340.2595 | 14.175 | 6755314 | 2 | 4 | 0 | 36.7 |
| No results | 348.1939 | 349.2011 | 20.79 | 25326848 | 132 | 3 | 0 | 51.1 |
| No results | 384.1216 | 385.1289 | 14.344 | 7197269 | 7 | 7 | 0 | 74.3 |
| No match | 369.0759 | 370.0832 | 11.682 | 15076254 | 43 | 6 | 0 | 51.9 |
| No match | 311.1094 | 312.1167 | 10.661 | 14788736 | 205 | 7 | 0 | 70.6 |
| No match | 342.1467 | 343.1539 | 24.284 | 6185946 | 165 | 1 | 0 | 91.7 |
| No results | 502.2293 | 503.2366 | 14.594 | 6783412 | 31 | 2 | 0 | 80.1 |
| No match | 374.2092 | 375.2164 | 21.602 | 14756200 | 111 | 2 | 0 | 41.9 |
| No results | 638.4606 | 656.4944 | 24.621 | 10761239 | 0 | 1 | 0 | 44.9 |
| No results | 448.1897 | 225.1022 | 13.187 | 13899267 | 95 | 3 | 0 | 78.4 |
| No match | 460.2463 | 461.2536 | 17.381 | 9204006 | 152 | 1 | 0 | 40.5 |
| No results | 786.4343 | 787.4416 | 23.602 | 13850900 | 4 | 1 | 0 | 44.3 |
| No results | 594.3975 | 612.4312 | 24.457 | 12050689 | 0 | 1 | 0 | 46.6 |
| No results | 331.0798 | 332.0871 | 11.986 | 11836823 | 65 | 3 | 0 | 65 |
| No results | 611.4241 | 612.4314 | 23.912 | 19263127 | 0 | 1 | 0 | 43.6 |
| No results | 430.1994 | 431.2067 | 20.785 | 10339141 | 239 | 2 | 0 | 45.4 |
| No results | 276.1364 | 277.1437 | 22.123 | 17344283 | 75 | 2 | 0 | 51.8 |
| No results | 271.1211 | 272.1284 | 14.68 | 9997200 | 242 | 2 | 0 | 63.6 |
| No results | 407.2884 | 408.2957 | 13.565 | 15575347 | 2 | 1 | 0 | 80.3 |
| No results | 379.2933 | 380.3006 | 21.626 | 39855900 | 1 | 1 | 0 | 30.2 |
| No results | 502.2933 | 503.3005 | 21.513 | 27485258 | 26 | 1 | 0 | 35.3 |
| No results | 350.1729 | 351.1802 | 19.157 | 12610640 | 55 | 1 | 0 | 47.1 |
| No results | 268.1101 | 269.1174 | 20.46 | 7714089 | 151 | 4 | 0 | 56.6 |
| No results | 378.2409 | 379.2481 | 23.727 | 10605703 | 76 | 3 | 0 | 44.6 |
| No match | 440.2677 | 441.275 | 14.302 | 14718977 | 50 | 1 | 0 | 90 |
| No results | 308.1413 | 309.1486 | 20.006 | 7237787 | 105 | 1 | 0 | 45.3 |
| No results | 487.2728 | 488.2801 | 15.048 | 13049680 | 35 | 1 | 0 | 51.3 |
| No results | 337.1426 | 338.1499 | 9.4 | 8612674 | 389 | 4 | 0 | 89.8 |
| No results | 334.1782 | 335.1855 | 16.873 | 28818655 | 45 | 2 | 0 | 60.4 |
| No results | 404.1294 | 405.1367 | 20.561 | 10651834 | 3 | 1 | 0 | 30.9 |
| No results | 669.4664 | 670.4736 | 24.408 | 10242605 | 0 | 1 | 0 | 52.9 |
| No results | 503.31 | 487.2906 | 22.49 | 6424007 | 2 | 1 | 0 | 65 |
| No results | 346.1779 | 347.1852 | 20.987 | 19360220 | 160 | 1 | 0 | 32.5 |
| No results | 457.2834 | 458.2907 | 12.841 | 9798784 | 30 | 1 | 0 | 46.3 |
| No results | 380.0865 | 381.0937 | 14.149 | 6287800 | 1 | 5 | 0 | 41.2 |
| No results | 366.2405 | 367.2478 | 22.122 | 9334958 | 31 | 2 | 0 | 47.1 |
| No results | 472.2463 | 473.2536 | 18.679 | 9334522 | 149 | 3 | 0 | 40.9 |

| | | | | | | | | |
|---|---|---|---|---|---|---|---|---|
| No results | 486.2344 | 487.2417 | 13.795 | 8272277 | 17 | 1 | 0 | 43.4 |
| No match | 435.2411 | 436.2484 | 15.026 | 13113715 | 59 | 1 | 0 | 67.8 |
| No results | 287.1776 | 305.2112 | 20.519 | 27950122 | 0 | 1 | 0 | 31.1 |
| No results | 444.2337 | 445.241 | 17.615 | 19182011 | 42 | 2 | 0 | 37.9 |
| No match | 304.1676 | 305.1748 | 20.309 | 31770987 | 77 | 1 | 0 | 33.1 |
| No results | 346.2548 | 347.2621 | 22.734 | 6263697 | 0 | 2 | 0 | 38.8 |
| No results | 460.2826 | 461.2899 | 22.17 | 14898324 | 75 | 2 | 0 | 37.6 |
| No results | 439.2432 | 440.2505 | 17.003 | 6800398 | 2 | 1 | 0 | 30.2 |
| No results | 472.319 | 473.3263 | 21.406 | 10471229 | 47 | 1 | 0 | 45.9 |
| No results | 318.1105 | 319.1177 | 20.733 | 7405362 | 80 | 2 | 0 | 35.9 |
| No results | 348.221 | 349.2282 | 18.215 | 6492044 | 179 | 2 | 0 | 35 |
| No results | 518.2973 | 519.3045 | 14.278 | 7150218 | 6 | 3 | 0 | 56.7 |
| No results | 387.1795 | 388.1868 | 12.183 | 13761185 | 88 | 1 | 0 | 34.6 |
| No results | 312.1725 | 313.1798 | 20.915 | 34470028 | 1 | 1 | 0 | 49.9 |
| No results | 419.2462 | 420.2535 | 13.223 | 16243330 | 41 | 4 | 0 | 67.4 |
| No results | 280.1677 | 281.175 | 19.433 | 24611311 | 95 | 6 | 0 | 50.1 |
| No results | 292.077 | 293.0843 | 20.912 | 7025741 | 27 | 2 | 0 | 37.9 |
| No results | 553.3824 | 554.3897 | 23.956 | 24426284 | 0 | 1 | 0 | 39.6 |
| No results | 554.3063 | 555.3136 | 19.437 | 5986504 | 4 | 1 | 0 | 42.4 |
| No results | 502.3299 | 503.3371 | 20.351 | 11434793 | 78 | 1 | 0 | 34.1 |
| No results | 394.1993 | 395.2066 | 18.818 | 8205157 | 1 | 1 | 0 | 38.9 |
| No results | 419.1315 | 420.1388 | 22.918 | 5556866 | 184 | 4 | 0 | 73.9 |
| No results | 350.2496 | 351.2569 | 21.812 | 18513917 | 1 | 1 | 0 | 32.5 |
| No results | 280.1677 | 281.1749 | 24.314 | 13907365 | 95 | 1 | 0 | 34.1 |
| No match | 322.1781 | 323.1854 | 21.34 | 12836250 | 57 | 3 | 0 | 39.8 |
| No match | 314.1519 | 315.1592 | 15.053 | 6839376 | 2 | 5 | 0 | 34.3 |
| No results | 362.0824 | 363.0896 | 17.958 | 6342178 | 76 | 8 | 0 | 42.4 |
| No results | 458.2306 | 459.2378 | 19.535 | 29870088 | 181 | 2 | 0 | 49.2 |
| No match | 378.2042 | 379.2114 | 22.758 | 32364573 | 47 | 1 | 0 | 31.2 |
| No match | 733.4609 | 734.4682 | 15.665 | 7286088 | 17 | 2 | 0 | 31.2 |
| No match | 314.2245 | 347.258 | 23.562 | 8148400 | 130 | 8 | 0 | 52.7 |
| No results | 269.0238 | 270.0311 | 10.442 | 8262346 | 3 | 2 | 0 | 46.6 |
| No match | 278.1057 | 279.113 | 18.714 | 8367260 | 306 | 4 | 0 | 59.1 |
| No results | 304.0947 | 305.102 | 16.277 | 7932846 | 71 | 4 | 0 | 79.2 |
| No match | 372.1508 | 373.1581 | 12.655 | 12127833 | 9 | 1 | 0 | 58.8 |
| No results | 528.2067 | 529.214 | 23.332 | 3696284 | 19 | 2 | 0 | 53.9 |
| No results | 516.2813 | 517.2886 | 13.285 | 9098616 | 3 | 3 | 0 | 67.3 |
| No results | 306.1469 | 307.1542 | 18.908 | 23782180 | 93 | 5 | 0 | 50.4 |
| No results | 451.1269 | 452.1342 | 17.915 | 4337446 | 13 | 1 | 0 | 59.9 |
| No results | 318.1103 | 319.1175 | 18.089 | 8078708 | 81 | 2 | 0 | 44.2 |
| No results | 488.2275 | 489.2348 | 10.789 | 8416484 | 42 | 5 | 0 | 78.8 |
| No results | 602.263 | 302.1388 | 13.009 | 8669409 | 12 | 3 | 0 | 39.4 |
| No match | 364.1886 | 365.1959 | 22.048 | 11040311 | 100 | 1 | 0 | 42.7 |
| No match | 1018.603 | 510.3088 | 24.816 | 9661747 | 0 | 2 | 0 | 57.4 |
| No match | 424.1802 | 425.1874 | 11.766 | 8274465 | 68 | 3 | 0 | 96.3 |
| No results | 388.1158 | 389.1231 | 19.335 | 7190583 | 100 | 5 | 0 | 58.7 |
| No results | 674.4207 | 675.428 | 24.889 | 5716624 | 4 | 1 | 0 | 80.3 |
| No results | 304.2039 | 305.2112 | 20.558 | 27950122 | 97 | 1 | 0 | 31.1 |
| No results | 482.267 | 483.2743 | 21.891 | 9235004 | 22 | 1 | 0 | 44 |
| No results | 421.2619 | 422.2691 | 13.959 | 12528492 | 38 | 1 | 0 | 32.9 |
| No results | 308.1413 | 309.1486 | 21.621 | 9543408 | 91 | 3 | 0 | 53.2 |

| | | | | | | | | |
|---|---|---|---|---|---|---|---|---|
| No results | 349.2252 | 350.2325 | 19.88 | 9208859 | 46 | 6 | 0 | 47.8 |
| No match | 309.1842 | 310.1914 | 24.19 | 2963174 | 224 | 3 | 0 | 56.7 |
| No results | 844.477 | 423.2458 | 23.622 | 9155856 | 2 | 1 | 0 | 46.9 |
| No match | 265.168 | 266.1753 | 9.464 | 7729137 | 126 | 2 | 0 | 51 |
| No match | 317.1993 | 318.2066 | 16.22 | 5928318 | 113 | 4 | 0 | 53.7 |
| No results | 286.157 | 287.1643 | 19.311 | 13977412 | 73 | 5 | 0 | 54.6 |
| No results | 291.241 | 292.2483 | 21.359 | 56622348 | 0 | 1 | 0 | 30.2 |
| No results | 456.2516 | 457.2589 | 19.626 | 12667202 | 75 | 1 | 0 | 39.8 |
| No results | 421.2677 | 422.275 | 14.016 | 7327083 | 4 | 2 | 0 | 76 |
| No results | 344.1622 | 345.1695 | 20.773 | 10666974 | 161 | 1 | 0 | 41.1 |
| No results | 428.2927 | 429.3 | 22.846 | 8557638 | 63 | 1 | 0 | 48 |
| No results | 630.2942 | 316.1544 | 13.675 | 6778392 | 10 | 3 | 0 | 59.8 |
| No results | 261.0688 | 262.0761 | 15.708 | 8309194 | 18 | 1 | 0 | 46 |
| No results | 447.2833 | 448.2906 | 22.758 | 4198552 | 5 | 1 | 0 | 47.8 |
| No results | 506.1463 | 507.1536 | 19.522 | 4290961 | 31 | 3 | 0 | 46.7 |
| No match | 326.1299 | 327.1372 | 11.36 | 14428850 | 47 | 4 | 0 | 52.7 |
| No match | 360.1322 | 361.1395 | 14.525 | 5536729 | 96 | 4 | 0 | 58.6 |
| No match | 314.2246 | 315.2318 | 19.239 | 6546881 | 130 | 8 | 0 | 49.8 |
| No match | 363.0873 | 364.0946 | 20.117 | 4806319 | 24 | 7 | 0 | 37.2 |
| No match | 330.2193 | 331.2266 | 21.407 | 21797060 | 170 | 2 | 0 | 50.5 |
| No results | 425.2064 | 426.2137 | 15.73 | 4788628 | 112 | 5 | 0 | 40.8 |
| No results | 502.2934 | 503.3006 | 18.553 | 12356177 | 51 | 1 | 0 | 34.7 |
| No results | 336.1686 | 337.1759 | 13.697 | 8596593 | 61 | 1 | 0 | 35.5 |
| No results | 391.1366 | 392.1439 | 23.662 | 3731277 | 41 | 2 | 0 | 44.9 |
| No results | 472.2465 | 473.2538 | 21.642 | 7535555 | 141 | 2 | 0 | 38.1 |
| No results | 238.0995 | 239.1067 | 16.684 | 4821201 | 79 | 2 | 0 | 46 |
| No results | 430.2718 | 431.2791 | 20.343 | 23756700 | 44 | 1 | 0 | 53.9 |
| No results | 415.2111 | 416.2184 | 10.595 | 10777915 | 62 | 1 | 0 | 42.5 |
| No results | 388.1884 | 389.1957 | 19.034 | 8764013 | 77 | 6 | 0 | 38.2 |
| No results | 391.1366 | 392.1439 | 23.425 | 3012419 | 187 | 2 | 0 | 38.8 |
| No results | 383.2098 | 384.2171 | 11.219 | 7489620 | 3 | 2 | 0 | 74.1 |
| No results | 232.1465 | 233.1538 | 21.01 | 52623901 | 125 | 13 | 0 | 77.7 |
| No results | 232.1466 | 233.1539 | 18.811 | 34491435 | 125 | 10 | 0 | 67 |
| No results | 232.1466 | 233.1539 | 18.941 | 13022781 | 125 | 10 | 0 | 67 |
| No results | 262.1571 | 263.1644 | 17.63 | 20602587 | 80 | 2 | 0 | 60.5 |
| No results | 262.1573 | 263.1645 | 20.955 | 37976777 | 80 | 2 | 0 | 59.3 |
| No results | 262.1572 | 263.1644 | 19.609 | 56205818 | 80 | 2 | 0 | 52.6 |
| No results | 246.1622 | 247.1695 | 18.753 | 52775689 | 50 | 5 | 0 | 71.5 |
| No results | 246.1622 | 247.1695 | 17.67 | 13463733 | 51 | 5 | 0 | 47.7 |
| No results | 348.1937 | 349.201 | 24.419 | 44648902 | 132 | 7 | 0 | 58.9 |
| No results | 348.1938 | 349.201 | 18.147 | 95449173 | 132 | 7 | 0 | 69.2 |
| No results | 348.1938 | 349.2011 | 18.385 | 36741550 | 132 | 4 | 0 | 60.5 |
| No results | 348.1938 | 349.2011 | 25.072 | 5049609 | 132 | 2 | 0 | 30.5 |
| No results | 349.2254 | 350.2327 | 15.111 | 13683237 | 46 | 1 | 0 | 53.5 |
| No results | 472.3189 | 455.3156 | 23.536 | 1.6E+09 | 32 | 1 | 0 | 38.5 |
| No results | 472.3189 | 473.3262 | 23.216 | 2.09E+08 | 32 | 1 | 0 | 41 |
| No results | 271.215 | 272.2222 | 20.591 | 25201580 | 8 | 2 | 0 | 34.2 |
| No results | 238.1208 | 239.1281 | 20.697 | 35899564 | 50 | 1 | 0 | 33.9 |
| No results | 342.2154 | 343.2227 | 20.007 | 17767379 | 10 | 1 | 0 | 37.4 |
| Partial mat | 248.1415 | 249.1487 | 19.776 | 94147694 | 245 | 15 | 0 | 48.6 |
| Partial mat | 248.1415 | 249.1488 | 17.742 | 11191547 | 245 | 17 | 0 | 66.7 |

| | | | | | | | | |
|---|---|---|---|---|---|---|---|---|
| No results | 293.1628 | 294.17 | 21.155 | 19210436 | 95 | 1 | 0 | 31.2 |
| No results | 472.319 | 455.3157 | 22.591 | 4.28E+08 | 32 | 1 | 0 | 40 |
| No results | 280.1676 | 281.1749 | 20.951 | 76778979 | 95 | 6 | 0 | 50.2 |
| No results | 280.1677 | 281.175 | 19.182 | 22674226 | 95 | 6 | 0 | 56.1 |
| Full match | 307.1785 | 308.1857 | 16.575 | 10012352 | 85 | 2 | 0 | 38.9 |
| No results | 322.1568 | 323.1641 | 23.262 | 10717806 | 60 | 2 | 0 | 39.8 |
| No results | 294.1831 | 295.1904 | 21.782 | 26870066 | 59 | 1 | 0 | 49.5 |
| No results | 448.1433 | 449.1506 | 17.784 | 21977019 | 8 | 4 | 0 | 70.4 |
| No results | 486.2985 | 487.3058 | 19.187 | 16677219 | 83 | 1 | 0 | 33.8 |
| No results | 428.2172 | 429.2245 | 21.04 | 4801554 | 1 | 1 | 0 | 32.7 |
| No results | 368.235 | 369.2423 | 24.328 | 8165325 | 34 | 2 | 0 | 56.3 |
| No results | 234.0894 | 235.0967 | 23.276 | 63637049 | 89 | 9 | 0 | 87.7 |
| No results | 435.1628 | 436.1701 | 22.317 | 4226199 | 2 | 3 | 0 | 44.1 |
| No results | 283.1574 | 284.1647 | 13.661 | 26680218 | 294 | 1 | 0 | 32.1 |
| No results | 264.1364 | 265.1436 | 19.326 | 38935467 | 231 | 17 | 0 | 57.2 |
| No results | 264.1364 | 265.1437 | 18.776 | 16191213 | 231 | 16 | 0 | 62.9 |
| Partial mat | 248.1414 | 249.1487 | 21.53 | 21267513 | 245 | 7 | 0 | 33.9 |
| No results | 457.2832 | 458.2905 | 14.963 | 20410615 | 10 | 1 | 0 | 38.2 |
| No results | 272.1415 | 273.1487 | 21.847 | 23619368 | 79 | 5 | 0 | 49.8 |
| No results | 324.1145 | 325.1217 | 14.488 | 22442529 | 97 | 3 | 0 | 54.3 |
| No results | 324.1144 | 325.1217 | 14.694 | 19230658 | 97 | 3 | 0 | 50.6 |
| No results | 306.1468 | 307.1541 | 19.865 | 20648512 | 93 | 1 | 0 | 32.1 |
| No results | 269.1993 | 270.2066 | 21.106 | 24116413 | 23 | 4 | 0 | 42.4 |
| No results | 269.1993 | 270.2066 | 22.351 | 11481321 | 23 | 4 | 0 | 44.3 |
| No results | 306.1468 | 307.1541 | 18.39 | 23544341 | 93 | 6 | 0 | 53.5 |
| No results | 307.0436 | 308.0508 | 12.912 | 35226028 | 36 | 2 | 0 | 86.9 |
| No match | 317.1296 | 301.1104 | 21.937 | 11104042 | 5 | 1 | 0 | 36.4 |
| No results | 344.126 | 345.1332 | 19.156 | 30918923 | 155 | 8 | 0 | 51.2 |
| Partial mat | 348.2301 | 331.2268 | 19.152 | 16027581 | 79 | 5 | 0 | 51.6 |
| No results | 280.2041 | 281.2114 | 22.124 | 25813313 | 43 | 1 | 0 | 31.8 |
| No results | 379.2724 | 380.2797 | 15.56 | 14820813 | 19 | 3 | 0 | 43.3 |
| No results | 381.2881 | 382.2953 | 16.113 | 26491297 | 20 | 2 | 0 | 59.5 |
| No results | 381.2881 | 382.2953 | 16.395 | 17288055 | 20 | 1 | 0 | 32.9 |
| No results | 466.2719 | 467.2792 | 23.604 | 10476879 | 19 | 4 | 0 | 46.4 |
| No results | 320.1624 | 321.1697 | 17.285 | 40442271 | 41 | 1 | 0 | 31.1 |
| No results | 320.1624 | 321.1696 | 18.6 | 37924603 | 41 | 2 | 0 | 39.7 |
| No results | 344.1623 | 345.1696 | 16.386 | 33959761 | 160 | 6 | 0 | 86.5 |
| No results | 298.2145 | 299.2218 | 24.94 | 15508900 | 13 | 1 | 0 | 40.5 |
| No results | 644.3099 | 323.1622 | 12.612 | 20820992 | 4 | 1 | 0 | 54.1 |
| No results | 472.2462 | 473.2535 | 21.033 | 17190066 | 139 | 2 | 0 | 40.1 |
| No results | 292.1676 | 293.1749 | 21.457 | 55620380 | 79 | 1 | 0 | 35.6 |
| No results | 288.1363 | 289.1435 | 22.359 | 7450195 | 85 | 8 | 0 | 55.8 |
| No results | 348.1936 | 349.2009 | 25.185 | 4967814 | 133 | 2 | 0 | 31.4 |
| No results | 312.205 | 313.2123 | 18.462 | 11070686 | 31 | 2 | 0 | 53.6 |
| Full match | 248.1778 | 249.1851 | 20.208 | 1.76E+08 | 50 | 3 | 0 | 69 |
| Full match | 248.1778 | 249.1851 | 21.604 | 1.42E+08 | 50 | 3 | 0 | 75.7 |
| Full match | 248.178 | 281.2115 | 22.218 | 44369123 | 52 | 3 | 0 | 77.8 |
| No results | 340.1586 | 341.1659 | 14.137 | 16526655 | 156 | 1 | 0 | 31.1 |
| No results | 352.1312 | 353.1385 | 20.003 | 8686195 | 7 | 1 | 0 | 38.1 |
| No results | 334.1781 | 335.1854 | 14.683 | 37595653 | 45 | 1 | 0 | 37.2 |
| No results | 353.0646 | 354.0718 | 14.652 | 3.19E+08 | 9 | 3 | 0 | 70.4 |

| | | | | | | | | |
|---|---|---|---|---|---|---|---|---|
| No results | 321.1689 | 322.1762 | 14.156 | 12644554 | 73 | 3 | 0 | 48.1 |
| No results | 321.169 | 322.1762 | 14.297 | 11532400 | 73 | 3 | 0 | 78.8 |
| No results | 427.0772 | 428.0844 | 17.408 | 8966728 | 3 | 1 | 0 | 32.7 |
| No results | 528.2429 | 529.2501 | 23.591 | 8846493 | 3 | 2 | 0 | 43.5 |
| No results | 254.0694 | 255.0767 | 18.617 | 21897946 | 94 | 4 | 0 | 82.1 |
| No results | 418.2356 | 419.2428 | 22.543 | 9107958 | 80 | 1 | 0 | 43.9 |
| No results | 413.1411 | 414.1483 | 12.452 | 1.81E+08 | 154 | 3 | 0 | 51.2 |
| No results | 452.1185 | 453.1258 | 23.985 | 22325723 | 2 | 2 | 0 | 67.7 |
| No results | 470.2007 | 471.208 | 23.685 | 37733232 | 10 | 4 | 0 | 48.9 |
| No results | 233.0513 | 234.0586 | 19.97 | 5811473 | 58 | 1 | 0 | 36 |
| Partial mat | 302.2457 | 303.253 | 25.227 | 1.59E+08 | 3 | 1 | 0 | 69.2 |
| Partial mat | 302.2457 | 303.253 | 25.128 | 91464778 | 3 | 1 | 0 | 71.9 |
| Partial mat | 302.2457 | 303.253 | 25.381 | 1.16E+08 | 3 | 1 | 0 | 69.2 |
| Partial mat | 302.2457 | 303.253 | 25.048 | 13414211 | 3 | 1 | 0 | 75.1 |
| Partial mat | 302.2458 | 303.253 | 24.976 | 11812868 | 3 | 1 | 0 | 73.5 |
| No results | 478.1601 | 479.1674 | 22.048 | 13943242 | 13 | 3 | 0 | 56.4 |
| No results | 230.1058 | 231.1131 | 16.514 | 7187965 | 251 | 6 | 0 | 72.8 |
| No results | 302.1519 | 303.1592 | 20.062 | 23804130 | 93 | 1 | 0 | 41 |
| Partial mat | 248.1415 | 249.1488 | 18.388 | 41062345 | 245 | 15 | 0 | 68 |
| No results | 187.0999 | 188.1071 | 19.832 | 10883753 | 60 | 3 | 0 | 52.4 |
| No results | 187.0999 | 188.1071 | 18.569 | 11556395 | 60 | 1 | 0 | 38.3 |
| No results | 342.1657 | 343.173 | 15.004 | 46304205 | 1 | 3 | 0 | 69 |
| No results | 394.1692 | 395.1764 | 20.017 | 4765813 | 78 | 4 | 0 | 38.7 |
| Full match | 260.1778 | 261.1851 | 20.809 | 15103477 | 45 | 2 | 0 | 38.5 |
| No match | 428.2325 | 429.2397 | 17.766 | 1.83E+08 | 79 | 2 | 0 | 95 |
| No results | 296.1413 | 297.1486 | 23.788 | 7632467 | 94 | 1 | 0 | 30.5 |
| No results | 418.2355 | 419.2428 | 21.244 | 23506139 | 188 | 2 | 0 | 53.8 |
| No results | 442.244 | 443.2513 | 15.669 | 10808790 | 5 | 1 | 0 | 68.7 |
| No results | 206.0845 | 207.0918 | 21.015 | 15844604 | 40 | 3 | 0 | 93.5 |
| No results | 263.1523 | 264.1596 | 19.319 | 10313047 | 200 | 2 | 0 | 50.1 |
| No results | 436.2674 | 454.3013 | 18.653 | 2.1E+08 | 1 | 2 | 0 | 31.6 |
| Full match | 235.1574 | 236.1647 | 24.654 | 4964929 | 193 | 1 | 0 | 31.3 |
| No results | 429.1108 | 430.118 | 15.37 | 13212601 | 24 | 1 | 0 | 31.2 |
| No results | 308.0798 | 309.0871 | 16.537 | 19341080 | 83 | 4 | 0 | 58.7 |
| No results | 242.0807 | 243.0879 | 13.923 | 19780895 | 87 | 4 | 0 | 86.1 |
| No results | 282.0893 | 283.0966 | 21.344 | 75058439 | 233 | 2 | 0 | 42.6 |
| No results | 280.2041 | 281.2114 | 22.209 | 44369123 | 43 | 1 | 0 | 31.8 |
| No results | 260.1626 | 261.1699 | 17.697 | 43672895 | 16 | 1 | 0 | 37.6 |
| No results | 260.1626 | 293.196 | 18.24 | 18949402 | 17 | 1 | 0 | 39.6 |
| No results | 260.1626 | 261.1699 | 18.333 | 13248062 | 17 | 1 | 0 | 34.2 |
| No results | 309.1366 | 310.1438 | 22.654 | 8052669 | 285 | 1 | 0 | 37.9 |
| No results | 242.2037 | 243.211 | 24.953 | 27037215 | 1 | 1 | 0 | 91.3 |
| No results | 264.1364 | 247.1331 | 14.89 | 9722340 | 231 | 16 | 0 | 45.4 |
| No results | 273.1368 | 274.144 | 14.206 | 17371194 | 203 | 2 | 0 | 81 |
| No results | 273.1368 | 274.144 | 19.284 | 9956794 | 203 | 2 | 0 | 68.8 |
| No results | 246.0716 | 247.0789 | 22.654 | 6188793 | 29 | 2 | 0 | 54.4 |
| Partial mat | 326.1518 | 327.1591 | 18.68 | 28003595 | 129 | 4 | 0 | 49.7 |
| No results | 414.117 | 415.1242 | 16.814 | 32413225 | 21 | 1 | 0 | 71.5 |
| Partial mat | 281.1628 | 282.1701 | 19.556 | 10521628 | 67 | 9 | 0 | 48.8 |
| No results | 267.0759 | 268.0832 | 14.031 | 31930593 | 17 | 2 | 0 | 93.2 |
| No results | 472.2914 | 473.2987 | 14.852 | 10381458 | 1 | 1 | 0 | 49.6 |

| | | | | | | | | |
|---|---|---|---|---|---|---|---|---|
| No match | 300.1032 | 301.1105 | 21.939 | 11104042 | 24 | 2 | 0 | 38.8 |
| Partial mat | 281.1629 | 282.1701 | 7.944 | 15568473 | 67 | 6 | 0 | 57.3 |
| No results | 238.1358 | 515.2346 | 22.228 | 32111990 | 17 | 2 | 0 | 45 |
| No results | 196.1578 | 197.1651 | 17.142 | 82183333 | 42 | 1 | 0 | 88.6 |
| No results | 340.1787 | 341.186 | 9.829 | 12235255 | 410 | 1 | 0 | 38.8 |
| Full match | 230.131 | 231.1383 | 19.663 | 27835803 | 78 | 5 | 0 | 42.7 |
| Full match | 230.1308 | 231.1381 | 22.756 | 31684510 | 78 | 7 | 0 | 57.5 |
| No results | 328.1675 | 329.1747 | 22.878 | 18968772 | 150 | 1 | 0 | 41.9 |
| No results | 359.0941 | 360.1014 | 11.298 | 13657141 | 159 | 2 | 0 | 54.2 |
| No results | 424.3038 | 425.3111 | 17.631 | 41305992 | 2 | 1 | 0 | 80.1 |
| Full match | 232.1215 | 233.1288 | 19.567 | 6272819 | 254 | 1 | 0 | 30.7 |
| No results | 261.1943 | 262.2015 | 19.1 | 27912013 | 5 | 1 | 0 | 44.6 |
| No results | 261.1942 | 262.2015 | 19.5 | 42980296 | 5 | 1 | 0 | 46.8 |
| No results | 354.1468 | 355.1541 | 19.683 | 7480590 | 177 | 1 | 0 | 34.8 |
| No results | 332.1988 | 333.2061 | 17.761 | 11397665 | 230 | 15 | 0 | 70.4 |
| No results | 281.1994 | 282.2066 | 21.27 | 10160141 | 38 | 2 | 0 | 50.8 |
| No results | 350.2457 | 351.253 | 24.586 | 1.54E+08 | 83 | 4 | 0 | 51.8 |
| No results | 386.1128 | 387.12 | 16.028 | 10900454 | 25 | 2 | 0 | 37.6 |
| No results | 395.1314 | 396.1387 | 21.788 | 11486583 | 241 | 4 | 0 | 68.7 |
| No results | 308.1413 | 309.1486 | 20.57 | 7810964 | 91 | 1 | 0 | 46.7 |
| No match | 330.1831 | 331.1904 | 18.549 | 71558095 | 173 | 2 | 0 | 48.2 |
| No results | 278.0695 | 279.0768 | 13.75 | 10407191 | 42 | 4 | 0 | 42.6 |
| No results | 276.184 | 277.1913 | 11.074 | 28678518 | 167 | 3 | 0 | 48.8 |
| No results | 276.184 | 277.1913 | 11.187 | 19312702 | 167 | 3 | 0 | 48.8 |
| No results | 276.184 | 277.1913 | 12.224 | 12866444 | 167 | 3 | 0 | 40.9 |
| Full match | 175.0999 | 176.1071 | 22.613 | 6956555 | 65 | 1 | 0 | 69.4 |
| Partial mat | 251.0948 | 252.1021 | 22.463 | 28215979 | 211 | 2 | 0 | 52.5 |
| No results | 288.1365 | 289.1438 | 23.986 | 5644388 | 85 | 1 | 0 | 30.8 |
| No results | 377.1626 | 378.1699 | 20.187 | 13965890 | 121 | 1 | 0 | 44 |
| No results | 446.267 | 447.2742 | 20.752 | 20376372 | 100 | 1 | 0 | 52 |
| No results | 406.1297 | 407.137 | 13.896 | 12095253 | 30 | 6 | 0 | 50.7 |
| No results | 350.2094 | 351.2167 | 23.147 | 38706375 | 143 | 14 | 0 | 60 |
| No results | 231.0686 | 232.0759 | 22.879 | 24925905 | 4 | 1 | 0 | 73.3 |
| Partial mat | 250.1571 | 251.1644 | 21.075 | 1.5E+08 | 228 | 11 | 0 | 58.3 |
| Partial mat | 250.1571 | 251.1644 | 20.779 | 1.31E+08 | 228 | 11 | 0 | 58.3 |
| Partial mat | 250.1571 | 251.1644 | 19.474 | 53803172 | 228 | 11 | 0 | 73.4 |
| Partial mat | 250.1572 | 251.1645 | 20.096 | 27760515 | 228 | 11 | 0 | 64 |
| No results | 218.0733 | 251.1068 | 21.593 | 10679692 | 5 | 1 | 0 | 30.3 |
| No results | 252.155 | 253.1622 | 22.109 | 14734557 | 1 | 3 | 0 | 65 |
| No results | 334.1778 | 335.1851 | 21.432 | 28060848 | 46 | 1 | 0 | 33.7 |
| No results | 472.319 | 473.3263 | 22.298 | 23794782 | 32 | 1 | 0 | 60.2 |
| No results | 472.319 | 473.3262 | 21.113 | 11388733 | 32 | 1 | 0 | 60.3 |
| No results | 418.1781 | 419.1854 | 22.312 | 22561607 | 1 | 1 | 0 | 39.4 |
| No results | 224.1179 | 225.1252 | 20.102 | 9306622 | 1 | 1 | 0 | 32.7 |
| No results | 318.1831 | 319.1904 | 17.616 | 19896859 | 62 | 1 | 0 | 36.8 |
| No results | 389.2777 | 390.285 | 22.882 | 19201074 | 1 | 2 | 0 | 46.7 |
| No results | 389.2777 | 390.285 | 22.767 | 12231929 | 1 | 2 | 0 | 40.6 |
| No results | 309.1214 | 310.1286 | 17.16 | 20056549 | 19 | 1 | 0 | 39.1 |
| No results | 220.0737 | 221.081 | 20.6 | 7545821 | 75 | 1 | 0 | 43.1 |
| No results | 369.1148 | 370.1221 | 13.269 | 27447085 | 248 | 3 | 0 | 35.1 |
| No results | 314.0903 | 315.0976 | 21.549 | 26368856 | 87 | 4 | 0 | 40.9 |

| | | | | | | | | |
|---|---|---|---|---|---|---|---|---|
| No results | 301.1064 | 302.1137 | 10.261 | 36408457 | 140 | 3 | 0 | 62.9 |
| No results | 645.3261 | 646.3334 | 19.638 | 44328964 | 5 | 1 | 0 | 63.7 |
| No results | 274.103 | 275.1102 | 24.293 | 5979838 | 11 | 2 | 0 | 33.6 |
| No results | 486.1958 | 487.2031 | 23.356 | 30591462 | 8 | 1 | 0 | 53.2 |
| No results | 321.0925 | 322.0998 | 15.863 | 45783932 | 2 | 2 | 0 | 50.8 |
| No results | 236.1415 | 237.1488 | 21.192 | 2.97E+08 | 57 | 1 | 0 | 33.9 |
| No results | 397.1461 | 398.1534 | 13.492 | 78173081 | 196 | 7 | 0 | 67.8 |
| No results | 332.2352 | 333.2425 | 19.52 | 6741037 | 132 | 3 | 0 | 55.1 |
| No results | 328.1246 | 329.1319 | 13.932 | 13923836 | 173 | 2 | 0 | 60.7 |
| No results | 355.2724 | 356.2797 | 14.994 | 56976929 | 14 | 2 | 0 | 47.1 |
| No results | 355.2723 | 356.2796 | 15.286 | 38793014 | 14 | 2 | 0 | 36.6 |
| No results | 393.1157 | 394.123 | 22.118 | 9352458 | 179 | 1 | 0 | 67.9 |
| No results | 390.273 | 391.2803 | 18.069 | 5254055 | 5 | 3 | 0 | 30.3 |
| No results | 301.2254 | 302.2326 | 22.092 | 9435154 | 3 | 3 | 0 | 55.1 |
| No results | 301.2254 | 302.2327 | 22.241 | 14757279 | 3 | 3 | 0 | 55 |
| No match | 255.1626 | 256.1699 | 12.943 | 30053876 | 103 | 1 | 0 | 92.5 |
| No match | 255.1628 | 256.1701 | 13.004 | 29621034 | 103 | 1 | 0 | 89.7 |
| No match | 547.236 | 548.2433 | 20.856 | 37244132 | 28 | 1 | 0 | 92.2 |
| Full match | 309.1842 | 310.1915 | 20.637 | 21113443 | 225 | 3 | 0 | 35 |
| Full match | 309.1843 | 310.1916 | 24.262 | 6875233 | 235 | 3 | 0 | 56.7 |
| Partial mat | 258.1371 | 259.1444 | 17.808 | 10788045 | 288 | 3 | 0 | 61.6 |
| No results | 363.22 | 364.2272 | 22.966 | 23777106 | 112 | 1 | 0 | 44.2 |
| No results | 320.1736 | 321.1809 | 15.683 | 1.99E+08 | 150 | 2 | 0 | 35.8 |
| No results | 320.1737 | 321.181 | 13.444 | 11694977 | 150 | 2 | 0 | 60.1 |
| No results | 377.1626 | 378.1699 | 20.39 | 6073581 | 122 | 1 | 0 | 31.8 |
| No results | 246.1371 | 247.1444 | 15.161 | 59592284 | 274 | 3 | 0 | 65 |
| No results | 384.3352 | 385.3424 | 18.987 | 53533727 | 1 | 2 | 0 | 49.8 |
| No results | 502.3023 | 503.3096 | 13.537 | 13167589 | 1 | 1 | 0 | 39.7 |
| No results | 484.2166 | 485.2239 | 22.801 | 11307759 | 5 | 3 | 0 | 44.5 |
| No results | 314.1631 | 315.1704 | 20.778 | 13496020 | 303 | 1 | 0 | 49 |
| No results | 374.1955 | 375.2028 | 18.09 | 7573278 | 56 | 8 | 0 | 52.6 |

| mzCloud_Best_Match_Confidence | mzVault_Best_Match | Mass_List_Match_EFS_HRAM_Cor | Mass_List_Match_Extractables_anc | MS2 | Reference_Ion | Area_A1 | Area_A2 | Area_A3 |
|---|---|---|---|---|---|---|---|---|
| 9 | | Single mat | No matche | PreferredD | [M+H]+1 | 5.9E+08 | 5.7E+08 | 5.7E+08 |
| 54.9 | | No matche | Single mat | PreferredD | [M+H]+1 | 98761752 | 1.73E+08 | 1.79E+08 |
| 43.8 | | No matche | Single mat | PreferredD | [M+H]+1 | 92808888 | 92635956 | 91767601 |
| 10 | | No matche | Single mat | PreferredD | [M+H]+1 | 68141862 | 70147540 | 66560735 |
| 9.8 | | No matche | No matche | PreferredD | [M+H]+1 | 1.7E+08 | 1.91E+08 | 1.49E+08 |
| 55.5 | | No matche | No matche | PreferredD | [M+H]+1 | 61956476 | 63376112 | 54958516 |
| 98 | 94.5 | No matche | Single mat | PreferredD | [M+H]+1 | 2.66E+08 | 2.5E+08 | 2.71E+08 |
| 58.8 | | No matche | Single mat | PreferredD | [M+H]+1 | 4503541 | 4764976 | 6500166 |
| | 78.1 | No matche | Single mat | PreferredD | [M+H]+1 | 2992915 | 3196964 | 2217133 |
| 60.2 | | Single mat | Single mat | PreferredD | [M+H]+1 | 3088375 | 3421106 | 4907328 |
| 37.8 | | No matche | Single mat | PreferredD | [M+H]+1 | 4661041 | 8361661 | 17320217 |
| 60.4 | | Single mat | Single mat | PreferredD | [M+H]+1 | 14454056 | 11533434 | 12864039 |
| 45.2 | | Single mat | Single mat | PreferredD | [M+H]+1 | 886734.7 | 700074.3 | 892652.4 |
| 9.3 | | No matche | Single mat | PreferredD | [M+H]+1 | 25528687 | 27613302 | 28027689 |
| 9.2 | | No matche | Single mat | PreferredD | [M+H]+1 | 15691382 | 17017573 | 17498508 |
| 9.2 | | No matche | Single mat | PreferredD | [M+H]+1 | 7349983 | 8229304 | 7415919 |
| 93.4 | 94.7 | Multiple m | Single mat | PreferredD | [M+H]+1 | 1.99E+08 | 2.31E+08 | 1.96E+08 |
| 8.4 | | No matche | No matche | PreferredD | [M+H]+1 | 81170744 | 80568177 | 78109009 |
| 60.5 | | No matche | No matche | PreferredD | [M+H]+1 | 8495685 | 48787541 | 30072649 |
| 38 | | No matche | Multiple m | PreferredD | [M+H]+1 | 8760804 | 6419704 | 6119129 |
| 41.5 | | No matche | No matche | PreferredD | [M+H]+1 | 7490353 | 18168115 | 7070355 |
| 63.8 | | No matche | No matche | PreferredD | [M+H]+1 | 3.12E+08 | 3.16E+08 | 3.27E+08 |
| 66.3 | | No matche | No matche | PreferredD | [M+H]+1 | 5.16E+08 | 5.13E+08 | 5.25E+08 |
| 64.5 | | No matche | No matche | PreferredD | [M+H]+1 | 32419417 | 30356364 | 27979335 |
| 64.7 | | No matche | No matche | PreferredD | [M+H]+1 | 9313427 | 10297042 | 10423166 |
| 61 | | Single mat | No matche | PreferredD | [M+H]+1 | 42339450 | 43753647 | 45182411 |
| 57.2 | | Single mat | No matche | PreferredD | [M+H]+1 | 15942157 | 13828286 | 18989341 |
| 67.6 | | Multiple m | Multiple m | PreferredD | [M+H]+1 | 14744572 | 13535133 | 12693809 |
| 61.2 | | No matche | No matche | PreferredD | [M+H]+1 | 8025877 | 7378231 | 7152199 |
| 9.7 | | No matche | No matche | PreferredD | [M+H]+1 | 33922396 | 36929107 | 34640245 |
| 57.4 | | No matche | No matche | PreferredD | [M+H]+1 | 7574511 | 8070427 | 8862753 |
| 7.4 | | No matche | No matche | PreferredD | [M+H]+1 | 27454652 | 28044795 | 29195190 |
| 56.6 | | No matche | No matche | PreferredD | [M+H]+1 | 18693831 | 24265628 | 13499387 |
| 7.2 | | No matche | No matche | PreferredD | [M+H]+1 | 11624013 | 17934247 | 13583076 |
| 58.7 | | No matche | No matche | PreferredD | [M+H]+1 | 26413218 | 22814184 | 22988905 |
| 61 | | Single mat | No matche | PreferredD | [M+H]+1 | 23008919 | 21316353 | 21424272 |
| 65.6 | | Single mat | No matche | PreferredD | [M+H]+1 | 5321794 | 33690232 | 8101226 |
| 65.5 | | Single mat | No matche | PreferredD | [M+H]+1 | 13017367 | 12479957 | 7371535 |
| 65.2 | | Single mat | No matche | PreferredD | [M+H]+1 | 10051946 | 13884071 | 14421441 |
| 63.3 | | Single mat | No matche | PreferredD | [M+H]+1 | 7906828 | 8742484 | 10316713 |

| | | | | | | | | |
|---|---|---|---|---|---|---|---|---|
| 61.2 | | No matche | No matche | PreferredD | [M+H]+1 | 12542203 | 13912871 | 15334105 |
| 9 | | Multiple m | No matche | PreferredD | [M+H]+1 | 24060418 | 23353181 | 23766178 |
| 57.7 | | No matche | No matche | PreferredD | [M+H]+1 | 35843717 | 32922363 | 33518916 |
| 63.5 | | No matche | No matche | PreferredD | [M+H]+1 | 30581545 | 14672866 | 27496935 |
| 41.5 | | No matche | No matche | PreferredD | [M+H]+1 | 4749477 | 6082893 | 5820364 |
| 67.8 | | No matche | No matche | PreferredD | [M+H]+1 | 1.27E+09 | 1.19E+09 | 1.21E+09 |
| 66.8 | | No matche | No matche | PreferredD | [M+H]+1 | 1.09E+08 | 1.05E+08 | 1.11E+08 |
| 66.6 | | No matche | No matche | PreferredD | [M+H]+1 | 12654044 | 12449586 | 10796171 |
| 64.4 | 57.5 | Single mat | No matche | PreferredD | [M+H]+1 | 25240911 | 26805628 | 26843860 |
| 60.6 | | No matche | No matche | PreferredD | [M+H]+1 | 2400184 | 906764.7 | 3206387 |
| 58.3 | | No matche | No matche | PreferredD | [M+H]+1 | 12356779 | 12173331 | 10386148 |
| 64.2 | | No matche | No matche | PreferredD | [M+H]+1 | 18480598 | 17603427 | 17832244 |
| 69.3 | 58.7 | No matche | No matche | PreferredD | [M+H]+1 | 58792420 | 57936190 | 57408264 |
| 65 | | Single mat | No matche | PreferredD | [M+H]+1 | 16315752 | 17558458 | 17820640 |
| 9.2 | | No matche | No matche | PreferredD | [M+H]+1 | 1811492 | 2037058 | 1667952 |
| 63.7 | | No matche | No matche | PreferredD | [M+H]+1 | 32001812 | 42774961 | 28078008 |
| 60.5 | | No matche | No matche | PreferredD | [M+H]+1 | 12282546 | 4401450 | 9987926 |
| 43 | | No matche | No matche | PreferredD | [M+H]+1 | 86861092 | 85530004 | 83253616 |
| 40.2 | | No matche | No matche | PreferredD | [M+H]+1 | 97882920 | 89896637 | 95565504 |
| 44.6 | | No matche | No matche | PreferredD | [M+H]+1 | 25814013 | 28720322 | 18185812 |
| 42.4 | | No matche | No matche | PreferredD | [M+H]+1 | 32680338 | 29699764 | 27390062 |
| 59.8 | | No matche | No matche | PreferredD | [M+H]+1 | 38543164 | 38649598 | 38624760 |
| 63.9 | | No matche | No matche | PreferredD | [M+H]+1 | 3765182 | 1115734 | 2490951 |
| 65.3 | | No matche | No matche | PreferredD | [M+H]+1 | 8502926 | 11376433 | 11509984 |
| 56.9 | | No matche | No matche | PreferredD | [M+H]+1 | 16874605 | 19352939 | 18693961 |
| 59.6 | | No matche | Multiple m | PreferredD | [M+H]+1 | 1616221 | 1717641 | 372027 |
| 6.9 | | No matche | No matche | PreferredD | [M+H]+1 | 10671315 | 9964022 | 13320041 |
| 9 | | No matche | No matche | PreferredD | [M+H]+1 | 6506669 | 6512832 | 6636311 |
| 46.9 | | Single mat | No matche | PreferredD | [M+H]+1 | 14397056 | 13662907 | 13462253 |
| 61.8 | | No matche | No matche | PreferredD | [M+H]+1 | 3646870 | 867631.4 | 6807481 |
| 83.8 | | No matche | Multiple m | PreferredD | [M+H]+1 | 1.68E+08 | 2.08E+08 | 2.87E+08 |
| 7.1 | | No matche | Single mat | PreferredD | [M+H]+1 | 18152546 | 18594945 | 26460072 |
| 38.1 | | No matche | No matche | PreferredD | [M+H]+1 | 7952066 | 7827616 | 8424092 |
| 66.7 | | Single mat | No matche | PreferredD | [M+H]+1 | 8525409 | 9763747 | 9322733 |
| 62.8 | | No matche | No matche | PreferredD | [M+H]+1 | 16931731 | 21494685 | 39529559 |
| 64.3 | | No matche | No matche | PreferredD | [M+H]+1 | 11731210 | 11670838 | 32422991 |
| 70.7 | | No matche | Single mat | PreferredD | [M+H]+1 | 13252425 | 12885896 | 12967327 |
| 7.6 | | No matche | No matche | PreferredD | [M+H]+1 | 21743374 | 20350434 | 19207992 |
| 82 | | No matche | No matche | PreferredD | [M+H]+1 | 28955649 | 23182731 | 86325104 |
| 62.3 | | No matche | No matche | PreferredD | [M+H]+1 | 16319657 | 11742489 | 23688709 |
| 59.7 | | No matche | No matche | PreferredD | [M+H]+1 | 8439776 | 7563706 | 8591524 |
| 63.2 | | No matche | No matche | PreferredD | [M+H]+1 | 10740113 | 11444267 | 11360060 |
| 61.2 | | No matche | No matche | PreferredD | [M+H]+1 | 65903571 | 48802727 | 65000436 |
| 64.7 | | No matche | No matche | PreferredD | [M+H]+1 | 64983794 | 80290671 | 76669353 |
| 63 | | No matche | No matche | PreferredD | [M+H]+1 | 16602295 | 8143255 | 7539987 |
| 47.4 | | No matche | No matche | PreferredD | [M+H]+1 | 7513670 | 12336569 | 7233538 |
| 43.2 | | No matche | No matche | PreferredD | [M+NH4]+ | 31412279 | 30523327 | 28879129 |
| 62.6 | | No matche | No matche | PreferredD | [M+H]+1 | 15916368 | 15847114 | 14581017 |
| 59.9 | | No matche | No matche | PreferredD | [M+H]+1 | 16442122 | 15068964 | 8961906 |
| 9.9 | | No matche | Single mat | PreferredD | [M+H]+1 | 8.91E+08 | 9.03E+08 | 8.31E+08 |
| 7.2 | | No matche | No matche | PreferredD | [M+H]+1 | 49644303 | 48189608 | 46859347 |

| | | | | | | | | |
|---|---|---|---|---|---|---|---|---|
| 65.8 | | Multiple m | Multiple m | PreferredD | [M+H]+1 | 7746233 | 8122252 | 8205323 |
| 59 | | No matche | No matche | PreferredD | [M+H]+1 | 17733735 | 16119119 | 15760986 |
| 44.1 | | No matche | No matche | PreferredD | [M+H]+1 | 5385169 | 5872908 | 4308200 |
| 42.5 | | No matche | No matche | PreferredD | [M+H]+1 | 21514343 | 21032726 | 20477314 |
| 43.7 | | No matche | No matche | PreferredD | [M+H]+1 | 31412279 | 30523327 | 28879129 |
| 7.8 | | No matche | No matche | PreferredD | [M+H]+1 | 15624874 | 11806393 | 11977653 |
| 60.8 | | Single mat | Multiple m | PreferredD | [M+H]+1 | 7227313 | 5268264 | 4711725 |
| 41.4 | | Single mat | No matche | PreferredD | [M+H]+1 | 16477531 | 20487375 | 23646297 |
| 41.7 | | No matche | No matche | PreferredD | [M+H]+1 | 6978803 | 7109623 | 7121566 |
| 58.3 | | No matche | No matche | PreferredD | [M+H]+1 | 16967944 | 11562471 | 13117914 |
| 7.5 | | No matche | Multiple m | PreferredD | [M+H]+1 | 41038091 | 38516412 | 39026301 |
| 47.9 | 53.1 | No matche | Multiple m | PreferredD | [M+H]+1 | 1.31E+08 | 1.22E+08 | 1.21E+08 |
| 7.3 | | No matche | No matche | PreferredD | [M+H]+1 | 13254554 | 13513537 | 9422750 |
| 63.4 | | Multiple m | No matche | PreferredD | [M+H]+1 | 6922054 | 8035362 | 7750226 |
| 63.4 | | Multiple m | No matche | PreferredD | [M+H]+1 | 12196279 | 11116087 | 12991367 |
| 9.8 | | Single mat | No matche | PreferredD | [M+H]+1 | 1.3E+08 | 1.15E+08 | 98774845 |
| 9.8 | | Single mat | No matche | PreferredD | [M+H]+1 | 55746778 | 53059311 | 50075296 |
| 9.9 | | Single mat | No matche | PreferredD | [M+H]+1 | 31261599 | 29082797 | 28801230 |
| 38.2 | | No matche | Single mat | PreferredD | [M+H]+1 | 1073102 | 898484.8 | 880309.5 |
| 72 | 80.6 | Single mat | No matche | PreferredD | [M+H]+1 | 10157718 | 9514309 | 9004219 |
| 63.8 | 56.8 | Single mat | No matche | PreferredD | [M+H]+1 | 5450557 | 4976437 | 4537731 |
| 84.9 | 77.9 | Multiple m | No matche | PreferredD | [M+H]+1 | 1.04E+08 | 95296846 | 1.01E+08 |
| 56.3 | | Single mat | Single mat | PreferredD | [M+H]+1 | 24196851 | 27061135 | 28095760 |
| 87.7 | 94.5 | Single mat | No matche | PreferredD | [M+H]+1 | 14724921 | 16588567 | 28485327 |
| 80.8 | 62.5 | Multiple m | No matche | PreferredD | [M+H]+1 | 11979620 | 11959466 | 11850865 |
| 9 | | No matche | No matche | PreferredD | [M+H]+1 | 44412449 | 46433862 | 19663578 |
| 81.9 | | Multiple m | No matche | PreferredD | [M+H]+1 | 24408937 | 25303381 | 24615316 |
| 57.7 | | No matche | No matche | PreferredD | [M+H]+1 | 1.13E+08 | 1.2E+08 | 1.16E+08 |
| 59 | | No matche | No matche | PreferredD | [M+H]+1 | 22426268 | 15162415 | 13212154 |
| 6.9 | | No matche | No matche | PreferredD | [M+H]+1 | 2.62E+08 | 2.65E+08 | 2.57E+08 |
| 71.3 | | No matche | Multiple m | PreferredD | [M+H]+1 | 4.11E+08 | 3.97E+08 | 4.04E+08 |
| 6.5 | | No matche | Multiple m | PreferredD | [M+H]+1 | 17636787 | 16580316 | 18158760 |
| 76.1 | | No matche | No matche | PreferredD | [M+H]+1 | 36033087 | 35561069 | 35103009 |
| 46.5 | | No matche | Single mat | PreferredD | [M+H]+1 | 33231949 | 33762727 | 35609557 |
| 44.1 | | No matche | Single mat | PreferredD | [M+H]+1 | 23063123 | 24364949 | 23942253 |
| 6.9 | | No matche | No matche | PreferredD | [M+H]+1 | 15597732 | 14522006 | 12755168 |
| 9.2 | 83.6 | No matche | Single mat | PreferredD | [M+H]+1 | 9232656 | 10831760 | 9185831 |
| 6.5 | | No matche | Single mat | PreferredD | [M+H]+1 | 71633051 | 72794367 | 74267350 |
| 8.7 | | No matche | Single mat | PreferredD | [M+H]+1 | 2912204 | 3254417 | 2866134 |
| 39.2 | | Multiple m | Single mat | PreferredD | [M+H]+1 | 4259938 | 5279024 | 5309517 |
| 38.1 | | No matche | Single mat | PreferredD | [M+H]+1 | 35395715 | 32834809 | 33400064 |
| 64.8 | | No matche | Multiple m | PreferredD | [M+H]+1 | 17666524 | 28123751 | 12999283 |
| 57.5 | | No matche | Multiple m | PreferredD | [M+H]+1 | 4723466 | 4912594 | 4917387 |
| 9.1 | | Multiple m | No matche | PreferredD | [M+H]+1 | 30461079 | 29576720 | 28854385 |
| 7.5 | | No matche | No matche | PreferredD | [M+H]+1 | 5593656 | 5039046 | 5041131 |
| 58 | | No matche | No matche | PreferredD | [M+H]+1 | 27537016 | 24634453 | 26759458 |
| 58.2 | | No matche | No matche | PreferredD | [M+H]+1 | 8572568 | 7714790 | 8218292 |
| 65.2 | 87.7 | Multiple m | No matche | PreferredD | [M+H]+1 | 1.04E+08 | 1.03E+08 | 95688323 |
| 9.3 | | Multiple m | No matche | PreferredD | [M+H]+1 | 19997550 | 20067823 | 19833709 |
| 53.7 | 74.5 | No matche | No matche | PreferredD | [M+H]+1 | 12137334 | 13000428 | 12653531 |
| 62.2 | | Multiple m | Single mat | PreferredD | [M+H]+1 | 21327446 | 17915559 | 18585370 |

| | | | | | | | | |
|---|---|---|---|---|---|---|---|---|
| 59.4 | | Multiple m | Single mat | PreferredD | [M+H]+1 | 9682655 | 9641663 | 8721762 |
| 52.6 | 71.4 | Multiple m | No matche | PreferredD | [M+H]+1 | 38281852 | 33503114 | 32609668 |
| 9.6 | | Single mat | No matche | PreferredD | [M+H]+1 | 14313648 | 12590700 | 14767139 |
| 47 | | No matche | No matche | PreferredD | [M+H]+1 | 53835953 | 42565831 | 39010521 |
| 7.4 | | Multiple m | No matche | PreferredD | [M+H]+1 | 14618166 | 9816921 | 9085098 |
| 83.2 | | No matche | No matche | PreferredD | [M+H]+1 | 58300885 | 57953321 | 56748955 |
| 9.3 | | No matche | No matche | PreferredD | [M+H]+1 | 15570482 | 14930023 | 16338241 |
| 9.5 | | No matche | No matche | PreferredD | [M+H]+1 | 2.17E+09 | 2.1E+09 | 2.04E+09 |
| 10 | | Multiple m | Multiple m | PreferredD | [M+H]+1 | 1.28E+08 | 1.3E+08 | 1.31E+08 |
| 8.1 | | Multiple m | Multiple m | PreferredD | [M+H]+1 | 11220134 | 12045835 | 6186723 |
| 9.3 | | Multiple m | No matche | PreferredD | [M+H]+1 | 23044667 | 23157964 | 23136980 |
| 45.2 | | No matche | No matche | PreferredD | [M+H]+1 | 27957949 | 27051854 | 26372994 |
| 58.5 | | No matche | No matche | PreferredD | [M+H-NH3 | 13939560 | 12408868 | 13391684 |
| 7 | 66.4 | No matche | No matche | OtherDDA | [M+2H]+2 | 1.44E+08 | 1.41E+08 | 95553191 |
| 79.7 | 88 | No matche | No matche | PreferredD | [M+H]+1 | 20619508 | 19815995 | 19764546 |
| 80.8 | 87.8 | Single mat | No matche | PreferredD | [M+2H]+2 | 29065043 | 28345669 | 29397158 |
| 7.9 | | No matche | No matche | PreferredD | [M+H]+1 | 9420474 | 11180912 | 11172847 |
| 78.2 | 86.6 | Single mat | No matche | PreferredD | [M+H]+1 | 5966338 | 5987592 | 6038987 |
| 6.6 | | No matche | No matche | PreferredD | [M+H]+1 | 978875.9 | 1065486 | 932743.4 |
| 55.7 | | Single mat | No matche | PreferredD | [M+H]+1 | 6852462 | 6685606 | 6904510 |
| 51.2 | | No matche | No matche | PreferredD | [M+H]+1 | 35665990 | 36167720 | 35814496 |
| 9.6 | | Single mat | No matche | PreferredD | [M+H]+1 | 90137100 | 88073595 | 85321950 |
| 8.2 | | Single mat | No matche | PreferredD | [M+H]+1 | 22510186 | 23336887 | 24705153 |
| 43.8 | | Multiple m | No matche | PreferredD | [M+H]+1 | 6407784 | 6597240 | 7086940 |
| 43.1 | | Multiple m | No matche | PreferredD | [M+H]+1 | 6701856 | 6462644 | 6251709 |
| 87.7 | 90.7 | Single mat | No matche | PreferredD | [M+H]+1 | 1.16E+08 | 1.09E+08 | 1.09E+08 |
| 10 | | Single mat | No matche | PreferredD | [M+H]+1 | 86795789 | 88142694 | 83200950 |
| 71.3 | 82.5 | Multiple m | No matche | PreferredD | [M+H]+1 | 45112379 | 51265175 | 48334613 |
| 57.7 | | Single mat | No matche | PreferredD | [M+H]+1 | 8200992 | 7449607 | 6448850 |
| 80.8 | | No matche | No matche | PreferredD | [M+H]+1 | 1.34E+08 | 1.32E+08 | 1.32E+08 |
| 59.1 | | Single mat | No matche | PreferredD | [M+H]+1 | 371241.6 | 381233.3 | 1617480 |
| 9.7 | 79.5 | Single mat | No matche | PreferredD | [M+H]+1 | 1.59E+08 | 1.57E+08 | 1.55E+08 |
| 93.4 | 89.3 | Single mat | No matche | PreferredD | [M+H]+1 | 1.78E+09 | 1.73E+09 | 1.73E+09 |
| 59 | | Single mat | No matche | PreferredD | [M+H]+1 | 695721.7 | 319362.2 | 195664.3 |
| 63.3 | | No matche | No matche | PreferredD | [M+H]+1 | 7789807 | 6965267 | 7732263 |
| 44 | | No matche | No matche | PreferredD | [M+H]+1 | 8579584 | 7754815 | 7884504 |
| 54.9 | | No matche | Single mat | PreferredD | [M+H]+1 | 3253687 | 3468063 | 5361160 |
| 64.9 | | Single mat | Multiple m | PreferredD | [M+H]+1 | 17480044 | 16633645 | 2694333 |
| 42.9 | | No matche | Single mat | PreferredD | [M+H]+1 | 672127.7 | 4108975 | 1424625 |
| 46.1 | 84 | No matche | No matche | PreferredD | [M+H]+1 | 9367956 | 9412502 | 9912443 |
| 8.3 | | Single mat | No matche | PreferredD | [M+H]+1 | 7797389 | 6330501 | 6711779 |
| 79.9 | | Single mat | No matche | PreferredD | [M+H]+1 | 13094727 | 11845808 | 12900933 |
| 9.4 | | No matche | No matche | PreferredD | [M+H]+1 | 12623377 | 12215212 | 12157502 |
| 100 | 73.4 | Single mat | No matche | PreferredD | [M+H]+1 | 2.53E+08 | 2.62E+08 | 2.58E+08 |
| 63.3 | | Single mat | No matche | PreferredD | [M+H]+1 | 39242712 | 35569165 | 6157300 |
| 85.4 | 75.4 | No matche | No matche | PreferredD | [M+H]+1 | 60220305 | 59977861 | 57967273 |
| 85.5 | | No matche | No matche | PreferredD | [M+H]+1 | 83152190 | 85480151 | 85626207 |
| 70.4 | 50.1 | Single mat | No matche | PreferredD | [M+H]+1 | 8752141 | 9199372 | 9343655 |
| 74.2 | 83.9 | No matche | No matche | PreferredD | [M+H]+1 | 79780756 | 79282233 | 80258776 |
| 83.4 | 80.6 | Single mat | No matche | PreferredD | [M+H]+1 | 18985743 | 19350041 | 19199912 |
| 87.4 | | Single mat | Single mat | PreferredD | [M+H]+1 | 85057151 | 82442920 | 84760274 |

| | | | | | | | | |
|---|---|---|---|---|---|---|---|---|
| 63.6 | | Single mat | No matche | PreferredD | [M+H]+1 | 12816046 | 11171536 | 12630298 |
| 88.9 | 87.7 | No matche | No matche | PreferredD | [M+H]+1 | 8.68E+08 | 8.51E+08 | 8.57E+08 |
| 82.1 | | No matche | Single mat | PreferredD | [M+H]+1 | 1.92E+08 | 1.74E+08 | 1.74E+08 |
| 84.2 | 92.1 | Single mat | No matche | PreferredD | [M+H]+1 | 76794600 | 78066303 | 76437222 |
| 6.7 | | No matche | No matche | PreferredD | [M+H]+1 | 8593023 | 8309790 | 8955331 |
| 40.8 | | No matche | No matche | PreferredD | [M+H]+1 | 3115381 | 3472444 | 3396064 |
| 82.9 | | No matche | Multiple m | PreferredD | [M+H]+1 | 1.43E+08 | 1.38E+08 | 1.25E+08 |
| 41.3 | | Single mat | No matche | PreferredD | [M+H]+1 | 86046745 | 86720465 | 86742246 |
| 9.3 | | Single mat | No matche | PreferredD | [M+H]+1 | 14003471 | 13641719 | 14019826 |
| | 54.6 | No matche | No matche | PreferredD | [M+H]+1 | 32187524 | 30354317 | 30750275 |
| 53 | | Multiple m | No matche | PreferredD | [M+H]+1 | 3.67E+08 | 3.36E+08 | 3.53E+08 |
| 9.4 | | Multiple m | No matche | PreferredD | [M+H]+1 | 70220325 | 71034231 | 72003788 |
| 46.7 | | Multiple m | No matche | PreferredD | [M+H]+1 | 3992896 | 2129438 | 1863712 |
| 60.1 | | Multiple m | No matche | PreferredD | [M+H]+1 | 1291970 | 6090200 | 5296294 |
| 58.1 | | Single mat | No matche | PreferredD | [M+H]+1 | 13888610 | 17099194 | 15215449 |
| 6.9 | | No matche | No matche | PreferredD | [M+H]+1 | 70551384 | 68470322 | 68951041 |
| 44 | | No matche | No matche | PreferredD | [M+H]+1 | 7363558 | 7711226 | 5494112 |
| 63.1 | | Multiple m | No matche | PreferredD | [M+H]+1 | 34397684 | 32375732 | 36239034 |
| 65.8 | | Multiple m | No matche | PreferredD | [M+H]+1 | 32253886 | 30705389 | 34747304 |
| 60.5 | | Multiple m | No matche | PreferredD | [M+H]+1 | 12020854 | 12879871 | 15433510 |
| 38.4 | | No matche | No matche | PreferredD | [M+H]+1 | 5969447 | 7609638 | 7223913 |
| 6.6 | | No matche | No matche | PreferredD | [M+H]+1 | 7921388 | 7829472 | 7220390 |
| 44.3 | | No matche | No matche | PreferredD | [M+H]+1 | 30741563 | 30896969 | 28164963 |
| 45.5 | | Single mat | No matche | PreferredD | [M+H]+1 | 46777516 | 47808798 | 48767776 |
| 47.4 | | Single mat | No matche | PreferredD | [M+H]+1 | 29674054 | 23704945 | 26152071 |
| 44.7 | | Single mat | No matche | PreferredD | [M+H]+1 | 11292713 | 10591369 | 10989821 |
| 44.7 | | No matche | No matche | PreferredD | [M+H]+1 | 58314221 | 33743809 | 36619920 |
| 56.5 | | Single mat | No matche | PreferredD | [M+H]+1 | 50696363 | 49968541 | 47348602 |
| 61.6 | | No matche | No matche | PreferredD | [M+H]+1 | 7190109 | 7911899 | 8322670 |
| 62.4 | | No matche | No matche | PreferredD | [M+H]+1 | 8219907 | 7741179 | 7833708 |
| 7.5 | | Single mat | No matche | PreferredD | [M+H]+1 | 9858944 | 12783351 | 4257663 |
| 66.4 | | Single mat | No matche | PreferredD | [M+H]+1 | 76010927 | 77390063 | 79700616 |
| 43.6 | | Single mat | No matche | PreferredD | [M+H]+1 | 6710440 | 8072785 | 7356181 |
| 65.9 | | Single mat | No matche | PreferredD | [M+H]+1 | 15675738 | 13711320 | 15134699 |
| 76.8 | | Multiple m | No matche | PreferredD | [M+H]+1 | 46269087 | 45661172 | 43092750 |
| 61.1 | | Multiple m | No matche | PreferredD | [M+H]+1 | 564195.1 | 836163.2 | 851620.2 |
| 69.6 | | Single mat | No matche | PreferredD | [M+H]+1 | 468313.3 | 470473.3 | 525980.1 |
| 46.7 | | Single mat | No matche | PreferredD | [M+H]+1 | 519744.3 | 529006.4 | 575120.4 |
| 84.1 | 87.6 | Multiple m | No matche | PreferredD | [M+H]+1 | 1.88E+08 | 1.89E+08 | 1.9E+08 |
| 62 | | Multiple m | No matche | PreferredD | [M+H]+1 | 10229606 | 10352426 | 10061733 |
| 91.8 | | Multiple m | Multiple m | PreferredD | [M+H]+1 | 42620538 | 46836175 | 45114985 |
| 61.9 | | Multiple m | Multiple m | PreferredD | [M+H]+1 | 11132747 | 13885928 | 15892142 |
| 69.2 | | Single mat | No matche | PreferredD | [M+H]+1 | 57399807 | 54070369 | 53198971 |
| 54.3 | 77.9 | Single mat | No matche | PreferredD | [M+H]+1 | 37997849 | 36686644 | 34846395 |
| 98.4 | 95.2 | Multiple m | Single mat | PreferredD | [M+H]+1 | 2.76E+08 | 2.7E+08 | 2.75E+08 |
| 91.3 | 89.3 | Multiple m | Single mat | PreferredD | [M+H]+1 | 44451645 | 51630420 | 57731169 |
| 6.7 | | Multiple m | Single mat | PreferredD | [M+H]+1 | 12040829 | 12085016 | 12792244 |
| 7.5 | | No matche | Single mat | PreferredD | [M+H]+1 | 23473402 | 22123498 | 21597422 |
| 38 | | No matche | Single mat | PreferredD | [M+H]+1 | 21104286 | 16718116 | 5499866 |
| 58.2 | 52 | Single mat | Single mat | PreferredD | [M+H]+1 | 14001476 | 14980024 | 13444606 |
| 67 | | Single mat | No matche | PreferredD | [M+H]+1 | 13453458 | 13002208 | 12768944 |

| | | | | | | | | |
|---|---|---|---|---|---|---|---|---|
| 59.3 | | Single mat | No matche | PreferredD | [M+H]+1 | 16542415 | 26381978 | 27303723 |
| 37.8 | | No matche | No matche | PreferredD | [M+H]+1 | 4372004 | 3968101 | 4107533 |
| 54 | 62.1 | Single mat | No matche | PreferredD | [M+H]+1 | 10127190 | 9614493 | 8301730 |
| 64.4 | | No matche | No matche | PreferredD | [M+H]+1 | 1501915 | 592974.2 | 348240 |
| 61.2 | | No matche | Single mat | PreferredD | [M+H]+1 | 37369834 | 29597272 | 29171419 |
| 60.4 | | No matche | Single mat | PreferredD | [M+H]+1 | 17189994 | 11774475 | 15682241 |
| 64.1 | | No matche | Single mat | PreferredD | [M+H]+1 | 11870531 | 15112448 | 15926107 |
| 57.7 | | No matche | Single mat | PreferredD | [M+H]+1 | 19302991 | 12830229 | 18069780 |
| 57.7 | | No matche | No matche | PreferredD | [M+H]+1 | 8521545 | 6847910 | 9442618 |
| 63.6 | | No matche | Single mat | PreferredD | [M+H+Me( | 24874655 | 22888313 | 22433953 |
| 63.5 | | No matche | Single mat | PreferredD | [M+NH4]+ | 14040141 | 14757279 | 12766918 |
| 57.6 | | No matche | No matche | PreferredD | [M+H]+1 | 8302278 | 36428489 | 39521302 |
| 56.7 | | No matche | No matche | PreferredD | [M+H]+1 | 13193978 | 8270409 | 9563453 |
| 70.6 | | No matche | No matche | PreferredD | [M+H]+1 | 32702318 | 54075306 | 20157388 |
| 42.4 | | No matche | No matche | PreferredD | [M+H]+1 | 371088.9 | 288154.5 | 287312.2 |
| 72 | | No matche | No matche | PreferredD | [M+H]+1 | 68917771 | 65985561 | 64040735 |
| 71.1 | | No matche | No matche | PreferredD | [M+H]+1 | 2261265 | 2608360 | 17363602 |
| 67.8 | | No matche | No matche | PreferredD | [M+H]+1 | 59523326 | 9487078 | 7598636 |
| 67.8 | | No matche | No matche | PreferredD | [M+H]+1 | 12544099 | 10287548 | 15682416 |
| 65.2 | | No matche | No matche | PreferredD | [M+H]+1 | 9038587 | 7026756 | 8558063 |
| 42 | | No matche | Single mat | PreferredD | [M+NH4]+ | 4891094 | 2189224 | 4700421 |
| 64.7 | | No matche | No matche | PreferredD | [M+H]+1 | 25098980 | 70219417 | 57739416 |
| 56.6 | | No matche | Single mat | PreferredD | [M+H]+1 | 40156855 | 34491473 | 51057528 |
| 61.7 | | No matche | No matche | PreferredD | [M+H]+1 | 13171872 | 12755992 | 7696013 |
| 60.3 | | Single mat | No matche | PreferredD | [M+H]+1 | 1178313 | 475490.5 | 2213420 |
| 60.2 | | Single mat | No matche | PreferredD | [M+H]+1 | 1895907 | 714796 | 1686387 |
| 60.2 | | Single mat | No matche | PreferredD | [M+H]+1 | 1084462 | 460800.2 | 668619.3 |
| 58.6 | | Single mat | No matche | PreferredD | [M+H]+1 | 9906887 | 9846003 | 10291757 |
| 58.3 | | No matche | No matche | PreferredD | [M+H]+1 | 3475283 | 3379798 | 3069049 |
| 38.7 | | No matche | No matche | PreferredD | [M+H]+1 | 1859695 | 1743500 | 2208078 |
| 65.4 | | No matche | No matche | PreferredD | [M+H]+1 | 86830091 | 83937220 | 81927282 |
| 43.3 | | Multiple m | No matche | PreferredD | [M+H]+1 | 3063264 | 11713914 | 12931630 |
| 64.6 | | No matche | No matche | PreferredD | [M+H]+1 | 6.29E+08 | 5.95E+08 | 6.09E+08 |
| 41.5 | | No matche | No matche | PreferredD | [M+H]+1 | 1.73E+08 | 1.82E+08 | 1.69E+08 |
| 9.1 | | No matche | No matche | PreferredD | [M+H]+1 | 2.59E+08 | 1.26E+08 | 2.74E+08 |
| 7.3 | | No matche | No matche | PreferredD | [M+H]+1 | 1.35E+08 | 1.26E+08 | 1.28E+08 |
| 9.1 | | No matche | No matche | PreferredD | [M+H]+1 | 4.31E+08 | 2.68E+08 | 1.01E+08 |
| 9 | | No matche | No matche | PreferredD | [M+H]+1 | 3.02E+08 | 2.7E+08 | 2.87E+08 |
| 55.1 | | No matche | No matche | PreferredD | [M+H]+1 | 1.58E+08 | 1.54E+08 | 1.54E+08 |
| 6.6 | | No matche | No matche | PreferredD | [M+H]+1 | 1904928 | 935945.8 | 1150221 |
| 7.4 | | No matche | No matche | PreferredD | [M+H]+1 | 87058934 | 83862047 | 88086369 |
| 6.7 | | No matche | No matche | PreferredD | [M+2H]+2 | 29953707 | 17817150 | 15016134 |
| 44.6 | | No matche | No matche | PreferredD | [M+H]+1 | 1E+08 | 97485949 | 99998214 |
| 7.5 | | No matche | No matche | PreferredD | [M+H]+1 | 66023573 | 64709096 | 61191457 |
| 41.2 | | No matche | No matche | PreferredD | [M+H]+1 | 1.04E+08 | 1.03E+08 | 1.07E+08 |
| 38.7 | | No matche | No matche | PreferredD | [M+H]+1 | 1.7E+08 | 1.57E+08 | 1.66E+08 |
| 39.2 | | No matche | No matche | PreferredD | [M+H]+1 | 67721811 | 61485734 | 63681034 |
| 67.5 | | No matche | No matche | PreferredD | [M+2H]+2 | 59001873 | 60212578 | 63745978 |
| 61.7 | | No matche | No matche | PreferredD | [M+H]+1 | 69207794 | 64274583 | 65079983 |
| 6.7 | | No matche | No matche | OtherDDA | [M+2H]+2 | 1.19E+08 | 1.46E+08 | 1.41E+08 |
| 6.9 | | No matche | No matche | PreferredD | [M+H]+1 | 22021372 | 20938462 | 21421561 |

| | | | | | | | | |
|---|---|---|---|---|---|---|---|---|
| 58.4 | | No matche | No matche | PreferredD | [M+H]+1 | 53299417 | 56503889 | 55526054 |
| 7.1 | | No matche | No matche | PreferredD | [M+H]+1 | 50893879 | 49432567 | 48421284 |
| 82.9 | | No matche | No matche | PreferredD | [M+H]+1 | 1.35E+08 | 1.19E+08 | 1.2E+08 |
| 82.9 | | No matche | No matche | PreferredD | [M+H]+1 | 1.1E+08 | 1.03E+08 | 1.17E+08 |
| 6.6 | | No matche | No matche | PreferredD | [M+H]+1 | 2.1E+08 | 1.98E+08 | 2.05E+08 |
| 82.4 | | No matche | No matche | PreferredD | [M+H]+1 | 96350860 | 98920601 | 1.01E+08 |
| 40.2 | | No matche | No matche | PreferredD | [M+H]+1 | 50023146 | 69187720 | 88411352 |
| 78.4 | | Single mat | No matche | PreferredD | [M+H]+1 | 29162736 | 29304974 | 26374138 |
| 67.1 | | No matche | No matche | PreferredD | [M+H]+1 | 24331748 | 52192769 | 50150435 |
| 44.6 | | No matche | No matche | PreferredD | [M+H]+1 | 34496080 | 30730151 | 30374892 |
| 7.4 | | No matche | No matche | PreferredD | [M+H]+1 | 38789418 | 43269454 | 44816095 |
| 8.2 | | No matche | No matche | PreferredD | [M+H]+1 | 76615863 | 78185407 | 80907356 |
| 65.6 | | No matche | No matche | PreferredD | [M+H]+1 | 1.1E+08 | 1.08E+08 | 1.1E+08 |
| 82.9 | | No matche | No matche | PreferredD | [M+H]+1 | 1.58E+08 | 1.68E+08 | 1.61E+08 |
| 82.7 | | No matche | No matche | PreferredD | [M+H]+1 | 1.26E+08 | 1.54E+08 | 1.07E+08 |
| 8.4 | | No matche | No matche | PreferredD | [M+H]+1 | 44286797 | 45699202 | 44789834 |
| 7.2 | | No matche | No matche | PreferredD | [M+H]+1 | 23934693 | 23874606 | 20623268 |
| 8 | | No matche | No matche | PreferredD | [M+H]+1 | 42925205 | 41385662 | 45958867 |
| 6.7 | | No matche | No matche | PreferredD | [M+H]+1 | 16237646 | 28283462 | 77245765 |
| 82.4 | | No matche | No matche | PreferredD | [M+H]+1 | 71959446 | 59891354 | 70820496 |
| 7.8 | | No matche | No matche | PreferredD | [M+H]+1 | 20203018 | 18738710 | 18853756 |
| 82.4 | | No matche | No matche | PreferredD | [M+H]+1 | 82025968 | 81042694 | 92356540 |
| 39.8 | | No matche | No matche | PreferredD | [M+H]+1 | 44216327 | 50164995 | 51780942 |
| 70.1 | | No matche | No matche | PreferredD | [M+H]+1 | 43471512 | 43927473 | 43327585 |
| 6.6 | | No matche | No matche | PreferredD | [M+H]+1 | 88642143 | 83945223 | 89342952 |
| 9.1 | | No matche | No matche | PreferredD | [M+H]+1 | 3498545 | 3744004 | 3933091 |
| 62 | | Multiple m | No matche | PreferredD | [M+H]+1 | 11501685 | 10545939 | 10494743 |
| 9.6 | | Single mat | No matche | PreferredD | [M+H]+1 | 3025480 | 8470315 | 9084452 |
| 60 | | No matche | No matche | PreferredD | [M+H]+1 | 24429099 | 5425066 | 14585447 |
| 41.9 | | No matche | No matche | PreferredD | [M+2H]+2 | 13957795 | 13656124 | 14790065 |
| 8.3 | | No matche | No matche | PreferredD | [M+H]+1 | 1.2E+08 | 1.16E+08 | 1.22E+08 |
| 61.7 | | Single mat | No matche | PreferredD | [M+H]+1 | 34148166 | 32722806 | 35070010 |
| 6.7 | | No matche | No matche | OtherDDA | [M+3H]+3 | 42394780 | 44065368 | 45697278 |
| 82.5 | | Single mat | No matche | PreferredD | [M+2H]+2 | 13329107 | 12474661 | 13422729 |
| 83.9 | | No matche | No matche | PreferredD | [M+H]+1 | 87133945 | 82768016 | 84895530 |
| 8.1 | | No matche | No matche | PreferredD | [M+H]+1 | 83074043 | 1.72E+08 | 1.81E+08 |
| 68.8 | | No matche | No matche | PreferredD | [M+H]+1 | 31138776 | 25422730 | 25741352 |
| 7.3 | | No matche | No matche | PreferredD | [M+H]+1 | 52192073 | 48223024 | 8983668 |
| 6.5 | | No matche | No matche | PreferredD | [M+H]+1 | 62268170 | 50858720 | 60773463 |
| 64.2 | | No matche | No matche | PreferredD | [M+NH4]+ | 18480598 | 16224659 | 17832244 |
| 7.2 | | No matche | No matche | PreferredD | [M+H]+1 | 4379153 | 28139269 | 7778816 |
| 60.8 | | No matche | No matche | PreferredD | [M+H]+1 | 27171631 | 27348681 | 27076168 |
| 79.9 | | No matche | No matche | PreferredD | [M+H]+1 | 70054024 | 66817642 | 65811602 |
| 94.9 | | No matche | No matche | PreferredD | [M+H]+1 | 97356162 | 93730870 | 98127043 |
| 62.6 | | No matche | No matche | PreferredD | [M+H]+1 | 26105136 | 24166820 | 25155645 |
| 66.1 | | Multiple m | No matche | PreferredD | [M+H]+1 | 12091888 | 14111461 | 14271970 |
| 39.9 | | No matche | No matche | PreferredD | [M+H]+1 | 32638380 | 30773139 | 30867408 |
| 41.6 | | No matche | No matche | PreferredD | [M+H]+1 | 23078234 | 21920353 | 20164798 |
| 7.2 | | No matche | No matche | PreferredD | [M+H]+1 | 31260660 | 31006523 | 30903649 |
| 59 | | No matche | No matche | PreferredD | [M+H]+1 | 13335991 | 12350744 | 11744195 |
| 63.7 | | No matche | No matche | PreferredD | [M+H-H2O | 19784534 | 25880868 | 25405577 |

| | | | | | | | |
|---|---|---|---|---|---|---|---|
| 6.5 | No matche | No matche | OtherDDA | [M+2H]+2 | 20275652 | 18531156 | 17831067 |
| 63.6 | No matche | No matche | PreferredD | [M+H]+1 | 25639593 | 25318537 | 25584240 |
| 8.7 | No matche | No matche | PreferredD | [M+H]+1 | 20343658 | 25328668 | 25362113 |
| 6.6 | No matche | No matche | PreferredD | [M+H]+1 | 25008318 | 6885591 | 31933329 |
| 56.7 | No matche | No matche | PreferredD | [2M+H]+1 | 19706647 | 21677117 | 20766198 |
| 80.1 | No matche | No matche | PreferredD | [M+H]+1 | 39244239 | 35669664 | 35501740 |
| 9.6 | No matche | No matche | PreferredD | [M+H]+1 | 58037368 | 53330386 | 61868179 |
| 42.9 | No matche | No matche | PreferredD | [M+H]+1 | 19176347 | 20312928 | 20140831 |
| 7.7 | No matche | No matche | PreferredD | [M+H]+1 | 18748528 | 23208333 | 21255317 |
| 7.8 | No matche | No matche | PreferredD | [M+H]+1 | 42499058 | 40079760 | 26123299 |
| 6.7 | No matche | No matche | PreferredD | [M+H]+1 | 17725937 | 18733831 | 17894912 |
| 62.8 | No matche | No matche | PreferredD | [M+H]+1 | 28406240 | 26615297 | 27598786 |
| 61.1 | No matche | No matche | PreferredD | [M+H]+1 | 19945776 | 29310368 | 31739508 |
| 63.5 | Single mat | No matche | PreferredD | [M+H]+1 | 35351719 | 38027784 | 34062096 |
| 7 | No matche | No matche | PreferredD | [M+H]+1 | 22492077 | 22377526 | 23749897 |
| 45.6 | No matche | No matche | PreferredD | [M+H]+1 | 22679476 | 24066274 | 21543978 |
| 7.8 | No matche | No matche | PreferredD | [M+H]+1 | 14444310 | 41620809 | 28716107 |
| 47.6 | No matche | No matche | PreferredD | [M+H]+1 | 1940595 | 1942511 | 2099063 |
| 38.1 | No matche | No matche | PreferredD | [M+H]+1 | 9573031 | 9369733 | 9651016 |
| 6.6 | No matche | No matche | PreferredD | [M+H]+1 | 22388670 | 21960479 | 20560633 |
| 59.4 | No matche | No matche | PreferredD | [M+NH4]+ | 30342568 | 27354075 | 29045069 |
| 61.5 | Single mat | No matche | PreferredD | [M+H]+1 | 11251991 | 12771634 | 13639104 |
| 62.8 | No matche | No matche | PreferredD | [M+H]+1 | 33364964 | 26175074 | 36479405 |
| 43.2 | No matche | No matche | PreferredD | [M+H]+1 | 16930394 | 17605866 | 17925267 |
| 55.4 | No matche | No matche | PreferredD | [M+H]+1 | 34481422 | 32086021 | 31626958 |
| 64.7 | No matche | No matche | PreferredD | [M+H]+1 | 16660898 | 15237276 | 14103467 |
| 42.2 | No matche | No matche | PreferredD | [M+NH4]+ | 9299936 | 8581356 | 8273072 |
| 41.5 | No matche | No matche | PreferredD | [M+NH4]+ | 24812533 | 22526373 | 23442446 |
| 65.3 | No matche | Single mat | PreferredD | [M+H]+1 | 723982.4 | 1002080 | 2380254 |
| 38.3 | No matche | No matche | PreferredD | [M+H]+1 | 1980556 | 1499753 | 2157358 |
| 43.5 | No matche | No matche | PreferredD | [M+H]+1 | 57029667 | 54121653 | 1.06E+08 |
| 41.6 | No matche | No matche | PreferredD | [M+H]+1 | 68550754 | 72288721 | 40795037 |
| 58.1 | No matche | No matche | PreferredD | [M+H-H2O | 8296129 | 3465897 | 1544975 |
| 9.6 | No matche | No matche | PreferredD | [M+H]+1 | 22709281 | 16935037 | 25892987 |
| 57.1 | No matche | No matche | PreferredD | [M+NH4]+ | 9042435 | 18985961 | 15978466 |
| 57.1 | No matche | No matche | PreferredD | [M+H]+1 | 16750871 | 16055303 | 17864398 |
| 40.9 | No matche | No matche | PreferredD | [M+H]+1 | 28976569 | 31116771 | 32319912 |
| 57 | No matche | No matche | PreferredD | [M+H]+1 | 9024357 | 12101442 | 10557182 |
| 56.3 | No matche | No matche | PreferredD | [M+H-H2O | 1800634 | 253690.8 | 270442 |
| 61.6 | No matche | No matche | PreferredD | [M+H]+1 | 36847749 | 79720290 | 66798742 |
| 50.7 | No matche | No matche | PreferredD | [M+H+Me( | 20619508 | 19815995 | 19764546 |
| 41 | No matche | No matche | PreferredD | [M+H]+1 | 10231830 | 9862690 | 9758179 |
| 49.5 | No matche | No matche | PreferredD | [M+H]+1 | 897819.3 | 1094616 | 1083217 |
| 6.8 | No matche | No matche | PreferredD | [M+H]+1 | 10924251 | 15125737 | 16174539 |
| 44.1 | No matche | No matche | PreferredD | [M+H]+1 | 48989635 | 47007150 | 36967547 |
| 42.8 | Single mat | No matche | PreferredD | [M+H]+1 | 20285938 | 20510722 | 20381954 |
| 8 | No matche | No matche | PreferredD | [M+H]+1 | 21811907 | 22167254 | 22409996 |
| 60.8 | No matche | No matche | PreferredD | [M+H]+1 | 24268113 | 21599694 | 22365420 |
| 59 | No matche | No matche | PreferredD | [M+NH4]+ | 24426284 | 23486428 | 20730866 |
| 57.6 | No matche | No matche | PreferredD | [M+H-H2O | 10830492 | 13411433 | 15703787 |
| 65.1 | No matche | No matche | PreferredD | [M+H]+1 | 12411961 | 13019089 | 13803296 |

| | | | | | | | |
|---|---|---|---|---|---|---|---|
| 7 | No matche | No matche | PreferredD | [M+H]+1 | 9747039 | 8608000 | 9176130 |
| 76.3 | No matche | No matche | PreferredD | [M+H]+1 | 14184113 | 14529530 | 14471259 |
| 41.5 | No matche | No matche | PreferredD | [M+H]+1 | 11063210 | 11892916 | 35193420 |
| 66.2 | No matche | Multiple m | PreferredD | [M+H]+1 | 38256757 | 19213351 | 19476217 |
| 6.8 | Single mat | No matche | PreferredD | [M+H]+1 | 12030900 | 12127074 | 20827587 |
| 37.9 | No matche | No matche | PreferredD | [M+H]+1 | 14299234 | 12408868 | 13391684 |
| 61.9 | No matche | No matche | PreferredD | [M+H]+1 | 9370039 | 13489003 | 13756513 |
| 43.3 | No matche | No matche | PreferredD | [M+NH4]+ | 19263127 | 17951356 | 17380903 |
| 44.4 | No matche | No matche | PreferredD | [M+H]+1 | 4834087 | 8142751 | 7536217 |
| 41.6 | No matche | No matche | PreferredD | [M+H]+1 | 45788309 | 44889715 | 41440329 |
| 57.2 | No matche | No matche | PreferredD | [M+H]+1 | 25320813 | 25507871 | 17019893 |
| 59.4 | No matche | No matche | PreferredD | [M+H]+1 | 30342568 | 27354075 | 29045069 |
| 8 | No matche | No matche | PreferredD | [M+H]+1 | 17927798 | 18490679 | 15802137 |
| 39.3 | No matche | No matche | PreferredD | [M+H]+1 | 6202427 | 6949812 | 6784103 |
| 57.7 | No matche | No matche | PreferredD | [M+H]+1 | 18435158 | 24904955 | 19727528 |
| 46.9 | No matche | No matche | PreferredD | [M+H]+1 | 37089032 | 35553381 | 35448468 |
| 8.6 | No matche | No matche | PreferredD | [M+H]+1 | 5732566 | 5167336 | 4807783 |
| 58.7 | No matche | No matche | PreferredD | [M+H]+1 | 8086663 | 6544672 | 5679995 |
| 57.7 | No matche | No matche | PreferredD | [M+H]+1 | 674794.7 | 426262.7 | 1267613 |
| 56.5 | No matche | No matche | PreferredD | [M+H]+1 | 22490415 | 23624848 | 22315694 |
| 66.3 | No matche | No matche | PreferredD | [M+H]+1 | 12135962 | 16318076 | 16112498 |
| 65.6 | No matche | No matche | PreferredD | [M+H]+1 | 10142932 | 9210315 | 9561659 |
| 59.1 | No matche | No matche | OtherDDA | [M+H]+1 | 13228806 | 11000190 | 15109990 |
| 57.2 | No matche | No matche | PreferredD | [M+H]+1 | 9991007 | 13043777 | 10704260 |
| 6.9 | No matche | Multiple m | PreferredD | [M+H]+1 | 9358386 | 9426659 | 8462358 |
| 6.6 | Multiple m | Single mat | PreferredD | [M+H]+1 | 15783794 | 16667558 | 11249495 |
| 46 | Single mat | No matche | PreferredD | [M+H]+1 | 651133.9 | 922174.7 | 892970.3 |
| 59.9 | No matche | No matche | PreferredD | [M+H]+1 | 11740797 | 12264239 | 12337599 |
| 64.2 | No matche | No matche | PreferredD | [M+H]+1 | 15279224 | 13249165 | 11899657 |
| 9 | No matche | No matche | PreferredD | [M+H]+1 | 8502880 | 9155149 | 9043370 |
| 45.4 | No matche | No matche | PreferredD | [M+H]+1 | 6987969 | 6507737 | 6251685 |
| 57 | No matche | No matche | PreferredD | [M+H]+1 | 4128172 | 2683640 | 3655958 |
| 57.9 | Single mat | No matche | PreferredD | [M+H-NH3 | 4552527 | 4760243 | 5225321 |
| 61.6 | No matche | No matche | PreferredD | [M+H]+1 | 41636883 | 26969132 | 17653448 |
| 60.7 | No matche | No matche | PreferredD | [M+H]+1 | 10818936 | 10586911 | 12283383 |
| 8.9 | No matche | No matche | PreferredD | [M+H]+1 | 10965715 | 10886037 | 11036678 |
| 63.4 | No matche | Multiple m | PreferredD | [M+H]+1 | 17199387 | 18882411 | 18243345 |
| 63.3 | No matche | No matche | PreferredD | [M+H]+1 | 6778711 | 6960344 | 9097527 |
| 60.6 | No matche | No matche | PreferredD | [M+H]+1 | 38290061 | 45678367 | 27489016 |
| 57.9 | No matche | Multiple m | PreferredD | [M+H]+1 | 16772693 | 1.56E+07 | 17513735 |
| 56.5 | No matche | No matche | PreferredD | [M+H]+1 | 7428487 | 8169281 | 8606569 |
| 43 | No matche | No matche | PreferredD | [M+H]+1 | 2846685 | 2494848 | 2090994 |
| 62.7 | No matche | No matche | PreferredD | [M+H]+1 | 11303638 | 17798693 | 12498318 |
| 8.8 | No matche | No matche | PreferredD | [M+H]+1 | 24471261 | 22740135 | 22218606 |
| 44.8 | No matche | Single mat | PreferredD | [M+H]+1 | 1189736 | 2656884 | 3276315 |
| 6.5 | No matche | No matche | PreferredD | [M+H]+1 | 84379.87 | 62923.88 | 71682.11 |
| 38 | No matche | No matche | PreferredD | [M+H]+1 | 8741395 | 10877987 | 11541497 |
| 61.5 | No matche | No matche | PreferredD | [M+H]+1 | 806321.1 | 2044184 | 5203457 |
| 9.4 | No matche | No matche | PreferredD | [M+H]+1 | 4244691 | 38090534 | 24658626 |
| 65.3 | No matche | No matche | PreferredD | [M+H]+1 | 11334204 | 11106896 | 13701765 |
| 42.4 | No matche | No matche | PreferredD | [M+H]+1 | 5149102 | 4511029 | 3956718 |

| | | | | | | | |
|---|---|---|---|---|---|---|---|
| 57.7 | No matche | No matche | PreferredD | [M+H]+1 | 10830492 | 13411433 | 15703787 |
| 43.3 | No matche | No matche | PreferredD | [M+H-NH3 | 999309.8 | 338542.2 | 193825.3 |
| 7 | No matche | No matche | PreferredD | [M+H]+1 | 12477974 | 11611330 | 9563147 |
| 7 | No matche | No matche | PreferredD | [M+H]+1 | 12333559 | 13797908 | 13487464 |
| 7.8 | No matche | No matche | PreferredD | [M+H]+1 | 6712764 | 5943623 | 6725925 |
| 38 | No matche | No matche | PreferredD | [M+H]+1 | 13481642 | 12952799 | 10526015 |
| 61.5 | No matche | No matche | PreferredD | [M+H]+1 | 3945603 | 1348547 | 2570895 |
| 7.9 | No matche | No matche | PreferredD | [M+H]+1 | 14537744 | 12357727 | 12927375 |
| 58.3 | No matche | No matche | PreferredD | [M+H]+1 | 16063609 | 16212445 | 15927407 |
| 56.3 | No matche | No matche | OtherDDA | [M+2H]+2 | 1280217 | 1110279 | 1070413 |
| 38.1 | No matche | No matche | PreferredD | [M+H]+1 | 16134333 | 14940110 | 13810717 |
| 64.7 | No matche | No matche | PreferredD | [M+ACN+H | 25098702 | 15009647 | 17420450 |
| 60.5 | No matche | No matche | PreferredD | [M+H]+1 | 209403.4 | 211390.5 | 254806.9 |
| 40.9 | No matche | No matche | PreferredD | [M+H]+1 | 4799800 | 4225236 | 5575289 |
| 42 | No matche | No matche | OtherDDA | [M+2H]+2 | 6647487 | 6244005 | 6490049 |
| 62.4 | No matche | No matche | PreferredD | [M+H]+1 | 13812935 | 14359909 | 13548127 |
| 46.7 | No matche | No matche | PreferredD | [M+H]+1 | 7045514 | 7197269 | 6825626 |
| 41.3 | Single mat | No matche | PreferredD | [M+H]+1 | 14774575 | 14948145 | 15076254 |
| 8.5 | Single mat | No matche | PreferredD | [M+H]+1 | 14116173 | 14788736 | 14333378 |
| 9.6 | No matche | Single mat | PreferredD | [M+H]+1 | 512751.1 | 268391.8 | 508602.3 |
| 61.9 | No matche | No matche | PreferredD | [M+H]+1 | 3215397 | 3697069 | 3677190 |
| 59.7 | Single mat | No matche | PreferredD | [M+H]+1 | 14092770 | 14756200 | 12334276 |
| 40.1 | No matche | No matche | PreferredD | [M+NH4]+ | 8670072 | 5527656 | 6453516 |
| 8.9 | No matche | No matche | PreferredD | [M+2H]+2 | 13899267 | 13047604 | 13481296 |
| 59.3 | Single mat | No matche | PreferredD | [M+H]+1 | 8463894 | 8412313 | 9204006 |
| 40 | No matche | No matche | PreferredD | [M+H]+1 | 13850900 | 10115551 | 2488058 |
| 43.8 | No matche | No matche | PreferredD | [M+NH4]+ | 12050689 | 11839991 | 10585335 |
| 8.3 | No matche | No matche | PreferredD | [M+H]+1 | 1548530 | 1664977 | 1485198 |
| 43.3 | No matche | No matche | PreferredD | [M+H]+1 | 19263127 | 17951356 | 17380903 |
| 43.6 | No matche | No matche | PreferredD | [M+H]+1 | 2155799 | 2345477 | 1774914 |
| 62.6 | No matche | No matche | PreferredD | [M+H]+1 | 4579275 | 5843601 | 3828087 |
| 8.2 | No matche | No matche | PreferredD | [M+H]+1 | 9621220 | 8991128 | 9997200 |
| 9 | No matche | No matche | PreferredD | [M+H]+1 | 15575347 | 14891445 | 15255965 |
| 37.7 | No matche | No matche | PreferredD | [M+H]+1 | 33659165 | 32711055 | 31106956 |
| 41.7 | No matche | No matche | PreferredD | [M+H]+1 | 16219406 | 15264718 | 27485258 |
| 61.2 | No matche | No matche | PreferredD | [M+H]+1 | 12610640 | 8042792 | 9687703 |
| 64 | No matche | No matche | PreferredD | [M+H]+1 | 7714089 | 7181132 | 7143347 |
| 60.5 | No matche | No matche | PreferredD | [M+H]+1 | 1800634 | 253690.8 | 406758 |
| 55.9 | Single mat | No matche | PreferredD | [M+H]+1 | 13441361 | 13568731 | 13603074 |
| 60.7 | No matche | No matche | PreferredD | [M+H]+1 | 6973288 | 7237787 | 6646161 |
| 44.7 | No matche | No matche | PreferredD | [M+H]+1 | 9848147 | 9046028 | 9932042 |
| 9.5 | No matche | No matche | PreferredD | [M+H]+1 | 6468541 | 6972968 | 6923315 |
| 65.1 | No matche | No matche | PreferredD | [M+H]+1 | 12270278 | 12601929 | 12534534 |
| 40.9 | No matche | No matche | PreferredD | [M+H]+1 | 9239164 | 10651834 | 6328762 |
| 45 | No matche | No matche | PreferredD | [M+H]+1 | 10242605 | 8535642 | 8347607 |
| 47.2 | No matche | No matche | PreferredD | [M+H-NH3 | 826145.7 | 153786.3 | 239158 |
| 57 | No matche | No matche | PreferredD | [M+H]+1 | 9012487 | 9313733 | 10881085 |
| 40.3 | No matche | No matche | PreferredD | [M+H]+1 | 2831942 | 2747839 | 2583907 |
| 7.1 | No matche | No matche | PreferredD | [M+H]+1 | 5552727 | 5858564 | 6287800 |
| 43.9 | No matche | No matche | PreferredD | [M+H]+1 | 1955928 | 2720206 | 601029.6 |
| 42.8 | No matche | No matche | PreferredD | [M+H]+1 | 7690946 | 8835657 | 9334522 |

| | | | | | | | |
|---|---|---|---|---|---|---|---|
| 43.2 | No matche | No matche | PreferredD | [M+H]+1 | 6456941 | 6252865 | 5900841 |
| 43.8 | Single mat | No matche | PreferredD | [M+H]+1 | 13113715 | 12966481 | 12658649 |
| 56.6 | No matche | No matche | PreferredD | [M+NH4]+ | 18915931 | 23004477 | 22809031 |
| 58.5 | No matche | No matche | PreferredD | [M+H]+1 | 11849285 | 11091474 | 11281237 |
| 57.1 | No matche | Single mat | PreferredD | [M+H]+1 | 22032317 | 7460609 | 21706074 |
| 58.8 | No matche | No matche | PreferredD | [M+H]+1 | 6263697 | 5547434 | 4765571 |
| 38.9 | No matche | No matche | PreferredD | [M+H]+1 | 9953303 | 12768799 | 14898324 |
| 56.3 | No matche | No matche | PreferredD | [M+H]+1 | 6800398 | 6346824 | 6218957 |
| 60.9 | No matche | No matche | PreferredD | [M+H]+1 | 10471229 | 10008487 | 10381719 |
| 58 | No matche | No matche | PreferredD | [M+H]+1 | 250821 | 233650.3 | 237560.6 |
| 57.7 | No matche | No matche | PreferredD | [M+H]+1 | 6143144 | 6491763 | 6492044 |
| 64 | No matche | No matche | PreferredD | [M+H]+1 | 1315969 | 1958859 | 2171314 |
| 38.4 | No matche | No matche | PreferredD | [M+H]+1 | 11784588 | 13761185 | 12817383 |
| 62.1 | No matche | No matche | PreferredD | [M+H]+1 | 14774214 | 29070752 | 34470028 |
| 43.8 | No matche | No matche | PreferredD | [M+H]+1 | 15983600 | 16243330 | 14968116 |
| 62.1 | No matche | No matche | PreferredD | [M+H]+1 | 6844329 | 7328302 | 2606224 |
| 42.2 | No matche | No matche | PreferredD | [M+H]+1 | 7025741 | 5766087 | 6898814 |
| 59 | No matche | No matche | PreferredD | [M+H]+1 | 24426284 | 23486428 | 20730866 |
| 59.9 | No matche | No matche | PreferredD | [M+H]+1 | 4664428 | 4416772 | 4218977 |
| 57.4 | No matche | No matche | PreferredD | [M+H]+1 | 9972782 | 9234020 | 11434793 |
| 58.8 | No matche | No matche | PreferredD | [M+H]+1 | 522620 | 490172.4 | 1741120 |
| 8.7 | No matche | No matche | PreferredD | [M+H]+1 | 4459698 | 4864734 | 4201105 |
| 56.9 | No matche | No matche | PreferredD | [M+H]+1 | 17475169 | 15017169 | 18513917 |
| 57.4 | No matche | No matche | PreferredD | [M+H]+1 | 13907365 | 5543677 | 5404736 |
| 42.6 | Multiple m | No matche | PreferredD | [M+H]+1 | 1349231 | 1040818 | 4811328 |
| 6.7 | No matche | Multiple m | PreferredD | [M+H]+1 | 5956619 | 5736406 | 6839376 |
| 7.1 | No matche | No matche | PreferredD | [M+H]+1 | 5407483 | 6023107 | 6342178 |
| 61.8 | No matche | No matche | PreferredD | [M+H]+1 | 6641404 | 6643938 | 29870088 |
| 56.6 | Single mat | No matche | PreferredD | [M+H]+1 | 386760.8 | 1692659 | 1777245 |
| 6.6 | Single mat | No matche | PreferredD | [M+H]+1 | 4484524 | 4565694 | 5653733 |
| 62.9 | Multiple m | Single mat | PreferredD | [M+H+MeO | 6730065 | 6609699 | 6946563 |
| 40.4 | No matche | No matche | PreferredD | [M+H]+1 | 6330324 | 6011159 | 8183719 |
| 64.7 | No matche | Single mat | PreferredD | [M+H]+1 | 7234494 | 7985514 | 8367260 |
| 60.7 | No matche | No matche | PreferredD | [M+H]+1 | 6798634 | 7932846 | 7395320 |
| 46 | Single mat | No matche | PreferredD | [M+H]+1 | 11903419 | 12015847 | 12127833 |
| 45.1 | No matche | No matche | PreferredD | [M+H]+1 | 2717764 | 2391809 | 2529322 |
| 43.8 | No matche | No matche | PreferredD | [M+H]+1 | 5369153 | 5584152 | 1699032 |
| 62.2 | No matche | No matche | PreferredD | [M+H]+1 | 23782180 | 11962919 | 15766391 |
| 8 | No matche | No matche | PreferredD | [M+H]+1 | 4337446 | 3337400 | 3407284 |
| 60.4 | No matche | No matche | PreferredD | [M+H]+1 | 3986192 | 6174340 | 8078708 |
| 49.4 | No matche | No matche | PreferredD | [M+H]+1 | 6770883 | 6990350 | 6826860 |
| 7 | No matche | No matche | OtherDDA | [M+2H]+2 | 7450018 | 8540405 | 8578351 |
| 43.1 | No matche | Single mat | PreferredD | [M+H]+1 | 4044485 | 2733573 | 2476921 |
| 7.9 | No matche | Single mat | OtherDDA | [M+2H]+2 | 9661747 | 8408302 | 7471955 |
| 9.8 | Single mat | No matche | PreferredD | [M+H]+1 | 2498407 | 3252170 | 3720277 |
| 64.6 | No matche | No matche | PreferredD | [M+H]+1 | 7190583 | 6441022 | 6086450 |
| 9 | No matche | No matche | PreferredD | [M+H]+1 | 3635429 | 5486119 | 5716624 |
| 56.6 | No matche | No matche | PreferredD | [M+H]+1 | 18915931 | 23004477 | 22809031 |
| 43.3 | No matche | No matche | PreferredD | [M+H]+1 | 6296616 | 9235004 | 8388314 |
| 38.2 | No matche | No matche | PreferredD | [M+H]+1 | 6627758 | 6021918 | 6183521 |
| 63 | No matche | No matche | PreferredD | [M+H]+1 | 6362265 | 6846117 | 7507311 |

| | | | | | | | |
|---|---|---|---|---|---|---|---|
| 61.4 | No matche | No matche | PreferredD | [M+H]+1 | 9208859 | 6811076 | 7374224 |
| 45.6 | Multiple m | No matche | PreferredD | [M+H]+1 | 227420.1 | 215759.1 | 210310.9 |
| 43.9 | No matche | No matche | OtherDDA | [M+2H]+2 | 9155856 | 6840568 | 7433943 |
| 7.5 | Single mat | No matche | PreferredD | [M+H]+1 | 7212666 | 7729137 | 7360980 |
| 41.5 | Single mat | No matche | PreferredD | [M+H]+1 | 215196.9 | 332134.1 | 762247.1 |
| 63.4 | No matche | No matche | PreferredD | [M+H]+1 | 9420046 | 10183737 | 12123771 |
| 40.8 | No matche | No matche | PreferredD | [M+H]+1 | 55230148 | 56055270 | 49560643 |
| 59.1 | No matche | No matche | PreferredD | [M+H]+1 | 11855152 | 12552848 | 12667202 |
| 8.8 | No matche | No matche | PreferredD | [M+H]+1 | 7240349 | 6865272 | 7327083 |
| 59.5 | No matche | No matche | PreferredD | [M+H]+1 | 10666974 | 8888228 | 8092345 |
| 61.5 | No matche | No matche | PreferredD | [M+H]+1 | 8557638 | 8122908 | 7456954 |
| 42.5 | No matche | No matche | OtherDDA | [M+2H]+2 | 5429327 | 3689736 | 4864305 |
| 7.3 | No matche | No matche | PreferredD | [M+H]+1 | 773667.5 | 845186.1 | 583085.7 |
| 61.4 | No matche | No matche | PreferredD | [M+H]+1 | 4004940 | 4076851 | 4198552 |
| 40.4 | No matche | No matche | PreferredD | [M+H]+1 | 982510.5 | 1830913 | 1983157 |
| 44.9 | Single mat | No matche | PreferredD | [M+H]+1 | 14039641 | 13851166 | 14428850 |
| 7.9 | Single mat | No matche | PreferredD | [M+H]+1 | 3856977 | 4749106 | 3989890 |
| 62 | Multiple m | Single mat | PreferredD | [M+H]+1 | 6546881 | 6063445 | 6334545 |
| 6.9 | Single mat | No matche | PreferredD | [M+H]+1 | 4740890 | 4622463 | 4806319 |
| 62.2 | Multiple m | No matche | PreferredD | [M+H]+1 | 21797060 | 19681155 | 13686792 |
| 7 | No matche | No matche | PreferredD | [M+H]+1 | 4788628 | 4504189 | 2758069 |
| 41.6 | No matche | No matche | PreferredD | [M+H]+1 | 11353797 | 12356177 | 11035694 |
| 57.8 | No matche | No matche | PreferredD | [M+H]+1 | 3693965 | 3293775 | 3424672 |
| 7.2 | No matche | No matche | PreferredD | [M+H]+1 | 3731277 | 3041239 | 3431480 |
| 42.3 | No matche | No matche | PreferredD | [M+H]+1 | 724041.9 | 4962109 | 1007873 |
| 60.9 | No matche | No matche | PreferredD | [M+H]+1 | 3867128 | 4641956 | 4821201 |
| 7.7 | No matche | No matche | PreferredD | [M+H]+1 | 23291995 | 20053896 | 23756700 |
| 7.1 | No matche | No matche | PreferredD | [M+H]+1 | 4108969 | 2823104 | 3898038 |
| 6.9 | No matche | No matche | PreferredD | [M+H]+1 | 6734167 | 7773798 | 6798524 |
| 39.1 | No matche | No matche | PreferredD | [M+H]+1 | 2807251 | 3012419 | 2502131 |
| 46.6 | No matche | No matche | PreferredD | [M+H]+1 | 7489620 | 7354140 | 7357554 |
| 73.5 | No matche | No matche | PreferredD | [M+H]+1 | 52623901 | 48182812 | 51465123 |
| 67 | No matche | No matche | PreferredD | [M+H]+1 | 34491435 | 12930883 | 12772687 |
| 67 | No matche | No matche | PreferredD | [M+H]+1 | 11204318 | 8559369 | 10544960 |
| 65.1 | No matche | No matche | PreferredD | [M+H]+1 | 20602587 | 20558171 | 12641363 |
| 64.8 | No matche | No matche | PreferredD | [M+H]+1 | 9582887 | 12787276 | 37976777 |
| 62.8 | No matche | No matche | PreferredD | [M+H]+1 | 39938428 | 11732283 | 40696981 |
| 69 | No matche | No matche | PreferredD | [M+H]+1 | 48036090 | 50658730 | 52775689 |
| 61.4 | No matche | No matche | PreferredD | [M+H]+1 | 7165769 | 13050239 | 5888260 |
| 46 | No matche | No matche | PreferredD | [M+H]+1 | 2678833 | 2778808 | 2603825 |
| 47.9 | No matche | No matche | PreferredD | [M+H]+1 | 36648449 | 33419886 | 29194957 |
| 65.2 | No matche | No matche | PreferredD | [M+H]+1 | 31142334 | 31408404 | 4949056 |
| 56.4 | No matche | No matche | PreferredD | [M+H]+1 | 65590.27 | 173335 | 85117.95 |
| 63.1 | No matche | No matche | PreferredD | [M+H]+1 | 444267.6 | 143068.7 | 451009.7 |
| 58.7 | No matche | No matche | PreferredD | [M+H-H2O | 1.6E+09 | 1.57E+09 | 1.51E+09 |
| 59.4 | No matche | No matche | PreferredD | [M+H]+1 | 2.09E+08 | 2.04E+08 | 2E+08 |
| 57.4 | No matche | No matche | PreferredD | [M+H]+1 | 25201580 | 24063187 | 24474038 |
| 57.4 | No matche | No matche | PreferredD | [M+H]+1 | 1812496 | 2799415 | 2259010 |
| 58.4 | No matche | No matche | PreferredD | [M+H]+1 | 17767379 | 15626618 | 15679315 |
| 61.7 | No matche | Single mat | PreferredD | [M+H]+1 | 37761250 | 60730441 | 30667906 |
| 67 | No matche | Single mat | PreferredD | [M+H]+1 | 9502904 | 9090325 | 9255559 |

| | | | | | | | |
|---|---|---|---|---|---|---|---|
| 41 | No matche | No matche | PreferredD | [M+H]+1 | 19102159 | 19210436 | 17224079 |
| 59.1 | No matche | No matche | PreferredD | [M+H-H2O | 4.28E+08 | 4.11E+08 | 4.06E+08 |
| 62.1 | No matche | No matche | PreferredD | [M+H]+1 | 71304973 | 74852036 | 76778979 |
| 63.8 | No matche | No matche | PreferredD | [M+H]+1 | 13667146 | 22674226 | 7155289 |
| 6.9 | Single mat | No matche | PreferredD | [M+H]+1 | 9176934 | 8405267 | 8260080 |
| 42.6 | No matche | No matche | PreferredD | [M+H]+1 | 9640715 | 10717806 | 9519408 |
| 44.3 | No matche | No matche | PreferredD | [M+H]+1 | 26870066 | 23244373 | 25324247 |
| 8.5 | No matche | No matche | PreferredD | [M+H]+1 | 262760.9 | 253571.2 | 258763.6 |
| 41.5 | No matche | No matche | PreferredD | [M+H]+1 | 16637649 | 16300069 | 16677219 |
| 57 | No matche | No matche | PreferredD | [M+H]+1 | 984049.9 | 534199.2 | 805010.8 |
| 42 | No matche | No matche | PreferredD | [M+H]+1 | 7420258 | 8165325 | 7106018 |
| 54.6 | No matche | No matche | PreferredD | [M+H]+1 | 63637049 | 59316225 | 58112684 |
| 7.2 | No matche | No matche | PreferredD | [M+H]+1 | 4141967 | 4226199 | 3997481 |
| 6.6 | No matche | No matche | PreferredD | [M+H]+1 | 24508517 | 26680218 | 25670424 |
| 64.2 | No matche | No matche | PreferredD | [M+H]+1 | 31539708 | 25384887 | 38935467 |
| 65.8 | No matche | No matche | PreferredD | [M+H]+1 | 12786501 | 15186480 | 12541992 |
| 57.4 | No matche | Single mat | PreferredD | [M+H]+1 | 15751368 | 19580821 | 18490313 |
| 58.6 | No matche | No matche | PreferredD | [M+H]+1 | 20075887 | 20410615 | 19844720 |
| 62 | No matche | No matche | PreferredD | [M+H]+1 | 22181309 | 21792495 | 20748504 |
| 45.2 | No matche | No matche | PreferredD | [M+H]+1 | 18630760 | 19003034 | 18094823 |
| 44.5 | No matche | No matche | PreferredD | [M+H]+1 | 16182323 | 17402607 | 16334154 |
| 56.8 | No matche | No matche | PreferredD | [M+H]+1 | 20648512 | 19381957 | 19622590 |
| 59.9 | No matche | No matche | PreferredD | [M+H]+1 | 17452141 | 17463344 | 24116413 |
| 60.4 | No matche | No matche | PreferredD | [M+H]+1 | 10230642 | 11481321 | 9817690 |
| 63.1 | No matche | No matche | PreferredD | [M+H]+1 | 23544341 | 23016214 | 23442282 |
| 71.3 | No matche | No matche | PreferredD | [M+H]+1 | 32386381 | 35226028 | 33699678 |
| 58.1 | Single mat | Single mat | PreferredD | [M+H-NH3 | 11104042 | 10125325 | 10780934 |
| 62.4 | No matche | No matche | PreferredD | [M+H]+1 | 27831739 | 30887923 | 30918923 |
| 62.5 | No matche | Single mat | PreferredD | [M+H-H2O | 16027581 | 13208684 | 13608543 |
| 56.8 | No matche | No matche | PreferredD | [M+H]+1 | 739170.4 | 2176573 | 2065526 |
| 60.1 | No matche | No matche | PreferredD | [M+H]+1 | 269959.5 | 245519.1 | 242432.8 |
| 64.8 | No matche | No matche | PreferredD | [M+H]+1 | 442879.2 | 436830.3 | 489889.1 |
| 6.6 | No matche | No matche | PreferredD | [M+H]+1 | 2206635 | 975054.7 | 1141960 |
| 7.3 | No matche | No matche | PreferredD | [M+H]+1 | 10476879 | 9063410 | 9056300 |
| 56.6 | No matche | No matche | PreferredD | [M+H]+1 | 36087025 | 40442271 | 40401027 |
| 59.1 | No matche | No matche | PreferredD | [M+H]+1 | 13742302 | 37924603 | 37448952 |
| 9.3 | No matche | No matche | PreferredD | [M+H]+1 | 17965828 | 17229885 | 16914231 |
| 59.3 | No matche | No matche | PreferredD | [M+H]+1 | 8327887 | 8693550 | 7244936 |
| 45.2 | No matche | No matche | OtherDDA | [M+2H]+2 | 14929034 | 20820992 | 19706114 |
| 42.6 | No matche | No matche | PreferredD | [M+H]+1 | 17190066 | 17130482 | 16935070 |
| 57.9 | No matche | No matche | PreferredD | [M+H]+1 | 52779736 | 33634833 | 55620380 |
| 63.8 | No matche | No matche | PreferredD | [M+H]+1 | 4289160 | 1135683 | 3883283 |
| 56.6 | No matche | No matche | PreferredD | [M+H]+1 | 229100 | 817926 | 438576.9 |
| 63.1 | No matche | No matche | PreferredD | [M+H]+1 | 11070686 | 10526888 | 11042575 |
| 67.6 | No matche | Multiple m | PreferredD | [M+H]+1 | 1.17E+08 | 1.06E+08 | 1.06E+08 |
| 72.1 | No matche | Multiple m | PreferredD | [M+H]+1 | 1.14E+08 | 1.09E+08 | 1.15E+08 |
| 73.5 | No matche | Multiple m | PreferredD | [M+H+Me( | 3744118 | 2030104 | 3864195 |
| 6.6 | No matche | No matche | PreferredD | [M+H]+1 | 9482727 | 7833607 | 7871170 |
| 58.6 | No matche | No matche | PreferredD | [M+H]+1 | 208474 | 1108019 | 354857.6 |
| 58.3 | No matche | No matche | PreferredD | [M+H]+1 | 34327337 | 37063462 | 37595653 |
| 8.5 | No matche | No matche | PreferredD | [M+H]+1 | 3.19E+08 | 3.13E+08 | 2.99E+08 |

| | | | | | | | |
|---|---|---|---|---|---|---|---|
| 7.4 | No matche | No matche | PreferredD | [M+H]+1 | 9014022 | 8728439 | 9093623 |
| 8.9 | No matche | No matche | PreferredD | [M+H]+1 | 8483953 | 7392849 | 8237524 |
| 38.1 | No matche | No matche | PreferredD | [M+H]+1 | 2862293 | 3584877 | 3295415 |
| 43.2 | No matche | No matche | PreferredD | [M+H]+1 | 8846493 | 7497120 | 7207081 |
| 51.3 | No matche | No matche | PreferredD | [M+H]+1 | 21414280 | 21638234 | 21897946 |
| 60.3 | No matche | No matche | PreferredD | [M+H]+1 | 9107958 | 7231472 | 8096976 |
| 41.1 | No matche | No matche | PreferredD | [M+H]+1 | 1.47E+08 | 1.48E+08 | 1.5E+08 |
| 8.4 | No matche | No matche | PreferredD | [M+H]+1 | 2705276 | 2915130 | 2222427 |
| 40.8 | No matche | No matche | PreferredD | [M+H]+1 | 36368924 | 37733232 | 31747358 |
| 58 | No matche | No matche | PreferredD | [M+H]+1 | 5811473 | 5693021 | 5164199 |
| 47.9 | No matche | Single mat | PreferredD | [M+H]+1 | 1.33E+08 | 1.37E+08 | 1.18E+08 |
| 50.7 | No matche | Single mat | PreferredD | [M+H]+1 | 91464778 | 85953468 | 75771712 |
| 47.9 | No matche | Single mat | PreferredD | [M+H]+1 | 1.06E+08 | 1.16E+08 | 1.01E+08 |
| 55.1 | No matche | Single mat | PreferredD | [M+H]+1 | 10780962 | 13414211 | 12309530 |
| 52.9 | No matche | Single mat | PreferredD | [M+H]+1 | 11812868 | 9844877 | 9874860 |
| 42 | No matche | No matche | PreferredD | [M+H]+1 | 13943242 | 13298725 | 13388830 |
| 8.6 | No matche | No matche | PreferredD | [M+H]+1 | 7187965 | 6944099 | 6735306 |
| 42.8 | No matche | No matche | PreferredD | [M+H]+1 | 10399105 | 12320361 | 11330467 |
| 67.3 | No matche | Single mat | PreferredD | [M+H]+1 | 10954306 | 17830470 | 35327905 |
| 44.9 | No matche | No matche | PreferredD | [M+H]+1 | 370951.4 | 230459 | 529236.5 |
| 42.3 | No matche | No matche | PreferredD | [M+H]+1 | 383004.7 | 530916.4 | 341846.7 |
| 8.4 | No matche | No matche | PreferredD | [M+H]+1 | 41479031 | 46304205 | 34742013 |
| 42.4 | No matche | No matche | PreferredD | [M+H]+1 | 4669193 | 4765813 | 2067371 |
| 39.1 | No matche | Single mat | PreferredD | [M+H]+1 | 12407604 | 11458570 | 14064372 |
| 9.7 | Single mat | No matche | PreferredD | [M+H]+1 | 1.83E+08 | 1.76E+08 | 1.72E+08 |
| 37.8 | No matche | No matche | PreferredD | [M+H]+1 | 7632467 | 6932411 | 6560508 |
| 63.2 | No matche | No matche | PreferredD | [M+H]+1 | 5729497 | 7736935 | 6988933 |
| 8.4 | No matche | No matche | PreferredD | [M+H]+1 | 10808790 | 10736876 | 10639808 |
| 9.7 | No matche | No matche | PreferredD | [M+H]+1 | 15844604 | 14947830 | 14950594 |
| 62.1 | No matche | No matche | PreferredD | [M+H]+1 | 9716282 | 9701859 | 10313047 |
| 6.6 | No matche | No matche | PreferredD | [M+NH4]+ | 2.1E+08 | 1.98E+08 | 2.05E+08 |
| 56.6 | No matche | Multiple m | PreferredD | [M+H]+1 | 4423154 | 2437027 | 1767236 |
| 56.6 | No matche | No matche | PreferredD | [M+H]+1 | 5646606 | 1561843 | 1972133 |
| 7.9 | No matche | No matche | PreferredD | [M+H]+1 | 18918394 | 17711198 | 19341080 |
| 53.6 | No matche | No matche | PreferredD | [M+H]+1 | 14997851 | 10528870 | 14376761 |
| 59.9 | No matche | No matche | PreferredD | [M+H]+1 | 72087082 | 73970517 | 75058439 |
| 56.8 | No matche | No matche | PreferredD | [M+H]+1 | 3744118 | 2030104 | 4007343 |
| 58.4 | No matche | No matche | PreferredD | [M+H]+1 | 43672895 | 43512105 | 43472050 |
| 59 | No matche | No matche | PreferredD | [M+H+Me( | 17866487 | 16414488 | 18949402 |
| 57.5 | No matche | No matche | PreferredD | [M+H]+1 | 13187129 | 13248062 | 11498836 |
| 42.2 | No matche | No matche | PreferredD | [M+H]+1 | 8052669 | 4230364 | 4787086 |
| 9.6 | No matche | No matche | PreferredD | [M+H]+1 | 10256887 | 9457245 | 9170492 |
| 60.7 | No matche | No matche | PreferredD | [M+H-H2O | 9186789 | 9722340 | 9519999 |
| 50.6 | No matche | No matche | PreferredD | [M+H]+1 | 13727270 | 13182675 | 14436898 |
| 44 | No matche | No matche | PreferredD | [M+H]+1 | 7703914 | 8737321 | 9792221 |
| 63.4 | No matche | No matche | PreferredD | [M+H]+1 | 6086769 | 5724853 | 6084842 |
| 7.5 | No matche | Single mat | PreferredD | [M+H]+1 | 14509458 | 15562035 | 16633132 |
| 45.1 | No matche | No matche | PreferredD | [M+H]+1 | 3061705 | 2934835 | 2263646 |
| 61.7 | Single mat | Single mat | PreferredD | [M+H]+1 | 10004141 | 9414113 | 9432976 |
| 9.7 | No matche | No matche | PreferredD | [M+H]+1 | 28524093 | 24897786 | 28444953 |
| 40.9 | No matche | No matche | PreferredD | [M+H]+1 | 4025974 | 4805682 | 4950675 |

| | | | | | | | |
|---|---|---|---|---|---|---|---|
| 58.8 | Single mat | No matche | PreferredD | [M+H]+1 | 11104042 | 10125325 | 10780934 |
| 64.2 | Single mat | Single mat | PreferredD | [M+H]+1 | 11573607 | 11552557 | 11662178 |
| 7.2 | No matche | No matche | PreferredD | [2M+K]+1 | 32087917 | 32111990 | 28732129 |
| 9.4 | No matche | No matche | PreferredD | [M+H]+1 | 75365878 | 79964728 | 82183333 |
| 39.1 | No matche | No matche | PreferredD | [M+H]+1 | 8730203 | 8697664 | 8683833 |
| 60 | No matche | Single mat | PreferredD | [M+H]+1 | 22458366 | 12307519 | 12154070 |
| 64.3 | No matche | Single mat | PreferredD | [M+H]+1 | 23173708 | 16613841 | 31684510 |
| 39.6 | No matche | No matche | PreferredD | [M+H]+1 | 11235784 | 10931719 | 10870512 |
| 7.7 | No matche | No matche | PreferredD | [M+H]+1 | 11044870 | 12833022 | 11872948 |
| 50.1 | No matche | No matche | PreferredD | [M+H]+1 | 41305992 | 39635210 | 38746050 |
| 56.4 | Single mat | Single mat | PreferredD | [M+H]+1 | 2242045 | 1145676 | 1833700 |
| 60.5 | No matche | No matche | PreferredD | [M+H]+1 | 24945087 | 27466700 | 27912013 |
| 61.1 | No matche | No matche | PreferredD | [M+H]+1 | 18280599 | 42980296 | 13186896 |
| 57.6 | No matche | No matche | PreferredD | [M+H]+1 | 536442.5 | 384168.3 | 687991.9 |
| 68.2 | No matche | No matche | PreferredD | [M+H]+1 | 11174300 | 11268656 | 11397665 |
| 62.3 | No matche | No matche | PreferredD | [M+H]+1 | 2125264 | 5378202 | 5551348 |
| 44.7 | No matche | No matche | PreferredD | [M+H]+1 | 5951053 | 6163313 | 6140501 |
| 6.9 | No matche | No matche | PreferredD | [M+H]+1 | 7979884 | 7460067 | 8230142 |
| 8.4 | No matche | No matche | PreferredD | [M+H]+1 | 10186805 | 9119232 | 9471409 |
| 61.1 | No matche | No matche | PreferredD | [M+H]+1 | 6028462 | 5646893 | 7810964 |
| 61.5 | No matche | Multiple m | PreferredD | [M+H]+1 | 8873493 | 9116596 | 11775332 |
| 39.8 | No matche | No matche | PreferredD | [M+H]+1 | 5234256 | 6306044 | 5973249 |
| 40.8 | No matche | No matche | PreferredD | [M+H]+1 | 19488002 | 28678518 | 20786877 |
| 40.8 | No matche | No matche | PreferredD | [M+H]+1 | 7671645 | 8362993 | 14336493 |
| 39.5 | No matche | No matche | PreferredD | [M+H]+1 | 12214587 | 12475211 | 12692685 |
| 67.8 | No matche | Single mat | PreferredD | [M+H]+1 | 4351190 | 3539920 | 3234726 |
| 41.4 | No matche | Single mat | PreferredD | [M+H]+1 | 20961891 | 22495267 | 24654985 |
| 56.4 | No matche | No matche | PreferredD | [M+H]+1 | 285898.3 | 198485.2 | 125420.7 |
| 43.3 | No matche | No matche | PreferredD | [M+H]+1 | 6794095 | 6181820 | 6400680 |
| 41.3 | No matche | No matche | PreferredD | [M+H]+1 | 20376372 | 19484171 | 18506803 |
| 7.5 | No matche | No matche | PreferredD | [M+H]+1 | 12095253 | 12013234 | 11802366 |
| 65 | No matche | No matche | PreferredD | [M+H]+1 | 2857649 | 1271533 | 5938178 |
| 52.5 | No matche | No matche | PreferredD | [M+H]+1 | 7394314 | 6863584 | 6695968 |
| 64.5 | No matche | Multiple m | PreferredD | [M+H]+1 | 91487533 | 99083143 | 1.5E+08 |
| 64.5 | No matche | Multiple m | PreferredD | [M+H]+1 | 45937539 | 24422332 | 53177165 |
| 70.4 | No matche | Multiple m | PreferredD | [M+H]+1 | 37633805 | 48151167 | 43211107 |
| 66.2 | No matche | Multiple m | PreferredD | [M+H]+1 | 17942983 | 17142888 | 17573039 |
| 6.5 | No matche | No matche | PreferredD | [M+H+Me( | 8331484 | 10443126 | 10679692 |
| 66.5 | No matche | No matche | PreferredD | [M+H]+1 | 12608406 | 11629110 | 13071564 |
| 57.3 | No matche | No matche | PreferredD | [M+H]+1 | 16945021 | 28060848 | 18309941 |
| 65.1 | No matche | No matche | PreferredD | [M+H]+1 | 23794782 | 23007251 | 21604817 |
| 65.1 | No matche | No matche | PreferredD | [M+H]+1 | 10890095 | 11388733 | 9921115 |
| 59 | No matche | No matche | PreferredD | [M+H]+1 | 22561607 | 21657419 | 20501661 |
| 41.3 | No matche | No matche | PreferredD | [M+H]+1 | 9306622 | 8597351 | 7065267 |
| 38.8 | No matche | No matche | PreferredD | [M+H]+1 | 16350213 | 12546166 | 12378593 |
| 61.1 | No matche | No matche | PreferredD | [M+H]+1 | 19201074 | 17501384 | 17564734 |
| 59.3 | No matche | No matche | PreferredD | [M+H]+1 | 12231929 | 12025082 | 11926983 |
| 58.9 | No matche | No matche | PreferredD | [M+H]+1 | 18892632 | 19366837 | 20056549 |
| 60.1 | No matche | No matche | PreferredD | [M+H]+1 | 7267793 | 7356448 | 7545821 |
| 6.8 | No matche | No matche | PreferredD | [M+H]+1 | 15901583 | 16881978 | 15321897 |
| 7 | No matche | No matche | PreferredD | [M+H]+1 | 12127705 | 6926132 | 8526994 |

| | | | | | | | | |
|---|---|---|---|---|---|---|---|---|
| 46.8 | | No matche | No matche | PreferredD | [M+H]+1 | 25733654 | 24701636 | 25604357 |
| 8.2 | | No matche | No matche | PreferredD | [M+H]+1 | 43922073 | 44025718 | 44328964 |
| 41.4 | | No matche | No matche | PreferredD | [M+H]+1 | 5475670 | 5382449 | 4814359 |
| 7.7 | | No matche | No matche | PreferredD | [M+H]+1 | 30517901 | 30591462 | 27817890 |
| 7.5 | | No matche | No matche | PreferredD | [M+H]+1 | 45783932 | 43107382 | 44481566 |
| 57.4 | | No matche | No matche | PreferredD | [M+H]+1 | 19946004 | 25570838 | 17908379 |
| 8.4 | | No matche | No matche | PreferredD | [M+H]+1 | 62167781 | 66148391 | 65785826 |
| 63.6 | | No matche | No matche | PreferredD | [M+H]+1 | 6690138 | 6254601 | 6741037 |
| 46.4 | | No matche | No matche | PreferredD | [M+H]+1 | 7297400 | 8109385 | 7623285 |
| 43.9 | | No matche | No matche | PreferredD | [M+H]+1 | 2592481 | 2623527 | 2641440 |
| 42 | | No matche | No matche | PreferredD | [M+H]+1 | 4891094 | 2189224 | 4700421 |
| 8.4 | | No matche | No matche | PreferredD | [M+H]+1 | 8493055 | 8593054 | 8701973 |
| 40.8 | | No matche | No matche | PreferredD | [M+H]+1 | 5254055 | 5092003 | 4952035 |
| 63.6 | | No matche | No matche | PreferredD | [M+H]+1 | 9435154 | 8979421 | 8919274 |
| 63.5 | | No matche | No matche | PreferredD | [M+H]+1 | 14040141 | 14757279 | 12766918 |
| 9.6 | | Single mat | No matche | PreferredD | [M+H]+1 | 29096105 | 29591105 | 30053876 |
| 9.5 | | Single mat | No matche | PreferredD | [M+H]+1 | 25289119 | 28136482 | 29621034 |
| 78.8 | | Single mat | No matche | PreferredD | [M+H]+1 | 14225506 | 14593757 | 14451156 |
| 41.7 | | Multiple m | No matche | PreferredD | [M+H]+1 | 15639578 | 14668178 | 5671432 |
| 45.6 | | Multiple m | No matche | PreferredD | [M+H]+1 | 259788.4 | 357223.4 | 146383.5 |
| 8.1 | | No matche | Single mat | PreferredD | [M+H]+1 | 4383658 | 4126995 | 4831742 |
| 40 | | No matche | No matche | PreferredD | [M+H]+1 | 421404.5 | 255351.3 | 171661.6 |
| 57.9 | | No matche | No matche | PreferredD | [M+H]+1 | 1.99E+08 | 1.94E+08 | 1.98E+08 |
| 65 | | No matche | No matche | PreferredD | [M+H]+1 | 9582154 | 10136384 | 10814773 |
| 41.1 | | No matche | No matche | PreferredD | [M+H]+1 | 4348929 | 4672339 | 4795748 |
| 47.2 | | No matche | No matche | PreferredD | [M+H]+1 | 24212924 | 24376676 | 23659359 |
| 7.5 | | No matche | No matche | PreferredD | [M+H]+1 | 52892782 | 53533727 | 42903532 |
| 7 | | No matche | No matche | PreferredD | [M+H]+1 | 7194819 | 4335399 | 8072982 |
| 7.2 | | No matche | No matche | PreferredD | [M+H]+1 | 11307759 | 10945038 | 10758200 |
| 61.8 | | No matche | No matche | PreferredD | [M+H]+1 | 11843853 | 10937221 | 12515076 |
| 44.9 | | No matche | No matche | PreferredD | [M+H]+1 | 153915.9 | 148494.7 | 156611.4 |

| Area_A_PB | Area_D1 | Area_D2 | Area_D3 | Area_D_PB | Area_Pool_QC_1 | Area_Pool_QC_2 | Area_Pool_QC_3 | Area_S1 |
|---|---|---|---|---|---|---|---|---|
| 18163831 | 1.04E+09 | 1.08E+09 | 1.08E+09 | 13517965 | 7.49E+08 | 7.28E+08 | 7.55E+08 | 5.95E+08 |
| 4790095 | 92303555 | 87657332 | 96137436 | 7138368 | 1.45E+08 | 1.11E+08 | 51173976 | 60635053 |
| 4553780 | 49759190 | 45412638 | 48996169 | 4614440 | 78930821 | 57503900 | 62885501 | 32331102 |
| 13308945 | 1.2E+09 | 1.26E+09 | 1.19E+09 | 1933441 | 4.64E+08 | 4.29E+08 | 4.44E+08 | 7867664 |
| 89430.34 | 1121271 | 595191.6 | 478463 | 411965.7 | 68182524 | 62433679 | 63376548 | 136797.1 |
| 89465.58 | 175370.1 | 184151.2 | 168730.5 | 209701 | 24116846 | 22261820 | 23057241 | 146251.6 |
| 7046612 | 5.19E+08 | 4.36E+08 | 4.35E+08 | 6056288 | 3.56E+08 | 2.91E+08 | 3.67E+08 | 2.18E+08 |
| 238143.7 | 2505479 | 2674797 | 2525209 | 168317.3 | 3657046 | 3575272 | 3875374 | 1689482 |
| 127523.4 | 2444737 | 2375549 | 2710208 | 124251.2 | 2256504 | 2239973 | 2166983 | 495476.2 |
| 237998.4 | 7847205 | 7832553 | 8275782 | 151621.9 | 6623919 | 3576534 | 3744709 | 2744671 |
| 678221.5 | 8958210 | 11139748 | 10608299 | 176040.2 | 23744410 | 15916476 | 28060308 | 19789311 |
| 88781.71 | 1691991 | 861354.8 | 788624.8 | 107459.5 | 7451359 | 4911189 | 5347645 | 1980935 |
| 49844.79 | 9858146 | 9988993 | 10134925 | 56210.93 | 3689901 | 3172316 | 3236300 | 87058.39 |
| 90816.07 | 41474773 | 39294098 | 41037651 | 107843 | 31629212 | 28850431 | 30041378 | 16890185 |
| 131450.7 | 30239134 | 27515258 | 29071978 | 133989 | 21666384 | 19617893 | 19807912 | 10909847 |
| 89991.12 | 17140682 | 14525478 | 16295079 | 90195.2 | 11626809 | 10324827 | 9741524 | 4328468 |
| 17498333 | 4.38E+08 | 3.61E+08 | 3.4E+08 | 21329507 | 2.47E+08 | 2.28E+08 | 2.25E+08 | 1.13E+08 |
| 242115.6 | 33009224 | 33389705 | 31550182 | 428372.9 | 46057273 | 41051399 | 42660571 | 5607715 |
| 660632 | 34568813 | 54863012 | 14372131 | 1033609 | 55846242 | 48785036 | 50727970 | 47072510 |
| 96212.48 | 99977.86 | 100206.5 | 94779.12 | 96397.58 | 2102153 | 2052919 | 830858.5 | 94686.9 |
| 95090.59 | 8320270 | 9563158 | 8674777 | 100939.5 | 5142797 | 4399884 | 11393866 | 1347846 |
| 411877.1 | 4441035 | 2426011 | 3510488 | 1829712 | 1.26E+08 | 1.23E+08 | 1.37E+08 | 1912093 |
| 1093880 | 2402389 | 1161077 | 1973523 | 2773986 | 2.27E+08 | 2.04E+08 | 2.03E+08 | 261577.3 |
| 93022.62 | 1776401 | 3149793 | 4262145 | 250319.3 | 13440788 | 13780692 | 22603765 | 3793159 |
| 161367.8 | 6153373 | 4443761 | 2854891 | 203796 | 5533640 | 6862981 | 6274718 | 1260058 |
| 78887.24 | 4846974 | 1672652 | 4753878 | 123868.3 | 20376024 | 16611678 | 15383081 | 2282494 |
| 80868.15 | 26738363 | 24984572 | 25539985 | 332973 | 20717769 | 18676346 | 17666609 | 12896794 |
| 313457.7 | 7092812 | 19272474 | 3201457 | 367379 | 11790819 | 10665062 | 13308450 | 5640398 |
| 83444.27 | 423441.6 | 304787.4 | 558593.6 | 41980.55 | 3187222 | 3107523 | 3612689 | 187428.1 |
| 76380.38 | 93086.22 | 71608.14 | 69401.79 | 72396.61 | 14847239 | 15292528 | 13443286 | 1534925 |
| 82263.89 | 8387542 | 6183739 | 7133646 | 84760.98 | 5128256 | 6014232 | 5169836 | 831570.6 |
| 334723 | 27655888 | 26068234 | 30603406 | 795896.8 | 33320471 | 28580247 | 29949699 | 34238195 |
| 1586170 | 7961864 | 11979784 | 15997900 | 358251.4 | 14181377 | 13514117 | 14416999 | 10046878 |
| 775003.2 | 12045005 | 40567027 | 9145103 | 502606.8 | 12345222 | 47762514 | 10753698 | 18361838 |
| 124417.6 | 7042771 | 683010.7 | 7000008 | 152804.2 | 15548997 | 10811888 | 13489800 | 4675881 |
| 2232564 | 19825351 | 20150673 | 11909996 | 544890.9 | 20136595 | 19568070 | 13798915 | 11172590 |
| 737404.1 | 4119941 | 4415098 | 4475950 | 205249.8 | 6692523 | 6439010 | 3843161 | 4553969 |
| 349449.1 | 8147373 | 8754712 | 8149424 | 933559.6 | 10440056 | 18966819 | 4213714 | 10198087 |
| 344725.3 | 11992705 | 24386378 | 24801892 | 1197863 | 6971269 | 8843491 | 9720569 | 3860246 |
| 407335.5 | 6717764 | 7020209 | 7980157 | 474154 | 9718341 | 6639062 | 8500351 | 5262944 |

| | | | | | | | | |
|---|---|---|---|---|---|---|---|---|
| 391014.4 | 5697938 | 13377861 | 9384919 | 171048.9 | 8648453 | 10137936 | 10748757 | 4296495 |
| 192824.4 | 6207952 | 3100996 | 6437877 | 675495.7 | 12744366 | 12516307 | 14658387 | 5249220 |
| 863264.3 | 21012412 | 22856989 | 27406188 | 423187.5 | 21283840 | 21670315 | 21824304 | 8228776 |
| 2396368 | 26795746 | 26739650 | 27379385 | 1136479 | 14929512 | 28335383 | 57706418 | 5694228 |
| 812718.9 | 26022267 | 20714748 | 20929372 | 587223 | 17306461 | 14047412 | 17314547 | 24696929 |
| 5619591 | 6275012 | 2805279 | 1925663 | 12703766 | 5.31E+08 | 5.16E+08 | 5.2E+08 | 2529943 |
| 90139.24 | 267030.6 | 238496.3 | 165500.9 | 1011370 | 45436789 | 42349254 | 42595457 | 141235.1 |
| 157296.3 | 5460734 | 901517.3 | 1621649 | 380688.8 | 6959944 | 5289558 | 6890642 | 447300.7 |
| 107703 | 728626.4 | 1709535 | 1317686 | 148071.1 | 12843483 | 12085566 | 14565820 | 516828.3 |
| 174598 | 16073264 | 10772706 | 11090830 | 120307.8 | 5770676 | 5136977 | 9580561 | 1760798 |
| 268123.2 | 10764097 | 12428740 | 11025079 | 382250.7 | 13824952 | 6746264 | 10068761 | 5478550 |
| 70466.59 | 158001.4 | 164110 | 163722.8 | 120819.7 | 7420869 | 7223159 | 7393170 | 388720.1 |
| 95480.53 | 210313.5 | 216097.9 | 191853.6 | 237246 | 24255531 | 20481612 | 20575975 | 158718 |
| 83422.22 | 126214.4 | 244883.4 | 276721.1 | 89155.73 | 6389481 | 5441519 | 6171462 | 107689 |
| 113896.9 | 6477964 | 9607074 | 8265468 | 113047 | 5570856 | 4531862 | 4528497 | 2930538 |
| 830633.6 | 24664281 | 39098819 | 21424171 | 450546.9 | 6561041 | 36333258 | 37726854 | 16790537 |
| 597178 | 11752456 | 14221282 | 15082557 | 185591.4 | 5155259 | 17659283 | 15080114 | 14893134 |
| 340737.6 | 34778582 | 27518264 | 33377220 | 1020071 | 53986820 | 47856161 | 47925988 | 27883406 |
| 213095.7 | 2726411 | 20501113 | 6804758 | 184668.7 | 52052100 | 49989003 | 45254163 | 5600856 |
| 186174.7 | 42368732 | 43891112 | 42845598 | 322487.5 | 28837961 | 40257859 | 31063850 | 38916430 |
| 395211.1 | 18315096 | 27509733 | 18255004 | 504206 | 18738097 | 22436913 | 20122772 | 8035016 |
| 3653169 | 14491029 | 16160917 | 13901630 | 2517067 | 22130228 | 21150945 | 20675377 | 8216625 |
| 1523075 | 6943900 | 3398846 | 9843329 | 196570.9 | 16817726 | 15385105 | 21428544 | 45494012 |
| 995056.2 | 21672409 | 23363953 | 23251351 | 1740968 | 13199801 | 20135865 | 5958358 | 10776587 |
| 1484527 | 9383244 | 10224336 | 11023295 | 1692812 | 12000779 | 16554011 | 12375546 | 16064135 |
| 124227.6 | 200185.3 | 389975.3 | 1415700 | 137829.8 | 3178715 | 2578250 | 2015357 | 5292006 |
| 134499.5 | 949702.8 | 1144863 | 1092697 | 132344.8 | 4823537 | 4257965 | 4432915 | 485479.7 |
| 71126.13 | 6784556 | 7001117 | 6717267 | 73757.62 | 6367303 | 6135408 | 4962178 | 3204851 |
| 858935.7 | 3449563 | 3899991 | 3932023 | 958228.2 | 8974771 | 8748051 | 9091984 | 7093001 |
| 622347.5 | 5039711 | 873804.8 | 2740066 | 257171.1 | 13321658 | 3917921 | 8159073 | 10589554 |
| 17945489 | 18564246 | 14247170 | 13886512 | 24700049 | 99920334 | 85089345 | 1.01E+08 | 11180547 |
| 3803749 | 12389445 | 64024035 | 13594995 | 3866098 | 61616015 | 10303630 | 12110043 | 15710650 |
| 67758.23 | 7697676 | 6695465 | 6548800 | 72408.5 | 7424941 | 6782698 | 6972374 | 4964266 |
| 263341 | 7323088 | 7033753 | 6773860 | 687600 | 9248426 | 8359193 | 7053688 | 7249217 |
| 395759.1 | 19107040 | 24838493 | 17007650 | 560811.1 | 12332794 | 21501619 | 16801556 | 14232315 |
| 258578.1 | 31886427 | 9415526 | 28231599 | 290518.5 | 9991576 | 8669236 | 13751007 | 11290290 |
| 543978.3 | 13497276 | 10273902 | 11787221 | 628589.2 | 12578658 | 8156415 | 13013251 | 6504392 |
| 145666.4 | 12670497 | 11317456 | 9891741 | 173363.4 | 16633598 | 4615362 | 14338360 | 9421915 |
| 6408190 | 59535299 | 21278899 | 28666283 | 5838149 | 69562693 | 23116496 | 26432453 | 4683047 |
| 112705.6 | 9283904 | 16257922 | 7068835 | 346170.7 | 15769908 | 9177212 | 9043338 | 3383989 |
| 112903.2 | 8237432 | 5609828 | 7005416 | 142855.6 | 7133984 | 5206001 | 5442372 | 3644523 |
| 162921.9 | 7458309 | 8229703 | 9249195 | 649174.2 | 7954631 | 7969709 | 3725659 | 4592339 |
| 5572272 | 13819823 | 74342849 | 41360444 | 6850236 | 50932216 | 19495655 | 40894892 | 23978878 |
| 3273287 | 18165056 | 66169414 | 28466769 | 3867484 | 21281483 | 15205644 | 1.3E+08 | 11752468 |
| 285084.9 | 4872438 | 6629811 | 10140595 | 156946.8 | 6503108 | 3751423 | 2864631 | 3443907 |
| 380151.5 | 6624748 | 6446594 | 7740775 | 609324 | 6368917 | 6337324 | 6358846 | 4360120 |
| 261284 | 15561093 | 14813986 | 15669975 | 458918 | 20492963 | 18070150 | 19744642 | 7445934 |
| 1443897 | 9153229 | 26438738 | 26271897 | 538072.7 | 45986038 | 44072552 | 56942912 | 32277205 |
| 882566.7 | 32528691 | 13572836 | 13700543 | 3246682 | 16369064 | 13427956 | 6265068 | 5149540 |
| 33667877 | 1.8E+08 | 1.47E+08 | 1.43E+08 | 45323872 | 4.35E+08 | 4E+08 | 3.6E+08 | 83828424 |
| 1717313 | 1.13E+08 | 1.21E+08 | 1.21E+08 | 791511.4 | 72143183 | 70089407 | 76233000 | 42136575 |

| | | | | | | | | |
|---|---|---|---|---|---|---|---|---|
| 433627.8 | 8343929 | 6833713 | 8377783 | 833482 | 6943401 | 6142346 | 7289524 | 4661365 |
| 123731.9 | 1139882 | 3170185 | 1626930 | 153100.2 | 12869653 | 8335471 | 9843144 | 1727048 |
| 89292.07 | 7983169 | 3970811 | 7302456 | 97804.81 | 7491002 | 6059322 | 6027893 | 4192454 |
| 1261587 | 9756899 | 9482363 | 9198851 | 455920.1 | 14128198 | 11679661 | 12168275 | 4130071 |
| 261284 | 15561093 | 14813986 | 15669975 | 458918 | 20492963 | 18070150 | 19744642 | 7445934 |
| 67005.63 | 351337.4 | 130231 | 271580.6 | 161665.6 | 6485364 | 3900802 | 5762412 | 216678.2 |
| 191170.9 | 4255475 | 4796227 | 3630119 | 593810.1 | 3928284 | 5460930 | 2228964 | 1822697 |
| 144776.4 | 12681892 | 13171933 | 12573466 | 123755.1 | 18058952 | 18078428 | 17125294 | 14199206 |
| 89082.91 | 337364.7 | 874573.4 | 643980.1 | 180957 | 2804855 | 2829577 | 6970353 | 488288.1 |
| 1143387 | 15209472 | 17045877 | 14524478 | 1425460 | 15603935 | 7922670 | 9846728 | 7324418 |
| 378918.3 | 1500960 | 11560403 | 1566044 | 339168.1 | 23483382 | 17603181 | 17687840 | 564259.9 |
| 5317745 | 52935985 | 54102570 | 53556287 | 6284114 | 84518125 | 79174349 | 81518246 | 49263696 |
| 149796 | 11829094 | 13901561 | 14819962 | 246009.6 | 11713848 | 15228840 | 24442143 | 9048991 |
| 369348.1 | 19978113 | 20392392 | 27122901 | 2526935 | 11033772 | 8464792 | 8107986 | 4228068 |
| 1119754 | 10206890 | 10086900 | 9969476 | 604299.5 | 10504997 | 10112128 | 6553376 | 6927571 |
| 9139297 | 1.42E+08 | 1.39E+08 | 1.37E+08 | 7884123 | 1.32E+08 | 1.24E+08 | 1.51E+08 | 1.06E+08 |
| 5080528 | 67393189 | 75285471 | 68271601 | 3413156 | 54544289 | 55828080 | 48806921 | 42039363 |
| 2329808 | 39919118 | 35949177 | 34958784 | 1703114 | 29989089 | 25761087 | 28197888 | 23820136 |
| 94986.03 | 6336630 | 8755859 | 6749999 | 129841.2 | 2374951 | 2955341 | 2477137 | 244237.9 |
| 3711327 | 3.56E+08 | 3.64E+08 | 3.38E+08 | 479998 | 1.31E+08 | 1.21E+08 | 1.17E+08 | 8156850 |
| 181394.5 | 3590243 | 4122317 | 3964723 | 240157.2 | 4785412 | 4197102 | 4466352 | 3466184 |
| 3339900 | 74556861 | 69580884 | 63537240 | 2410354 | 75433613 | 72130922 | 78093875 | 35681334 |
| 2503199 | 14686419 | 12676940 | 14794610 | 2298733 | 18056746 | 16525626 | 16255221 | 5097406 |
| 1095569 | 1099157 | 1173563 | 820921.6 | 1983889 | 10031509 | 9792560 | 10914677 | 1730382 |
| 571370.7 | 3.22E+08 | 3.16E+08 | 3.26E+08 | 207908.2 | 1.3E+08 | 1.08E+08 | 99992794 | 18069042 |
| 159732.4 | 58616322 | 48223809 | 50316303 | 267016 | 37663983 | 33021452 | 18887359 | 33602681 |
| 91820.76 | 30114865 | 26826715 | 25911348 | 89990.37 | 21861335 | 20803021 | 19016744 | 8875052 |
| 983024.4 | 372727.8 | 3189564 | 1302928 | 1680134 | 51686806 | 47229357 | 48890847 | 399972.1 |
| 407809.7 | 866603.2 | 724975 | 260458.5 | 254295.5 | 7391285 | 6919616 | 6043223 | 203578.9 |
| 5790710 | 7463234 | 8281493 | 7124400 | 12799157 | 1.06E+08 | 1.05E+08 | 1.12E+08 | 8014432 |
| 32532366 | 4.62E+08 | 4.6E+08 | 4.44E+08 | 20299902 | 5.23E+08 | 4.66E+08 | 4.59E+08 | 5.14E+08 |
| 394107.7 | 7374513 | 8233759 | 5551151 | 1145566 | 11146101 | 6754674 | 8725641 | 1364925 |
| 3543416 | 42514606 | 40262528 | 39383046 | 2903887 | 66793656 | 59016137 | 59344443 | 87473862 |
| 2689998 | 3633045 | 2751519 | 3352306 | 178056.4 | 16834716 | 16731225 | 16950871 | 4274629 |
| 248585.5 | 17319858 | 18130188 | 17254140 | 1119825 | 22764854 | 18389636 | 18735459 | 13466334 |
| 261409.3 | 11055011 | 10657412 | 11600059 | 222256.6 | 24609373 | 23628443 | 22723584 | 39111457 |
| 800294.4 | 20566226 | 16879846 | 15862173 | 898916.8 | 11531864 | 11098789 | 10269640 | 5224716 |
| 1158821 | 10303159 | 8743881 | 8413791 | 1187117 | 32925410 | 26899723 | 27777875 | 1461842 |
| 58572.53 | 8760316 | 10216219 | 9564444 | 67792.44 | 4372699 | 4000918 | 4101548 | 841731 |
| 95026.28 | 12374859 | 11225945 | 12532400 | 104061.2 | 9407834 | 7991195 | 9069805 | 8695335 |
| 180514.5 | 37614947 | 37927779 | 36663577 | 196373.9 | 37807169 | 30761183 | 30641719 | 20585193 |
| 2565750 | 24613739 | 20800366 | 17072429 | 2631866 | 15795155 | 14576085 | 14299920 | 9384035 |
| 1270354 | 10576455 | 10920990 | 11230407 | 1223082 | 5541514 | 5964456 | 30019510 | 4582700 |
| 93337.84 | 20275774 | 18039031 | 17810089 | 105860.2 | 59179867 | 55496273 | 54880709 | 1.01E+08 |
| 50470.95 | 90602.85 | 95200.79 | 77396.56 | 58104.38 | 2065910 | 1777599 | 1828927 | 71604 |
| 157924.8 | 25512477 | 22320925 | 20150424 | 189018.8 | 44092900 | 25360549 | 10823209 | 10149176 |
| 83555.59 | 9162629 | 8329179 | 8268867 | 83090.91 | 7460565 | 6740392 | 5532685 | 1583372 |
| 3190472 | 87982228 | 87628798 | 88352177 | 3599090 | 91562405 | 81493781 | 82748857 | 59016232 |
| 175433.7 | 13061630 | 12833592 | 13586752 | 163010.6 | 16927197 | 13287604 | 13882672 | 8881665 |
| 294163.8 | 516959.3 | 637701.2 | 567588.3 | 702652.3 | 5415449 | 4732599 | 5230912 | 167601.2 |
| 257254.8 | 20199590 | 22723761 | 21323797 | 133850.9 | 19174593 | 17581274 | 16571754 | 5113067 |

| | | | | | | | | |
|---|---|---|---|---|---|---|---|---|
| 138908.7 | 3269207 | 6037509 | 6056546 | 410506.6 | 6971030 | 7626998 | 5734190 | 3522859 |
| 569817.2 | 33669191 | 34399392 | 32933900 | 437811.3 | 32635172 | 30863932 | 28190090 | 15195363 |
| 94607.05 | 5723868 | 5342069 | 4510440 | 109846.7 | 8832886 | 9199933 | 9686874 | 4525486 |
| 2086493 | 63592442 | 58256767 | 45249571 | 1626727 | 55577369 | 52438831 | 37433502 | 40836136 |
| 94130.1 | 19279186 | 19373389 | 13588638 | 90187.1 | 13028562 | 12479531 | 8912236 | 7511162 |
| 274410.3 | 1.32E+08 | 1.31E+08 | 1.29E+08 | 208168.1 | 75593422 | 64032755 | 56999057 | 11294748 |
| 87228.12 | 23601981 | 19027516 | 18875973 | 92505.67 | 13228654 | 12580527 | 12433887 | 231047.4 |
| 24147819 | 1.28E+09 | 1.29E+09 | 1.26E+09 | 31136842 | 1.38E+09 | 1.25E+09 | 1.24E+09 | 1.83E+08 |
| 103663.9 | 1.27E+08 | 99109714 | 1.08E+08 | 95749.12 | 1.2E+08 | 1.1E+08 | 91393263 | 63647769 |
| 204957.5 | 10870484 | 5533443 | 6316887 | 145635.4 | 13036760 | 10290690 | 3297823 | 2162960 |
| 1626508 | 15235214 | 14447455 | 15057500 | 1664623 | 15034123 | 13484380 | 14122369 | 1844486 |
| 122989 | 23016747 | 20751556 | 19101618 | 163069.4 | 23245960 | 22613350 | 20656042 | 15822470 |
| 313228.1 | 35123935 | 37037138 | 37092644 | 105539.9 | 27025704 | 25925395 | 26116616 | 32817002 |
| 438296.3 | 2.12E+08 | 67927036 | 69197327 | 414046.6 | 1.07E+08 | 1.46E+08 | 18864634 | 34297724 |
| 100885.3 | 15516807 | 19022348 | 18210286 | 102290.7 | 17282538 | 13965423 | 14688382 | 1786214 |
| 95513.06 | 21289045 | 20414546 | 19395882 | 100368.9 | 28482501 | 25769988 | 24157726 | 20977138 |
| 89776.15 | 11828466 | 9034733 | 9260889 | 93154 | 9877477 | 8826442 | 8213663 | 3102342 |
| 64968.76 | 7276425 | 4487444 | 5709667 | 61918.89 | 6404667 | 6235122 | 5478663 | 4224877 |
| 618769 | 1.22E+08 | 1.45E+08 | 1.39E+08 | 179734.6 | 45531085 | 41978311 | 43324429 | 225826.2 |
| 90835.6 | 8243305 | 7575122 | 7340282 | 95669.28 | 7803642 | 7264982 | 7498203 | 6756470 |
| 74666.48 | 40229942 | 34487132 | 32446612 | 96080.35 | 41957486 | 38195235 | 35067240 | 36961237 |
| 166941.9 | 86282506 | 75383455 | 72006862 | 143743.3 | 78100879 | 71539672 | 67487467 | 38024459 |
| 155661.6 | 20093382 | 19071577 | 19280158 | 308486.8 | 25849168 | 25665256 | 23748574 | 26830233 |
| 77029.86 | 5874249 | 5600854 | 4684164 | 81704.74 | 8599928 | 7566288 | 6235269 | 8432773 |
| 154931.8 | 6788221 | 5566180 | 4591442 | 161210 | 8834699 | 6832897 | 5633761 | 9061004 |
| 372609.2 | 1.11E+08 | 1.04E+08 | 1.03E+08 | 104463.1 | 1.24E+08 | 1.1E+08 | 1.25E+08 | 94247595 |
| 104072.6 | 1.12E+08 | 94073698 | 1.04E+08 | 165674.4 | 99991485 | 84912217 | 78689796 | 59757494 |
| 136613.5 | 35665618 | 37499501 | 39426937 | 90296.89 | 37647329 | 36327729 | 36859091 | 19872314 |
| 104961.2 | 5785044 | 6541626 | 5707497 | 88473.62 | 6463577 | 5216932 | 4614495 | 1909883 |
| 403645.7 | 1.55E+08 | 1.46E+08 | 1.49E+08 | 670809.6 | 1.35E+08 | 1.09E+08 | 1.07E+08 | 42565015 |
| 4909158 | 1.05E+09 | 1.11E+09 | 1.11E+09 | 173665.8 | 4.07E+08 | 3.41E+08 | 3.63E+08 | 2064603 |
| 1004243 | 1.67E+08 | 1.71E+08 | 1.67E+08 | 1242353 | 1.7E+08 | 1.44E+08 | 1.56E+08 | 97032864 |
| 30385575 | 1.7E+09 | 1.76E+09 | 1.72E+09 | 39523742 | 2.15E+09 | 1.87E+09 | 1.94E+09 | 2E+09 |
| 98658.78 | 1383506 | 1489735 | 982596.4 | 50500.94 | 2151594 | 1194685 | 4075241 | 4620803 |
| 152331.2 | 3326474 | 1344020 | 2772187 | 155673.6 | 4616613 | 4914806 | 5866568 | 2639776 |
| 152002.9 | 6766821 | 8322347 | 4697414 | 160013.4 | 5720640 | 5158097 | 3735378 | 248267.7 |
| 753921.7 | 1875220 | 1781790 | 1988885 | 2032661 | 8322822 | 3374320 | 18567018 | 20459545 |
| 330486 | 15093348 | 14648016 | 3010679 | 566805.9 | 41938776 | 38866358 | 38228482 | 89901977 |
| 136498.4 | 338252.5 | 2063686 | 334876.6 | 260650.5 | 12982179 | 11429206 | 11422156 | 29864052 |
| 56571.14 | 11426080 | 10780952 | 10722074 | 75388.1 | 10951936 | 9122456 | 7961011 | 9174307 |
| 79854.26 | 1748501 | 4106481 | 2617283 | 99516.56 | 6259036 | 3224021 | 2624088 | 3334756 |
| 97405.37 | 10571458 | 8653343 | 8240125 | 95678.83 | 11506428 | 10010485 | 7951263 | 4541599 |
| 168799.2 | 8347654 | 7699142 | 8120144 | 168416.9 | 7731116 | 6381590 | 6401746 | 172120.1 |
| 349878.4 | 3.05E+08 | 2.49E+08 | 2.5E+08 | 686872.2 | 2.4E+08 | 2.31E+08 | 1.98E+08 | 1.46E+08 |
| 314532.7 | 9993145 | 6870052 | 6588304 | 375649.3 | 8145358 | 7076114 | 21499083 | 7297765 |
| 152813.9 | 41207920 | 41156123 | 44410780 | 154793.2 | 45993987 | 35514825 | 7845698 | 7139026 |
| 177948.3 | 83858981 | 94265363 | 90948257 | 143571.7 | 84672865 | 74367948 | 72954817 | 44499249 |
| 84520.47 | 9315749 | 8046020 | 8239198 | 84604.79 | 11729302 | 10827603 | 9103735 | 12514044 |
| 1214072 | 91553933 | 92563425 | 82367255 | 1286850 | 70106307 | 61760339 | 64850887 | 15860066 |
| 161263.7 | 12151787 | 11481305 | 11673412 | 176355.8 | 13914443 | 11796554 | 13019336 | 3804582 |
| 128957.6 | 57096776 | 53712516 | 55071928 | 125077.1 | 63764127 | 53636628 | 62926726 | 22271856 |

| | | | | | | | | |
|---|---|---|---|---|---|---|---|---|
| 156813.3 | 12300655 | 11853595 | 11502451 | 151513.5 | 9596784 | 8212597 | 7239158 | 451829.1 |
| 10236669 | 6.4E+08 | 6.63E+08 | 5.24E+08 | 9547755 | 7.65E+08 | 6.89E+08 | 6.97E+08 | 4.94E+08 |
| 1440424 | 39993535 | 43166834 | 41136083 | 3481326 | 97856987 | 87649658 | 88553750 | 18558042 |
| 1176423 | 1.02E+08 | 1.05E+08 | 98587054 | 1185485 | 75697048 | 67171289 | 68192775 | 31088449 |
| 96420.08 | 879032.3 | 1398183 | 4189535 | 107496.1 | 3935059 | 4064855 | 5967272 | 3897713 |
| 64055.64 | 3628909 | 3483409 | 3703364 | 116231.9 | 2457450 | 2668566 | 2799631 | 502749.6 |
| 6289120 | 90677822 | 89840575 | 95643601 | 6446483 | 1.07E+08 | 99611264 | 1.07E+08 | 72666853 |
| 201093.6 | 243840.3 | 223508.6 | 225282.7 | 204994.9 | 35743120 | 29120962 | 27938279 | 215782 |
| 185972.6 | 14126573 | 14783800 | 13757736 | 203015.8 | 15003994 | 12896220 | 12937098 | 8096021 |
| 98585.65 | 30923049 | 31467887 | 30878199 | 160785 | 25943119 | 25525422 | 25239628 | 7590879 |
| 1077950 | 899029.6 | 1496486 | 3051754 | 1738057 | 1.52E+08 | 1.39E+08 | 1.41E+08 | 979685.8 |
| 195271.2 | 1.19E+08 | 1.35E+08 | 1.26E+08 | 215063.7 | 84817593 | 69920258 | 63647937 | 8051614 |
| 272282.4 | 10408980 | 6855309 | 6713166 | 93237.09 | 5251613 | 5039278 | 4745530 | 5363646 |
| 158359.3 | 5809939 | 6611041 | 6059640 | 107572.7 | 7433843 | 3592696 | 8264103 | 11751727 |
| 104004.6 | 13567602 | 18403532 | 15331527 | 148570.9 | 14910993 | 13966920 | 10618831 | 8287385 |
| 176689 | 52303623 | 52082145 | 53594895 | 252272.3 | 63975171 | 50797804 | 35983489 | 28795302 |
| 121176.3 | 4375032 | 3549121 | 4123754 | 117078.1 | 3856098 | 3796587 | 5445609 | 3114321 |
| 417167.3 | 24054370 | 24231786 | 28717222 | 297357.2 | 26483401 | 25939770 | 27406073 | 23510590 |
| 952264 | 27454001 | 31570364 | 31065533 | 192425.6 | 29630107 | 26177579 | 23352679 | 19079378 |
| 381972.7 | 12426575 | 12414499 | 12997036 | 774696.9 | 7332312 | 5206930 | 6733258 | 8239416 |
| 139238.6 | 2057549 | 3233294 | 3756087 | 134532.8 | 4383916 | 4168591 | 3240448 | 1804300 |
| 100759.7 | 11072401 | 12315484 | 11632006 | 115552.6 | 9705044 | 8926656 | 8369031 | 4628698 |
| 408503.9 | 29088855 | 26683773 | 26298321 | 404305.4 | 35014450 | 25522976 | 23114345 | 33426031 |
| 109287.5 | 174614.1 | 274403.3 | 185102 | 120223.7 | 21632696 | 19271188 | 20046997 | 150545.9 |
| 99322.22 | 297614.4 | 368902.2 | 318088.5 | 155761.7 | 14388819 | 11193485 | 13177117 | 301222.5 |
| 225349.3 | 8075228 | 7072656 | 989074.3 | 209575.4 | 5908993 | 6915504 | 5761649 | 2706668 |
| 98787.39 | 200678.9 | 204987.6 | 196704.8 | 140836.6 | 14849317 | 14302417 | 14181577 | 169510.7 |
| 421458.9 | 57170645 | 58727587 | 56187996 | 1445758 | 46863487 | 44770625 | 44971266 | 19965903 |
| 74940.02 | 389475.1 | 236261.5 | 393088.8 | 88477.5 | 3257733 | 3172509 | 3408711 | 147603.7 |
| 139533.1 | 496667.1 | 798940.7 | 1853068 | 60651.02 | 4625739 | 2814079 | 3042690 | 998603.1 |
| 94042.72 | 1800297 | 1472114 | 5969571 | 113291.1 | 4215802 | 1362514 | 1001644 | 586713.2 |
| 374250.2 | 56519637 | 65298764 | 64073996 | 225854.5 | 71748439 | 60791557 | 61330476 | 37209818 |
| 1278946 | 27154460 | 26379018 | 27500987 | 883195.5 | 13665376 | 12269142 | 10494498 | 2564431 |
| 129938.8 | 11543171 | 12227358 | 12343435 | 272149.2 | 13361066 | 10479492 | 4426623 | 3811222 |
| 1214225 | 1.04E+08 | 1.06E+08 | 1.07E+08 | 241214.6 | 70423848 | 57601475 | 60216953 | 30878265 |
| 741484.1 | 43642363 | 54850519 | 53972984 | 177230.8 | 20884254 | 18364547 | 8584987 | 1100637 |
| 371032.1 | 35788572 | 40903210 | 42103635 | 215144 | 13447236 | 10796149 | 11491102 | 274688.7 |
| 234167.5 | 25761553 | 32258334 | 30965571 | 217734.4 | 11286170 | 8770371 | 7640118 | 270152.9 |
| 521120.3 | 2.42E+08 | 2.38E+08 | 2.3E+08 | 476334.5 | 2.2E+08 | 1.86E+08 | 1.87E+08 | 1.21E+08 |
| 193204.2 | 15780080 | 19314102 | 19154567 | 202799.4 | 12962777 | 3985384 | 10855285 | 2720516 |
| 84953.94 | 53530005 | 43551751 | 37119255 | 86432.52 | 54806705 | 52214662 | 35626119 | 47895031 |
| 323942.7 | 8346280 | 10029367 | 10615486 | 295211.8 | 13416645 | 12645068 | 10921715 | 10559335 |
| 92388.61 | 57850171 | 55602455 | 56039613 | 125836.6 | 50869574 | 46570405 | 48404704 | 23683063 |
| 710109 | 33684467 | 42596886 | 42003111 | 573429.8 | 38170558 | 32793810 | 34155251 | 19787415 |
| 207910 | 1.86E+08 | 1.65E+08 | 1.66E+08 | 203983.9 | 2.48E+08 | 2.14E+08 | 1.48E+08 | 1.57E+08 |
| 92179.57 | 32474911 | 32412202 | 30933842 | 166077.1 | 44403853 | 42004727 | 44033711 | 14724497 |
| 102902.4 | 10137951 | 9549515 | 9084575 | 105169.5 | 10629702 | 9597434 | 9192738 | 6469866 |
| 114615.4 | 234422.5 | 244645.8 | 236777.2 | 146156.9 | 9249494 | 6899889 | 8448028 | 194257.8 |
| 185913.9 | 13033252 | 17698208 | 16666869 | 130239 | 15135759 | 13735843 | 15839592 | 2552613 |
| 79510.3 | 580438.7 | 2683783 | 617783.1 | 88122.83 | 7692058 | 7247525 | 6453965 | 1175105 |
| 75214.13 | 10175499 | 10045738 | 10501186 | 90160.74 | 8635216 | 7948513 | 6386257 | 122478.4 |

| | | | | | | | | |
|---|---|---|---|---|---|---|---|---|
| 156489.8 | 19063838 | 10635934 | 12490131 | 166134.5 | 21802019 | 8720829 | 14590861 | 4538647 |
| 281580 | 12792116 | 13038531 | 13147292 | 114343.2 | 8841431 | 8560870 | 7946710 | 8550802 |
| 134476.9 | 13696897 | 12644262 | 13029912 | 137920.5 | 12884245 | 11084916 | 10863271 | 7455274 |
| 202250.1 | 2235021 | 5569394 | 8422488 | 54038.66 | 3266848 | 2931669 | 3465313 | 659022 |
| 1148449 | 32770091 | 25100194 | 26700242 | 1320923 | 26461240 | 24032197 | 29790332 | 16015921 |
| 1446631 | 16822127 | 45199486 | 27627758 | 1511794 | 18746349 | 43320468 | 21148023 | 20307233 |
| 796574.4 | 14508289 | 12179832 | 16644056 | 890711.7 | 11555170 | 9440843 | 5327581 | 10716002 |
| 1549700 | 17573576 | 18778840 | 15880919 | 1461512 | 16733417 | 16616736 | 16994082 | 8942406 |
| 175599.6 | 4073433 | 3141986 | 4353276 | 178208.5 | 4194760 | 4551964 | 5337102 | 1043220 |
| 224835.1 | 30262725 | 15266381 | 11971044 | 447798.3 | 16447939 | 15180746 | 13086053 | 5486737 |
| 206773.6 | 8203103 | 8435579 | 8775335 | 742715.3 | 9299712 | 9198250 | 9018023 | 2987970 |
| 1260986 | 6527050 | 5644871 | 9123095 | 876574.4 | 26551371 | 16191806 | 20714112 | 20334206 |
| 987964 | 9677452 | 9545407 | 6314554 | 1111244 | 7995328 | 5678522 | 5575583 | 2040770 |
| 694913.9 | 23875926 | 26741921 | 17243324 | 1112289 | 32027461 | 18546295 | 28936727 | 19497175 |
| 624708.6 | 1077018 | 742195.8 | 364749.2 | 161845.3 | 27156099 | 22081875 | 25476397 | 75989579 |
| 145633.2 | 1640863 | 993161.5 | 3546028 | 366723.6 | 33486071 | 29753526 | 32565811 | 3077983 |
| 4875171 | 14053842 | 1485550 | 14800091 | 217357.1 | 32481721 | 33229808 | 38806609 | 59823832 |
| 694913.9 | 9310202 | 9579258 | 9023403 | 1231010 | 98211219 | 62480187 | 74526766 | 25956014 |
| 881969.6 | 12616398 | 11357713 | 12250268 | 1093235 | 11497631 | 11102691 | 11066788 | 9289052 |
| 112909.3 | 5707769 | 6031793 | 5701532 | 123363.5 | 4092833 | 6719480 | 6785423 | 4509073 |
| 198700.2 | 1838158 | 3658861 | 780458 | 167706.9 | 15059919 | 12692335 | 13572038 | 29995476 |
| 4703383 | 46148296 | 41776106 | 40617432 | 6075546 | 50286639 | 45749244 | 45119532 | 48292334 |
| 3479897 | 34597020 | 34481446 | 25652344 | 3624765 | 36151958 | 37243917 | 39433008 | 11803860 |
| 108148.3 | 11232560 | 12514032 | 12288855 | 142779.7 | 11325385 | 10081683 | 11854542 | 5681081 |
| 95987.35 | 664586.3 | 1345069 | 370862.2 | 94503.15 | 6395169 | 4074939 | 5025186 | 12969862 |
| 82130.7 | 505980.2 | 577538.2 | 367101.2 | 86158.44 | 6927686 | 6006845 | 4633664 | 17417594 |
| 99776.27 | 641209.4 | 573172.7 | 413290.8 | 99348.04 | 3565743 | 2803411 | 3604687 | 14671816 |
| 183125 | 8649321 | 8258545 | 2493079 | 255901.6 | 5831715 | 6301953 | 6183584 | 16855144 |
| 156102.1 | 12272141 | 13217295 | 13530315 | 69487.36 | 7337248 | 5969282 | 4740104 | 1049592 |
| 102310.5 | 6874978 | 8563114 | 7052926 | 145859.3 | 3446444 | 3511145 | 2743931 | 356371.1 |
| 484184.9 | 1786411 | 514237.6 | 597120.9 | 1173778 | 38297660 | 32583606 | 34500159 | 392719.6 |
| 121435.4 | 10014829 | 10490912 | 3698856 | 392417.9 | 13964675 | 8864812 | 8549287 | 12720173 |
| 3208289 | 5305743 | 2443921 | 2753433 | 7020504 | 2.66E+08 | 2.4E+08 | 2.51E+08 | 1136592 |
| 525889 | 518336.9 | 920810.8 | 1116984 | 1212299 | 74119624 | 72073462 | 66851619 | 1056778 |
| 179288.5 | 21127938 | 98663646 | 1.1E+08 | 3321395 | 2.31E+08 | 1.43E+08 | 1.41E+08 | 65957533 |
| 637048.8 | 1.26E+08 | 1.32E+08 | 1.29E+08 | 398439.8 | 1.28E+08 | 1.17E+08 | 1.2E+08 | 82468880 |
| 2108701 | 59877318 | 54358390 | 59398159 | 5489137 | 1.3E+08 | 69901252 | 1.41E+08 | 65556517 |
| 938766.1 | 34807639 | 41619597 | 41153302 | 202695.9 | 1.83E+08 | 97390018 | 84419817 | 30876193 |
| 238260.5 | 34068135 | 32562629 | 31952321 | 574203 | 80645066 | 70016909 | 72360073 | 4407389 |
| 1496885 | 55106603 | 70612002 | 69708039 | 105745.4 | 23399046 | 21372422 | 18671305 | 2082816 |
| 520527.3 | 79637911 | 83665338 | 79458626 | 312708.1 | 83750379 | 72754027 | 74027485 | 47435089 |
| 155277.6 | 17392209 | 14714151 | 16077006 | 168361.9 | 19106933 | 20108569 | 10703500 | 2554590 |
| 1082847 | 1912417 | 3731004 | 422434 | 1832504 | 43198758 | 38600960 | 39724673 | 3905707 |
| 945317.8 | 44351399 | 49410656 | 52469424 | 856315.4 | 44199835 | 40641832 | 37264506 | 5516653 |
| 1041799 | 2686702 | 2749480 | 8816351 | 2181403 | 45006620 | 39326475 | 38996843 | 1797128 |
| 437593.4 | 2839353 | 680676.6 | 1603212 | 1094565 | 71067966 | 65869465 | 67572564 | 612179.9 |
| 108094.8 | 582991.8 | 395054.9 | 402295.4 | 408897.6 | 30379106 | 28425249 | 29299007 | 370137.7 |
| 107397.9 | 73252392 | 61567778 | 59670367 | 133556.5 | 55017471 | 53712221 | 51930761 | 28243116 |
| 200212.9 | 496886.5 | 249701.8 | 406013.1 | 677351.8 | 27091642 | 24801835 | 26530912 | 177686.3 |
| 368585.9 | 1.91E+08 | 1.08E+08 | 69008976 | 414842.7 | 1.59E+08 | 1.47E+08 | 22899965 | 19653621 |
| 392955.5 | 14868903 | 15351230 | 14995351 | 222256.6 | 31152304 | 29208578 | 32468833 | 57593732 |

| | | | | | | | | |
|---|---|---|---|---|---|---|---|---|
| 88778.88 | 755673.3 | 499410.4 | 612835.4 | 243769.6 | 27873869 | 25596911 | 25461966 | 352445.2 |
| 85971.01 | 19453622 | 19358483 | 15631784 | 282622.5 | 29096379 | 25434921 | 27249619 | 6044486 |
| 10505916 | 1.09E+08 | 83414730 | 96540955 | 9120583 | 90028781 | 84139009 | 94112799 | 39303240 |
| 2586887 | 22571231 | 17701563 | 22072147 | 2579826 | 58339530 | 48133748 | 57703131 | 25729099 |
| 1014445 | 1.47E+08 | 1.47E+08 | 1.46E+08 | 1785856 | 1.46E+08 | 1.36E+08 | 1.43E+08 | 44713032 |
| 5148706 | 78493547 | 73625052 | 73614814 | 5232573 | 68352706 | 63508023 | 89085368 | 29823139 |
| 219341.9 | 1461227 | 10481786 | 14232391 | 198643.9 | 49962096 | 9722412 | 3610299 | 1221611 |
| 312377.5 | 16436463 | 17564810 | 18373346 | 374135.4 | 23770223 | 19035302 | 20051423 | 13284873 |
| 548162.4 | 11527062 | 10086963 | 9308353 | 795926.8 | 17560997 | 38677247 | 50292962 | 4537269 |
| 1082847 | 1286111 | 3731004 | 1191548 | 343057.1 | 14291938 | 12089203 | 11906037 | 3905707 |
| 265462.4 | 36486887 | 30212307 | 34691689 | 314363.4 | 41291410 | 35745797 | 37079805 | 30495925 |
| 204308.2 | 10486197 | 8858797 | 8655980 | 231080.5 | 37258073 | 34917139 | 36800236 | 5450494 |
| 241891 | 11742072 | 10433438 | 6590851 | 721664.4 | 49865329 | 47535615 | 48263803 | 6026908 |
| 10484467 | 1.11E+08 | 64631582 | 63163931 | 9616473 | 1.18E+08 | 87065779 | 70136798 | 40930796 |
| 7384350 | 95593127 | 1E+08 | 96591391 | 8335036 | 1.02E+08 | 97621160 | 68292849 | 70110989 |
| 79600.1 | 18370919 | 28720439 | 25242581 | 188569.7 | 29581194 | 25187979 | 21392215 | 4562864 |
| 551607.6 | 21423806 | 22748438 | 25490295 | 450375 | 21865133 | 18642550 | 22419228 | 11802497 |
| 159827.9 | 9675582 | 10255205 | 9924800 | 238077.8 | 21548733 | 19532117 | 18481508 | 177244.4 |
| 45917.27 | 2094018 | 9303150 | 4640631 | 96444.18 | 8249167 | 9045982 | 19202491 | 1854870 |
| 1484660 | 15341737 | 15019429 | 14780067 | 1722669 | 38283852 | 33379837 | 40151046 | 14280526 |
| 84840.68 | 21921579 | 23109373 | 20254345 | 103613.3 | 25669059 | 23157931 | 22499402 | 27656349 |
| 1203238 | 16420818 | 17707149 | 15715506 | 1837044 | 48084754 | 48451008 | 39294661 | 19225571 |
| 237342.2 | 38305322 | 37169932 | 36616017 | 230839 | 45469621 | 42910197 | 41758769 | 33811105 |
| 87107.8 | 37596073 | 39769237 | 43085265 | 84195.64 | 38563918 | 32629038 | 38895597 | 12735781 |
| 608625.6 | 3422370 | 9000189 | 2396435 | 1076268 | 39792801 | 35412481 | 35452568 | 691150.1 |
| 65225.75 | 2210903 | 2245375 | 2159422 | 85511.73 | 12610866 | 10802951 | 11051297 | 25791495 |
| 619711.8 | 1674030 | 11313963 | 11241319 | 151310.5 | 20932076 | 18457007 | 12264144 | 30088911 |
| 94433.15 | 724284.1 | 125672.6 | 190372.8 | 96626.13 | 17241115 | 14923999 | 10831218 | 36638498 |
| 118759.2 | 16874596 | 7449699 | 15867733 | 148777.1 | 26865034 | 30761432 | 12291472 | 9980829 |
| 118892 | 13176826 | 11280562 | 11421667 | 115382.1 | 11501654 | 9693432 | 10654310 | 3397605 |
| 310486.3 | 86364062 | 77821732 | 49842969 | 1661418 | 43439771 | 83163427 | 77699324 | 33453280 |
| 92602.47 | 1107816 | 2613673 | 2297867 | 159128.7 | 17200795 | 14522194 | 18576587 | 2584673 |
| 118286.7 | 10773487 | 11715703 | 12120307 | 136319.1 | 25734512 | 24587354 | 25917751 | 9152981 |
| 93399 | 26739365 | 30648042 | 26457112 | 117606.1 | 15485197 | 13754413 | 15844819 | 698101.7 |
| 2946156 | 15542966 | 14574825 | 13829337 | 2360219 | 44430858 | 37943629 | 31513168 | 16209473 |
| 216198.9 | 48942143 | 86692062 | 91193122 | 3166393 | 76699814 | 1.09E+08 | 1.11E+08 | 25222383 |
| 515080.9 | 10625299 | 9315196 | 9164539 | 565128.2 | 15397952 | 14440795 | 14261050 | 1984341 |
| 72472.19 | 60447949 | 53907573 | 51823888 | 271798.9 | 39155730 | 39774713 | 31608688 | 12648651 |
| 91657.45 | 2441842 | 5168097 | 6388814 | 80652.2 | 10749664 | 29676425 | 55881084 | 654179.5 |
| 81160.98 | 207972.2 | 336647.3 | 201290.6 | 120819.7 | 7420869 | 7223159 | 7393170 | 274548.9 |
| 298644.5 | 5280048 | 7565055 | 6999548 | 80588.78 | 9596461 | 7785946 | 7863133 | 2600830 |
| 3574102 | 19843288 | 22084040 | 19996597 | 241415.7 | 37704146 | 31576855 | 69289837 | 43927857 |
| 6216894 | 14986374 | 15513198 | 9434735 | 3931412 | 35791741 | 32090946 | 31292729 | 16452043 |
| 8039954 | 16812740 | 12948988 | 21227964 | 6113616 | 52254166 | 54608876 | 62018896 | 24485364 |
| 185360.7 | 211784.7 | 255194.8 | 364607.5 | 332747.5 | 10495317 | 9391402 | 11484985 | 205666.6 |
| 60286.33 | 16949810 | 17218891 | 16320645 | 89983.08 | 9979162 | 10074019 | 5331493 | 2622965 |
| 98223.39 | 7283409 | 7129633 | 7370550 | 104019.2 | 17487180 | 14507071 | 15337158 | 4308527 |
| 109470.4 | 7872326 | 11033560 | 11068496 | 127751.4 | 14506163 | 14283881 | 14857378 | 3704799 |
| 91244.41 | 13478570 | 13203133 | 13853223 | 102706.5 | 20057647 | 16610145 | 17588217 | 2661581 |
| 76155.94 | 10274843 | 10691775 | 11240448 | 92266.36 | 15696485 | 14133922 | 15097713 | 18533255 |
| 84755.41 | 1484555 | 392617.6 | 573729 | 86925.81 | 12717986 | 10883157 | 10800161 | 284622.8 |

| | | | | | | | | |
|---|---|---|---|---|---|---|---|---|
| 134187.7 | 18648425 | 16384946 | 16798410 | 143410.3 | 26282590 | 22865166 | 28319932 | 39879973 |
| 103410.1 | 4423462 | 4510583 | 4918508 | 174988.7 | 17935400 | 10016757 | 12307490 | 3375427 |
| 78411.32 | 281862.4 | 147966 | 136118.7 | 86904.81 | 9854874 | 9233251 | 10314747 | 278076 |
| 54589.85 | 2432869 | 4996880 | 731373 | 67362.18 | 4411727 | 16915200 | 46288682 | 1278174 |
| 839501.3 | 18808012 | 22298729 | 19544266 | 619825.3 | 19821013 | 20046487 | 19623413 | 14428609 |
| 164812.4 | 9878117 | 8930916 | 10155601 | 542934.8 | 20563213 | 16618338 | 11285681 | 2236805 |
| 428719.8 | 64675013 | 18635396 | 24158448 | 1900789 | 57043226 | 7173043 | 2493877 | 4954965 |
| 92769.93 | 590382.1 | 459969.4 | 1010799 | 109296.1 | 8518634 | 8498141 | 8523407 | 125481.6 |
| 119973.5 | 22134984 | 23872163 | 22572120 | 92630.4 | 20164796 | 17410395 | 16734552 | 7241539 |
| 64690.17 | 11131882 | 10328563 | 1924539 | 56457.73 | 23903510 | 21987484 | 20867021 | 12279561 |
| 73655.82 | 148870 | 145451 | 188434.2 | 74681.91 | 7232313 | 6687962 | 7498304 | 339612.8 |
| 88919.98 | 2253932 | 2031247 | 410870.3 | 299714.5 | 13087188 | 10122385 | 10306616 | 944776.1 |
| 159701.8 | 13185247 | 6078717 | 12867849 | 244287.4 | 20793562 | 22217659 | 21927935 | 26448881 |
| 169330.5 | 36380084 | 40054312 | 40491654 | 122668.7 | 34057176 | 32216852 | 35528168 | 21137513 |
| 77483.69 | 266823.5 | 216580.1 | 869645.7 | 88036.53 | 9358524 | 8426659 | 9331021 | 206401.6 |
| 447436.6 | 4773604 | 2925719 | 4334767 | 632003.4 | 13306698 | 11369313 | 7479694 | 3109350 |
| 1921017 | 20274768 | 18827321 | 18649643 | 2469224 | 21207280 | 14407098 | 20201353 | 7190986 |
| 78775.68 | 7609429 | 6422436 | 6337933 | 82295.41 | 8580116 | 7624046 | 8579448 | 13698672 |
| 102700.5 | 14095751 | 15758403 | 15769126 | 109122.3 | 9607338 | 8891176 | 8461587 | 2235832 |
| 97334.48 | 5897199 | 5450421 | 5714647 | 124178.3 | 10498672 | 9165592 | 10748730 | 139860.6 |
| 92441.2 | 12227548 | 13177076 | 13073325 | 196697 | 17907048 | 17700830 | 17043653 | 6008790 |
| 74305.34 | 16776191 | 15947593 | 17027468 | 82132.78 | 12651825 | 11262854 | 13275980 | 8318666 |
| 2515129 | 12778873 | 12185008 | 12097746 | 2546539 | 20823087 | 22418142 | 17961499 | 7844704 |
| 124066.4 | 5678320 | 4997516 | 5181505 | 199237.3 | 8968545 | 8284529 | 8224693 | 822423.9 |
| 456507.7 | 5173964 | 5266668 | 5098677 | 803029.3 | 17191547 | 18508120 | 17180188 | 7819094 |
| 70910.59 | 704411 | 777793.6 | 254629 | 93394.91 | 7277436 | 6668339 | 6618364 | 150778 |
| 341472.6 | 7730884 | 8424340 | 6810299 | 372235.9 | 7098619 | 6786764 | 7791670 | 3405042 |
| 232361.6 | 12057933 | 19407739 | 12097788 | 257350.4 | 26069390 | 23422704 | 12514167 | 3601311 |
| 50720.56 | 7990321 | 1045820 | 2046836 | 56352.58 | 9955167 | 10293102 | 10426801 | 16821133 |
| 1370479 | 31679107 | 5062579 | 2156150 | 2407829 | 1718058 | 4269299 | 1537461 | 5338104 |
| 37098.56 | 21043377 | 48124332 | 48830051 | 75576.74 | 39891983 | 18346855 | 34412969 | 5738558 |
| 381696.3 | 19864437 | 20025130 | 19376886 | 426701.7 | 39968559 | 37086444 | 38799790 | 13179907 |
| 192519.9 | 973112.9 | 1785310 | 20180091 | 849738.9 | 10205317 | 8740865 | 9277432 | 16059693 |
| 318349 | 12775700 | 17335002 | 10459925 | 542196 | 18707061 | 19413804 | 12913753 | 13448984 |
| 1601061 | 17176019 | 14335503 | 16345660 | 1728301 | 11889710 | 10197889 | 17052413 | 10541835 |
| 89764.53 | 256526.3 | 223841.7 | 426258.3 | 124975.2 | 7732825 | 7895131 | 8819219 | 168292.1 |
| 1114756 | 30521287 | 25757391 | 27919312 | 2193293 | 28271166 | 25118812 | 21429371 | 13311751 |
| 79899.84 | 4115171 | 2632326 | 3703502 | 83867.93 | 8458469 | 8063575 | 6732881 | 7940500 |
| 39215.9 | 1295224 | 627431.8 | 1135459 | 38053.75 | 3287737 | 2692138 | 2676730 | 10605703 |
| 410241.2 | 23743069 | 26019469 | 23739092 | 520460.9 | 55646845 | 43203522 | 40312239 | 7452402 |
| 111090.3 | 15516807 | 19022348 | 18210286 | 110511.8 | 17282538 | 13965423 | 14688382 | 1786214 |
| 55752.13 | 1530265 | 1760640 | 1846475 | 82367.97 | 4791637 | 3849417 | 4141448 | 342850.9 |
| 59155.48 | 690396.7 | 887044 | 830615.9 | 73721.17 | 3740527 | 3548580 | 3426970 | 9093820 |
| 75110.28 | 352744.8 | 510996 | 267602.5 | 100550.7 | 9453453 | 7645409 | 9235881 | 226598.4 |
| 378218.9 | 38055347 | 42390221 | 38186692 | 172797.6 | 10627832 | 19722259 | 38007157 | 12568962 |
| 139047.6 | 20477258 | 20992793 | 19999713 | 140957.3 | 22471265 | 19968878 | 19817116 | 16416159 |
| 165584.9 | 28659385 | 29017151 | 16928668 | 156850.8 | 25046087 | 22362962 | 22375060 | 4185805 |
| 279373.4 | 12104629 | 23968915 | 17460843 | 189641.5 | 12096867 | 21993607 | 38085233 | 17934983 |
| 106524.7 | 10715920 | 10061293 | 11055684 | 120223.8 | 15783268 | 13227899 | 14298803 | 4096392 |
| 76252.78 | 337297.7 | 559082.5 | 352887.1 | 83056.46 | 6735708 | 6216147 | 9113392 | 2789097 |
| 98546.26 | 2076627 | 719393.5 | 680629 | 145330 | 10579702 | 9211838 | 8650414 | 391057 |

| | | | | | | | | |
|---|---|---|---|---|---|---|---|---|
| 75970.37 | 6529645 | 7948228 | 8090360 | 168712.1 | 6164695 | 5615635 | 5796694 | 661798.6 |
| 484357.5 | 14223428 | 14041593 | 14413416 | 224168.1 | 17524762 | 14259554 | 16370236 | 16656916 |
| 176659.2 | 18775059 | 19407739 | 6940072 | 85799.39 | 26069390 | 23422704 | 12514167 | 3396028 |
| 2448063 | 18089939 | 20165166 | 18542833 | 2518251 | 30212365 | 27760515 | 28131265 | 16588134 |
| 94401.48 | 393588.4 | 275065.3 | 467297 | 134583.8 | 7066324 | 8370684 | 8276897 | 943477.4 |
| 102200.9 | 35123935 | 37037138 | 21529922 | 108626 | 27362312 | 25925395 | 266395 | 32817002 |
| 145621.5 | 17302316 | 17478315 | 17200068 | 79566.58 | 14098403 | 14053647 | 13751498 | 9405048 |
| 72717.55 | 7515762 | 6440638 | 8333748 | 74122.7 | 12515368 | 10573176 | 11497961 | 2726248 |
| 60589.82 | 7881151 | 10024215 | 8592150 | 67645.66 | 6073664 | 5835299 | 5207922 | 124954 |
| 209159.7 | 21533503 | 22138604 | 19260569 | 294543.9 | 28994074 | 28023871 | 23109302 | 10548560 |
| 217537 | 19887604 | 17549097 | 21041577 | 219996.2 | 20472639 | 15731210 | 18092520 | 1265964 |
| 70738.16 | 12227548 | 13177076 | 13073325 | 187947.3 | 17907048 | 17700830 | 17043653 | 5045452 |
| 858580.4 | 6197549 | 7439115 | 3381279 | 172044.9 | 6766085 | 10269745 | 9668770 | 1882484 |
| 86648.65 | 8953596 | 7251262 | 7749317 | 54025.85 | 7453784 | 5705175 | 6524213 | 4290029 |
| 115287.3 | 628046.7 | 5938217 | 668174.1 | 148793.7 | 14978336 | 15977687 | 13965552 | 681621.9 |
| 178190 | 32114160 | 35566252 | 22645538 | 236573.2 | 18695647 | 33150241 | 19357413 | 19893519 |
| 124170.6 | 5483081 | 5350263 | 6356271 | 59851.85 | 5724570 | 5053230 | 6236163 | 5142392 |
| 81043.47 | 2429152 | 2306338 | 2449727 | 50607.04 | 4402868 | 3254271 | 3675766 | 1593457 |
| 73371.97 | 9871772 | 9861318 | 10173170 | 84425.39 | 3930743 | 3870923 | 3791428 | 280077.1 |
| 208351.9 | 16967349 | 20042148 | 20381505 | 228943.5 | 18637687 | 17389468 | 16175334 | 6797519 |
| 107478.1 | 16155933 | 14356971 | 14472536 | 103845.7 | 13962227 | 12193593 | 12100087 | 4439392 |
| 90538.66 | 15294328 | 5211977 | 7455784 | 95131.25 | 8766455 | 7256244 | 5423279 | 1076355 |
| 114532.9 | 3834199 | 523948 | 1401996 | 107461.3 | 3324564 | 5494321 | 5608668 | 386329.6 |
| 70553.45 | 551838.6 | 2507141 | 829376.4 | 78821.88 | 5744360 | 3975089 | 5015536 | 169204.4 |
| 665197.1 | 10821982 | 12473569 | 12128601 | 521465.5 | 10592602 | 9764823 | 10507307 | 9571983 |
| 98833.52 | 15033109 | 12840369 | 8616916 | 89959.87 | 8727107 | 8426401 | 6459256 | 4739141 |
| 79664.07 | 363310 | 184160 | 138011.1 | 86333.89 | 4466494 | 3518393 | 3129270 | 9083373 |
| 91956.1 | 9696517 | 8749061 | 9181118 | 106962.2 | 9922921 | 7671541 | 6859466 | 2111573 |
| 84358.06 | 911203 | 592334.2 | 526556.7 | 108875.4 | 6554970 | 6530906 | 6702975 | 324027.7 |
| 80777.1 | 11073528 | 11803560 | 9867061 | 89131.33 | 7509868 | 6940963 | 5870198 | 170751.5 |
| 95815.99 | 7120549 | 7246179 | 7093371 | 67612.75 | 7309310 | 6543802 | 6360459 | 5195690 |
| 80976.26 | 8107261 | 8493049 | 8559950 | 108875.2 | 4357412 | 4393955 | 4278334 | 335722.1 |
| 181595.2 | 6729406 | 8405599 | 8901907 | 80555.66 | 5281344 | 5181749 | 6447257 | 2920721 |
| 1242082 | 14377295 | 96188247 | 67357298 | 4543975 | 17740520 | 24749657 | 16083176 | 17860746 |
| 66531.8 | 9872635 | 9305966 | 10500953 | 76842.51 | 9598123 | 8734343 | 8877002 | 3541919 |
| 252011.1 | 2529699 | 3379079 | 2622373 | 230980.5 | 5483718 | 6092732 | 3848843 | 1417904 |
| 137389 | 4195716 | 563534.9 | 4929152 | 227519.4 | 11874674 | 10536853 | 9660811 | 6496528 |
| 249068.7 | 18595317 | 11634354 | 12145532 | 124199.1 | 10231109 | 11361179 | 13911550 | 8120317 |
| 589791.2 | 14508198 | 14983790 | 15592603 | 849606.1 | 30727783 | 28806261 | 21782059 | 14834118 |
| 4915688 | 55800317 | 18579795 | 27393392 | 980462 | 12229995 | 68734030 | 8368186 | 7033322 |
| 68202.16 | 9460598 | 7238983 | 6596359 | 73802.06 | 6376070 | 5283354 | 4498673 | 296850.9 |
| 431616.1 | 1165775 | 1158757 | 1195949 | 342353.1 | 2773241 | 2436715 | 3066930 | 3548228 |
| 96979.88 | 606054.8 | 2952889 | 1377977 | 102090 | 6142246 | 4122512 | 5706089 | 577880.5 |
| 210561.2 | 5771286 | 8226945 | 8659383 | 157511.3 | 19114286 | 11897899 | 10431566 | 11965772 |
| 332866.3 | 464517.6 | 976665 | 2421887 | 122115.8 | 6179180 | 2978711 | 4614957 | 11533283 |
| 241315.6 | 100807.6 | 228594.1 | 135343.9 | 33884.49 | 1246835 | 1119507 | 1626030 | 3763766 |
| 101168.2 | 3534521 | 6908048 | 7085572 | 101574.3 | 6710427 | 6267435 | 5054718 | 1385459 |
| 550969.5 | 3177000 | 4868816 | 2060471 | 228130.7 | 4225890 | 5876300 | 4186943 | 11228820 |
| 1187988 | 12552979 | 7211895 | 9402777 | 783527.2 | 28682459 | 26079679 | 12276138 | 2263031 |
| 86845.99 | 2043288 | 2694553 | 2433436 | 97147.49 | 7450039 | 8660843 | 7215152 | 2587842 |
| 213665 | 1496244 | 1472750 | 1324518 | 356639.7 | 1948449 | 2023379 | 2050576 | 85899.47 |

| | | | | | | | | |
|---|---|---|---|---|---|---|---|---|
| 83851.09 | 337297.7 | 559082.5 | 352887.1 | 91467.78 | 6735708 | 6216147 | 9113392 | 2644523 |
| 56263.19 | 558617.6 | 565859.7 | 447094.6 | 58548.69 | 2591096 | 2518540 | 1857177 | 5481289 |
| 1158799 | 3183651 | 3772107 | 5373900 | 880824.3 | 8048248 | 6615372 | 6020742 | 521511.6 |
| 176329.1 | 10675960 | 8992616 | 7250196 | 168835.9 | 15306032 | 14302246 | 13046606 | 16024895 |
| 86945.07 | 10254806 | 7332165 | 6845002 | 90055.42 | 6177545 | 6601669 | 5171437 | 2680914 |
| 75201.56 | 14521826 | 14555268 | 14704669 | 83602.91 | 15220873 | 11329432 | 10571556 | 4295234 |
| 149002.7 | 1432115 | 4548165 | 1830170 | 47501.59 | 3154552 | 2449157 | 2916308 | 6993395 |
| 48188.73 | 6838837 | 12499404 | 10792155 | 55342.8 | 8608491 | 10172051 | 19204724 | 824339.7 |
| 114771.4 | 15317056 | 15313301 | 15585467 | 115576 | 15920214 | 15317233 | 14343216 | 6115125 |
| 131045 | 6193561 | 6387276 | 6209103 | 83044.85 | 3191900 | 2423821 | 3151787 | 3247788 |
| 161298.2 | 7612087 | 7562372 | 7809537 | 226942.4 | 11631979 | 10055882 | 12297013 | 3331044 |
| 400190.2 | 45082465 | 12714922 | 25215875 | 202349.6 | 17488078 | 24843704 | 13327667 | 11622711 |
| 156370.3 | 219546.2 | 201355.8 | 207760.2 | 172554.3 | 7180977 | 6373832 | 6323981 | 16542540 |
| 64933.31 | 9889308 | 9916131 | 10830675 | 67365.99 | 4045609 | 3617111 | 3305285 | 3595631 |
| 73271.19 | 5814196 | 4607740 | 4889066 | 78683.95 | 6436265 | 6101669 | 6490439 | 6341713 |
| 1768873 | 11504292 | 11621243 | 15239315 | 180334.8 | 14671542 | 13161960 | 14149968 | 10486504 |
| 72734.86 | 4731275 | 4136083 | 4802992 | 76645.98 | 6059465 | 4753912 | 5617215 | 2280848 |
| 127913.4 | 13908991 | 11031873 | 10995178 | 128265.2 | 15037195 | 11915684 | 9879339 | 7073211 |
| 154144.6 | 12706436 | 14041770 | 13757226 | 141473.8 | 14719205 | 12722421 | 11277689 | 8749702 |
| 425765.1 | 442097.5 | 515536.3 | 402557.3 | 98910.8 | 6185946 | 4925256 | 5800713 | 366344.8 |
| 66607.54 | 6783412 | 4271195 | 3724951 | 70222.9 | 2862792 | 2834612 | 2520132 | 99965.47 |
| 69734.02 | 982539.3 | 9530064 | 676558.8 | 134762.6 | 7176396 | 9834843 | 9653708 | 1176419 |
| 51756.71 | 10761239 | 8906519 | 7822161 | 73264.76 | 7971643 | 6320362 | 6780120 | 3782271 |
| 121783.2 | 8872507 | 9712725 | 9485671 | 122712.9 | 10894390 | 9279546 | 8166135 | 4984132 |
| 79489.07 | 311185.5 | 147267.8 | 1345553 | 93848.31 | 3526886 | 4409781 | 5011776 | 187091.1 |
| 26019.72 | 620098.2 | 2094470 | 1799900 | 88875.18 | 1792777 | 5494583 | 559271 | 351871.6 |
| 64045.86 | 5976996 | 5157795 | 3062261 | 75682.33 | 8079678 | 6620405 | 9454611 | 1666181 |
| 135296.3 | 11836823 | 11164561 | 10792711 | 130723.7 | 5032699 | 3646354 | 3704061 | 130354.5 |
| 73983.66 | 7515762 | 6440638 | 8333748 | 76090.15 | 12515368 | 10573176 | 11497961 | 2726248 |
| 75832.53 | 3420509 | 2947702 | 2171938 | 100545.3 | 3706158 | 3794034 | 5097787 | 8318826 |
| 1481696 | 3787725 | 3869463 | 577650.9 | 184738.2 | 11244042 | 8900277 | 9383437 | 15083157 |
| 96679.11 | 9886446 | 8514473 | 8625771 | 94264.63 | 7217733 | 6344701 | 6259898 | 227631.8 |
| 125766.4 | 2317556 | 2547732 | 2327187 | 140604.5 | 6869238 | 5818852 | 4090045 | 143354.3 |
| 373809.3 | 29944786 | 29364940 | 29433641 | 518531.7 | 28081656 | 29478022 | 28024883 | 34524918 |
| 244665.8 | 1205778 | 11604498 | 7719191 | 233836 | 6491149 | 13325497 | 8645553 | 6295444 |
| 87945.12 | 1038248 | 794165.2 | 812117.1 | 98374.94 | 8712236 | 2910636 | 3543319 | 3036247 |
| 92401.11 | 1110274 | 3192387 | 3510005 | 127611.4 | 4719912 | 4385624 | 3269483 | 785499.5 |
| 56757.72 | 1295224 | 627431.8 | 1135459 | 55288 | 3287737 | 2692138 | 2676730 | 10605703 |
| 160864.9 | 14577215 | 13981251 | 11745600 | 167039.7 | 14718977 | 12811066 | 11062895 | 9849280 |
| 72446.88 | 2290914 | 2820633 | 2889348 | 90555.75 | 3891456 | 3871454 | 3457169 | 1430619 |
| 118278.6 | 10828530 | 10189053 | 10058579 | 124511 | 11675711 | 9955928 | 10935888 | 10860928 |
| 86001.1 | 8612674 | 5758680 | 6827085 | 82548.41 | 5273549 | 5591613 | 4450125 | 1599963 |
| 365633.4 | 736413.8 | 9778244 | 2310926 | 139297 | 28818655 | 12503452 | 20622028 | 14273609 |
| 272861.9 | 7656019 | 7255272 | 7635412 | 545920.5 | 7762107 | 6648523 | 6420680 | 2727392 |
| 75876.02 | 2610516 | 3140031 | 3363262 | 51004.03 | 6188604 | 4625235 | 7118368 | 1236252 |
| 43044 | 188412.9 | 349982.5 | 209318.2 | 51747.74 | 2054896 | 1769535 | 2056567 | 6424007 |
| 358787.3 | 10972287 | 4629314 | 10491537 | 350267.1 | 12559415 | 19360220 | 15008195 | 4802039 |
| 121900.7 | 5065722 | 5188258 | 4586457 | 121432.6 | 5968437 | 5378260 | 5557887 | 8722354 |
| 72925.16 | 5653436 | 5097157 | 4097746 | 78312.36 | 4146160 | 3706653 | 2966295 | 80345.4 |
| 189899.9 | 9334958 | 8379187 | 7590104 | 95843.36 | 3288632 | 4115884 | 4737997 | 905945.7 |
| 90455.22 | 503052.9 | 1691728 | 761570.5 | 100417.2 | 4368368 | 4326028 | 4255849 | 947908.3 |

| | | | | | | | | |
|---|---|---|---|---|---|---|---|---|
| 83446.25 | 8272277 | 7442537 | 6139418 | 92076.53 | 5191246 | 4382491 | 3614268 | 100112.2 |
| 118936.6 | 4556653 | 5618403 | 5489123 | 125203.7 | 7612888 | 6827169 | 6042115 | 1508408 |
| 690136.3 | 20880196 | 18125109 | 25199391 | 197243.8 | 27950122 | 22197398 | 14123918 | 9211722 |
| 151376.7 | 19027199 | 19182011 | 18859218 | 166433.4 | 14979868 | 6117230 | 14139679 | 10702826 |
| 649115.9 | 8971431 | 24046093 | 20818151 | 321723.5 | 31770987 | 18135223 | 19641513 | 22516011 |
| 48767.13 | 2594083 | 2764314 | 2438211 | 128912.1 | 3653166 | 2938648 | 3380088 | 1000518 |
| 105862.9 | 2420727 | 7288670 | 869436.1 | 170321.3 | 5641678 | 3119758 | 2668269 | 2484956 |
| 76876.54 | 6073829 | 5649213 | 5635021 | 82907.7 | 5508688 | 4929236 | 4670539 | 1875712 |
| 172917.7 | 1164031 | 618701.8 | 561968.6 | 252291.3 | 4798297 | 3663076 | 4361720 | 782164.3 |
| 74672.86 | 241508 | 182044.5 | 174116.1 | 100787.9 | 1451181 | 1439853 | 1640215 | 5127282 |
| 81318.98 | 155582.6 | 151415.5 | 147669.4 | 88825.44 | 2379565 | 2066613 | 1195466 | 117449.2 |
| 88541.98 | 2137261 | 118730.6 | 1179803 | 91940.66 | 4330717 | 3789485 | 2827630 | 7150218 |
| 150647.2 | 3029010 | 5570102 | 6054343 | 153225.4 | 8444281 | 6958096 | 4652001 | 2781213 |
| 157265 | 16517687 | 15728319 | 4373337 | 324879.6 | 11044851 | 9202642 | 10214929 | 9427667 |
| 164212.4 | 13621406 | 14099674 | 14266666 | 161355.3 | 12075510 | 9942649 | 9604718 | 1438004 |
| 451748.3 | 6605922 | 6500306 | 6304726 | 833251.8 | 10205317 | 8740865 | 9277432 | 16059693 |
| 104818.7 | 2005945 | 2298392 | 2953588 | 136688.7 | 4021636 | 3468584 | 3688586 | 1197066 |
| 126922.7 | 10715920 | 10061293 | 11055684 | 168614.6 | 15783268 | 13227899 | 14298803 | 4096392 |
| 76348.3 | 5986504 | 5482796 | 5533117 | 86746.5 | 4580505 | 4559836 | 4696108 | 3099360 |
| 86119.48 | 433590.7 | 678652.5 | 1721205 | 114336.1 | 7745516 | 5142093 | 2863184 | 2120142 |
| 100572.3 | 346062.2 | 565852 | 610553.3 | 109923.8 | 1377034 | 4138412 | 2748316 | 6669231 |
| 112629.8 | 4600728 | 4473521 | 5556866 | 125750.7 | 3243558 | 3103421 | 2839962 | 459874.6 |
| 215226 | 8265385 | 6536423 | 6056611 | 260064 | 11908083 | 6731028 | 5622274 | 2756866 |
| 363886.7 | 1511800 | 1752245 | 146940.7 | 100820.8 | 1838829 | 2606789 | 2302894 | 1235714 |
| 432357 | 3446980 | 3964796 | 533813.7 | 119398.4 | 8225158 | 3534336 | 3031702 | 10507907 |
| 84226.93 | 558545.2 | 529700.6 | 1831387 | 89336.32 | 2239598 | 2654315 | 3583695 | 314998.7 |
| 78737.52 | 3600254 | 3797703 | 4174457 | 88552.32 | 4715575 | 3823361 | 3780871 | 2037112 |
| 120317.8 | 11895482 | 11262815 | 12480759 | 187143.4 | 6812520 | 15395721 | 6477053 | 6392817 |
| 181771.8 | 1596625 | 789218.9 | 483864.5 | 152104.5 | 11524364 | 9856306 | 10249437 | 28141810 |
| 74035.35 | 7286088 | 7204083 | 7041945 | 76616.55 | 4697219 | 3560623 | 4360653 | 266250.6 |
| 95038.5 | 3706560 | 8148400 | 2688854 | 118127.3 | 2154900 | 4453952 | 3822621 | 2058891 |
| 98571.53 | 6885359 | 5866237 | 5566595 | 96116.27 | 8262346 | 7687412 | 5977177 | 6639978 |
| 99908.21 | 193166.5 | 196692.2 | 195606.6 | 111069.7 | 2881160 | 2587143 | 3092556 | 157358.2 |
| 87768.63 | 7884971 | 6228134 | 6269109 | 96759.32 | 6320812 | 5428064 | 4646843 | 1149858 |
| 143220.3 | 8858715 | 8344320 | 9225636 | 159348.1 | 10922446 | 5641819 | 9114239 | 4352487 |
| 115687.4 | 3670175 | 3696284 | 3177785 | 112986.3 | 3010727 | 2105477 | 2502899 | 798071.3 |
| 97841.93 | 9098616 | 7470894 | 6661616 | 95891.51 | 5664223 | 4817219 | 3840061 | 1451944 |
| 197089.7 | 5197107 | 10454063 | 2992546 | 236862.3 | 9865321 | 8808473 | 9480217 | 8211444 |
| 59168.7 | 589040.4 | 297915.5 | 318069 | 65577.83 | 2104453 | 1493723 | 1168322 | 498952.7 |
| 78357.86 | 6115007 | 4783084 | 4402385 | 86879.7 | 3875717 | 3736973 | 3481430 | 1273857 |
| 92376.02 | 8416484 | 6841448 | 5560840 | 89184.28 | 6126499 | 5782767 | 3293418 | 1629449 |
| 117540.2 | 7936841 | 8669409 | 8605301 | 124069.2 | 6964684 | 6601444 | 5398771 | 2903644 |
| 226542.7 | 1578761 | 1643915 | 1760328 | 169524.8 | 5374260 | 3501342 | 5520902 | 9508196 |
| 94370.8 | 2013734 | 1807937 | 1712786 | 140411.8 | 5694275 | 5593060 | 6506325 | 835510.4 |
| 89833.92 | 8274465 | 7468453 | 7630323 | 84900.42 | 5751229 | 6217070 | 6903154 | 3183096 |
| 85002.51 | 4702050 | 5023034 | 5113798 | 96443.08 | 4075973 | 4062681 | 4141360 | 597109.5 |
| 42539.89 | 1690887 | 2239002 | 2766665 | 52082.93 | 4122621 | 1486700 | 1502472 | 586255.9 |
| 690136.3 | 20880196 | 18125109 | 25199391 | 1249329 | 27950122 | 22197398 | 14123918 | 2118551 |
| 73129.25 | 776410.3 | 295976.7 | 245846.7 | 90807.35 | 2729021 | 2170303 | 2295181 | 334899.7 |
| 131474.7 | 7708386 | 12528492 | 8654225 | 137753.9 | 10758350 | 6279927 | 6060652 | 4168256 |
| 136888.4 | 4996509 | 8054907 | 9543408 | 94389.32 | 4460307 | 4451018 | 7893524 | 2482901 |

| | | | | | | | | |
|---|---|---|---|---|---|---|---|---|
| 156084 | 5303126 | 5297433 | 5914499 | 118926.9 | 7330939 | 7443006 | 4537648 | 3759208 |
| 158413.2 | 60821.15 | 240719.8 | 100020.1 | 31295.91 | 900174.6 | 879708.6 | 1153699 | 2501922 |
| 122590.9 | 706920.8 | 1364415 | 1283287 | 63735.25 | 4889696 | 6290097 | 4853909 | 346679.2 |
| 90868.66 | 4071731 | 2120654 | 3111104 | 88380.75 | 4062960 | 4401260 | 3082436 | 212948.9 |
| 76877.03 | 229383.3 | 230984.4 | 348030.1 | 79227.37 | 1860875 | 1980806 | 2096974 | 4463013 |
| 112542.1 | 7181548 | 10620121 | 10459421 | 135754.4 | 7791287 | 9144910 | 8336614 | 13977412 |
| 1005636 | 37711654 | 33493612 | 53200104 | 1232510 | 56622348 | 50992731 | 47398024 | 42801459 |
| 105297.4 | 549845.9 | 1099492 | 232404.3 | 98796.05 | 6536738 | 5866930 | 6448093 | 265376.1 |
| 93437.96 | 1048588 | 1054581 | 711803.1 | 99091.23 | 2806267 | 2312789 | 2446426 | 108869 |
| 160492.4 | 4513381 | 4835428 | 1027361 | 142691.5 | 5711301 | 2333032 | 5824132 | 3788106 |
| 50090.72 | 488230.1 | 1010305 | 302654 | 60037.79 | 5151527 | 4716353 | 2073058 | 429135 |
| 86480.02 | 6778392 | 4430489 | 6461148 | 90716.76 | 4733808 | 4089623 | 4738570 | 531027.7 |
| 104631.4 | 575613.7 | 327574.1 | 557689.2 | 109938.9 | 3392794 | 1338811 | 2670167 | 5974412 |
| 46354.57 | 2135347 | 2334083 | 2373515 | 56859.67 | 2763407 | 2591968 | 2696485 | 498411.4 |
| 75270.18 | 2237942 | 2417038 | 2016032 | 83824.36 | 2551148 | 2675006 | 2674536 | 3537779 |
| 172427.2 | 11105309 | 8887875 | 8143535 | 160172.9 | 11060970 | 9565651 | 2253679 | 4254165 |
| 69143.56 | 5536729 | 3044780 | 3012449 | 70758.63 | 4480383 | 3839188 | 2858774 | 2666818 |
| 178263.9 | 500441 | 534787.2 | 587878.3 | 117458.7 | 2215428 | 2082106 | 1355265 | 441341.7 |
| 82151.57 | 3588478 | 3754919 | 3764316 | 98102.27 | 3818241 | 3847512 | 3744375 | 1575131 |
| 589759.4 | 2995336 | 6617952 | 7590740 | 1255057 | 6192520 | 11605154 | 13373004 | 3125146 |
| 77964.63 | 4090937 | 2779737 | 4126123 | 81919.4 | 3529882 | 3288159 | 3347141 | 1874798 |
| 112622.6 | 3073176 | 3206492 | 1939721 | 126672.6 | 6378857 | 6379209 | 4987720 | 1509612 |
| 78613.08 | 8596593 | 3880098 | 3776261 | 81871.56 | 3701107 | 3520707 | 3682450 | 3208465 |
| 48537.07 | 1341718 | 1714533 | 1675954 | 93633.53 | 2103310 | 2044664 | 2091942 | 178324.7 |
| 94495.92 | 372725.9 | 445253.1 | 1457240 | 75289.79 | 4020591 | 3752315 | 2097810 | 6621698 |
| 68596.48 | 4440530 | 3627382 | 3325312 | 74718.6 | 3124134 | 2874117 | 2580273 | 97293.94 |
| 352173.6 | 2166699 | 4923303 | 2525263 | 483429.8 | 12142502 | 14360187 | 13990222 | 3574731 |
| 137724.8 | 3864271 | 3025085 | 3257486 | 122584.5 | 4595845 | 4370977 | 3820500 | 4913591 |
| 87282.38 | 582107.3 | 745188.7 | 356072.9 | 95700.39 | 2986723 | 2843991 | 7226727 | 430149 |
| 49312.38 | 1090511 | 1319688 | 1179216 | 53630.19 | 1496432 | 1428500 | 1312007 | 66363.52 |
| 94129.41 | 182058 | 180905.4 | 93560.35 | 97901.12 | 2444628 | 2550325 | 2499612 | 97172.22 |
| 2253249 | 15763360 | 38229184 | 26342244 | 5070914 | 44229842 | 36555859 | 39693414 | 22372109 |
| 875765.3 | 26366422 | 11385349 | 12091832 | 873706.3 | 10646845 | 10805028 | 10406610 | 17748559 |
| 586588 | 11981323 | 11747735 | 10311839 | 644661.9 | 9881758 | 13022781 | 10153365 | 6268337 |
| 2054858 | 8496745 | 8498359 | 8616430 | 2048330 | 9760790 | 11228765 | 8501854 | 5544874 |
| 3126560 | 30239363 | 30240983 | 31083365 | 3340334 | 33906502 | 30391560 | 7531993 | 27284395 |
| 3591347 | 20564260 | 23520840 | 35219108 | 3550934 | 56205818 | 34446230 | 36327401 | 28813368 |
| 1317092 | 8671937 | 9737863 | 7342921 | 3186197 | 23763799 | 23956413 | 25447205 | 5215564 |
| 1162471 | 6286239 | 6211052 | 13398172 | 854210.7 | 13463733 | 6006225 | 5828829 | 5445687 |
| 1323805 | 3180047 | 2724554 | 2883423 | 556138.5 | 14020162 | 11252327 | 15216640 | 34001350 |
| 2606693 | 38839689 | 11515580 | 20854947 | 698925.3 | 13543287 | 33304490 | 95449173 | 6466239 |
| 2963146 | 22446802 | 22023364 | 21938753 | 312454.1 | 16942056 | 14696465 | 29244946 | 26891284 |
| 114361.6 | 79804.33 | 157238.8 | 72903.09 | 29791.81 | 1406407 | 1314911 | 1459935 | 3398087 |
| 87721.72 | 330239.9 | 279942 | 128429.4 | 93750.74 | 3494200 | 2979358 | 3294293 | 9919092 |
| 52857344 | 18388213 | 32846013 | 10617406 | 92571411 | 8.54E+08 | 7.54E+08 | 7.36E+08 | 6248400 |
| 1781149 | 5239943 | 5102164 | 4049333 | 3446904 | 83876064 | 72710325 | 79642265 | 2703747 |
| 592078 | 21566640 | 23925844 | 22728487 | 114561.5 | 23063987 | 20974796 | 21593871 | 14224219 |
| 1347585 | 1874724 | 2936112 | 3302147 | 1514405 | 10853804 | 11393101 | 9202157 | 25911020 |
| 90859.47 | 903453.1 | 1449287 | 1240751 | 115863.4 | 7802572 | 6629580 | 6125410 | 566144 |
| 1207078 | 94147694 | 12052901 | 13539675 | 1945254 | 18747245 | 53391552 | 26962256 | 16102117 |
| 420316 | 10173247 | 10202096 | 11191547 | 241196.1 | 9946637 | 5660934 | 7756803 | 3913872 |

| | | | | | | | | |
|---|---|---|---|---|---|---|---|---|
| 83024.23 | 12573272 | 12847396 | 14382839 | 105962.5 | 14047022 | 10940867 | 12248285 | 7027962 |
| 5858305 | 1364579 | 2851955 | 1236105 | 11917446 | 1.79E+08 | 1.62E+08 | 1.7E+08 | 901154.7 |
| 934232.7 | 11592837 | 12281221 | 38862941 | 1354906 | 18319647 | 66715200 | 68194677 | 33892209 |
| 302081.5 | 20297281 | 20393269 | 20180091 | 341527.3 | 15033423 | 11472928 | 16818724 | 8086090 |
| 83437.31 | 10012352 | 8679052 | 9577284 | 91798.62 | 9265676 | 8185385 | 8459111 | 3137776 |
| 295566.9 | 1581972 | 5158265 | 2479054 | 267240.8 | 2664688 | 5860210 | 2451096 | 4089120 |
| 1055898 | 21831687 | 25940691 | 20833634 | 2023057 | 18417850 | 20915642 | 21140626 | 13920182 |
| 145359.2 | 17141914 | 21977019 | 21698373 | 118284.8 | 3085949 | 6485426 | 5305941 | 176905.6 |
| 113690.2 | 274908.9 | 288487 | 493891.9 | 124260.8 | 5886282 | 5867193 | 7686709 | 261964.7 |
| 73744.69 | 4616260 | 4400290 | 4801554 | 105518.4 | 2051193 | 1836091 | 1816814 | 584121.9 |
| 247096.5 | 242490.9 | 4383435 | 2778727 | 424717.8 | 4791184 | 3071902 | 3400758 | 263867.9 |
| 884543.4 | 520896.2 | 438793.1 | 290146.3 | 1595398 | 25773597 | 22506505 | 24752221 | 2740109 |
| 58532.25 | 633712.2 | 539219 | 669720.2 | 69149.32 | 2075178 | 1377653 | 1579336 | 118290.6 |
| 183454.7 | 18639756 | 19921082 | 20752992 | 672807.7 | 18546778 | 15524165 | 15849531 | 208260.3 |
| 152134.8 | 7781619 | 25520826 | 23445831 | 181502.1 | 5128916 | 30601286 | 30750197 | 29114668 |
| 227882.2 | 3182534 | 7746540 | 7626704 | 249071.2 | 16191213 | 10842177 | 15789858 | 6999403 |
| 828803.4 | 19425239 | 21267513 | 21182300 | 248366.6 | 18826078 | 15321644 | 12461159 | 8846772 |
| 211244.7 | 19687601 | 18083395 | 18417232 | 78220.45 | 18705855 | 16066816 | 17132058 | 9919863 |
| 409071.6 | 16093466 | 20422516 | 23619368 | 184884.5 | 19602325 | 20708527 | 22459199 | 12604746 |
| 88363.73 | 20727244 | 16581271 | 18874613 | 92054.38 | 22442529 | 17681597 | 20280066 | 14890336 |
| 92586.66 | 18395237 | 14442179 | 18512727 | 92013.52 | 18721971 | 16545617 | 18740409 | 13467153 |
| 125412.4 | 2751587 | 2838870 | 300196.1 | 156545.4 | 13010198 | 15080566 | 14382066 | 2713331 |
| 708514.2 | 17225935 | 13327950 | 6426617 | 365490.8 | 16953895 | 10096757 | 9434142 | 4917207 |
| 600327.1 | 4195096 | 528197.1 | 3002726 | 114108.3 | 3679288 | 4042653 | 1790817 | 500008.6 |
| 140749 | 6865749 | 8479153 | 9854677 | 179566 | 13356351 | 14440136 | 15072416 | 4992165 |
| 87460.26 | 32282232 | 26183889 | 26297221 | 91071.32 | 29045559 | 26233821 | 25587290 | 14388890 |
| 57090.19 | 3088114 | 2988413 | 2798137 | 97879.07 | 5917210 | 5333309 | 5062680 | 1468572 |
| 186740.7 | 29694551 | 25519615 | 25783976 | 578800.6 | 24913870 | 22283238 | 22583295 | 10472524 |
| 93649.25 | 375438.3 | 1234261 | 1599718 | 266639.2 | 9309660 | 5187202 | 8175591 | 196687.7 |
| 164230.1 | 2968780 | 2629457 | 2732707 | 1480702 | 22932469 | 17389049 | 25813313 | 9230889 |
| 120963.2 | 216115.5 | 216181.8 | 216825.1 | 128304.3 | 4294947 | 3561855 | 2684986 | 11306739 |
| 185429.8 | 678218.5 | 457784.8 | 723339 | 193545 | 7765075 | 7137308 | 8021655 | 21278255 |
| 249001.3 | 1011335 | 1173630 | 1173548 | 252934.9 | 6740964 | 5667758 | 5885252 | 13408568 |
| 180232.3 | 553938.2 | 918678.5 | 1439125 | 392565 | 3334286 | 4352019 | 4091745 | 405178.5 |
| 129996.1 | 13166254 | 10538120 | 13070933 | 173819.8 | 24753976 | 22643828 | 22913749 | 8586413 |
| 143397.2 | 15157723 | 2253682 | 15589523 | 162187.8 | 18744121 | 13456426 | 27309927 | 7439233 |
| 185155.7 | 33959761 | 29884630 | 30918073 | 82202.17 | 23903266 | 20986926 | 20704305 | 11789434 |
| 448642.2 | 13397657 | 15508900 | 14343342 | 377819.3 | 8822082 | 9319800 | 9261312 | 2752107 |
| 150997.5 | 16979730 | 17634825 | 16661346 | 158379.2 | 18009932 | 12568877 | 15750917 | 3882167 |
| 89078.28 | 940490.6 | 427322.8 | 391106.2 | 112888.9 | 8114283 | 8143426 | 8660410 | 2422351 |
| 1022617 | 43366633 | 50440901 | 6251009 | 1758326 | 50306626 | 32965921 | 41358100 | 4923773 |
| 122907.9 | 7450195 | 4542744 | 6843867 | 96073.21 | 4290261 | 3968623 | 3170705 | 2872089 |
| 147343.8 | 66105.27 | 63383.07 | 74404.51 | 176861.2 | 1064702 | 1051401 | 1390243 | 4214199 |
| 87961.59 | 163059.7 | 156691.1 | 156994.9 | 103318.5 | 3547991 | 3665585 | 4257458 | 140136.7 |
| 12768319 | 1.48E+08 | 1.55E+08 | 1.48E+08 | 12198382 | 1.56E+08 | 1.4E+08 | 1.38E+08 | 1.36E+08 |
| 10074628 | 1.04E+08 | 1.21E+08 | 1.16E+08 | 10782490 | 1.23E+08 | 1.2E+08 | 1.12E+08 | 1.15E+08 |
| 308594.7 | 8426750 | 6563745 | 5796408 | 1765633 | 22932469 | 17389049 | 25813313 | 39259557 |
| 165013.1 | 15581300 | 16526655 | 15323851 | 177203.1 | 11578630 | 9544073 | 9602613 | 4640681 |
| 75210.62 | 868082.2 | 220051.9 | 297100.9 | 94010.32 | 3599380 | 3079676 | 1945138 | 6379705 |
| 92473.51 | 958743.3 | 995493.5 | 383817.1 | 90164.06 | 16559441 | 14942124 | 16155655 | 2030761 |
| 2226131 | 2.83E+08 | 2.55E+08 | 2.72E+08 | 2131421 | 2.86E+08 | 2.15E+08 | 2.5E+08 | 94765709 |

| | | | | | | | | |
|---|---|---|---|---|---|---|---|---|
| 83850.79 | 12644554 | 11295949 | 8586710 | 90045.1 | 9243703 | 7335484 | 6814507 | 2885824 |
| 80281.05 | 11532400 | 8778254 | 6327697 | 82333.05 | 7352138 | 7008945 | 4916229 | 2048432 |
| 93071.3 | 2960723 | 2850528 | 3098943 | 108334.3 | 5181473 | 4075741 | 4270613 | 6004318 |
| 214949.4 | 989053.3 | 1109928 | 1246142 | 381669.9 | 3748991 | 2445463 | 2867036 | 115503.1 |
| 460172.3 | 14362860 | 16305260 | 16074548 | 910702.9 | 15490745 | 16300662 | 16804981 | 7144739 |
| 85537.86 | 3915849 | 3080652 | 2126599 | 68726.5 | 2816582 | 4000320 | 4143412 | 4124488 |
| 138688.6 | 1.81E+08 | 1.52E+08 | 1.56E+08 | 125165.1 | 1.43E+08 | 1.33E+08 | 1.21E+08 | 43453848 |
| 803426.8 | 18925513 | 21038436 | 22325723 | 208052.8 | 7819128 | 8065393 | 6661454 | 224249.8 |
| 2519166 | 25508100 | 29673417 | 34874992 | 2507917 | 27896308 | 23357142 | 24599071 | 4773100 |
| 101301.3 | 4114695 | 5029734 | 4157335 | 108641 | 4601974 | 3809423 | 4680328 | 2189107 |
| 6298538 | 1.59E+08 | 77265541 | 82175358 | 7879004 | 95428576 | 83521118 | 94189025 | 45942315 |
| 2558204 | 61130340 | 63573796 | 61372719 | 4389361 | 61035661 | 56834571 | 71755486 | 34495105 |
| 3170694 | 70816363 | 72752091 | 66820208 | 11449831 | 70894023 | 73108775 | 73125072 | 32827151 |
| 685061.7 | 10013242 | 9826613 | 9285947 | 733082.7 | 10394726 | 10143000 | 10940654 | 5590388 |
| 685061.7 | 9096403 | 9623326 | 8996400 | 1028228 | 8976478 | 9406596 | 10542971 | 4673441 |
| 704030.8 | 12772287 | 13548902 | 12414692 | 681066.5 | 13051741 | 11100375 | 9734560 | 7514069 |
| 89054.72 | 6590019 | 6429095 | 6663148 | 88842.53 | 6607262 | 6140803 | 6345947 | 5119259 |
| 350504.7 | 11468831 | 12773092 | 11480974 | 358705.5 | 15167559 | 23804130 | 10374587 | 6760018 |
| 1531951 | 41062345 | 9464048 | 10642954 | 372186.8 | 10450717 | 17470106 | 9214020 | 12379391 |
| 401999.3 | 8660024 | 10883753 | 9802574 | 353788.4 | 2472940 | 3021245 | 2471649 | 309639.6 |
| 130140.9 | 9341668 | 9007052 | 11556395 | 149569.2 | 4039979 | 3478600 | 3647039 | 691016.9 |
| 232204.9 | 19766566 | 17392093 | 16338483 | 368856.4 | 24716782 | 6148066 | 26570301 | 7322825 |
| 165645.3 | 2128767 | 3780035 | 3129410 | 84386.3 | 2853039 | 2674673 | 2718390 | 520181.8 |
| 114725 | 11819049 | 12423854 | 12388880 | 153583.2 | 10020533 | 9778065 | 13974369 | 8094453 |
| 832489.9 | 58658560 | 59642542 | 61347513 | 1136235 | 1.14E+08 | 96912796 | 93367449 | 32705922 |
| 196098.3 | 1871111 | 1992748 | 304539.3 | 133298 | 3227625 | 2790205 | 3351642 | 235581.4 |
| 88124.04 | 12689175 | 11463579 | 13024421 | 91685.42 | 15381204 | 11630683 | 14871940 | 20612400 |
| 146811.9 | 5312504 | 6704434 | 5804539 | 159042.7 | 7353633 | 6512379 | 5195223 | 658839.3 |
| 275768.4 | 11383796 | 12794415 | 12073970 | 295799.5 | 12242177 | 11461013 | 11813742 | 4640883 |
| 95941.42 | 288760.6 | 690417 | 927124.6 | 300964.4 | 2892400 | 3119584 | 3688398 | 761316.8 |
| 1014445 | 1.47E+08 | 1.47E+08 | 1.46E+08 | 1833712 | 1.46E+08 | 1.36E+08 | 1.43E+08 | 44713032 |
| 223092.1 | 2789318 | 4964929 | 3436254 | 217952.4 | 2206548 | 1834689 | 2106405 | 619286.8 |
| 71812.7 | 7516874 | 13212601 | 6797438 | 72446.61 | 4849910 | 4085320 | 2946899 | 182238 |
| 119509.9 | 11388825 | 12335346 | 11531758 | 213004.8 | 13345910 | 11643498 | 12050299 | 2634062 |
| 97254.95 | 19780895 | 16496801 | 16216067 | 102991.4 | 13788524 | 12306756 | 13107460 | 5986492 |
| 734628.1 | 769417.6 | 601941.9 | 844415.5 | 488932.6 | 30351490 | 28906188 | 26263098 | 362087.5 |
| 308594.7 | 9211281 | 6563745 | 5796408 | 1480702 | 22932469 | 17389049 | 25813313 | 39259557 |
| 218029.8 | 29252996 | 25949950 | 25465684 | 366816.6 | 30139713 | 26764107 | 27143596 | 6685020 |
| 1349308 | 11531886 | 12314113 | 11692085 | 1298527 | 11652134 | 10955236 | 10810147 | 2689437 |
| 84668.03 | 4061076 | 2501287 | 2773287 | 131719.4 | 6567779 | 5373687 | 5472026 | 218180.6 |
| 149745.4 | 355506.7 | 151640.5 | 131776.8 | 245076.2 | 1809371 | 2011855 | 4459308 | 144180.6 |
| 398595.4 | 11142178 | 11320712 | 10346667 | 286355.7 | 14354290 | 14138098 | 15373944 | 21928107 |
| 116011.7 | 8935989 | 7937632 | 7868638 | 83299.99 | 5983711 | 5480118 | 5955650 | 842857.8 |
| 87794.6 | 17371194 | 14121269 | 13774507 | 97095.72 | 10798664 | 9852715 | 9339741 | 173119.5 |
| 159981.7 | 9956794 | 9793547 | 9653829 | 122884.2 | 9465787 | 8275178 | 9117772 | 7578738 |
| 305109.5 | 4510708 | 6188793 | 4107327 | 199538.2 | 4421844 | 3860693 | 4517272 | 2640913 |
| 187832.3 | 27263857 | 28003595 | 28000744 | 104236.8 | 18271036 | 17468681 | 16411989 | 3518325 |
| 456466.8 | 25080108 | 32180557 | 32413225 | 110617.5 | 12333032 | 10021824 | 10146305 | 154590.6 |
| 90830.17 | 10200539 | 10521628 | 10108722 | 106994.1 | 9396155 | 5625496 | 8999165 | 1503905 |
| 213732.2 | 31930593 | 29718698 | 29049833 | 255415.3 | 25412911 | 23640715 | 23405280 | 9564525 |
| 105901.8 | 2913126 | 780435.2 | 1696281 | 108932.6 | 6291569 | 1847225 | 2913628 | 10381458 |

| | | | | | | | | |
|---|---|---|---|---|---|---|---|---|
| 58175.39 | 3088114 | 2988413 | 2798137 | 97879.07 | 5917210 | 5333309 | 5062680 | 1468572 |
| 59376.13 | 15568473 | 11499364 | 12037164 | 56952.19 | 9953574 | 10112700 | 8360646 | 3460864 |
| 1207665 | 19683537 | 23842812 | 23925677 | 1089795 | 28140082 | 22365231 | 22428905 | 13464323 |
| 102308.6 | 3206434 | 2833136 | 3188315 | 147835.4 | 34104486 | 29735276 | 31062361 | 1174025 |
| 90960.05 | 12235255 | 9336925 | 9739832 | 85058.74 | 8183316 | 7941310 | 6343821 | 1599830 |
| 328055.3 | 6022337 | 15003954 | 27835803 | 291884.9 | 11154006 | 7935848 | 11319986 | 8555641 |
| 1462559 | 15423073 | 12910100 | 24898237 | 2292824 | 19929253 | 23079948 | 9724402 | 15876584 |
| 1078007 | 5033432 | 8758084 | 5464914 | 148466.4 | 18968772 | 3936074 | 4269008 | 5531593 |
| 103079 | 13657141 | 10231028 | 10010793 | 91158.43 | 13203150 | 11402401 | 9996577 | 9420898 |
| 459408.3 | 8268349 | 8139849 | 7851657 | 920044.2 | 20903732 | 20558912 | 23713345 | 8871908 |
| 87300.35 | 1611704 | 707494.9 | 1443524 | 102836.1 | 3097702 | 2584676 | 2572565 | 4369178 |
| 964872.7 | 12956566 | 11890014 | 10811731 | 371505.2 | 15314750 | 15956133 | 15684979 | 5317070 |
| 572233.2 | 11606595 | 12482362 | 8741925 | 3058979 | 10474298 | 12473242 | 13080744 | 5457697 |
| 88119.83 | 1011148 | 1043468 | 641348.6 | 97721.38 | 2095733 | 3230861 | 3476287 | 6810737 |
| 108495.5 | 7827468 | 6316233 | 5155342 | 114268.9 | 6731813 | 6863313 | 10027503 | 2083603 |
| 82831.7 | 6556275 | 7091462 | 6567697 | 133792.9 | 7730402 | 10160141 | 9359550 | 6912465 |
| 1910012 | 5367941 | 5554673 | 3914022 | 313803.7 | 49181451 | 42527948 | 48602231 | 1.14E+08 |
| 73241.61 | 10900454 | 9221633 | 9827273 | 74151.2 | 8021180 | 6851593 | 7957936 | 2929100 |
| 61995.9 | 9761795 | 11486583 | 10165805 | 74155.81 | 8299500 | 6994248 | 7360035 | 1301629 |
| 142507 | 3943095 | 4656700 | 4002590 | 136205.5 | 4792404 | 3805432 | 3404684 | 1647112 |
| 1560532 | 9326805 | 30198601 | 21892879 | 636020.3 | 71558095 | 7755733 | 32591473 | 18691118 |
| 110039.7 | 3942827 | 3852178 | 5080602 | 117067 | 8300567 | 6907379 | 6720752 | 6152011 |
| 97268.16 | 10958119 | 11752387 | 10256872 | 98560.34 | 21445245 | 12138734 | 7481217 | 14148330 |
| 129939.7 | 19312702 | 11639338 | 9781654 | 129846.1 | 13829912 | 8346127 | 13500407 | 5176648 |
| 125359.7 | 10418332 | 12819650 | 12866444 | 134394.6 | 12220521 | 10945877 | 10710349 | 7809905 |
| 131494.2 | 6956555 | 6903722 | 6258156 | 154323.5 | 4697838 | 4736161 | 4701513 | 3295686 |
| 359798.3 | 27817975 | 28215979 | 28202657 | 219046.2 | 26219857 | 23445544 | 13787395 | 15881979 |
| 351223.7 | 158690.8 | 410586 | 381987.4 | 134761.9 | 1215207 | 1551776 | 2221207 | 3436287 |
| 91818.61 | 13376978 | 13813415 | 13965890 | 112163.6 | 8366543 | 8052315 | 7708398 | 3530646 |
| 107610.4 | 494517.6 | 3134324 | 887346.9 | 136518.7 | 9609475 | 7956512 | 10270554 | 1170077 |
| 146004.5 | 7437741 | 9452782 | 9436840 | 149790.9 | 10385386 | 9135945 | 9110429 | 6481208 |
| 590671.7 | 608263.4 | 1308863 | 696750.4 | 146190.3 | 13610568 | 13163718 | 14997584 | 32532757 |
| 562997.4 | 23409390 | 24248049 | 24925905 | 291280.3 | 15945472 | 15352312 | 15694101 | 16718658 |
| 7499643 | 53617349 | 33310022 | 54599014 | 7950793 | 22337050 | 20100784 | 22990236 | 12269195 |
| 3884855 | 63552349 | 51393346 | 1.31E+08 | 3752846 | 28409252 | 47943099 | 26231742 | 21828388 |
| 3975319 | 33814454 | 22632446 | 30119409 | 4289785 | 24225240 | 28459334 | 53803172 | 18623674 |
| 1653737 | 12822091 | 14094748 | 12927084 | 1670217 | 16573694 | 27760515 | 26040788 | 11648606 |
| 67028.22 | 1469244 | 243778.3 | 1299659 | 108284.8 | 3767930 | 3939502 | 2573264 | 143571.4 |
| 78785.04 | 14678873 | 13970381 | 14734557 | 90994.68 | 11562268 | 10618042 | 9260397 | 2491838 |
| 145788.1 | 9060738 | 25887140 | 15755479 | 261863.4 | 21397141 | 14144167 | 13657415 | 2490045 |
| 252685 | 3039067 | 2847395 | 667694.4 | 436840.9 | 11348297 | 7858374 | 8435215 | 477138.7 |
| 158334.6 | 539649.6 | 419818.5 | 417063.4 | 226554.7 | 5239945 | 4357106 | 4715250 | 371293.7 |
| 396671.7 | 1576621 | 1646984 | 1380808 | 1006043 | 11865775 | 9474397 | 11021958 | 1666585 |
| 92950.09 | 176544.6 | 187492.4 | 172092.1 | 110997.4 | 3057913 | 3268153 | 2982867 | 154180.1 |
| 326295.6 | 13674132 | 13845990 | 9853259 | 366351.5 | 16508011 | 13533847 | 16316165 | 14074089 |
| 113253.1 | 13086129 | 12524437 | 12438218 | 154336.4 | 12323006 | 11683240 | 12215059 | 3165771 |
| 49357.83 | 8264752 | 7698870 | 7828235 | 57664.15 | 8840035 | 7444911 | 9620055 | 2417562 |
| 239069.2 | 14734452 | 13532637 | 12798118 | 241771.8 | 13910600 | 11414019 | 10882858 | 1445324 |
| 429679.3 | 496861.4 | 346690.4 | 378882 | 744569.6 | 2814367 | 3141378 | 2965829 | 623684.4 |
| 83870.41 | 27447085 | 16773484 | 15784015 | 82198.5 | 15597451 | 14505157 | 13429013 | 10129510 |
| 291542.4 | 23950215 | 26368856 | 23157485 | 110561.9 | 12860547 | 12557598 | 13178147 | 6818714 |

| | | | | | | | | |
|---|---|---|---|---|---|---|---|---|
| 98854.42 | 36408457 | 25652988 | 26681205 | 96148.16 | 26992417 | 23077034 | 17305233 | 7650433 |
| 235651.7 | 5667181 | 2875853 | 3200731 | 230162.4 | 20752697 | 19180420 | 20248376 | 1294158 |
| 590310.2 | 5416011 | 5707835 | 5853294 | 192038.3 | 5979838 | 4868200 | 5317156 | 4849568 |
| 2021194 | 20099028 | 27363203 | 27128666 | 2422796 | 24188629 | 18561855 | 20754710 | 4643937 |
| 87472.49 | 40127139 | 37819570 | 39226939 | 126615.1 | 37363294 | 33970839 | 35535215 | 15115379 |
| 5217623 | 20710147 | 20876258 | 15467516 | 16761219 | 1.03E+08 | 94059665 | 95843615 | 2.39E+08 |
| 124711.7 | 78173081 | 65662853 | 71111417 | 135668.5 | 65047429 | 52535988 | 57939899 | 22338849 |
| 112082.8 | 240574.8 | 244851.8 | 222129.6 | 125377.8 | 2967650 | 3015902 | 2088062 | 206425.8 |
| 135356.5 | 6465794 | 13923836 | 11922655 | 141821.1 | 10229905 | 8080623 | 8306691 | 6888864 |
| 395398.8 | 1066656 | 2041639 | 2069651 | 422172.7 | 20524222 | 16598184 | 17296072 | 42735286 |
| 198700.2 | 1838158 | 3658861 | 880822.5 | 195740.6 | 16110638 | 12692335 | 13572038 | 29995476 |
| 58365.49 | 8561156 | 8463195 | 9352458 | 137288.8 | 6964539 | 5732965 | 5806289 | 311733.5 |
| 71682.53 | 119699.7 | 122323.2 | 125306.9 | 83723.32 | 1965486 | 1682971 | 1586603 | 107047 |
| 336590 | 5363668 | 5272746 | 5884824 | 338620.1 | 7342265 | 5576973 | 5583585 | 2848921 |
| 206773.6 | 8203103 | 8435579 | 8775335 | 742715.3 | 9299712 | 9198250 | 9018023 | 2987970 |
| 174929 | 26514799 | 24989418 | 24336896 | 186042.5 | 26430309 | 22314601 | 20729280 | 8957572 |
| 163926.2 | 26219046 | 25151915 | 23930474 | 172143.3 | 24246570 | 21994875 | 20709191 | 8957572 |
| 1937041 | 33098617 | 23890973 | 37244132 | 1072989 | 21360222 | 20496126 | 20556896 | 17890873 |
| 108811.6 | 4066158 | 4086328 | 3379474 | 152054.9 | 9090742 | 9181524 | 9464498 | 17997155 |
| 158413.2 | 115261.1 | 107484.2 | 295323.4 | 49089.6 | 2671782 | 1488567 | 1966008 | 4905280 |
| 80453.5 | 3798480 | 3579418 | 3313543 | 89891.19 | 7681517 | 6502014 | 6871501 | 9226129 |
| 45315.91 | 987659.6 | 945058.5 | 1025725 | 66070.02 | 5034058 | 4409605 | 4552415 | 10638677 |
| 1349268 | 1.85E+08 | 1.73E+08 | 1.7E+08 | 1458542 | 1.87E+08 | 1.72E+08 | 1.83E+08 | 1.28E+08 |
| 70653.81 | 11694977 | 8522009 | 9130869 | 73172.59 | 9719458 | 9138072 | 8806032 | 6034128 |
| 86147.41 | 5249446 | 4511236 | 6073581 | 116984.3 | 5197526 | 4296224 | 4956084 | 3633931 |
| 86230.94 | 18572280 | 15747165 | 13536670 | 95202.75 | 35607490 | 31991210 | 26492843 | 50159222 |
| 67284.18 | 7934700 | 6838409 | 6855777 | 467911.5 | 23341746 | 19181060 | 19728656 | 908868.2 |
| 105132.3 | 11931743 | 5566310 | 7989210 | 117535.9 | 11938790 | 10794576 | 7686168 | 7796783 |
| 149844.7 | 2690943 | 2657948 | 2875177 | 269965.9 | 5391515 | 4650088 | 5081483 | 113428.6 |
| 112568.9 | 13496020 | 12886721 | 13364427 | 149253.5 | 11688238 | 10629771 | 10757637 | 6486562 |
| 78210.08 | 133999 | 127187.5 | 125696.8 | 86715.85 | 2278728 | 2273597 | 2026646 | 6645830 |

| Area_S2 | Area_S3 | Area_S_PB | Group_Area_Avedøre | Group_Area_Hvidovre | Group_Area_Skævinge | Group_CV_pct_Avedøre | Group_CV_pct_Hvidovre | Group_CV_pct_Skævinge |
|---|---|---|---|---|---|---|---|---|
| 6.62E+08 | 6.63E+08 | 18883258 | 5.7E+08 | 1.08E+09 | 6.62E+08 | 2.006289 | 2.412113 | 6.10E+00 |
| 75355604 | 80355104 | 2544986 | 1.73E+08 | 92303555 | 75355604 | 29.70995 | 4.614152 | 14.21556 |
| 40342102 | 44085603 | 3147335 | 92635956 | 48996169 | 40342102 | 0.603875 | 4.829347 | 15.42917 |
| 6195283 | 5602647 | 25396394 | 68141862 | 1.2E+09 | 6195283 | 2.632537 | 3.182996 | 17.91889 |
| 153437 | 160714.1 | 101836.4 | 1.7E+08 | 595191.6 | 153437 | 12.21282 | 46.80422 | 8.16E+00 |
| 164609.4 | 167001.8 | 108017.4 | 61956476 | 175370.1 | 164609.4 | 7.498444 | 4.392828 | 7.127163 |
| 2.6E+08 | 2.84E+08 | 8705711 | 2.66E+08 | 4.36E+08 | 2.6E+08 | 4.232572 | 10.39209 | 13.28715 |
| 2234112 | 2560015 | 223118.2 | 4764976 | 2525209 | 2234112 | 20.64568 | 3.60E+00 | 2.04E+01 |
| 555625.7 | 488089.6 | 121609.2 | 2992915 | 2444737 | 495476.2 | 18.44778 | 7.037894 | 7.220199 |
| 2795621 | 2793002 | 210766.7 | 3421106 | 7847205 | 2793002 | 25.44976 | 3.15E+00 | 1.03E+00 |
| 17167897 | 22325794 | 165556 | 8361661 | 10608299 | 19789311 | 64.35471 | 1.11E+01 | 1.31E+01 |
| 5053042 | 711505.3 | 101097.8 | 12864039 | 861354.8 | 1980935 | 11.29091 | 45.05268 | 86.46093 |
| 86208.86 | 92615.5 | 54055.38 | 886734.7 | 9988993 | 87058.39 | 13.25087 | 1.385406 | 3.93E+00 |
| 18812667 | 18522849 | 107209.4 | 27613302 | 41037651 | 18522849 | 4.950001 | 2.841519 | 5.734159 |
| 13233623 | 13583081 | 132587.7 | 17017573 | 29071978 | 13233623 | 5.592374 | 4.721744 | 11.55464 |
| 7694329 | 7314035 | 90601.58 | 7415919 | 16295079 | 7314035 | 6.389416 | 8.347554 | 28.59826 |
| 1.05E+08 | 1.09E+08 | 20341271 | 1.99E+08 | 3.61E+08 | 1.09E+08 | 9.220559 | 13.69654 | 3.507148 |
| 6077468 | 6601869 | 206855 | 80568177 | 33009224 | 6077468 | 2.028754 | 2.97409 | 8.158677 |
| 47405880 | 52610675 | 1009715 | 30072649 | 34568813 | 47405880 | 69.24387 | 58.51066 | 6.33E+00 |
| 92406.76 | 91517.06 | 91586.18 | 6419704 | 99977.86 | 92406.76 | 20.36983 | 3.122043 | 1.76E+00 |
| 2091784 | 1737932 | 91553.79 | 7490353 | 8674777 | 1737932 | 57.65156 | 7.23E+00 | 2.16E+01 |
| 3681483 | 1946750 | 137755.5 | 3.16E+08 | 3510488 | 1946750 | 2.404375 | 29.15407 | 4.03E+01 |
| 249640.8 | 366936.7 | 364107.9 | 5.16E+08 | 1973523 | 261577.3 | 1.240141 | 34.15879 | 2.21E+01 |
| 3040152 | 560040.3 | 110483.9 | 30356364 | 3149793 | 3040152 | 7.344679 | 40.65439 | 68.64398 |
| 2235510 | 2401450 | 176411.6 | 10297042 | 4443761 | 2235510 | 6.069002 | 36.78872 | 31.37284 |
| 1509177 | 3090154 | 83865.17 | 43753647 | 4753878 | 2282494 | 3.248482 | 48.07079 | 34.46255 |
| 15691617 | 16031456 | 171996.5 | 15942157 | 25539985 | 15691617 | 15.96328 | 3.480287 | 1.16E+01 |
| 8186261 | 9624664 | 289597.7 | 13535133 | 7092812 | 8186261 | 7.54785 | 85.07013 | 25.81034 |
| 267946.1 | 168927.7 | 106327.9 | 7378231 | 423441.6 | 187428.1 | 6.031273 | 29.60605 | 25.29868 |
| 1473767 | 1485425 | 71366.44 | 34640245 | 71608.14 | 1485425 | 4.465574 | 1.68E+01 | 2.17E+00 |
| 749215.7 | 717481.8 | 82937.1 | 8070427 | 7133646 | 749215.7 | 7.953987 | 15.27842 | 7.686615 |
| 38166249 | 39198401 | 975819 | 28044795 | 27655888 | 38166249 | 3.135392 | 8.187059 | 7.035319 |
| 6169125 | 5592399 | 225230.7 | 18693831 | 11979784 | 6169125 | 28.60963 | 3.35E+01 | 3.33E+01 |
| 6247478 | 9742880 | 251367.7 | 13583076 | 12045005 | 9742880 | 22.45967 | 84.35446 | 54.45191 |
| 5685491 | 5900969 | 140032.3 | 22988905 | 7000008 | 5685491 | 8.430287 | 74.55343 | 1.21E+01 |
| 16686795 | 14260020 | 341105.1 | 21424272 | 19825351 | 14260020 | 4.323622 | 26.9823 | 19.68469 |
| 5104791 | 5888162 | 788022 | 8101226 | 4415098 | 5104791 | 99.57737 | 4.39E+00 | 1.29E+01 |
| 10109444 | 11965421 | 345200.6 | 12479957 | 8149424 | 10198087 | 28.44114 | 4.192046 | 9.731663 |
| 6367182 | 5336194 | 200937.7 | 13884071 | 24386378 | 5336194 | 18.63627 | 35.68955 | 24.28803 |
| 6248145 | 4175597 | 210153.6 | 8742484 | 7020209 | 5262944 | 13.61335 | 9.104606 | 19.82624 |

| | | | | | | | | |
|---|---|---|---|---|---|---|---|---|
| 2006135 | 6177736 | 168773.4 | 13912871 | 9384919 | 4296495 | 10.02193 | 40.48714 | 50.21828 |
| 482984.4 | 4296602 | 66221.99 | 23766178 | 6207952 | 4296602 | 1.497377 | 3.55E+01 | 7.54E+01 |
| 12770030 | 10249998 | 507231.4 | 33518916 | 22856989 | 10249998 | 4.527141 | 13.85122 | 21.84264 |
| 22844929 | 22193484 | 538261.8 | 27496935 | 26795746 | 22193484 | 34.78947 | 1.313492 | 5.75E+01 |
| 26229530 | 25060597 | 983277.4 | 5820364 | 20929372 | 25060597 | 12.72522 | 1.33E+01 | 3.161525 |
| 2983328 | 1054185 | 2396868 | 1.21E+09 | 2805279 | 2529943 | 3.218492 | 6.27E+01 | 46.07779 |
| 158963.2 | 160738.2 | 118474.7 | 1.09E+08 | 238496.3 | 158963.2 | 2.826687 | 2.34E+01 | 7.02E+00 |
| 638330 | 417234.9 | 78066.56 | 12449586 | 1621649 | 447300.7 | 8.513395 | 92.09683 | 23.93749 |
| 508819 | 4067252 | 104719.2 | 26805628 | 1317686 | 516828.3 | 3.478089 | 39.43828 | 1.21E+02 |
| 1179540 | 3736877 | 113917.9 | 2400184 | 11090830 | 1760798 | 53.74202 | 23.5078 | 60.23022 |
| 6121896 | 6647900 | 368334.5 | 12173331 | 11025079 | 6121896 | 9.353739 | 7.849428 | 9.628088 |
| 142911.7 | 251821.9 | 90735.75 | 17832244 | 163722.8 | 251821.9 | 2.531702 | 2.11E+00 | 47.16406 |
| 176875.7 | 185301.5 | 104285.3 | 57936190 | 210313.5 | 176875.7 | 1.203427 | 6.144156 | 7.82E+00 |
| 117044.2 | 258647.2 | 86242.13 | 17558458 | 244883.4 | 117044.2 | 4.665392 | 36.73165 | 52.49568 |
| 2887286 | 2849857 | 108887.9 | 1811492 | 8265468 | 2887286 | 10.11869 | 19.34056 | 1.39744 |
| 4208996 | 11128772 | 896326.5 | 32001812 | 24664281 | 11128772 | 22.19586 | 33.13747 | 58.83825 |
| 7932297 | 16689441 | 202180.6 | 9987926 | 14221282 | 14893134 | 45.593 | 1.26E+01 | 3.51E+01 |
| 32412111 | 35762730 | 202049.8 | 85530004 | 33377220 | 32412111 | 2.14078 | 1.21E+01 | 1.23E+01 |
| 7834730 | 18444808 | 173325.5 | 95565504 | 6804758 | 7834730 | 4.35018 | 93.00945 | 64.57361 |
| 45607268 | 39433536 | 180973.9 | 25814013 | 42845598 | 39433536 | 22.44538 | 1.809427 | 9.009569 |
| 11763094 | 13875568 | 526838.6 | 29699764 | 18315096 | 11763094 | 8.863361 | 24.93433 | 26.34654 |
| 9022210 | 7591022 | 3840441 | 38624760 | 14491029 | 8216625 | 0.144233 | 7.891044 | 8.67E+00 |
| 47500802 | 42301363 | 218822.6 | 2490951 | 6943900 | 45494012 | 53.92304 | 47.96812 | 5.81E+00 |
| 11964179 | 12068201 | 1685350 | 11376433 | 23251351 | 11964179 | 16.23691 | 4.155001 | 6.184374 |
| 18262552 | 18533679 | 1457277 | 18693961 | 10224336 | 18262552 | 7.0117 | 8.032245 | 7.686248 |
| 7590690 | 5637031 | 139838.6 | 1616221 | 389975.3 | 5637031 | 60.66002 | 97.80024 | 20.08034 |
| 439909.9 | 297496.7 | 153525.5 | 10671315 | 1092697 | 439909.9 | 15.63046 | 9.510497 | 24.05644 |
| 3605381 | 3779931 | 73873.3 | 6512832 | 6784556 | 3605381 | 1.116234 | 2.17E+00 | 8.35E+00 |
| 7873839 | 8171082 | 339876.7 | 13662907 | 3899991 | 7873839 | 3.555593 | 7.173928 | 7.219636 |
| 12164166 | 14337256 | 54448.92 | 3646870 | 2740066 | 12164166 | 78.74852 | 72.34138 | 1.52E+01 |
| 10205824 | 10610514 | 9772088 | 2.08E+08 | 14247170 | 10610514 | 27.40426 | 16.72129 | 4.59132 |
| 18215642 | 19267863 | 3582850 | 18594945 | 13594995 | 18215642 | 22.18348 | 9.82E+01 | 1.03E+01 |
| 5594042 | 5433460 | 72104.44 | 7952066 | 6695465 | 5433460 | 3.900183 | 8.957344 | 6.139063 |
| 10110125 | 6977298 | 830195.6 | 9322733 | 7033753 | 7249217 | 6.819392 | 3.90E+00 | 2.14E+01 |
| 23273818 | 18315925 | 271536.2 | 21494685 | 19107040 | 18315925 | 45.98546 | 1.99E+01 | 2.43E+01 |
| 23566658 | 14959897 | 292217.6 | 11731210 | 28231599 | 14959897 | 64.29304 | 5.20E+01 | 3.79E+01 |
| 8093709 | 7459502 | 362735.4 | 12967327 | 11787221 | 7459502 | 1.476495 | 1.36E+01 | 1.09E+01 |
| 3425751 | 3987217 | 168693.6 | 20350434 | 11317456 | 3987217 | 6.213937 | 12.30415 | 59.01543 |
| 16138640 | 16320177 | 5006525 | 28955649 | 28666283 | 16138640 | 75.63361 | 55.60927 | 53.84958 |
| 4567899 | 6891321 | 494072.5 | 16319657 | 9283904 | 4567899 | 34.93993 | 44.11624 | 3.61E+01 |
| 3114002 | 3601009 | 79784.48 | 8439776 | 7005416 | 3601009 | 6.767442 | 18.9134 | 8.529514 |
| 3834790 | 3418148 | 131499.3 | 11360060 | 8229703 | 3834790 | 3.439141 | 10.80677 | 15.07655 |
| 15527066 | 15353408 | 5388767 | 65000436 | 41360444 | 15527066 | 16.06462 | 7.02E+01 | 2.70E+01 |
| 13727928 | 12152229 | 2842333 | 76669353 | 28466769 | 12152229 | 10.81308 | 6.72E+01 | 8.33E+00 |
| 4177137 | 1506510 | 130043.3 | 8143255 | 6629811 | 3443907 | 47.08266 | 37.18 | 45.35118 |
| 5628301 | 8447406 | 581223.3 | 7513670 | 6624748 | 5628301 | 31.77685 | 1.01E+01 | 3.40E+01 |
| 7670397 | 8104578 | 349405.2 | 30523327 | 15561093 | 7670397 | 4.245575 | 3.04E+00 | 4.33E+00 |
| 36612144 | 37441896 | 1761774 | 15847114 | 26271897 | 36612144 | 4.866409 | 4.82E+01 | 7.83E+00 |
| 6157447 | 6654579 | 1154672 | 15068964 | 13700543 | 6157447 | 29.51558 | 54.71795 | 1.28E+01 |
| 61646986 | 60882133 | 14993171 | 8.91E+08 | 1.47E+08 | 61646986 | 4.365694 | 12.79722 | 1.89E+01 |
| 46839421 | 46822519 | 1814891 | 48189608 | 1.21E+08 | 46822519 | 2.888057 | 4.24188 | 5.987525 |

| | | | | | | | | |
|---|---|---|---|---|---|---|---|---|
| 4631087 | 4574354 | 591646.9 | 8122252 | 8343929 | 4631087 | 3.048464 | 11.23129 | 9.56E-01 |
| 2308436 | 8786020 | 135852.6 | 16119119 | 1626930 | 2308436 | 6.354781 | 53.55998 | 91.68499 |
| 5601082 | 6129392 | 93376.47 | 5385169 | 7302456 | 5601082 | 15.43011 | 33.45133 | 18.86441 |
| 4597820 | 4651460 | 172317.1 | 21032726 | 9482363 | 4597820 | 2.470243 | 2.943616 | 6.430722 |
| 7670397 | 8104578 | 349405.2 | 30523327 | 15561093 | 7670397 | 4.245575 | 3.03593 | 4.325932 |
| 153453.9 | 263946.5 | 65461.39 | 11977653 | 271580.6 | 216678.2 | 16.4191 | 44.60229 | 26.22922 |
| 5233340 | 430937.8 | 87199.3 | 5268264 | 4255475 | 1822697 | 23.03702 | 13.80477 | 99.00858 |
| 17713775 | 16668126 | 114036.4 | 20487375 | 12681892 | 16668126 | 17.7828 | 2.489382 | 11.14431 |
| 260360.7 | 265568.7 | 107722 | 7109623 | 643980.1 | 265568.7 | 1.12026 | 4.36E+01 | 3.85E+01 |
| 13270003 | 5539551 | 657117.8 | 13117914 | 15209472 | 7324418 | 20.04443 | 8.361167 | 46.463 |
| 5560099 | 9743252 | 289701.7 | 39026301 | 1566044 | 5560099 | 3.373146 | 118.7317 | 86.88431 |
| 56378395 | 53963720 | 4797466 | 1.22E+08 | 53556287 | 53963720 | 4.502916 | 1.09E+00 | 6.80E+00 |
| 19969546 | 12325434 | 187761.9 | 13254554 | 13901561 | 12325434 | 18.98865 | 1.13E+01 | 4.07E+01 |
| 6296579 | 4803912 | 290103.9 | 7750226 | 20392392 | 4803912 | 7.640239 | 17.82754 | 20.89399 |
| 7521054 | 7727314 | 508068.8 | 12196279 | 10086900 | 7521054 | 7.778085 | 1.18E+00 | 5.62E+00 |
| 1.38E+08 | 1.3E+08 | 8478615 | 1.15E+08 | 1.39E+08 | 1.3E+08 | 13.73042 | 1.76E+00 | 1.30E+01 |
| 56600296 | 59931174 | 5142241 | 53059311 | 68271601 | 56600296 | 5.356889 | 6.151297 | 1.80E+01 |
| 30910682 | 26453062 | 1742267 | 29082797 | 35949177 | 26453062 | 4.531661 | 7.105877 | 1.32E+01 |
| 489406.4 | 353296.9 | 122843.3 | 898484.8 | 6749999 | 353296.9 | 11.19787 | 1.78E+01 | 3.39E+01 |
| 9135744 | 8385528 | 7953737 | 9514309 | 3.56E+08 | 8385528 | 6.047151 | 3.763968 | 5.98267 |
| 4734372 | 4542320 | 198553.9 | 4976437 | 3964723 | 4542320 | 9.152069 | 7.021451 | 16.09197 |
| 44499969 | 45694045 | 2405015 | 1.01E+08 | 69580884 | 44499969 | 4.315784 | 7.971722 | 13.03392 |
| 7288931 | 5387258 | 2323493 | 27061135 | 14686419 | 5387258 | 7.635701 | 8.486885 | 20.09373 |
| 1902261 | 2078093 | 415831.9 | 16588567 | 1099157 | 1902261 | 37.45043 | 1.80E+01 | 9.13E+00 |
| 26336614 | 28005380 | 828641.8 | 11959466 | 3.22E+08 | 26336614 | 0.580518 | 1.49E+00 | 2.20E+01 |
| 10032194 | 34050397 | 118456.7 | ######## | ######## | ######## | 40.46679 | 10.49253 | 53.0583 |
| 10240434 | 10716909 | 88535.28 | ######## | ######## | ######## | 1.890307 | 8.004149 | 9.61E+00 |
| 433358.6 | 547550.8 | 389321.8 | ######## | ######## | ######## | 2.798976 | 88.49913 | 16.81287 |
| 219443.1 | 1152252 | 151702.9 | ######## | ######## | ######## | 28.67493 | 51.36214 | 1.03E+02 |
| 8695682 | 9809878 | 1867425 | ######## | ######## | ######## | 1.439248 | 7.80357 | 10.25318 |
| 6.31E+08 | 6.01E+08 | 26630206 | ######## | ######## | ######## | 1.734939 | 2.14E+00 | 1.04E+01 |
| 1656816 | 708128.5 | 533790.9 | ######## | ######## | ######## | 4.60611 | 1.94E+01 | 39.08175 |
| 1.01E+08 | 1.06E+08 | 2787453 | ######## | ######## | ######## | 1.307598 | 3.96644 | 9.628373 |
| 3536542 | 1787337 | 2707995 | ######## | ######## | ######## | 3.64907 | 1.39E+01 | 3.99E+01 |
| 15513052 | 16570660 | 962772.6 | ######## | ######## | ######## | 2.791559 | 2.78E+00 | 1.04E+01 |
| 44104963 | 46697618 | 149035.7 | ######## | ######## | ######## | 10.04235 | 4.261843 | 8.904069 |
| 4910616 | 5052567 | 886815.4 | ######## | ######## | ######## | 9.610717 | 13.92832 | 3.106908 |
| 1448929 | 1476489 | 1178371 | ######## | ######## | ######## | 1.811043 | 11.02436 | 0.942892 |
| 785474.5 | 1010746 | 92614.78 | ######## | ######## | ######## | 7.045378 | 7.665598 | 1.33E+01 |
| 9256680 | 10374415 | 99772.49 | 5279024 | 12374859 | 9256680 | 12.06925 | 5.921153 | 9.052669 |
| 24725951 | 24760673 | 189358.5 | 33400064 | 37614947 | 24725951 | 3.971399 | 1.76E+00 | 1.03E+01 |
| 19077524 | 17943515 | 2398271 | 17666524 | 20800366 | 17943515 | 39.52101 | 18.10343 | 34.26089 |
| 3425299 | 3720393 | 1251383 | 4912594 | 10920990 | 3720393 | 2.279924 | 3.00E+00 | 1.54E+01 |
| 1.07E+08 | 1.24E+08 | 95229.46 | 29576720 | 18039031 | 1.07E+08 | 2.715788 | 7.281752 | 10.81816 |
| 80513.57 | 84424.62 | 45010.18 | 5041131 | 90602.85 | 80513.57 | 6.117275 | 1.05E+01 | 8.33E+00 |
| 12726904 | 16465656 | 177774.4 | 26759458 | 22320925 | 12726904 | 5.710705 | 11.90216 | 2.42E+01 |
| 1904650 | 1628960 | 81461.79 | 8218292 | 8329179 | 1628960 | 5.276914 | 5.82E+00 | 1.02E+01 |
| 66710097 | 67345591 | 3528805 | 1.03E+08 | 87982228 | 66710097 | 4.447327 | 0.411104 | 7.204166 |
| 9246749 | 10325263 | 71500.75 | 19997550 | 13061630 | 9246749 | 0.601676 | 2.93E+00 | 7.91E+00 |
| 100067.6 | 152372.8 | 52380.58 | 12653531 | 567588.3 | 152372.8 | 3.447664 | 1.06E+01 | 2.53E+01 |
| 7107240 | 8810472 | 442739 | 18585370 | 21323797 | 7107240 | 9.378383 | 5.904978 | 26.39858 |

| | | | | | | | | |
|---|---|---|---|---|---|---|---|---|
| 3661568 | 4011114 | 230114.1 | 9641663 | 6037509 | 3661568 | 5.811775 | 31.31763 | 6.74E+00 |
| 18755533 | 18981419 | 711602.3 | 33503114 | 33669191 | 18755533 | 8.764288 | 2.18E+00 | 1.20E+01 |
| 6147777 | 5500157 | 95248.82 | 14313648 | 5342069 | 5500157 | 8.266561 | 11.94992 | 15.14748 |
| 53199627 | 41652100 | 1805220 | 42565831 | 58256767 | 41652100 | 17.14772 | 16.93912 | 1.53E+01 |
| 9477306 | 5285608 | 91343.14 | 9816921 | 19279186 | 7511162 | 26.89976 | 1.90E+01 | 2.82E+01 |
| 6966128 | 13582264 | 421085.8 | 57953321 | 1.31E+08 | 11294748 | 1.412282 | 1.170354 | 3.17E+01 |
| 115414.2 | 114688.7 | 87347.26 | 15570482 | 19027516 | 115414.2 | 4.515921 | 1.31E+01 | 4.36E+01 |
| 2.09E+08 | 2.15E+08 | 18935603 | 2.1E+09 | 1.28E+09 | 2.09E+08 | 3.060208 | 1.440074 | 8.195736 |
| 74335698 | 68023400 | 116702.3 | 1.3E+08 | 1.08E+08 | 68023400 | 1.047179 | 12.92239 | 7.824684 |
| 6222882 | 6064191 | 110466.7 | 11220134 | 6316887 | 6064191 | 32.3032 | 38.05226 | 4.77E+01 |
| 2009368 | 2258691 | 1643988 | 23136980 | 15057500 | 2009368 | 0.260779 | 2.770521 | 10.23462 |
| 19434391 | 17442634 | 151133.9 | 27051854 | 20751556 | 17442634 | 2.931288 | 9.379391 | 1.03E+01 |
| 38924907 | 39858508 | 144523.9 | 13391684 | 37037138 | 38924907 | 5.854865 | 3.08E+00 | 1.03E+01 |
| 58219767 | 8608103 | 675941.8 | 1.41E+08 | 69197327 | 34297724 | 21.39729 | 71.2164 | 7.36E+01 |
| 2296894 | 2657296 | 105500 | 19815995 | 18210286 | 2296894 | 2.389289 | 10.43606 | 19.48087 |
| 24813285 | 25490328 | 99426.86 | 29065043 | 20414546 | 24813285 | 1.857545 | 4.652229 | 1.02E+01 |
| 4536325 | 3793501 | 92113.81 | 11172847 | 9260889 | 3793501 | 9.574448 | 15.4541 | 18.81917 |
| 4856778 | 5278986 | 60692.61 | 5987592 | 5709667 | 4856778 | 0.622785 | 24.00258 | 11.08278 |
| 254454.5 | 252893.5 | 1244833 | 978875.9 | 1.39E+08 | 252893.5 | 6.79102 | 8.59E+00 | 6.59E+00 |
| 8108196 | 8273502 | 95151.55 | 6852462 | 7575122 | 8108196 | 1.678255 | 6.07E+00 | 10.79066 |
| 38907878 | 38910544 | 125585.7 | 35814496 | 34487132 | 38907878 | 0.718261 | 11.29791 | 2.94E+00 |
| 43335222 | 43842407 | 104233.6 | 88073595 | 75383455 | 43335222 | 2.750046 | 9.578571 | 7.721697 |
| 31725486 | 20332748 | 166092.7 | 23336887 | 19280158 | 26830233 | 4.713795 | 2.77E+00 | 2.17E+01 |
| 10083647 | 7019890 | 80186.76 | 6597240 | 5600854 | 8432773 | 5.232903 | 11.57256 | 1.80E+01 |
| 10499134 | 7722620 | 163638.1 | 6462644 | 5566180 | 9061004 | 3.479902 | 1.95E+01 | 1.53E+01 |
| 1.1E+08 | 1.15E+08 | 193059 | 1.09E+08 | 1.04E+08 | 1.1E+08 | 3.479566 | 4.051859 | 1.00E+01 |
| 73361892 | 80595029 | 288681.4 | 86795789 | 1.04E+08 | 73361892 | 2.968935 | 8.857035 | 14.8514 |
| 15329629 | 17145926 | 93129.86 | 48334613 | 37499501 | 17145926 | 6.380009 | 5.011508 | 13.1036 |
| 2438063 | 2080171 | 198769.4 | 7449607 | 5785044 | 2080171 | 11.93275 | 7.665995 | 12.58155 |
| 51395961 | 46127106 | 601111.9 | 1.32E+08 | 1.49E+08 | 46127106 | 0.641266 | 3.011975 | 9.514464 |
| 279857.1 | 1446717 | 8036698 | 381233.3 | 1.11E+09 | 1446717 | 90.7168 | 3.008305 | 7.17E+01 |
| 1.19E+08 | 1.25E+08 | 1142594 | 1.57E+08 | 1.67E+08 | 1.19E+08 | 1.255882 | 1.30E+00 | 1.30E+01 |
| 2.4E+09 | 2.43E+09 | 30573952 | 1.73E+09 | 1.72E+09 | 2.4E+09 | 1.713415 | 1.913826 | 10.55481 |
| 4614303 | 6325113 | 58465.32 | 319362.2 | 1383506 | 4620803 | 64.53447 | 20.80933 | 19.00742 |
| 358493.7 | 574765.1 | 113710.8 | 7732263 | 2772187 | 574765.1 | 6.141284 | 41.22807 | 105.7352 |
| 788990 | 529471.5 | 144401.8 | 7884504 | 6766821 | 529471.5 | 5.493747 | 27.57213 | 51.78309 |
| 20352785 | 24367042 | 128057.4 | 3468063 | 1875220 | 20459545 | 28.79675 | 5.510843 | 10.52834 |
| 97843076 | 99612586 | 157073.7 | 16633645 | 14648016 | 97843076 | 67.67268 | 62.75329 | 5.399386 |
| 33889530 | 28992926 | 159181.8 | 1424625 | 338252.5 | 29864052 | 87.33787 | 109.3047 | 8.45E+00 |
| 10325908 | 11082509 | 82807.93 | 9412502 | 10780952 | 10325908 | 3.160942 | 3.56E+00 | 9.43E+00 |
| 1158069 | 1119457 | 96234.2 | 6711779 | 2617283 | 1158069 | 10.95657 | 42.22656 | 6.78E+01 |
| 6398632 | 4139310 | 95569.74 | 12900933 | 8653343 | 4541599 | 5.32861 | 1.36E+01 | 2.40E+01 |
| 175658.2 | 182874.5 | 187040.5 | 12215212 | 8120144 | 175658.2 | 2.059338 | 4.08E+00 | 3.10E+00 |
| 1.72E+08 | 1.65E+08 | 976741.4 | 2.58E+08 | 2.5E+08 | 1.65E+08 | 1.746497 | 11.86746 | 8.335413 |
| 8407185 | 8546491 | 444767.8 | 35569165 | 6870052 | 8407185 | 67.19101 | 24.17385 | 8.464969 |
| 10263303 | 10814929 | 154747.7 | 59977861 | 41207920 | 10263303 | 2.082485 | 4.411691 | 21.07566 |
| 54325990 | 59908253 | 147916.3 | 85480151 | 90948257 | 54325990 | 1.637861 | 5.926932 | 14.74419 |
| 15272189 | 15671337 | 87473.15 | 9199372 | 8239198 | 15272189 | 3.389786 | 8.02E+00 | 1.19E+01 |
| 19094133 | 16541315 | 1317423 | 79780756 | 91553933 | 16541315 | 0.612114 | 6.32E+00 | 9.93E+00 |
| 4031802 | 5072387 | 175417.6 | 19199912 | 11673412 | 4031802 | 0.954633 | 2.93E+00 | 1.57E+01 |
| 20455683 | 27560763 | 126312.2 | 84760274 | 55071928 | 22271856 | 1.702222 | 3.079912 | 15.75487 |

| | | | | | | | | |
|---|---|---|---|---|---|---|---|---|
| 613418.4 | 288893.9 | 151498.9 | 12630298 | 11853595 | 451829.1 | 7.378679 | 3.37E+00 | 3.59E+01 |
| 5.8E+08 | 5.88E+08 | 5538447 | 8.57E+08 | 6.4E+08 | 5.8E+08 | 0.98897 | 12.27802 | 9.40907 |
| 21601364 | 23367353 | 1374610 | 1.74E+08 | 41136083 | 21601364 | 5.714681 | 3.879193 | 11.48853 |
| 35790372 | 33060018 | 1436656 | 76794600 | 1.02E+08 | 33060018 | 1.110563 | 3.08E+00 | 7.09E+00 |
| 571043.5 | 4343332 | 102691.8 | 8593023 | 1398183 | 3897713 | 3.754064 | 82.59838 | 70.17756 |
| 651025.8 | 599321.4 | 165361.5 | 3396064 | 3628909 | 599321.4 | 5.649718 | 3.103097 | 12.87909 |
| 83888231 | 79084838 | 5567895 | 1.38E+08 | 90677822 | 79084838 | 6.905231 | 3.41E+00 | 7.17E+00 |
| 217212 | 225341 | 213579.9 | 86720465 | 225282.7 | 217212 | 0.457105 | 4.877644 | 2.349517 |
| 10017124 | 10722694 | 192231.2 | 14003471 | 14126573 | 10017124 | 1.538954 | 3.65E+00 | 1.41E+01 |
| 9671918 | 10014586 | 103269.8 | 30750275 | 30923049 | 9671918 | 3.10198 | 1.06E+00 | 1.44E+01 |
| 375463.1 | 497854.8 | 333395 | 3.53E+08 | 1496486 | 497854.8 | 4.492538 | 6.12E+01 | 51.71603 |
| 8526255 | 13685435 | 200480.2 | 71034231 | 1.26E+08 | 8526255 | 1.25603 | 6.435548 | 30.9751 |
| 8258691 | 5433841 | 193342.1 | 2129438 | 6855309 | 5433841 | 43.58387 | 2.62E+01 | 2.60E+01 |
| 13226442 | 10291028 | 124892.6 | 5296294 | 6059640 | 11751727 | 60.85666 | 6.65E+00 | 1.25E+01 |
| 9591671 | 9698970 | 336322.2 | 15215449 | 15331527 | 9591671 | 10.47538 | 15.52094 | 8.55E+00 |
| 34944151 | 40827717 | 182956.5 | 68951041 | 52303623 | 34944151 | 1.571705 | 1.55144 | 17.26171 |
| 3143545 | 3078286 | 117742.1 | 7363558 | 4123754 | 3114321 | 17.3917 | 10.54226 | 1.050385 |
| 28062604 | 25398887 | 1288134 | 34397684 | 24231786 | 25398887 | 5.627537 | 10.29451 | 8.913575 |
| 15316885 | 24519885 | 215938.9 | 32253886 | 31065533 | 19079378 | 6.26145 | 7.476151 | 23.56023 |
| 1114417 | 9439097 | 616058.6 | 12879871 | 12426575 | 8239416 | 13.20269 | 2.639373 | 71.83705 |
| 1001644 | 1336731 | 134519.8 | 7223913 | 3233294 | 1336731 | 12.3672 | 28.84742 | 29.19462 |
| 4412478 | 5170770 | 117959.2 | 7829472 | 11632006 | 4628698 | 4.975387 | 5.333274 | 8.246039 |
| 32672452 | 33980757 | 171530.8 | 30741563 | 26683773 | 33426031 | 5.125962 | 5.53E+00 | 1.97E+00 |
| 1461528 | 773029.3 | 121975.9 | 47808798 | 185102 | 773029.3 | 2.082986 | 2.59E+01 | 8.25E+01 |
| 261694.6 | 368928.8 | 132487.9 | 26152071 | 318088.5 | 301222.5 | 11.31875 | 11.18341 | 1.75E+01 |
| 7037610 | 2138943 | 125432 | 10989821 | 7072656 | 2706668 | 3.210039 | 71.29004 | 67.64422 |
| 191229.7 | 197221.1 | 135709.3 | 36619920 | 200678.9 | 191229.7 | 31.31695 | 2.06E+00 | 7.84E+00 |
| 23830670 | 20501368 | 1544618 | 49968541 | 57170645 | 20501368 | 3.568749 | 2.232437 | 9.769842 |
| 195130.7 | 1916509 | 81951.72 | 7911899 | 389475.1 | 195130.7 | 7.342949 | 2.64E+01 | 1.34E+02 |
| 534490 | 2292931 | 87168.69 | 7833708 | 798940.7 | 998603.1 | 3.201537 | 67.846 | 71.45534 |
| 3378039 | 6545588 | 109670.8 | 9858944 | 1800297 | 3378039 | 48.31589 | 81.38661 | 85.09951 |
| 46192577 | 52838986 | 217785.9 | 77390063 | 64073996 | 46192577 | 2.399385 | 7.67E+00 | 1.73E+01 |
| 2307826 | 1322344 | 2211342 | 7356181 | 27154460 | 2307826 | 9.234391 | 2.126822 | 3.18E+01 |
| 6397281 | 5221954 | 134263.8 | 15134699 | 12227358 | 5221954 | 6.837313 | 3.59E+00 | 2.52E+01 |
| 38421444 | 40707753 | 409924.4 | 45661172 | 1.06E+08 | 38421444 | 3.746017 | 1.666965 | 1.40E+01 |
| 962414.8 | 479273.4 | 475282.9 | 836163.2 | 53972984 | 962414.8 | 21.53676 | 12.26472 | 38.4979 |
| 300886.8 | 310886.3 | 745207.4 | 470473.3 | 40903210 | 300886.8 | 6.694906 | 8.47E+00 | 6.33E+00 |
| 295773.4 | 295886 | 419849.9 | 529006.4 | 30965571 | 295773.4 | 5.479758 | 1.16E+01 | 5.16E+00 |
| 1.53E+08 | 1.59E+08 | 627161 | 1.89E+08 | 2.38E+08 | 1.53E+08 | 0.704076 | 2.630996 | 1.40E+01 |
| 2801740 | 3727436 | 197328.8 | 10229606 | 19154567 | 2801740 | 1.428619 | 11.03754 | 1.81E+01 |
| 58588407 | 42763491 | 82118.5 | 45114985 | 43551751 | 47895031 | 4.725225 | 1.85E+01 | 1.62E+01 |
| 14162308 | 13824012 | 282866.7 | 13885928 | 10029367 | 13824012 | 17.52187 | 12.18958 | 15.48597 |
| 29144012 | 33671819 | 97727.99 | 54070369 | 56039613 | 29144012 | 4.039075 | 2.11E+00 | 1.73E+01 |
| 25277552 | 25720595 | 435453.4 | 36686644 | 42003111 | 25277552 | 4.336066 | 12.63826 | 1.40E+01 |
| 1.44E+08 | 1.23E+08 | 344628.4 | 2.75E+08 | 1.66E+08 | 1.44E+08 | 1.212214 | 6.93234 | 12.24209 |
| 60525285 | 64738967 | 91413.41 | 51630420 | 32412202 | 60525285 | 12.96452 | 2.730712 | 5.94E+01 |
| 7882942 | 7096485 | 99118.97 | 12085016 | 9549515 | 7096485 | 3.426391 | 5.504235 | 9.90E+00 |
| 231450.9 | 218370.5 | 139010.5 | 22123498 | 236777.2 | 218370.5 | 4.320317 | 2.243671 | 8.788013 |
| 15176091 | 6528032 | 429056.6 | 16718116 | 16666869 | 6528032 | 55.72855 | 15.50981 | 7.98E+01 |
| 533715.3 | 1461684 | 89279.42 | 14001476 | 617783.1 | 1175105 | 5.496378 | 9.30E+01 | 4.50E+01 |
| 148569.1 | 145473.8 | 87931.99 | 13002208 | 10175499 | 145473.8 | 2.661548 | 2.291251 | 10.26661 |

| | | | | | | | | |
|---|---|---|---|---|---|---|---|---|
| 4171894 | 13137565 | 155373.3 | 26381978 | 12490131 | 4538647 | 25.48035 | 3.15E+01 | 6.97E+01 |
| 10865907 | 10985806 | 404198.3 | 4107533 | 13038531 | 10865907 | 4.944364 | 1.400633 | 13.54376 |
| 9653876 | 9175783 | 142055.3 | 9614493 | 13029912 | 9175783 | 10.07186 | 4.057902 | 13.19762 |
| 2683735 | 385708.1 | 302334.6 | 592974.2 | 5569394 | 659022 | 74.64237 | 5.73E+01 | 1.01E+02 |
| 25746429 | 20036049 | 1209861 | 29597272 | 26700242 | 20036049 | 14.40215 | 14.35324 | 2.37E+01 |
| 17365213 | 18093998 | 1487590 | 15682241 | 27627758 | 18093998 | 18.78071 | 4.79E+01 | 8.24E+00 |
| 11113683 | 8242509 | 834569.2 | 15112448 | 14508289 | 10716002 | 15.00052 | 1.55E+01 | 1.55E+01 |
| 16019353 | 17927490 | 1459380 | 18069780 | 17573576 | 16019353 | 20.53753 | 8.36E+00 | 33.11213 |
| 226518.6 | 2375944 | 201074.2 | 8521545 | 4073433 | 1043220 | 15.90457 | 16.44567 | 89.28264 |
| 7269043 | 5955394 | 252606.9 | 22888313 | 15266381 | 5955394 | 5.547307 | 5.09E+01 | 1.48E+01 |
| 3141716 | 5420732 | 1126336 | 14040141 | 8435579 | 3141716 | 7.275786 | 3.40E+00 | 3.54E+01 |
| 20437870 | 24518180 | 1505277 | 36428489 | 6527050 | 20437870 | 61.24887 | 2.55E+01 | 1.10E+01 |
| 2972592 | 2587193 | 976650.4 | 9563453 | 9545407 | 2587193 | 24.68029 | 22.37419 | 18.4812 |
| 24509454 | 20858860 | 1142360 | 32702318 | 23875926 | 20858860 | 48.11156 | 2.15E+01 | 1.20E+01 |
| 79442018 | 85706270 | 298934.9 | 288154.5 | 742195.8 | 79442018 | 15.25336 | 48.9496 | 6.128053 |
| 961023.2 | 2772956 | 118843.9 | 65985561 | 1640863 | 2772956 | 3.702226 | 6.44E+01 | 5.04E+01 |
| 64591441 | 75014218 | 1605219 | 2608360 | 14053842 | 64591441 | 116.3244 | 73.97332 | 1.17E+01 |
| 30285053 | 30388620 | 2089778 | 9487078 | 9310202 | 30285053 | 115.3208 | 2.987596 | 8.760735 |
| 10498378 | 10446854 | 494433.4 | 12544099 | 12250268 | 10446854 | 21.10464 | 5.361831 | 6.785176 |
| 4373715 | 3799255 | 120175.6 | 8558063 | 5707769 | 4373715 | 12.8007 | 3.249245 | 8.914941 |
| 35962548 | 38793014 | 163268.6 | 4700421 | 1838158 | 35962548 | 38.39911 | 69.58011 | 12.86191 |
| 34057703 | 39150925 | 2221904 | 57739416 | 41776106 | 39150925 | 45.66665 | 6.807614 | 17.80879 |
| 15055672 | 16394909 | 1327782 | 40156855 | 34481446 | 15055672 | 20.09399 | 16.24975 | 16.37511 |
| 2138086 | 7264977 | 149904.2 | 12755992 | 12288855 | 5681081 | 27.19973 | 5.695915 | 52.20891 |
| 15510492 | 15258322 | 88790.87 | 1178313 | 664586.3 | 15258322 | 67.81945 | 62.97809 | 9.60E+00 |
| 19921746 | 20693901 | 97401.2 | 1686387 | 505980.2 | 19921746 | 43.9972 | 22.12809 | 8.85E+00 |
| 18180347 | 10142408 | 84866.31 | 668619.3 | 573172.7 | 14671816 | 43.03217 | 2.16E+01 | 2.81E+01 |
| 20721970 | 23419472 | 242967.8 | 9906887 | 8258545 | 20721970 | 2.413457 | 5.33E+01 | 1.62E+01 |
| 1261181 | 1158410 | 116668.2 | 3379798 | 13217295 | 1158410 | 6.42102 | 5.036057 | 9.149935 |
| 198049.6 | 204254.7 | 178758.3 | 1859695 | 7052926 | 204254.7 | 12.48089 | 12.37232 | 35.45762 |
| 2026222 | 3870052 | 175405.4 | 83937220 | 597120.9 | 2026222 | 2.926005 | 7.37E+01 | 8.30E+01 |
| 15302978 | 14766171 | 301865.9 | 11713914 | 10014829 | 14766171 | 58.25447 | 46.99242 | 9.56E+00 |
| 4561235 | 2129044 | 787089.7 | 6.09E+08 | 2753433 | 2129044 | 2.779309 | 44.86014 | 6.75E+01 |
| 400438.5 | 2869034 | 120744.7 | 1.73E+08 | 920810.8 | 1056778 | 3.739692 | 35.81864 | 8.87E+01 |
| 50019707 | 73575342 | 798636.4 | 2.59E+08 | 98663646 | 65957533 | 37.18202 | 63.0837 | 19.02408 |
| 97688337 | 1.02E+08 | 522437 | 1.28E+08 | 1.29E+08 | 97688337 | 3.699325 | 2.384058 | 1.10E+01 |
| 26874580 | 43048868 | 657144.7 | 2.68E+08 | 59398159 | 43048868 | 61.88998 | 5.282555 | 43.01859 |
| 22991768 | 22683677 | 352212.1 | 2.87E+08 | 41153302 | 22991768 | 5.591611 | 9.709329 | 18.19779 |
| 5514164 | 5875192 | 163070.5 | 1.54E+08 | 32562629 | 5514164 | 1.636131 | 3.313995 | 1.45E+01 |
| 1901605 | 1837448 | 2482274 | 1150221 | 69708039 | 1901605 | 38.25941 | 1.34E+01 | 6.56E+00 |
| 55807250 | 53136061 | 551886.7 | 87058934 | 79637911 | 53136061 | 2.551725 | 2.939524 | 8.20409 |
| 3498368 | 4321476 | 166085.2 | 17817150 | 16077006 | 3498368 | 37.93836 | 8.337523 | 25.56658 |
| 341424.7 | 324971.8 | 159650.9 | 99998214 | 1912417 | 341424.7 | 1.481197 | 81.95065 | 1.35E+02 |
| 6247797 | 6474183 | 741852.8 | 64709096 | 49410656 | 6247797 | 3.905239 | 8.411092 | 8.231664 |
| 3302765 | 5609072 | 177904.6 | 1.04E+08 | 2749480 | 3302765 | 1.988667 | 74.11255 | 5.38E+01 |
| 740463.9 | 476156.4 | 215413.6 | 1.66E+08 | 1603212 | 612179.9 | 4.046146 | 63.42438 | 21.68186 |
| 493281 | 633456.8 | 98854.02 | 63681034 | 402295.4 | 493281 | 4.919767 | 23.14141 | 26.40527 |
| 32148848 | 28916786 | 103576.8 | 60212578 | 61567778 | 28916786 | 4.041852 | 11.34547 | 7.013378 |
| 298186.1 | 203562.4 | 135721.9 | 65079983 | 406013.1 | 203562.4 | 3.998506 | 3.25E+01 | 2.80E+01 |
| 83081521 | 20685818 | 827305.2 | 1.41E+08 | 1.08E+08 | 20685818 | 10.53757 | 5.08E+01 | 8.83E+01 |
| 67254833 | 60008349 | 116168.9 | 21421561 | 14995351 | 60008349 | 2.527914 | 1.659339 | 8.16E+00 |

| | | | | | | | | |
|---|---|---|---|---|---|---|---|---|
| 1630008 | 2834261 | 120342.2 | 55526054 | 612835.4 | 1630008 | 2.980035 | 20.62388 | 7.73E+01 |
| 7087890 | 6967123 | 99922.94 | 49432567 | 19358483 | 6967123 | 2.507139 | 12.01013 | 8.52E+00 |
| 46486291 | 44531704 | 8351729 | 1.2E+08 | 96540955 | 44531704 | 7.187975 | 13.50297 | 8.55E+00 |
| 22518671 | 28963979 | 2372143 | 1.1E+08 | 22072147 | 25729099 | 6.520333 | 12.89156 | 12.52139 |
| 49007000 | 51278762 | 321976.8 | 2.05E+08 | 1.47E+08 | 49007000 | 2.93671 | 0.553388 | 6.89874 |
| 41321677 | 57555445 | 4827900 | 98920601 | 73625052 | 41321677 | 2.408935 | 3.739526 | 3.25E+01 |
| 9401529 | 8476715 | 59090.89 | 69187720 | 10481786 | 8476715 | 27.73419 | 75.23436 | 70.36132 |
| 15224224 | 15199563 | 282439.1 | 29162736 | 17564810 | 15199563 | 5.843541 | 5.57235 | 7.636707 |
| 4577703 | 4693141 | 611538.4 | 50150435 | 10086963 | 4577703 | 36.77824 | 10.92092 | 1.7574 |
| 758075.1 | 743362.2 | 100550.3 | 30730151 | 1286111 | 758075.1 | 7.166443 | 69.56251 | 101.0634 |
| 36087400 | 41224025 | 122668.3 | 43269454 | 34691689 | 36087400 | 7.40113 | 9.561694 | 14.93124 |
| 5303581 | 5152887 | 231408.7 | 78185407 | 8858797 | 5303581 | 2.763643 | 10.74891 | 2.80646 |
| 12109943 | 9511131 | 107530.8 | 1.1E+08 | 10433438 | 9511131 | 1.185252 | 27.92293 | 33.11892 |
| 42188470 | 36636723 | 7645854 | 1.61E+08 | 64631582 | 40930796 | 3.064459 | 34.0994 | 7.292269 |
| 59416041 | 53659986 | 7965746 | 1.26E+08 | 96591391 | 59416041 | 18.24362 | 2.413549 | 13.67152 |
| 4336475 | 3854139 | 81412.77 | 44789834 | 25242581 | 4336475 | 1.593484 | 2.18E+01 | 8.51E+00 |
| 13336727 | 13472109 | 613971.3 | 23874606 | 22748438 | 13336727 | 8.30633 | 8.93E+00 | 7.21E+00 |
| 156532.7 | 144889 | 152658.3 | 42925205 | 9924800 | 156532.7 | 5.35871 | 2.92E+00 | 1.03E+01 |
| 2237103 | 3078505 | 174938.6 | 28283462 | 4640631 | 2237103 | 79.60791 | 6.84E+01 | 26.1912 |
| 17640330 | 16088846 | 1489688 | 70820496 | 15019429 | 16088846 | 9.862931 | 1.87E+00 | 1.05E+01 |
| 32167045 | 29194089 | 101772.5 | 18853756 | 21921579 | 29194089 | 4.226501 | 6.590494 | 7.73E+00 |
| 20950579 | 25749603 | 202316.8 | 82025968 | 16420818 | 20950579 | 7.361276 | 6.08E+00 | 1.54E+01 |
| 39449059 | 46301414 | 137562 | 50164995 | 37169932 | 39449059 | 8.17669 | 2.304828 | 1.57E+01 |
| 16265871 | 15290985 | 84457.25 | 43471512 | 39769237 | 15290985 | 0.718701 | 6.885033 | 12.34774 |
| 2764964 | 6917193 | 257353.8 | 88642143 | 3422370 | 2764964 | 3.361656 | 7.19E+01 | 9.17E+01 |
| 29889094 | 29390063 | 87062.1 | 3744004 | 2210903 | 29390063 | 5.84884 | 1.96E+00 | 7.88E+00 |
| 35663899 | 36096549 | 101555.4 | 10545939 | 11241319 | 35663899 | 5.228479 | 68.65362 | 9.869301 |
| 34193773 | 33971445 | 98429.51 | 8470315 | 190372.8 | 34193773 | 48.615 | 94.73762 | 4.235988 |
| 9648789 | 9878239 | 205807 | 14585447 | 15867733 | 9878239 | 64.1594 | 3.86E+01 | 1.73E+00 |
| 5300555 | 4908137 | 115906.3 | 13957795 | 11421667 | 4908137 | 4.155007 | 8.83329 | 2.22E+01 |
| 17116567 | 38482070 | 1069346 | 1.2E+08 | 77821732 | 33453280 | 2.506718 | 26.77646 | 37.63095 |
| 813421.8 | 2878846 | 94459.89 | 34148166 | 2297867 | 2584673 | 3.480157 | 39.5773 | 53.39911 |
| 8891757 | 10267631 | 126155.2 | 44065368 | 11715703 | 9152981 | 3.748454 | 5.99022 | 7.742767 |
| 1008206 | 614203.4 | 111192.3 | 13329107 | 26739365 | 698101.7 | 3.995581 | 8.381265 | 26.83136 |
| 15490750 | 17840286 | 2239922 | 84895530 | 14574825 | 16209473 | 2.570511 | 5.865379 | 7.290473 |
| 44299040 | 46370479 | 230361.5 | 1.72E+08 | 86692062 | 44299040 | 37.26872 | 3.07E+01 | 3.02E+01 |
| 862092.4 | 853005.2 | 262347.6 | 25741352 | 9315196 | 862092.4 | 11.70847 | 8.281231 | 5.28E+01 |
| 1198795 | 7245082 | 239795.8 | 48223024 | 53907573 | 7245082 | 65.49401 | 8.123506 | 81.46867 |
| 4512801 | 2651809 | 64234.4 | 60773463 | 5168097 | 2651809 | 10.69745 | 4.33E+01 | 7.40E+01 |
| 179910.5 | 210751.4 | 110497.8 | 17832244 | 207972.2 | 210751.4 | 6.632186 | 30.68436 | 2.18E+01 |
| 2677198 | 2687479 | 308186.1 | 7778816 | 6999548 | 2677198 | 95.65997 | 17.99093 | 1.782886 |
| 38179604 | 46231811 | 443155.6 | 27171631 | 19996597 | 43927857 | 0.508393 | 6.064496 | 9.69E+00 |
| 17078654 | 12840057 | 3603645 | 66817642 | 14986374 | 16452043 | 3.281135 | 2.53E+01 | 1.48E+01 |
| 26816536 | 30032037 | 6554950 | 97356162 | 16812740 | 26816536 | 2.435003 | 24.37286 | 1.03E+01 |
| 781853.3 | 209911.3 | 56044.64 | 25155645 | 255194.8 | 209911.3 | 3.85492 | 28.40991 | 83.03851 |
| 2434352 | 2724935 | 102279.8 | 14111461 | 16949810 | 2622965 | 9.005412 | 2.739165 | 5.68E+00 |
| 5288550 | 5986300 | 98557.48 | 30867408 | 7283409 | 5288550 | 3.34351 | 1.679954 | 1.62E+01 |
| 5883571 | 5820686 | 127251.5 | 21920353 | 11033560 | 5820686 | 6.753331 | 18.36876 | 2.41E+01 |
| 1359464 | 3616385 | 102719.3 | 31006523 | 13478570 | 2661581 | 0.591715 | 2.41499 | 44.50081 |
| 23301067 | 24458256 | 89846.06 | 12350744 | 10691775 | 23301067 | 6.438828 | 4.511101 | 14.21203 |
| 590900.6 | 477305.2 | 89531.78 | 25405577 | 573729 | 477305.2 | 14.31324 | 71.63041 | 34.33502 |

| | | | | | | | | |
|---|---|---|---|---|---|---|---|---|
| 41577267 | 43263875 | 138723.6 | 18531156 | 16798410 | 41577267 | 6.668295 | 6.98E+00 | 4.07E+00 |
| 660539.4 | 1011219 | 93348.1 | 25584240 | 4510583 | 1011219 | 0.672682 | 5.72E+00 | 8.78E+01 |
| 221212.2 | 128832.6 | 87530.9 | 25328668 | 147966 | 221212.2 | 12.19606 | 42.90625 | 35.97528 |
| 3153113 | 1342883 | 49703.67 | 25008318 | 2432869 | 1342883 | 60.79368 | 78.93173 | 55.2967 |
| 16298977 | 14643888 | 716159.1 | 20766198 | 19544266 | 14643888 | 4.76027 | 9.101394 | 6.76673 |
| 13302624 | 10638896 | 389158.6 | 35669664 | 9878117 | 10638896 | 5.743549 | 6.650907 | 66.18718 |
| 1254135 | 4929658 | 818039.9 | 58037368 | 24158448 | 4929658 | 7.405593 | 70.175 | 57.35129 |
| 323942.5 | 1669457 | 84124.26 | 20140831 | 590382.1 | 323942.5 | 3.081993 | 41.89755 | 118.9315 |
| 8453459 | 9422508 | 107868.8 | 21255317 | 22572120 | 8453459 | 10.6101 | 3.952812 | 1.31E+01 |
| 10481551 | 11171043 | 261644.9 | 40079760 | 10328563 | 11171043 | 24.39507 | 65.4241 | 8.019881 |
| 306290.6 | 735259.7 | 77024.98 | 17894912 | 148870 | 339612.8 | 2.97922 | 14.84643 | 51.83207 |
| 351609.5 | 457368.2 | 102240.7 | 27598786 | 2031247 | 457368.2 | 3.256748 | 64.2661 | 54.12139 |
| 22622313 | 30470307 | 448583.7 | 29310368 | 12867849 | 26448881 | 23.06595 | 37.48127 | 14.80133 |
| 23267398 | 23449161 | 114306 | 35351719 | 40054312 | 23267398 | 5.648186 | 5.793863 | 5.68E+00 |
| 180909.2 | 170859.8 | 88139.75 | 22492077 | 266823.5 | 180909.2 | 3.328913 | 80.57643 | 9.847287 |
| 5597100 | 5579099 | 462036.9 | 22679476 | 4334767 | 5579099 | 5.549443 | 24.06812 | 30.05418 |
| 7355269 | 9454807 | 1677609 | 28716107 | 18827321 | 7355269 | 48.10256 | 4.630578 | 15.77767 |
| 17665082 | 17680351 | 82630.02 | 1942511 | 6422436 | 17665082 | 4.560731 | 10.47081 | 14.03489 |
| 2219158 | 2602922 | 103676.4 | 9573031 | 15758403 | 2235832 | 1.523613 | 6.332576 | 9.219999 |
| 157246.3 | 153042.1 | 129752.3 | 21960479 | 5714647 | 153042.1 | 4.418851 | 3.949588 | 6.05E+00 |
| 7245261 | 5848744 | 119916.1 | 29045069 | 13073325 | 6008790 | 5.182828 | 4.060894 | 12.00263 |
| 9499870 | 11384252 | 79534.59 | 12771634 | 16776191 | 9499870 | 9.624743 | 3.407388 | 1.59E+01 |
| 9140055 | 9485099 | 2207685 | 33364964 | 12185008 | 9140055 | 16.5116 | 3.00E+00 | 9.80E+00 |
| 645168.7 | 831006.9 | 83825.16 | 17605866 | 5181505 | 822423.9 | 2.904672 | 6.662726 | 13.69144 |
| 6365797 | 7377275 | 646572.9 | 32086021 | 5173964 | 7377275 | 4.682921 | 1.624515 | 1.04E+01 |
| 396465.5 | 164398.6 | 87742.98 | 15237276 | 704411 | 164398.6 | 8.356982 | 48.92558 | 58.21066 |
| 4253810 | 4792222 | 260700.1 | 8581356 | 7730884 | 4253810 | 6.043923 | 10.5769 | 16.85041 |
| 5210800 | 3635303 | 116428.9 | 23442446 | 12097788 | 3635303 | 4.876576 | 29.14337 | 22.16324 |
| 16781770 | 18229908 | 69407.24 | 1002080 | 2046836 | 16821133 | 64.79796 | 101.6141 | 4.774703 |
| 2302919 | 3747084 | 2591861 | 1980556 | 5062579 | 3747084 | 18.10922 | 1.25E+02 | 4.00E+01 |
| 5614003 | 8225161 | 284047.4 | 57029667 | 48124332 | 5738558 | 40.04416 | 40.2792 | 22.57026 |
| 13387607 | 13417024 | 479225.8 | 68550754 | 19864437 | 13387607 | 28.41801 | 1.708773 | 9.70E-01 |
| 24484609 | 24611311 | 103241.8 | 3465897 | 1785310 | 24484609 | 78.4208 | 1.42E+02 | 2.26E+01 |
| 1432365 | 1536254 | 521981 | 22709281 | 12775700 | 1536254 | 20.78652 | 25.86613 | 1.26E+02 |
| 12241680 | 11527238 | 1581822 | 15978466 | 16345660 | 11527238 | 34.76366 | 9.155508 | 7.462803 |
| 197666.1 | 197186.9 | 121695.4 | 16750871 | 256526.3 | 197186.9 | 5.402893 | 35.95725 | 8.961695 |
| 14740706 | 12389211 | 2920405 | 31116771 | 27919312 | 13311751 | 5.497318 | 8.499015 | 8.788963 |
| 8782831 | 8112350 | 80553.7 | 10557182 | 3703502 | 8112350 | 14.56819 | 21.97328 | 5.38E+00 |
| 9746550 | 9334724 | 39736.31 | 270442 | 1135459 | 9746550 | 114.635 | 34.20765 | 6.553143 |
| 8097600 | 8030761 | 507120.2 | 66798742 | 23743069 | 8030761 | 35.98155 | 5.37E+00 | 4.51E+00 |
| 2296894 | 2657296 | 113711.2 | 19815995 | 18210286 | 2296894 | 2.389289 | 1.04E+01 | 1.95E+01 |
| 353491.4 | 135215.3 | 60575.11 | 9862690 | 1760640 | 342850.9 | 2.500692 | 9.548713 | 44.39808 |
| 10454311 | 10184823 | 71289.62 | 1083217 | 830615.9 | 10184823 | 10.77597 | 12.61456 | 7.268694 |
| 708999.9 | 487449 | 93191.1 | 15125737 | 352744.8 | 487449 | 19.74039 | 3.28E+01 | 5.09E+01 |
| 13427193 | 12546698 | 220701.3 | 47007150 | 38186692 | 12568962 | 14.54226 | 6.24E+00 | 3.91E+00 |
| 18241680 | 18893512 | 137226.9 | 20381954 | 20477258 | 18241680 | 0.55308 | 2.423929 | 7.19E+00 |
| 15456751 | 18903711 | 160432.9 | 22167254 | 28659385 | 15456751 | 1.359284 | 27.65894 | 5.99E+01 |
| 20216937 | 21029393 | 189987.3 | 22365420 | 17460843 | 20216937 | 6.040987 | 33.29517 | 8.132492 |
| 5027348 | 4939437 | 114228.8 | 23486428 | 10715920 | 4939437 | 8.393857 | 4.76333 | 10.96465 |
| 412559.8 | 324420.5 | 83518.88 | 13411433 | 352887.1 | 412559.8 | 18.31039 | 29.72772 | 118.9622 |
| 495696.8 | 789119.1 | 127338.6 | 13019089 | 719393.5 | 495696.8 | 5.333669 | 68.60274 | 36.94023 |

| | | | | | | | | |
|---|---|---|---|---|---|---|---|---|
| 694606.5 | 733216.1 | 67705.99 | 9176130 | 7948228 | 694606.5 | 6.205913 | 1.15E+01 | 5.13E+00 |
| 18955747 | 19650290 | 218462.2 | 14471259 | 14223428 | 18955747 | 1.28458 | 1.306935 | 8.504977 |
| 3643910 | 8758824 | 105227.6 | 11892916 | 18775059 | 3643910 | 70.67132 | 46.69052 | 57.48285 |
| 17264497 | 15760886 | 2202627 | 19476217 | 18542833 | 16588134 | 42.57367 | 5.763325 | 4.553596 |
| 1797004 | 1128865 | 129684.1 | 12127074 | 393588.4 | 1128865 | 33.68571 | 25.61271 | 34.80749 |
| 23876198 | 17065486 | 144523.9 | 13391684 | 35123935 | 23876198 | 7.073104 | 27.07333 | 32.13069 |
| 12765504 | 11469341 | 120804.6 | 13489003 | 17302316 | 11469341 | 20.14676 | 0.812281 | 15.11417 |
| 3184129 | 2951740 | 75499.09 | 17951356 | 7515762 | 2951740 | 5.303394 | 12.77867 | 7.750365 |
| 135855.4 | 74825.13 | 67285.79 | 7536217 | 8592150 | 124954 | 25.76119 | 12.35847 | 29.09281 |
| 13680742 | 13237457 | 127172.2 | 44889715 | 21533503 | 13237457 | 5.211939 | 7.233561 | 1.36E+01 |
| 1164563 | 3335480 | 245980.8 | 25320813 | 19887604 | 1265964 | 21.43349 | 9.128542 | 63.74378 |
| 3388819 | 4862971 | 70162.46 | 29045069 | 13073325 | 4862971 | 5.182828 | 4.060894 | 20.49388 |
| 684257.9 | 378127.6 | 99100.6 | 17927798 | 6197549 | 684257.9 | 8.145934 | 3.67E+01 | 8.10E+01 |
| 5260671 | 5622214 | 108462.9 | 6784103 | 7749317 | 5260671 | 5.906474 | 10.96145 | 1.36E+01 |
| 659466.4 | 5645734 | 149627 | 19727528 | 668174.1 | 681621.9 | 16.2863 | 1.27E+02 | 123.3371 |
| 17874708 | 17988691 | 226509.2 | 35553381 | 32114160 | 17988691 | 2.548948 | 22.2187 | 6.101969 |
| 6258341 | 5748348 | 52778.62 | 5167336 | 5483081 | 5748348 | 8.90369 | 9.538212 | 9.773033 |
| 1687449 | 1585973 | 39932.55 | 6544672 | 2429152 | 1593457 | 18.00644 | 3.237127 | 3.485852 |
| 233713.4 | 124532.9 | 88762.41 | 674794.7 | 9871772 | 233713.4 | 54.74667 | 1.776623 | 37.53184 |
| 8176501 | 8017587 | 227231.1 | 22490415 | 20042148 | 8017587 | 3.116099 | 9.831868 | 9.844593 |
| 5382639 | 5353610 | 100993.4 | 16112498 | 14472536 | 5353610 | 15.86913 | 6.72E+00 | 1.06E+01 |
| 1263651 | 1209569 | 93015.39 | 9561659 | 7455784 | 1209569 | 4.886846 | 56.79371 | 8.15E+00 |
| 505121.1 | 1383804 | 103859 | 13228806 | 1401996 | 505121.1 | 15.68937 | 89.31309 | 71.83994 |
| 360092.2 | 434148.2 | 80225.07 | 10704260 | 829376.4 | 360092.2 | 14.19971 | 81.62191 | 42.5653 |
| 10697443 | 10771179 | 1106443 | 9358386 | 12128601 | 10697443 | 5.924764 | 7.378065 | 6.50E+00 |
| 4959342 | 5422347 | 92063.59 | 15783794 | 12840369 | 4959342 | 19.95464 | 26.81158 | 6.918665 |
| 11593917 | 11890703 | 88922.78 | 892970.3 | 184160 | 11593917 | 18.09689 | 52.08572 | 14.20683 |
| 2309823 | 2662214 | 106086.9 | 12264239 | 9181118 | 2309823 | 2.686598 | 5.15E+00 | 1.18E+01 |
| 316945.5 | 397991.6 | 109485.1 | 13249165 | 592334.2 | 324027.7 | 12.62365 | 3.04E+01 | 1.30E+01 |
| 745098 | 410171.1 | 88858.79 | 9043370 | 11073528 | 410171.1 | 3.919193 | 8.960096 | 65.26904 |
| 6542222 | 5545309 | 102541.9 | 6507737 | 7120549 | 5545309 | 5.678527 | 1.14E+00 | 1.21E+01 |
| 430923.5 | 275105.3 | 94056.56 | 3655958 | 8493049 | 335722.1 | 21.10918 | 2.913499 | 22.61948 |
| 2841514 | 3743336 | 167530.7 | 4760243 | 8405599 | 2920721 | 7.108971 | 14.20813 | 1.58E+01 |
| 20385262 | 18109604 | 2454653 | 26969132 | 67357298 | 18109604 | 42.05047 | 7.00E+01 | 7.41E+00 |
| 5165406 | 4496608 | 73502.35 | 10818936 | 9872635 | 4496608 | 8.190965 | 6.042121 | 18.53829 |
| 335437.4 | 265396.4 | 165312.5 | 10965715 | 2622373 | 335437.4 | 0.687436 | 1.64E+01 | 9.60E+01 |
| 1343014 | 782814.3 | 112718.2 | 18243345 | 4195716 | 1343014 | 4.691696 | 72.3868 | 1.10E+02 |
| 9729088 | 8422777 | 122590.6 | 6960344 | 12145532 | 8422777 | 16.94043 | 27.46732 | 9.763094 |
| 16369425 | 17251593 | 858965.8 | 38290061 | 14983790 | 16369425 | 24.62253 | 3.62E+00 | 7.57E+00 |
| 21491780 | 8055368 | 2201954 | 16772693 | 27393392 | 8055368 | 5.826953 | 5.73E+01 | 6.62E+01 |
| 366443 | 317018.3 | 73964.31 | 8169281 | 7238983 | 317018.3 | 7.381172 | 1.94E+01 | 1.10E+01 |
| 4826401 | 5022583 | 187685.3 | 2494848 | 1165775 | 4826401 | 15.26305 | 1.683928 | 17.92802 |
| 2156707 | 1481331 | 100951.6 | 12498318 | 1377977 | 1481331 | 24.93023 | 7.27E+01 | 5.64E+01 |
| 6279627 | 10262156 | 146211.5 | 22740135 | 8226945 | 10262156 | 5.095255 | 20.62471 | 30.70967 |
| 12879531 | 12795679 | 113355.3 | 2656884 | 976665 | 12795679 | 45.13332 | 78.8291 | 6.081005 |
| 4425162 | 4408446 | 36882.55 | 71682.11 | 135343.9 | 4408446 | 14.77918 | 42.67054 | 8.981019 |
| 1477581 | 1643138 | 99587.01 | 10877987 | 6908048 | 1477581 | 14.08696 | 34.24647 | 8.692862 |
| 11970141 | 12877981 | 254727.4 | 2044184 | 3177000 | 11970141 | 84.45986 | 41.97266 | 6.868488 |
| 4652633 | 1618946 | 552756.2 | 24658626 | 9402777 | 2263031 | 76.31682 | 2.76E+01 | 5.62E+01 |
| 4174400 | 1361574 | 97324.64 | 11334204 | 2433436 | 2587842 | 11.92793 | 13.71119 | 5.21E+01 |
| 40783.98 | 54459.07 | 158352.6 | 4511029 | 1472750 | 54459.07 | 13.14582 | 6.505739 | 38.31242 |

| | | | | | | | | |
|---|---|---|---|---|---|---|---|---|
| 412559.8 | 324420.5 | 91905.81 | 13411433 | 352887.1 | 412559.8 | 18.31039 | 29.72772 | 1.17E+02 |
| 6658611 | 5725013 | 61130.38 | 338542.2 | 558617.6 | 5725013 | 84.106 | 12.70899 | 10.43558 |
| 2234856 | 2049768 | 817454.4 | 11611330 | 3772107 | 2049768 | 13.3434 | 27.58033 | 58.69588 |
| 20602148 | 17753577 | 167532.5 | 13487464 | 8992616 | 17753577 | 5.842655 | 19.0904 | 1.28E+01 |
| 3397288 | 2691959 | 87041.03 | 6712764 | 7332165 | 2691959 | 6.932788 | 2.26E+01 | 1.40E+01 |
| 1350178 | 10716812 | 83202 | 12952799 | 14555268 | 4295234 | 12.79292 | 6.67E-01 | 8.78E+01 |
| 6510809 | 6717062 | 54992.95 | 2570895 | 1830170 | 6717062 | 49.55875 | 65.13824 | 3.592346 |
| 5905107 | 6967540 | 50278.52 | 12927375 | 10792155 | 5905107 | 8.517624 | 28.91037 | 7.19E+01 |
| 11347505 | 5787252 | 115710 | 16063609 | 15317056 | 6115125 | 0.887275 | 1.013044 | 40.25666 |
| 3925739 | 3419654 | 85083.48 | 1110279 | 6209103 | 3419654 | 9.658152 | 1.718513 | 9.98E+00 |
| 3778402 | 3479305 | 77067.85 | 14940110 | 7612087 | 3479305 | 7.766211 | 1.706418 | 6.456224 |
| 25405913 | 15209191 | 237994.2 | 17420450 | 25215875 | 15209191 | 27.47522 | 58.98884 | 41.06741 |
| 19546267 | 20866487 | 160853.4 | 211390.5 | 207760.2 | 19546267 | 11.394 | 4.40E+00 | 1.17E+01 |
| 3196109 | 459495.4 | 69156.97 | 4799800 | 9916131 | 3196109 | 13.9212 | 5.247958 | 7.06E+01 |
| 6567438 | 6755314 | 78104.29 | 6490049 | 4889066 | 6567438 | 3.147676 | 12.36775 | 3.159332 |
| 25326848 | 12424723 | 377479.9 | 13812935 | 11621243 | 12424723 | 2.976802 | 16.60474 | 50.16981 |
| 2943620 | 3419817 | 77480.68 | 7045514 | 4731275 | 2943620 | 2.660749 | 8.034123 | 19.85217 |
| 8509640 | 8185402 | 122288 | 14948145 | 11031873 | 8185402 | 1.013924 | 1.40E+01 | 9.51E+00 |
| 10707385 | 11089017 | 145811.9 | 14333378 | 13757226 | 10707385 | 2.381481 | 5.21E+00 | 1.23E+01 |
| 264800.6 | 519732 | 66040.53 | 508602.3 | 442097.5 | 366344.8 | 32.54099 | 12.64472 | 33.4548 |
| 128157.4 | 196240.2 | 68180.98 | 3677190 | 4271195 | 128157.4 | 7.720812 | 33.10947 | 34.9905 |
| 1059088 | 1324193 | 92015.23 | 14092770 | 982539.3 | 1176419 | 9.116882 | 1.35E+02 | 1.12E+01 |
| 6998825 | 5146703 | 50503.58 | 6453516 | 8906519 | 5146703 | 23.45789 | 16.2198 | 3.04E+01 |
| 5937858 | 5564478 | 127979.1 | 13481296 | 9485671 | 5564478 | 3.16009 | 4.645134 | 8.75E+00 |
| 239065.9 | 158048.9 | 86926.4 | 8463894 | 311185.5 | 187091.1 | 5.095192 | 108.0431 | 21.0778 |
| 247910.3 | 716017.5 | 25480.38 | 10115551 | 1799900 | 351871.6 | 65.67638 | 51.84808 | 56.04419 |
| 2344484 | 2515013 | 56080.61 | 11839991 | 5157795 | 2344484 | 6.893795 | 31.76477 | 2.06E+01 |
| 178573.6 | 209804.3 | 127420.4 | 1548530 | 11164561 | 178573.6 | 5.822098 | 4.697949 | 23.14864 |
| 3184129 | 2966288 | 77791.3 | 17951356 | 7515762 | 2966288 | 5.303394 | 12.77867 | 7.740401 |
| 9578723 | 10339141 | 89878.85 | 2155799 | 2947702 | 9578723 | 13.88928 | 2.21E+01 | 1.08E+01 |
| 17209124 | 17344283 | 1289803 | 4579275 | 3787725 | 17209124 | 21.44246 | 68.39397 | 7.67E+00 |
| 227686.2 | 185323.8 | 115039 | 9621220 | 8625771 | 227631.8 | 5.330628 | 8.458468 | 11.44583 |
| 144874.9 | 143315.8 | 133283.6 | 15255965 | 2327187 | 143354.3 | 2.245266 | 5.43E+00 | 6.18E-01 |
| 39855900 | 38568999 | 481332.6 | 32711055 | 29433641 | 38568999 | 3.970403 | 1.07E+00 | 7.39E+00 |
| 4127015 | 4394619 | 161070.9 | 16219406 | 7719191 | 4394619 | 34.57754 | 76.78358 | 23.93768 |
| 448883.1 | 1623171 | 97890.09 | 9687703 | 812117.1 | 1623171 | 22.87518 | 1.54E+01 | 7.61E+01 |
| 1292252 | 2255394 | 142800.8 | 7181132 | 3192387 | 1292252 | 4.344705 | 50.0536 | 51.69428 |
| 6660015 | 9334724 | 54540.36 | 406758 | 1135459 | 9334724 | 103.9035 | 34.20765 | 22.71425 |
| 10771675 | 9487813 | 167672.6 | 13568731 | 13981251 | 9849280 | 0.629351 | 11.1118 | 6.596474 |
| 1580141 | 1715753 | 87866.63 | 6973288 | 2820633 | 1580141 | 4.262766 | 12.27901 | 9.05E+00 |
| 12456255 | 13049680 | 123122.5 | 9848147 | 10189053 | 12456255 | 5.090409 | 3.977941 | 9.337757 |
| 2146233 | 1902766 | 89735.96 | 6923315 | 6827085 | 1902766 | 4.09542 | 2.04E+01 | 1.45E+01 |
| 16103525 | 17234050 | 143238.7 | 12534534 | 2310926 | 16103525 | 1.40584 | 112.9859 | 9.413288 |
| 2816116 | 3179767 | 123847.9 | 9239164 | 7635412 | 2816116 | 25.22166 | 3.002543 | 8.243709 |
| 1573607 | 1561988 | 50862.42 | 8535642 | 3140031 | 1561988 | 11.5466 | 12.72636 | 13.14131 |
| 5075954 | 4851142 | 50389.08 | 239158 | 209318.2 | 5075954 | 90.07692 | 35.256 | 15.60732 |
| 5680078 | 6235461 | 316019.3 | 9313733 | 10491537 | 5680078 | 10.30473 | 40.60294 | 1.30E+01 |
| 9558583 | 9798784 | 127677.5 | 2747839 | 5065722 | 9558583 | 4.635434 | 6.42907 | 6.036902 |
| 86088.22 | 86818.97 | 76330.18 | 5858564 | 5097157 | 86088.22 | 6.258943 | 15.92691 | 4.20E+00 |
| 4069831 | 803859.2 | 65571.7 | 1955928 | 8379187 | 905945.7 | 61.01105 | 10.35897 | 96.38193 |
| 399798.9 | 320551.8 | 100037.2 | 8835657 | 761570.5 | 399798.9 | 9.775344 | 63.43931 | 61.43531 |

| | | | | | | | | |
|---|---|---|---|---|---|---|---|---|
| 97931.27 | 101460.9 | 91794.98 | 6252865 | 7442537 | 100112.2 | 4.534676 | 14.75892 | 1.78E+00 |
| 1383508 | 2447830 | 123807.5 | 12966481 | 5489123 | 1508408 | 1.798259 | 11.09473 | 32.68653 |
| 7076301 | 5462754 | 289784.5 | 22809031 | 20880196 | 7076301 | 10.68837 | 1.67E+01 | 2.59E+01 |
| 13857739 | 11549292 | 164037 | 11281237 | 19027199 | 11549292 | 3.456792 | 0.848672 | 13.56641 |
| 27927428 | 14796338 | 302179.1 | 21706074 | 20818151 | 22516011 | 48.7533 | 44.23102 | 30.34623 |
| 1191491 | 1297654 | 54528.58 | 5547434 | 2594083 | 1191491 | 13.56064 | 6.275966 | 1.29E+01 |
| 5480554 | 548633.6 | 71686.6 | 12768799 | 2420727 | 2484956 | 19.7799 | 9.50E+01 | 87.55479 |
| 2447003 | 2575103 | 81796.58 | 6346824 | 5649213 | 2447003 | 4.733203 | 4.309525 | 16.19488 |
| 384024 | 480085.7 | 255070.7 | 10381719 | 618701.8 | 480085.7 | 2.385885 | 42.53452 | 3.79E+01 |
| 6094204 | 7405362 | 95979.66 | 237560.6 | 182044.5 | 6094204 | 3.739296 | 1.85E+01 | 1.84E+01 |
| 135891.9 | 136864.2 | 86018.62 | 6491763 | 151415.5 | 135891.9 | 3.158212 | 2.611902 | 8.41E+00 |
| 6090597 | 6659871 | 91865.74 | 1958859 | 1179803 | 6659871 | 24.53247 | 88.16372 | 7.994195 |
| 1367452 | 1271554 | 152790.6 | 12817383 | 5570102 | 1367452 | 7.731107 | 3.33E+01 | 4.68E+01 |
| 11314124 | 11387013 | 218506 | 29070752 | 15728319 | 11314124 | 38.98614 | 55.66849 | 10.37187 |
| 1917379 | 1773225 | 156235.2 | 15983600 | 14099674 | 1773225 | 4.283704 | 2.39E+00 | 1.44E+01 |
| 24484609 | 24611311 | 474954 | 6844329 | 6500306 | 24484609 | 46.44912 | 2.361875 | 2.26E+01 |
| 1161990 | 1586129 | 123323.8 | 6898814 | 2298392 | 1197066 | 10.5664 | 20.0576 | 17.90069 |
| 5027348 | 4939437 | 124000.8 | 23486428 | 10715920 | 4939437 | 8.393857 | 4.76333 | 10.96465 |
| 3067041 | 3239114 | 86848.21 | 4416772 | 5533117 | 3099360 | 5.034295 | 4.90E+00 | 2.92E+00 |
| 1024508 | 431071.9 | 109472.2 | 9972782 | 678652.5 | 1024508 | 10.96567 | 72.39202 | 7.19E+01 |
| 7728031 | 8205157 | 109948.4 | 522620 | 565852 | 7728031 | 77.67705 | 27.8972 | 10.43392 |
| 553644.5 | 650758.9 | 110977.5 | 4459698 | 4600728 | 553644.5 | 7.419218 | 12.14208 | 17.20512 |
| 4980750 | 3172849 | 253397.9 | 17475169 | 6536423 | 3172849 | 10.56186 | 16.70932 | 32.50955 |
| 300101.1 | 299243.1 | 215624.3 | 5543677 | 1511800 | 300101.1 | 58.77163 | 76.14806 | 8.83E+01 |
| 7848170 | 12836250 | 227498.2 | 1349231 | 3446980 | 10507907 | 87.21502 | 69.83529 | 24.0047 |
| 1040637 | 403341.4 | 88340.06 | 5956619 | 558545.2 | 403341.4 | 9.449002 | 76.38038 | 67.52509 |
| 2459187 | 2838258 | 89294.06 | 6023107 | 3797703 | 2459187 | 8.019966 | 7.56273 | 16.39219 |
| 8263273 | 6299993 | 131413.1 | 6643938 | 11895482 | 6392817 | 93.22365 | 5.127456 | 15.85712 |
| 29954317 | 32364573 | 148083.9 | 1692659 | 789218.9 | 29954317 | 60.63737 | 60.10504 | 7.02544 |
| 243159.1 | 231517 | 77987.59 | 4565694 | 7204083 | 243159.1 | 13.32038 | 1.73E+00 | 7.16E+00 |
| 2249160 | 3458322 | 158764.6 | 6730065 | 3706560 | 2249160 | 2.524402 | 5.99E+01 | 2.93E+01 |
| 7826124 | 7791025 | 99599.27 | 6330324 | 5866237 | 7791025 | 17.14621 | 11.32185 | 9.097099 |
| 184919.8 | 180017.4 | 110537.1 | 7985514 | 195606.6 | 180017.4 | 7.330147 | 9.25E-01 | 8.45E+00 |
| 1797121 | 1416051 | 96451.14 | 7395320 | 6269109 | 1416051 | 7.692432 | 1.39E+01 | 2.24E+01 |
| 5633291 | 2852673 | 143434.2 | 12015847 | 8858715 | 4352487 | 0.933838 | 5.03E+00 | 3.25E+01 |
| 987932.8 | 756452.5 | 171634.2 | 2529322 | 3670175 | 798071.3 | 6.426556 | 8.31E+00 | 1.46E+01 |
| 1584311 | 1212104 | 94341.01 | 5369153 | 7470894 | 1451944 | 51.77678 | 16.02842 | 13.32316 |
| 2942939 | 11588628 | 1145958 | 15766391 | 5197107 | 8211444 | 35.1384 | 61.68411 | 57.47506 |
| 335535.1 | 437108.1 | 65376.25 | 3407284 | 318069 | 437108.1 | 15.11345 | 4.05E+01 | 1.95E+01 |
| 1692311 | 1907563 | 83324.09 | 6174340 | 4783084 | 1692311 | 33.68392 | 17.63203 | 19.83514 |
| 1591639 | 1269829 | 89383.66 | 6826860 | 6841448 | 1591639 | 1.661707 | 20.61144 | 1.32E+01 |
| 3110011 | 2164785 | 119490.3 | 8540405 | 8605301 | 2903644 | 7.824204 | 4.83E+00 | 1.82E+01 |
| 10334170 | 11040311 | 163933.4 | 2733573 | 1643915 | 10334170 | 27.25436 | 5.54E+00 | 7.45E+00 |
| 821252.8 | 681973.8 | 85473.02 | 8408302 | 1807937 | 821252.8 | 12.90481 | 8.34E+00 | 1.09E+01 |
| 1860390 | 5186070 | 87737.01 | 3252170 | 7630323 | 3183096 | 19.52755 | 5.47E+00 | 4.91E+01 |
| 907611.1 | 916359.7 | 97435.93 | 6441022 | 5023034 | 907611.1 | 8.576683 | 4.37E+00 | 2.25E+01 |
| 1311147 | 1256851 | 54712.65 | 5486119 | 2239002 | 1256851 | 23.06634 | 24.09843 | 38.40108 |
| 7076301 | 5462754 | 406125.2 | 22809031 | 20880196 | 5462754 | 10.68837 | 16.6616 | 51.75577 |
| 169165 | 552000.8 | 84130.6 | 8388314 | 295976.7 | 334899.7 | 18.96972 | 66.66287 | 54.53963 |
| 5557308 | 6194365 | 130266.8 | 6183521 | 8654225 | 5557308 | 4.99727 | 26.52044 | 1.95E+01 |
| 4648535 | 3133181 | 176998.8 | 6846117 | 8054907 | 3133181 | 8.324236 | 30.77934 | 32.47779 |

| | | | | | | | | |
|---|---|---|---|---|---|---|---|---|
| 4130492 | 1938670 | 116814.1 | 7374224 | 5303126 | 3759208 | 16.07863 | 6.441963 | 35.80608 |
| 2766407 | 2963174 | 31788.78 | 215759.1 | 100020.1 | 2766407 | 4.012562 | 70.67519 | 8.435373 |
| 1035928 | 890064.3 | 62331.36 | 7433943 | 1283287 | 890064.3 | 15.3981 | 32.05909 | 4.79E+01 |
| 99908.99 | 162150.2 | 85346.31 | 7360980 | 3111104 | 162150.2 | 3.576947 | 3.15E+01 | 3.58E+01 |
| 5386263 | 5928318 | 79688.13 | 332134.1 | 230984.4 | 5386263 | 65.99344 | 25.25116 | 14.08714 |
| 7556551 | 5377290 | 132819.8 | 10183737 | 10459421 | 7556551 | 13.17966 | 20.59937 | 49.84154 |
| 51290802 | 44322348 | 1188304 | 55230148 | 37711654 | 44322348 | 6.594446 | 2.50E+01 | 9.81E+00 |
| 288337.7 | 2075436 | 95051.79 | 12552848 | 549845.9 | 288337.7 | 3.556777 | 6.99E+01 | 118.4954 |
| 110066.2 | 113026.8 | 93565.58 | 7240349 | 1048588 | 110066.2 | 3.435642 | 20.90918 | 1.934243 |
| 4242021 | 9522812 | 158090.8 | 8888228 | 4513381 | 4242021 | 14.30374 | 6.11E+01 | 54.48641 |
| 172879.7 | 258505.2 | 81686.15 | 8122908 | 488230.1 | 258505.2 | 6.890213 | 6.11E+01 | 4.55E+01 |
| 1391408 | 1825293 | 90185.93 | 4864305 | 6461148 | 1391408 | 19.03867 | 2.16E+01 | 5.27E+01 |
| 7718435 | 8309194 | 107493.7 | 773667.5 | 557689.2 | 7718435 | 18.45861 | 28.40527 | 1.66E+01 |
| 657496.1 | 201952.1 | 52138.82 | 4076851 | 2334083 | 498411.4 | 2.39083 | 5.60E+00 | 5.11E+01 |
| 4038504 | 4290961 | 82211.28 | 1830913 | 2237942 | 4038504 | 33.72246 | 9.033878 | 9.690967 |
| 5753417 | 5593806 | 157947.9 | 14039641 | 8887875 | 5593806 | 2.088371 | 1.64E+01 | 1.58E+01 |
| 2959984 | 2339791 | 70212.59 | 3989890 | 3044780 | 2666818 | 11.46348 | 3.75E+01 | 1.17E+01 |
| 149742.8 | 298812.7 | 94753.63 | 6334545 | 534787.2 | 298812.7 | 3.837117 | 8.142215 | 4.92E+01 |
| 1754179 | 1627442 | 109583.6 | 4740890 | 3754919 | 1627442 | 1.973066 | 2.67E+00 | 5.57E+00 |
| 1067652 | 2023662 | 281352.6 | 19681155 | 6617952 | 2023662 | 22.87786 | 4.22E+01 | 4.97E+01 |
| 2134690 | 1481298 | 80097.39 | 4504189 | 4090937 | 1874798 | 27.37069 | 20.93466 | 1.80E+01 |
| 4320907 | 1055507 | 127561.4 | 11353797 | 3073176 | 1509612 | 5.950305 | 2.54E+01 | 7.71E+01 |
| 3110706 | 3866000 | 81258.69 | 3424672 | 3880098 | 3208465 | 5.878881 | 50.82525 | 1.21E+01 |
| 86873.3 | 135009.3 | 52068.48 | 3431480 | 1675954 | 135009.3 | 10.17265 | 12.99715 | 34.29237 |
| 5891888 | 7535555 | 73100.48 | 1007873 | 445253.1 | 6621698 | 106.1769 | 7.99E+01 | 1.23E+01 |
| 188025.7 | 119953.2 | 71128.8 | 4641956 | 3627382 | 119953.2 | 11.41177 | 15.18788 | 34.95577 |
| 1100654 | 2558026 | 422786.1 | 23291995 | 2525263 | 2558026 | 9.017953 | 46.76248 | 51.57575 |
| 10777915 | 10410401 | 118280.6 | 3898038 | 3257486 | 10410401 | 19.10275 | 12.81053 | 37.7538 |
| 739896.5 | 8764013 | 96333.44 | 6798524 | 582107.3 | 739896.5 | 8.202329 | 34.82388 | 142.6813 |
| 75880.92 | 82234.91 | 44137.5 | 2807251 | 1179216 | 75880.92 | 9.256548 | 9.66E+00 | 10.67545 |
| 192159.7 | 96010.13 | 92268.78 | 7357554 | 180905.4 | 97172.22 | 1.043899 | 33.35951 | 4.30E+01 |
| 32211334 | 28593329 | 4500970 | 51465123 | 26342244 | 28593329 | 4.538474 | 41.97157 | 17.94976 |
| 9681120 | 8612333 | 2167188 | 12930883 | 12091832 | 9681120 | 62.26716 | 5.09E+01 | 4.16E+01 |
| 7238616 | 8503495 | 565256.9 | 10544960 | 11747735 | 7238616 | 13.62756 | 7.97E+00 | 15.27652 |
| 6558073 | 6419026 | 1686149 | 20558171 | 8498359 | 6419026 | 25.55836 | 8.04E-01 | 8.90E+00 |
| 25760101 | 35008995 | 2887847 | 12787276 | 30240983 | 27284395 | 77.30773 | 1.595013 | 16.89455 |
| 45366292 | 34892569 | 3137017 | 39938428 | 23520840 | 34892569 | 53.61666 | 29.31669 | 23.03003 |
| 6075566 | 6069189 | 2348577 | 50658730 | 8671937 | 6069189 | 4.702482 | 1.40E+01 | 8.55E+00 |
| 6984591 | 8891307 | 849550.1 | 7165769 | 6286239 | 6984591 | 43.90051 | 47.82249 | 24.28635 |
| 44648902 | 40749359 | 436100.7 | 2678833 | 2883423 | 40749359 | 3.266947 | 7.892296 | 13.53498 |
| 36390613 | 37573970 | 393251.9 | 33419886 | 20854947 | 36390613 | 11.29671 | 58.50902 | 65.75234 |
| 36313030 | 36741550 | 405987 | 31142334 | 22023364 | 36313030 | 67.55614 | 1.229675 | 1.67E+01 |
| 4764987 | 5049609 | 54658.57 | 85117.95 | 79804.33 | 4764987 | 53.14637 | 4.53E+01 | 2.00E+01 |
| 13683237 | 9038108 | 90686.97 | 444267.6 | 279942 | 9919092 | 50.81424 | 42.66802 | 2.27E+01 |
| 3770797 | 2634994 | 7094017 | 1.57E+09 | 18388213 | 3770797 | 2.784294 | 54.71494 | 43.80585 |
| 2835254 | 378150.3 | 206305.4 | 2.04E+08 | 5102164 | 2703747 | 2.296407 | 13.57638 | 70.07823 |
| 16809561 | 16912793 | 127798.4 | 24474038 | 22728487 | 16809561 | 2.345408 | 5.187467 | 9.531378 |
| 26839302 | 35899564 | 1310471 | 2259010 | 2936112 | 26839302 | 21.57805 | 27.41541 | 18.67505 |
| 529820.8 | 641007.8 | 110198.2 | 15679315 | 1240751 | 566144 | 7.46459 | 22.99462 | 9.79E+00 |
| 10336334 | 5839698 | 710420.1 | 37761250 | 13539675 | 10336334 | 36.49997 | 1.18E+02 | 4.78E+01 |
| 5131984 | 5732756 | 239404.6 | 9255559 | 10202096 | 5131984 | 2.236865 | 5.509892 | 18.81242 |

| | | | | | | | | |
|---|---|---|---|---|---|---|---|---|
| 6887400 | 6305136 | 241344.3 | 19102159 | 12847396 | 6887400 | 6.0332 | 7.35E+00 | 5.686002 |
| 282232.6 | 296779 | 713615.9 | 4.11E+08 | 1364579 | 296779 | 2.728456 | 4.94E+01 | 7.16E+01 |
| 34629727 | 46246130 | 3898350 | 74852036 | 12281221 | 34629727 | 3.736509 | 74.35561 | 1.81E+01 |
| 11032702 | 11841159 | 474329.1 | 13667146 | 20297281 | 11032702 | 53.74778 | 0.526188 | 1.92E+01 |
| 3876036 | 3786614 | 91359.46 | 8405267 | 9577284 | 3786614 | 5.720975 | 7.22E+00 | 1.12E+01 |
| 2843022 | 4731775 | 135934.1 | 9640715 | 2479054 | 4089120 | 6.623666 | 60.54742 | 2.47E+01 |
| 16339966 | 20192909 | 383043.4 | 25324247 | 21831687 | 16339966 | 7.23524 | 1.18E+01 | 1.88E+01 |
| 207937.1 | 200495.4 | 368361 | 258763.6 | 21698373 | 200495.4 | 1.78344 | 1.34E+01 | 8.30E+00 |
| 470505 | 270228.1 | 124724.8 | 16637649 | 288487 | 270228.1 | 1.25328 | 34.81499 | 35.33097 |
| 596484.9 | 347664.4 | 87554.67 | 805010.8 | 4616260 | 584121.9 | 29.24514 | 4.360097 | 2.75E+01 |
| 1324764 | 460037.7 | 73307.45 | 7420258 | 2778727 | 460037.7 | 7.192872 | 8.46E+01 | 8.27E+01 |
| 402948.5 | 7909132 | 4986117 | 59316225 | 438793.1 | 2740109 | 4.813281 | 28.07486 | 1.04E+02 |
| 128730.1 | 128072.4 | 66935.02 | 4141967 | 633712.2 | 128072.4 | 2.806343 | 10.97323 | 4.676159 |
| 316171.1 | 562867 | 321136.9 | 25670424 | 19921082 | 316171.1 | 4.241803 | 5.38434 | 5.02E+01 |
| 35275864 | 4426322 | 176034.5 | 31539708 | 23445831 | 29114668 | 21.23331 | 51.27058 | 71.17017 |
| 8749421 | 8088766 | 249127 | 12786501 | 7626704 | 8088766 | 10.8207 | 42.05363 | 11.12173 |
| 6795402 | 10055015 | 407372.5 | 18490313 | 21182300 | 8846772 | 10.99711 | 5.041992 | 1.92E+01 |
| 12048382 | 12022705 | 166175.7 | 20075887 | 18417232 | 12022705 | 1.414802 | 4.52E+00 | 1.08E+01 |
| 10692967 | 13956427 | 142254.4 | 21792495 | 20422516 | 12604746 | 3.434431 | 1.88E+01 | 1.32E+01 |
| 17808606 | 20339528 | 95017.07 | 18630760 | 18874613 | 17808606 | 2.457751 | 1.11E+01 | 1.54E+01 |
| 15728221 | 19230658 | 88843.14 | 16334154 | 18395237 | 15728221 | 3.996761 | 13.53624 | 17.99 |
| 3388863 | 3497868 | 151334.5 | 19622590 | 2751587 | 3388863 | 3.382706 | 7.34E+01 | 1.33E+01 |
| 7923588 | 5440070 | 177529.1 | 17463344 | 13327950 | 5440070 | 19.53718 | 44.36516 | 26.35897 |
| 1400424 | 2385947 | 171147.5 | 10230642 | 3002726 | 1400424 | 8.242297 | 7.26E+01 | 6.60E+01 |
| 9123308 | 7985142 | 187626.3 | 23442282 | 8479153 | 7985142 | 1.200547 | 1.78E+01 | 2.90E+01 |
| 17883042 | 17956987 | 90729.95 | 33699678 | 26297221 | 17883042 | 4.208248 | 12.34717 | 12.17843 |
| 1949629 | 2037356 | 64178.15 | 10780934 | 2988413 | 1949629 | 4.673651 | 4.98E+00 | 1.68E+01 |
| 12974974 | 11003906 | 513183.5 | 30887923 | 25783976 | 11003906 | 5.935519 | 8.66E+00 | 1.15E+01 |
| 859574.3 | 229402.4 | 104239.2 | 13608543 | 1234261 | 229402.4 | 10.67963 | 5.87E+01 | 8.72E+01 |
| 9749998 | 10087023 | 159735.6 | 2065526 | 2732707 | 9749998 | 48.16591 | 6.263642 | 4.45E+00 |
| 14820813 | 12756840 | 125920.6 | 245519.1 | 216181.8 | 12756840 | 5.969334 | 1.81E-01 | 1.36E+01 |
| 24794638 | 26491297 | 199016.5 | 442879.2 | 678218.5 | 24794638 | 6.362136 | 2.29E+01 | 10.99273 |
| 17288055 | 15179725 | 266748.8 | 1141960 | 1173548 | 15179725 | 46.35697 | 8.36779 | 12.70056 |
| 247180 | 237799.7 | 158948.6 | 9063410 | 918678.5 | 247180 | 8.582774 | 4.58E+01 | 31.69504 |
| 6731284 | 8441964 | 146759 | 40401027 | 13070933 | 8441964 | 6.420955 | 1.22E+01 | 1.30E+01 |
| 14382304 | 25357123 | 161440.2 | 37448952 | 15157723 | 14382304 | 46.54523 | 6.89E+01 | 5.74E+01 |
| 13771635 | 14450303 | 88419.05 | 17229885 | 30918073 | 13771635 | 3.106595 | 6.71E+00 | 10.36675 |
| 6132194 | 5189901 | 1320858 | 8327887 | 14343342 | 5189901 | 9.313176 | 7.34E+00 | 3.72E+01 |
| 4701580 | 4422765 | 155178.5 | 19706114 | 16979730 | 4422765 | 16.9319 | 2.903994 | 9.60945 |
| 4365994 | 609759.8 | 93670.45 | 17130482 | 427322.8 | 2422351 | 0.780738 | 5.24E+01 | 7.62E+01 |
| 5921774 | 37071853 | 3998799 | 52779736 | 43366633 | 5921774 | 25.25714 | 7.12E+01 | 1.14E+02 |
| 4509017 | 1128259 | 141188.8 | 3883283 | 6843867 | 2872089 | 55.29168 | 24.42842 | 5.96E+01 |
| 4967814 | 4639605 | 19855.87 | 438576.9 | 66105.27 | 4639605 | 60.27229 | 8.45E+00 | 8.20E+00 |
| 163817.2 | 155354.2 | 102142.8 | 11042575 | 156994.9 | 155354.2 | 2.814051 | 2.260582 | 7.84E+00 |
| 1.69E+08 | 1.76E+08 | 13188587 | 1.06E+08 | 1.48E+08 | 1.69E+08 | 5.801751 | 2.69E+00 | 1.34E+01 |
| 1.42E+08 | 1.38E+08 | 10584200 | 1.14E+08 | 1.16E+08 | 1.38E+08 | 3.208253 | 7.71292 | 1.11E+01 |
| 44369123 | 42031651 | 976791.4 | 3744118 | 6563745 | 42031651 | 31.93495 | 19.52195 | 6.11E+00 |
| 5672134 | 4788251 | 175897.7 | 7871170 | 15581300 | 4788251 | 11.21346 | 4.00579 | 11.08156 |
| 8471646 | 8686195 | 91218.62 | 354857.6 | 297100.9 | 8471646 | 86.63835 | 76.66588 | 16.24097 |
| 825792.3 | 318074.2 | 89644.05 | 37063462 | 958743.3 | 825792.3 | 4.82713 | 4.40E+01 | 8.31E+01 |
| 1.14E+08 | 1.3E+08 | 1704092 | 3.13E+08 | 2.72E+08 | 1.14E+08 | 3.294159 | 5.28E+00 | 1.55E+01 |

| | | | | | | | | |
|---|---|---|---|---|---|---|---|---|
| 4224728 | 3177229 | 89381.74 | 9014022 | 11295949 | 3177229 | 2.146698 | 19.06027 | 2.05E+01 |
| 3170549 | 2669521 | 83678.49 | 8237524 | 8778254 | 2669521 | 7.119015 | 29.32419 | 2.14E+01 |
| 7352387 | 8966728 | 102580.4 | 3295415 | 2960723 | 7352387 | 11.1982 | 4.19E+00 | 19.93239 |
| 79751.05 | 106459.4 | 62742.02 | 7497120 | 1109928 | 106459.4 | 11.14478 | 11.53507 | 18.48354 |
| 8302697 | 9263218 | 791362 | 21638234 | 16074548 | 8302697 | 1.118021 | 6.810478 | 12.87831 |
| 5080941 | 575839.1 | 68457.56 | 8096976 | 3080652 | 4124488 | 11.53013 | 2.94E+01 | 7.28E+01 |
| 83070194 | 83396007 | 123804.6 | 1.48E+08 | 1.56E+08 | 83070194 | 1.032553 | 9.508072 | 32.8227 |
| 58964.74 | 92344.11 | 2076461 | 2705276 | 21038436 | 92344.11 | 13.58707 | 8.268133 | 69.81591 |
| 5631400 | 5391763 | 2791793 | 36368924 | 29673417 | 5391763 | 8.891396 | 15.63348 | 8.411076 |
| 2689571 | 2643157 | 108444.6 | 5693021 | 4157335 | 2643157 | 6.202715 | 1.16E+01 | 1.10E+01 |
| 57599154 | 54589047 | 6195229 | 1.33E+08 | 82175358 | 54589047 | 7.509887 | 43.04154 | 1.15E+01 |
| 41982991 | 43052455 | 3482938 | 85953468 | 61372719 | 41982991 | 9.433457 | 2.170437 | 1.17E+01 |
| 39225058 | 30187905 | 10066040 | 1.06E+08 | 70816363 | 32827151 | 7.247245 | 4.31E+00 | 13.63562 |
| 6163191 | 7546818 | 649768.5 | 12309530 | 9826613 | 6163191 | 10.8668 | 3.890739 | 15.63431 |
| 7223526 | 6354244 | 797150.2 | 9874860 | 9096403 | 6354244 | 10.72856 | 3.65E+00 | 2.13E+01 |
| 8548063 | 8460328 | 626369.9 | 13388830 | 12772287 | 8460328 | 2.577016 | 4.49E+00 | 7.01E+00 |
| 6561926 | 6219713 | 94073.37 | 6944099 | 6590019 | 6219713 | 3.257085 | 1.825087 | 1.26E+01 |
| 9418052 | 9028873 | 391082.5 | 11330467 | 11480974 | 9028873 | 8.46501 | 6.294572 | 1.71E+01 |
| 14878861 | 6905324 | 388893.4 | 17830470 | 10642954 | 12379391 | 58.80233 | 87.85119 | 35.81178 |
| 255305.3 | 273697.4 | 384377.8 | 370951.4 | 9802574 | 273697.4 | 39.66147 | 11.36774 | 9.885808 |
| 232837.5 | 2396917 | 143876.3 | 383004.7 | 9341668 | 691016.9 | 23.75385 | 13.89804 | 103.0255 |
| 8296275 | 8456199 | 238219.4 | 41479031 | 17392093 | 8296275 | 14.21923 | 9.85E+00 | 7.64E+00 |
| 430927.8 | 664926.5 | 81880.39 | 4669193 | 3129410 | 520181.8 | 39.9261 | 2.76E+01 | 2.19E+01 |
| 9100248 | 15103477 | 231079.6 | 12407604 | 12388880 | 9100248 | 10.43082 | 2.780691 | 35.20163 |
| 39763877 | 43034766 | 462582.9 | 1.76E+08 | 59642542 | 39763877 | 3.178784 | 2.27192 | 13.71079 |
| 2516192 | 394153.4 | 121023.5 | 6932411 | 1871111 | 394153.4 | 7.729373 | 6.78E+01 | 1.21E+02 |
| 23506139 | 23430784 | 94601.74 | 6988933 | 12689175 | 23430784 | 14.87901 | 6.63032 | 7.33E+00 |
| 1784534 | 959157.7 | 156639.2 | 10736876 | 5804539 | 959157.7 | 0.790439 | 11.88208 | 5.14E+01 |
| 5949861 | 5808476 | 137158.9 | 14950594 | 12073970 | 5808476 | 3.390397 | 5.83714 | 13.14229 |
| 1109626 | 827870.2 | 286905.7 | 9716282 | 690417 | 827870.2 | 3.51934 | 50.78643 | 2.06E+01 |
| 49007000 | 51278762 | 1012270 | 2.05E+08 | 1.47E+08 | 49007000 | 2.93671 | 0.553388 | 6.89874 |
| 768269.1 | 629955.1 | 270668.8 | 2437027 | 3436254 | 629955.1 | 48.03026 | 29.95009 | 12.3578 |
| 225428.1 | 136701 | 74375.38 | 1972133 | 7516874 | 182238 | 73.50101 | 38.30328 | 2.45E+01 |
| 3617832 | 3657965 | 126244.8 | 18918394 | 11531758 | 3617832 | 4.533569 | 4.34176 | 17.55563 |
| 7505234 | 7630216 | 99638.47 | 14376761 | 16496801 | 7505234 | 18.20048 | 11.32757 | 12.99684 |
| 220754.2 | 281671 | 148420.5 | 73970517 | 769417.6 | 281671 | 2.039638 | 16.80783 | 24.60016 |
| 44369123 | 42031651 | 1073625 | 3744118 | 6563745 | 42031651 | 32.9294 | 2.49E+01 | 6.11E+00 |
| 8445806 | 9045130 | 258018.7 | 43512105 | 25949950 | 8445806 | 0.244071 | 7.664996 | 15.22292 |
| 4163071 | 3046716 | 1317244 | 17866487 | 11692085 | 3046716 | 7.168427 | 3.488184 | 23.29607 |
| 556442.5 | 165631.4 | 100269.3 | 13187129 | 2773287 | 218180.6 | 7.851475 | 26.77475 | 67.67284 |
| 457955.2 | 261634.1 | 71505.58 | 4787086 | 151640.5 | 261634.1 | 36.29052 | 5.81E+01 | 5.51E+01 |
| 26016436 | 27037215 | 318515.4 | 9457245 | 11142178 | 26016436 | 5.847546 | 4.741618 | 10.81736 |
| 609610.2 | 813631.9 | 87969.18 | 9519999 | 7937632 | 813631.9 | 2.853699 | 7.242457 | 16.82253 |
| 109135.1 | 106250.9 | 98598.23 | 13727270 | 14121269 | 109135.1 | 4.563238 | 13.1489 | 2.92E+01 |
| 8819723 | 7808108 | 97490.31 | 8737321 | 9793547 | 7808108 | 11.94092 | 1.547072 | 8.183382 |
| 2897406 | 2528087 | 100441.7 | 6084842 | 4510708 | 2640913 | 3.493393 | 22.36545 | 7.038808 |
| 7352205 | 7760688 | 103842.3 | 15562035 | 28000744 | 7352205 | 6.820634 | 1.54E+00 | 3.77E+01 |
| 169846.8 | 166592 | 647186.8 | 2934835 | 32180557 | 166592 | 15.57543 | 1.39E+01 | 4.91E+00 |
| 1443973 | 2662630 | 130196.6 | 9432976 | 10200539 | 1503905 | 3.486921 | 2.109594 | 36.7317 |
| 11014490 | 11207039 | 274946.1 | 28444953 | 29718698 | 11014490 | 7.589819 | 4.99E+00 | 8.474453 |
| 10240039 | 8715143 | 106768.5 | 4805682 | 1696281 | 10240039 | 10.82544 | 5.95E+01 | 9.45E+00 |

| | | | | | | | | |
|---|---|---|---|---|---|---|---|---|
| 1949629 | 2037356 | 65398.08 | 10780934 | 2988413 | 1949629 | 4.673651 | 4.980255 | 16.83904 |
| 2451426 | 3432064 | 54396.07 | 11573607 | 12037164 | 3432064 | 0.50166 | 1.70E+01 | 1.84E+01 |
| 15498844 | 15208956 | 1047757 | 32087917 | 23842812 | 15208956 | 6.277011 | 1.08E+01 | 7.47E+00 |
| 1822604 | 1861654 | 108050.5 | 79964728 | 3188315 | 1822604 | 4.39211 | 6.843007 | 23.84941 |
| 2304805 | 2202537 | 88481.77 | 8697664 | 9739832 | 2202537 | 0.273508 | 1.50E+01 | 1.87E+01 |
| 12967560 | 14012751 | 178080.9 | 12307519 | 15003954 | 12967560 | 37.75828 | 67.31119 | 2.45E+01 |
| 6484008 | 8767784 | 208496.6 | 23173708 | 15423073 | 8767784 | 31.71737 | 35.62957 | 4.72E+01 |
| 1145852 | 6331648 | 76070.57 | 10931719 | 5464914 | 5531593 | 1.776402 | 31.73993 | 64.38277 |
| 12047678 | 11533973 | 91668.61 | 11872948 | 10231028 | 11533973 | 7.509369 | 1.81E+01 | 1.27E+01 |
| 10810336 | 10237051 | 720414 | 39635210 | 8139849 | 10237051 | 3.257755 | 2.638744 | 9.98E+00 |
| 6272819 | 5712214 | 101468 | 1833700 | 1443524 | 5712214 | 31.83606 | 38.34231 | 17.94498 |
| 8767397 | 6540598 | 762808.3 | 27466700 | 11890014 | 6540598 | 5.975714 | 9.022491 | 25.44435 |
| 5324013 | 3901225 | 430682.1 | 18280599 | 11606595 | 5324013 | 64.21536 | 1.79E+01 | 1.76E+01 |
| 6450188 | 7480590 | 95326.9 | 536442.5 | 1011148 | 6810737 | 28.33116 | 24.86149 | 7.56E+00 |
| 727711.1 | 2993532 | 117660.6 | 11268656 | 6316233 | 2083603 | 0.994041 | 20.82827 | 58.92671 |
| 6963116 | 7227396 | 131999.1 | 5378202 | 6567697 | 6963116 | 44.35169 | 4.537325 | 2.404079 |
| 1.54E+08 | 1.39E+08 | 352045.2 | 6140501 | 5367941 | 1.39E+08 | 1.914932 | 18.16163 | 1.46E+01 |
| 3525889 | 3456891 | 74525.78 | 7979884 | 9827273 | 3456891 | 4.978711 | 8.52E+00 | 9.88E+00 |
| 1240747 | 1556530 | 73076.31 | 9471409 | 10165805 | 1301629 | 5.67097 | 8.61E+00 | 1.23E+01 |
| 2319991 | 1591164 | 131762.9 | 6028462 | 4002590 | 1647112 | 17.78389 | 9.425458 | 21.89189 |
| 22785096 | 23209976 | 664569.8 | 9116596 | 21892879 | 22785096 | 16.22482 | 5.13E+01 | 1.16E+01 |
| 7133819 | 10407191 | 113943 | 5973249 | 3942827 | 7133819 | 9.396828 | 15.95028 | 28.21158 |
| 22574641 | 7254785 | 99939.25 | 20786877 | 10958119 | 14148330 | 21.63973 | 6.808907 | 5.23E+01 |
| 12480929 | 6626870 | 129698.1 | 8362993 | 11639338 | 6626870 | 36.19934 | 37.21192 | 47.77226 |
| 6889051 | 8236306 | 127886.2 | 12475211 | 12819650 | 7809905 | 1.921006 | 11.6338 | 9.01E+00 |
| 3520664 | 3157903 | 252686.3 | 3539920 | 6903722 | 3295686 | 15.55924 | 5.798677 | 5.507745 |
| 23042363 | 23621582 | 161538.9 | 22495267 | 28202657 | 23042363 | 8.172015 | 0.805019 | 20.67758 |
| 5644388 | 5433583 | 60832.04 | 198485.2 | 381987.4 | 5433583 | 39.52693 | 43.4956 | 2.52E+01 |
| 3995382 | 4049662 | 125334.6 | 6400680 | 13813415 | 3995382 | 4.803583 | 2.227997 | 7.393398 |
| 775739.6 | 672167.3 | 93507.34 | 19484171 | 887346.9 | 775739.6 | 4.806323 | 9.46E+01 | 3.01E+01 |
| 6397144 | 7088381 | 148044.9 | 12013234 | 9436840 | 6481208 | 1.262237 | 1.32E+01 | 5.67E+00 |
| 37279064 | 38706375 | 149120.5 | 2857649 | 696750.4 | 37279064 | 70.70973 | 4.38E+01 | 8.94E+00 |
| 19938672 | 19543882 | 675389.1 | 6863584 | 24248049 | 19543882 | 5.219573 | 3.14E+00 | 9.37E+00 |
| 14929868 | 16639398 | 7328015 | 99083143 | 53617349 | 14929868 | 27.98882 | 25.47479 | 1.51E+01 |
| 28579027 | 27895745 | 3531017 | 45937539 | 63552349 | 27895745 | 36.32035 | 5.25E+01 | 1.42E+01 |
| 18275949 | 22706662 | 3871129 | 43211107 | 30119409 | 18623674 | 12.23734 | 1.97E+01 | 1.24E+01 |
| 15007519 | 12133086 | 1513968 | 17573039 | 12927084 | 12133086 | 2.281237 | 5.32E+00 | 1.40E+01 |
| 161592.7 | 161567.8 | 87096.52 | 10443126 | 1299659 | 161567.8 | 13.16822 | 66.12092 | 6.68E+00 |
| 2236326 | 4262224 | 132104.8 | 12608406 | 14678873 | 2491838 | 5.9218 | 2.946029 | 36.81646 |
| 19387947 | 19597821 | 172765 | 18309941 | 15755479 | 19387947 | 28.72384 | 50.1239 | 7.10E+01 |
| 2088008 | 2182184 | 138735.5 | 23007251 | 2847395 | 2088008 | 4.864763 | 60.29483 | 60.56327 |
| 346390.2 | 346395 | 213259.1 | 10890095 | 419818.5 | 346395 | 6.952789 | 15.25429 | 4.05E+00 |
| 1659512 | 2066533 | 138537.8 | 21657419 | 1576621 | 1666585 | 4.786089 | 8.98664 | 12.96094 |
| 169276.4 | 170048.2 | 112034.1 | 8597351 | 176544.6 | 169276.4 | 13.7638 | 4.434653 | 5.438828 |
| 10497050 | 19896859 | 371156.9 | 12546166 | 13674132 | 14074089 | 16.32615 | 18.11901 | 32.00777 |
| 3847966 | 3577560 | 96086.43 | 17564734 | 12524437 | 3577560 | 5.3267 | 2.77E+00 | 9.73E+00 |
| 2523388 | 2604808 | 56819.84 | 12025082 | 7828235 | 2523388 | 1.290666 | 3.738781 | 3.732732 |
| 1837137 | 990857.2 | 270328.5 | 19366837 | 13532637 | 1445324 | 3.010877 | 7.141224 | 29.73285 |
| 138065.7 | 211095.2 | 146992.9 | 7356448 | 378882 | 211095.2 | 1.921805 | 19.40327 | 8.07E+01 |
| 8827806 | 8772926 | 80869.4 | 15901583 | 16773484 | 8827806 | 4.917761 | 3.23E+01 | 8.31E+00 |
| 8189196 | 7731705 | 246137.7 | 8526994 | 23950215 | 7731705 | 28.97762 | 6.830256 | 9.205209 |

| | | | | | | | | |
|---|---|---|---|---|---|---|---|---|
| 10304645 | 9310377 | 95583.71 | 25604357 | 26681205 | 9310377 | 2.218212 | 2.01E+01 | 1.48E+01 |
| 1631571 | 1781639 | 224522.8 | 44025718 | 3200731 | 1631571 | 0.479557 | 38.99408 | 15.91133 |
| 5915847 | 5165615 | 492075.1 | 5382449 | 5707835 | 5165615 | 6.851726 | 3.94E+00 | 1.03E+01 |
| 5913954 | 4254743 | 2344831 | 30517901 | 27128666 | 4643937 | 5.331939 | 16.6023 | 17.57347 |
| 18333576 | 18227074 | 150756.9 | 44481566 | 39226939 | 18227074 | 3.010587 | 2.977725 | 10.61261 |
| 2.97E+08 | 2.7E+08 | 2995966 | 19946004 | 20710147 | 2.7E+08 | 18.77192 | 16.17369 | 10.78933 |
| 28444256 | 31597492 | 130821.7 | 65785826 | 71111417 | 28444256 | 3.401852 | 8.754362 | 17.14158 |
| 199944 | 312200.1 | 125606.4 | 6690138 | 240574.8 | 206425.8 | 4.074487 | 5.119685 | 26.31199 |
| 7528849 | 6595658 | 134113 | 7623285 | 11922655 | 6888864 | 5.322846 | 35.83914 | 6.812996 |
| 56345719 | 56976929 | 408376.1 | 2623527 | 2041639 | 56345719 | 0.945781 | 33.09218 | 15.46809 |
| 35962548 | 38793014 | 190560.3 | 4700421 | 1838158 | 35962548 | 38.39911 | 66.37992 | 1.29E+01 |
| 617040.6 | 822680.8 | 68229.64 | 8593054 | 8561156 | 617040.6 | 1.21557 | 5.545822 | 44.03573 |
| 116627.2 | 117050.1 | 85467.83 | 5092003 | 122323.2 | 116627.2 | 2.963984 | 2.291297 | 4.98E+00 |
| 2781660 | 3267391 | 127008.1 | 8979421 | 5363668 | 2848921 | 3.096031 | 5.997368 | 8.87E+00 |
| 3141716 | 5420732 | 1126336 | 14040141 | 8435579 | 3141716 | 7.275786 | 3.397185 | 35.38427 |
| 12177603 | 12382103 | 180075.8 | 29591105 | 24989418 | 12177603 | 1.619236 | 4.421308 | 17.19272 |
| 11245655 | 12454540 | 165790.2 | 28136482 | 25151915 | 11245655 | 7.952385 | 4.56E+00 | 1.63E+01 |
| 16631945 | 17895348 | 2664291 | 14451156 | 33098617 | 17890873 | 1.287345 | 21.75854 | 4.17E+00 |
| 20638027 | 21113443 | 140372.2 | 14668178 | 4066158 | 20638027 | 45.82813 | 10.46846 | 8.42963 |
| 6875233 | 6755755 | 42203.65 | 259788.4 | 115261.1 | 6755755 | 41.46767 | 6.15E+01 | 1.79E+01 |
| 10788045 | 10497070 | 91966.55 | 4383658 | 3579418 | 10497070 | 8.019856 | 6.814174 | 8.166987 |
| 12493274 | 23777106 | 51883.4 | 255351.3 | 987659.6 | 12493274 | 44.94775 | 4.092122 | 45.47622 |
| 1.49E+08 | 1.5E+08 | 963821.1 | 1.98E+08 | 1.73E+08 | 1.49E+08 | 1.351768 | 4.517021 | 8.74E+00 |
| 7496279 | 7420172 | 74467.43 | 10136384 | 9130869 | 7420172 | 6.065678 | 1.72E+01 | 1.18E+01 |
| 3514536 | 3827584 | 105831 | 4672339 | 5249446 | 3633931 | 5.010116 | 14.80775 | 4.32E+00 |
| 59592284 | 46419455 | 89831.66 | 24212924 | 15747165 | 50159222 | 1.560841 | 1.58E+01 | 1.30E+01 |
| 1125835 | 1113732 | 76569.63 | 52892782 | 6855777 | 1113732 | 11.97537 | 8.710443 | 11.61741 |
| 13167589 | 12080514 | 113337 | 7194819 | 7989210 | 12080514 | 29.90869 | 3.78E+01 | 2.58E+01 |
| 148131.3 | 81927.08 | 56755.59 | 10945038 | 2690943 | 113428.6 | 2.539435 | 4.270161 | 28.9225 |
| 8089965 | 8050111 | 138329.3 | 11843853 | 13364427 | 8050111 | 6.730328 | 2.419912 | 12.12426 |
| 7545044 | 7573278 | 83166.95 | 153915.9 | 127187.5 | 7545044 | 2.701814 | 3.432158 | 7.271139 |

| Ratio_Hvidovre_Avedøre_ | Ratio_Skævinge_Avedøre_ | Ratio_Skævinge_Hvidovre_ | Log2_Fold_Change_Hvidovre_Aved | Log2_Fold_Change_Skævinge_Ave | Log2_Fold_Change_Skævinge_Hvid | P_value_Hvidovre_Avedøre_ | P_value_Skævinge_Avedøre_ | P_value_Skævinge_Hvidovre_ |
|---|---|---|---|---|---|---|---|---|
| 1.89E+00 | 1.161 | 6.14E-01 | 9.20E-01 | 0.22 | -0.7 | 3.38E-06 | 0.046401 | 1.05E-05 |
| 0.533 | 0.435 | 0.816 | -0.91 | -1.2 | -0.29 | 0.086423 | 0.015403 | 0.379184 |
| 0.529 | 0.435 | 0.823 | -0.92 | -1.2 | -0.28 | 0.000405 | 7.94E-05 | 0.072344 |
| 17.595 | 0.091 | 0.005 | 4.14 | -3.46 | -7.6 | 1.76E-07 | 5.37E-07 | 9.74E-11 |
| 4.00E-03 | 0.001 | 0.258 | -8.16 | -10.11 | -1.96 | 8.46E-07 | 2.65E-07 | 0.001119 |
| 0.003 | 0.003 | 0.939 | -8.46 | -8.56 | -0.09 | 6.42E-14 | 6.42E-14 | 0.224451 |
| 1.641 | 0.977 | 0.596 | 0.71 | -0.03 | -0.75 | 0.001145 | 0.892283 | 0.000807 |
| 0.53 | 4.69E-01 | 8.85E-01 | -0.92 | -1.09 | -0.18 | 0.005282 | 0.001577 | 0.418058 |
| 8.17E-01 | 0.166 | 0.203 | -0.29 | -2.59 | -2.3 | 0.623429 | 7.76E-06 | 1.13E-05 |
| 2.29E+00 | 8.16E-01 | 3.56E-01 | 1.20E+00 | -0.29 | -1.49 | 0.001442 | 0.096406 | 0.00024 |
| 1.269 | 2.37E+00 | 1.87E+00 | 0.34 | 1.24 | 0.9 | 0.888559 | 0.100226 | 0.181889 |
| 0.067 | 0.154 | 2.3 | -3.9 | -2.7 | 1.2 | 0.006057 | 0.021805 | 0.493609 |
| 1.13E+01 | 0.098 | 9.00E-03 | 3.49E+00 | -3.35 | -6.84 | 1.05E-07 | 2.30E-07 | 6.67E-12 |
| 1.486 | 0.671 | 0.451 | 0.57 | -0.58 | -1.15 | 0.000108 | 0.000112 | 1.84E-06 |
| 1.708 | 0.778 | 0.455 | 0.77 | -0.36 | -1.14 | 0.000416 | 0.011256 | 3.72E-05 |
| 2.20E+00 | 0.986 | 0.449 | 1.14 | -0.02 | -1.16 | 0.008431 | 0.451874 | 0.00249 |
| 1.812 | 0.547 | 0.302 | 0.86 | -0.87 | -1.73 | 0.000628 | 0.000404 | 8.97E-06 |
| 0.41 | 0.075 | 0.184 | -1.29 | -3.73 | -2.44 | 1.70E-06 | 1.27E-10 | 5.19E-08 |
| 1.15E+00 | 1.576 | 1.37E+00 | 0.2 | 0.66 | 0.46 | 0.879141 | 0.3982 | 0.653069 |
| 0.016 | 0.014 | 0.924 | -6 | -6.12 | -0.11 | 1.29E-08 | 1.10E-08 | 0.821082 |
| 1.158 | 2.32E-01 | 2.00E-01 | 0.21 | -2.11 | -2.32 | 0.914149 | 0.001582 | 0.002225 |
| 1.10E-02 | 0.006 | 5.55E-01 | -6.49 | -7.34 | -0.85 | 2.32E-06 | 1.62E-06 | 0.360397 |
| 0.004 | 0.001 | 1.33E-01 | -8.03 | -10.95 | -2.92 | 5.29E-07 | 1.01E-07 | 0.000272 |
| 0.104 | 0.1 | 0.965 | -3.27 | -3.32 | -0.05 | 0.011112 | 0.004901 | 0.710621 |
| 0.432 | 0.217 | 0.503 | -1.21 | -2.2 | -0.99 | 0.032518 | 0.001299 | 0.038439 |
| 0.109 | 0.052 | 0.48 | -3.2 | -4.26 | -1.06 | 0.000626 | 0.000265 | 0.452496 |
| 1.60E+00 | 0.984 | 6.14E-01 | 0.68 | -0.02 | -0.7 | 0.006373 | 0.664417 | 0.00276 |
| 0.524 | 0.605 | 1.154 | -0.93 | -0.73 | 0.21 | 0.437507 | 0.44305 | 0.999927 |
| 0.057 | 0.025 | 0.443 | -4.12 | -5.3 | -1.18 | 1.03E-05 | 2.68E-06 | 0.019708 |
| 2.00E-03 | 4.30E-02 | 2.07E+01 | -8.92 | -4.54 | 4.37 | 3.81E-13 | 5.44E-08 | 9.23E-08 |
| 8.84E-01 | 0.093 | 0.105 | -0.18 | -3.43 | -3.25 | 0.382599 | 6.45E-07 | 8.13E-07 |
| 0.986 | 1.361 | 1.38 | -0.02 | 0.44 | 0.46 | 0.992462 | 0.004984 | 0.004457 |
| 6.41E-01 | 3.30E-01 | 5.15E-01 | -6.40E-01 | -1.6 | -0.96 | 0.25667 | 0.024612 | 0.221725 |
| 0.887 | 0.717 | 0.809 | -0.17 | -0.48 | -0.31 | 0.943131 | 0.789959 | 0.606329 |
| 0.304 | 0.247 | 8.12E-01 | -1.72 | -2.02 | -0.3 | 0.045538 | 0.125095 | 0.714553 |
| 0.925 | 0.666 | 0.719 | -0.11 | -0.59 | -0.48 | 0.338624 | 0.081856 | 0.530916 |
| 5.45E-01 | 6.30E-01 | 1.16E+00 | -0.88 | -0.67 | 0.21 | 0.173065 | 0.276754 | 0.925926 |
| 0.653 | 0.817 | 1.251 | -0.61 | -0.29 | 0.32 | 0.341893 | 0.997821 | 0.317384 |
| 1.76E+00 | 0.384 | 0.219 | 0.81 | -1.38 | -2.19 | 0.27124 | 0.024145 | 0.003994 |
| 0.803 | 0.602 | 0.75 | -0.32 | -0.73 | -0.42 | 0.264572 | 0.009735 | 0.073855 |

| | | | | | | | | |
|---|---|---|---|---|---|---|---|---|
| 0.675 | 0.309 | 0.458 | -0.57 | -1.7 | -1.13 | 0.450553 | 0.020304 | 0.097133 |
| 2.61E-01 | 1.81E-01 | 6.92E-01 | -1.94 | -2.47 | -0.53 | 0.117843 | 0.025568 | 0.474759 |
| 0.682 | 0.306 | 0.448 | -0.55 | -1.71 | -1.16 | 0.056611 | 0.000168 | 0.001254 |
| 0.974 | 0.807 | 8.28E-01 | -0.04 | -0.31 | -0.27 | 0.928418 | 0.517813 | 0.344642 |
| 3.60E+00 | 4.31E+00 | 1.197 | 1.85E+00 | 2.11 | 0.26 | 9.27E-06 | 5.52E-06 | 0.40978 |
| 2.00E-03 | 2.00E-03 | 0.902 | -8.75E+00 | -8.9 | -0.15 | 1.18E-05 | 7.26E-06 | 0.472598 |
| 0.002 | 1.00E-03 | 6.67E-01 | -8.84 | -9.42 | -0.59 | 3.72E-09 | 1.61E-09 | 0.06187 |
| 0.13 | 0.036 | 0.276 | -2.94 | -4.8 | -1.86 | 0.016956 | 0.000945 | 0.046465 |
| 4.90E-02 | 0.019 | 3.92E-01 | -4.35E+00 | -5.7 | -1.35 | 0.004981 | 0.003977 | 0.969468 |
| 4.621 | 0.734 | 0.159 | 2.21 | -0.45 | -2.66 | 0.011318 | 0.996256 | 0.012363 |
| 0.906 | 0.503 | 0.555 | -0.14 | -0.99 | -0.85 | 0.963866 | 0.000299 | 0.000353 |
| 9.00E-03 | 1.40E-02 | 1.538 | -6.77E+00 | -6.15 | 0.62 | 2.41E-06 | 4.02E-06 | 0.288063 |
| 4.00E-03 | 0.003 | 8.41E-01 | -8.11E+00 | -8.36 | -0.25 | 6.42E-14 | 6.42E-14 | 0.026708 |
| 0.014 | 0.007 | 0.478 | -6.16 | -7.23 | -1.07 | 1.56E-05 | 1.02E-05 | 0.570574 |
| 4.56E+00 | 1.594 | 0.349 | 2.19 | 0.67 | -1.52 | 2.01E-05 | 0.011738 | 0.00017 |
| 0.771 | 0.348 | 0.451 | -0.38 | -1.52 | -1.15 | 0.853037 | 0.03303 | 0.064266 |
| 1.424 | 1.49E+00 | 1.05E+00 | 0.51 | 0.58 | 0.07 | 0.321356 | 0.429614 | 0.965563 |
| 0.39 | 3.79E-01 | 9.71E-01 | -1.36 | -1.4 | -0.04 | 5.66E-05 | 5.79E-05 | 0.998805 |
| 0.071 | 0.082 | 1.151 | -3.81 | -3.61 | 0.2 | 0.008773 | 0.014173 | 0.895681 |
| 1.66 | 1.528 | 0.92 | 0.73 | 0.61 | -0.12 | 0.006367 | 0.009165 | 0.932242 |
| 0.617 | 0.396 | 0.642 | -0.7 | -1.34 | -0.64 | 0.19488 | 0.003191 | 0.024751 |
| 3.75E-01 | 0.213 | 5.67E-01 | -1.41E+00 | -2.23 | -0.82 | 5.16E-06 | 5.14E-07 | 9.85E-05 |
| 2.79E+00 | 18.264 | 6.552 | 1.48 | 4.19 | 2.71 | 0.083428 | 0.00058 | 0.005206 |
| 2.044 | 1.052 | 0.515 | 1.03 | 0.07 | -0.96 | 0.000277 | 0.464009 | 0.000647 |
| 0.547 | 0.977 | 1.786 | -0.87 | -0.03 | 0.84 | 0.000215 | 0.817811 | 0.000315 |
| 0.241 | 3.488 | 14.455 | -2.05 | 1.8 | 3.85 | 0.502807 | 0.06477 | 0.015912 |
| 0.102 | 0.041 | 0.403 | -3.29 | -4.6 | -1.31 | 9.36E-06 | 1.34E-06 | 0.00146 |
| 1.042 | 5.54E-01 | 5.31E-01 | 0.06 | -0.85 | -0.91 | 0.59798 | 1.39E-05 | 9.24E-06 |
| 0.285 | 0.576 | 2.019 | -1.81 | -0.8 | 1.01 | 8.19E-07 | 6.92E-05 | 2.10E-05 |
| 0.751 | 3.336 | 4.439 | -0.41 | 1.74 | 2.15 | 0.953547 | 0.137675 | 0.094511 |
| 0.068 | 0.051 | 0.745 | -3.87 | -4.3 | -0.43 | 4.85E-06 | 2.26E-06 | 0.105717 |
| 0.731 | 9.80E-01 | 1.34E+00 | -0.45 | -0.03 | 0.42 | 0.98928 | 0.932524 | 0.874895 |
| 0.842 | 0.683 | 0.812 | -0.25 | -0.55 | -0.3 | 0.076558 | 0.000611 | 0.005976 |
| 7.54E-01 | 7.78E-01 | 1.03E+00 | -0.41 | -0.36 | 0.04 | 0.091844 | 0.422587 | 0.4774 |
| 0.889 | 0.852 | 0.959 | -0.17 | -0.23 | -0.06 | 0.739276 | 0.529435 | 0.926209 |
| 2.407 | 1.28E+00 | 5.30E-01 | 1.27 | 0.35 | -0.92 | 0.886283 | 0.996503 | 0.849042 |
| 0.909 | 5.75E-01 | 6.33E-01 | -0.14 | -0.8 | -0.66 | 0.485797 | 0.00108 | 0.002981 |
| 0.556 | 0.196 | 0.352 | -0.85 | -2.35 | -1.51 | 0.140629 | 0.004557 | 0.053431 |
| 9.90E-01 | 0.557 | 0.563 | -0.01 | -0.84 | -0.83 | 0.954776 | 0.116499 | 0.168817 |
| 0.569 | 0.28 | 0.492 | -0.81 | -1.84 | -1.02 | 0.331511 | 0.015992 | 0.10451 |
| 0.83 | 0.427 | 0.514 | -0.27 | -1.23 | -0.96 | 0.289267 | 0.000423 | 0.001451 |
| 0.724 | 0.338 | 0.466 | -0.47 | -1.57 | -1.1 | 0.033317 | 5.29E-05 | 0.00036 |
| 0.636 | 2.39E-01 | 3.75E-01 | -0.65 | -2.07 | -1.41 | 0.474006 | 0.069314 | 0.329839 |
| 0.371 | 1.59E-01 | 4.27E-01 | -1.43 | -2.66 | -1.23 | 0.090752 | 0.003311 | 0.053409 |
| 0.814 | 0.423 | 0.519 | -0.3 | -1.24 | -0.94 | 0.591842 | 0.031033 | 0.109905 |
| 8.82E-01 | 7.49E-01 | 8.50E-01 | -1.80E-01 | -0.42 | -0.24 | 0.5527 | 0.243935 | 0.760997 |
| 0.51 | 2.51E-01 | 4.93E-01 | -0.97 | -1.99 | -1.02 | 1.73E-06 | 2.78E-08 | 1.66E-06 |
| 1.66E+00 | 2.31E+00 | 1.39E+00 | 7.30E-01 | 1.21 | 0.48 | 0.811593 | 0.065809 | 0.145809 |
| 0.909 | 0.409 | 4.49E-01 | -0.14 | -1.29 | -1.15 | 0.519277 | 0.078114 | 0.019474 |
| 1.65E-01 | 0.069 | 4.20E-01 | -2.6 | -3.85 | -1.25 | 7.81E-06 | 9.76E-07 | 0.000554 |
| 2.515 | 0.972 | 0.386 | 1.33 | -0.04 | -1.37 | 1.05E-06 | 0.27751 | 7.88E-07 |

| | | | | | | | | |
|---|---|---|---|---|---|---|---|---|
| 1.03E+00 | 0.57 | 5.55E-01 | 4.00E-02 | -0.81 | -0.85 | 0.894428 | 0.000172 | 0.000225 |
| 0.101 | 0.143 | 1.419 | -3.31 | -2.8 | 0.5 | 0.008476 | 0.033848 | 0.472155 |
| 1.356 | 1.04 | 0.767 | 0.44 | 0.06 | -0.38 | 0.705986 | 0.996054 | 0.753483 |
| 0.451 | 0.219 | 0.485 | -1.15 | -2.19 | -1.04 | 1.40E-06 | 2.34E-08 | 1.78E-06 |
| 0.51 | 0.251 | 0.493 | -0.97 | -1.99 | -1.02 | 1.73E-06 | 2.78E-08 | 1.66E-06 |
| 0.023 | 0.018 | 0.798 | -5.46 | -5.79 | -0.33 | 1.90E-05 | 1.60E-05 | 0.9141 |
| 0.808 | 0.346 | 0.428 | -0.31 | -1.53 | -1.22 | 0.879341 | 0.173253 | 0.317162 |
| 0.619 | 0.814 | 1.314 | -0.69 | -0.3 | 0.39 | 0.011228 | 0.168669 | 0.138565 |
| 9.10E-02 | 0.037 | 4.12E-01 | -3.46E+00 | -4.74 | -1.28 | 0.00029 | 9.00E-05 | 0.187252 |
| 1.159 | 0.558 | 0.482 | 0.21 | -0.84 | -1.05 | 0.852844 | 0.14239 | 0.071238 |
| 0.04 | 0.142 | 3.55 | -4.64 | -2.81 | 1.83 | 0.064414 | 0.068033 | 0.998951 |
| 0.44 | 4.43E-01 | 1.01E+00 | -1.18 | -1.17 | 0.01 | 1.57E-06 | 1.51E-06 | 0.978762 |
| 1.049 | 9.30E-01 | 8.87E-01 | 0.07 | -0.1 | -0.17 | 0.844476 | 0.907056 | 0.989625 |
| 2.631 | 0.62 | 0.236 | 1.4 | -0.69 | -2.09 | 0.00041 | 0.047844 | 6.77E-05 |
| 0.827 | 6.17E-01 | 7.46E-01 | -0.27 | -0.7 | -0.42 | 0.018462 | 9.74E-05 | 0.001226 |
| 1.206 | 1.125 | 9.33E-01 | 0.27 | 0.17 | -0.1 | 0.157045 | 0.662497 | 0.461629 |
| 1.29E+00 | 1.067 | 8.29E-01 | 3.60E-01 | 0.09 | -0.27 | 0.06129 | 0.990785 | 0.052177 |
| 1.24E+00 | 0.91 | 7.36E-01 | 3.10E-01 | -0.14 | -0.44 | 0.057011 | 0.422326 | 0.011923 |
| 7.513 | 3.93E-01 | 5.20E-02 | 2.91 | -1.35 | -4.26 | 9.74E-05 | 0.004534 | 9.04E-06 |
| 37.413 | 0.881 | 0.024 | 5.23 | -0.18 | -5.41 | 9.08E-14 | 0.098329 | 7.26E-14 |
| 0.797 | 0.913 | 1.146 | -0.33 | -0.13 | 0.2 | 0.095608 | 0.270905 | 0.703081 |
| 0.69 | 0.441 | 0.64 | -0.54 | -1.18 | -0.64 | 0.007257 | 6.99E-05 | 0.001484 |
| 0.543 | 0.199 | 0.367 | -0.88 | -2.33 | -1.45 | 0.002412 | 1.86E-05 | 0.000425 |
| 6.60E-02 | 1.15E-01 | 1.73E+00 | -3.92 | -3.12 | 0.79 | 1.25E-05 | 5.16E-05 | 0.041728 |
| 26.895 | 2.202 | 8.20E-02 | 4.75 | 1.14 | -3.61 | 4.17E-07 | 0.002096 | 1.15E-06 |
| 1.133 | 0.757 | 0.668 | 0.18 | -0.4 | -0.58 | 0.584759 | 0.581475 | 0.175907 |
| 1.09E+00 | 0.416 | 0.382 | 0.12 | -1.27 | -1.39 | 0.255524 | 1.20E-05 | 6.04E-06 |
| 0.011 | 0.004 | 0.333 | -6.48 | -8.06 | -1.59 | 0.000264 | 9.32E-05 | 0.244734 |
| 0.048 | 0.014 | 3.03E-01 | -4.39 | -6.11 | -1.72 | 0.00236 | 0.001342 | 0.784426 |
| 0.028 | 0.033 | 1.165 | -5.13 | -4.91 | 0.22 | 3.17E-10 | 6.84E-10 | 0.112681 |
| 1.14E+00 | 1.49E+00 | 1.31E+00 | 0.19 | 0.57 | 0.38 | 0.132 | 0.001077 | 0.008369 |
| 0.418 | 0.077 | 0.185 | -1.26 | -3.69 | -2.43 | 0.01744 | 5.94E-05 | 0.00062 |
| 1.132 | 2.845 | 2.513 | 0.18 | 1.51 | 1.33 | 0.081892 | 2.30E-06 | 5.33E-06 |
| 0.099 | 0.105 | 1.06E+00 | -3.33 | -3.26 | 0.08 | 0.000114 | 9.63E-05 | 0.946918 |
| 0.723 | 6.48E-01 | 8.96E-01 | -0.47 | -0.63 | -0.16 | 0.003075 | 0.000358 | 0.069636 |
| 0.761 | 3.037 | 3.99 | -0.39 | 1.6 | 2 | 0.023348 | 7.41E-06 | 2.22E-06 |
| 1.828 | 0.547 | 0.299 | 0.87 | -0.87 | -1.74 | 0.000686 | 0.000419 | 9.62E-06 |
| 0.12 | 0.02 | 0.167 | -3.06 | -5.64 | -2.58 | 4.79E-08 | 6.01E-13 | 1.35E-07 |
| 3.284 | 0.289 | 0.088 | 1.72 | -1.79 | -3.51 | 1.53E-05 | 9.88E-06 | 3.50E-07 |
| 2.344 | 1.753 | 0.748 | 1.23 | 0.81 | -0.42 | 6.58E-05 | 0.000404 | 0.045863 |
| 1.126 | 7.40E-01 | 6.57E-01 | 0.17 | -0.43 | -0.61 | 0.236002 | 0.001084 | 0.000298 |
| 1.177 | 1.016 | 0.863 | 0.24 | 0.02 | -0.21 | 0.929144 | 0.689445 | 0.487409 |
| 2.223 | 7.57E-01 | 3.41E-01 | 1.15 | -0.4 | -1.55 | 7.82E-05 | 0.050412 | 1.88E-05 |
| 0.61 | 3.617 | 5.93 | -0.71 | 1.85 | 2.57 | 0.000719 | 1.68E-06 | 4.59E-07 |
| 0.018 | 1.60E-02 | 8.89E-01 | -5.8 | -5.97 | -0.17 | 3.83E-10 | 2.36E-10 | 0.359764 |
| 8.34E-01 | 0.476 | 0.57 | -0.26 | -1.07 | -0.81 | 0.507495 | 0.003649 | 0.011747 |
| 1.013 | 1.98E-01 | 1.96E-01 | 0.02 | -2.33 | -2.35 | 0.696937 | 6.90E-07 | 6.03E-07 |
| 0.854 | 0.648 | 0.758 | -0.23 | -0.63 | -0.4 | 0.036348 | 8.06E-05 | 0.000607 |
| 0.653 | 0.462 | 7.08E-01 | -0.61 | -1.11 | -0.5 | 0.000102 | 3.13E-06 | 0.000388 |
| 0.045 | 0.012 | 2.68E-01 | -4.48 | -6.38 | -1.9 | 1.41E-06 | 2.43E-07 | 0.000123 |
| 1.147 | 0.382 | 0.333 | 0.2 | -1.39 | -1.59 | 0.734105 | 0.000749 | 0.000436 |

| | | | | | | | | |
|---|---|---|---|---|---|---|---|---|
| 6.26E-01 | 0.38 | 6.06E-01 | -0.68 | -1.4 | -0.72 | 0.023365 | 0.004307 | 0.309405 |
| 1.005 | 5.60E-01 | 5.57E-01 | 0.01 | -0.84 | -0.84 | 0.906807 | 0.000199 | 0.000257 |
| 0.373 | 0.384 | 1.03 | -1.42 | -1.38 | 0.04 | 0.000166 | 0.000203 | 0.938254 |
| 1.37E+00 | 0.979 | 7.15E-01 | 4.50E-01 | -0.03 | -0.48 | 0.329286 | 0.999496 | 0.341229 |
| 1.964 | 7.65E-01 | 3.90E-01 | 0.97 | -0.39 | -1.36 | 0.15196 | 0.192771 | 0.013595 |
| 2.267 | 0.195 | 0.086 | 1.18 | -2.36 | -3.54 | 0.00575 | 0.000101 | 1.02E-05 |
| 1.222 | 7.00E-03 | 6.00E-03 | 0.29 | -7.08 | -7.37 | 0.427431 | 1.12E-06 | 8.95E-07 |
| 0.61 | 0.1 | 0.163 | -0.71 | -3.33 | -2.61 | 5.77E-05 | 9.41E-10 | 2.36E-08 |
| 0.829 | 0.521 | 0.628 | -0.27 | -0.94 | -0.67 | 0.146671 | 0.000251 | 0.001174 |
| 0.563 | 0.54 | 0.96 | -0.83 | -0.89 | -0.06 | 0.769415 | 0.174242 | 0.409838 |
| 0.651 | 0.087 | 0.133 | -0.62 | -3.53 | -2.91 | 0.00029 | 5.46E-09 | 4.98E-08 |
| 7.67E-01 | 0.645 | 8.41E-01 | -3.80E-01 | -0.63 | -0.25 | 0.018823 | 0.001485 | 0.08485 |
| 2.766 | 2.91E+00 | 1.05E+00 | 1.47 | 1.54 | 0.07 | 5.80E-06 | 5.20E-06 | 0.951062 |
| 0.49 | 0.243 | 0.496 | -1.03 | -2.04 | -1.01 | 0.920462 | 0.072261 | 0.11864 |
| 0.919 | 0.116 | 0.126 | -0.12 | -3.11 | -2.99 | 0.466476 | 2.10E-06 | 2.97E-06 |
| 7.02E-01 | 0.854 | 1.22E+00 | -0.51 | -0.23 | 0.28 | 0.001691 | 0.025236 | 0.075005 |
| 0.829 | 0.34 | 0.41 | -0.27 | -1.56 | -1.29 | 0.887778 | 0.000385 | 0.000529 |
| 0.954 | 0.811 | 0.851 | -0.07 | -0.3 | -0.23 | 0.921776 | 0.23966 | 0.381001 |
| 141.689 | 2.58E-01 | 2.00E-03 | 7.15 | -1.95 | -9.1 | 1.32E-13 | 1.21E-06 | 6.42E-14 |
| 1.11E+00 | 1.18E+00 | 1.07 | 1.40E-01 | 0.24 | 0.1 | 0.180633 | 0.19537 | 0.99788 |
| 9.63E-01 | 1.086 | 1.13E+00 | -0.05 | 0.12 | 0.17 | 0.986107 | 0.506369 | 0.427018 |
| 0.856 | 0.492 | 0.575 | -0.22 | -1.02 | -0.8 | 0.176095 | 3.82E-05 | 0.000108 |
| 0.826 | 1.15 | 1.39E+00 | -0.28 | 0.2 | 0.48 | 0.272305 | 0.671059 | 0.089473 |
| 8.49E-01 | 1.278 | 1.51E+00 | -0.24 | 0.35 | 0.59 | 0.168312 | 0.152385 | 0.012137 |
| 8.61E-01 | 1.40E+00 | 1.63E+00 | -2.20E-01 | 0.49 | 0.7 | 0.468039 | 0.068829 | 0.01557 |
| 9.54E-01 | 1.005 | 1.054 | -0.07 | 0.01 | 0.08 | 0.681242 | 0.65915 | 0.999167 |
| 1.196 | 0.845 | 0.707 | 0.26 | -0.24 | -0.5 | 0.160497 | 0.128667 | 0.010132 |
| 0.776 | 0.355 | 0.457 | -0.37 | -1.5 | -1.13 | 0.031076 | 1.92E-05 | 9.95E-05 |
| 0.777 | 0.279 | 0.36 | -0.36 | -1.84 | -1.48 | 0.14018 | 2.13E-05 | 6.01E-05 |
| 1.12E+00 | 0.348 | 0.31 | 0.17 | -1.52 | -1.69 | 0.091419 | 1.36E-06 | 8.63E-07 |
| 2.90E+03 | 3.795 | 1.00E-03 | 1.15E+01 | 1.92 | -9.58 | 5.84E-05 | 0.786452 | 8.22E-05 |
| 1.064 | 7.58E-01 | 7.12E-01 | 0.09 | -0.4 | -0.49 | 0.567393 | 0.005178 | 0.001977 |
| 0.994 | 1.384 | 1.392 | -0.01 | 0.47 | 0.48 | 0.977949 | 0.006303 | 0.005175 |
| 4.332 | 14.469 | 3.34 | 2.12 | 3.85 | 1.74 | 0.019276 | 0.000463 | 0.012939 |
| 0.359 | 0.074 | 0.207 | -1.48 | -3.75 | -2.27 | 0.15641 | 0.015188 | 0.2128 |
| 0.858 | 0.067 | 0.078 | -0.22 | -3.9 | -3.68 | 0.752101 | 0.000232 | 0.00037 |
| 0.541 | 5.899 | 10.91 | -0.89 | 2.56 | 3.45 | 0.004474 | 4.44E-05 | 5.03E-06 |
| 0.881 | 5.882 | 6.68 | -0.18 | 2.56 | 2.74 | 0.99633 | 0.029021 | 0.026371 |
| 2.37E-01 | 20.963 | 8.83E+01 | -2.07 | 4.39 | 6.46 | 0.3823 | 0.009375 | 0.002388 |
| 1.145 | 1.10E+00 | 9.58E-01 | 0.2 | 0.13 | -0.06 | 0.075142 | 0.486552 | 0.345493 |
| 3.90E-01 | 0.173 | 4.42E-01 | -1.36 | -2.53 | -1.18 | 0.082652 | 0.016162 | 0.414431 |
| 0.671 | 3.52E-01 | 5.25E-01 | -0.58 | -1.51 | -0.93 | 0.095118 | 0.000788 | 0.007145 |
| 0.665 | 0.014 | 0.022 | -0.59 | -6.12 | -5.53 | 7.80E-06 | 6.42E-14 | 6.42E-14 |
| 0.968 | 0.639 | 0.661 | -0.05 | -0.65 | -0.6 | 0.873135 | 0.00108 | 0.00074 |
| 0.193 | 0.236 | 1.224 | -2.37 | -2.08 | 0.29 | 0.206842 | 0.23369 | 0.994764 |
| 0.687 | 0.171 | 0.249 | -0.54 | -2.55 | -2.01 | 0.046622 | 5.99E-06 | 2.09E-05 |
| 1.064 | 0.636 | 0.597 | 0.09 | -0.65 | -0.74 | 0.761798 | 0.001973 | 0.001102 |
| 8.96E-01 | 1.66E+00 | 1.85E+00 | -1.60E-01 | 0.73 | 0.89 | 0.642931 | 0.001521 | 0.000742 |
| 1.15E+00 | 2.07E-01 | 1.81E-01 | 2.00E-01 | -2.27 | -2.47 | 0.211574 | 5.30E-07 | 3.84E-07 |
| 0.608 | 2.10E-01 | 3.45E-01 | -0.72 | -2.25 | -1.53 | 0.001328 | 2.03E-06 | 2.15E-05 |
| 0.65 | 0.263 | 0.404 | -0.62 | -1.93 | -1.31 | 0.003194 | 5.44E-06 | 5.90E-05 |

| | | | | | | | | |
|---|---|---|---|---|---|---|---|---|
| 9.39E-01 | 3.60E-02 | 3.80E-02 | -9.00E-02 | -4.8 | -4.71 | 0.989662 | 3.89E-06 | 4.07E-06 |
| 0.746 | 0.677 | 0.907 | -0.42 | -0.56 | -0.14 | 0.008561 | 0.002677 | 0.485856 |
| 0.236 | 0.124 | 0.525 | -2.08 | -3.01 | -0.93 | 1.21E-06 | 1.97E-07 | 0.000104 |
| 1.327 | 0.43 | 0.324 | 0.41 | -1.22 | -1.62 | 0.000679 | 1.22E-06 | 3.45E-07 |
| 0.163 | 0.454 | 2.788 | -2.62 | -1.14 | 1.48 | 0.109729 | 0.164872 | 0.945884 |
| 1.069 | 0.176 | 0.165 | 0.1 | -2.5 | -2.6 | 0.514178 | 8.36E-07 | 6.93E-07 |
| 0.655 | 5.71E-01 | 8.72E-01 | -0.61 | -0.81 | -0.2 | 0.000635 | 8.95E-05 | 0.042805 |
| 0.003 | 0.003 | 0.964 | -8.59 | -8.64 | -0.05 | 6.42E-14 | 6.42E-14 | 0.197308 |
| 1.009 | 7.15E-01 | 7.09E-01 | 0.01 | -0.48 | -0.5 | 0.943197 | 0.004664 | 0.003437 |
| 1.01E+00 | 3.15E-01 | 3.13E-01 | 0.01 | -1.67 | -1.68 | 1 | 6.12E-06 | 6.12E-06 |
| 4.00E-03 | 1.00E-03 | 0.333 | -7.88 | -9.47 | -1.59 | 1.63E-05 | 5.48E-06 | 0.070811 |
| 1.774 | 0.12 | 0.068 | 0.83 | -3.06 | -3.89 | 0.01499 | 1.94E-05 | 4.01E-06 |
| 3.219 | 2.552 | 7.93E-01 | 1.69 | 1.35 | -0.34 | 0.009769 | 0.026708 | 0.655298 |
| 1.144 | 2.22E+00 | 1.94E+00 | 0.19 | 1.15 | 0.96 | 0.399027 | 0.056679 | 0.328265 |
| 1.01E+00 | 0.63 | 6.26E-01 | 1.00E-02 | -0.67 | -0.68 | 0.9785 | 0.004308 | 0.00358 |
| 0.759 | 0.507 | 0.668 | -0.4 | -0.98 | -0.58 | 0.037332 | 0.000382 | 0.005404 |
| 0.56 | 0.423 | 0.755 | -0.84 | -1.24 | -0.41 | 0.00463 | 0.000596 | 0.102461 |
| 0.704 | 0.738 | 1.048 | -0.51 | -0.44 | 0.07 | 0.012272 | 0.012344 | 0.999984 |
| 0.963 | 0.592 | 0.614 | -0.05 | -0.76 | -0.7 | 0.783338 | 0.011466 | 0.024424 |
| 0.965 | 0.64 | 0.663 | -0.05 | -0.64 | -0.59 | 0.994165 | 0.205804 | 0.23411 |
| 0.448 | 0.185 | 0.413 | -1.16 | -2.43 | -1.27 | 0.015654 | 0.000596 | 0.023959 |
| 1.486 | 0.591 | 0.398 | 0.57 | -0.76 | -1.33 | 0.000446 | 0.000212 | 5.11E-06 |
| 0.868 | 1.09E+00 | 1.25E+00 | -0.2 | 0.12 | 0.33 | 0.108186 | 0.056419 | 0.003916 |
| 4.00E-03 | 1.60E-02 | 4.18E+00 | -8.01E+00 | -5.95 | 2.06 | 0.000176 | 0.000538 | 0.265723 |
| 0.012 | 0.012 | 9.47E-01 | -6.36 | -6.44 | -0.08 | 5.23E-08 | 4.61E-08 | 0.848312 |
| 0.644 | 0.246 | 0.383 | -0.64 | -2.02 | -1.39 | 0.291449 | 0.235693 | 0.983662 |
| 0.005 | 0.005 | 9.53E-01 | -7.51 | -7.58 | -0.07 | 9.96E-08 | 8.88E-08 | 0.853318 |
| 1.144 | 0.41 | 0.359 | 0.19 | -1.29 | -1.48 | 0.050079 | 6.11E-06 | 2.29E-06 |
| 0.049 | 2.50E-02 | 5.01E-01 | -4.34 | -5.34 | -1 | 0.008262 | 0.010253 | 0.976568 |
| 0.102 | 0.127 | 1.25 | -3.29 | -2.97 | 0.32 | 0.008282 | 0.012135 | 0.930385 |
| 0.183 | 0.343 | 1.876 | -2.45 | -1.55 | 0.91 | 0.319776 | 0.287176 | 0.995593 |
| 0.828 | 0.597 | 0.721 | -0.27 | -0.74 | -0.47 | 0.104499 | 0.002438 | 0.030779 |
| 3.691 | 0.314 | 0.085 | 1.88 | -1.67 | -3.56 | 0.000717 | 0.000688 | 1.31E-05 |
| 0.808 | 3.45E-01 | 0.427 | -0.31 | -1.54 | -1.23 | 0.308299 | 0.000378 | 0.001217 |
| 2.319 | 0.841 | 3.63E-01 | 1.21 | -0.25 | -1.46 | 5.16E-05 | 0.057258 | 1.39E-05 |
| 64.548 | 1.151 | 0.018 | 6.01 | 0.2 | -5.81 | 5.53E-06 | 0.946053 | 6.21E-06 |
| 86.941 | 6.40E-01 | 7.00E-03 | 6.44 | -0.64 | -7.09 | 1.24E-12 | 0.000372 | 9.68E-14 |
| 5.85E+01 | 0.559 | 1.00E-02 | 5.87 | -0.84 | -6.71 | 1.91E-10 | 0.000191 | 6.53E-12 |
| 1.26E+00 | 0.81 | 6.42E-01 | 3.40E-01 | -0.3 | -0.64 | 0.042411 | 0.017429 | 0.00091 |
| 1.87E+00 | 0.274 | 1.46E-01 | 9.00E-01 | -1.87 | -2.77 | 0.002875 | 4.38E-05 | 4.21E-06 |
| 0.965 | 1.06E+00 | 1.10E+00 | -0.05 | 0.09 | 0.14 | 0.992891 | 0.705833 | 0.641216 |
| 0.722 | 0.996 | 1.378 | -0.47 | -0.01 | 0.46 | 0.086319 | 0.897651 | 0.152531 |
| 1.036 | 0.539 | 5.20E-01 | 0.05 | -0.89 | -0.94 | 0.938914 | 0.000664 | 0.000522 |
| 1.145 | 0.689 | 6.02E-01 | 0.2 | -0.54 | -0.73 | 0.741637 | 0.008311 | 0.00397 |
| 0.602 | 0.523 | 0.869 | -0.73 | -0.94 | -0.2 | 0.001123 | 0.000151 | 0.05348 |
| 6.28E-01 | 1.172 | 1.87E+00 | -0.67 | 0.23 | 0.9 | 0.510054 | 0.775503 | 0.884131 |
| 7.90E-01 | 0.587 | 0.743 | -0.34 | -0.77 | -0.43 | 0.009853 | 0.000159 | 0.004329 |
| 0.011 | 0.01 | 0.922 | -6.55 | -6.66 | -0.12 | 6.44E-14 | 6.42E-14 | 0.138668 |
| 9.97E-01 | 0.39 | 3.92E-01 | 0 | -1.36 | -1.35 | 0.909596 | 0.472125 | 0.292691 |
| 0.044 | 0.084 | 1.902 | -4.5 | -3.57 | 0.93 | 0.003493 | 0.003387 | 0.999372 |
| 0.783 | 0.011 | 0.014 | -0.35 | -6.48 | -6.13 | 0.008401 | 6.88E-14 | 1.03E-13 |

| | | | | | | | | |
|---|---|---|---|---|---|---|---|---|
| 4.73E-01 | 1.72E-01 | 3.63E-01 | -1.08E+00 | -2.54 | -1.46 | 0.380733 | 0.026453 | 0.157859 |
| 3.174 | 2.645 | 0.833 | 1.67 | 1.4 | -0.26 | 8.59E-06 | 3.95E-05 | 0.026669 |
| 1.355 | 0.954 | 0.704 | 0.44 | -0.07 | -0.51 | 0.014613 | 0.710141 | 0.006269 |
| 9.392 | 1.11E+00 | 1.18E-01 | 3.23 | 0.15 | -3.08 | 0.061357 | 0.919499 | 0.100766 |
| 9.02E-01 | 0.677 | 7.50E-01 | -0.15 | -0.56 | -0.41 | 0.670451 | 0.0468 | 0.142144 |
| 1.76E+00 | 1.154 | 6.55E-01 | 0.82 | 0.21 | -0.61 | 0.10439 | 0.650218 | 0.329261 |
| 0.96 | 0.709 | 7.39E-01 | -0.06 | -0.5 | -0.44 | 0.996938 | 0.074686 | 0.067998 |
| 0.973 | 0.887 | 0.912 | -0.04 | -0.17 | -0.13 | 0.965483 | 0.665523 | 0.52503 |
| 0.478 | 0.122 | 0.256 | -1.06 | -3.03 | -1.97 | 0.429835 | 0.016634 | 0.081922 |
| 0.667 | 2.60E-01 | 3.90E-01 | -0.58 | -1.94 | -1.36 | 0.51015 | 0.003401 | 0.010766 |
| 6.01E-01 | 2.24E-01 | 3.72E-01 | -7.30E-01 | -2.16 | -1.42 | 0.050566 | 0.000431 | 0.00506 |
| 0.179 | 5.61E-01 | 3.13E+00 | -2.48 | -0.83 | 1.65 | 0.074367 | 0.991719 | 0.086869 |
| 0.998 | 0.271 | 0.271 | 0 | -1.89 | -1.88 | 0.573265 | 0.000657 | 0.00147 |
| 7.30E-01 | 6.38E-01 | 0.874 | -4.50E-01 | -0.65 | -0.19 | 0.35791 | 0.308304 | 0.991239 |
| 2.576 | 275.692 | 107.036 | 1.36 | 8.11 | 6.74 | 0.071701 | 2.02E-06 | 4.63E-06 |
| 0.025 | 4.20E-02 | 1.69E+00 | -5.33 | -4.57 | 0.76 | 0.000376 | 0.000451 | 0.960213 |
| 5.388 | 24.763 | 4.596 | 2.43 | 4.63 | 2.2 | 0.896863 | 0.04112 | 0.07162 |
| 0.981 | 3.192 | 3.253 | -0.03 | 1.67 | 1.7 | 0.57907 | 0.564264 | 0.166877 |
| 0.977 | 0.833 | 0.853 | -0.03 | -0.26 | -0.23 | 0.900382 | 0.164969 | 0.285782 |
| 0.667 | 0.511 | 0.766 | -0.58 | -0.97 | -0.38 | 0.010967 | 0.000347 | 0.014182 |
| 3.91E-01 | 7.651 | 19.564 | -1.35 | 2.94 | 4.29 | 0.25926 | 0.004643 | 0.001027 |
| 0.724 | 0.678 | 0.937 | -0.47 | -0.56 | -0.09 | 0.945409 | 0.845365 | 0.96906 |
| 0.859 | 0.375 | 0.437 | -0.22 | -1.42 | -1.2 | 0.220118 | 0.000877 | 0.004317 |
| 0.963 | 0.445 | 0.462 | -0.05 | -1.17 | -1.11 | 0.956254 | 0.08426 | 0.05883 |
| 5.64E-01 | 12.949 | 22.959 | -0.83 | 3.69 | 4.52 | 0.647482 | 0.003787 | 0.00169 |
| 0.3 | 11.813 | 39.373 | -1.74 | 3.56 | 5.3 | 0.024172 | 0.000172 | 2.68E-05 |
| 0.857 | 21.943 | 2.56E+01 | -0.22 | 4.46 | 4.68 | 0.614575 | 7.50E-05 | 4.62E-05 |
| 8.34E-01 | 2.09E+00 | 2.51E+00 | -0.26 | 1.06 | 1.33 | 0.284615 | 0.182025 | 0.022643 |
| 3.911 | 0.343 | 0.088 | 1.97 | -1.54 | -3.51 | 1.10E-06 | 4.21E-06 | 3.57E-08 |
| 3.793 | 0.11 | 0.029 | 1.92 | -3.19 | -5.11 | 0.000611 | 5.55E-05 | 2.67E-06 |
| 0.007 | 2.40E-02 | 3.39E+00 | -7.14 | -5.37 | 1.76 | 0.000875 | 0.001777 | 0.662291 |
| 0.855 | 1.261 | 1.474 | -0.23 | 0.33 | 0.56 | 0.991412 | 0.451589 | 0.393087 |
| 5.00E-03 | 0.003 | 7.73E-01 | -7.79E+00 | -8.16 | -0.37 | 2.35E-05 | 1.52E-05 | 0.590983 |
| 0.005 | 0.006 | 1.148 | -7.55 | -7.35 | 0.2 | 9.60E-05 | 0.00013 | 0.85157 |
| 0.381 | 0.255 | 0.669 | -1.39 | -1.97 | -0.58 | 0.102332 | 0.107839 | 0.999068 |
| 1.01E+00 | 0.762 | 0.755 | 0.01 | -0.39 | -0.41 | 0.999776 | 0.003173 | 0.003231 |
| 0.221 | 0.16 | 0.725 | -2.18 | -2.64 | -0.46 | 0.035842 | 0.014897 | 0.73737 |
| 0.143 | 0.08 | 0.559 | -2.8 | -3.64 | -0.84 | 2.18E-06 | 8.99E-07 | 0.010434 |
| 0.211 | 0.036 | 0.169 | -2.24 | -4.81 | -2.56 | 1.72E-06 | 1.09E-08 | 8.42E-07 |
| 60.604 | 1.653 | 2.70E-02 | 5.92 | 0.73 | -5.2 | 1.80E-06 | 0.13904 | 3.34E-06 |
| 0.915 | 0.61 | 0.667 | -0.13 | -0.71 | -0.58 | 0.35732 | 5.84E-05 | 0.000128 |
| 0.902 | 0.196 | 0.218 | -0.15 | -2.35 | -2.2 | 0.582341 | 0.000405 | 0.000843 |
| 1.90E-02 | 0.003 | 1.79E-01 | -5.71E+00 | -8.19 | -2.49 | 0.006089 | 0.002983 | 0.739811 |
| 0.764 | 0.097 | 0.126 | -0.39 | -3.37 | -2.98 | 0.008695 | 5.49E-08 | 1.46E-07 |
| 2.60E-02 | 0.032 | 1.20E+00 | -5.25E+00 | -4.98 | 0.26 | 0.000573 | 0.000397 | 0.857544 |
| 0.01 | 0.004 | 0.382 | -6.69 | -8.08 | -1.39 | 2.74E-05 | 9.63E-06 | 0.101976 |
| 0.006 | 0.008 | 1.226 | -7.31 | -7.01 | 0.29 | 3.76E-07 | 4.06E-07 | 0.897398 |
| 1.023 | 0.48 | 0.47 | 0.03 | -1.06 | -1.09 | 0.662723 | 7.54E-05 | 4.83E-05 |
| 0.006 | 3.00E-03 | 5.01E-01 | -7.32 | -8.32 | -1 | 8.98E-07 | 6.05E-07 | 0.109497 |
| 0.762 | 1.46E-01 | 1.92E-01 | -0.39 | -2.77 | -2.38 | 0.915992 | 0.046568 | 0.076807 |
| 7.00E-01 | 2.801 | 4.00E+00 | -5.10E-01 | 1.49 | 2 | 0.000307 | 7.24E-07 | 1.55E-07 |

| | | | | | | | | |
|---|---|---|---|---|---|---|---|---|
| 0.011 | 0.029 | 2.66 | -6.5 | -5.09 | 1.41 | 0.000318 | 0.000755 | 0.467695 |
| 3.92E-01 | 0.141 | 3.60E-01 | -1.35E+00 | -2.83 | -1.47 | 2.14E-05 | 5.68E-07 | 2.35E-05 |
| 8.02E-01 | 0.37 | 4.61E-01 | -0.32 | -1.44 | -1.12 | 0.044527 | 3.57E-05 | 0.000185 |
| 0.2 | 0.233 | 1.166 | -2.32 | -2.1 | 0.22 | 4.04E-06 | 9.48E-06 | 0.127003 |
| 0.716 | 0.239 | 0.333 | -0.48 | -2.07 | -1.58 | 0.000214 | 4.71E-08 | 3.13E-07 |
| 7.44E-01 | 0.418 | 5.61E-01 | -4.30E-01 | -1.26 | -0.83 | 0.26575 | 0.003451 | 0.02052 |
| 0.151 | 0.123 | 0.809 | -2.72 | -3.03 | -0.31 | 0.054796 | 0.03649 | 0.941278 |
| 0.602 | 0.521 | 0.865 | -0.73 | -0.94 | -0.21 | 0.000252 | 4.04E-05 | 0.03304 |
| 0.201 | 0.091 | 0.454 | -2.31 | -3.45 | -1.14 | 0.001535 | 0.000118 | 0.020035 |
| 0.042 | 0.025 | 0.589 | -4.58 | -5.34 | -0.76 | 0.00433 | 0.002538 | 0.831496 |
| 0.802 | 0.834 | 1.04 | -0.32 | -0.26 | 0.06 | 0.108433 | 0.237797 | 0.815266 |
| 0.113 | 0.068 | 0.599 | -3.14 | -3.88 | -0.74 | 4.57E-08 | 2.93E-09 | 9.91E-05 |
| 0.095 | 0.087 | 0.912 | -3.39 | -3.53 | -0.13 | 7.63E-05 | 6.79E-05 | 0.971761 |
| 0.402 | 0.255 | 0.633 | -1.31 | -1.97 | -0.66 | 0.006963 | 0.000248 | 0.012899 |
| 0.767 | 0.472 | 0.615 | -0.38 | -1.08 | -0.7 | 0.100274 | 0.001094 | 0.010703 |
| 0.564 | 9.70E-02 | 1.72E-01 | -0.83 | -3.37 | -2.54 | 0.003628 | 2.13E-06 | 1.43E-05 |
| 0.953 | 5.59E-01 | 5.86E-01 | -0.07 | -0.84 | -0.77 | 0.963501 | 0.000363 | 0.000306 |
| 2.31E-01 | 4.00E-03 | 1.60E-02 | -2.11E+00 | -8.1 | -5.99 | 6.83E-07 | 6.42E-14 | 1.11E-12 |
| 1.64E-01 | 0.079 | 0.482 | -2.61 | -3.66 | -1.05 | 0.021426 | 0.005763 | 0.475561 |
| 0.212 | 2.27E-01 | 1.07E+00 | -2.24 | -2.14 | 0.1 | 1.63E-06 | 1.97E-06 | 0.699065 |
| 1.16E+00 | 1.548 | 1.33E+00 | 2.20E-01 | 0.63 | 0.41 | 0.124924 | 0.000398 | 0.002354 |
| 0.2 | 0.255 | 1.28E+00 | -2.32 | -1.97 | 0.35 | 2.63E-06 | 7.98E-06 | 0.039069 |
| 7.41E-01 | 0.786 | 1.06E+00 | -4.30E-01 | -0.35 | 0.09 | 0.047411 | 0.109601 | 0.79 |
| 0.915 | 0.352 | 0.384 | -0.13 | -1.51 | -1.38 | 0.483475 | 9.09E-06 | 1.49E-05 |
| 0.039 | 0.031 | 8.08E-01 | -4.69 | -5 | -0.31 | 0.007319 | 0.003094 | 0.658398 |
| 5.91E-01 | 7.85E+00 | 1.33E+01 | -7.60E-01 | 2.97 | 3.73 | 8.69E-05 | 2.99E-08 | 1.61E-09 |
| 1.066 | 3.382 | 3.173 | 0.09 | 1.76 | 1.67 | 0.525783 | 0.153332 | 0.036666 |
| 0.022 | 4.037 | 179.615 | -5.48 | 2.01 | 7.49 | 0.002163 | 0.036074 | 0.000199 |
| 1.088 | 0.677 | 6.23E-01 | 0.12 | -0.56 | -0.68 | 0.999656 | 0.843804 | 0.831308 |
| 8.18E-01 | 0.352 | 4.30E-01 | -2.90E-01 | -1.51 | -1.22 | 0.398844 | 0.000185 | 0.000448 |
| 0.65 | 0.279 | 0.43 | -0.62 | -1.84 | -1.22 | 0.150093 | 0.002599 | 0.024054 |
| 0.067 | 0.076 | 1.125 | -3.89 | -3.72 | 0.17 | 0.000821 | 0.000774 | 0.996513 |
| 0.266 | 0.208 | 0.781 | -1.91 | -2.27 | -0.36 | 5.92E-07 | 2.94E-07 | 0.014977 |
| 2.006 | 0.052 | 0.026 | 1 | -4.25 | -5.26 | 0.002576 | 1.42E-06 | 5.23E-07 |
| 0.172 | 0.191 | 1.112 | -2.54 | -2.39 | 0.15 | 6.78E-08 | 1.18E-07 | 0.087121 |
| 0.505 | 2.58E-01 | 5.11E-01 | -0.98 | -1.95 | -0.97 | 0.1784 | 0.012802 | 0.153094 |
| 3.62E-01 | 0.033 | 9.30E-02 | -1.47E+00 | -4.9 | -3.43 | 0.01125 | 2.64E-05 | 0.000255 |
| 1.118 | 0.15 | 0.134 | 0.16 | -2.73 | -2.9 | 0.661492 | 0.119373 | 0.038873 |
| 0.085 | 4.40E-02 | 5.13E-01 | -3.56 | -4.52 | -0.96 | 0.006525 | 0.00172 | 0.369119 |
| 1.20E-02 | 0.012 | 1.01E+00 | -6.42E+00 | -6.4 | 0.02 | 8.67E-07 | 7.88E-07 | 0.831788 |
| 0.9 | 0.344 | 0.382 | -0.15 | -1.54 | -1.39 | 0.661711 | 0.064311 | 0.200755 |
| 0.736 | 1.617 | 2.197 | -0.44 | 0.69 | 1.14 | 0.005453 | 0.000424 | 2.72E-05 |
| 2.24E-01 | 2.46E-01 | 1.10E+00 | -2.16 | -2.02 | 0.13 | 8.56E-05 | 0.000157 | 0.547972 |
| 1.73E-01 | 0.275 | 1.60E+00 | -2.53E+00 | -1.86 | 0.67 | 2.15E-05 | 0.000139 | 0.020549 |
| 0.01 | 0.008 | 0.823 | -6.62 | -6.9 | -0.28 | 5.50E-05 | 6.95E-05 | 0.888715 |
| 1.201 | 0.186 | 0.155 | 0.26 | -2.43 | -2.69 | 0.013116 | 3.09E-07 | 1.39E-07 |
| 2.36E-01 | 0.171 | 7.26E-01 | -2.08E+00 | -2.55 | -0.46 | 3.93E-06 | 1.31E-06 | 0.012235 |
| 0.503 | 0.266 | 5.28E-01 | -0.99 | -1.91 | -0.92 | 0.006022 | 0.000223 | 0.012506 |
| 0.435 | 0.086 | 0.197 | -1.2 | -3.54 | -2.34 | 0.028858 | 8.64E-05 | 0.000768 |
| 0.866 | 1.887 | 2.179 | -0.21 | 0.92 | 1.12 | 0.22038 | 0.000912 | 0.00025 |
| 0.023 | 0.019 | 0.832 | -5.47 | -5.73 | -0.27 | 0.000203 | 9.91E-05 | 0.46096 |

| | | | | | | | | |
|---|---|---|---|---|---|---|---|---|
| 0.906 | 2.24E+00 | 2.48E+00 | -0.14 | 1.17 | 1.31 | 0.241863 | 8.27E-06 | 4.26E-06 |
| 1.76E-01 | 4.00E-02 | 2.24E-01 | -2.50E+00 | -4.66 | -2.16 | 0.012336 | 0.000751 | 0.045426 |
| 0.006 | 0.009 | 1.495 | -7.42 | -6.84 | 0.58 | 4.27E-06 | 4.92E-06 | 0.912768 |
| 0.097 | 0.054 | 0.552 | -3.36 | -4.22 | -0.86 | 0.037037 | 0.02733 | 0.964723 |
| 0.941 | 0.705 | 0.749 | -0.09 | -0.5 | -0.42 | 0.89131 | 0.003576 | 0.005533 |
| 0.277 | 0.298 | 1.077 | -1.85 | -1.75 | 0.11 | 0.060719 | 0.024467 | 0.742367 |
| 0.416 | 0.085 | 0.204 | -1.26 | -3.56 | -2.29 | 0.449445 | 0.00235 | 0.007919 |
| 0.029 | 0.016 | 0.549 | -5.09 | -5.96 | -0.87 | 0.004422 | 0.002304 | 0.760689 |
| 1.062 | 0.398 | 0.375 | 0.09 | -1.33 | -1.42 | 0.587259 | 7.36E-05 | 4.43E-05 |
| 0.258 | 0.279 | 1.082 | -1.96 | -1.84 | 0.11 | 0.025223 | 0.122772 | 0.453028 |
| 0.008 | 0.019 | 2.281 | -6.91 | -5.72 | 1.19 | 2.30E-06 | 9.22E-06 | 0.014317 |
| 0.074 | 0.017 | 0.225 | -3.76 | -5.92 | -2.15 | 0.002182 | 0.000606 | 0.300297 |
| 0.439 | 0.902 | 2.055 | -1.19 | -0.15 | 1.04 | 0.019193 | 0.999692 | 0.019713 |
| 1.13E+00 | 0.658 | 5.81E-01 | 1.80E-01 | -0.6 | -0.78 | 0.248778 | 0.000163 | 6.19E-05 |
| 0.012 | 0.008 | 0.678 | -6.4 | -6.96 | -0.56 | 6.19E-05 | 2.51E-05 | 0.211008 |
| 0.191 | 0.246 | 1.287 | -2.39 | -2.02 | 0.36 | 0.00032 | 0.000539 | 0.729734 |
| 0.656 | 0.256 | 0.391 | -0.61 | -1.97 | -1.36 | 0.538218 | 0.010038 | 0.035335 |
| 3.306 | 9.094 | 2.751 | 1.73 | 3.18 | 1.46 | 1.98E-05 | 9.89E-07 | 0.000138 |
| 1.646 | 0.234 | 0.142 | 0.72 | -2.1 | -2.82 | 0.000286 | 6.60E-07 | 1.36E-07 |
| 2.60E-01 | 0.007 | 0.027 | -1.94 | -7.16 | -5.22 | 2.14E-07 | 6.42E-14 | 6.51E-14 |
| 4.50E-01 | 0.207 | 0.46 | -1.15 | -2.27 | -1.12 | 3.36E-05 | 1.01E-06 | 7.71E-05 |
| 1.31E+00 | 0.744 | 0.566 | 0.39 | -0.43 | -0.82 | 0.044506 | 0.060035 | 0.002174 |
| 0.365 | 0.274 | 0.75 | -1.45 | -1.87 | -0.41 | 0.000148 | 2.50E-05 | 0.027083 |
| 0.294 | 0.047 | 0.159 | -1.76 | -4.42 | -2.66 | 9.46E-06 | 3.63E-08 | 7.72E-07 |
| 1.61E-01 | 0.23 | 1.43E+00 | -2.63E+00 | -2.12 | 0.51 | 1.99E-07 | 5.62E-07 | 0.002586 |
| 0.046 | 0.011 | 0.233 | -4.44 | -6.53 | -2.1 | 0.000306 | 8.19E-05 | 0.134644 |
| 0.901 | 0.496 | 0.55 | -0.15 | -1.01 | -0.86 | 0.433655 | 0.000711 | 0.002013 |
| 0.516 | 0.155 | 0.3 | -0.95 | -2.69 | -1.73 | 0.04777 | 9.93E-05 | 0.000685 |
| 2.043 | 16.786 | 8.218 | 1.03 | 4.07 | 3.04 | 0.42407 | 0.008008 | 0.035659 |
| 2.556 | 1.89E+00 | 7.40E-01 | 1.35 | 0.92 | -0.43 | 0.206486 | 0.624562 | 0.612644 |
| 0.844 | 0.101 | 0.119 | -0.24 | -3.31 | -3.07 | 0.177377 | 0.000575 | 0.002966 |
| 2.90E-01 | 0.195 | 6.74E-01 | -1.79E+00 | -2.36 | -0.57 | 0.000827 | 0.000149 | 0.085923 |
| 0.515 | 7.06E+00 | 1.37E+01 | -0.96 | 2.82 | 3.78 | 0.995389 | 0.173722 | 0.154624 |
| 5.63E-01 | 0.068 | 0.12 | -0.83 | -3.89 | -3.06 | 0.726133 | 0.046627 | 0.124895 |
| 1.023 | 0.721 | 0.705 | 0.03 | -0.47 | -0.5 | 0.790069 | 0.567378 | 0.269281 |
| 0.015 | 0.012 | 0.769 | -6.03 | -6.41 | -0.38 | 9.71E-07 | 6.35E-07 | 0.087993 |
| 0.897 | 0.428 | 0.477 | -0.16 | -1.22 | -1.07 | 0.356326 | 2.97E-05 | 6.01E-05 |
| 0.351 | 0.768 | 2.19 | -1.51 | -0.38 | 1.13 | 0.000368 | 0.250292 | 0.001348 |
| 4.199 | 36.039 | 8.584 | 2.07 | 5.17 | 3.1 | 0.493011 | 0.004123 | 0.013952 |
| 0.355 | 1.20E-01 | 3.38E-01 | -1.49 | -3.06 | -1.56 | 0.010067 | 0.000118 | 0.002603 |
| 9.19E-01 | 1.16E-01 | 1.26E-01 | -1.20E-01 | -3.11 | -2.99 | 0.466476 | 2.10E-06 | 2.97E-06 |
| 0.179 | 0.035 | 0.195 | -2.49 | -4.85 | -2.36 | 0.001274 | 2.01E-05 | 0.000838 |
| 0.767 | 9.402 | 12.262 | -0.38 | 3.23 | 3.62 | 0.069946 | 7.48E-07 | 4.74E-07 |
| 0.023 | 3.20E-02 | 1.38E+00 | -5.42 | -4.96 | 0.47 | 8.02E-05 | 0.000104 | 0.878209 |
| 0.812 | 2.67E-01 | 3.29E-01 | -0.3 | -1.9 | -1.6 | 0.417794 | 1.08E-05 | 1.90E-05 |
| 1.01E+00 | 0.895 | 8.91E-01 | 1.00E-02 | -0.16 | -0.17 | 0.991399 | 0.023401 | 0.020275 |
| 1.29E+00 | 0.697 | 5.39E-01 | 0.37 | -0.52 | -0.89 | 0.975701 | 0.259059 | 0.199007 |
| 0.781 | 0.904 | 1.158 | -0.36 | -0.15 | 0.21 | 0.293053 | 0.687022 | 0.710914 |
| 0.456 | 0.21 | 0.461 | -1.13 | -2.25 | -1.12 | 9.01E-05 | 1.34E-06 | 6.11E-05 |
| 0.026 | 0.031 | 1.169 | -5.25 | -5.02 | 0.23 | 0.00229 | 0.005715 | 0.606439 |
| 0.055 | 0.038 | 0.689 | -4.18 | -4.72 | -0.54 | 0.000709 | 0.000209 | 0.233588 |

| | | | | | | | | |
|---|---|---|---|---|---|---|---|---|
| 0.866 | 0.076 | 8.70E-02 | -0.21 | -3.72 | -3.52 | 0.055481 | 7.82E-08 | 1.45E-07 |
| 0.983 | 1.31 | 1.333 | -0.02 | 0.39 | 0.41 | 0.957484 | 0.002709 | 0.00212 |
| 1.579 | 0.306 | 0.194 | 0.66 | -1.71 | -2.37 | 0.90903 | 0.089552 | 0.152854 |
| 0.952 | 0.852 | 0.895 | -0.07 | -0.23 | -0.16 | 0.433925 | 0.183896 | 0.764508 |
| 0.032 | 0.093 | 2.868 | -4.95 | -3.43 | 1.52 | 1.55E-05 | 0.00016 | 0.006942 |
| 2.623 | 1.783 | 0.68 | 1.39 | 0.83 | -0.56 | 0.018995 | 0.077007 | 0.51364 |
| 1.283 | 0.85 | 0.663 | 0.36 | -0.23 | -0.59 | 0.06077 | 0.815222 | 0.028656 |
| 0.419 | 0.164 | 0.393 | -1.26 | -2.6 | -1.35 | 5.17E-05 | 9.88E-07 | 4.60E-05 |
| 1.14 | 0.017 | 0.015 | 0.19 | -5.91 | -6.1 | 0.438391 | 2.58E-06 | 1.84E-06 |
| 4.80E-01 | 0.295 | 6.15E-01 | -1.06E+00 | -1.76 | -0.7 | 0.000206 | 9.01E-06 | 0.001412 |
| 7.85E-01 | 0.05 | 0.064 | -0.35 | -4.32 | -3.97 | 0.896839 | 0.000339 | 0.000457 |
| 0.45 | 0.167 | 0.372 | -1.15 | -2.58 | -1.43 | 0.000709 | 5.18E-06 | 0.000147 |
| 0.346 | 3.80E-02 | 1.10E-01 | -1.53 | -4.71 | -3.18 | 0.077408 | 0.000901 | 0.010161 |
| 1.14E+00 | 0.775 | 6.79E-01 | 1.90E-01 | -0.37 | -0.56 | 0.181463 | 0.045099 | 0.004837 |
| 0.034 | 0.035 | 1.02 | -4.88 | -4.86 | 0.03 | 0.039902 | 0.040253 | 0.999971 |
| 0.903 | 0.506 | 0.56 | -0.15 | -0.98 | -0.84 | 0.27864 | 0.00297 | 0.016304 |
| 1.061 | 1.112 | 1.048 | 0.09 | 0.15 | 0.07 | 0.507292 | 0.525494 | 0.999342 |
| 0.371 | 0.243 | 0.656 | -1.43 | -2.04 | -0.61 | 5.36E-05 | 7.78E-06 | 0.009844 |
| 14.629 | 0.346 | 0.024 | 3.87 | -1.53 | -5.4 | 0.000477 | 0.01923 | 5.16E-05 |
| 0.891 | 0.356 | 0.4 | -0.17 | -1.49 | -1.32 | 0.091256 | 9.57E-06 | 2.83E-05 |
| 0.898 | 0.332 | 0.37 | -0.15 | -1.59 | -1.43 | 0.984194 | 9.24E-05 | 8.44E-05 |
| 0.78 | 0.127 | 1.62E-01 | -0.36 | -2.98 | -2.62 | 0.86581 | 0.000499 | 0.00072 |
| 0.106 | 0.038 | 0.36 | -3.24 | -4.71 | -1.47 | 0.01901 | 0.004596 | 0.413097 |
| 0.077 | 0.034 | 0.434 | -3.69 | -4.89 | -1.2 | 0.004201 | 0.000438 | 0.066456 |
| 1.296 | 1.143 | 8.82E-01 | 0.37 | 0.19 | -0.18 | 0.007567 | 0.122205 | 0.117796 |
| 0.814 | 0.314 | 0.386 | -0.3 | -1.67 | -1.37 | 0.537561 | 0.00212 | 0.005916 |
| 0.206 | 12.984 | 62.956 | -2.28 | 3.7 | 5.98 | 0.004882 | 0.00015 | 1.27E-05 |
| 0.749 | 0.188 | 2.52E-01 | -0.42 | -2.41 | -1.99 | 0.010091 | 6.54E-07 | 1.41E-06 |
| 0.045 | 0.024 | 5.47E-01 | -4.48 | -5.35 | -0.87 | 3.22E-06 | 1.22E-06 | 0.015899 |
| 1.224 | 0.045 | 0.037 | 0.29 | -4.46 | -4.75 | 0.839149 | 0.00026 | 0.000183 |
| 1.094 | 0.852 | 0.779 | 0.13 | -0.23 | -0.36 | 0.420071 | 0.14937 | 0.027845 |
| 2.323 | 0.092 | 0.04 | 1.22 | -3.44 | -4.66 | 0.002407 | 1.12E-05 | 1.66E-06 |
| 1.77E+00 | 0.614 | 3.47E-01 | 0.82 | -0.7 | -1.53 | 0.007725 | 0.01504 | 0.000289 |
| 2.50E+00 | 6.71E-01 | 2.69E-01 | 1.32E+00 | -0.57 | -1.9 | 0.606487 | 0.768185 | 0.279341 |
| 0.913 | 0.416 | 0.455 | -0.13 | -1.27 | -1.13 | 0.476588 | 0.000214 | 0.000475 |
| 0.239 | 3.10E-02 | 1.28E-01 | -2.06 | -5.03 | -2.97 | 0.046503 | 0.000954 | 0.017188 |
| 2.30E-01 | 0.074 | 3.20E-01 | -2.12 | -3.76 | -1.64 | 0.079603 | 0.059553 | 0.971104 |
| 1.745 | 1.21 | 0.693 | 0.8 | 0.28 | -0.53 | 0.016826 | 0.622168 | 0.052512 |
| 0.391 | 0.428 | 1.092 | -1.35 | -1.23 | 0.13 | 0.001108 | 0.001725 | 0.84943 |
| 1.633 | 4.80E-01 | 2.94E-01 | 0.71 | -1.06 | -1.77 | 0.332198 | 0.531606 | 0.080321 |
| 0.886 | 0.039 | 4.40E-02 | -0.17 | -4.69 | -4.51 | 0.896701 | 3.90E-07 | 4.21E-07 |
| 0.467 | 1.935 | 4.14 | -1.1 | 0.95 | 2.05 | 0.001691 | 0.005578 | 6.62E-05 |
| 1.10E-01 | 1.19E-01 | 1.08E+00 | -3.18E+00 | -3.08 | 0.1 | 0.008986 | 0.007388 | 0.980084 |
| 0.362 | 0.451 | 1.247 | -1.47 | -1.15 | 0.32 | 0.002486 | 0.006933 | 0.550798 |
| 0.368 | 4.816 | 13.101 | -1.44 | 2.27 | 3.71 | 0.313093 | 0.022587 | 0.004229 |
| 1.888 | 61.5 | 32.572 | 0.92 | 5.94 | 5.03 | 0.037321 | 2.97E-06 | 9.51E-06 |
| 0.635 | 0.136 | 0.214 | -0.66 | -2.88 | -2.23 | 0.051643 | 0.000187 | 0.00155 |
| 1.554 | 5.856 | 3.768 | 0.64 | 2.55 | 1.91 | 0.658798 | 0.024998 | 0.074801 |
| 0.381 | 9.20E-02 | 2.41E-01 | -1.39 | -3.45 | -2.05 | 0.698016 | 0.05839 | 0.167878 |
| 0.215 | 0.228 | 1.063 | -2.22 | -2.13 | 0.09 | 0.00275 | 0.003038 | 0.99315 |
| 0.326 | 0.012 | 0.037 | -1.61 | -6.37 | -4.76 | 0.002246 | 1.22E-06 | 6.35E-06 |

| | | | | | | | | |
|---|---|---|---|---|---|---|---|---|
| 2.60E-02 | 0.031 | 1.17E+00 | -5.25E+00 | -5.02 | 0.23 | 0.002018 | 0.004919 | 0.610391 |
| 1.65 | 16.911 | 10.249 | 0.72 | 4.08 | 3.36 | 0.805357 | 0.0013 | 0.002204 |
| 0.325 | 0.177 | 0.543 | -1.62 | -2.5 | -0.88 | 0.103022 | 0.004972 | 0.081273 |
| 0.667 | 1.316 | 1.974 | -0.58 | 0.4 | 0.98 | 0.028616 | 0.072053 | 0.001801 |
| 1.092 | 4.01E-01 | 3.67E-01 | 0.13 | -1.32 | -1.45 | 0.267272 | 0.001684 | 0.000457 |
| 1.124 | 0.332 | 2.95E-01 | 0.17 | -1.59 | -1.76 | 0.933684 | 0.133759 | 0.085108 |
| 0.712 | 2.613 | 3.67 | -0.49 | 1.39 | 1.88 | 0.992181 | 0.07957 | 0.068459 |
| 8.35E-01 | 0.457 | 5.47E-01 | -2.60E-01 | -1.13 | -0.87 | 0.859829 | 0.111995 | 0.2196 |
| 0.954 | 0.381 | 0.399 | -0.07 | -1.39 | -1.32 | 0.969208 | 0.010686 | 0.013809 |
| 5.592 | 3.08 | 0.551 | 2.48 | 1.62 | -0.86 | 7.04E-07 | 5.50E-06 | 0.000275 |
| 0.51 | 0.233 | 0.457 | -0.97 | -2.1 | -1.13 | 2.14E-05 | 3.76E-07 | 8.49E-06 |
| 1.447 | 0.873 | 0.603 | 0.53 | -0.2 | -0.73 | 0.770215 | 0.939546 | 0.580115 |
| 9.83E-01 | 9.25E+01 | 9.41E+01 | -0.02 | 6.53 | 6.56 | 0.682443 | 7.50E-10 | 5.73E-10 |
| 2.07E+00 | 0.666 | 3.22E-01 | 1.05E+00 | -0.59 | -1.63 | 0.416952 | 0.22997 | 0.041535 |
| 0.753 | 1.012 | 1.343 | -0.41 | 0.02 | 0.43 | 0.017563 | 0.969278 | 0.0135 |
| 0.841 | 0.899 | 1.069 | -0.25 | -0.15 | 0.1 | 0.91859 | 0.954069 | 0.778548 |
| 0.672 | 0.418 | 0.622 | -0.57 | -1.26 | -0.68 | 0.014241 | 0.000328 | 0.009931 |
| 0.738 | 5.48E-01 | 7.42E-01 | -0.44 | -0.87 | -0.43 | 0.062784 | 0.000463 | 0.004736 |
| 9.60E-01 | 0.747 | 7.78E-01 | -0.06 | -0.42 | -0.36 | 0.604754 | 0.004284 | 0.011663 |
| 0.869 | 0.72 | 0.829 | -0.2 | -0.47 | -0.27 | 0.928119 | 0.896801 | 0.706043 |
| 1.162 | 0.035 | 0.03 | 0.22 | -4.84 | -5.06 | 0.41758 | 1.52E-05 | 8.79E-06 |
| 0.07 | 8.30E-02 | 1.20E+00 | -3.84 | -3.58 | 0.26 | 0.057882 | 0.026074 | 0.793333 |
| 1.38E+00 | 0.798 | 5.78E-01 | 4.60E-01 | -0.33 | -0.79 | 0.35275 | 0.400892 | 0.061484 |
| 0.704 | 0.413 | 5.87E-01 | -0.51 | -1.28 | -0.77 | 0.000779 | 4.06E-06 | 9.28E-05 |
| 0.037 | 0.022 | 0.601 | -4.77 | -5.5 | -0.73 | 0.002983 | 0.000987 | 0.42763 |
| 0.178 | 0.035 | 0.195 | -2.49 | -4.85 | -2.35 | 0.067533 | 0.006669 | 0.182107 |
| 4.36E-01 | 0.198 | 0.455 | -1.20E+00 | -2.34 | -1.14 | 0.008259 | 0.000365 | 0.020879 |
| 7.21 | 0.115 | 0.016 | 2.85 | -3.12 | -5.97 | 7.26E-06 | 3.50E-06 | 1.44E-07 |
| 0.419 | 0.165 | 0.395 | -1.26 | -2.6 | -1.34 | 5.18E-05 | 9.93E-07 | 4.65E-05 |
| 1.367 | 4.44E+00 | 3.25E+00 | 0.45 | 2.15 | 1.7 | 0.159889 | 8.80E-05 | 0.000306 |
| 8.27E-01 | 3.758 | 4.54E+00 | -0.27 | 1.91 | 2.18 | 0.323218 | 0.116149 | 0.017122 |
| 0.897 | 0.024 | 0.026 | -0.16 | -5.4 | -5.24 | 0.716377 | 2.13E-09 | 2.68E-09 |
| 0.153 | 9.00E-03 | 6.20E-02 | -2.71 | -6.73 | -4.02 | 1.49E-11 | 6.42E-14 | 6.42E-14 |
| 0.9 | 1.18E+00 | 1.31E+00 | -0.15 | 0.24 | 0.39 | 0.130484 | 0.026174 | 0.002516 |
| 0.476 | 0.271 | 0.569 | -1.07 | -1.88 | -0.81 | 0.13065 | 0.13591 | 0.999473 |
| 0.084 | 1.68E-01 | 2.00E+00 | -3.58 | -2.58 | 1 | 0.005344 | 0.012561 | 0.696625 |
| 0.445 | 0.18 | 0.405 | -1.17 | -2.47 | -1.3 | 0.057999 | 0.010944 | 0.379811 |
| 2.791 | 22.949 | 8.221 | 1.48 | 4.52 | 3.04 | 0.596561 | 0.005013 | 0.014165 |
| 1.03 | 0.726 | 0.704 | 0.04 | -0.46 | -0.51 | 0.98005 | 0.006935 | 0.008423 |
| 0.404 | 0.227 | 0.56 | -1.31 | -2.14 | -0.84 | 3.82E-05 | 2.74E-06 | 0.00117 |
| 1.035 | 1.265 | 1.223 | 0.05 | 0.34 | 0.29 | 0.404295 | 0.012995 | 0.066143 |
| 9.86E-01 | 0.275 | 2.79E-01 | -2.00E-02 | -1.86 | -1.84 | 0.972446 | 9.30E-05 | 8.26E-05 |
| 0.184 | 1.285 | 6.968 | -2.44 | 0.36 | 2.8 | 0.091176 | 0.921055 | 0.05592 |
| 0.826 | 0.305 | 0.369 | -0.28 | -1.71 | -1.44 | 0.625185 | 0.000465 | 0.000927 |
| 0.368 | 0.183 | 0.497 | -1.44 | -2.45 | -1.01 | 0.000108 | 4.94E-06 | 0.000976 |
| 0.875 | 21.224 | 24.25 | -0.19 | 4.41 | 4.6 | 0.830028 | 0.001658 | 0.001036 |
| 1.13E+00 | 0.61 | 5.41E-01 | 1.70E-01 | -0.71 | -0.89 | 0.749225 | 0.129311 | 0.326347 |
| 1.844 | 3.479 | 1.887 | 0.88 | 1.8 | 0.92 | 3.89E-05 | 7.36E-07 | 2.64E-05 |
| 0.87 | 0.015 | 0.017 | -0.2 | -6.09 | -5.89 | 0.15975 | 4.24E-09 | 7.63E-09 |
| 4.284 | 0.463 | 0.108 | 2.1 | -1.11 | -3.21 | 0.050555 | 0.99891 | 0.047863 |
| 0.086 | 0.045 | 0.525 | -3.54 | -4.47 | -0.93 | 0.002923 | 0.000926 | 0.399962 |

| | | | | | | | | |
|---|---|---|---|---|---|---|---|---|
| 1.19E+00 | 0.016 | 1.30E-02 | 0.25 | -5.96 | -6.22 | 0.180314 | 9.02E-10 | 4.75E-10 |
| 0.423 | 0.116 | 0.275 | -1.24 | -3.1 | -1.86 | 0.002619 | 3.14E-05 | 0.000936 |
| 0.915 | 3.10E-01 | 3.39E-01 | -0.13 | -1.69 | -1.56 | 0.995934 | 0.00091 | 0.000972 |
| 1.687 | 1.024 | 0.607 | 0.75 | 0.03 | -0.72 | 0.000534 | 0.748522 | 0.000915 |
| 0.959 | 1.037 | 1.082 | -0.06 | 0.05 | 0.11 | 0.981367 | 0.733226 | 0.832607 |
| 4.68E-01 | 0.215 | 0.459 | -1.10E+00 | -2.22 | -1.12 | 0.000526 | 7.44E-06 | 0.000346 |
| 1.90E-01 | 0.195 | 1.027 | -2.4 | -2.36 | 0.04 | 0.162235 | 0.10797 | 0.945886 |
| 0.89 | 0.386 | 0.433 | -0.17 | -1.38 | -1.21 | 0.456283 | 4.63E-05 | 8.79E-05 |
| 0.06 | 0.046 | 0.776 | -4.07 | -4.43 | -0.37 | 0.000115 | 5.70E-05 | 0.418562 |
| 0.766 | 2.57E+01 | 3.35E+01 | -0.38 | 4.68 | 5.07 | 0.302004 | 6.58E-07 | 5.04E-07 |
| 2.30E-02 | 0.021 | 8.97E-01 | -5.42 | -5.58 | -0.16 | 8.84E-14 | 6.93E-14 | 0.031915 |
| 0.602 | 3.4 | 5.645 | -0.73 | 1.77 | 2.5 | 0.4287 | 0.247927 | 0.04611 |
| 0.435 | 1.07E-01 | 2.45E-01 | -1.2 | -3.23 | -2.03 | 0.024184 | 0.000766 | 0.022946 |
| 0.541 | 0.389 | 0.719 | -0.89 | -1.36 | -0.48 | 0.179906 | 0.192486 | 0.998427 |
| 0.882 | 1.11E-01 | 1.26E-01 | -0.18 | -3.17 | -2.99 | 0.325134 | 3.68E-07 | 4.80E-07 |
| 9.50E-01 | 3.577 | 3.77E+00 | -7.00E-02 | 1.84 | 1.91 | 0.705825 | 0.006862 | 0.016288 |
| 0.333 | 0.174 | 0.521 | -1.59 | -2.53 | -0.94 | 0.000682 | 4.75E-05 | 0.009239 |
| 0.456 | 0.21 | 0.461 | -1.13 | -2.25 | -1.12 | 9.01E-05 | 1.34E-06 | 6.11E-05 |
| 1.253 | 7.02E-01 | 5.60E-01 | 0.33 | -0.51 | -0.84 | 0.001114 | 0.000168 | 7.02E-06 |
| 6.80E-02 | 0.103 | 1.51E+00 | -3.88E+00 | -3.28 | 0.59 | 0.005572 | 0.008369 | 0.914181 |
| 1.083 | 14.787 | 13.657 | 0.11 | 3.89 | 3.77 | 0.502664 | 0.002012 | 0.000787 |
| 1.032 | 0.124 | 0.12 | 0.04 | -3.01 | -3.05 | 0.760712 | 2.30E-06 | 1.92E-06 |
| 0.374 | 0.182 | 0.485 | -1.42 | -2.46 | -1.04 | 0.004739 | 0.000239 | 0.018659 |
| 2.73E-01 | 0.054 | 1.99E-01 | -1.87E+00 | -4.21 | -2.33 | 0.061561 | 0.032566 | 0.863836 |
| 2.555 | 7.788 | 3.048 | 1.35 | 2.96 | 1.61 | 0.999183 | 0.097486 | 0.102384 |
| 0.094 | 0.068 | 0.722 | -3.41 | -3.88 | -0.47 | 0.009496 | 0.003401 | 0.574436 |
| 0.631 | 0.408 | 0.648 | -0.67 | -1.29 | -0.63 | 0.00923 | 0.000194 | 0.006361 |
| 1.79 | 0.962 | 0.537 | 0.84 | -0.06 | -0.9 | 0.980234 | 0.546989 | 0.448705 |
| 0.466 | 17.697 | 37.954 | -1.1 | 4.15 | 5.25 | 0.903635 | 0.001271 | 0.00091 |
| 1.58E+00 | 5.30E-02 | 3.40E-02 | 0.66 | -4.23 | -4.89 | 0.003516 | 2.69E-08 | 6.50E-09 |
| 5.51E-01 | 3.34E-01 | 6.07E-01 | -0.86 | -1.58 | -0.72 | 0.360283 | 0.037857 | 0.245253 |
| 0.927 | 1.231 | 1.328 | -0.11 | 0.3 | 0.41 | 0.576998 | 0.691768 | 0.221085 |
| 0.024 | 0.023 | 9.20E-01 | -5.35 | -5.47 | -0.12 | 9.30E-12 | 4.11E-12 | 0.156974 |
| 8.48E-01 | 1.91E-01 | 2.26E-01 | -0.24 | -2.38 | -2.15 | 0.786795 | 3.47E-05 | 4.76E-05 |
| 0.737 | 0.362 | 4.91E-01 | -0.44 | -1.47 | -1.03 | 0.218631 | 0.00149 | 0.008471 |
| 1.45E+00 | 3.16E-01 | 2.17E-01 | 5.40E-01 | -1.66 | -2.2 | 0.019784 | 2.80E-05 | 5.87E-06 |
| 1.391 | 0.27 | 0.194 | 0.48 | -1.89 | -2.36 | 0.152776 | 0.061283 | 0.005393 |
| 0.33 | 0.521 | 1.58 | -1.6 | -0.94 | 0.66 | 0.128165 | 0.207802 | 0.924261 |
| 0.093 | 0.128 | 1.37E+00 | -3.42 | -2.96 | 0.46 | 0.000101 | 0.000127 | 0.906907 |
| 0.775 | 0.274 | 0.354 | -0.37 | -1.87 | -1.5 | 0.778766 | 0.002067 | 0.003809 |
| 1.00E+00 | 0.233 | 2.33E-01 | 0 | -2.1 | -2.1 | 0.999634 | 3.13E-05 | 3.17E-05 |
| 1.008 | 3.40E-01 | 3.37E-01 | 0.01 | -1.56 | -1.57 | 0.961188 | 8.31E-05 | 7.23E-05 |
| 0.601 | 3.78E+00 | 6.29E+00 | -0.73 | 1.92 | 2.65 | 0.008827 | 0.000197 | 1.99E-05 |
| 2.15E-01 | 9.80E-02 | 4.54E-01 | -2.22 | -3.36 | -1.14 | 5.97E-06 | 6.48E-07 | 0.000175 |
| 2.346 | 9.79E-01 | 4.17E-01 | 1.23 | -0.03 | -1.26 | 0.029472 | 0.999775 | 0.030184 |
| 0.78 | 0.141 | 0.181 | -0.36 | -2.83 | -2.47 | 0.134762 | 5.93E-06 | 1.45E-05 |
| 0.408 | 0.229 | 0.561 | -1.29 | -2.13 | -0.83 | 0.058331 | 0.002589 | 0.058689 |
| 0.915 | 0.239 | 0.262 | -0.13 | -2.06 | -1.93 | 0.999008 | 0.005338 | 0.005563 |
| 0.035 | 0.04 | 1.132 | -4.82 | -4.65 | 0.18 | 0.000831 | 0.000588 | 0.885072 |
| 1.40E+00 | 0.899 | 6.42E-01 | 4.80E-01 | -0.15 | -0.64 | 0.088087 | 0.514673 | 0.02148 |
| 1.177 | 0.458 | 0.389 | 0.23 | -1.13 | -1.36 | 0.96824 | 0.036818 | 0.027602 |

| | | | | | | | | |
|---|---|---|---|---|---|---|---|---|
| 0.719 | 0.51 | 0.709 | -0.48 | -0.97 | -0.5 | 0.306501 | 0.011583 | 0.077983 |
| 0.464 | 12.822 | 27.659 | -1.11 | 3.68 | 4.79 | 0.201827 | 0.000642 | 0.000179 |
| 0.173 | 0.12 | 0.694 | -2.53 | -3.06 | -0.53 | 0.002564 | 0.000862 | 0.42366 |
| 4.23E-01 | 0.022 | 5.20E-02 | -1.24E+00 | -5.5 | -4.26 | 0.02067 | 7.80E-06 | 3.89E-05 |
| 0.695 | 16.217 | 23.319 | -0.52 | 4.02 | 4.54 | 0.551912 | 0.000519 | 0.000253 |
| 1.027 | 0.742 | 0.722 | 0.04 | -0.43 | -0.47 | 0.880131 | 0.645796 | 0.901144 |
| 0.683 | 8.03E-01 | 1.18E+00 | -0.55 | -0.32 | 0.23 | 0.15287 | 0.493073 | 0.615279 |
| 4.40E-02 | 2.30E-02 | 0.524 | -4.51E+00 | -5.44 | -0.93 | 0.007193 | 0.007658 | 0.997896 |
| 0.145 | 0.015 | 0.105 | -2.79 | -6.04 | -3.25 | 3.13E-06 | 6.90E-08 | 2.55E-06 |
| 0.508 | 0.477 | 0.94 | -0.98 | -1.07 | -0.09 | 0.108573 | 0.538184 | 0.429635 |
| 0.06 | 0.032 | 5.29E-01 | -4.06 | -4.97 | -0.92 | 0.000678 | 0.000195 | 0.217703 |
| 1.328 | 2.86E-01 | 2.15E-01 | 0.41 | -1.81 | -2.22 | 0.783845 | 0.013172 | 0.006508 |
| 7.21E-01 | 9.976 | 1.38E+01 | -0.47 | 3.32 | 3.79 | 0.14539 | 5.19E-05 | 1.89E-05 |
| 0.573 | 1.22E-01 | 2.14E-01 | -0.8 | -3.03 | -2.23 | 0.1923 | 0.000529 | 0.002525 |
| 1.222 | 2.206 | 1.805 | 0.29 | 1.14 | 0.85 | 0.208119 | 0.00625 | 0.054553 |
| 6.33E-01 | 3.98E-01 | 6.29E-01 | -6.60E-01 | -1.33 | -0.67 | 0.020727 | 0.000228 | 0.004028 |
| 0.763 | 6.68E-01 | 8.76E-01 | -0.39 | -0.58 | -0.19 | 0.788691 | 0.098161 | 0.229713 |
| 8.40E-02 | 0.047 | 5.59E-01 | -3.57 | -4.41 | -0.84 | 0.000202 | 4.89E-05 | 0.084828 |
| 7.92E-01 | 3.43E-01 | 4.33E-01 | -3.40E-01 | -1.54 | -1.21 | 0.000496 | 1.61E-07 | 6.65E-07 |
| 0.336 | 0.103 | 3.06E-01 | -1.57 | -3.28 | -1.71 | 0.036163 | 0.002038 | 0.067672 |
| 9.08E-01 | 0.416 | 0.458 | -0.14 | -1.26 | -1.13 | 0.917063 | 0.019159 | 0.030401 |
| 2.71E-01 | 1.33E-01 | 4.91E-01 | -1.89 | -2.91 | -1.03 | 0.017639 | 0.006674 | 0.649459 |
| 1.13E+00 | 0.937 | 8.27E-01 | 1.80E-01 | -0.09 | -0.27 | 0.311634 | 0.993117 | 0.272181 |
| 0.488 | 0.039 | 0.081 | -1.03 | -4.67 | -3.63 | 0.015273 | 5.35E-06 | 2.77E-05 |
| 0.442 | 6.57E+00 | 1.49E+01 | -1.18 | 2.72 | 3.89 | 0.354716 | 0.110208 | 0.017865 |
| 0.781 | 0.026 | 0.033 | -0.36 | -5.27 | -4.92 | 0.672341 | 2.78E-06 | 3.64E-06 |
| 0.108 | 0.11 | 1.013 | -3.21 | -3.19 | 0.02 | 0.003182 | 0.001444 | 0.646315 |
| 0.836 | 2.671 | 3.196 | -0.26 | 1.42 | 1.68 | 0.968644 | 0.029406 | 0.022216 |
| 0.086 | 0.109 | 1.271 | -3.55 | -3.2 | 0.35 | 0.037081 | 0.175413 | 0.475777 |
| 4.20E-01 | 2.70E-02 | 0.064 | -1.25E+00 | -5.21 | -3.96 | 0.000117 | 1.75E-08 | 1.76E-07 |
| 2.50E-02 | 0.013 | 5.37E-01 | -5.35E+00 | -6.24 | -0.9 | 1.27E-05 | 9.61E-06 | 0.775357 |
| 0.512 | 0.556 | 1.085 | -0.97 | -0.85 | 0.12 | 0.049527 | 0.07968 | 0.924131 |
| 0.935 | 0.749 | 0.801 | -0.1 | -0.42 | -0.32 | 0.923605 | 0.525084 | 0.738577 |
| 1.114 | 0.686 | 0.616 | 0.16 | -0.54 | -0.7 | 0.524708 | 0.05109 | 0.013508 |
| 4.13E-01 | 3.12E-01 | 7.55E-01 | -1.27 | -1.68 | -0.4 | 0.005115 | 0.000718 | 0.1248 |
| 2.365 | 2.134 | 0.902 | 1.24 | 1.09 | -0.15 | 0.273739 | 0.322863 | 0.989719 |
| 0.589 | 0.874 | 1.483 | -0.76 | -0.19 | 0.57 | 0.994525 | 0.732574 | 0.676112 |
| 0.171 | 1.20E-01 | 7.00E-01 | -2.55 | -3.06 | -0.51 | 1.54E-06 | 6.68E-07 | 0.007362 |
| 0.877 | 0.975 | 1.111 | -0.19 | -0.04 | 0.15 | 0.998301 | 0.860821 | 0.886379 |
| 1.076 | 15.212 | 14.132 | 0.11 | 3.93 | 3.82 | 0.545938 | 1.45E-07 | 1.80E-07 |
| 0.624 | 1.089 | 1.745 | -0.68 | 0.12 | 0.8 | 0.714395 | 0.696271 | 0.999435 |
| 7.07E-01 | 1.166 | 1.65E+00 | -0.5 | 0.22 | 0.72 | 0.860612 | 0.439295 | 0.725737 |
| 0.938 | 5.60E+01 | 5.97E+01 | -0.09 | 5.81 | 5.9 | 0.998169 | 5.99E-05 | 5.82E-05 |
| 0.63 | 22.327 | 3.54E+01 | -0.67 | 4.48 | 5.15 | 0.757622 | 0.000295 | 0.000189 |
| 0.012 | 0.002 | 0.205 | -6.41 | -8.7 | -2.29 | 2.89E-05 | 4.60E-06 | 0.008779 |
| 0.025 | 0.013 | 0.53 | -5.32 | -6.24 | -0.92 | 0.001135 | 0.000245 | 0.147892 |
| 0.929 | 0.687 | 0.74 | -0.11 | -0.54 | -0.44 | 0.36949 | 0.000466 | 0.001381 |
| 1.3 | 11.881 | 9.141 | 0.38 | 3.57 | 3.19 | 0.719981 | 2.82E-05 | 4.05E-05 |
| 7.90E-02 | 0.036 | 4.56E-01 | -3.66E+00 | -4.79 | -1.13 | 1.95E-06 | 6.93E-07 | 0.003347 |
| 0.359 | 2.74E-01 | 7.63E-01 | -1.48 | -1.87 | -0.39 | 0.704566 | 0.129712 | 0.359275 |
| 1.102 | 0.554 | 0.503 | 0.14 | -0.85 | -0.99 | 0.450899 | 0.001321 | 0.000507 |

| | | | | | | | | |
|---|---|---|---|---|---|---|---|---|
| 6.73E-01 | 3.61E-01 | 0.536 | -0.57 | -1.47 | -0.9 | 0.001701 | 2.83E-06 | 3.24E-05 |
| 3.00E-03 | 0.001 | 2.17E-01 | -8.24 | -10.44 | -2.2 | 1.57E-05 | 3.91E-06 | 0.024291 |
| 1.64E-01 | 0.463 | 2.82 | -2.61 | -1.11 | 1.5 | 0.011836 | 0.186401 | 0.13346 |
| 1.49E+00 | 0.807 | 5.44E-01 | 5.70E-01 | -0.31 | -0.88 | 0.342957 | 0.685151 | 0.118004 |
| 1.139 | 4.51E-01 | 3.95E-01 | 0.19 | -1.15 | -1.34 | 0.457601 | 3.88E-05 | 2.19E-05 |
| 0.257 | 0.424 | 1.65E+00 | -1.96 | -1.24 | 0.72 | 0.013532 | 0.047312 | 0.559227 |
| 0.862 | 6.45E-01 | 7.48E-01 | -0.21 | -0.63 | -0.42 | 0.662047 | 0.021412 | 0.062735 |
| 83.854 | 0.775 | 9.00E-03 | 6.39 | -0.37 | -6.76 | 6.52E-10 | 0.024957 | 2.02E-10 |
| 0.017 | 0.016 | 0.937 | -5.85 | -5.94 | -0.09 | 4.59E-06 | 4.22E-06 | 0.966974 |
| 5.73E+00 | 0.726 | 1.27E-01 | 2.52E+00 | -0.46 | -2.98 | 0.000299 | 0.188933 | 9.26E-05 |
| 0.374 | 0.062 | 1.66E-01 | -1.42 | -4.01 | -2.59 | 0.192697 | 0.044134 | 0.511507 |
| 7.00E-03 | 0.046 | 6.245 | -7.08E+00 | -4.44 | 2.64 | 0.00112 | 0.008317 | 0.141908 |
| 0.153 | 0.031 | 0.202 | -2.71 | -5.02 | -2.31 | 2.50E-07 | 2.51E-10 | 6.32E-07 |
| 7.76E-01 | 0.012 | 1.60E-02 | -3.70E-01 | -6.34 | -5.98 | 0.552507 | 3.91E-06 | 5.71E-06 |
| 0.743 | 0.923 | 1.242 | -0.43 | -0.12 | 0.31 | 0.602361 | 0.595311 | 0.999912 |
| 0.596 | 0.633 | 1.061 | -0.75 | -0.66 | 0.08 | 0.033298 | 0.166511 | 0.451538 |
| 1.146 | 0.478 | 0.418 | 0.2 | -1.06 | -1.26 | 0.451199 | 0.001255 | 0.000485 |
| 0.917 | 5.99E-01 | 6.53E-01 | -0.12 | -0.74 | -0.62 | 0.466476 | 0.000133 | 0.000282 |
| 0.937 | 5.78E-01 | 6.17E-01 | -0.09 | -0.79 | -0.7 | 0.739043 | 0.006072 | 0.013336 |
| 1.013 | 9.56E-01 | 9.44E-01 | 0.02 | -0.07 | -0.08 | 0.998857 | 0.811325 | 0.787168 |
| 1.126 | 0.963 | 0.855 | 0.17 | -0.05 | -0.23 | 0.977291 | 0.927579 | 0.837346 |
| 0.14 | 1.73E-01 | 1.23E+00 | -2.83 | -2.53 | 0.3 | 0.010521 | 0.054432 | 0.385886 |
| 0.763 | 0.312 | 0.408 | -0.39 | -1.68 | -1.29 | 0.220653 | 0.013713 | 0.133833 |
| 0.294 | 0.137 | 4.66E-01 | -1.77 | -2.87 | -1.1 | 0.083105 | 0.034439 | 0.7649 |
| 0.362 | 3.41E-01 | 9.42E-01 | -1.47 | -1.55 | -0.09 | 0.002345 | 0.001135 | 0.670001 |
| 0.78 | 0.531 | 0.68 | -0.36 | -0.91 | -0.56 | 0.157338 | 0.000392 | 0.001977 |
| 2.77E-01 | 1.81E-01 | 6.52E-01 | -1.85E+00 | -2.47 | -0.62 | 1.79E-05 | 2.46E-06 | 0.003584 |
| 0.835 | 3.56E-01 | 4.27E-01 | -0.26 | -1.49 | -1.23 | 0.388636 | 2.74E-05 | 5.31E-05 |
| 9.10E-02 | 1.70E-02 | 1.86E-01 | -3.46E+00 | -5.89 | -2.43 | 0.004864 | 0.000995 | 0.231583 |
| 1.32E+00 | 4.72 | 3.57E+00 | 0.4 | 2.24 | 1.84 | 0.161142 | 0.001582 | 0.011785 |
| 0.881 | 51.959 | 59.01 | -0.18 | 5.7 | 5.88 | 0.148284 | 5.95E-10 | 2.94E-10 |
| 1.53E+00 | 5.60E+01 | 36.558 | 6.10E-01 | 5.81 | 5.19 | 0.152369 | 3.65E-07 | 5.28E-07 |
| 1.028 | 13.293 | 12.935 | 0.04 | 3.73 | 3.69 | 0.675159 | 7.54E-05 | 4.88E-05 |
| 1.01E-01 | 2.70E-02 | 0.269 | -3.30E+00 | -5.2 | -1.89 | 0.000296 | 3.07E-05 | 0.012597 |
| 0.324 | 2.09E-01 | 6.46E-01 | -1.63 | -2.26 | -0.63 | 3.94E-05 | 5.63E-06 | 0.007812 |
| 0.405 | 3.84E-01 | 9.49E-01 | -1.3 | -1.38 | -0.08 | 0.239912 | 0.603207 | 0.702275 |
| 1.794 | 0.799 | 0.445 | 0.84 | -0.32 | -1.17 | 0.000159 | 0.010853 | 1.85E-05 |
| 1.722 | 0.623 | 3.62E-01 | 0.78 | -0.68 | -1.47 | 0.070227 | 0.063163 | 0.003114 |
| 0.862 | 0.224 | 0.26 | -0.21 | -2.16 | -1.94 | 0.768826 | 1.38E-05 | 1.86E-05 |
| 0.025 | 0.141 | 5.67E+00 | -5.33 | -2.82 | 2.5 | 0.001481 | 0.013445 | 0.124379 |
| 0.822 | 1.12E-01 | 1.37E-01 | -0.28 | -3.16 | -2.87 | 0.685965 | 0.205286 | 0.550204 |
| 1.762 | 0.74 | 0.42 | 0.82 | -0.44 | -1.25 | 0.288392 | 0.983728 | 0.233257 |
| 0.151 | 1.06E+01 | 7.02E+01 | -2.73 | 3.4 | 6.13 | 0.002159 | 0.000602 | 2.18E-05 |
| 0.014 | 0.014 | 0.99 | -6.14 | -6.15 | -0.02 | 6.42E-14 | 6.42E-14 | 0.630224 |
| 1.4 | 1.59E+00 | 1.14E+00 | 0.49 | 0.67 | 0.19 | 0.010644 | 0.004734 | 0.713087 |
| 1.009 | 1.203 | 1.19E+00 | 0.01 | 0.27 | 0.25 | 0.998438 | 0.147979 | 0.158367 |
| 1.75E+00 | 11.226 | 6.40E+00 | 0.81 | 3.49 | 2.68 | 0.015345 | 2.69E-05 | 0.00022 |
| 1.98 | 0.608 | 0.307 | 0.99 | -0.72 | -1.7 | 0.000351 | 0.001162 | 1.14E-05 |
| 0.837 | 23.873 | 28.514 | -0.26 | 4.58 | 4.83 | 0.971414 | 0.003964 | 0.003209 |
| 0.026 | 2.20E-02 | 8.61E-01 | -5.27 | -5.49 | -0.22 | 0.000591 | 0.000706 | 0.966997 |
| 0.869 | 3.64E-01 | 4.18E-01 | -0.2 | -1.46 | -1.26 | 0.263622 | 3.53E-05 | 8.30E-05 |

| | | | | | | | | |
|---|---|---|---|---|---|---|---|---|
| 1.25E+00 | 0.352 | 0.281 | 0.33 | -1.5 | -1.83 | 0.420281 | 0.000812 | 0.000319 |
| 1.07E+00 | 0.324 | 3.04E-01 | 9.00E-02 | -1.63 | -1.72 | 0.916525 | 0.001773 | 0.00128 |
| 0.898 | 2.231 | 2.483 | -0.15 | 1.16 | 1.31 | 0.730625 | 0.000751 | 0.000435 |
| 0.148 | 0.014 | 0.096 | -2.76 | -6.14 | -3.38 | 7.37E-06 | 1.01E-07 | 2.10E-06 |
| 0.743 | 0.384 | 0.517 | -0.43 | -1.38 | -0.95 | 0.007677 | 2.09E-05 | 0.000231 |
| 0.38 | 0.509 | 1.339 | -1.39 | -0.97 | 0.42 | 0.272473 | 0.159376 | 0.905424 |
| 1.05 | 0.56 | 0.533 | 0.07 | -0.84 | -0.91 | 0.882035 | 0.01115 | 0.006781 |
| 7.777 | 0.034 | 0.004 | 2.96 | -4.87 | -7.83 | 0.001811 | 0.000172 | 8.98E-06 |
| 0.816 | 0.148 | 0.182 | -0.29 | -2.75 | -2.46 | 0.257271 | 2.24E-06 | 3.77E-06 |
| 0.73 | 4.64E-01 | 6.36E-01 | -0.45 | -1.11 | -0.65 | 0.068944 | 0.000159 | 0.00103 |
| 6.19E-01 | 0.411 | 6.64E-01 | -6.90E-01 | -1.28 | -0.59 | 0.461765 | 0.009617 | 0.040188 |
| 7.14E-01 | 0.488 | 0.684 | -0.49 | -1.03 | -0.55 | 0.014268 | 0.000126 | 0.00219 |
| 6.66E-01 | 0.309 | 0.464 | -0.59 | -1.69 | -1.11 | 0.002936 | 1.08E-05 | 0.000163 |
| 0.798 | 0.501 | 0.627 | -0.33 | -1 | -0.67 | 0.108962 | 0.000984 | 0.008664 |
| 0.921 | 6.43E-01 | 6.99E-01 | -0.12 | -0.64 | -0.52 | 0.565234 | 0.007508 | 0.023921 |
| 0.954 | 6.32E-01 | 6.62E-01 | -0.07 | -0.66 | -0.59 | 0.516475 | 4.59E-05 | 8.19E-05 |
| 0.949 | 0.896 | 9.44E-01 | -0.08 | -0.16 | -0.08 | 0.654871 | 0.105366 | 0.32891 |
| 1.01E+00 | 0.797 | 7.86E-01 | 0.02 | -0.33 | -0.35 | 0.8747 | 0.045942 | 0.025336 |
| 0.597 | 0.694 | 1.163 | -0.74 | -0.53 | 0.22 | 0.940836 | 0.545813 | 0.733398 |
| 26.425 | 0.738 | 0.028 | 4.72 | -0.44 | -5.16 | 9.54E-06 | 0.507705 | 6.09E-06 |
| 24.39 | 1.804 | 0.074 | 4.61 | 0.85 | -3.76 | 0.003209 | 0.596408 | 0.008532 |
| 4.19E-01 | 0.2 | 4.77E-01 | -1.25E+00 | -2.32 | -1.07 | 0.000235 | 4.28E-06 | 0.000285 |
| 0.67 | 1.11E-01 | 1.66E-01 | -0.58 | -3.17 | -2.59 | 0.76669 | 0.001249 | 0.002246 |
| 0.998 | 0.733 | 0.735 | 0 | -0.45 | -0.45 | 0.979963 | 0.50057 | 0.604563 |
| 0.339 | 0.226 | 0.667 | -1.56 | -2.15 | -0.58 | 1.00E-05 | 1.40E-06 | 0.001553 |
| 2.70E-01 | 5.70E-02 | 2.11E-01 | -1.89E+00 | -4.14 | -2.25 | 0.106797 | 0.045482 | 0.782682 |
| 1.82E+00 | 3.353 | 1.85E+00 | 0.86 | 1.75 | 0.88 | 0.001037 | 2.10E-05 | 0.001102 |
| 0.541 | 0.089 | 0.165 | -0.89 | -3.48 | -2.6 | 0.109458 | 0.000183 | 0.000935 |
| 8.08E-01 | 0.389 | 0.481 | -0.31 | -1.36 | -1.06 | 0.040492 | 1.76E-05 | 7.91E-05 |
| 7.10E-02 | 0.085 | 1.199 | -3.81E+00 | -3.55 | 0.26 | 0.000193 | 0.000495 | 0.366883 |
| 0.716 | 0.239 | 0.333 | -0.48 | -2.07 | -1.58 | 0.000214 | 4.71E-08 | 3.13E-07 |
| 1.41 | 0.258 | 0.183 | 0.5 | -1.95 | -2.45 | 0.520972 | 0.004749 | 0.001707 |
| 3.81E+00 | 0.092 | 2.40E-02 | 1.93E+00 | -3.44 | -5.37 | 0.043376 | 0.001037 | 0.000129 |
| 0.61 | 0.191 | 0.314 | -0.71 | -2.39 | -1.67 | 0.005982 | 3.32E-06 | 2.21E-05 |
| 1.147 | 0.522 | 0.455 | 0.2 | -0.94 | -1.14 | 0.132163 | 0.005166 | 0.000748 |
| 0.01 | 0.004 | 0.366 | -6.59 | -8.04 | -1.45 | 2.52E-07 | 6.55E-08 | 0.001363 |
| 1.753 | 11.226 | 6.40E+00 | 0.81 | 3.49 | 2.68 | 0.019859 | 4.42E-05 | 0.000377 |
| 0.596 | 0.194 | 0.325 | -0.75 | -2.37 | -1.62 | 0.002658 | 2.05E-06 | 1.54E-05 |
| 0.654 | 0.171 | 0.261 | -0.61 | -2.55 | -1.94 | 0.027161 | 1.29E-05 | 6.41E-05 |
| 0.21 | 0.017 | 0.079 | -2.25 | -5.92 | -3.67 | 0.011102 | 5.50E-05 | 0.000747 |
| 0.032 | 5.50E-02 | 1.73E+00 | -4.98 | -4.19 | 0.79 | 0.000418 | 0.000696 | 0.75536 |
| 1.178 | 2.751 | 2.335 | 0.24 | 1.46 | 1.22 | 0.189149 | 1.33E-05 | 3.16E-05 |
| 0.834 | 0.085 | 0.103 | -0.26 | -3.55 | -3.29 | 0.339966 | 5.40E-07 | 6.93E-07 |
| 1.029 | 0.008 | 0.008 | 0.04 | -6.97 | -7.02 | 0.828106 | 2.38E-07 | 2.13E-07 |
| 1.121 | 0.894 | 0.797 | 0.16 | -0.16 | -0.33 | 0.268228 | 0.528669 | 0.063787 |
| 0.741 | 0.434 | 0.585 | -0.43 | -1.2 | -0.77 | 0.218275 | 0.000761 | 0.003635 |
| 1.799 | 0.472 | 2.63E-01 | 0.85 | -1.08 | -1.93 | 0.074518 | 0.008537 | 0.000778 |
| 10.965 | 5.70E-02 | 5.00E-03 | 3.45 | -4.14 | -7.59 | 1.32E-06 | 6.64E-07 | 5.21E-09 |
| 1.081 | 0.159 | 0.147 | 0.11 | -2.65 | -2.76 | 0.912737 | 0.000118 | 9.48E-05 |
| 1.045 | 0.387 | 0.371 | 0.06 | -1.37 | -1.43 | 0.26804 | 9.42E-06 | 4.92E-06 |
| 0.353 | 2.13E+00 | 6.037 | -1.5 | 1.09 | 2.59 | 0.035587 | 0.121239 | 0.003009 |

| | | | | | | | | |
|---|---|---|---|---|---|---|---|---|
| 0.277 | 0.181 | 0.652 | -1.85 | -2.47 | -0.62 | 1.79E-05 | 2.46E-06 | 0.003584 |
| 1.04 | 2.97E-01 | 2.85E-01 | 0.06 | -1.75 | -1.81 | 0.663043 | 8.25E-05 | 5.26E-05 |
| 0.743 | 4.74E-01 | 6.38E-01 | -0.43 | -1.08 | -0.65 | 0.008976 | 0.000105 | 0.002401 |
| 0.04 | 0.023 | 0.572 | -4.65 | -5.46 | -0.81 | 8.22E-07 | 3.51E-07 | 0.005146 |
| 1.12E+00 | 0.253 | 2.26E-01 | 0.16 | -1.98 | -2.14 | 0.355029 | 3.75E-05 | 1.92E-05 |
| 1.22E+00 | 1.054 | 8.64E-01 | 2.90E-01 | 0.08 | -0.21 | 0.971215 | 0.817944 | 0.923116 |
| 6.66E-01 | 0.378 | 5.68E-01 | -5.90E-01 | -1.4 | -0.81 | 0.618029 | 0.068349 | 0.234964 |
| 0.5 | 0.506 | 1.012 | -1 | -0.98 | 0.02 | 0.487573 | 0.103764 | 0.459092 |
| 0.862 | 9.71E-01 | 1.13E+00 | -0.21 | -0.04 | 0.17 | 0.841249 | 0.732444 | 0.977481 |
| 0.205 | 0.258 | 1.258 | -2.28 | -1.95 | 0.33 | 3.33E-07 | 6.47E-07 | 0.017303 |
| 0.787 | 3.115 | 3.957 | -0.35 | 1.64 | 1.98 | 0.469988 | 0.013945 | 0.00395 |
| 0.433 | 0.238 | 0.55 | -1.21 | -2.07 | -0.86 | 0.00181 | 9.79E-05 | 0.011034 |
| 0.635 | 0.291 | 0.459 | -0.66 | -1.78 | -1.12 | 0.143385 | 0.007179 | 0.093974 |
| 1.89E+00 | 12.696 | 6.74E+00 | 9.10E-01 | 3.67 | 2.75 | 0.077393 | 2.54E-05 | 9.41E-05 |
| 0.561 | 0.185 | 0.33 | -0.84 | -2.44 | -1.6 | 0.315959 | 0.004283 | 0.022744 |
| 1.221 | 1.295 | 1.06 | 0.29 | 0.37 | 0.08 | 0.186206 | 0.150369 | 0.984552 |
| 8.74E-01 | 22.66 | 2.59E+01 | -0.19 | 4.5 | 4.7 | 0.220416 | 6.42E-07 | 4.71E-07 |
| 1.232 | 4.33E-01 | 3.52E-01 | 0.3 | -1.21 | -1.51 | 0.029552 | 3.02E-05 | 7.10E-06 |
| 1.073 | 1.37E-01 | 1.28E-01 | 0.1 | -2.86 | -2.97 | 0.512655 | 6.81E-07 | 5.64E-07 |
| 0.664 | 0.273 | 0.412 | -0.59 | -1.87 | -1.28 | 0.043192 | 0.000207 | 0.002006 |
| 2.401 | 2.50E+00 | 1.04E+00 | 1.26 | 1.32 | 0.06 | 0.17683 | 0.090836 | 0.863386 |
| 0.66 | 1.194 | 1.809 | -0.6 | 0.26 | 0.86 | 0.184597 | 0.240954 | 0.019641 |
| 0.527 | 0.681 | 1.291 | -0.92 | -0.56 | 0.37 | 0.099848 | 0.228097 | 0.798863 |
| 1.392 | 0.792 | 0.569 | 0.48 | -0.34 | -0.81 | 0.646918 | 0.711144 | 0.269086 |
| 1.03E+00 | 0.626 | 0.609 | 0.04 | -0.68 | -0.71 | 0.851854 | 0.001193 | 0.00186 |
| 1.95 | 0.931 | 0.477 | 0.96 | -0.1 | -1.07 | 0.000771 | 0.463313 | 0.000323 |
| 1.254 | 1.024 | 0.817 | 0.33 | 0.03 | -0.29 | 0.21285 | 0.668887 | 0.069264 |
| 1.93E+00 | 27.375 | 1.42E+01 | 9.40E-01 | 4.77 | 3.83 | 0.485346 | 0.000208 | 0.000454 |
| 2.158 | 0.624 | 0.289 | 1.11 | -0.68 | -1.79 | 5.37E-06 | 5.31E-05 | 4.28E-07 |
| 0.046 | 4.00E-02 | 8.74E-01 | -4.46 | -4.65 | -0.19 | 0.002065 | 0.00128 | 0.834505 |
| 7.86E-01 | 5.40E-01 | 6.87E-01 | -0.35 | -0.89 | -0.54 | 0.009795 | 0.000393 | 0.019691 |
| 0.244 | 1.30E+01 | 5.35E+01 | -2.04 | 3.71 | 5.74 | 0.057857 | 0.001981 | 0.000237 |
| 3.533 | 2.85E+00 | 8.06E-01 | 1.82 | 1.51 | -0.31 | 1.17E-06 | 3.81E-06 | 0.007044 |
| 5.41E-01 | 0.151 | 2.78E-01 | -8.90E-01 | -2.73 | -1.84 | 0.009928 | 0.000115 | 0.002555 |
| 1.383 | 6.07E-01 | 4.39E-01 | 0.47 | -0.72 | -1.19 | 0.167007 | 0.435915 | 0.032131 |
| 0.697 | 4.31E-01 | 6.18E-01 | -0.52 | -1.21 | -0.69 | 0.041886 | 0.002174 | 0.063862 |
| 0.736 | 0.69 | 9.39E-01 | -0.44 | -0.53 | -0.09 | 0.016386 | 0.009941 | 0.890634 |
| 0.124 | 0.015 | 0.124 | -3.01 | -6.01 | -3.01 | 0.004432 | 0.000323 | 0.035142 |
| 1.164 | 0.198 | 0.17 | 0.22 | -2.34 | -2.56 | 0.651471 | 0.000289 | 0.000166 |
| 8.60E-01 | 1.059 | 1.23E+00 | -0.22 | 0.08 | 0.3 | 0.893388 | 0.504412 | 0.75804 |
| 0.124 | 0.091 | 0.733 | -3.01 | -3.46 | -0.45 | 0.010569 | 0.00593 | 0.841856 |
| 3.90E-02 | 0.032 | 8.25E-01 | -4.70E+00 | -4.97 | -0.28 | 5.17E-08 | 2.45E-08 | 0.044188 |
| 0.073 | 0.077 | 1.057 | -3.78 | -3.7 | 0.08 | 1.64E-07 | 2.41E-07 | 0.186586 |
| 0.021 | 0.02 | 0.959 | -5.61 | -5.67 | -0.06 | 2.76E-09 | 2.00E-09 | 0.540911 |
| 1.09 | 1.122 | 1.029 | 0.12 | 0.17 | 0.04 | 0.856173 | 0.964907 | 0.720542 |
| 0.713 | 0.204 | 2.86E-01 | -0.49 | -2.3 | -1.81 | 0.001494 | 3.64E-07 | 1.07E-06 |
| 0.651 | 0.21 | 0.322 | -0.62 | -2.25 | -1.63 | 7.60E-06 | 1.22E-10 | 1.57E-08 |
| 0.699 | 0.075 | 0.107 | -0.52 | -3.74 | -3.23 | 0.126891 | 5.10E-06 | 1.24E-05 |
| 5.20E-02 | 0.029 | 5.57E-01 | -4.28 | -5.12 | -0.84 | 0.000609 | 0.000285 | 0.532321 |
| 1.055 | 5.55E-01 | 5.26E-01 | 0.08 | -0.85 | -0.93 | 0.462047 | 0.023911 | 0.006164 |
| 2.809 | 0.907 | 0.323 | 1.49 | -0.14 | -1.63 | 0.001053 | 0.510694 | 0.000451 |

| | | | | | | | | |
|---|---|---|---|---|---|---|---|---|
| 1.042 | 3.64E-01 | 3.49E-01 | 0.06 | -1.46 | -1.52 | 0.481335 | 0.00027 | 0.000131 |
| 7.30E-02 | 0.037 | 0.51 | -3.78 | -4.75 | -0.97 | 3.03E-05 | 4.71E-06 | 0.008402 |
| 1.06 | 0.96 | 9.05E-01 | 0.08 | -0.06 | -0.14 | 0.432008 | 0.969544 | 0.553583 |
| 0.889 | 0.152 | 0.171 | -0.17 | -2.72 | -2.55 | 0.330232 | 1.21E-05 | 2.33E-05 |
| 0.882 | 0.41 | 0.465 | -0.18 | -1.29 | -1.11 | 0.126239 | 5.77E-06 | 1.43E-05 |
| 1.038 | 13.543 | 13.043 | 0.05 | 3.76 | 3.71 | 0.71206 | 2.44E-06 | 1.98E-06 |
| 1.081 | 0.432 | 0.4 | 0.11 | -1.21 | -1.32 | 0.573642 | 0.000236 | 0.000127 |
| 0.036 | 0.031 | 0.858 | -4.8 | -5.02 | -0.22 | 5.77E-07 | 5.73E-07 | 0.999023 |
| 1.564 | 0.904 | 0.578 | 0.65 | -0.15 | -0.79 | 0.364385 | 0.886665 | 0.205357 |
| 0.778 | 21.477 | 27.598 | -0.36 | 4.42 | 4.79 | 0.120421 | 1.16E-05 | 4.75E-06 |
| 3.91E-01 | 7.651 | 1.96E+01 | -1.35E+00 | 2.94 | 4.29 | 0.256829 | 0.003443 | 0.000802 |
| 0.996 | 0.072 | 0.072 | -0.01 | -3.8 | -3.79 | 0.995406 | 5.81E-05 | 5.55E-05 |
| 2.40E-02 | 0.023 | 9.53E-01 | -5.38 | -5.45 | -0.07 | 6.42E-14 | 6.42E-14 | 0.094861 |
| 5.97E-01 | 0.317 | 5.31E-01 | -7.40E-01 | -1.66 | -0.91 | 0.000162 | 1.52E-06 | 4.96E-05 |
| 0.601 | 0.224 | 0.372 | -0.73 | -2.16 | -1.42 | 0.050566 | 0.000431 | 0.00506 |
| 0.844 | 0.412 | 0.487 | -0.24 | -1.28 | -1.04 | 0.254962 | 7.89E-05 | 0.000213 |
| 0.894 | 4.00E-01 | 4.47E-01 | -0.16 | -1.32 | -1.16 | 0.568633 | 0.000116 | 0.000215 |
| 2.29E+00 | 1.238 | 0.541 | 1.2 | 0.31 | -0.89 | 0.001119 | 0.269191 | 0.005009 |
| 0.277 | 1.407 | 5.076 | -1.85 | 0.49 | 2.34 | 0.020946 | 0.155031 | 0.002391 |
| 0.444 | 2.60E+01 | 5.86E+01 | -1.17 | 4.7 | 5.87 | 0.474739 | 0.000229 | 0.000112 |
| 0.817 | 2.395 | 2.933 | -0.29 | 1.26 | 1.55 | 0.029919 | 3.03E-05 | 7.15E-06 |
| 3.868 | 48.926 | 12.649 | 1.95 | 5.61 | 3.66 | 0.009784 | 2.27E-05 | 0.000225 |
| 8.73E-01 | 0.752 | 8.61E-01 | -2.00E-01 | -0.41 | -0.22 | 0.121246 | 0.00119 | 0.010245 |
| 9.01E-01 | 7.32E-01 | 8.13E-01 | -0.15 | -0.45 | -0.3 | 0.886999 | 0.02228 | 0.038818 |
| 1.12E+00 | 0.778 | 0.692 | 0.17 | -0.36 | -0.53 | 0.2834 | 0.054724 | 0.008053 |
| 0.65 | 2.07E+00 | 3.19E+00 | -0.62 | 1.05 | 1.67 | 0.011199 | 0.000513 | 4.28E-05 |
| 0.13 | 0.021 | 0.162 | -2.95 | -5.57 | -2.62 | 1.73E-06 | 2.91E-08 | 1.73E-06 |
| 1.11 | 1.68E+00 | 1.51E+00 | 0.15 | 0.75 | 0.6 | 0.653076 | 0.205178 | 0.581717 |
| 0.246 | 0.01 | 0.042 | -2.02 | -6.59 | -4.57 | 0.000159 | 2.43E-07 | 1.29E-06 |
| 1.128 | 0.68 | 0.602 | 0.17 | -0.56 | -0.73 | 0.262162 | 0.001476 | 0.000406 |
| 0.826 | 49.021 | 59.322 | -0.28 | 5.62 | 5.89 | 0.013252 | 6.44E-14 | 6.42E-14 |

| Adj_P_value_Hvidovre_Avedøre_ | Adj_P_value_Skævinge_Avedøre_ | Adj_P_value_Skævinge_Hvidovre_ | Peak_Rating_A1 | Peak_Rating_A2 | Peak_Rating_A3 | Peak_Rating_A_PB | Peak_Rating_D1 | Peak_Rating_D2 |
|---|---|---|---|---|---|---|---|---|
| 3.75E-05 | 0.071712 | 1.25E-04 | 6.60E+00 | 6.6 | 6.60E+00 | 6.40E+00 | 6.6 | 6.6 |
| 0.16156 | 0.026353 | 5.62E-01 | 4.70E+00 | 5.5 | 5.5 | 1 | 4.7 | 3.9 |
| 0.001772 | 0.000285 | 0.147059 | 3.6 | | | | | |
| 5.47E-06 | 6.72E-06 | 1.59E-08 | 6.60E+00 | 6.60E+00 | 6.60E+00 | 4.8 | 6.60E+00 | 6.6 |
| 1.48E-05 | 4.27E-06 | 0.004674 | 6.20E+00 | 6.20E+00 | 6.6 | | 2.9 | 4.2 |
| 7.40E-12 | 5.45E-12 | 0.369928 | 6.20E+00 | 6.20E+00 | 6.2 | | | |
| 0.004282 | 0.949326 | 3.58E-03 | 5.8 | 6.60E+00 | 6.60E+00 | 6.1 | 6.60E+00 | 6.2 |
| 0.015338 | 0.003525 | 0.60585 | 3.6 | 4 | 3.1 | 3.8 | 4.4 | 3.6 |
| 0.777435 | 4.25E-05 | 0.000133 | 3.6 | 3.3 | 3.6 | | 5.5 | 5.2 |
| 0.005219 | 0.13834 | 1.34E-03 | 3.30E+00 | 3.6 | 2.10E+00 | 1.50E+00 | 3.5 | 2.7 |
| 0.985576 | 0.14307 | 3.11E-01 | 2.00E-01 | 0.5 | | 4.00E-01 | 0.8 | |
| 0.017245 | 0.036083 | 6.86E-01 | 3.00E+00 | 3.3 | 3.00E+00 | | 0.6 | 1.5 |
| 3.60E-06 | 3.95E-06 | 1.19E-09 | 3.00E+00 | 4.5 | 3.4 | | 5.8 | 5.8 |
| 0.000582 | 0.000378 | 3.69E-05 | 5.80E+00 | 5.5 | 5.80E+00 | | 6.2 | 5.8 |
| 0.00181 | 0.019941 | 0.000317 | 5.2 | 5.2 | 5.2 | | 5.8 | 5.8 |
| 0.02275 | 0.54491 | 9.12E-03 | 3.50E+00 | 3.50E+00 | 5.00E+00 | | 4.80E+00 | 5.2 |
| 0.002563 | 0.001101 | 1.14E-04 | 6.20E+00 | 5.8 | 6.2 | 6.00E+00 | 6.2 | 5.8 |
| 2.27E-05 | 5.96E-09 | 3.79E-06 | 6.2 | 6.2 | 6.6 | 2.3 | 6.6 | 6.2 |
| 0.979597 | 0.489265 | 8.44E-01 | 1.2 | 2.00E+00 | 9.00E-01 | | 2.40E+00 | 2 |
| 5.86E-07 | 3.34E-07 | 9.74E-01 | 4.20E+00 | 4.6 | 4.6 | | | |
| 1 | 0.003531 | 0.008288 | 4.20E+00 | 3 | 4.2 | | 2.3 | 2.3 |
| 2.79E-05 | 1.29E-05 | 5.40E-01 | 6.20E+00 | 6.6 | 6.6 | 0.4 | 0.7 | 0.7 |
| 1.21E-05 | 2.14E-06 | 0.001484 | 6.60E+00 | 6.60E+00 | 6.6 | | 7.00E-01 | 1.5 |
| 0.028522 | 0.009424 | 0.895164 | 5.50E+00 | 4.30E+00 | 5.1 | | 3.3 | 0.2 |
| 0.070662 | 0.002974 | 8.72E-02 | 3.10E+00 | 2 | 8.00E-01 | | 0.5 | 0.2 |
| 0.002556 | 0.000773 | 0.644579 | 6.2 | 6.6 | 6.2 | | 0.2 | 0.6 |
| 0.018023 | 0.751623 | 0.009861 | 4.5 | 4.1 | 3.3 | | 5.5 | 5.5 |
| 0.597123 | 0.536007 | 1 | 2.2 | 3 | 2.6 | 0.4 | 0.8 | 1.4 |
| 9.00E-05 | 1.88E-05 | 4.93E-02 | 5.00E+00 | 4.6 | 5.00E+00 | 0.00E+00 | 4.5 | 4.2 |
| 3.71E-11 | 1.32E-06 | 5.90E-06 | 6.20E+00 | 6.2 | 6.2 | | 5.4 | |
| 0.538638 | 7.54E-06 | 2.28E-05 | 5.00E+00 | 5.00E+00 | 5 | | 4.6 | 5.4 |
| 1 | 0.00957 | 0.014423 | 4.30E+00 | 4.70E+00 | 3.9 | 4.50E+00 | 5.80E+00 | 5.5 |
| 0.392825 | 0.040224 | 3.66E-01 | 1.80E+00 | 1.7 | 7.00E-01 | 6.00E-01 | 0.8 | 1.4 |
| 1 | 0.86473 | 0.799087 | 1.80E+00 | 2.20E+00 | 1.4 | 1.10E+00 | 7.00E-01 | 0.9 |
| 0.093921 | 0.174672 | 8.98E-01 | 5.10E+00 | 4.50E+00 | 5.20E+00 | | | 3 |
| 0.488832 | 0.119643 | 7.27E-01 | 2.20E+00 | 3.70E+00 | 3.00E+00 | | 2.60E+00 | 2.6 |
| 0.286737 | 0.353851 | 1 | | 1.3 | 2.3 | | | |
| 0.49199 | 1 | 0.491441 | 7.00E-01 | 7.00E-01 | 0.8 | 4.00E-01 | | |
| 0.40931 | 0.039523 | 1.33E-02 | 1.20E+00 | 1.1 | 2.60E+00 | 0.00E+00 | 0.7 | 0.9 |
| 0.402063 | 0.017488 | 1.49E-01 | 1.20E+00 | 2 | 1.60E+00 | | 2.7 | 1.2 |

| | | | | | | | | |
|---|---|---|---|---|---|---|---|---|
| 0.61058 | 0.033805 | 0.187135 | 2.2 | 2.2 | 1.8 | | 4.6 | 1.8 |
| 0.209419 | 0.041587 | 0.666095 | 5.2 | 5.2 | 4.8 | 1.9 | 1.2 | 3.6 |
| 0.112945 | 0.000529 | 5.16E-03 | 5.80E+00 | 5.8 | 5.80E+00 | 4.50E+00 | 3.7 | 3.7 |
| 1 | 0.61 | 0.522267 | 1.3 | 2.2 | 3.2 | | | |
| 8.33E-05 | 3.24E-05 | 0.598814 | 3.3 | 0.8 | 0.8 | 2.3 | 6.6 | 6 |
| 0.0001 | 4.05E-05 | 0.66329 | 6.60E+00 | 6.6 | 6.6 | 3.9 | 1.6 | 2.1 |
| 1.83E-07 | 5.94E-08 | 1.30E-01 | 6.60E+00 | 6.6 | 6.60E+00 | 4.50E+00 | 3 | 1.9 |
| 0.040689 | 0.002271 | 1.02E-01 | 4.80E+00 | 4.10E+00 | 5.00E+00 | 4.50E+00 | 5.00E-01 | 2.3 |
| 0.014618 | 0.007863 | 1 | 6.2 | 6.2 | 6.6 | | 1.9 | 0.2 |
| 0.028923 | 1 | 0.03363 | 1.7 | 4.5 | 0.6 | | 3.7 | 3.9 |
| 1 | 0.000852 | 0.001826 | 1.1 | | 0.8 | | | 1.8 |
| 2.88E-05 | 2.56E-05 | 0.453763 | 1.4 | 1.8 | 2.6 | | | |
| 7.40E-12 | 5.45E-12 | 0.063608 | 6.6 | 6.20E+00 | 6.6 | | | |
| 0.000126 | 5.26E-05 | 7.62E-01 | 6.00E+00 | 6 | 5.60E+00 | | | |
| 0.000154 | 0.020686 | 0.001027 | 3.3 | 2.5 | 3.3 | | 3.9 | 4.2 |
| 0.965665 | 0.052494 | 0.134118 | 2 | 2.8 | 2 | 0 | | 0.9 |
| 0.46853 | 0.522039 | 1 | 1.1 | 0.6 | | 0 | | |
| 0.000348 | 0.000219 | 1.00E+00 | 6.20E+00 | 6.2 | 6.20E+00 | 8.00E-01 | 4.7 | 4.3 |
| 0.023476 | 0.024503 | 1.00E+00 | 5.80E+00 | 5.80E+00 | 5.50E+00 | 1.90E+00 | 2.1 | |
| 0.018023 | 0.016587 | 1 | 1.7 | 1.3 | 1.8 | 3 | 5.1 | 5.1 |
| 0.314729 | 0.006471 | 0.059681 | 4.3 | 3.9 | 4.3 | | 1.8 | 1.3 |
| 5.27E-05 | 6.54E-06 | 6.66E-04 | 6.20E+00 | 5.80E+00 | 5.10E+00 | 1.70E+00 | 3.70E+00 | 3.3 |
| 0.156533 | 0.001499 | 0.016433 | 2.5 | 4.8 | 2.9 | | 0.5 | 1.4 |
| 0.001297 | 0.557246 | 3.01E-03 | | 0.7 | 7.00E-01 | 1.10E+00 | 0.7 | 0.7 |
| 0.001044 | 0.888711 | 1.67E-03 | 6.00E+00 | 6.4 | 6.40E+00 | | 6.1 | 5.8 |
| 0.661681 | 0.096989 | 4.15E-02 | 6.00E-01 | 0.6 | 1.9 | | | 1.5 |
| 8.38E-05 | 1.11E-05 | 0.005807 | 3.5 | 5 | 3.7 | | 1.9 | 2.5 |
| 0.753967 | 6.72E-05 | 0.000115 | 4.60E+00 | 5.00E+00 | 5 | | 5.00E+00 | 4.6 |
| 1.48E-05 | 0.000255 | 0.000214 | 3.7 | 4.1 | 5.2 | 1.5 | 3.3 | 3.3 |
| 1 | 0.190612 | 0.182599 | 0.6 | 2.3 | | | 2 | 3 |
| 5.00E-05 | 1.67E-05 | 0.200188 | 6.60E+00 | 6.60E+00 | 6.6 | 6.40E+00 | 7.40E+00 | 7 |
| 1 | 0.977657 | 1.00E+00 | 3.00E+00 | 1.8 | | | 1.1 | 1.3 |
| 0.145509 | 0.001563 | 1.84E-02 | 5.00E+00 | 5.8 | 5.40E+00 | | 5 | 5.4 |
| 0.170063 | 0.514137 | 6.68E-01 | | | 0.5 | 0.8 | | |
| 0.879946 | 0.620884 | 1 | 2.6 | 0.7 | 2.4 | 0.4 | 4.1 | 3.2 |
| 0.984385 | 1 | 0.990589 | 1.10E+00 | 1.10E+00 | 0.9 | | 1.70E+00 | 0.8 |
| 0.64385 | 0.00255 | 0.010483 | 1.1 | | | | | 0.8 |
| 0.241412 | 0.008872 | 0.115053 | 2.6 | 1.4 | 1.8 | | | 1.4 |
| 1 | 0.163787 | 0.292112 | 1.7 | 1.1 | 1.3 | | 3.2 | 2.6 |
| 0.480597 | 0.02728 | 1.99E-01 | 2.60E+00 | 3.3 | 2.60E+00 | | 0.8 | 3 |
| 0.432053 | 0.001141 | 0.00578 | 3.5 | 2.7 | 3.5 | | 3.5 | 5.4 |
| 0.07194 | 0.000203 | 1.85E-03 | 2.00E+00 | 2.2 | 2.20E+00 | 1.50E+00 | 2.3 | 2.3 |
| 0.633762 | 0.103003 | 5.06E-01 | 9.00E-01 | 2.40E+00 | 1.70E+00 | | 2.6 | 0.9 |
| 0.168575 | 0.006688 | 1.15E-01 | 1.30E+00 | 0.9 | 1.7 | | 1.4 | 1.3 |
| 0.74784 | 0.049571 | 0.20659 | 2.6 | 2.7 | 2.3 | 0 | 2.9 | 3.5 |
| 0.711558 | 0.315935 | 9.32E-01 | 3.10E+00 | 3.3 | 3.5 | 1.90E+00 | 5 | 4.2 |
| 2.27E-05 | 7.58E-07 | 3.52E-05 | 6.20E+00 | 6.20E+00 | 6.2 | 1.90E+00 | 6 | 6.4 |
| 0.939933 | 0.0984 | 2.59E-01 | 1.10E+00 | 1.8 | 7.00E-01 | 6.00E-01 | 0.8 | |
| 0.677692 | 0.114829 | 4.89E-02 | | | 8.00E-01 | 1.50E+00 | 0.9 | |
| 7.22E-05 | 9.45E-06 | 0.002616 | 6.6 | 6.6 | 6.6 | 5.8 | 6.6 | 5.8 |
| 1.70E-05 | 0.354596 | 2.24E-05 | 6.6 | 6.6 | 6.2 | 3.6 | 6.6 | 6.6 |

| | | | | | | | | |
|---|---|---|---|---|---|---|---|---|
| 0.98994 | 0.000539 | 0.001268 | 5.4 | 4.6 | 3.1 | 3.8 | 3.1 | 3.1 |
| 0.022814 | 0.053711 | 0.663123 | 3.3 | 4.8 | 3 | 0.4 | 1.7 | 0.2 |
| 0.852849 | 1 | 0.927468 | 2.70E+00 | 3.10E+00 | 2.5 | | 2.30E+00 | 3.6 |
| 2.03E-05 | 6.76E-07 | 3.61E-05 | 5.6 | 5.6 | 6 | 0.2 | 5.8 | 5 |
| 2.27E-05 | 7.58E-07 | 3.52E-05 | 6.20E+00 | 6.2 | 6.20E+00 | 1.90E+00 | 6 | 6.4 |
| 0.000149 | 7.58E-05 | 1.00E+00 | 3.70E+00 | 4.1 | 5.60E+00 | | | |
| 0.979597 | 0.234557 | 0.491283 | 3.1 | 3.5 | 2.1 | 1.9 | 1 | 1.4 |
| 0.028729 | 0.228729 | 2.49E-01 | 5.60E+00 | 5.2 | 5.20E+00 | 0.00E+00 | 4.8 | 4.5 |
| 0.001344 | 0.000316 | 0.317831 | 0.8 | 2.7 | 4.6 | | 2.3 | 0 |
| 0.965665 | 0.196343 | 0.1451 | 3.3 | 3 | 3.7 | 1 | 0.7 | 0.7 |
| 0.125868 | 0.101467 | 1.00E+00 | 5.50E+00 | 4.7 | 5.50E+00 | 0.00E+00 | 3.6 | |
| 2.17E-05 | 1.22E-05 | 1 | 6.6 | 6.2 | 6.6 | 3.9 | 6.2 | 6.6 |
| 0.961425 | 0.960799 | 1 | 2.2 | 1.1 | 1.2 | 2.3 | 1.8 | 2.2 |
| 0.001793 | 0.073664 | 0.000501 | 2.3 | | 1.6 | | 0.7 | 0.7 |
| 0.043588 | 0.000337 | 5.07E-03 | 1.10E+00 | 1.4 | 0.7 | | | |
| 0.264477 | 0.749865 | 0.653147 | 5.50E+00 | 5.50E+00 | 6.2 | 5.8 | 5.5 | 5.8 |
| 0.12081 | 1 | 0.113202 | 6.2 | 5.5 | 5.5 | 4.2 | 5.8 | 5.8 |
| 0.113578 | 0.513972 | 0.032606 | 6.2 | 5.8 | 6.2 | 4.4 | 6.2 | 5.8 |
| 0.00054 | 0.008831 | 1.14E-04 | 1.10E+00 | 2.3 | 1.90E+00 | | 4.2 | 3.5 |
| 9.78E-12 | 0.140477 | 2.29E-11 | 4.7 | 4.4 | 4.7 | 4.5 | 6.2 | 6.6 |
| 0.175918 | 0.347023 | 8.89E-01 | 5.80E+00 | 4.00E+00 | 3.30E+00 | 1.90E+00 | 3.60E+00 | 4.8 |
| 0.020169 | 0.000257 | 5.89E-03 | 6.60E+00 | 6.2 | 6.2 | 1.70E+00 | 6.6 | 6.6 |
| 0.007943 | 8.57E-05 | 0.002117 | 6.2 | 5.8 | 6.2 | | 5.2 | 5.2 |
| 0.000105 | 0.0002 | 9.33E-02 | 6.40E+00 | 6.40E+00 | 6.60E+00 | 4.50E+00 | 6.00E+00 | 5.7 |
| 1.04E-05 | 0.004527 | 2.77E-05 | 4.10E+00 | 3.7 | 3 | | 6.2 | 6.6 |
| 0.740718 | 0.67058 | 0.302326 | 3.2 | 3.9 | 3.7 | | 4.3 | 3.9 |
| 0.391803 | 5.99E-05 | 8.56E-05 | 6.20E+00 | 6.20E+00 | 5.5 | | 6.6 | 6.2 |
| 0.001248 | 0.000324 | 0.397596 | 6.60E+00 | 6.60E+00 | 6.2 | 2.30E+00 | 4.50E+00 | |
| 0.007805 | 0.003061 | 9.50E-01 | 5.20E+00 | 4.8 | 4.80E+00 | 0.00E+00 | 0 | 1.5 |
| 1.88E-08 | 2.72E-08 | 0.210981 | 5.8 | 6.2 | 6.2 | 5 | | |
| 0.22889 | 0.002545 | 0.024372 | 6.2 | 6.2 | 6.2 | 2.4 | 6.2 | 6.2 |
| 0.041633 | 0.000223 | 0.002894 | 3.3 | 3 | 3.3 | 0.4 | 1.6 | 2 |
| 0.154288 | 1.68E-05 | 7.79E-05 | 5.80E+00 | 5.8 | 5.50E+00 | | 5.5 | 5.8 |
| 0.000612 | 0.000334 | 1 | 4.70E+00 | 5.5 | 4.3 | | 1.4 | 2.1 |
| 0.009707 | 0.000993 | 1.43E-01 | 4.10E+00 | 4.7 | 4.1 | 2.70E+00 | 4.1 | 4.5 |
| 0.053528 | 4.12E-05 | 4.20E-05 | 4.8 | 3.7 | 5.2 | 4.2 | 4.8 | 5 |
| 0.00276 | 0.001135 | 0.000118 | 5.8 | 5.6 | 5.8 | 3.8 | 6 | 5.6 |
| 1.88E-06 | 4.17E-11 | 8.05E-06 | 2.00E+00 | 2 | | | 0.8 | 2 |
| 0.000125 | 5.13E-05 | 1.43E-05 | 5.20E+00 | 4 | 3.3 | | 4.6 | 5.4 |
| 0.000393 | 0.001102 | 0.101166 | 4.00E+00 | 3.1 | 3.9 | | 5.2 | 5.6 |
| 0.365429 | 0.002555 | 1.60E-03 | 5.50E+00 | 4.3 | 4.3 | | 5.1 | 5.1 |
| 1 | 0.77416 | 6.79E-01 | 4.5 | 1.7 | 3.70E+00 | | 5.5 | 4.1 |
| 0.000453 | 0.077298 | 1.97E-04 | | | | | 3.1 | 3 |
| 0.00287 | 1.32E-05 | 1.67E-05 | 5.8 | 6.20E+00 | 6.2 | | 6 | 5.6 |
| 2.24E-08 | 1.05E-08 | 0.539514 | 3.1 | 5.8 | 5 | | | |
| 0.666566 | 0.007274 | 3.23E-02 | 1.70E+00 | 1.7 | 2.00E+00 | 3.40E+00 | 3.2 | 3 |
| 0.84566 | 7.72E-06 | 1.89E-05 | 5 | 5 | 4.6 | | 3.1 | 4.6 |
| 0.077842 | 0.000288 | 0.002841 | 6.6 | 6.6 | 6.6 | 5.2 | 6.6 | 6.6 |
| 0.000559 | 2.14E-05 | 0.001961 | 6 | 4.5 | 5.6 | 2.3 | 5.6 | 5.2 |
| 2.04E-05 | 4.03E-06 | 0.00079 | 5.6 | 5.6 | 6 | | 6.9 | |
| 0.875451 | 0.001864 | 2.16E-03 | 4.8 | 5.20E+00 | 4.5 | 1.9 | 5.2 | 5.2 |

| | | | | | | | | |
|---|---|---|---|---|---|---|---|---|
| 0.053528 | 0.00843 | 4.81E-01 | 2.70E+00 | 1.60E+00 | 1.60E+00 | 4.50E+00 | 3.3 | 2 |
| 0.99726 | 0.000608 | 0.001416 | 5.1 | 5.5 | 5.8 | 0.8 | 5.8 | 5.5 |
| 0.000841 | 0.000619 | 1 | 4.5 | 5.6 | 4.5 | | 4.2 | 3.5 |
| 0.478218 | 1 | 5.19E-01 | 6.2 | 6.2 | 5.8 | 0.2 | 6.6 | 6.2 |
| 0.257183 | 0.256819 | 3.64E-02 | 5.20E+00 | 5.4 | 5.00E+00 | | 5.6 | 5.2 |
| 0.016484 | 0.000347 | 0.000123 | 5.10E+00 | 4.7 | 4.7 | | 6.2 | 6.2 |
| 0.586625 | 1.03E-05 | 2.34E-05 | 6.00E+00 | 6 | 5.60E+00 | | 6.4 | 5.2 |
| 0.000353 | 3.60E-08 | 1.90E-06 | 6.60E+00 | 6.6 | 6.6 | 6.6 | 6.6 | 6.6 |
| 0.249365 | 0.000735 | 0.004888 | 6.6 | 6.6 | 6.6 | 4.9 | 6.2 | 6.2 |
| 0.905329 | 0.235583 | 0.598814 | 3.3 | 3 | 3.1 | | 3 | 3.9 |
| 0.001344 | 1.75E-07 | 3.76E-06 | 5.6 | 5.6 | 5.2 | | 5.2 | 5.6 |
| 0.044416 | 0.003343 | 0.16737 | 5.8 | 4.3 | 4.3 | | 4.5 | 5.2 |
| 5.78E-05 | 3.11E-05 | 1.00E+00 | | 5.2 | 4.80E+00 | | 6.2 | 5.8 |
| 1 | 0.10707 | 2.19E-01 | 3.60E+00 | 2.4 | | 3.40E+00 | 4.7 | 2 |
| 0.626139 | 1.58E-05 | 5.04E-05 | 4.8 | 4.1 | 4.8 | | 4.8 | 5.6 |
| 0.005899 | 0.041145 | 0.151117 | 5.1 | 4.7 | 5.1 | | 5.2 | 4.8 |
| 0.984977 | 0.001058 | 0.002528 | 4.60E+00 | 6 | 5.6 | | 4.8 | 5.4 |
| 1 | 0.311779 | 0.564333 | 5.4 | 5.4 | 4.2 | | 5 | 5.2 |
| 1.35E-11 | 1.06E-05 | 2.22E-11 | | | | 6.70E+00 | 6.2 | 6.2 |
| 0.295923 | 0.259943 | 1 | 4.2 | 4.6 | 4.6 | | 4.6 | 5 |
| 1 | 0.599327 | 6.16E-01 | 5.50E+00 | 6.6 | 5.8 | | 6.6 | 6.2 |
| 0.290699 | 0.000156 | 0.000707 | 6.60E+00 | 6.6 | 5.8 | 8.00E-01 | 6.2 | 6.6 |
| 0.410311 | 0.757045 | 1.74E-01 | 5.60E+00 | 6 | 5.80E+00 | | 6 | 5.6 |
| 0.279655 | 0.20886 | 3.31E-02 | 5.80E+00 | 4.6 | 4.20E+00 | | 4.2 | 4.2 |
| 0.627424 | 0.102318 | 0.040834 | 4.2 | 3.5 | 5 | | 2.7 | 2.7 |
| 0.83105 | 0.747109 | 1 | 6.6 | 6.2 | 6.6 | 0.8 | 6.2 | 6.6 |
| 0.268962 | 0.179292 | 2.85E-02 | 6.2 | 6.2 | 6.2 | | 6.2 | 6.6 |
| 0.067998 | 8.84E-05 | 0.000671 | 6.20E+00 | 6.20E+00 | 5.8 | | 6.60E+00 | 6.2 |
| 0.240829 | 9.49E-05 | 4.62E-04 | 4.20E+00 | 3.9 | 4.2 | 4.50E+00 | 3.1 | 3.1 |
| 0.169429 | 1.12E-05 | 2.34E-05 | 5.80E+00 | 5.8 | 5.80E+00 | 1.90E+00 | 6.2 | 6.2 |
| 0.000356 | 0.861582 | 0.00059 | | | | 4.4 | 6.2 | 6.2 |
| 0.725475 | 0.009901 | 0.007531 | 6.2 | 6.2 | 6.2 | 3.4 | 6.6 | 6.2 |
| 1 | 0.011831 | 0.016374 | 5.8 | 6.2 | 6.2 | 6.6 | 6.6 | 6.2 |
| 0.045302 | 0.001233 | 0.034964 | 4.00E-01 | 1.50E+00 | 4.5 | | 2.00E-01 | |
| 0.263623 | 0.02605 | 0.353429 | 2.7 | 2 | 3.1 | | | 0.6 |
| 0.891976 | 0.000691 | 0.00189 | 2.7 | 3.1 | 3.1 | 1.9 | 2.3 | 3.1 |
| 0.013388 | 0.000177 | 7.43E-05 | 6.00E-01 | 2.00E-01 | 2.00E+00 | | | |
| 1 | 0.046593 | 0.062989 | | | 4.8 | 2.3 | | |
| 0.538638 | 0.016923 | 0.008817 | | | 1.7 | | | |
| 0.143485 | 0.580742 | 5.23E-01 | 5.00E+00 | 5.8 | 5.40E+00 | | 6 | 5.8 |
| 0.155362 | 0.027558 | 6.03E-01 | 3.10E+00 | 2 | 3.50E+00 | | 2.9 | 0.2 |
| 0.175175 | 0.001941 | 2.13E-02 | 4.1 | 5.2 | 5.6 | | 4.2 | 4.2 |
| 7.22E-05 | 5.45E-12 | 2.22E-11 | 4.5 | 3.7 | 3.7 | | 3.1 | 2.7 |
| 0.978006 | 0.00255 | 0.003379 | 7 | 6.6 | 6.2 | 1.9 | 6.2 | 6.6 |
| 0.329114 | 0.305761 | 1.00E+00 | 9.00E-01 | 0.9 | 3.90E+00 | 8.00E-01 | 0.5 | 1.2 |
| 0.095818 | 3.46E-05 | 0.000213 | 5.8 | 4.7 | 5.8 | | 4.3 | 5.1 |
| 0.899277 | 0.004292 | 4.61E-03 | 6.20E+00 | 6.6 | 6.2 | 3.00E+00 | 5.1 | 6.6 |
| 0.796658 | 0.003413 | 3.38E-03 | 3.90E+00 | 4.2 | 4.60E+00 | | 5.8 | 4.2 |
| 0.334951 | 6.68E-06 | 1.54E-05 | 6.6 | 6.6 | 6.6 | 4 | 6.6 | 6.6 |
| 0.004837 | 1.54E-05 | 0.000217 | 4.5 | 4.5 | 4.1 | | 3.7 | 4.1 |
| 0.010051 | 3.22E-05 | 0.000455 | 5.5 | 5.8 | 5.8 | | 6.6 | 5.5 |

| | | | | | | | | |
|---|---|---|---|---|---|---|---|---|
| 1 | 2.52E-05 | 6.38E-05 | 4.1 | 3.7 | 4.1 | | 3.7 | 4.8 |
| 0.023012 | 0.005558 | 0.677485 | 6.6 | 6.6 | 6.6 | 4.6 | 6.6 | 6.2 |
| 1.89E-05 | 3.53E-06 | 0.000692 | 6.2 | 6.6 | 6.2 | 1.7 | 6.2 | 6.6 |
| 0.002743 | 1.06E-05 | 1.43E-05 | 6.6 | 6.6 | 6.2 | 3.3 | 6.2 | 6.6 |
| 0.197464 | 0.224356 | 1 | 3.10E+00 | 2.70E+00 | 2.3 | | 3.80E+00 | 1.4 |
| 0.672421 | 8.62E-06 | 2.01E-05 | 5.2 | 5.5 | 5.2 | 3.4 | 4.8 | 4 |
| 0.002586 | 0.000315 | 0.095448 | 6.6 | 6.6 | 6.6 | 5 | 6.6 | 6.6 |
| 7.40E-12 | 5.45E-12 | 3.32E-01 | 5.50E+00 | 5.50E+00 | 4.70E+00 | | | |
| 1 | 0.009033 | 0.011755 | 3.30E+00 | 3.3 | 4.1 | | 4.8 | 4.1 |
| 1 | 3.50E-05 | 8.60E-05 | 6.6 | 6.2 | 6.6 | | 6.6 | 6.2 |
| 0.000131 | 3.23E-05 | 1.44E-01 | 6.60E+00 | 6.6 | 6.6 | 2.70E+00 | 1.9 | 0.2 |
| 0.036617 | 8.90E-05 | 6.33E-05 | 5.50E+00 | 5.5 | 5.1 | | 5.5 | 4.3 |
| 0.02562 | 0.043202 | 0.846306 | 4.8 | 4.8 | 5.5 | 1.5 | 5.8 | 5.4 |
| 0.557358 | 0.086133 | 0.504474 | 2.5 | 0.5 | 0.8 | 1.5 | 0.5 | 0.8 |
| 1 | 0.00843 | 0.012158 | 4.8 | 3.3 | 3.7 | | 4.1 | 4.1 |
| 0.079478 | 0.001052 | 0.017006 | 5.8 | 6.2 | 5.8 | | 5.1 | 6.2 |
| 0.013777 | 0.001534 | 0.195472 | 3.5 | 3.1 | 2.3 | | 2.1 | 4.8 |
| 0.030955 | 0.021654 | 1 | 3.2 | 2.8 | 2.8 | 0.8 | 5.2 | 5.5 |
| 0.916439 | 0.02026 | 5.90E-02 | 4.30E+00 | 3.6 | 3.90E+00 | 8.00E-01 | 3.6 | 3.6 |
| 1 | 0.272585 | 3.83E-01 | 7.00E-01 | 0.7 | 1.10E+00 | 0.00E+00 | | |
| 0.037966 | 0.001534 | 0.05804 | 2.7 | 4.20E+00 | 3.5 | | 1.7 | 1.4 |
| 0.001917 | 0.000639 | 7.50E-05 | 3.90E+00 | 2.70E+00 | 4.2 | | 5.60E+00 | 4.8 |
| 0.19546 | 0.085802 | 0.013065 | 5.50E+00 | 6.2 | 6.2 | 1.10E+00 | 5.5 | 6.2 |
| 0.00088 | 0.001404 | 0.425282 | 6.6 | 6.6 | 6.6 | | | 2.8 |
| 2.02E-06 | 1.18E-06 | 9.90E-01 | 5.80E+00 | 6 | 5.8 | | 5.1 | 3.2 |
| 0.434777 | 0.307792 | 1 | 3.30E+00 | 3.5 | 2.6 | 0 | 0.5 | 0.5 |
| 3.52E-06 | 1.94E-06 | 9.93E-01 | 3.60E+00 | 6.2 | 6.6 | | | |
| 0.101698 | 3.50E-05 | 4.25E-05 | 6.2 | 6.6 | 5.8 | 4.2 | 6.6 | 6.2 |
| 0.022401 | 0.018313 | 1 | 3.9 | 4.2 | 3.5 | | 2.3 | 2.7 |
| 0.022439 | 0.021313 | 1 | 3.1 | 3.5 | 3.5 | | 2.3 | 1.9 |
| 0.466892 | 0.365919 | 1.00E+00 | 1.20E+00 | 2.2 | 4.40E+00 | | 2.5 | 1.7 |
| 0.190057 | 0.005163 | 7.18E-02 | 5.50E+00 | 5.5 | 5.80E+00 | | 5.5 | 5.5 |
| 0.002865 | 0.001735 | 0.000148 | 3.10E+00 | 3.10E+00 | 4.6 | 2.00E-01 | 6.6 | 6.2 |
| 0.454674 | 0.001044 | 5.04E-03 | 5.20E+00 | 4.5 | 4.10E+00 | | 4.5 | 4.8 |
| 0.000327 | 0.086852 | 1.55E-04 | 5.10E+00 | 4.7 | 3.9 | | 5.8 | 6.2 |
| 5.54E-05 | 0.987039 | 8.69E-05 | 1.50E+00 | 2.30E+00 | 2.30E+00 | 4.90E+00 | 5.5 | 5.5 |
| 1.18E-10 | 0.001029 | 2.81E-11 | | | | 6.20E+00 | 4.3 | 4.7 |
| 1.21E-08 | 0.000591 | 1.19E-09 | | | | 1.30E+00 | 5.1 | 4.7 |
| 0.088542 | 0.029437 | 0.003966 | 6.20E+00 | 6.2 | 6.2 | 2.70E+00 | 5.8 | 6.2 |
| 0.009211 | 0.000175 | 6.48E-05 | 2.3 | 2.3 | 2 | | 3.7 | 3.3 |
| 1 | 0.789635 | 8.33E-01 | 6.20E+00 | 6.20E+00 | 6.6 | | 5.5 | 6.2 |
| 0.161513 | 0.953798 | 2.68E-01 | 3.00E+00 | 4.1 | 4.50E+00 | 1.50E+00 | 3.5 | 3.5 |
| 1 | 0.001686 | 0.002497 | 6.2 | 6.6 | 5.8 | | 6.2 | 6.2 |
| 0.882121 | 0.015153 | 1.32E-02 | 4.30E+00 | 3.9 | 4.70E+00 | 3.00E+00 | 4.7 | 5.1 |
| 0.00421 | 0.000482 | 1.15E-01 | 6.20E+00 | 6.2 | 5.80E+00 | 1.90E+00 | 6.2 | 6.2 |
| 0.668595 | 0.851639 | 1.00E+00 | | | | | | |
| 0.025754 | 0.000507 | 1.41E-02 | 4.50E+00 | 4.1 | 5.20E+00 | | 3.9 | 4.6 |
| 7.40E-12 | 5.45E-12 | 0.248777 | 5.6 | 5.2 | 5.2 | | | |
| 0.998093 | 0.565666 | 4.59E-01 | 1.80E+00 | 1.1 | 2.30E+00 | 4.00E-01 | 1.8 | 2.2 |
| 0.010858 | 0.006818 | 1.00E+00 | 5.20E+00 | 5.2 | 4.10E+00 | | 2.3 | 0.2 |
| 0.022715 | 5.45E-12 | 2.81E-11 | 5.6 | 5.6 | 4.8 | | 5 | 5.4 |

| | | | | | | | | |
|---|---|---|---|---|---|---|---|---|
| 0.53712 | 0.042873 | 0.27548 | 3 | 3.2 | 3.9 | | 1.1 | 0.5 |
| 7.81E-05 | 0.00016 | 0.063553 | 5.50E+00 | 5.5 | 4.4 | 5.3 | 6.4 | 6 |
| 0.035867 | 0.793698 | 0.019197 | 3.9 | 3.1 | 2.7 | | 3.3 | 4.1 |
| 0.120826 | 0.970213 | 1.93E-01 | 1.40E+00 | 4.50E+00 | 4.2 | 3.40E+00 | | 5.4 |
| 0.821558 | 0.07222 | 2.54E-01 | 3.90E+00 | 3.6 | 3.20E+00 | | 3.2 | 2.4 |
| 0.189944 | 0.738416 | 5.05E-01 | 1.40E+00 | 2.60E+00 | 1.8 | | 1.1 | 1.3 |
| 1 | 0.110384 | 1.40E-01 | 1.40E+00 | 1.1 | 7.00E-01 | | 1.4 | 2.6 |
| 1 | 0.752541 | 7.20E-01 | | 1.1 | | | 0.7 | |
| 0.589332 | 0.028269 | 1.62E-01 | 3.10E+00 | 2.7 | 2.70E+00 | | 1.4 | 4.8 |
| 0.668595 | 0.006832 | 0.029922 | 2.4 | | 4.8 | | 2 | 2.6 |
| 0.102534 | 0.001161 | 1.61E-02 | 1.10E+00 | 1.1 | 1.1 | 3.80E+00 | 3.9 | 3.5 |
| 0.142137 | 1 | 1.70E-01 | 1.2 | 5.5 | 5.1 | | 1.6 | 2 |
| 0.73174 | 0.001671 | 0.005837 | 2.60E+00 | 3.10E+00 | 2.3 | | 3.10E+00 | 3.5 |
| 0.509486 | 0.389928 | 1 | 9.00E-01 | 2.80E+00 | 1.4 | 4.00E-01 | 7.00E-01 | 1.3 |
| 0.137749 | 1.54E-05 | 7.00E-05 | 2.3 | | | 1.9 | | 1.5 |
| 0.00166 | 0.001207 | 1 | 6.6 | 6.6 | 6.6 | 3.8 | 1.7 | 1.5 |
| 0.991563 | 0.064205 | 0.145731 | 3.60E+00 | 2.50E+00 | 1.1 | 4.40E+00 | | 4.8 |
| 0.735789 | 0.653124 | 2.89E-01 | | 2 | 1.20E+00 | 0.4 | 1.6 | 2 |
| 0.993575 | 0.22438 | 0.451034 | | 1.20E+00 | 1.4 | | 1.10E+00 | 2.6 |
| 0.02822 | 0.000967 | 0.037678 | 2.30E+00 | 2 | 2 | | 2.3 | 2 |
| 0.395753 | 0.009 | 4.37E-03 | 6.00E-01 | 6.00E-01 | 2.00E-01 | | 1.70E+00 | 0.2 |
| 1 | 0.909572 | 1 | 0.9 | 0.9 | 0.9 | | 0.9 | |
| 0.345273 | 0.002135 | 0.014072 | 5.50E+00 | 5.80E+00 | 6.2 | 5.90E+00 | 4.3 | 4.3 |
| 1 | 0.122806 | 0.124858 | 3.3 | 2.2 | 2.7 | | 2.6 | 3.3 |
| 0.800843 | 0.007522 | 0.0066 | 2.10E+00 | 1.5 | 1.7 | 1.9 | 1.9 | 0.6 |
| 0.055128 | 0.000539 | 2.54E-04 | 2.00E-01 | 1.5 | 2.00E-01 | | 2.3 | 0 |
| 0.769205 | 0.000272 | 3.79E-04 | 4.00E-01 | 1.5 | | | 0 | 0 |
| 0.425882 | 0.245297 | 5.55E-02 | | | | | | |
| 1.77E-05 | 2.65E-05 | 2.76E-06 | 1.7 | 3.3 | 4 | | 5.2 | 5.6 |
| 0.002507 | 0.000212 | 4.74E-05 | 2.50E+00 | 2.1 | 4.4 | | 5.4 | 3.9 |
| 0.003409 | 0.003918 | 8.52E-01 | 5.80E+00 | 5.8 | 5.80E+00 | 2.30E+00 | 0.2 | 1.9 |
| 1 | 0.544728 | 0.578261 | 0.6 | 0.7 | | | | |
| 0.000175 | 7.28E-05 | 7.82E-01 | 6.6 | 6.20E+00 | 6.60E+00 | 5.2 | 0.8 | 2.5 |
| 0.000534 | 0.000425 | 9.92E-01 | 6.20E+00 | 6.60E+00 | 6.20E+00 | 4.90E+00 | 2.00E+00 | 2.4 |
| 0.186779 | 0.152978 | 1 | | 3.6 | 3.6 | 4.5 | 2.6 | 3.9 |
| 1 | 0.006443 | 0.011172 | 6.60E+00 | 6.6 | 6.2 | 1.9 | 6.2 | 6.2 |
| 0.076907 | 0.025615 | 0.913955 | 3.9 | 3.6 | 3.6 | 1.4 | 4.3 | |
| 2.67E-05 | 8.98E-06 | 0.02913 | 3.2 | 2.8 | 3.2 | 0 | 3.2 | 3.6 |
| 2.27E-05 | 3.34E-07 | 2.33E-05 | 6.20E+00 | 6.20E+00 | 6.20E+00 | 2.30E+00 | 6.20E+00 | 6.6 |
| 2.35E-05 | 0.192308 | 5.48E-05 | 4.50E+00 | 4.70E+00 | | 5.70E+00 | 6.20E+00 | 6.6 |
| 0.508889 | 0.00022 | 0.000818 | 6.2 | 6.2 | 6.6 | 2.7 | 6.2 | 6.2 |
| 0.738798 | 0.001104 | 0.003716 | 2 | 3 | 5.2 | | 3 | 3.3 |
| 0.017319 | 0.006126 | 0.915594 | 6.2 | 6.20E+00 | 6.2 | 1.5 | 2.00E-01 | 1 |
| 0.023343 | 1.32E-06 | 8.20E-06 | 6.6 | 6.60E+00 | 6.2 | 4.9 | 6.2 | 6.6 |
| 0.00238 | 0.001086 | 0.993771 | 4.7 | 5.8 | 5.5 | 0 | | |
| 0.000199 | 5.02E-05 | 0.194998 | 6.6 | 6.2 | 6.6 | 4.5 | 0.6 | 2.3 |
| 9.74E-06 | 5.71E-06 | 1 | 6.6 | 6.20E+00 | 6.6 | 1.9 | 5.6 | 2.9 |
| 0.814038 | 0.000272 | 3.93E-04 | 6.20E+00 | 6.60E+00 | 6.2 | 1.50E+00 | 5.80E+00 | 6.2 |
| 1.55E-05 | 7.18E-06 | 2.06E-01 | 5.80E+00 | 6.2 | 6.20E+00 | 3.40E+00 | 0.5 | |
| 1 | 0.071944 | 1.54E-01 | | 3.9 | 3.20E+00 | | | 4.3 |
| 0.001406 | 7.87E-06 | 8.59E-06 | 4.8 | 4.1 | 4.5 | 0.4 | 4.5 | 5.2 |

| | | | | | | | | |
|---|---|---|---|---|---|---|---|---|
| 0.001452 | 0.001871 | 6.59E-01 | 6.20E+00 | 6.60E+00 | 6.20E+00 | | 4.5 | 2.3 |
| 0.000163 | 6.94E-06 | 2.34E-04 | 6.60E+00 | 6.60E+00 | 5.80E+00 | | 5.60E+00 | 6 |
| 0.092249 | 0.000147 | 0.001093 | 2 | 2.4 | 2.4 | 3.1 | 4.3 | 4.3 |
| 4.31E-05 | 4.97E-05 | 0.232016 | 4.3 | 3.6 | 3.9 | | | 3.3 |
| 0.001042 | 1.19E-06 | 1.38E-05 | 5.1 | 5.5 | 5.5 | 1.1 | 5.5 | 5.5 |
| 0.403404 | 0.006921 | 0.0511 | 3.9 | 3.2 | 3.2 | 1.6 | 4.7 | 4.3 |
| 0.110021 | 0.057524 | 1 | 3.2 | 3.2 | 2.8 | | 4.4 | 2 |
| 0.001193 | 0.000163 | 7.63E-02 | 6.60E+00 | 6.60E+00 | 6.60E+00 | | 6.40E+00 | 6 |
| 0.005468 | 0.000394 | 0.049983 | 3.9 | 3.9 | 4.3 | | | |
| 0.013035 | 0.005336 | 0.978828 | 5.5 | 5.5 | 5.5 | 1.5 | 1.7 | 1 |
| 0.195821 | 0.310045 | 0.970531 | 5.8 | 5.80E+00 | 5.1 | 1.1 | 5.5 | 6.2 |
| 1.82E-06 | 1.03E-07 | 0.00067 | 5.50E+00 | 5.1 | 5.5 | | 2.7 | 3.1 |
| 0.000445 | 0.000251 | 1 | 6.2 | 6.2 | 6.6 | 3 | 2.6 | 1.6 |
| 0.019446 | 0.00073 | 0.034881 | 2 | 2.4 | 2.4 | | 4.3 | |
| 0.183679 | 0.002577 | 0.029788 | 1.3 | 3.2 | 1.3 | | 4.3 | 4.3 |
| 0.011211 | 1.60E-05 | 0.000159 | 6.6 | 6.6 | 5.8 | | 6 | 6.6 |
| 1 | 0.001004 | 1.63E-03 | 6.4 | 6.4 | 6 | 4.5 | 6.4 | 6.4 |
| 1.38E-05 | 5.45E-12 | 2.39E-10 | 6.2 | 6.2 | 6.2 | | 5.4 | 5.4 |
| 0.049642 | 0.010907 | 6.67E-01 | 2.60E+00 | 2.8 | 1.70E+00 | 4.50E+00 | 2.5 | 1.6 |
| 2.23E-05 | 1.52E-05 | 0.88469 | 4.7 | 4.3 | 2.4 | | | |
| 0.21969 | 0.001087 | 0.008712 | 5.6 | 6 | 5.6 | | 6 | 6 |
| 3.10E-05 | 4.34E-05 | 0.088272 | 3.9 | 4.3 | 3.9 | | | |
| 0.097099 | 0.155208 | 0.952093 | 5.8 | 6.2 | 6.2 | 0.8 | 5.1 | 5.5 |
| 0.642227 | 4.82E-05 | 0.000164 | 6.6 | 6.2 | 6.2 | | 6.6 | 6.2 |
| 0.020287 | 0.006307 | 0.848169 | 6.2 | 6.2 | 6.2 | 1.1 | 0.6 | |
| 0.000495 | 8.01E-07 | 1.69E-07 | 3.6 | 4.00E+00 | 2.90E+00 | | 4.00E+00 | 3.3 |
| 0.68335 | 0.210017 | 0.083628 | | | | 1.5 | 4.4 | |
| 0.007263 | 0.056934 | 1.16E-03 | 2.50E+00 | 3.5 | 3.90E+00 | | 3.4 | |
| 1 | 0.908894 | 9.79E-01 | 1.70E+00 | 1.6 | 0.7 | | 1.1 | 6.1 |
| 0.557358 | 0.000573 | 2.21E-03 | 3.30E+00 | 4.5 | 6.00E+00 | | 3.7 | 4.5 |
| 0.254232 | 0.005422 | 0.058237 | 4.3 | | 3.6 | 0.4 | 4.3 | 3.9 |
| 0.003236 | 0.001913 | 1 | 6.2 | 6.2 | 6.2 | | 1.4 | 0.6 |
| 1.26E-05 | 4.56E-06 | 0.039478 | 5.1 | 5.5 | 4.7 | | 2 | 2.6 |
| 0.008396 | 1.16E-05 | 1.77E-05 | 5.6 | 5.6 | 4.8 | | 5.8 | 6.6 |
| 2.50E-06 | 2.42E-06 | 1.71E-01 | 4.70E+00 | 4.7 | 3.9 | | | |
| 0.293353 | 0.022337 | 0.268883 | 3.6 | 3.6 | 3.6 | | 3.6 | 3.9 |
| 0.028768 | 0.000115 | 1.41E-03 | 5.50E+00 | 5.8 | 6.6 | 2.3 | 4.2 | 5.4 |
| 0.813497 | 0.16731 | 0.087926 | 2 | 3.2 | 3.1 | | 3.6 | 2.8 |
| 0.018364 | 0.003805 | 0.551074 | 3.2 | 2.8 | 3.2 | 1.5 | 2.9 | 2.3 |
| 1.50E-05 | 8.26E-06 | 0.978828 | 6 | 5.6 | 5.2 | | | 1.5 |
| 0.813497 | 0.096337 | 0.336798 | 4.00E+00 | 6.6 | 5 | | 3.9 | 5 |
| 0.015699 | 0.001142 | 2.56E-04 | | | | | | 0.7 |
| 0.000489 | 0.0005 | 7.42E-01 | 4.70E+00 | 3.9 | 3.60E+00 | | | |
| 0.000163 | 0.000453 | 0.051142 | 4.7 | 4.7 | 4.7 | | 3.3 | 1.8 |
| 0.000342 | 0.000256 | 1 | 6.2 | 6.2 | 6.2 | 3 | 5.6 | 3.3 |
| 0.032883 | 4.72E-06 | 8.11E-06 | 2.6 | 2.6 | 3.3 | | 4.8 | 5.2 |
| 4.23E-05 | 1.10E-05 | 0.033327 | 5.8 | 5.5 | 5.8 | | 4.2 | 3.9 |
| 0.017156 | 0.000668 | 0.033974 | 5.20E+00 | 6.00E+00 | 5.2 | | 3.5 | 3.7 |
| 0.063758 | 0.000306 | 3.46E-03 | 6.20E+00 | 6.6 | 5.80E+00 | | 5.6 | 4.1 |
| 0.345476 | 0.002201 | 0.001385 | 6 | 5.2 | 3.3 | | 4.2 | 5.8 |
| 0.001001 | 0.000341 | 0.652548 | 5.6 | 5.8 | 5.8 | | 0.2 | 4.5 |

| | | | | | | | | |
|---|---|---|---|---|---|---|---|---|
| 0.373389 | 4.46E-05 | 6.53E-05 | 3.30E+00 | 3 | 3.70E+00 | | 3.3 | 3 |
| 0.031097 | 0.001865 | 0.100363 | 5.5 | 5.8 | 5.1 | | | |
| 4.49E-05 | 2.97E-05 | 1 | 6 | 6.2 | 6.6 | | 1.1 | 2.7 |
| 0.079056 | 0.044085 | 1 | 2.4 | 0.8 | | | 5.2 | |
| 0.987941 | 0.007139 | 0.01732 | 5.6 | 6.4 | 6 | 1.9 | 6 | 5.6 |
| 0.1199 | 0.040004 | 0.917136 | 3.2 | | 3.6 | | | |
| 0.609519 | 0.005006 | 0.02334 | 2 | 2 | 1.7 | 4.5 | 2.8 | |
| 0.013281 | 0.004923 | 0.932398 | 6.4 | 5.6 | 5.6 | | 0.4 | 1.5 |
| 0.743655 | 0.000269 | 0.000369 | 6.00E+00 | 5.6 | 4.8 | | 5.6 | 6 |
| 0.057147 | 0.171603 | 0.644887 | | | 3.6 | 2.3 | | 1.2 |
| 2.79E-05 | 4.87E-05 | 0.037963 | 5.6 | 5.6 | 5.6 | | | |
| 0.007307 | 0.001551 | 0.468706 | 5.1 | 5.1 | 5.1 | | 0.2 | 0.2 |
| 0.045132 | 1 | 0.049301 | 1.4 | 3.9 | 4.3 | 1.9 | | 0.5 |
| 0.382749 | 0.000515 | 0.00047 | 6.2 | 6.6 | 6.2 | 0.8 | 6.2 | 6.6 |
| 0.000373 | 0.00011 | 3.51E-01 | 6.00E+00 | 5.60E+00 | 5.60E+00 | | 0.4 | 3 |
| 0.001455 | 0.001405 | 9.09E-01 | 3.70E+00 | 4.1 | 3.30E+00 | | | 2.1 |
| 0.697078 | 0.017986 | 8.10E-02 | 3.30E+00 | 4.7 | 9.00E-01 | 1.00E+00 | 1.4 | 1.1 |
| 0.000153 | 9.48E-06 | 0.000859 | 2.9 | 2.5 | 3.6 | | 5.4 | 3.9 |
| 0.001332 | 7.56E-06 | 8.05E-06 | 3.5 | 5 | 3.5 | | 6 | 6 |
| 6.39E-06 | 5.45E-12 | 2.22E-11 | 6 | 6 | 5.6 | | 4.2 | 4.2 |
| 0.000231 | 9.57E-06 | 5.61E-04 | 5.10E+00 | 4.7 | 4.30E+00 | | 4.8 | 4.5 |
| 0.092249 | 0.090561 | 8.13E-03 | 4.50E+00 | 5.2 | 5.2 | | 5.2 | 6 |
| 0.000765 | 0.000109 | 0.064388 | 3.9 | 5.1 | 4.3 | | | |
| 8.42E-05 | 9.58E-07 | 2.22E-05 | 6 | 5.6 | 5.6 | 4.9 | 5 | 3.6 |
| 6.04E-06 | 6.90E-06 | 0.009328 | 3.2 | 5.1 | 3.6 | | 1.2 | 2.3 |
| 0.001403 | 0.000292 | 0.242834 | 4.8 | 4.8 | 5.6 | | | |
| 0.593774 | 0.001785 | 0.007623 | 5 | 6.1 | 5.8 | 4.9 | 6.1 | 6.1 |
| 0.097687 | 0.000342 | 3.17E-03 | 4.7 | 4.10E+00 | 4.5 | | 4.5 | 2.6 |
| 0.583578 | 0.014636 | 0.081649 | 3.4 | 1.9 | 0.2 | 4.9 | 0.8 | 4.5 |
| 0.328803 | 0.71284 | 0.804672 | | | | | 1.7 | 2 |
| 0.291991 | 0.001487 | 0.01044 | 2 | 2.4 | | | 0.7 | 2.8 |
| 0.003256 | 0.000478 | 0.169037 | 2.40E+00 | 2.4 | 1.7 | | | |
| 1 | 0.2351 | 0.270988 | 5.00E-01 | 1.7 | 2.1 | 1.10E+00 | 3 | 3.6 |
| 0.868728 | 0.072007 | 2.29E-01 | 1.1 | 7.00E-01 | 2.00E+00 | 4.5 | 1.10E+00 | 2.2 |
| 0.921404 | 0.656171 | 4.30E-01 | 4.60E+00 | 2.2 | 2.2 | | 3.3 | 2.6 |
| 1.62E-05 | 7.51E-06 | 0.172118 | 4.8 | 4.5 | 5.2 | | 1.5 | |
| 0.507762 | 0.000126 | 0.000462 | 5.80E+00 | 5.5 | 4.7 | 0.6 | 5.1 | 4.3 |
| 0.00163 | 0.322434 | 0.005466 | 5 | 6 | 4.6 | | 5.5 | 5.5 |
| 0.651342 | 0.008126 | 3.73E-02 | 6.00E-01 | 4.5 | 4.50E+00 | | 2.9 | 2.3 |
| 0.026267 | 0.000393 | 9.36E-03 | 1.70E+00 | 2 | 1.30E+00 | | 1.4 | 1.3 |
| 0.626139 | 1.58E-05 | 5.04E-05 | 4.8 | 4.1 | 4.8 | | 4.8 | 5.6 |
| 0.004668 | 9.14E-05 | 0.003701 | 5.00E+00 | 5.4 | 5 | | 3.3 | 4 |
| 0.134757 | 7.98E-06 | 1.67E-05 | 2.70E+00 | 4.20E+00 | 3.4 | | 1.90E+00 | 3 |
| 0.000461 | 0.000357 | 1 | 5.60E+00 | 4.1 | 4.5 | | 4.5 | 1.5 |
| 0.578048 | 5.53E-05 | 0.000197 | 4.3 | 3.2 | 3.2 | | | 3.6 |
| 1 | 0.038414 | 0.050551 | 3.70E+00 | 4.10E+00 | 4.8 | | 5.60E+00 | 4.5 |
| 1 | 0.332995 | 0.334868 | 3.3 | 3.00E+00 | 4.1 | 2.3 | 3.6 | 3.2 |
| 0.436752 | 0.771651 | 8.95E-01 | 2.80E+00 | | | 8.00E-01 | 1.40E+00 | 0.7 |
| 0.000507 | 1.11E-05 | 0.000467 | 4.70E+00 | 3.70E+00 | 4.8 | | 2.70E+00 | 2.7 |
| 0.00763 | 0.010821 | 0.799087 | 4.5 | 4.5 | 4.5 | | 4.5 | 4.5 |
| 0.002837 | 0.000631 | 3.82E-01 | 4.5 | 3.7 | 3.7 | | 0.2 | 1.5 |

| | | | | | | | | |
|---|---|---|---|---|---|---|---|---|
| 0.111235 | 1.72E-06 | 8.20E-06 | 5 | 5.4 | 5.4 | | 5.4 | 4.6 |
| 1 | 0.005612 | 0.007964 | 4.1 | 4.8 | 4.8 | 3.8 | 4.8 | 4.8 |
| 0.99774 | 0.129687 | 0.268692 | 4.8 | 3 | 4.7 | | 4.5 | 2.6 |
| 0.593872 | 0.247492 | 0.934322 | 1.7 | 2.2 | 3 | | | |
| 0.000125 | 0.000508 | 0.020851 | 4.50E+00 | 4.50E+00 | 3.7 | | 1.50E+00 | 4.5 |
| 0.044771 | 0.113282 | 0.707313 | 5.2 | 5.2 | 4.8 | 4.5 | 6.2 | 5.8 |
| 0.1199 | 0.886695 | 0.067618 | 3.5 | 2.6 | 3 | 4.5 | 4.1 | 4.8 |
| 0.000328 | 9.48E-06 | 0.000378 | 4.50E+00 | 4.8 | 4.5 | | 3.9 | 3.1 |
| 0.597859 | 1.83E-05 | 3.69E-05 | 5.5 | 5.4 | 5.8 | | 5.8 | 5.8 |
| 0.001013 | 4.79E-05 | 5.65E-03 | 5.80E+00 | 6.2 | 5.5 | | 4.8 | 4.5 |
| 0.991563 | 0.000947 | 0.002237 | 3.9 | 3.6 | 4.1 | | 4.1 | 5.6 |
| 0.002837 | 3.09E-05 | 0.000906 | 5.1 | 4.7 | 4.3 | | 4.8 | 4.5 |
| 0.146806 | 0.002185 | 2.85E-02 | 4.5 | 3.70E+00 | 3.70E+00 | | 1.20E+00 | 2 |
| 0.29657 | 0.069938 | 0.015495 | 5.8 | 5 | 4.2 | | 5.4 | 5.8 |
| 0.084162 | 0.062923 | 1 | 3.7 | 3.6 | 3.7 | | 4.2 | |
| 0.418317 | 0.006104 | 4.23E-02 | 2.00E+00 | 2.8 | 2.00E+00 | | 3.2 | 3.2 |
| 0.666512 | 0.617147 | 1.00E+00 | 5.00E+00 | 5.4 | 4.80E+00 | 4.50E+00 | 5.8 | 6.1 |
| 0.000335 | 4.25E-05 | 0.027872 | 5.80E+00 | 5.40E+00 | 5.8 | | 5.20E+00 | 4 |
| 0.002026 | 0.032213 | 4.11E-04 | 1.90E+00 | 4.5 | 6.00E-01 | | 5.4 | 5.8 |
| 0.169204 | 5.01E-05 | 0.000262 | 4.5 | 3.3 | 4.5 | | 3 | 3 |
| 1 | 0.000322 | 0.000601 | 5.2 | 5.2 | 5.2 | | 5.2 | 5.2 |
| 0.973827 | 0.001312 | 0.003308 | 3.5 | 3.9 | 3.1 | | 5.6 | 2 |
| 0.04478 | 0.008925 | 0.601847 | 3.3 | 4.8 | 3 | | 0.2 | 2.3 |
| 0.012722 | 0.001178 | 0.137622 | 5 | 3.7 | 5 | | 1.5 | 0.2 |
| 0.020905 | 0.170928 | 0.218018 | 4.6 | 3.9 | 4.2 | 4.5 | 5.8 | 5.2 |
| 0.696448 | 0.004573 | 0.018265 | 4.5 | 5.20E+00 | 4.8 | | 5.20E+00 | 5.2 |
| 0.014399 | 0.000481 | 0.000145 | 0.8 | 1.9 | 2.3 | | 2.7 | 1.9 |
| 0.026305 | 7.54E-06 | 3.23E-05 | 5.2 | 4.8 | 4.8 | | 5.8 | 3.9 |
| 3.63E-05 | 1.06E-05 | 4.15E-02 | 4.1 | 4.10E+00 | 3.7 | | 0.8 | 1.5 |
| 0.9582 | 0.00076 | 1.09E-03 | 5.4 | 4.60E+00 | 5.40E+00 | | 5.60E+00 | 5.2 |
| 0.579337 | 0.205139 | 6.59E-02 | 5.00E+00 | 5.4 | 5.80E+00 | | 6.1 | 5.4 |
| 0.007935 | 5.66E-05 | 3.52E-05 | 2.90E+00 | 4.40E+00 | 4.00E+00 | | 3.90E+00 | 4.2 |
| 0.021196 | 0.025827 | 0.001556 | 5.2 | 5.2 | 5 | | 5.4 | 5.8 |
| 0.761877 | 0.845416 | 0.443092 | 1.3 | 0.9 | 0.7 | 1 | 1.1 | 2.4 |
| 0.635347 | 0.000644 | 0.002317 | 3.9 | 3.5 | 5.2 | | 4.2 | 4.6 |
| 0.095716 | 0.002289 | 0.044228 | 3.7 | 3 | 4.1 | 3 | 3.3 | 1.4 |
| 0.150323 | 0.089867 | 1.00E+00 | 4.1 | 3.70E+00 | 3.70E+00 | 0.8 | 2.00E-01 | 4.5 |
| 0.040425 | 0.710701 | 1.14E-01 | 2.7 | 1.20E+00 | 1.60E+00 | 3.8 | 3.70E+00 | 3.3 |
| 0.00417 | 0.003813 | 9.91E-01 | 9.00E-01 | 0.9 | 9.00E-01 | | 1.1 | 1.4 |
| 0.481252 | 0.622895 | 0.16025 | 1.1 | 0.7 | 2.2 | | 3.2 | 1.1 |
| 0.991563 | 5.54E-06 | 1.66E-05 | 5.40E+00 | 5.00E+00 | 5.8 | | 5.40E+00 | 4.6 |
| 0.005899 | 0.010592 | 0.000492 | 4.8 | 5.9 | 5.5 | 5.3 | 5.5 | 5.5 |
| 0.023857 | 0.013612 | 1 | 4.5 | 4.5 | 4.8 | | 3.8 | 0.2 |
| 0.008143 | 0.012852 | 0.743723 | 2.8 | 3 | 3 | | 1.2 | 1.2 |
| 0.458934 | 0.037241 | 0.01385 | 1.7 | 0.2 | 0.2 | | 4.5 | 1.5 |
| 0.079478 | 2.04E-05 | 0.000118 | 5.3 | 3.80E+00 | 1.9 | 4.9 | 4.9 | 5.3 |
| 0.10446 | 0.00058 | 0.006114 | 5.80E+00 | 5.6 | 6 | | 3.3 | 4.2 |
| 0.811164 | 0.040774 | 0.150855 | 1.5 | 1 | 0.8 | 2.7 | 0.2 | 1 |
| 0.846469 | 0.088358 | 2.91E-01 | 2.90E+00 | 2.4 | 1.70E+00 | 2.10E+00 | 2.6 | 2.7 |
| 0.008871 | 0.006215 | 1 | 3.70E+00 | 3.7 | 4.1 | | 0.6 | 1 |
| 0.007506 | 1.06E-05 | 8.73E-05 | 4.60E+00 | 5.90E+00 | 5.5 | 4.90E+00 | 5.90E+00 | 5.5 |

| | | | | | | | | |
|---|---|---|---|---|---|---|---|---|
| 0.006868 | 0.009454 | 0.803001 | 4.5 | 4.5 | 4.5 | | 4.5 | 4.5 |
| 0.934561 | 0.002975 | 0.008225 | 4.50E+00 | 4.50E+00 | 4.9 | | 4.9 | 3.8 |
| 0.187705 | 0.009552 | 1.62E-01 | 3.70E+00 | | 2.70E+00 | 2.10E+00 | 4.80E+00 | 4.8 |
| 0.063429 | 0.106839 | 0.006975 | 3.70E+00 | 3.7 | 3.3 | | 3.9 | 3.9 |
| 0.404664 | 0.00373 | 0.002237 | 5.4 | 4.2 | 5.8 | | 5 | 4.2 |
| 1 | 0.18577 | 1.68E-01 | 3.30E+00 | 2.6 | 2.30E+00 | | 3.7 | 2.2 |
| 1 | 0.116801 | 0.140768 | 2.1 | 2.9 | 2.5 | | 2.5 | 2.5 |
| 0.97027 | 0.158016 | 0.363396 | 3 | 3.3 | 3.3 | | 4.6 | 3 |
| 1 | 0.019037 | 0.036903 | 3 | 2.6 | 3.7 | | 2.6 | 4.5 |
| 1.39E-05 | 3.23E-05 | 0.001495 | 1.4 | 3.3 | 4.5 | 3 | 5.8 | 5 |
| 0.000163 | 5.40E-06 | 0.00011 | 3.3 | 3.3 | 4.5 | 0.8 | 3.9 | 5 |
| 0.905329 | 0.982694 | 0.77085 | 2.40E+00 | 2.20E+00 | 1.8 | 4.00E-01 | 2.80E+00 | 4.1 |
| 0.831772 | 2.92E-08 | 6.70E-08 | | | 1.5 | | | |
| 0.577347 | 0.301182 | 0.092871 | 1 | 2.5 | 2.3 | | 3.5 | 3.9 |
| 0.041853 | 1 | 0.0363 | 5.4 | 5.4 | 5.8 | | 3.9 | 5.5 |
| 1 | 0.993629 | 0.944687 | | | | | | |
| 0.035123 | 0.000923 | 0.028021 | 5.4 | 5.40E+00 | 5.4 | | 4 | 4 |
| 0.123299 | 0.001233 | 0.015218 | 4.5 | 5.6 | 4.5 | | 4.8 | 4.1 |
| 0.760633 | 0.008396 | 0.032118 | 4.8 | 4.1 | 4.5 | | 4.5 | 4.5 |
| 1 | 0.953142 | 0.890219 | 2.5 | 5.2 | 3.3 | 4.8 | 2.3 | 2.3 |
| 0.577949 | 7.27E-05 | 0.000113 | 5.20E+00 | 5.50E+00 | 5.2 | | 5.40E+00 | 4.4 |
| 0.114978 | 0.042343 | 9.55E-01 | 3.70E+00 | 2.6 | 3.30E+00 | | 4.5 | |
| 0.503893 | 0.491834 | 0.129751 | 3.1 | 1.6 | 1.2 | | 2 | 3.9 |
| 0.003088 | 2.58E-05 | 0.000647 | 3.30E+00 | 3.30E+00 | 2.6 | | 3.90E+00 | 3.9 |
| 0.00948 | 0.002359 | 0.615825 | 4.2 | 3.5 | 4.6 | | 1.5 | |
| 0.130847 | 0.012439 | 0.310644 | 2.6 | 2 | 2.9 | | 4.5 | 1.4 |
| 0.022401 | 0.001009 | 0.051837 | 1.40E+00 | 3.70E+00 | 3.9 | | 2.70E+00 | 2 |
| 6.99E-05 | 2.33E-05 | 8.20E-06 | 1.00E+00 | 1.00E+00 | 1.4 | | 3.00E+00 | 3.3 |
| 0.000328 | 9.48E-06 | 0.00038 | 4.5 | 4.8 | 4.5 | | 3.9 | 3.1 |
| 0.268273 | 0.00031 | 0.001632 | 1 | 2.1 | 4.4 | | 1.7 | 2.9 |
| 0.470909 | 0.163463 | 0.044087 | 0.6 | 1.6 | 0.6 | 1.7 | | |
| 0.861588 | 7.58E-08 | 2.50E-07 | 4.60E+00 | 5.00E+00 | 5.4 | | 5 | 4.6 |
| 1.07E-09 | 5.45E-12 | 2.22E-11 | 4.8 | 4.5 | 4.8 | | 2.1 | 1 |
| 0.22674 | 0.042455 | 0.009196 | 1.30E+00 | 2.00E+00 | 1.7 | | 2.00E+00 | 2 |
| 0.226933 | 0.18836 | 1 | 1.8 | 3 | 2.8 | | 0.6 | 1.1 |
| 0.015461 | 0.021966 | 0.882147 | 2.20E+00 | 2.70E+00 | 3.5 | | 4.20E+00 | 3.8 |
| 0.115136 | 0.019446 | 0.563182 | 3.50E+00 | 2.70E+00 | 2.7 | | 4.8 | 2.1 |
| 0.752642 | 0.009617 | 0.037656 | 0.6 | 4.5 | 1.5 | | 2.9 | 2.3 |
| 1 | 0.012852 | 0.024456 | 1.8 | 2.6 | 2.6 | | 3.3 | 3.7 |
| 0.000258 | 1.91E-05 | 0.004878 | 3.9 | 3.5 | 3.9 | | 3.3 | 5.2 |
| 0.562718 | 0.02262 | 0.137248 | 2.7 | 2.3 | 3.9 | | 5.2 | 4.2 |
| 1 | 0.000324 | 5.90E-04 | 5.00E+00 | 4.20E+00 | 4.6 | | 5.00E+00 | 4.2 |
| 0.169204 | 0.971105 | 1.20E-01 | | | | | 4.5 | |
| 0.778676 | 0.001235 | 0.004013 | 3.1 | 4.2 | 2.7 | 2.3 | 4.2 | 3.9 |
| 0.000582 | 2.98E-05 | 0.00419 | 3.9 | 4.2 | 3.5 | | 3.6 | 2.1 |
| 0.953996 | 0.003683 | 0.004398 | 3.40E+00 | | 4.9 | | 3.00E+00 | 5.3 |
| 0.889533 | 0.180067 | 0.502468 | 3.10E+00 | 2.70E+00 | 0.7 | 0.00E+00 | 1.80E+00 | 1 |
| 0.000262 | 7.89E-06 | 0.000251 | 2.1 | 1.7 | 1 | | 3.5 | 3.5 |
| 0.26815 | 1.40E-07 | 6.38E-07 | 5.4 | 5.8 | 4.6 | | 5.4 | 5.4 |
| 0.102534 | 1 | 0.105068 | 0.2 | 0.2 | 4.5 | | 3.1 | 4.2 |
| 0.009349 | 0.002233 | 0.587319 | 3.5 | 4.6 | 3.9 | | 1.9 | 0.2 |

| | | | | | | | | |
|---|---|---|---|---|---|---|---|---|
| 0.295637 | 3.48E-08 | 5.90E-08 | 3.9 | 4.2 | 4.6 | | 4.6 | 4.6 |
| 0.008501 | 0.000132 | 0.004042 | 4.1 | 3.3 | 3.7 | | 1.7 | 2.7 |
| 1 | 0.002199 | 0.004179 | 1.4 | 1.1 | 2.2 | 0 | 2.6 | 1.4 |
| 0.002231 | 0.82823 | 0.003973 | 1.8 | 3 | 1.4 | | 1.8 | 2.6 |
| 1 | 0.813945 | 0.979151 | | 2 | | | 2.7 | 1.4 |
| 0.002205 | 4.13E-05 | 0.001804 | 5 | 4.2 | 4.8 | | 4 | 2.9 |
| 0.27132 | 0.15311 | 1 | 3.5 | 3 | 3 | | 0.2 | |
| 0.616712 | 0.000184 | 0.000623 | 4.6 | 5.4 | 4.6 | | 5.4 | 4.2 |
| 0.000617 | 0.000216 | 0.606219 | 2 | 0.8 | 0.8 | | | 0.4 |
| 0.447553 | 7.56E-06 | 1.72E-05 | | | | | | |
| 9.78E-12 | 5.45E-12 | 0.074238 | 3.1 | 3.5 | 5 | | | |
| 0.587972 | 0.319891 | 0.101601 | 3.60E+00 | 2.10E+00 | 3.6 | | 4.00E+00 | |
| 0.055128 | 0.001894 | 0.056118 | 4.5 | 4.1 | 4.5 | | 2.1 | 2.3 |
| 0.295205 | 0.256522 | 1 | 1.4 | 2.4 | 2.4 | | | 1.1 |
| 0.473196 | 5.30E-06 | 1.68E-05 | 3.7 | 3.7 | 4.1 | | 4.1 | 4.1 |
| 0.852849 | 0.012746 | 0.042258 | | | 0.6 | 0 | | |
| 0.002747 | 0.000187 | 0.026414 | 2.7 | 2.3 | 3.9 | | 5.2 | 1 |
| 0.000507 | 1.11E-05 | 0.000467 | 4.7 | 3.7 | 4.8 | | 2.7 | 2.7 |
| 0.004189 | 0.000529 | 9.51E-05 | 2.9 | 2.9 | 5.2 | | 3.9 | 4.2 |
| 0.016021 | 0.015235 | 1 | 2.3 | 3.1 | 2.6 | | 3 | 1.9 |
| 0.661681 | 0.004371 | 0.003522 | 1.5 | 1.1 | 0.2 | | 1.5 | 1.5 |
| 0.898286 | 1.68E-05 | 3.81E-05 | 4 | 3.6 | 4 | 4.5 | 4.4 | 5.5 |
| 0.014057 | 0.000709 | 0.047326 | 1.8 | 3.00E+00 | 1.8 | | 1.6 | 0.5 |
| 0.121111 | 0.051858 | 0.996152 | 3.00E+00 | 2.70E+00 | 3.5 | 1.5 | 1.4 | |
| 1 | 0.139743 | 0.195472 | 4.4 | 2.3 | 0.2 | | 1 | 0.2 |
| 0.024985 | 0.006832 | 7.66E-01 | 4.60E+00 | 5.00E+00 | 3.50E+00 | | 1.90E+00 | 1.5 |
| 0.024426 | 0.000597 | 0.019359 | 4.20E+00 | 3.90E+00 | 5 | | 5.20E+00 | 4.8 |
| 1 | 0.63726 | 0.640292 | 3.1 | 3.9 | | | 3.7 | 3.3 |
| 0.996169 | 0.002929 | 0.003966 | | | | 1.9 | | 1.5 |
| 0.01092 | 7.48E-07 | 5.66E-07 | 2.90E+00 | 1.4 | 2 | | 3.1 | 4.6 |
| 0.511975 | 0.059495 | 0.398282 | 2 | 2.7 | 2.7 | | 0.6 | 0.5 |
| 0.734618 | 0.776343 | 3.65E-01 | 5.00E+00 | 5.00E+00 | 4.20E+00 | | 4.6 | 4.6 |
| 6.92E-10 | 2.40E-10 | 2.74E-01 | 3.90E+00 | 5.00E+00 | 3.90E+00 | | | |
| 0.919013 | 0.000144 | 0.000388 | 5.00E+00 | 3.10E+00 | 3.5 | | 3.90E+00 | 3.9 |
| 0.343467 | 0.003352 | 2.46E-02 | 4.80E+00 | 3.70E+00 | 4.10E+00 | | 3.5 | 3.1 |
| 0.046363 | 0.00012 | 8.41E-05 | 4.80E+00 | 4.40E+00 | 5.20E+00 | | 5.9 | 5.9 |
| 0.258351 | 0.092173 | 0.016983 | 3.90E+00 | 2.3 | 3.6 | | 4.6 | 5 |
| 0.224042 | 0.274876 | 1.00E+00 | 2.20E+00 | 1.40E+00 | 2.20E+00 | | 0.8 | |
| 0.000551 | 0.000417 | 1.00E+00 | 5.20E+00 | 5.50E+00 | 5.20E+00 | | 3.4 | |
| 0.912498 | 0.004474 | 1.28E-02 | 5.20E+00 | 5.00E+00 | 3.50E+00 | | 3.1 | 3.3 |
| 1 | 0.000132 | 2.83E-04 | 4.60E+00 | 4.2 | 3.9 | | 5 | 5.4 |
| 1 | 0.000296 | 5.30E-04 | 2.70E+00 | 2.30E+00 | 3.10E+00 | | 3.50E+00 | 3.5 |
| 0.023573 | 0.000606 | 2.03E-04 | 1.40E+00 | | 2.00E-01 | 0.00E+00 | | |
| 5.92E-05 | 7.54E-06 | 1.05E-03 | 1.20E+00 | 1.20E+00 | 2.30E+00 | 3.40E+00 | 1.7 | 1.7 |
| 0.064939 | 1 | 7.07E-02 | 4.80E+00 | 3.60E+00 | 4.00E+00 | | 4.6 | 4.6 |
| 0.233284 | 3.44E-05 | 0.00016 | 3.9 | 3.1 | 3.1 | | 4.4 | 3.5 |
| 0.11559 | 0.005408 | 1.25E-01 | 3.60E+00 | 2.70E+00 | 3.90E+00 | | 4.00E+00 | 5.2 |
| 1 | 0.010179 | 1.74E-02 | 1.40E+00 | 1.1 | 2.2 | 0.00E+00 | 2.6 | 1.4 |
| 0.003267 | 0.001517 | 1.00E+00 | 3.90E+00 | 2.3 | 3.10E+00 | | 0.8 | 4.5 |
| 0.16422 | 0.6067 | 5.32E-02 | 2.7 | 2 | 2 | | 2.7 | 4.5 |
| 1 | 0.057996 | 0.065433 | 2.3 | 2 | 1.6 | 0 | 2.1 | 2.7 |

| | | | | | | | | |
|---|---|---|---|---|---|---|---|---|
| 0.452512 | 0.020448 | 1.56E-01 | 3.10E+00 | 3.1 | 3.1 | | 2.7 | 2.3 |
| 0.323524 | 0.001637 | 0.001072 | 3.8 | 3 | 3.8 | 2.3 | | 0.8 |
| 0.008372 | 0.002102 | 0.612019 | 2.3 | 0.5 | 2 | 1.5 | 4.9 | 0.2 |
| 0.048163 | 4.25E-05 | 3.31E-04 | 4.60E+00 | 4.2 | 4.60E+00 | | 5.2 | 4.8 |
| 0.711209 | 0.00136 | 1.40E-03 | 3.00E+00 | 1.50E+00 | 4.00E-01 | | 1.5 | 4.9 |
| 0.979945 | 0.734413 | 1 | 0.8 | 1.6 | 2.2 | | 0.8 | 1.2 |
| 0.258402 | 0.586815 | 0.807098 | | 1.7 | | | 2.4 | 2 |
| 0.02002 | 0.014039 | 1 | 4.8 | 4.8 | 5.2 | | 2.3 | 0.2 |
| 3.54E-05 | 1.58E-06 | 4.55E-05 | 4.6 | 4.2 | 3.9 | | 1.5 | 1.5 |
| 0.1959 | 0.628982 | 0.617817 | 1.2 | 0.8 | 1.6 | 1.5 | | |
| 0.002741 | 0.000599 | 3.61E-01 | 3.10E+00 | 3.9 | 2.70E+00 | | 4.5 | 0 |
| 0.916613 | 0.022899 | 1.97E-02 | 2.70E+00 | 2.9 | 2.10E+00 | | 2.7 | 3.6 |
| 0.247598 | 0.0002 | 1.97E-04 | 1.10E+00 | 0.4 | 4.50E+00 | | 4.5 | |
| 0.311547 | 0.001384 | 9.20E-03 | 5.20E+00 | 4 | 3.30E+00 | | 2.5 | 4 |
| 0.330504 | 0.011742 | 0.117039 | 4.5 | 3.3 | 2.9 | | 4 | 5.5 |
| 0.048242 | 0.000683 | 0.013386 | 3.7 | 3.3 | 3.3 | | 4.5 | 3.1 |
| 0.920702 | 0.140465 | 3.78E-01 | 4.40E+00 | 3.6 | 4.80E+00 | | 3.9 | 5.2 |
| 0.000996 | 0.000192 | 1.67E-01 | 2.70E+00 | 3.10E+00 | 3.10E+00 | | | |
| 0.002096 | 3.03E-06 | 1.99E-05 | 4.00E+00 | 4 | 3.30E+00 | | 3.6 | 2.9 |
| 0.077513 | 0.004417 | 1.40E-01 | 2.20E+00 | 2.2 | 1.80E+00 | | 0.6 | |
| 1 | 0.032107 | 0.071045 | 2.9 | 4.4 | 2.9 | | 5.5 | 3.6 |
| 0.041937 | 0.012442 | 8.41E-01 | 1.40E+00 | 2.20E+00 | 1.80E+00 | | | |
| 0.457615 | 1 | 4.33E-01 | 4.40E+00 | 4.8 | 5.20E+00 | | 3.1 | 4.8 |
| 0.037139 | 3.19E-05 | 0.000259 | 4 | 5.2 | 4.8 | | 4.8 | 5.9 |
| 0.505996 | 0.155906 | 4.57E-02 | 1.90E+00 | | 8.00E-01 | | 1.9 | 2.3 |
| 0.823627 | 1.93E-05 | 5.88E-05 | 3.30E+00 | 2.9 | 3.60E+00 | | 3.3 | 3.6 |
| 0.01002 | 0.003257 | 0.838408 | 1.40E+00 | 1.10E+00 | 1.4 | | 6.00E-01 | 0.2 |
| 1 | 0.047175 | 5.48E-02 | 2.90E+00 | 4.80E+00 | 4.00E+00 | | 3.30E+00 | 3.3 |
| 0.079108 | 0.23701 | 6.67E-01 | 2.3 | 2.30E+00 | 2.30E+00 | | 1.90E+00 | 3 |
| 0.000625 | 5.11E-07 | 9.10E-06 | 4.8 | 4.8 | 4.8 | | 3 | 5.2 |
| 0.000107 | 5.02E-05 | 0.941932 | 3.50E+00 | 4.2 | 4.2 | | | |
| 0.100776 | 0.11688 | 1.00E+00 | 3.9 | 4.3 | 3.9 | | 1.8 | 3.9 |
| 1 | 0.616842 | 9.15E-01 | 2.40E+00 | 1.80E+00 | 3.30E+00 | | 2.8 | 3.3 |
| 0.682604 | 0.078225 | 0.0363 | | 2 | 0.8 | | | 2.6 |
| 0.014935 | 0.001797 | 0.228605 | 3.70E+00 | 4.10E+00 | 4.8 | | | |
| 0.412169 | 0.405336 | 1 | 0.5 | 7.00E-01 | | | | |
| 1 | 0.813581 | 0.863651 | | 1.1 | | | 1.4 | 0.7 |
| 2.15E-05 | 7.57E-06 | 2.19E-02 | 6.20E+00 | 6.2 | 5.80E+00 | | | 1 |
| 1 | 0.92329 | 1 | 2.7 | 2.6 | 3.9 | | 4.6 | 2 |
| 0.705515 | 2.79E-06 | 9.16E-06 | 2.80E+00 | 1.20E+00 | 2.40E+00 | 2.40E+00 | 1.4 | 2.1 |
| 0.860682 | 0.780721 | 1 | 3.90E+00 | 2.8 | 3.6 | 0.2 | | 0.7 |
| 0.970638 | 0.53225 | 0.905843 | | | 1.7 | | | |
| 1 | 0.000224 | 0.000451 | 4.50E+00 | | 1.9 | 4.50E+00 | 3.4 | 1.9 |
| 0.89567 | 0.000842 | 0.001108 | | 1.90E+00 | | | | |
| 0.000206 | 2.83E-05 | 0.025276 | 6.2 | 6.2 | 6.6 | 7 | 5.6 | 6.6 |
| 0.004252 | 0.000722 | 0.26194 | 6.6 | 6.2 | 6.2 | 4.4 | 2.3 | 3.1 |
| 0.52288 | 0.001236 | 0.005558 | 6.2 | 6 | 5.8 | | 5.6 | 5.6 |
| 0.864302 | 0.000121 | 0.000342 | 1.9 | 1.1 | 1.9 | | | 2.2 |
| 2.46E-05 | 7.72E-06 | 0.011485 | 5.6 | 6 | 5.6 | | 3 | 2.9 |
| 0.851888 | 0.180503 | 0.539036 | 9.00E-01 | 9.00E-01 | 1.7 | 1.40E+00 | 1.30E+00 | 2.2 |
| 0.610847 | 0.003021 | 2.44E-03 | 1.20E+00 | | | 0.00E+00 | 2 | 1.6 |

| | | | | | | | | |
|---|---|---|---|---|---|---|---|---|
| 0.005927 | 1.96E-05 | 2.88E-04 | 4.80E+00 | 4.8 | 4.50E+00 | | 4.8 | 4.5 |
| 0.000126 | 2.52E-05 | 0.058705 | 6.6 | 6.6 | 6.6 | 5.4 | 4.8 | 1.7 |
| 0.030005 | 0.250205 | 2.41E-01 | | | | | 0.7 | 0.7 |
| 0.493 | 0.770183 | 2.18E-01 | 0.7 | 7.00E-01 | 5.00E-01 | | 0.7 | 1.1 |
| 0.617929 | 0.000158 | 2.20E-04 | 3.50E+00 | 2.7 | 3.5 | | 4.6 | 3.9 |
| 0.03376 | 0.072983 | 7.51E-01 | 3.50E+00 | 2.7 | 3.10E+00 | 0.00E+00 | 2.1 | 1.6 |
| 0.813497 | 0.035518 | 0.131871 | 9.00E-01 | | 0.9 | 1.5 | | 1.3 |
| 3.66E-08 | 0.04074 | 3.06E-08 | | | | 5.6 | 4.5 | 3.7 |
| 4.77E-05 | 2.66E-05 | 1 | 3.00E+00 | 3 | 3 | | | |
| 0.001376 | 0.252884 | 0.000647 | 1.9 | 4.5 | 2.3 | | 4 | 3.6 |
| 0.311819 | 0.068571 | 7.05E-01 | 5.00E+00 | 4.2 | 3.10E+00 | 2.70E+00 | 4.5 | 0.6 |
| 0.004202 | 0.015153 | 0.253479 | 6.6 | 6.6 | 6.2 | 2.7 | 2.3 | 4.5 |
| 7.26E-06 | 1.11E-08 | 1.93E-05 | 4.00E+00 | 2.10E+00 | 2.9 | | 2.3 | 2.3 |
| 0.711533 | 2.52E-05 | 8.26E-05 | 5.50E+00 | 4.7 | 5.10E+00 | | 4.1 | 4.5 |
| 0.758556 | 0.684608 | 1.00E+00 | 2.80E+00 | 3.2 | 2.80E+00 | 1.90E+00 | 0.5 | |
| 0.071938 | 0.226177 | 0.643662 | 1.80E+00 | 1.10E+00 | 1.8 | | 1 | |
| 0.610849 | 0.002896 | 0.00236 | 3.00E+00 | 3.00E+00 | 3 | 0.00E+00 | 3.00E+00 | 2.6 |
| 0.626139 | 0.000435 | 1.53E-03 | 6.00E+00 | 6.00E+00 | 6.00E+00 | | 6.4 | 6 |
| 0.879946 | 0.01145 | 3.59E-02 | 1.80E+00 | 2.2 | 2.6 | 1.10E+00 | 2.6 | 2.2 |
| 1 | 0.88363 | 0.951481 | 6 | 5.6 | 5.6 | | 6 | 5.6 |
| 1 | 0.97522 | 0.982161 | 6 | 6 | 5.6 | | 6 | 5.2 |
| 0.027277 | 0.083059 | 0.570331 | 5.2 | 3.3 | 3.7 | | | |
| 0.345714 | 0.023779 | 0.241838 | 1.8 | 1.4 | 1.1 | | 2.2 | 1.4 |
| 0.156069 | 0.054606 | 9.34E-01 | 3.10E+00 | 4.1 | 3.90E+00 | 0.00E+00 | 0.2 | 4.5 |
| 0.007767 | 0.002663 | 0.858524 | 4.80E+00 | 4.8 | 4.8 | | 1.2 | 2 |
| 0.264752 | 0.001074 | 7.53E-03 | 6.60E+00 | 6.2 | 6.60E+00 | | 6.2 | 6.2 |
| 0.000141 | 1.76E-05 | 0.012158 | 5.2 | 5.8 | 5.4 | | 4.8 | 5.2 |
| 0.54545 | 0.000118 | 0.000422 | 5.8 | 5.8 | 5.8 | 4.5 | 6.2 | 5.1 |
| 0.014357 | 0.002376 | 0.380001 | 4.80E+00 | 4.10E+00 | 4.5 | | | |
| 0.269933 | 0.003531 | 0.032338 | 3.80E+00 | 2.10E+00 | 2.9 | 2.30E+00 | | |
| 0.251689 | 2.48E-08 | 4.30E-08 | | | | | | |
| 0.257769 | 5.28E-06 | 1.77E-05 | | | | | 0 | 0.8 |
| 0.826092 | 0.000272 | 3.94E-04 | 2.00E-01 | 2.3 | 6.00E-01 | | 1.9 | 0.6 |
| 0.001362 | 0.000129 | 3.42E-02 | 5.00E+00 | 3.9 | 4.2 | 2.70E+00 | 0 | 0 |
| 0.000264 | 3.30E-05 | 2.31E-02 | 5.80E+00 | 6.2 | 6.20E+00 | | 3.7 | 2.3 |
| 0.370781 | 0.692378 | 0.887928 | 3 | 3.9 | 3.9 | | | 1.4 |
| 0.000811 | 0.019302 | 0.000196 | 5.6 | 4.8 | 5.6 | | 6.2 | 6.2 |
| 0.135153 | 0.094756 | 0.010858 | 4.6 | 3.5 | 5.4 | 3.4 | 6.4 | 5.2 |
| 0.905256 | 6.71E-05 | 1.97E-04 | 2.60E+00 | 3.3 | 3.30E+00 | | 2.6 | 3 |
| 0.005315 | 0.023343 | 2.28E-01 | 4.80E+00 | 4.8 | 4.1 | | 1.5 | 1.9 |
| 0.835319 | 0.272075 | 0.743467 | 2.4 | 1.3 | 2 | 0 | 1.7 | 2 |
| 0.430904 | 1 | 0.382135 | 1.7 | 1.7 | 1.7 | 0.4 | 3.5 | 5.2 |
| 0.007258 | 0.001543 | 0.000219 | 1.5 | | | 1.9 | 1.9 | 4.5 |
| 7.40E-12 | 5.45E-12 | 0.821438 | 4.80E+00 | 4.20E+00 | 4.8 | | | |
| 0.027502 | 0.009142 | 8.97E-01 | 6.20E+00 | 6.20E+00 | 6.20E+00 | | 6.2 | 6.2 |
| 1 | 0.203571 | 2.76E-01 | 5.50E+00 | 5.8 | 5.80E+00 | | 5.8 | 4.7 |
| 0.037275 | 0.000116 | 1.25E-03 | 2.50E+00 | 3.6 | 1 | 4.00E-01 | | 0.5 |
| 0.001571 | 0.002716 | 0.000133 | 2.7 | 2.3 | 1.6 | | 4.5 | 4.5 |
| 1 | 0.007844 | 1.11E-02 | 4.50E+00 | 1.4 | 1.9 | | 0 | 4.5 |
| 0.002442 | 0.001773 | 1 | 6.2 | 6.2 | 6.2 | | 2.3 | 1.9 |
| 0.401214 | 0.000146 | 0.000592 | 6.2 | 6.6 | 6.6 | 1.4 | 6.6 | 6.6 |

| | | | | | | | | |
|---|---|---|---|---|---|---|---|---|
| 0.579337 | 0.001992 | 0.001683 | 5.4 | 5.4 | 5 | | 5.2 | 5.6 |
| 1 | 0.00391 | 5.23E-03 | 5.80E+00 | 5 | 5.8 | | 5.6 | 5 |
| 0.872572 | 0.001865 | 0.002159 | 4 | 5.2 | 4.4 | | 4 | 4 |
| 7.02E-05 | 2.14E-06 | 4.01E-05 | 5 | 4.60E+00 | 5.80E+00 | 4.9 | 4.2 | 4.8 |
| 0.021092 | 9.41E-05 | 1.30E-03 | 4.80E+00 | 4.5 | 4.50E+00 | 8.00E-01 | 4.5 | 4.5 |
| 0.410413 | 0.217495 | 1.00E+00 | 2.30E+00 | 2.7 | 2.70E+00 | 2.70E+00 | 2.1 | 1 |
| 0.981264 | 0.019763 | 0.020442 | 6.6 | 6.2 | 6.6 | 3.4 | 5.5 | 5.8 |
| 0.006255 | 0.000539 | 0.000114 | 3.3 | 5.5 | 5.2 | 4.9 | 6.4 | 6.4 |
| 0.393303 | 1.66E-05 | 6.05E-05 | 6.6 | 6.6 | 6.2 | 5.9 | 6.6 | 7 |
| 0.13314 | 0.000506 | 4.38E-03 | 3.50E+00 | 2.7 | 4.20E+00 | | 4 | 4.2 |
| 0.621983 | 0.017303 | 9.04E-02 | 3.60E+00 | 3.2 | 3.90E+00 | 1.60E+00 | 2.4 | 2.4 |
| 0.035161 | 0.000416 | 0.008181 | 5.8 | 5.8 | 5.5 | 5.2 | 5.8 | 5.5 |
| 0.009377 | 5.53E-05 | 9.90E-04 | 2.00E+00 | 2.00E+00 | 2.00E+00 | 4.80E+00 | 3.2 | 1.3 |
| 0.196343 | 0.002355 | 0.024998 | 5.00E+00 | 5.20E+00 | 5.2 | 1.90E+00 | 3.10E+00 | 4.2 |
| 0.723373 | 0.013791 | 0.057983 | 4.8 | 5 | 5 | 1.9 | 5 | 4.6 |
| 0.67468 | 0.000182 | 5.88E-04 | 6.00E+00 | 5.60E+00 | 6.00E+00 | 4.20E+00 | 6 | 6 |
| 0.807057 | 0.149988 | 0.505085 | 3.90E+00 | 4.6 | 4.2 | | 5 | 4.6 |
| 0.978034 | 0.071057 | 0.060765 | 0.5 | 0.7 | 1.4 | 0.4 | 1.8 | 2.2 |
| 1 | 0.636626 | 9.12E-01 | 2.20E+00 | | 1.30E+00 | 2.00E-01 | 1.3 | 3.1 |
| 8.47E-05 | 0.600388 | 8.60E-05 | | | | 0.5 | 4.2 | 4.5 |
| 0.010083 | 0.685676 | 2.47E-02 | 1.90E+00 | 1.10E+00 | 4.50E+00 | 3.00E+00 | 3.1 | 3.1 |
| 0.001124 | 2.68E-05 | 1.54E-03 | 2.80E+00 | 3.60E+00 | 2.80E+00 | | 2.2 | 3 |
| 0.90352 | 0.002887 | 8.36E-03 | 3.30E+00 | 3.3 | 4.8 | | 2.9 | 3.6 |
| 1 | 0.59401 | 0.797384 | 2.2 | 2.2 | 1.8 | | 2.6 | 2.6 |
| 8.81E-05 | 1.15E-05 | 6.12E-03 | 6.20E+00 | 6.2 | 6.20E+00 | 2.70E+00 | 6.6 | 6.6 |
| 0.1936 | 0.070479 | 9.49E-01 | 3.10E+00 | 3.9 | 3.50E+00 | 1.90E+00 | 0.2 | 0.2 |
| 0.003957 | 9.41E-05 | 0.004613 | 3.90E+00 | 2.30E+00 | 2 | | 3.7 | 3.7 |
| 0.197147 | 0.000568 | 0.004042 | 2 | 2 | 2.7 | | 1.6 | 1.6 |
| 0.085144 | 8.19E-05 | 5.73E-04 | 6.00E+00 | 5.20E+00 | 5.20E+00 | 2.70E+00 | 5.60E+00 | 5.2 |
| 0.000956 | 0.001305 | 0.548583 | 3.5 | 2.3 | 3.1 | | 4.5 | 1.5 |
| 0.001042 | 1.19E-06 | 1.38E-05 | 5.1 | 5.50E+00 | 5.50E+00 | 1.1 | 5.50E+00 | 5.5 |
| 0.679255 | 0.009162 | 6.65E-03 | 5.90E+00 | 4.8 | 5.50E+00 | 2.30E+00 | 5.5 | 5.2 |
| 0.090326 | 0.002466 | 8.20E-04 | 5.80E+00 | 4.4 | 5.20E+00 | | 5 | 6 |
| 0.01708 | 2.23E-05 | 2.21E-04 | 4.80E+00 | 4.8 | 4.10E+00 | | 4.1 | 5.2 |
| 0.229074 | 0.009882 | 0.003402 | 5.6 | 5.4 | 5.2 | | 6 | 5.2 |
| 7.27E-06 | 1.51E-06 | 5.52E-03 | 6.60E+00 | 6.6 | 6.60E+00 | | 1 | 2.1 |
| 0.046511 | 0.000177 | 1.91E-03 | 2.50E+00 | 3.6 | 6.00E-01 | 4.00E-01 | | 0.5 |
| 0.008616 | 1.56E-05 | 0.000168 | 5.8 | 5.5 | 5.8 | | 5.8 | 6.2 |
| 0.060862 | 6.35E-05 | 4.80E-04 | 5.60E+00 | 5.2 | 4.8 | | 4.5 | 4.8 |
| 0.028515 | 0.00021 | 0.003401 | 5.2 | 4.5 | 4.8 | | 1.7 | 2.1 |
| 0.001815 | 0.001753 | 9.29E-01 | 2.70E+00 | 3.3 | 1.40E+00 | 0.00E+00 | 1.1 | 3 |
| 0.307292 | 6.50E-05 | 0.000283 | 6.10E+00 | 6.10E+00 | 6.1 | 5.30E+00 | 6.40E+00 | 6.4 |
| 0.489905 | 6.72E-06 | 2.01E-05 | 5 | 5 | 5.4 | | 5 | 4.6 |
| 0.952589 | 4.03E-06 | 1.04E-05 | 6.00E+00 | 6 | 5.6 | | 6 | 5.6 |
| 0.405662 | 0.620163 | 0.13349 | 3.50E+00 | 3.90E+00 | 2.7 | | 3.9 | 5.4 |
| 0.343039 | 0.001883 | 0.012288 | 3.9 | 4.2 | 3.9 | 2.3 | 4 | 4.2 |
| 0.142359 | 0.01552 | 0.003488 | 3.7 | 3.7 | 4.5 | | 5.1 | 5.5 |
| 1.97E-05 | 7.56E-06 | 4.64E-07 | 1.4 | 1.4 | 2.1 | 2.7 | 5.5 | 5.5 |
| 1 | 0.000395 | 0.000655 | 2.3 | 4.2 | 3.5 | | 3.9 | 3.9 |
| 0.405527 | 4.94E-05 | 7.30E-05 | 4.7 | 5.8 | 5.8 | | 6.2 | 5.8 |
| 0.07648 | 0.169693 | 1.06E-02 | 1.40E+00 | 1.7 | 2.10E+00 | | 1.7 | 0.4 |

| | | | | | | | | |
|---|---|---|---|---|---|---|---|---|
| 0.000141 | 1.76E-05 | 0.012158 | 5.2 | 5.8 | 5.4 | | 4.8 | 5.2 |
| 0.814043 | 0.000294 | 4.18E-04 | 6.40E+00 | 6.40E+00 | 5.6 | | 6.40E+00 | 5.6 |
| 0.023846 | 0.00036 | 0.008839 | 6.2 | 6.2 | 6.2 | 4 | 6 | 6 |
| 1.48E-05 | 5.21E-06 | 0.016292 | 6.60E+00 | 6.6 | 6.6 | | 4 | 5.2 |
| 0.506265 | 0.000154 | 0.000199 | 5.40E+00 | 3.90E+00 | 4.6 | | 5.60E+00 | 5 |
| 1 | 0.888711 | 1 | 0.7 | | | 1.5 | 0.8 | 0.7 |
| 0.77179 | 0.101754 | 0.38355 | 1.1 | 1.4 | 2.4 | 0.2 | 3 | 1.8 |
| 0.645368 | 0.147759 | 6.51E-01 | 3.70E+00 | 3 | 3.3 | | 0.8 | 1.6 |
| 0.959352 | 0.813581 | 1 | 5.2 | 6 | 5.6 | | 6 | 4.2 |
| 9.20E-06 | 7.54E-06 | 0.044495 | 3.9 | 4.3 | 3.6 | | | |
| 0.629418 | 0.024129 | 0.013158 | 1.7 | 4.8 | 4 | | 2.5 | 3.4 |
| 0.006255 | 0.000338 | 0.030564 | 3.2 | 4.3 | 3.6 | 0 | 1.8 | 3 |
| 0.244999 | 0.013269 | 0.181733 | 1.1 | 0.9 | 1.8 | | 4.5 | 4.8 |
| 0.146806 | 0.000111 | 6.54E-04 | 1.90E+00 | 3.8 | 2.30E+00 | | 0.4 | 1.1 |
| 0.462473 | 0.008396 | 0.055723 | 2.60E+00 | 3.30E+00 | 3.7 | | 0.5 | 1.6 |
| 0.303595 | 0.206304 | 1 | 2.9 | 0.8 | 0.5 | | 1.2 | 1.2 |
| 0.345476 | 7.54E-06 | 1.67E-05 | 3.9 | 3.5 | 3.5 | 5.5 | 4.6 | 5 |
| 0.065082 | 0.000128 | 9.59E-05 | 4.6 | 5 | 5.4 | | 6 | 5.8 |
| 0.671385 | 7.66E-06 | 1.83E-05 | 5.8 | 5.8 | 5 | | 5.8 | 6 |
| 0.089988 | 0.000627 | 0.007605 | 2.70E+00 | 2.3 | 2.3 | | 0.6 | 3.3 |
| 0.29123 | 0.131315 | 0.996101 | 1.6 | 0.5 | 1.8 | 1.7 | 0.5 | |
| 0.301333 | 0.313225 | 4.93E-02 | 4.20E+00 | 3.5 | 2.30E+00 | | 3.3 | 3.6 |
| 0.18298 | 0.298921 | 0.959386 | 5.20E+00 | 4.7 | 3 | | 4.1 | 5.2 |
| 0.800388 | 0.794385 | 4.30E-01 | 2.30E+00 | 2.7 | | | 4.1 | 3.3 |
| 0.965529 | 0.002773 | 0.007185 | 5.2 | 4.5 | 4.1 | | 4.6 | 4.8 |
| 0.003061 | 0.556574 | 1.70E-03 | 3.30E+00 | 4.80E+00 | 5.20E+00 | 2.30E+00 | 5.80E+00 | 5.8 |
| 0.336322 | 0.755011 | 0.141925 | | 5.2 | 5.1 | | 5.5 | 6.2 |
| 0.643805 | 0.00063 | 2.23E-03 | 0 | 4.20E+00 | | 2.3 | 3 | 3.4 |
| 5.45E-05 | 0.000204 | 1.66E-05 | 4.20E+00 | 2.3 | 3.10E+00 | | 5.2 | 6 |
| 0.007004 | 0.002942 | 0.980897 | 4.10E+00 | 4.1 | 4.8 | 5.3 | 4.2 | 0.6 |
| 0.025639 | 0.001077 | 0.049301 | 4.1 | 3.7 | 3.7 | | 3.5 | 4.2 |
| 0.114978 | 0.004306 | 0.001328 | 0.6 | 1.7 | | 0 | 1.9 | 0.2 |
| 1.84E-05 | 2.50E-05 | 0.021111 | 5.8 | 5.8 | 4.6 | 4.2 | 6 | 6.6 |
| 0.025922 | 0.000387 | 9.24E-03 | 1.30E+00 | 2 | 1.30E+00 | | | 2 |
| 0.277825 | 0.528367 | 0.074571 | 9.00E-01 | 1.70E+00 | 0.9 | | 9.00E-01 | 0.9 |
| 0.08758 | 0.004676 | 0.13349 | 1.3 | 1.3 | 0.9 | | 0.9 | 1.1 |
| 0.039438 | 0.017818 | 1 | | | | | | |
| 0.013302 | 0.00091 | 0.080645 | 2.7 | 3.5 | 4.6 | 1.9 | | 4.5 |
| 0.803592 | 0.000828 | 0.001002 | 4.10E+00 | 4.50E+00 | 4.5 | | 5.20E+00 | 5.6 |
| 0.989324 | 0.597529 | 0.930292 | 2.60E+00 | 2.8 | 1.8 | 0.4 | 2 | 2.4 |
| 0.027352 | 0.011198 | 0.985858 | 4.1 | 2.6 | 2.2 | 0.8 | | |
| 2.01E-06 | 6.93E-07 | 0.098053 | 1.4 | 1.1 | 2 | | | |
| 5.11E-06 | 4.03E-06 | 0.317095 | 5.6 | 5.6 | 5.6 | 4.9 | 4.3 | 4.3 |
| 1.41E-07 | 7.19E-08 | 7.37E-01 | 3.50E+00 | 3.5 | 2.70E+00 | | | |
| 0.967478 | 0.99873 | 0.903874 | 1.4 | 1.8 | 1.8 | | 1.1 | 0.7 |
| 0.005351 | 5.28E-06 | 2.63E-05 | 6.00E+00 | 6 | 6 | 4.00E-01 | 6 | 6 |
| 7.07E-05 | 5.86E-09 | 1.28E-06 | 5.2 | 5.6 | 5.6 | 4.5 | 5.4 | 5.4 |
| 0.222576 | 3.06E-05 | 0.000143 | 3.7 | 4.10E+00 | 3.7 | | 4.1 | 3.3 |
| 0.002503 | 0.000821 | 7.28E-01 | 3.10E+00 | 3.1 | 3.90E+00 | 1.10E+00 | 3.2 | 2.8 |
| 0.622038 | 0.039172 | 1.89E-02 | 5.6 | 5.60E+00 | 5.60E+00 | | 5.50E+00 | 5.6 |
| 0.004005 | 0.603574 | 0.002215 | 4.5 | 3.9 | 4.6 | | 5.6 | 5.8 |

| | | | | | | | | |
|---|---|---|---|---|---|---|---|---|
| 0.640518 | 0.000784 | 8.25E-04 | 5.10E+00 | 5.1 | 5.10E+00 | | 6.2 | 5.8 |
| 0.000213 | 2.89E-05 | 2.44E-02 | 4.70E+00 | 3.6 | 2.80E+00 | | 0.5 | 2.5 |
| 0.591915 | 1 | 0.746744 | 5.4 | 4.6 | 4 | 4.2 | 5.4 | 5.4 |
| 0.479083 | 6.00E-05 | 0.000232 | 6.6 | 6.2 | 6.6 | 5.5 | 6.4 | 6.6 |
| 0.221622 | 3.37E-05 | 1.59E-04 | 6.20E+00 | 6.2 | 6.2 | | 6.2 | 6.2 |
| 0.859427 | 1.76E-05 | 3.85E-05 | 1.8 | 2.4 | | 2 | 4.6 | 4.2 |
| 0.73174 | 0.000702 | 8.14E-04 | 6.60E+00 | 6.2 | 6.20E+00 | | 6.2 | 6.2 |
| 1.24E-05 | 6.96E-06 | 1 | 1.2 | 3.5 | 2.7 | | | |
| 0.516727 | 0.94433 | 3.43E-01 | 2.7 | 2.70E+00 | 4.2 | | 3.5 | 5.2 |
| 0.213237 | 5.82E-05 | 7.07E-05 | | | | | 0 | |
| 0.392921 | 0.006911 | 0.003569 | 0.6 | 0.6 | 0.2 | | 1.7 | 0.2 |
| 1 | 0.000219 | 0.000434 | 5.40E+00 | 4.20E+00 | 5.8 | | 5.8 | 5.8 |
| 7.40E-12 | 5.45E-12 | 0.183017 | 4.2 | 3.50E+00 | 2.9 | | | |
| 0.000825 | 1.22E-05 | 0.000399 | 5.00E+00 | 4.60E+00 | 5 | 4.00E-01 | 3.10E+00 | 3.9 |
| 0.102534 | 0.001161 | 0.016071 | 4.8 | 5.20E+00 | 4.5 | 3.8 | 3.9 | 3.5 |
| 0.391088 | 0.000283 | 0.00122 | 4.70E+00 | 3.90E+00 | 5.5 | | 5.10E+00 | 5.1 |
| 0.726586 | 0.00039 | 0.001227 | | | | | | |
| 0.004202 | 0.344935 | 1.59E-02 | 4.50E+00 | 4.5 | 5.2 | 3.3 | 6.2 | 5.6 |
| 0.048668 | 0.211848 | 0.00882 | 3.7 | 3.3 | 4.2 | | 4 | 3.6 |
| 0.634121 | 0.000684 | 7.27E-04 | 4.20E+00 | 2.3 | 1.90E+00 | 8.00E-01 | 5.3 | 4.5 |
| 0.065802 | 0.000128 | 9.61E-05 | 4 | 4.4 | 3.3 | | 2.9 | 5.2 |
| 0.025627 | 0.000101 | 0.001268 | 1.5 | 4.5 | | | 3 | 3.4 |
| 0.214326 | 0.002766 | 0.02873 | 6.2 | 6.6 | 6.6 | | 6.6 | 6.6 |
| 0.984913 | 0.036824 | 0.087851 | 5.8 | 5.80E+00 | 5.8 | | 6 | 5.4 |
| 0.424375 | 0.083472 | 0.023633 | 2.5 | 2.1 | 2.5 | | 3.9 | 4.8 |
| 0.028672 | 0.001345 | 3.60E-04 | 6.20E+00 | 6.2 | 6.00E+00 | | 6 | 6 |
| 2.27E-05 | 7.90E-07 | 3.58E-05 | 6.60E+00 | 6.60E+00 | 5.8 | | 5 | 5.4 |
| 0.805087 | 0.27202 | 0.772227 | 4.2 | 3.6 | 3.5 | | 4.5 | 4.2 |
| 0.000811 | 4.03E-06 | 3.05E-05 | 6.00E+00 | 6.00E+00 | 5.8 | 3.40E+00 | 5.20E+00 | 5.5 |
| 0.39929 | 0.003327 | 0.00204 | 3.70E+00 | 4.10E+00 | 3.3 | | 3.00E+00 | 5.2 |
| 0.033163 | 5.45E-12 | 2.22E-11 | | | | | | |

| Peak_Rating_D3 | Peak_Rating_D_PB | Peak_Rating_Pool_QC_1 | Peak_Rating_Pool_QC_2 | Peak_Rating_Pool_QC_3 | Peak_Rating_S1 | Peak_Rating_S2 | Peak_Rating_S3 | Peak_Rating_S_PB |
|---|---|---|---|---|---|---|---|---|
| 6.6 | 5.6 | 10 | 9.6 | 9.6 | 6.6 | 6.6 | 6.6 | 6 |
| 4.3 | 2 | 5.1 | 5.1 | 1.3 | 4.3 | 4.7 | 4.3 | 1.4 |
| | | | | 4.3 | | | | |
| 6.6 | 2.9 | 9.6 | 9.2 | 9.6 | 4.6 | 3.5 | 3.5 | 6.2 |
| 4.9 | 0.4 | 9.2 | 9.2 | 8.9 | | | | |
| | | 9 | 9 | 8.6 | | | | |
| 7 | 6.1 | 7.9 | 8.2 | 9 | 6.6 | 6.6 | 6.6 | 6.1 |
| 4.4 | 2.3 | 7 | 5.2 | 7 | 1.7 | 3.3 | 2.1 | 1.5 |
| 5.9 | | 8.6 | 8.2 | 7.8 | 4.9 | | 4.2 | 2.7 |
| 3.1 | 2.3 | 3.6 | 3.4 | 4.1 | 2.5 | 4.8 | 2.9 | 4.5 |
| | 0.4 | 2.1 | 2.1 | 2.3 | 3.3 | 3 | 3.7 | |
| 1.9 | 2.7 | 2.2 | 3.1 | 3 | 0.2 | 0.5 | 2.3 | |
| 5.8 | | 6.2 | 6.9 | 7.7 | | | | |
| 6.2 | 3 | 9.2 | 8.9 | 8.9 | 5.2 | 5.2 | 6 | 4.9 |
| 5.8 | | 8.1 | 6.6 | 8.1 | 5.2 | 4.1 | 5.2 | |
| 5.6 | | 7.4 | 6.4 | 6.8 | 4.8 | 3.1 | 3.1 | |
| 6.2 | 6 | 8.4 | 8.7 | 8.7 | 6.2 | 5.8 | 6.2 | 5.6 |
| 6.2 | 2.7 | 8.7 | 9.1 | 9.1 | 4.6 | 4.2 | 4.2 | 3 |
| 0.7 | | 4.2 | 4.2 | 4.6 | 2 | | 1.7 | |
| | | 2.9 | 1.4 | 1.9 | | | | |
| 2.3 | | 2 | 3.6 | 2.2 | 2.5 | 1.7 | 1.7 | |
| 0.7 | 1.5 | 9.1 | 9.1 | 9.1 | 0.2 | 0.2 | 0.2 | 2.7 |
| 1.1 | | 9.1 | 9.1 | 8.7 | 5.6 | 5.6 | 2.9 | 2.9 |
| 0.2 | 1.5 | 4.6 | 4.2 | 3.5 | 0.2 | 1 | 1.5 | |
| | 1.9 | 3.6 | 2.5 | 2.9 | 0.2 | | | |
| 0.2 | | 7.2 | 6.9 | 8 | | 0.6 | 0.2 | |
| 5.5 | | 7.8 | 7.4 | 6.2 | 3.7 | 3.7 | 3 | 0 |
| 2.9 | 0 | 4.2 | 4.7 | 4.6 | 4.2 | 3.1 | 3.5 | 0 |
| 1.9 | 4.5 | 6.9 | 5.4 | 6.2 | 2.4 | 2.4 | 2.8 | |
| | | 8.5 | 8.9 | 8.1 | 2.9 | 4 | 2.5 | |
| 5 | | 7.9 | 6.8 | 7.9 | 0.8 | 1.1 | 2.3 | |
| 6.2 | 0 | 8.4 | 8.4 | 8.7 | 6.2 | 6.6 | 6.2 | 0.8 |
| 0.7 | 0.4 | 4.1 | 4.1 | 4.8 | 0.8 | 2.3 | 2.3 | |
| | | 1.1 | 1.7 | 1.6 | 2.2 | 0.8 | 0.8 | |
| | | 4.5 | 4.2 | 4.8 | | | | |
| 1.1 | 0 | 3.7 | 4.1 | 3.3 | 3 | 2.6 | 3 | 0 |
| 1 | 3 | | | 3.1 | 4.4 | 3.5 | 2.3 | |
| | 0 | 0.5 | 0.7 | 2.1 | 2.7 | 2 | 2.2 | 0 |
| 0.9 | 0.2 | 4.2 | 3.5 | 2.3 | 0.6 | 0.8 | 1.2 | 3 |
| 1.2 | | 3.1 | 3.5 | 3.9 | 2 | 1.2 | 1.7 | 0 |

| | | | | | | | | |
|---|---|---|---|---|---|---|---|---|
| 2.7 | | 2.9 | 2.9 | 2.9 | | 0.6 | 0.5 | |
| 0.8 | 1.5 | 7.8 | 7.4 | 7.4 | 0.5 | 4.5 | 0.2 | |
| 3.6 | 1.5 | 7.5 | 8.3 | 6.7 | 0.5 | 1.1 | 0.8 | 0 |
| | 0.2 | 1.1 | 0.9 | 1.7 | 0.8 | | | 0 |
| 6 | 2.3 | 8.4 | 8 | 8.4 | 6.2 | 6.2 | 5.8 | 3 |
| 2.5 | 4.8 | 9.6 | 9.6 | 9.6 | 1.5 | 0.7 | 0.5 | 4.1 |
| | 0.4 | 9.2 | 9.6 | 9.2 | | | | |
| 1 | 2.3 | 5.1 | 7.4 | 4.4 | 1.9 | 2.3 | 1.9 | 2.7 |
| 0.6 | | 8.5 | 8.1 | 7.4 | 4.5 | 1.9 | 0.2 | |
| 3.7 | | 3.6 | 3.2 | 2.5 | 0.2 | 0.2 | | |
| 1.1 | | 1.2 | 2.1 | 1.3 | | | | |
| | | 6.1 | 5 | 6.9 | | 4.9 | | |
| | 5.6 | 9.1 | 7.4 | 7.8 | | | | |
| | | 7.9 | 6.8 | 6.8 | | | | |
| 4.6 | | 4.7 | 4.5 | 4.9 | 2.9 | 1.4 | 2.1 | |
| 1.1 | 0.4 | 2.7 | 0.9 | 0.9 | | 0.6 | 0.7 | 1.1 |
| | 0.4 | 0.8 | 1.4 | 1.1 | 3.3 | 4.6 | 3.7 | 1.5 |
| 4.7 | 0.4 | 8.4 | 8 | 7.2 | 5.8 | 5.5 | 5.5 | 1.9 |
| 0.5 | 4.5 | 8 | 7.2 | 8.4 | 0.5 | 0.5 | | |
| 5.5 | 0.8 | 6.2 | 5.5 | 7 | 5.1 | 4.7 | 5.1 | 0.4 |
| 0.7 | | 5.9 | 5.1 | 6.2 | 0.5 | 0.7 | 0.7 | |
| 3.3 | 0.6 | 8.3 | 8.6 | 7.9 | 3.2 | 3.2 | 3.6 | 2.2 |
| 0.8 | 4.5 | 5.2 | 6 | 5.6 | 5.8 | 5.8 | 5.8 | 4.2 |
| 1.1 | | 0.7 | 0.7 | 1.2 | 3.5 | 3.3 | 2.6 | 0.6 |
| 6.4 | | 7.5 | 7.9 | 7.5 | 4.8 | 3.7 | 3 | |
| | | 4.6 | 1.6 | 3.5 | 5 | 3.1 | 3.9 | |
| 0.8 | | 7.7 | 5 | 3.9 | 4.5 | 2.3 | 3 | |
| 5 | | 7 | 7 | 6.4 | 4.8 | 5.2 | 4 | |
| 1.7 | 0.8 | 6.5 | 6.5 | 7.3 | 4.2 | 3.1 | 3.1 | 1.5 |
| 1.7 | 0.4 | 0.7 | 2.5 | 1.6 | 5 | 5.2 | 4.1 | 4.9 |
| 6.2 | 7.6 | 9.1 | 9.1 | 9.1 | 8.9 | 8.7 | 7.9 | 8.3 |
| 1.4 | | 1.3 | 1.2 | 1.1 | | | | |
| 5.8 | | 7.3 | 8 | 8.4 | 4.8 | 5.4 | 5.4 | |
| | | 2.9 | | 2.9 | 2.3 | 1.6 | 3.5 | 0 |
| 4.8 | 1.9 | 4 | 4 | 3.6 | 2.2 | 2.6 | 2.2 | 1.5 |
| 0.9 | | 1.8 | 1.5 | 1.7 | 0.7 | 1.1 | 0.7 | |
| 1.1 | 0 | 2.1 | 2.2 | 1.7 | | | 0.8 | |
| 0.5 | | 1.4 | 0.6 | 1.4 | | 0.2 | 0.2 | |
| | | 1.7 | 3.3 | 1.7 | 3.6 | | | |
| 2.3 | | 3.1 | 2.9 | 3.2 | 3.6 | 4.4 | 2 | |
| 2.3 | | 5.4 | 5 | 4.6 | 1.4 | 2.5 | 2.1 | |
| 2 | | 2.5 | 2.1 | 3.4 | 1.4 | 1.4 | 3.6 | |
| | | 1.7 | 1.1 | 0.9 | 0.7 | 0.7 | 1.1 | |
| | | 1.1 | 1.8 | 2 | 1.1 | 3.7 | 1.4 | |
| 2 | 3.4 | 0.8 | 2.5 | 4.8 | 2.1 | 1.4 | 4 | 2.7 |
| 4.6 | 3.4 | 6.9 | 7.3 | 7.3 | 5.2 | 2.3 | 3.9 | 1.9 |
| 5.6 | 4.5 | 8.5 | 8.5 | 8.5 | 5 | 5 | 4.2 | 4.2 |
| | 0.4 | 4.8 | 4.4 | 4.8 | 5.1 | 5.5 | 5.5 | 0.2 |
| | 0.2 | 0.7 | | 0.8 | 2.3 | 2 | 2 | 0.2 |
| 6.2 | 5.1 | 9.1 | 8.7 | 9.1 | 6.6 | 5.5 | 5.1 | 3.7 |
| 6.6 | 3.4 | 9.6 | 9.6 | 9.6 | 6.6 | 6.6 | 6.6 | 4 |

| | | | | | | | | |
|---|---|---|---|---|---|---|---|---|
| 3.5 | 1.9 | 6.4 | 5.6 | 6 | 3.6 | 2.9 | 2.9 | 0.8 |
| 2.5 | 0 | 3.6 | 3.4 | 3.7 | 1.7 | 1 | 0.5 | |
| 2.3 | | 4.7 | 5.9 | 6.3 | 4 | 3.1 | 3.5 | |
| 5.4 | 1.9 | 7.2 | 7.2 | 7.6 | 4.8 | 4 | 2.1 | 4.5 |
| 5.6 | 4.5 | 8.5 | 8.5 | 8.5 | 5 | 5 | 4.2 | 4.2 |
| | 1.9 | 4.5 | 5 | 4.1 | | | | |
| 0.6 | | 1 | 2 | 2.9 | 0.6 | 0.5 | 4.5 | 4.5 |
| 5.6 | | 8.3 | 7.9 | 8.6 | 6 | 5.2 | 6 | |
| 0.8 | 1.9 | 2.5 | 2.5 | 1.2 | 0 | | | |
| 0.7 | 0.2 | 1.6 | 2.5 | 1.7 | 0.8 | 0.7 | 0.5 | 1.5 |
| 2.1 | 0 | 6.7 | 4.8 | 5.2 | 4.5 | 0.5 | | |
| 6.6 | 4.2 | 9.6 | 9.6 | 9.6 | 6.6 | 6.6 | 6.2 | 3.3 |
| 2.2 | | 3.1 | 2.7 | 3.7 | 2 | 3.3 | 3 | |
| 0.9 | 0.2 | 2.2 | 2 | 2 | | 1.2 | | |
| | | 1.5 | 2.2 | 1.8 | | | | |
| 5.8 | 5.4 | 8.2 | 8.2 | 7.9 | 5.5 | 5.5 | 5.8 | 5.4 |
| 5.8 | 4.8 | 9.1 | 8 | 8.4 | 5.5 | 6.2 | 5.5 | 3.9 |
| 6.2 | 5.2 | 8.4 | 8.4 | 8.4 | 6 | 5.8 | 6.2 | 5.2 |
| 4.2 | | 4.9 | 3.4 | 4.5 | | | | |
| 6.2 | | 9.1 | 8.7 | 8.4 | 6.8 | 7.1 | 7.1 | 6.8 |
| 3.3 | 4.2 | 7.3 | 7.7 | 7.7 | 5.5 | 4 | 3.6 | |
| 5.1 | 0.6 | 9.6 | 9.6 | 9.6 | 6.2 | 6.6 | 5.8 | 0.6 |
| 6 | | 8.1 | 8.5 | 8.5 | 4.2 | 3.9 | 4.6 | |
| 5.4 | 6.4 | 8.3 | 8.3 | 8.5 | 5.2 | 5.5 | 5.5 | 5.3 |
| 6.2 | | 8.6 | 8.6 | 7.9 | 4.1 | 5.1 | 5.1 | 0.4 |
| 3.9 | 0.4 | 3.3 | 4.8 | 4.6 | 3.6 | 2.7 | 3.6 | 1.9 |
| 5.8 | | 8.9 | 8.1 | 7.8 | 5 | 5.4 | 5.4 | |
| 0.6 | 2.5 | 9.2 | 9.2 | 9.6 | 4.5 | 3.8 | 3.8 | 2.7 |
| 1.9 | 1.5 | 5.9 | 7.4 | 5.1 | 1.5 | | 1.4 | 1.9 |
| | 5.9 | 9.2 | 9.2 | 9.2 | | | | 2.5 |
| 5.8 | | 8.4 | 8.7 | 8.7 | 6.2 | 6.2 | 6.2 | 2 |
| 1.2 | 0.2 | 4 | 3.4 | 3.4 | 1 | 1.4 | 0 | 1.5 |
| 6.2 | | 9.1 | 8.7 | 8.7 | 6.6 | 4.7 | 6.6 | |
| 3.3 | | 7.1 | 6 | 7.1 | 2.6 | 1.1 | 2.6 | 4.9 |
| | 1.7 | 6.5 | 5.7 | | 4.8 | 3.7 | 3.7 | 3.4 |
| 5.2 | 1.5 | 7.3 | 8.3 | 8.3 | 5.5 | 5.5 | 5.5 | |
| 5.6 | 4.5 | 8.5 | 8.1 | 7.9 | 5.8 | 5.2 | 5.8 | 3.4 |
| | | | 4.4 | | | | | |
| 4.2 | | 8.2 | 7.4 | 6.7 | 3 | 3.4 | 3.4 | |
| 5.2 | | 6.8 | 6.4 | 6.8 | 5 | 4.6 | 5 | |
| 5.1 | | 6.7 | 6.7 | 6 | 4.8 | 4.7 | 5.5 | |
| 4.8 | | 7.8 | 7.8 | 4.3 | 2 | 4.1 | 6 | |
| 4.8 | | 2.3 | 1.2 | 1.7 | 0.6 | 1 | 0.2 | |
| 5.6 | | 9.6 | 9.6 | 8.9 | 6.6 | 6.6 | 6.6 | |
| | | 7.3 | 5.4 | 6.2 | | | | |
| 3 | 1.5 | 3.2 | 2.4 | 2.2 | 4.2 | 3 | 3.7 | |
| 4.6 | | 6.6 | 5.9 | 5.9 | 1.4 | 2.9 | 3.3 | |
| 6.6 | 5.2 | 9.1 | 8.7 | 9.1 | 6.6 | 6.2 | 6.6 | 5.5 |
| 5.2 | 2.3 | 7.6 | 7.6 | 7.6 | 5 | 5 | 5 | 4.9 |
| | 5.8 | 8.3 | 8.1 | 7.9 | | | | |
| 5.2 | 1.9 | 7.8 | 7.8 | 7 | 5 | 2.7 | 4.2 | 1.5 |

| | | | | | | | | |
|---|---|---|---|---|---|---|---|---|
| 2 | 0.4 | 4.4 | 4 | 4.7 | 1 | 1.4 | 1 | 1.5 |
| 6.2 | 1.5 | 8.7 | 8.4 | 8 | 5.2 | 6 | 5.6 | 3.8 |
| 4.4 | | 8.4 | 6.5 | 7.3 | 3.6 | 3.1 | 3.9 | |
| 5.8 | | 7.6 | 6.8 | 6.5 | 5.8 | 5.8 | 5.5 | |
| 6 | | 6.7 | 7.1 | 5 | 4.6 | 5 | 4.6 | |
| 6.2 | 1.9 | 7.1 | 7.5 | 7.5 | 4.8 | 2.7 | 3.3 | 2.7 |
| 5.6 | | 8.6 | 8.3 | 9 | | | | |
| 6.6 | 5.8 | 9.1 | 9.1 | 9.1 | 6.6 | 6.6 | 6.6 | 5.6 |
| 6.6 | | 8.6 | 8.2 | 6.7 | 5.5 | 5.5 | 5.1 | 1.5 |
| 3.1 | | 3.3 | 3.1 | 2.5 | 4.4 | 3.5 | 2.7 | |
| 5.2 | | 8.1 | 7.4 | 7 | 2.6 | 3.4 | 3.4 | |
| 4.1 | | 7 | 7.4 | 8.1 | 5.2 | 4.8 | 5.6 | |
| 6.2 | | | 9.2 | 9.2 | 6.2 | 6.2 | 6.6 | 4.5 |
| 0.9 | 5.3 | | 4.7 | 5.2 | 1.7 | 3.2 | 5.8 | 3.4 |
| 5.6 | | 7.2 | 7.2 | 7.6 | 2.9 | 2.9 | 2.1 | |
| 4.5 | | 8 | 7.6 | 8 | 6.4 | 5.1 | 5.5 | |
| 5.4 | | 7.9 | 7.9 | 7.1 | 5.5 | 4 | 5.5 | |
| 5 | | 7.9 | 7.1 | 6.8 | 4 | 3.6 | 3.5 | |
| 6.6 | | 7.3 | 8.5 | 8.5 | | | | 5.5 |
| 4.2 | | 7.3 | 8 | 7.7 | 5.4 | 4.6 | 4.6 | |
| 6.2 | | 8.7 | 8.7 | 7.2 | 5.8 | 6.6 | 5.8 | |
| 6.6 | 4.9 | 9.1 | 9.1 | 8.7 | 6.6 | 6.6 | 6.2 | 4.5 |
| 6 | | 9.6 | 9.6 | 9 | 6.2 | 6.6 | 6.7 | |
| 2.9 | | 6.1 | 5.4 | 6.5 | 4.6 | 5 | 4.2 | |
| 3.6 | | 4.5 | 3.7 | 4.9 | 3.5 | 4.2 | 3.9 | |
| 6.6 | | 8.7 | 9.1 | 9.1 | 6.2 | 6.2 | 6.6 | 3 |
| 6.2 | | 8.2 | 8.2 | 6.3 | 5.5 | 5.5 | 4.3 | |
| 6.6 | | 9.2 | 8.9 | 9.6 | 5.6 | 4.5 | 5.2 | |
| 3.9 | | 4.6 | 4.6 | 5.2 | 4.8 | 1.7 | 3.3 | |
| 5.8 | 1.1 | 7.9 | 7.9 | 7.9 | 5.8 | 5.5 | 5.1 | 4.2 |
| 6.2 | | 8.4 | 8.4 | 8.7 | 1.4 | | 2.1 | 5.8 |
| 6.2 | 3.3 | 8.7 | 8.7 | 8.7 | 6.6 | 6.2 | 6.2 | 2.5 |
| 6.2 | 6.2 | 8.7 | 8.7 | 8.7 | 6.6 | 6.2 | 6.2 | 5.8 |
| 0 | | 3.3 | 2.5 | 1 | 5.2 | 3.6 | 3.1 | |
| 0.2 | | 3.4 | 4.5 | 2.9 | | 4.5 | 1.5 | |
| 3.3 | | 3.7 | 3.4 | 5.8 | 1.5 | 1.1 | 0.8 | |
| | 3.1 | 4.2 | 4.4 | 2.6 | 5.2 | 3 | 3.6 | 4.5 |
| 2.9 | 0.4 | 9.2 | 8.9 | 8.5 | 6.6 | 6.2 | 6.6 | 1.5 |
| | | 4 | 4.3 | 3.6 | 4.3 | 4.3 | 4.3 | |
| 5.4 | | 7.5 | 6.5 | 6.9 | 5 | 5.8 | 5.6 | 2.3 |
| 0.6 | | 1.2 | 1.7 | 2.1 | 0.2 | 2.1 | 1.7 | |
| 5.4 | | 6.7 | 5.8 | 5 | 3.3 | 4.2 | 2.5 | |
| 3.5 | | 4.4 | 4 | 4.4 | | | | |
| 6.6 | 1.5 | 9.1 | 8.7 | 7.6 | 5.5 | 5.5 | 5.8 | |
| 1.2 | | 1.2 | 1.6 | 2.6 | 3.1 | 3.5 | 3.1 | 0.8 |
| 5.8 | | 5.5 | 5.1 | 5 | 3.9 | 2.7 | 3.9 | |
| 6.2 | | 8.7 | 8.4 | 6.5 | 5.1 | 5.8 | 5.5 | 3 |
| 5 | | 7.6 | 8 | 5.5 | 4.5 | 4.8 | 3.7 | |
| 6.6 | 2.1 | 8.7 | 8.7 | 9.1 | 4.8 | 6 | 5.6 | 2.9 |
| 3.7 | | 7.4 | 6.6 | 6.6 | 2.5 | 2.9 | 2 | |
| 6.2 | | 8.4 | 8 | 5.7 | 4.5 | 4.1 | 3.9 | |

| | | | | | | | | |
|---|---|---|---|---|---|---|---|---|
| 4.1 | | 5.1 | 4.7 | 6.3 | 2.3 | | 1.9 | |
| 6.6 | 5 | 9.1 | 9.1 | 9.1 | 6.6 | 6.6 | 6.6 | 3.9 |
| 6.6 | 2.9 | 9.1 | 9.1 | 8.7 | 6 | 6 | 6 | 5.2 |
| 6.6 | 4.8 | 8.7 | 9.1 | 8.7 | 6.6 | 6.6 | 6.2 | 4.4 |
| | | 3.5 | 3.1 | 3.4 | | 4.5 | | |
| 4.4 | | 6.9 | 8.1 | 7.7 | 3.4 | 2.7 | 3.4 | |
| 6.6 | 4.6 | 9.6 | 9.6 | 9.6 | 6.6 | 6.6 | 6.6 | 5 |
| | | 7.1 | 7.1 | | | | | |
| 4.1 | | 5.5 | 6.2 | 5.1 | 3.1 | 2.7 | 2.3 | |
| 6.2 | 3.8 | 9.2 | 9.6 | 8.9 | 4.6 | 5.4 | 4.6 | |
| 0.2 | 4 | 9.6 | 9.6 | 9.6 | 0.4 | 1.9 | 1.5 | 1.9 |
| 5.5 | | 6.3 | 6.7 | 6.3 | 3.1 | 1.6 | 3.7 | |
| 5.8 | | 8.3 | 7.5 | 7.3 | 5.8 | 6.1 | 5.8 | 3 |
| 0.5 | | 1 | 1.5 | 1.7 | 3 | 3.7 | 3.9 | |
| 3.7 | | 4.5 | 5.6 | 4.6 | 2.7 | 3.5 | 3.1 | |
| 5.1 | 4.2 | 6.8 | 7.2 | 5.7 | 4.3 | 4.7 | 3.9 | |
| 3.3 | | 4.3 | 4.6 | 4.9 | 2.1 | 1.7 | 2.1 | |
| 4.3 | 1.1 | 7.3 | 7.7 | 7.3 | 5.2 | 5.5 | 5.8 | |
| 2.8 | | 5.2 | 5.2 | 4.6 | 0.7 | 0.7 | 0.9 | 1.1 |
| | | 2.3 | 3.1 | 2 | | 4.8 | | |
| 2.9 | | 3.3 | 4.4 | 2.9 | 2.1 | 1.1 | 2.1 | |
| 4.5 | | 7.1 | 7.5 | 6.4 | 3.3 | 3.6 | 5.8 | |
| 5.1 | 0.8 | 6.8 | 6.5 | 6.6 | 5.8 | 5.5 | 5.8 | 0.8 |
| | | 8.5 | 8.5 | 8.1 | | | 1.5 | |
| 5.1 | | 6.1 | 7.6 | 5.4 | 4.7 | 2 | 2.8 | |
| 4.5 | 1.5 | 4 | 4 | 3.6 | 0.6 | 0.5 | 0.2 | |
| | | 7.5 | 9 | 8.3 | | | | |
| 6.6 | 4.4 | 9.6 | 9.6 | 9.6 | 6 | 6 | 6.4 | 4.4 |
| 1.1 | | 7 | 6.7 | 7.8 | 1.5 | 4.5 | 1 | |
| 0.2 | | 3.5 | 5 | 3.9 | 0 | 2.3 | | |
| | | 1.7 | 2.1 | 4.5 | 4.5 | 0.2 | 0.5 | |
| 6.2 | | 8.4 | 8 | 7.6 | 5.5 | 4.7 | 5.1 | |
| 6.6 | 2.3 | 7.6 | 6.9 | 7 | 3.1 | 4.3 | 5.8 | 1.2 |
| 4.5 | 0.8 | 4.5 | 4.2 | 2.9 | 1.7 | 3.9 | 2.7 | |
| 5.8 | | 7.5 | 6.3 | 7.5 | 5.5 | 5.1 | 5.5 | 3.8 |
| 5.8 | | 3.7 | 3 | 3.9 | 0.6 | 0 | | 2.7 |
| 5.1 | | 4.2 | 3.6 | 4.2 | | | | 2.7 |
| 5.1 | | 2.8 | 2.6 | 2.6 | | | | 4.4 |
| 5.8 | 2.7 | 8.7 | 8.7 | 7.6 | 5.8 | 5.8 | 5.8 | 2.7 |
| 4.5 | | 3 | 2.9 | 3 | 1.4 | 1 | 1.7 | |
| 6.6 | | 7.6 | 7.2 | 6.8 | 6.2 | 5.8 | 3.6 | |
| 4.2 | 2.7 | 6.5 | 5.4 | 5 | 3.9 | 4.8 | 4.5 | 0.8 |
| 6.6 | | 9.6 | 9.6 | 9.6 | 6.4 | 6.2 | 6.2 | |
| 5.1 | 5.7 | 6.8 | 7.6 | 6.8 | 4.8 | 5.1 | 4.3 | |
| 6.6 | 2.7 | 7.2 | 7.6 | 3.8 | 6.2 | 5.1 | 5.5 | 3 |
| | 1.9 | | | 7 | 3.3 | 5.5 | 4.3 | |
| 4.6 | | 7.5 | 6 | 6.4 | 4.2 | 3.1 | 3.5 | |
| | | 5.9 | 5.5 | 5.1 | | | | |
| 2.6 | | 4 | 4.7 | 3.6 | 2.9 | 1.8 | 0.5 | |
| 1.9 | | 6.8 | 6.4 | 5.6 | 0.2 | 4.5 | 0.2 | |
| 3.9 | | 6.9 | 6.1 | 6.9 | | | | |

| | | | | | | | | |
|---|---|---|---|---|---|---|---|---|
| 1.4 | | 1.4 | 0.5 | 1.8 | 1.4 | 0.6 | 1.4 | |
| 6 | 3 | 8.3 | 8.7 | 8.3 | 5.8 | 6.4 | 6.4 | 0.8 |
| 4.5 | | 6.6 | 5.9 | 7 | 3.5 | 3.9 | 3.9 | |
| 5.8 | 1.9 | 4.6 | 5.8 | 6.5 | 1.9 | | 1.9 | 4.5 |
| 1.7 | | 4.8 | 5.4 | 4.8 | 3.3 | 2.8 | 2.6 | |
| 1.3 | | 1.4 | 0.9 | | 0.7 | 1.8 | 1.8 | |
| 0.7 | | 1.6 | | 3.6 | 0.8 | 0.7 | 0.5 | |
| 1.1 | | 4.1 | | | 0.8 | 0.7 | 0.7 | |
| 3.3 | | 6.8 | 3.8 | 4 | 0.8 | 4.5 | | |
| 0.7 | 0.8 | | 5 | 6.5 | | 0.8 | | 0.8 |
| 3.9 | 4.9 | 8.8 | 6.5 | 8 | 4.8 | 4.8 | 2.3 | 1 |
| 2 | 0 | 6.5 | 5.1 | 4.7 | 5.2 | 4.8 | 6.2 | |
| 3.1 | | 3.4 | 6 | 4.1 | 1 | 0.6 | 2.5 | |
| 1.1 | | | 1.7 | | 1.4 | 1.7 | 1.8 | |
| 2.3 | | 7.5 | 8 | 7.9 | 6.6 | 6.6 | 6.6 | |
| 0.2 | 2.3 | 8.4 | 8 | 8.7 | 0.6 | 1.9 | 0.6 | |
| | 4.2 | 7.2 | 6.8 | 7.6 | 6.2 | 5.8 | 6.2 | 0.2 |
| 2.3 | 0.2 | 2.7 | 2.3 | 3.4 | 5.1 | 5.5 | 5.5 | |
| 1.8 | 0 | | 4.5 | | | | | |
| 1.2 | | 3.5 | 2.2 | 3.4 | 1.4 | 1.4 | 2.1 | |
| 1.5 | | 5.5 | 5.9 | 6.2 | 4.3 | 4.7 | 4.7 | |
| | | | | 3.4 | 1.3 | | | 0.2 |
| 3.2 | 5.9 | 7.7 | 7.3 | 5.8 | 3.3 | 3 | 3.7 | 5.9 |
| 2.2 | | 5.5 | 5.3 | 5.5 | 1.2 | 2.1 | 2 | 2.7 |
| 4.9 | | 3 | 3.9 | 4.1 | 5.2 | 5.2 | 4.5 | |
| 2.7 | | 3.9 | 2.7 | 3.3 | 3.7 | 3.3 | 3.7 | |
| | | | | | 4.5 | 3.7 | 4.6 | |
| 0.6 | | 3.9 | 3.5 | 3.9 | 1.4 | 2.2 | 1.8 | |
| 5.6 | | 4.9 | 2.2 | 2.4 | 2.7 | 4 | 3.3 | 4.5 |
| 3.9 | | 6.8 | 4.9 | 6.8 | 0.9 | | | |
| 1.9 | 3.6 | 8.4 | 8.4 | 8.7 | 4.5 | 1.7 | 0.6 | 1.5 |
| 0.2 | | 1.7 | 1.5 | 1.8 | 3 | 2.6 | 3.3 | |
| 0.6 | 5.4 | 9.1 | 8.7 | 9.1 | 2.1 | | 0.2 | 1.9 |
| 1.9 | 5.3 | 8.7 | 9.1 | 8.7 | 0 | 4.2 | | |
| 4.3 | 0.2 | 5.3 | 3.8 | 4.9 | 3.2 | 3.6 | 4.7 | 0.4 |
| 6.6 | 4.2 | 9.6 | 9.2 | 9.2 | 6.6 | 6.2 | 6.6 | 3.4 |
| | 1.6 | | 4.4 | 4.4 | 3.2 | 5.1 | | 1.1 |
| 2.4 | 1.9 | 3.2 | 3.9 | 3.6 | 2.8 | 2.2 | 3 | |
| 6.6 | 2.7 | 8.7 | 8.7 | 8.4 | 3.6 | 3.1 | 3.9 | |
| 6.6 | | 8.4 | 8.4 | 8 | 4.9 | 6.8 | 5.7 | 6.8 |
| 6.6 | 4.5 | 8.7 | 9.1 | 8.7 | 6.2 | 6.2 | 6.2 | 1.5 |
| 3.7 | | 3.1 | 1.9 | 3.2 | 1 | 0.2 | 0.6 | |
| 4.5 | 1.7 | 8 | 8 | 8.7 | 0.2 | 4.5 | 4.5 | |
| 6.6 | 4.5 | 8.7 | 9.1 | 8.7 | 5 | 5 | 5 | 3 |
| 0.5 | 2.1 | 8 | 8.4 | 6.1 | | 0.2 | 0.8 | |
| 1.7 | 4.9 | 9.6 | 9.6 | 9.6 | 1.1 | 3.8 | 1.9 | |
| 5.6 | 2.1 | 9.2 | 9.2 | 9.2 | 2.3 | 4.5 | 2.7 | |
| 6.6 | 3 | 8.9 | 9.6 | 9.2 | 6.2 | 5.8 | 6.2 | |
| 2 | 3.5 | 9.2 | 8.9 | 8.5 | | 4.7 | | |
| 3.9 | | 3.9 | 4.7 | 0.7 | 3 | | 3 | |
| 4.5 | 1.5 | 8.4 | 8 | 7.2 | 5.8 | 6.2 | 6.2 | |

| | | | | | | | | |
|---|---|---|---|---|---|---|---|---|
| 1.5 | 2.7 | 4.9 | 4.9 | 7.6 | 4.5 | 0.2 | 0.2 | |
| 5.6 | 1.9 | 8.7 | 8.4 | 9.1 | 5.4 | 4.6 | 5 | |
| 4.7 | 2 | 5.3 | 6.5 | 5.7 | 2.8 | 2.8 | 2.4 | 2.3 |
| 3.3 | | 6 | 5.2 | 6.7 | 2.8 | 3.3 | 3.6 | |
| 5.8 | | 8.1 | 8.1 | 8.5 | 5.1 | 3.9 | 3.6 | |
| 3.9 | 1.6 | 4.3 | 4.3 | 4.7 | 3.9 | 2 | 3.6 | 1.4 |
| | | 2.8 | 1.2 | 3.3 | 1.7 | | | |
| 6.4 | | 8.4 | 8.4 | 8.4 | 6.4 | 6.4 | 6.4 | 2.3 |
| | 1.9 | 5.2 | 4.7 | 4.3 | | | | 1.9 |
| | 4.5 | 5.7 | 7.2 | 7.2 | 0.2 | 1.5 | 2.7 | |
| 5.8 | 1.1 | 8.7 | 8.7 | 8.4 | 6.2 | 5.8 | 5.8 | |
| 5 | | 7 | 7.3 | 7.7 | 1.6 | 0.8 | 2.3 | |
| 1.2 | 0.4 | 8.9 | 9.2 | 8.9 | 2.7 | 3 | 2.3 | |
| 4.7 | 2.3 | 3.8 | 4.2 | 4.2 | 3.2 | 1.3 | 2.4 | 3.1 |
| 3.9 | 2 | 3.8 | 4.2 | 3.4 | 3.9 | 2 | 2.4 | 3.5 |
| 6.6 | | 7.7 | 7.7 | 7.5 | 4.8 | 3.3 | 3.3 | |
| 6.6 | 4.9 | 8.5 | 8.5 | 8.9 | 6.4 | 6.4 | 6.4 | 4.9 |
| 5.4 | | 8.1 | 8.1 | 8.1 | | | | |
| | 2.3 | 3.1 | | 2.6 | 5.2 | 4.8 | 5.5 | 1.1 |
| | | 6.1 | 6.1 | 7.2 | 3 | 3.3 | 4.5 | |
| 6 | | 8.7 | 8.5 | 8.5 | 6.6 | 6.2 | 6.6 | |
| | | 5.6 | 5.6 | 6.7 | 1.8 | 3.3 | 4.7 | 1.9 |
| 6.2 | 1.1 | 8.9 | 8.9 | 8.5 | 5.5 | 5.8 | 6.2 | 4.5 |
| 6.6 | | 9.1 | 8.7 | 9.1 | 5.2 | 3.7 | 2.6 | |
| 3.3 | 2.3 | 8.4 | 8.4 | 8 | 4.5 | 1 | | 0 |
| 4 | | 8.1 | 8.3 | 8.5 | 6.2 | 6.2 | 5.1 | |
| | 1.5 | 5.9 | 5.1 | 5.9 | 6.6 | 6.6 | 6.2 | 4.5 |
| 4.5 | | 6.2 | 5.9 | 7 | 6.6 | 5.5 | 5.8 | |
| 6 | | 1.7 | 2 | 1.1 | 0.5 | 0.8 | 0.8 | 2.3 |
| 4.8 | | 7.8 | 7.5 | 7.1 | 2.1 | 3.5 | 4.8 | |
| 4.3 | 0.6 | 4.8 | 4.8 | 5.2 | 3.2 | 3.7 | | 1.1 |
| 1 | | 6.5 | 7.6 | 6.5 | 0.2 | 1.5 | 0.2 | |
| 1.8 | 2.7 | 6.6 | 6.6 | 6.2 | 2.7 | 2.7 | 2.7 | |
| 5.8 | | 7 | 6.6 | 7 | 3 | 3.8 | 2.7 | |
| | | 4.7 | 4.7 | 5.5 | 3.3 | 3 | 3 | |
| 3.6 | | 5.1 | 5.1 | 5.1 | 2.4 | 2.8 | 2.8 | |
| 4.6 | 3.4 | 8.3 | 8.3 | 8.3 | 4.8 | 2.3 | 1.5 | 3 |
| 3.9 | | 4.8 | 4.1 | 4.1 | 2.2 | 4.8 | 0.5 | |
| | 4.9 | 3.1 | | 3.9 | 3 | 2.1 | 2.9 | |
| | | 6.1 | 7.3 | 5.8 | | | | |
| 4.6 | 4.5 | 7.8 | 7.4 | 6.6 | 3.6 | 5.2 | 4 | |
| 0.7 | | 1.7 | 1.7 | 2.8 | 5.8 | 5.5 | 3.9 | |
| 0.8 | | 6.1 | 6.5 | 6.5 | | 3.3 | 2.6 | |
| | | 6.1 | 6.5 | 6.5 | 3.9 | 3.6 | 3.2 | |
| 2.1 | 4 | 7 | 6.3 | 7.2 | 3.4 | 1.1 | 4.5 | |
| 4.8 | 4.9 | 4.7 | 4 | 5.5 | 2.9 | 4.8 | 3.3 | 4.5 |
| 5 | | 8.1 | 8.1 | 7.4 | 5.2 | 4.6 | 3.5 | |
| 3.3 | | 6.4 | 7.1 | 7.5 | 1.4 | 2 | 2.7 | |
| 5.6 | | 8.1 | 7.8 | 7.8 | 4.8 | 5.2 | 2.9 | |
| 5.2 | | 8.5 | 8.5 | 8.5 | 5.6 | 6 | 6.2 | |
| 1.5 | | 7.8 | 6.2 | 6 | 2.3 | 1.5 | 1.5 | |

| | | | | | | | | |
|---|---|---|---|---|---|---|---|---|
| 3.3 | | 5.6 | 5.4 | 5.2 | 3.6 | 3.9 | 5.5 | |
| | | 5 | 4.4 | 5.4 | | 4.5 | 1.9 | |
| 2.3 | | 7.5 | 7.9 | 7.9 | 0.8 | 1.5 | | |
| 2.7 | | 4.8 | | 3.6 | 4.8 | 1.7 | 5.5 | |
| 6 | 4.5 | 8.6 | 9 | 9 | 6 | 6 | 6 | 3.8 |
| | | 2.8 | 3.2 | 3.2 | 1.7 | | | |
| 2.4 | 0.6 | 2.4 | 0.5 | 2.1 | 1.7 | 4.4 | 2.9 | 1.5 |
| | | 6.9 | 7.7 | 7.7 | | 1.9 | 0.2 | |
| 5.6 | | 7.6 | 7.6 | 8 | 3.5 | 4.2 | 5.4 | |
| 1.7 | | | | | 2.6 | 0.8 | 1.1 | |
| 2 | | 7.9 | 7.9 | 7.1 | 1.5 | 1.5 | 1.5 | |
| 4.5 | 1.5 | 5.7 | 5.5 | 6.6 | 0 | 2.7 | 1.1 | |
| | 0.4 | 5.6 | 5.6 | 6 | 5.5 | 4.8 | 5.5 | |
| 6.6 | 5.3 | 9.2 | 9.2 | 9.2 | 5.2 | 6 | 5.6 | 4.5 |
| 0 | | 7.5 | 7.5 | 7.1 | 2.4 | 5.1 | 5.1 | |
| | | 3.2 | | 3 | 1.4 | | | |
| 1.8 | | 3 | 4.5 | 4.8 | 2 | 1.6 | 1.2 | |
| 4.2 | | 7.9 | 7.5 | 6.8 | 6 | 5.6 | 6 | |
| 6 | | 8.3 | 7.1 | 8.3 | 3.3 | 5.5 | 4.8 | |
| 3.9 | | 8.3 | 8.3 | 6.8 | | | | |
| 4.8 | | 6.7 | 7.5 | 7.1 | 2.3 | 3.1 | 2.3 | |
| 5.2 | | 8.1 | 7.8 | 7.8 | 5.4 | 4.6 | 5.2 | |
| | | 5 | 4.6 | 4.6 | | | | |
| 4.2 | | 8 | 8.8 | 7.7 | 2.3 | 1.9 | 3.8 | |
| 0.8 | | 6.7 | 6.4 | 5.2 | | 2.3 | 2 | |
| 1.9 | | 6 | 7.9 | 6.8 | | | | |
| 6.1 | 5.3 | 8.3 | 7.5 | 8.3 | 5.9 | 5.9 | 5.5 | 4.5 |
| 3.3 | | 4.1 | 3.8 | 5.4 | 2.9 | 2 | 4 | |
| 0.6 | 2.3 | 5.8 | 6.1 | 5.8 | 6 | 6 | 5.6 | 4.5 |
| | | | 2.1 | | 2.5 | 3.8 | 2.2 | |
| 2.8 | 1.9 | 3.3 | 2.7 | 2.6 | 1.2 | 0.5 | 0.5 | |
| | | 5.8 | 5.1 | 5.1 | | 1.4 | | |
| 0.7 | 0 | 3 | 4.9 | 4.9 | 4.5 | 5.1 | 5.1 | |
| 2.3 | 0.4 | 2.8 | 2.8 | 1.7 | 2.2 | 3.3 | 1.7 | 5.3 |
| 3.7 | | 4.3 | 3.4 | 4.7 | 3.9 | 4.5 | 3.7 | |
| | | 6.4 | 8.3 | 6.4 | | | | |
| 5.1 | 1.7 | 7.1 | 7.5 | 6.5 | 4.5 | 4.1 | 4.1 | 1.7 |
| 5.5 | | 7.4 | 7 | 7.4 | 5.8 | 4.6 | 5.4 | |
| 2.1 | | 6.4 | 6.8 | 7.2 | 3.9 | 5.4 | 4.6 | |
| 1.8 | | 3.9 | 3.2 | 2.8 | 2.7 | 2.3 | 2.3 | |
| 5.6 | | 7.2 | 7.2 | 7.6 | 2.9 | 2.9 | 2.1 | |
| 5.2 | | 6.4 | 7.6 | 6 | 1.1 | 2.7 | 4.9 | |
| 3.4 | | 8.2 | 8.2 | 8.2 | 5.4 | 5.4 | 5.8 | |
| 4.5 | | 4.7 | 4 | 4.4 | 4.5 | 0 | 4.2 | |
| 3.6 | | 4.2 | 1.8 | 4.3 | | 2.2 | 1.4 | |
| 4.8 | | 6.6 | 7.8 | 7.4 | 5.6 | 4.8 | 3.3 | |
| 2.2 | | 6.1 | 6.2 | 5.5 | 1 | 1.8 | 1.8 | |
| 0.7 | | 2.2 | 1.4 | 2.8 | 5.2 | 4.8 | 4.5 | |
| 3.3 | | 5.9 | 5.9 | 6.2 | 1.7 | 1.2 | 1 | |
| 4.5 | | 4.5 | 3.7 | 3.7 | | 1.5 | 3 | |
| 1.9 | | 4.4 | 4 | 4.7 | 3.4 | 1.5 | 1.9 | |

| | | | | | | | | |
|---|---|---|---|---|---|---|---|---|
| 5 | | 8.8 | 8 | 8.4 | 3.4 | 3 | 3.4 | |
| 5.6 | 4.5 | 6.9 | 7.2 | 7.2 | 5.6 | 5.6 | 5.2 | 4.5 |
| 4.6 | | 4.1 | 3.8 | 5.4 | 3.6 | 3.6 | 2 | |
| | | 4.7 | 4.3 | 3.9 | 2.2 | 3.7 | 3.3 | |
| 4.2 | | 7.1 | 5.6 | 5.3 | 0 | 0.2 | 0.2 | |
| | | 5.8 | 6.2 | 4.5 | 6.2 | | | 4.5 |
| 5.2 | | 6.7 | 6.4 | 6.4 | 3.1 | 5.2 | 4.5 | 4.5 |
| 3.9 | | 7 | 6.8 | 7 | 2.1 | 2.5 | 2.9 | |
| 5.4 | | 8.3 | 7.9 | 8.3 | | | | |
| 4.1 | | 7.1 | 7.5 | 6.9 | 3.5 | 4.1 | 3.7 | |
| 5.2 | | 6.1 | 5.4 | 6.1 | 1.4 | 2.5 | 2.5 | |
| 4.8 | | 6 | 7.5 | 7.1 | 3.5 | 2.5 | 1.4 | |
| 1 | | 4.9 | 3.4 | 4.1 | 0.2 | 1.5 | 1.5 | |
| 5.8 | | 7.4 | 7.8 | 7.4 | 5.9 | 5.8 | 5.8 | 2.3 |
| 1.5 | | 5.9 | 5.5 | 4.3 | 1.5 | 1.5 | | |
| 3 | | 3.8 | 3.3 | 2.3 | 3 | 2.6 | 4.5 | |
| 4.6 | | 7.4 | 7.8 | 7.4 | 5.8 | 6.1 | 5.4 | |
| 5.5 | 4.9 | 5.2 | 6.3 | 6.7 | 5.5 | 4.4 | 4 | |
| 5.8 | | 8.6 | 7.4 | 5.9 | | 1.9 | | |
| 4.5 | | 6.6 | 4.7 | 4.7 | 3.1 | 2.7 | 1.6 | |
| 4.8 | | 7.8 | 7 | 7 | 3.6 | 3.5 | 2.7 | |
| 3.9 | | 5.3 | 4.5 | 3 | 2.7 | 0.6 | 2.9 | |
| 0.6 | | 3.1 | 3 | 3.7 | 4.5 | 1.9 | 0.2 | |
| 0.4 | | 5 | 4 | 4.6 | 1.9 | 4.5 | 4.5 | |
| 5.2 | 4.9 | 8.4 | 8 | 7.7 | 4.6 | 4.6 | 5.4 | 3.3 |
| 4.6 | | 6.9 | 6.9 | 3.9 | 5.2 | 2.9 | 1.6 | |
| | | 4.8 | 6.3 | 6.3 | 5.4 | 5.6 | 5.6 | |
| 4.2 | | 6.5 | 5.4 | 5.8 | 2.5 | 1.7 | 4 | |
| 1.5 | | 6.1 | 6.5 | 5.8 | 3.8 | 1.9 | 1.9 | |
| 5.8 | | 7.8 | 7.4 | 5.9 | 4.9 | 0.8 | 3 | |
| 5 | | 8.7 | 8.3 | 8.3 | 5.8 | 5.8 | 5.8 | 4.9 |
| 5.4 | | 8.2 | 8.6 | 6.3 | | 4.5 | 4.5 | |
| 5.4 | | 7.4 | 7 | 6.3 | 3.6 | 5.5 | 4.8 | 4.9 |
| 2 | | 1.7 | 2.3 | 2.1 | 2.6 | 3 | 3 | |
| 4.6 | | 6.8 | 6.8 | 5.3 | 4 | 2.3 | 4.4 | |
| 3.6 | 1.9 | 3.7 | 4.1 | 5 | 2.1 | 4.9 | 2.3 | 2.3 |
| 0.2 | | 4.6 | 3.6 | 4.4 | 0.5 | 1.7 | 4.5 | |
| 3 | | 4.6 | 3.7 | 3.7 | 3.5 | 3.1 | 3.5 | |
| 0.7 | | 2.8 | 2.4 | 2.2 | 2.6 | 1.4 | 1.4 | |
| 1.3 | 0.4 | 1.8 | 2.8 | 1.2 | 0.5 | 0.7 | 0.5 | |
| 5.8 | | 6.5 | 6.5 | 5.9 | 2.3 | 0.8 | | |
| 5.9 | 5.3 | 7.6 | 7.2 | 7.6 | 5.9 | 4.8 | 5.8 | 3.8 |
| 0.6 | | 4.6 | 5.5 | 5.8 | 1.5 | 0.2 | 0.6 | |
| 1.6 | | 3.1 | 3.1 | 2.1 | 1.8 | 1.6 | 1.2 | |
| 0.2 | | 2.9 | 3.8 | 2.6 | 3.7 | 3 | 3.7 | |
| 2.7 | 0.8 | 7 | 5.9 | 7 | 5.2 | 5.2 | 5.5 | 2.3 |
| 4.6 | | 6.6 | 6.6 | 6.3 | 2.5 | 1.7 | 2.5 | |
| 0.6 | 1.9 | 3.5 | 2.6 | 4.3 | 5.2 | 4.8 | 5.2 | 1.5 |
| 2 | 4.5 | 3.3 | 2.9 | 2.3 | 2.1 | 1.4 | 2.5 | 4.5 |
| 0.6 | | 6 | 6 | 5.3 | 0.2 | 0.2 | 2.9 | |
| 5.2 | 4.9 | 8.6 | 8.2 | 8.6 | | 4.9 | | 5.3 |

| | | | | | | | | |
|---|---|---|---|---|---|---|---|---|
| 4.5 | | 4.5 | 3.7 | 3.7 | | 1.5 | 3 | |
| 1.5 | | 7 | 6.7 | 6.3 | 5.8 | 5.8 | 5.8 | |
| | 2.7 | | | | 4.5 | | | 3.8 |
| 3.5 | | 7 | 7.8 | 6.2 | 5.2 | 4.5 | 4.8 | |
| 5.4 | | 7.4 | 7.4 | 6.3 | 5.2 | 2.9 | 4 | |
| 2.6 | | 3.6 | 4.7 | 2.6 | 1.7 | 2.1 | 2.7 | |
| 5.2 | | 6 | 7.2 | 6 | 3.5 | 5.8 | 5.4 | |
| 3.9 | | 1.6 | 2.3 | 3.3 | 3.4 | 2.3 | 2.7 | |
| 3.3 | | 5.5 | 6.6 | 6.6 | 3.5 | 3 | 2 | |
| 4.6 | | 7.2 | 6.8 | 6 | 4 | 4 | 3.3 | 4.9 |
| 3.9 | 1.1 | 5.4 | 5.5 | 5.4 | 4.4 | 4 | 3.6 | |
| 3.6 | 1.1 | 4.2 | 2.9 | 2.3 | 3.7 | 2.4 | 3.7 | 1.5 |
| | | 5.6 | 4.9 | 4.9 | 4.5 | 4.5 | 4.5 | |
| 4.1 | | 6 | 6.8 | 6.8 | | 0.6 | 1.9 | |
| 3.6 | | 7.7 | 8.4 | 7.7 | 4.2 | 5.4 | 4.6 | |
| 0.7 | | | 3.2 | | 2.3 | 1.7 | 3 | |
| 4.8 | | 6.6 | 7.6 | 7 | 4.4 | 5.5 | 4 | |
| 5.2 | | 5.1 | 4.7 | 4.9 | 3.5 | 3.1 | 3.1 | |
| 4.5 | | 6.5 | 6.5 | 6.5 | 3.9 | 3.9 | 3.3 | |
| 3.8 | 2.3 | 6.6 | 6.8 | 7.8 | 1.9 | 4.9 | 1.5 | 4.2 |
| 4.8 | | 7.7 | 6.2 | 7.3 | | | 3 | |
| 4.5 | | 2 | 2 | 2.3 | 1.7 | 1.1 | 1.4 | |
| 3.1 | | 6.6 | 4.7 | 5.1 | 3.3 | 3.1 | 2.3 | |
| 3.5 | | 6.5 | 5.9 | 4.4 | 3.3 | 3.1 | 2.3 | |
| | | 5.5 | 3.3 | 5.2 | 5.1 | 2.4 | 2 | |
| 2.1 | | 4.8 | 0.8 | 1.5 | 4.2 | 1.9 | 1.5 | |
| 2.9 | | 5.4 | 3.1 | 4.6 | 1.7 | 2.5 | 2.1 | |
| 2.7 | | 3.9 | 4.8 | 4.4 | | | | |
| 3.9 | | 7 | 6.8 | 7 | 2.1 | 2.5 | 3.3 | |
| 4.4 | | 6.7 | 5.5 | 3.9 | 4.2 | 3.5 | 3.9 | |
| | | 3.8 | 3.6 | 3.2 | 3.3 | 3 | 3.3 | 0.6 |
| 5 | | 7.5 | 6.4 | 7.1 | | | | |
| 2.9 | | 4.9 | 4.1 | 2 | | | | |
| 2 | | 5.5 | 4.7 | 4.7 | 2.8 | 2.8 | 3.6 | |
| 0.5 | | 1.3 | 2.7 | 2.1 | 0.5 | 0.2 | 0.2 | |
| 2.7 | | 1.6 | 4.4 | 1.4 | 0.6 | 4.5 | 0.2 | |
| 2.1 | | 4.8 | 4 | 4.8 | 4.2 | 2.1 | 1 | |
| 2.1 | | 6.4 | 6.8 | 7.2 | 3.9 | 5 | 4.6 | |
| 3.3 | | 5 | 5.4 | 5.7 | 3.9 | 2.7 | 2.7 | |
| 4.8 | | 6.9 | 3.9 | 5.4 | 2.5 | 3.3 | 4.4 | |
| 4.6 | | 6.6 | 6.8 | 6.6 | 4.1 | 4.1 | 4.5 | |
| 4.6 | | 5.1 | 7 | 4.9 | 2.9 | 3.3 | 2.5 | |
| 2.5 | | 1.8 | | 1.2 | 3.3 | 3 | 3 | |
| 4.6 | 0.8 | 5.1 | 5.5 | 7 | 1.7 | 3.3 | 3.3 | |
| 1.4 | | 4.5 | 4.3 | 6 | 1.7 | 1.7 | 1.4 | |
| 3 | | 4.6 | 6.5 | 8.1 | 5.4 | 3.5 | 4.4 | |
| 0.5 | | 1.7 | 1.7 | 2.1 | 0.2 | | 0.5 | |
| 2.9 | | 5.6 | 5.3 | 5.6 | 4.6 | 4.6 | 4.2 | |
| 5.2 | | 6.7 | 7 | 6.3 | | | | |
| 3.9 | 4.5 | 6.3 | 3.6 | 3.6 | 3.4 | 0.6 | 3.4 | |
| 0.8 | | 4.8 | 4.4 | 3.3 | 0.8 | 3 | 3.8 | |

| | | | | | | | | |
|---|---|---|---|---|---|---|---|---|
| 4.6 | | 5.8 | 4.8 | 6.3 | | | | |
| 2 | | 5.1 | 4.4 | 3.6 | | 1 | | |
| 2.8 | | 2.2 | 2.7 | 1.2 | 1.6 | 0.8 | 1.6 | 0.4 |
| 2.2 | | 2.2 | 3.1 | 2.2 | 2 | 3.3 | 2.6 | |
| 1.4 | | 2.6 | 1.6 | 1.6 | 2.2 | 2 | 1.8 | |
| 3.6 | | 6.4 | 6 | 5.7 | 4.9 | 2.5 | 4 | |
| 3.8 | 2.7 | 0.8 | 2.9 | 2.9 | 0.2 | 0.5 | 1.5 | |
| 4.6 | | 8.3 | 7.7 | 6.9 | 5.2 | 4.4 | 3.6 | |
| | | 2.6 | 3.4 | 2.2 | | | 1.5 | |
| | | 4.6 | 3.9 | 6.2 | 3.1 | 4.6 | 3.9 | |
| | | 4.5 | 3 | 5.3 | | | | |
| 2.5 | | 3.9 | 4.6 | 3.9 | 4.6 | 5 | 4.2 | |
| 3.1 | | 4.9 | 4.5 | 3.5 | 3.3 | 2.1 | 1.4 | |
| 1.4 | | 3.2 | 3.4 | 3 | | | | |
| 4.5 | | 6.5 | 5.5 | 4.4 | 2.9 | 1.4 | 1 | |
| | 0 | 3 | 4.9 | 4.9 | 4.5 | 5.1 | 5.1 | |
| 2.1 | | 6.2 | 7.7 | 5 | 3.3 | 3.3 | 1.4 | |
| 3.3 | | 5.9 | 5.9 | 6.2 | 1.7 | 1.2 | 1 | |
| 5.8 | | 7.4 | 6.7 | 7.4 | 4 | 4 | 4 | |
| 0.6 | | 1.6 | 1.2 | 4.4 | 0.2 | 0.4 | 4.5 | |
| 1.5 | | 2.9 | 0.6 | 2.5 | 3.1 | 4.6 | 3.1 | |
| 3.9 | 3.8 | 6.9 | 5.4 | 6.2 | 4.9 | 4.5 | 2.7 | 3 |
| 1.6 | | 1.1 | 2 | 2.3 | 1.7 | | 2.1 | |
| 1.5 | 1.5 | 5.5 | 2.1 | 3.6 | 0.2 | 1.9 | 2.3 | 0 |
| 4.5 | | 1.2 | 0.6 | 2.5 | 2.3 | 2.7 | 3 | |
| 0.2 | | 2.4 | 3.1 | 3.1 | 1.5 | 0.8 | 1.9 | |
| 3.6 | | 6 | 7.2 | 7.2 | 5.5 | 4 | 4.4 | |
| 3.3 | | 2.3 | | 1.6 | 2.3 | 2.7 | 2 | |
| 4.5 | | 4.7 | 5.3 | 5.3 | 4.7 | 3.2 | 4.7 | |
| 3.5 | | 4.9 | 4.5 | 5.3 | 1.9 | | | |
| 0.2 | | 3.8 | 1.9 | 1.9 | 0.2 | 0.2 | 0.2 | |
| 3.9 | | 5.8 | 6.5 | 4.6 | 5 | 3.9 | 3.9 | |
| | | 4.6 | 4.6 | 4.6 | | | | |
| 4.2 | | 5 | 4.2 | 4.8 | 2.1 | 1.4 | 1.7 | |
| 3.5 | | 4.2 | 5.5 | 4.7 | 3.3 | 3.5 | 3.3 | |
| 5.5 | | 5.5 | 6.7 | 6.3 | 4.9 | 3.4 | 3 | |
| 3.9 | | 5.4 | 4.8 | 5.2 | 1.4 | 3.3 | 2.9 | |
| 0.6 | | 3.4 | 3.4 | 4.1 | | 0.2 | | |
| 2.7 | | 6.2 | 6.2 | 6.2 | | 4.9 | 4.9 | |
| 3.6 | | 5 | 5.4 | 7.3 | 2.9 | 3.3 | 3.3 | |
| 5 | | 4.7 | 4.7 | 3 | 1.4 | 2.5 | 1.4 | |
| 3.9 | | 5.1 | 4.7 | 4.4 | 1.4 | 1.4 | 1.4 | |
| | | 2.2 | 1.6 | 1.5 | 2.7 | 1.2 | 2.6 | |
| 1 | 1.1 | 4.1 | 4.5 | 4.1 | 1.9 | 1.1 | 1.5 | |
| 4.6 | | 7.9 | 6.4 | 5.6 | 4.8 | 5.2 | 3.1 | |
| 3.1 | | 8.2 | 7 | 6.7 | 2.3 | 1.1 | 3.8 | |
| 4.8 | | 2.9 | 4 | 4 | 2.3 | 2.9 | 1 | |
| 2.8 | | 2.2 | 2.7 | 1.2 | 2.5 | 0.8 | 1.6 | |
| 3 | | 3.4 | 5.7 | 3.8 | 1.5 | 4.9 | 0.4 | |
| 3.1 | | 3.2 | 2.5 | 2.9 | 2.9 | 2.3 | 3.1 | |
| 3.5 | | 2.2 | 2.2 | 3.2 | 0.2 | 0.2 | 0.6 | 3.4 |

| | | | | | | | | |
|---|---|---|---|---|---|---|---|---|
| 4.6 | | 2.2 | 2.6 | 3.1 | 0.6 | 1 | 0.2 | |
| 4.9 | | 4.2 | 6.1 | 4.8 | 5.2 | 5.2 | 4.8 | |
| 2.9 | | 2.9 | 2.3 | 2.1 | 4.2 | 1.1 | 0 | |
| 2.9 | | 5.2 | 5.5 | 5.5 | | 4.5 | | |
| | | 6.2 | 4.6 | 5.4 | 4.8 | 4.6 | 5.4 | |
| 1.6 | | 4.5 | 4.5 | 4.5 | 1.8 | 0.5 | 0.5 | |
| 1.3 | | | 4.2 | 4.6 | | 2.4 | | |
| 4.5 | | 5.3 | 5.3 | 4.9 | 4.5 | 3 | 0.6 | |
| 3 | | 6 | 4.1 | 4.9 | | | | |
| 4.5 | | 1.6 | 1.7 | 0.8 | 0.6 | | 0.5 | 1.5 |
| 4.5 | | 2 | 1.7 | 2.1 | 0 | 4.5 | 1.5 | |
| 3.1 | | 5.4 | 5.4 | 5 | 2.3 | 2.5 | 1 | |
| 0.4 | | 2.5 | 1 | 2.9 | 3.9 | 3.9 | 4.6 | |
| 5.2 | | 5.9 | 8.2 | 7.8 | 5.3 | 4.9 | 4.5 | |
| 3.6 | | 6.3 | 5.9 | 7.4 | 3.6 | 3.3 | 4.4 | |
| 2 | | 3.3 | 3.1 | 4.8 | 1.4 | 2 | 2 | |
| 4 | | 5.8 | 6.2 | 6.2 | 5.5 | 5.2 | 5.2 | |
| | | 2.8 | 2.4 | 4.3 | | | | |
| 4 | | 7.4 | 5.5 | 7.4 | 3.6 | 2.9 | 2.1 | |
| 1.6 | 0.6 | 1.3 | 1.9 | 1.9 | 0.2 | 0.4 | 0.2 | |
| 4.4 | | 8.2 | 5.2 | 7 | 1.4 | 4 | 2.1 | |
| 1.4 | | 2.9 | 2.9 | 4.5 | 0.2 | | 0 | |
| 3.6 | | 8.2 | 7 | 6.3 | 4 | 4.8 | 2.9 | |
| 4 | | 8.2 | 7 | 8.2 | 3 | | | |
| 0.2 | | 0.7 | 0.7 | 0.7 | 3.1 | 2.7 | 3.1 | |
| 4 | | 5.8 | 6.9 | 5.8 | | 2.7 | | |
| 0.2 | | 3.6 | 4 | 4 | | 0.2 | 0.2 | |
| 2.9 | | 5 | 4.6 | 3.9 | 1 | 3.1 | 2.3 | |
| 4.5 | | 1.4 | 0.2 | 0.8 | 1.5 | 1.5 | 0.5 | |
| 5.2 | | 7.3 | 7.3 | 7.3 | | | | |
| | | 5.5 | 4.8 | 5.5 | | | | |
| 3.2 | 0.5 | 6.8 | 7.2 | 6.5 | 4.8 | 4.7 | 3.9 | 0.6 |
| 1.8 | | 5 | 5.4 | 6.9 | 1.4 | 2.3 | 2.7 | 0.6 |
| 2.7 | | 3.5 | 2.6 | 5 | 1.2 | 1.2 | 2 | |
| | | 4 | 2.7 | 3.2 | | | | |
| | | | | 0.8 | 2.8 | 2.8 | 2.4 | |
| | | 1.9 | | | 1.7 | 2 | 2 | |
| | 2.2 | 7.1 | 8.3 | 8.5 | | | | 0.7 |
| 2.2 | | 2.6 | 1.6 | 3.1 | 3.9 | 3.1 | 2 | |
| 1.7 | 0.8 | 4.8 | 5.6 | 4.8 | 5.1 | 5.1 | 5.5 | 0 |
| 0.7 | | 1.1 | 0.9 | 2.8 | 0.5 | | | |
| | | 1.6 | 1.2 | | 3.2 | 2.8 | 3.2 | |
| 4.5 | 3.4 | 7.3 | 5.8 | 7.3 | 5.2 | 4.8 | 5 | |
| | | 7.7 | 5.8 | 5.4 | 5.4 | 6 | 5.4 | |
| 4.6 | 6.6 | 8.7 | 8.7 | 8.7 | 3.9 | 3.6 | 2.9 | 5.4 |
| 3.3 | 6.3 | 8.7 | 8.7 | 8.7 | 0.2 | | 1.9 | 3.8 |
| 6 | | 8.6 | 8.3 | 8.3 | 5.6 | 6 | 5.6 | 1.9 |
| 0.7 | | 6.1 | 5.4 | 5.9 | 5.5 | 5.5 | 5.5 | |
| 4 | | 7.8 | 5.9 | 7.4 | 0.4 | 4.5 | 3 | |
| 1.1 | | 1.1 | 0.9 | 1.3 | 0.7 | 0.5 | 0.8 | |
| 1.8 | | 2.2 | 2.6 | 3 | 1.4 | | | |

| | | | | | | | | |
|---|---|---|---|---|---|---|---|---|
| 4.8 | | 6.9 | 6.5 | 5.7 | 3.9 | 4.6 | 3.9 | |
| 1.4 | 6 | 9.1 | 9.1 | 9.1 | 0.4 | 1.9 | 4.9 | 3 |
| 1.3 | | 1.8 | 1.7 | 2 | 5.1 | 5.1 | 5.5 | |
| 0.7 | | 2.2 | 2.2 | 2.2 | 2 | 1.8 | 2.2 | |
| 5 | | 7.1 | 6.4 | 6.4 | 4 | 3.6 | 4 | |
| 1.7 | 0.4 | 4.8 | 2 | 3.3 | | 0.2 | | 1.9 |
| 1.1 | | 3.6 | 4.3 | | 1.4 | 1.4 | | |
| 5.2 | | 3.8 | 2.9 | 2.5 | | | | 5.4 |
| 0.4 | | 3.1 | 2.7 | 3.9 | 1.5 | 1.5 | 2.7 | |
| 5.2 | | | 7.7 | 6.5 | 4.2 | 3.4 | 3.8 | |
| 0.6 | 2.7 | 4.3 | 5.4 | 6.2 | 3 | 0.2 | 2.3 | 1.9 |
| 2.3 | 4 | 9.1 | 8.5 | 9.1 | 4.8 | 4.5 | 5.4 | |
| 4.9 | | 5.4 | 6.2 | 5 | | | | |
| 5.2 | | 6.6 | 7 | 6.6 | | 0.4 | | |
| | 0.8 | 1.2 | | | | | 0.2 | 0.4 |
| | | 2.1 | 2.1 | 1.7 | 0.5 | 0.5 | 0.5 | |
| 3 | 1.9 | 4 | 4 | 3.2 | | 0.5 | 0.5 | 0 |
| 6.4 | 4.9 | 8.9 | 8.1 | 8.9 | 5.8 | 6 | 5.6 | 3.8 |
| 2.6 | 0.8 | 5.9 | 4.7 | 4.7 | 2.2 | 2.3 | 3 | |
| 5.6 | | 7.6 | 8 | 8 | 4.5 | 5.6 | 5.2 | |
| 6 | | 8.1 | 8.1 | 8.5 | 5.6 | 6 | 6 | |
| 3.8 | | 4.3 | 4.7 | 4 | | | | |
| 2 | 1.5 | 2.7 | 1.3 | 2.5 | 2.5 | 3.5 | 1.6 | |
| 0.2 | 1.9 | 5.7 | 2.2 | 5.3 | 4.5 | 1.4 | 1 | 4.2 |
| 2.3 | | 5.9 | 5.9 | 5.5 | 2.1 | 2.3 | 2.7 | |
| 6.2 | | 9.1 | 8.7 | 8 | 5.2 | 6 | 6 | |
| 5.2 | | 7.9 | 7.1 | 7.5 | 3.6 | 5.2 | 4.4 | |
| 5.8 | 3.4 | 9.1 | 8.1 | 8.5 | 4.2 | 5.2 | 4.8 | 4.9 |
| 1 | 0.4 | 3.4 | 3 | 4.1 | | | | |
| | 1.2 | 4.8 | 4.8 | 5.5 | 3.9 | 1.2 | 4.2 | 1.5 |
| | | 2 | 2.4 | 2 | 3.7 | 3.7 | 4.1 | |
| 2.3 | | 4.1 | 4.9 | 4.1 | 4.8 | 3.6 | 3.6 | |
| 0.6 | | 3.4 | 3.4 | 3 | 3.7 | 2.6 | 3 | |
| | 1.9 | 5.3 | 4.9 | 6 | 1.3 | 3.5 | 2.4 | |
| 2.6 | | 8.5 | 7.5 | 6.7 | 1.6 | 0.8 | 2 | |
| | | 1.9 | 1.6 | 2.2 | 0.5 | | 0.9 | |
| 6.2 | | 8.1 | 8.5 | 8.5 | 6 | 5.2 | 5.6 | |
| 5.6 | 3.4 | 7.7 | 8 | 8 | 3.3 | 4.2 | 4.2 | 2.9 |
| 3.3 | | 4.8 | 4.8 | 4.8 | 1.7 | 1.7 | 1.4 | |
| 4.5 | | 5.4 | 5.8 | 6.5 | 0.6 | | 1.5 | |
| 0.5 | 1.4 | 2.7 | 2.7 | 3.1 | 0.2 | 0.8 | | |
| 3.5 | 2.3 | 2.5 | 3.3 | 5.2 | 2.5 | 1.7 | 4.8 | 3.4 |
| 1.1 | | 6.4 | 6.1 | 4.4 | 5.5 | 5.2 | 3.6 | 4.5 |
| | | 5.8 | 6.2 | 5.4 | | | | |
| 6.2 | | 8.7 | 8.7 | 8.7 | 5.8 | 6.2 | 6.2 | |
| 5.1 | | 8.9 | 8.5 | 8.9 | 5.8 | 5.8 | 5.1 | |
| 0.5 | | 4.8 | 4.8 | 5.5 | 5.5 | 5.5 | 5.5 | |
| 4.5 | | 5.4 | 6.3 | 5.1 | 2.9 | 2.7 | 1.7 | |
| 3.4 | | 2 | 2.4 | 3.9 | 4.2 | 4.6 | 3.1 | |
| 4.5 | | 8.5 | 8.1 | 7.4 | 0.2 | 0 | 4.2 | |
| 6.6 | 1.4 | 8.6 | 8.6 | 8.6 | 6.2 | 6.6 | 6.2 | 1.4 |

| | | | | | | | | |
|---|---|---|---|---|---|---|---|---|
| 5.4 | | 6.9 | 6.9 | 6.9 | 5.2 | 3.6 | 3.3 | |
| 5.4 | | 5.3 | 5.6 | 4.6 | 3.3 | 4 | 5.2 | |
| 4.8 | | 7 | 5.3 | 6 | 4.2 | 5 | 4.6 | |
| 4.8 | 4.5 | 5 | 5 | 6.5 | | | | |
| 4.5 | 0.8 | 7.5 | 7.5 | 7.1 | 3.5 | 2.7 | 3.1 | 0.4 |
| 3.3 | | 4.4 | 5.2 | 4.8 | | | 3.4 | |
| 6.6 | | 8.7 | 8.4 | 8 | 5.8 | 6.2 | 5.8 | |
| 6.4 | 2.7 | 8.3 | 8.3 | 7.1 | 2.7 | | | 5.5 |
| 6.6 | 5.5 | 9.1 | 8.9 | 9.1 | 6.9 | 7.1 | 7.1 | 6.9 |
| 4 | | 4.5 | 5.7 | 5.3 | 2.5 | 2.9 | 2.9 | |
| 4.7 | 2.7 | 6.5 | 6.8 | 5.3 | 5.1 | 5.5 | 2.4 | 3.5 |
| 5.8 | 4 | 7.9 | 7.9 | 7.9 | 4.3 | 5.5 | 5.8 | 4.4 |
| 0.9 | 2.2 | 8.1 | 8.5 | 7.7 | 4.7 | 2 | 5.5 | 2.3 |
| 2.3 | | 7.7 | 7.7 | 7.5 | 5 | 5 | 2.3 | |
| 4.6 | | 7.5 | 6.8 | 7.5 | 4.8 | 4.6 | 5.4 | |
| 6 | 2.7 | 8 | 7.6 | 7.4 | 5 | 5.8 | 5.8 | 2.7 |
| 5 | | 8.4 | 6.5 | 8.4 | 3.9 | 4.2 | 5.4 | |
| 0.7 | | 0.7 | 1.1 | 0.8 | 2 | 2.3 | 2 | |
| 2 | | 2.9 | 1.2 | 2.9 | 1.4 | 1.4 | 1.2 | |
| 3.9 | | 7.2 | 4.5 | 3.8 | | | | 1.5 |
| 3.3 | 0.4 | 3.5 | 3.5 | 3.1 | 0.4 | 4.5 | | 1.5 |
| 2.6 | 0 | 3.2 | 3.1 | 3.2 | 2 | 0.8 | 1.6 | |
| 4 | | 7.4 | 7 | 8.2 | 3 | 4.2 | 4.5 | |
| 2.6 | | 2.6 | 2.6 | 2.8 | 0.5 | 0.5 | 1.1 | |
| 6.2 | 3.3 | 8.6 | 8.2 | 8.6 | 5.8 | 6.6 | 6.2 | 4.5 |
| 4.5 | 4.5 | 5 | 4.3 | 5.4 | 4.5 | | 1.5 | 4.9 |
| 3.7 | | 6.1 | 5.4 | 6.1 | 5.6 | 5.2 | 5.2 | |
| | | 4.6 | 4.2 | 3.9 | 1.5 | 0.2 | 0.4 | |
| 5.2 | 2.7 | 7.5 | 8.3 | 8.3 | 4.8 | 4.6 | 4.6 | 2.3 |
| 0.4 | | 4.5 | 3 | 3.8 | 0 | 0.2 | 0 | |
| 5.8 | | 8.1 | 8.1 | 8.5 | 5.1 | 3.9 | 3.6 | 0 |
| 5.2 | 3.4 | 7.7 | 8.1 | 7.3 | 4.3 | 5.4 | 4.7 | 5.4 |
| 5.4 | | 4.6 | 5 | 6.5 | | 4.9 | 3 | |
| 3.7 | | 7 | 6.6 | 6.6 | 1.7 | 2.1 | 3.3 | |
| 5.6 | | 8.1 | 8.1 | 7.8 | 4.2 | 4.2 | 4.6 | |
| 1.8 | 1.8 | 8.4 | 8.4 | 8.4 | 1.6 | | 2 | |
| 0.5 | 0.2 | 4.8 | 4.8 | 5.5 | 5.5 | 5.5 | 5.5 | |
| 5.1 | | 8 | 8.4 | 8.4 | 4.2 | 3.1 | 3.9 | |
| 4.8 | | 7.9 | 8.3 | 6.5 | 0.6 | 1.4 | 2.1 | |
| 3.6 | | 5.5 | 4 | 4.4 | 1.5 | 0 | 4.5 | |
| | 2.7 | 5.2 | 5.2 | 2.5 | 4.9 | 0.4 | 4.5 | |
| 6.1 | 3.4 | 9.4 | 9.4 | 9 | 6.4 | 6.6 | 5.8 | 4.9 |
| 5.4 | | 7.3 | 6.9 | 7.3 | 3.4 | 2.3 | 2.3 | |
| 5.2 | | 7.5 | 7.9 | 7.9 | | | | |
| 4.6 | 2.3 | 7.1 | 7.1 | 7.1 | 3.9 | 3.5 | 4.2 | |
| 4 | 1.9 | 6.9 | 6.5 | 6.9 | 4.4 | 3.3 | 4.8 | 4.5 |
| 5.8 | | 7 | 6.6 | 7 | 1.7 | 2.3 | 1.6 | |
| 5.5 | | 7.2 | 5.5 | 5.1 | | | | 3 |
| 3.1 | | 3.4 | 4.5 | 4.9 | 3.3 | 2.9 | 1 | |
| 5.1 | | 8.9 | 8.3 | 8.6 | 3.5 | 3.7 | 4.1 | 0 |
| 1.7 | | 2.3 | 4.8 | 2.5 | 4.6 | 3.5 | 4.2 | |

| | | | | | | | | |
|---|---|---|---|---|---|---|---|---|
| 5.2 | | 7.9 | 7.1 | 7.5 | 3.6 | 5.2 | 4.4 | |
| 6.4 | | 7.4 | 8.2 | 5.9 | 4.4 | 5.2 | 3.6 | |
| 6 | 4.9 | 8.2 | 8.4 | 8.4 | 6 | 6.4 | 6 | 4.9 |
| 3.6 | | 8.7 | 9.1 | 8.7 | 2.5 | 4.4 | 4 | |
| 5 | | 7.4 | 7.4 | 5.5 | 3.3 | 3.6 | 2.9 | |
| 1.3 | 0.8 | 2.6 | 2 | 3 | 2.3 | 3.3 | 3 | |
| 2.8 | 0.2 | 2.7 | 3.1 | 1.3 | 1.4 | 2.3 | 1.6 | 1.5 |
| 2.3 | 1.9 | 2.2 | 2.9 | 4 | | 2.5 | 1.2 | 4.5 |
| 5.4 | | 7.2 | 7.6 | 7 | 4.2 | 5.6 | 5.2 | |
| | | 5.1 | 5.1 | 5.1 | 1.6 | 1.6 | 3.1 | |
| 1.7 | | 3.4 | 4.5 | 6 | 2.9 | 4.6 | 4.2 | |
| 0.8 | 0.4 | 6.7 | 4.8 | 4.8 | 1.6 | 1.6 | 2.3 | |
| 2.3 | 1.7 | 3.2 | 5.4 | 5.7 | | 0.8 | 3.6 | |
| 3.8 | | 2.4 | 2 | 1.6 | 4.6 | 2.7 | 3.1 | |
| 3.5 | | 4.1 | 1.8 | 2.2 | 0.6 | 4.5 | 0.2 | |
| 0.8 | | 4.7 | 2.9 | 2.5 | 2 | 2 | 2 | |
| 3.3 | 4.2 | 9.1 | 9.1 | 9.1 | 6.6 | 5.8 | 6.6 | 0.8 |
| 5 | | 8.3 | 8.3 | 7.9 | 4 | 5.5 | 5.2 | |
| 5.8 | | 7.1 | 6.8 | 8.3 | 3.3 | 4.8 | 3.3 | |
| 3.3 | | 3.6 | 4 | 5.9 | 1.4 | 1 | 4.4 | |
| 0.7 | | 2 | 0.8 | | | | | |
| 2.3 | | 6.3 | 6.3 | 5.1 | 2.7 | 3.5 | 4.2 | |
| 4.6 | | 5.2 | 4.1 | 3.5 | 4.1 | 4.5 | 5 | |
| 2.7 | | | 3.4 | 4.3 | 2 | | 1.6 | |
| 4.8 | | 7.4 | 7 | 7.1 | 4.2 | 2.7 | 4.2 | |
| 5 | 1.9 | 8.6 | 6.7 | 7.8 | 4.8 | 4 | 4.4 | 1.5 |
| 6.2 | 1.9 | | 5.7 | 4.6 | 4.8 | 5.2 | 5.6 | 4.5 |
| 1.1 | 1.1 | 3 | 3.8 | 4.5 | 4 | 5.8 | 4.2 | 2.3 |
| 5.6 | | 6.9 | 8.8 | 7.7 | 4 | 4.8 | 5.5 | |
| 1.9 | 3 | 6.3 | 5.1 | 5.5 | 1.4 | 1.9 | 1.9 | |
| 3.9 | | 6.4 | 6.8 | 6.4 | 3.9 | 3.9 | 2.7 | |
| 1.5 | 2.3 | 7 | 7 | 7.8 | 6.6 | 6.2 | 6.6 | |
| 6.6 | 1.9 | 9.4 | 9 | 8.6 | 6.4 | 6 | 6.4 | 4.5 |
| | | 4.3 | 4.3 | 3.2 | 1.9 | 1.6 | 4.2 | |
| 2 | | 1.8 | 1.4 | 2.2 | 2.2 | 1.7 | 1.7 | |
| 1.3 | | 2 | | 1.3 | | 1.4 | | |
| | | | 2.3 | 2.7 | 1.8 | 1.8 | 4.1 | |
| | 2.7 | 5 | 2.4 | 4.3 | | | | |
| 5.2 | | 6.9 | 6.3 | 5.1 | 2.1 | 2.9 | 2.1 | |
| | 0.4 | | 2.1 | 3.2 | 1.7 | | | |
| 1.9 | 0 | 3.2 | 2.6 | 2.6 | 1.1 | 0.2 | | |
| | | 3 | 2.8 | 2.8 | | | | |
| 5.8 | 6.3 | 7.6 | 7.4 | 7.6 | 3.3 | 3.6 | 3.3 | |
| | | 5.9 | 5.9 | 4.8 | | | | |
| 2.7 | | 3.5 | 3.1 | 3.5 | 0.7 | 0.8 | 0.7 | |
| 5.6 | 3.4 | 8.6 | 5.2 | 8.6 | 5.2 | 5.5 | 4.8 | |
| 5.4 | 4.5 | 7.4 | 7.4 | 7.4 | 5.2 | 5.2 | 4 | |
| 5.2 | | 6.1 | 5.4 | 6.1 | 1 | 2.1 | 1.5 | |
| 3.5 | 3.5 | 6.2 | 5 | 4.3 | | | 4.5 | 3 |
| 5.6 | | 8.1 | 8.1 | 8.1 | 5 | 5.4 | 5.4 | |
| 6 | | 9 | 7.5 | 8.3 | 3.9 | 5.4 | 4.6 | 2.3 |

| | | | | | | | | |
|---|---|---|---|---|---|---|---|---|
| 5.8 | | 5.7 | 5.9 | 5.5 | 4.2 | 3.9 | 3.9 | |
| 1.7 | | 6 | 4.8 | 5.2 | 2.1 | 1.4 | 0.2 | |
| 6.1 | 3.4 | 7 | 7.2 | 7.4 | 4.8 | 5.8 | 5.8 | 4.2 |
| 6.6 | 5.5 | 8.6 | 8.4 | 8 | 4.9 | 5.9 | 6 | 6 |
| 6.6 | | 9.6 | 9.2 | 9.2 | 5.6 | 6.4 | 6 | |
| 3.8 | 4.2 | 9.6 | 9.2 | 9.6 | 6.6 | 6.6 | 6.6 | |
| 6.6 | | 8.6 | 7.5 | 7.1 | 5.6 | 5.5 | 5.8 | |
| | | 1.7 | 2.1 | 3.3 | | | | |
| 5.6 | | 5.5 | 6.6 | 5.9 | 4.6 | 4.2 | 3.5 | |
| | | 4.2 | 4.2 | 4.2 | 3.9 | 3.9 | 4.3 | |
| 1.9 | | 4.2 | 5.4 | 5.7 | 4.3 | 4.7 | 4.7 | |
| 5 | | 7.8 | 6.3 | 5.9 | 4.5 | 2.3 | 1.5 | |
| | | 5.7 | 5.7 | 5.7 | | | | |
| 3.1 | 4.5 | 4.6 | 5 | 5.8 | 4 | 3.6 | 4 | 4.5 |
| 3.9 | 4.9 | 8.8 | 6.5 | 8 | 4.8 | 4.8 | 2.3 | 1 |
| 5.5 | | 6.3 | 6.1 | | 3.5 | 4.1 | | |
| 5.2 | | | | 6.2 | 3.5 | 3.7 | 3.3 | |
| 6.6 | 1.9 | 7.9 | 7.9 | 8.3 | 6 | 6 | 4.5 | 4.4 |
| 3.3 | | 8.4 | 6.1 | 7.7 | 5.6 | 5.6 | 5.2 | |
| 1.9 | | 5.4 | 5.4 | 3.9 | 1.7 | 5 | 5.4 | |
| 4.4 | | 6.8 | 6.4 | 5.6 | 4.6 | 3.9 | 4.2 | |
| 2.7 | | 7.1 | 7.3 | 7.3 | 5.8 | 6 | 6.4 | |
| 6.6 | 2.9 | 9.6 | 9.6 | 9.6 | 6.6 | 6.6 | 6.6 | 3 |
| 5.4 | | 8.7 | 8.3 | 7.9 | 5.4 | 5.4 | 5.4 | |
| 4.6 | | 5.6 | 6.5 | 5.8 | 2.5 | 4.8 | 3.6 | |
| 5.6 | | 8.6 | 8.2 | 8.2 | 6.6 | 6.6 | 6.6 | |
| 3.1 | 2.3 | 8.4 | 7.6 | 8.4 | 4.5 | 4.4 | 5.2 | |
| 4.6 | | 5.5 | 5.3 | 4.1 | 3.9 | 3.7 | 4.1 | |
| 4.8 | 2.7 | 7.9 | 7.7 | 7.9 | | 3.2 | | |
| 4.1 | | 5.9 | 5.3 | 4.9 | 2.3 | 2.3 | 4.2 | |
| | | 6.9 | 8.1 | 6.5 | 5.8 | 5.4 | 5.4 | |
