## Supplementary Metabolite Catalogue 1 for "Non-target Analysis of Wastewater Treatment Plant Effluents: Chemical Fingerprinting as a Monitoring Tool"

### 2-ethylhexyl salicylate (octisalate)

Chromatogram showing prevalence in samples

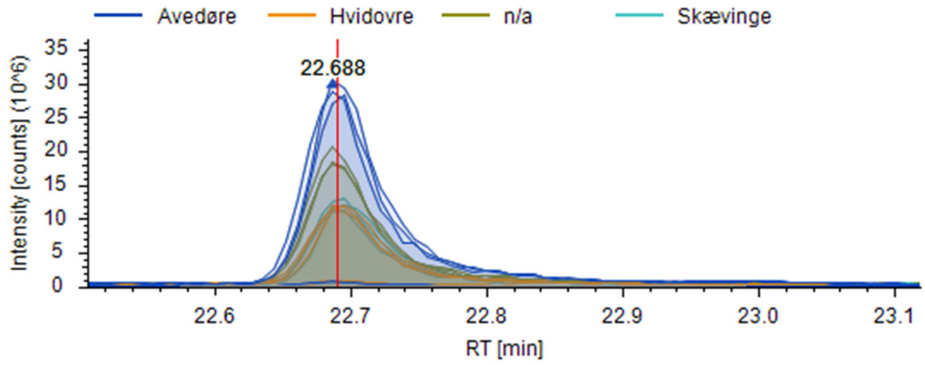


FISh coverage

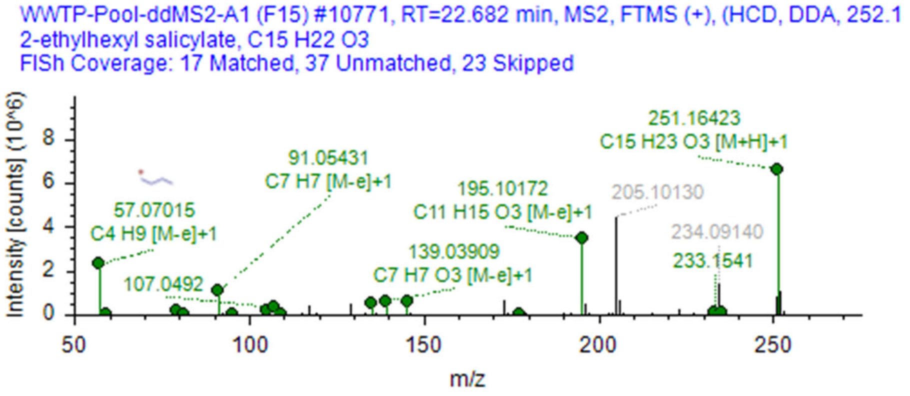


mzVault reference.

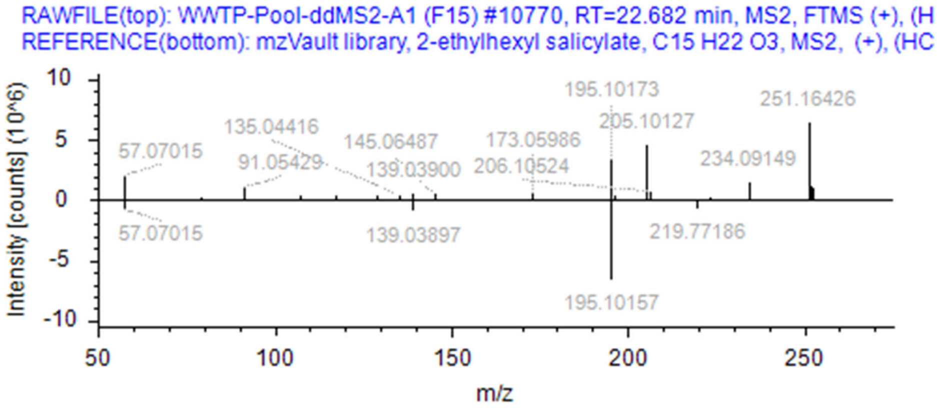

### Metabolites:

2-ethyl-hydroxyhexyl salicylate, $C_{15}H_{22}O_4$,

ref: 10.1007/s00204-019-02537-z

10.1016/j.toxlet.2019.04.001

2-ethyl-5-hydroxyhexyl 2-hydroxybenzoate

2-ethyl-5-oxohexyl 2-hydroxybenzoate, $C_{15}H_{20}O_4$

5-(((2-hydroxybenzoyl)oxy)methyl)heptanoic acid,

ref: 10.1007/s00204-019-02537-z

None identified in dataset OR by in silico fragmentation.

### Aldosterone

Chromatogram showing prevalence in samples

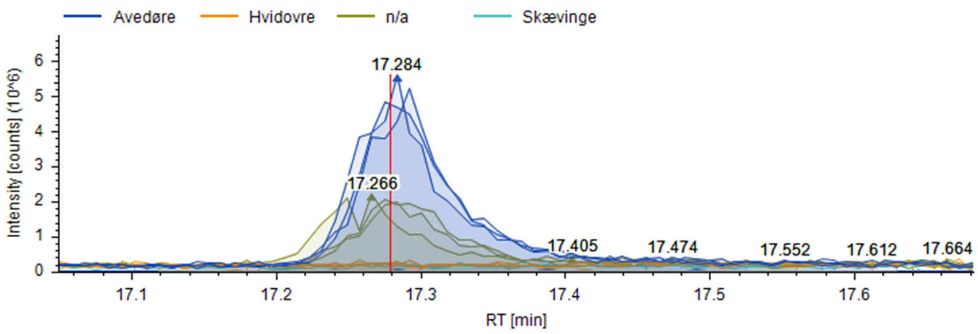


FISh coverage

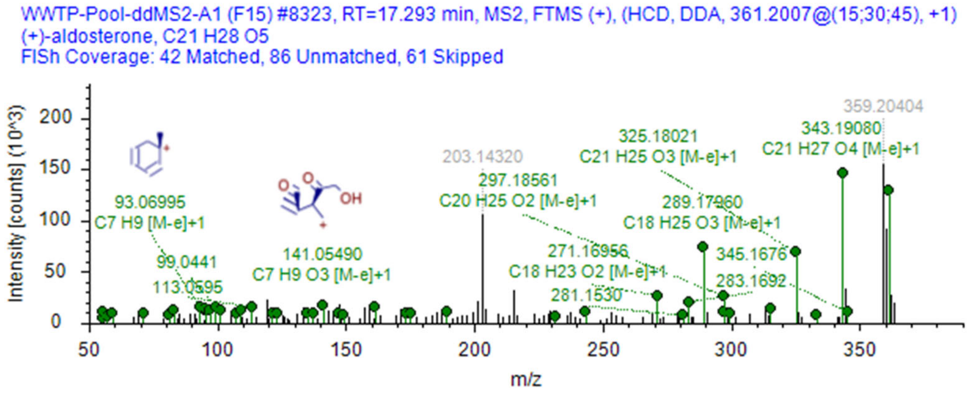


mzVault reference.

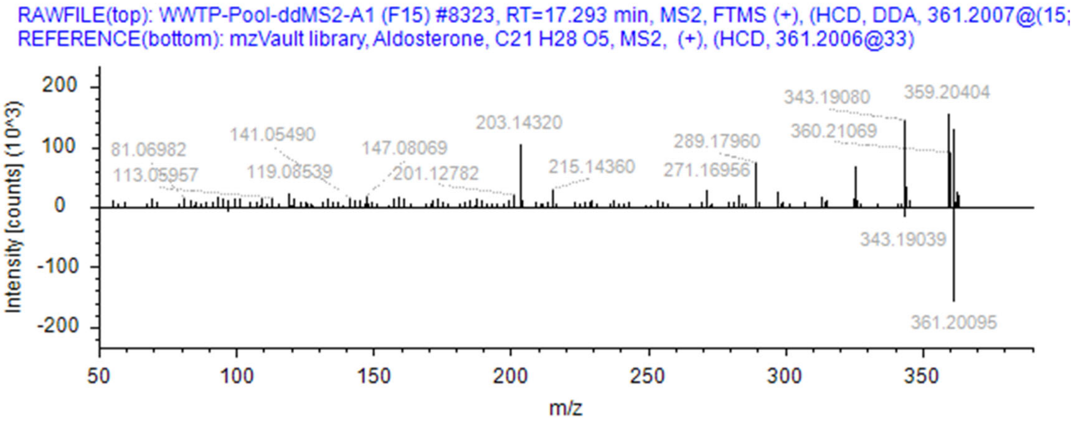

#### Metabolites:

5 alpha-dihydroaldosterone, 5 beta-dihydroaldosterone.

ref: https://doi.org/10.1152/ajprenal.1987.252.3.F365

None identified in dataset OR by in silico fragmentation. There is, however, a number of unidentified steroid metabolites in the dataset. While it would be outside the scope of this paper, a number of these may be identified as the above stated metabolites upon further analysis.

### Candesartan

Chromatogram showing prevalence in samples

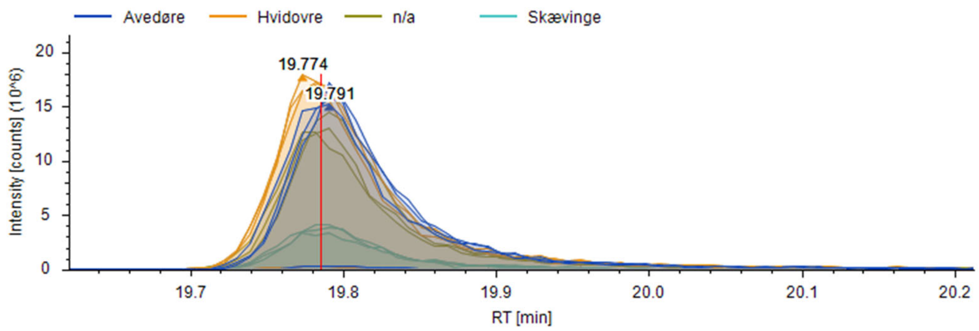


FISh coverage

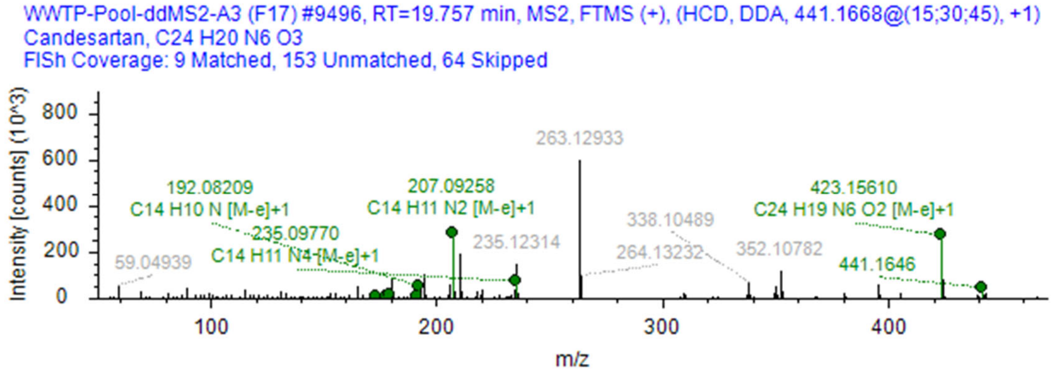


mzVault reference.

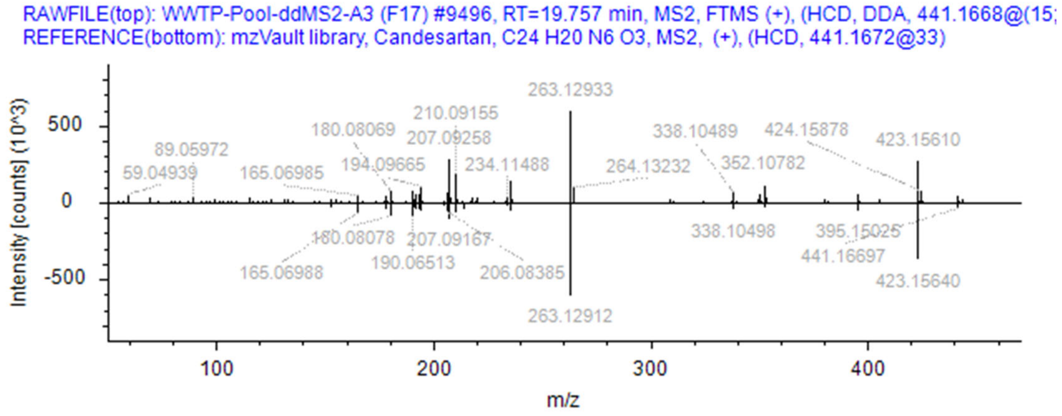

### Metabolites:

(2S,3S,4S,5R)-6-[2-Ethoxy-3-[[4-[2-(2H-tetrazol-5-yl)phenyl]phenyl]methyl]benzimidazole-4-carbonyl]oxy-3,4,5-trihydroxyoxane-2-carboxylic acid, $C_{30}H_{28}N_6O_9$;

3-[[4-[2-[2-[(3R,4S,5S,6S)-6-carboxy-3,4,5-trihydroxyoxan-2-yl]tetrazol-5-yl]phenyl]phenyl]methyl]-2-ethoxy-1H-benzimidazol-3-ium-4-carboxylic acid,$C_{30}H_{29}N_6O_9$,

ref: DOI:10.5281/zenodo.4056560

None identified in dataset OR by in silico fragmentation.

### Clarithromycin

Chromatogram showing prevalence in samples

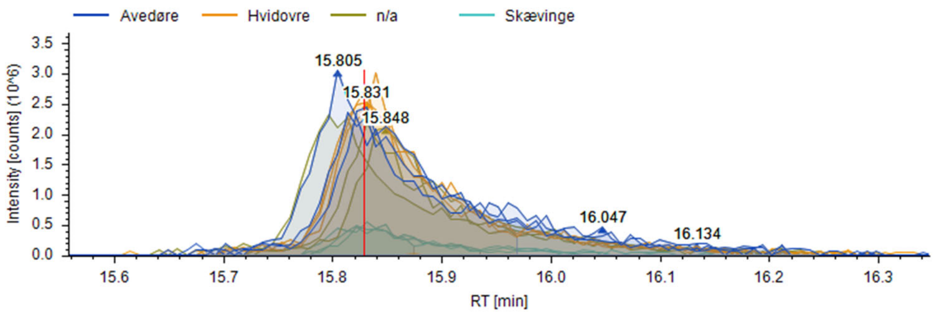


FISh coverage

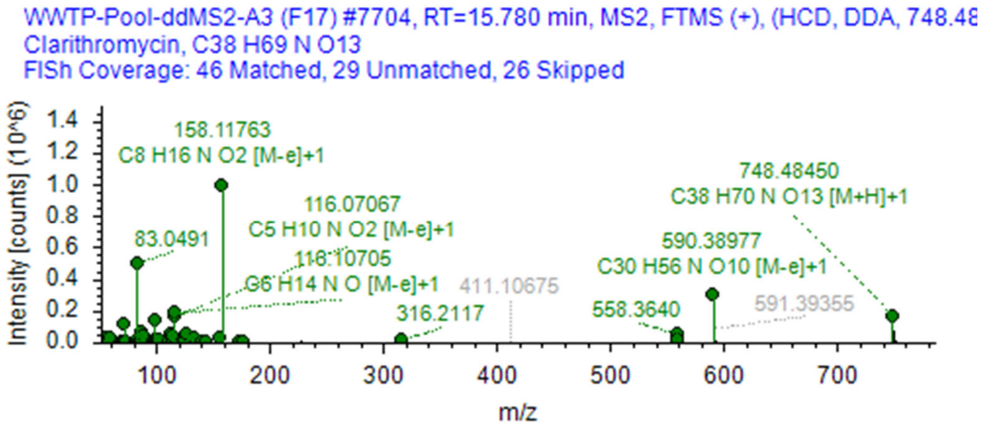


mzVault reference.

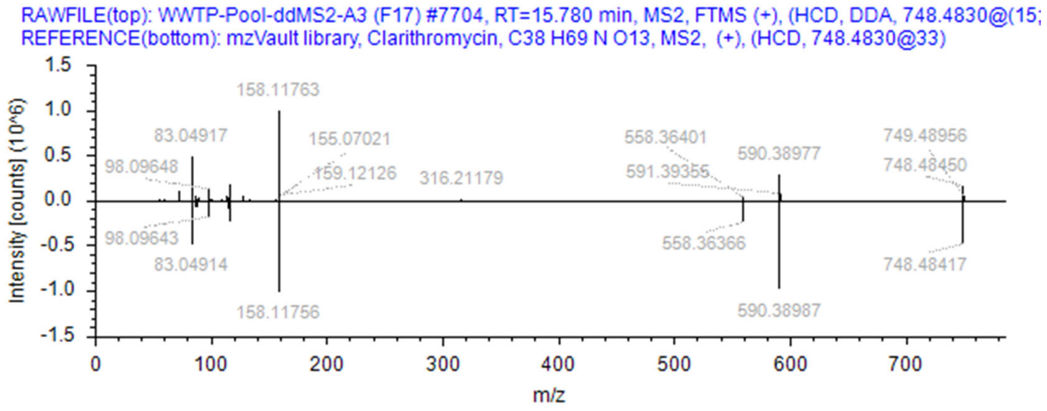

### Metabolites:

14-hydroxyclarithromycin, C38H69NO14, CAS 110671-78-8. Identified in dataset by in silico fragmentation. Pubchem ID 84020, Chemspider ID 75811, Metfrag hit 29/43 peaks, database: NormanSusDat_20Nov2019.
InChl key: BLPFDXNVUDZBII-KNPZYKNQSA-N
Ref: DOI: 10.2165/00003088-199937050-00003

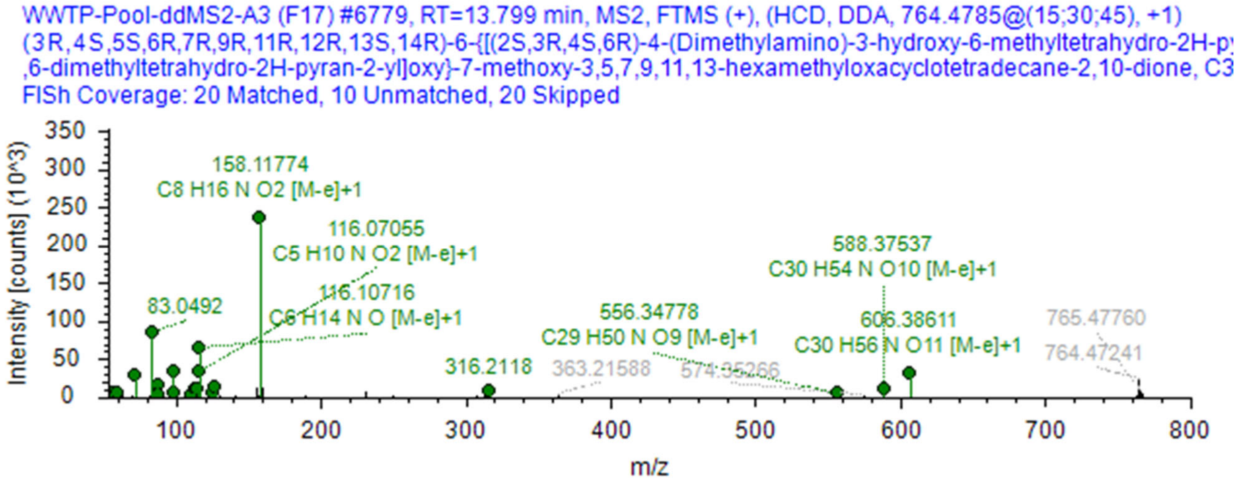

### DEET

Chromatogram showing prevalence in samples

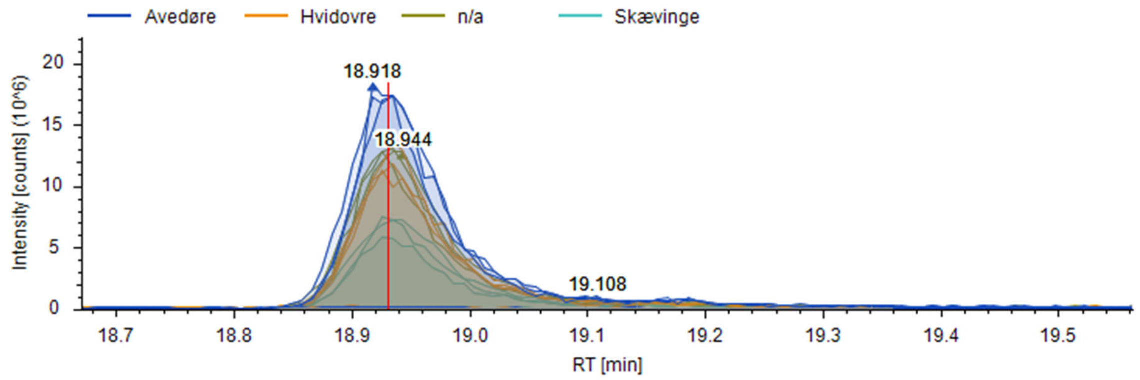


FISh coverage

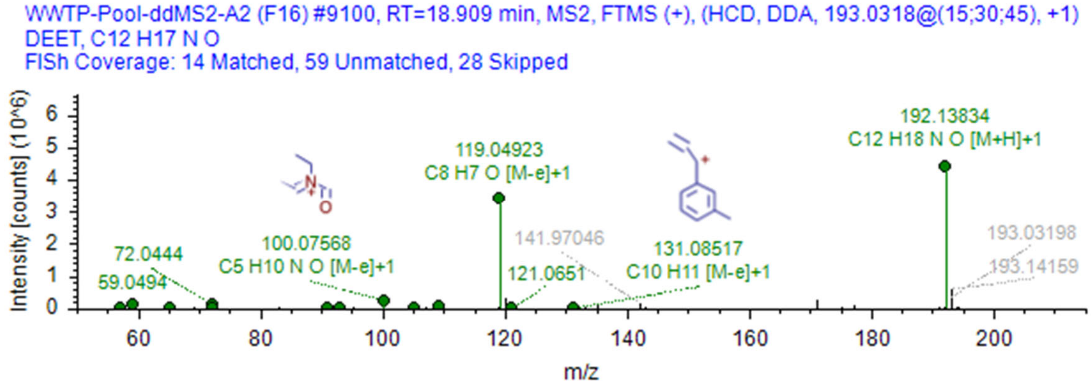


mzVault reference.

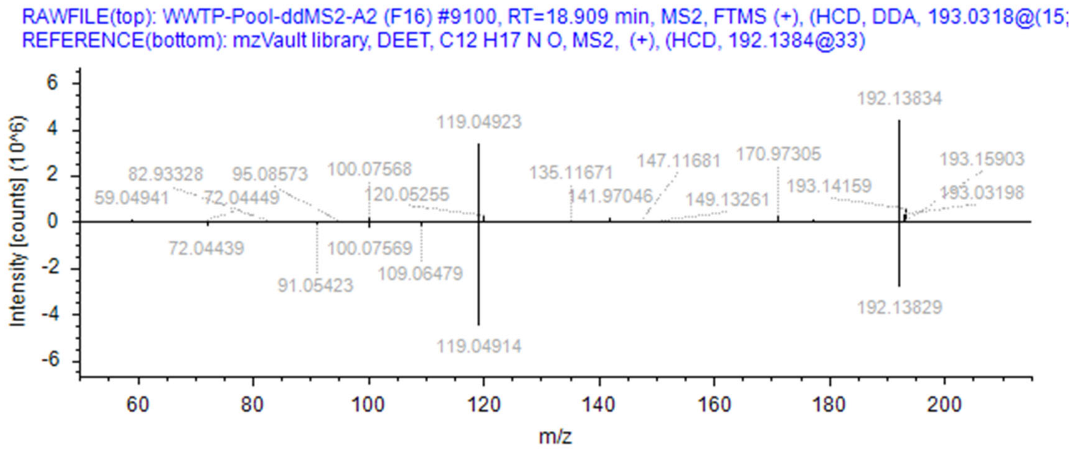

### Metabolites:

Identified via EnviPath.

N,N-Diethyl-m-toluamide Pathway:
m-Toluamide, $C_{10}H_{13}NO$; N,N-diethyl-3-(hydroxymethyl)benzamide, $C_{12}H_{17}NO_2$; N,N-diethyl-3-formylbenzamide, $C_{12}H_{15}NO_2$; 3-(diethylcarbamoyl)benzoate; m-Methylbenzoate; Diethylamine; Ethyl amine; Acetaldehyde

DEET Pathway:
DEET_TP_M222; DEET_TP_M164, $C_{10}H_{13}NO$.

None identified in dataset OR by in silico fragmentation.

### Diltiazem

Chromatogram showing prevalence in samples

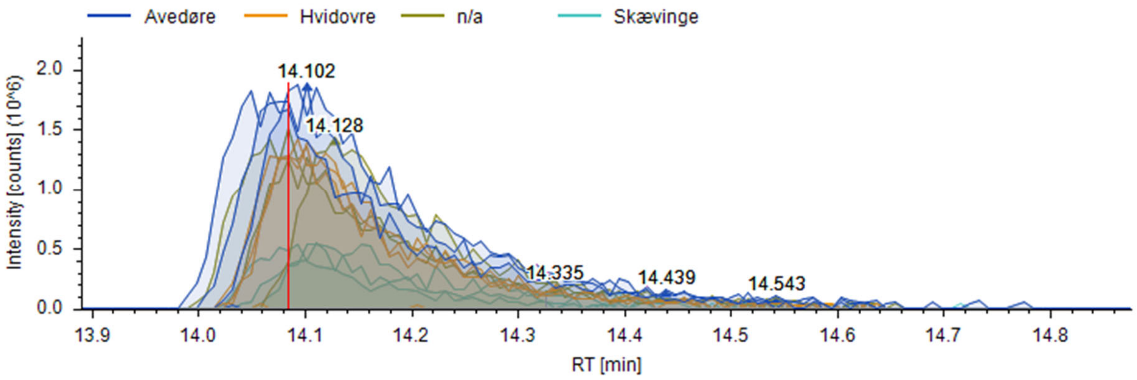


FISh coverage

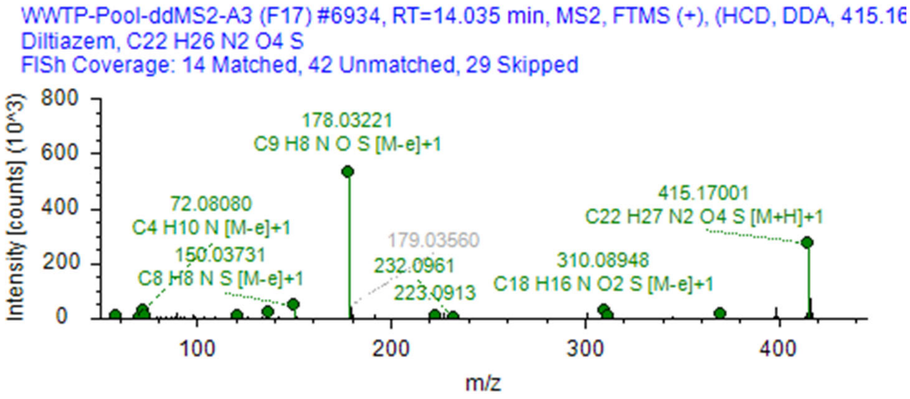


mzVault reference.

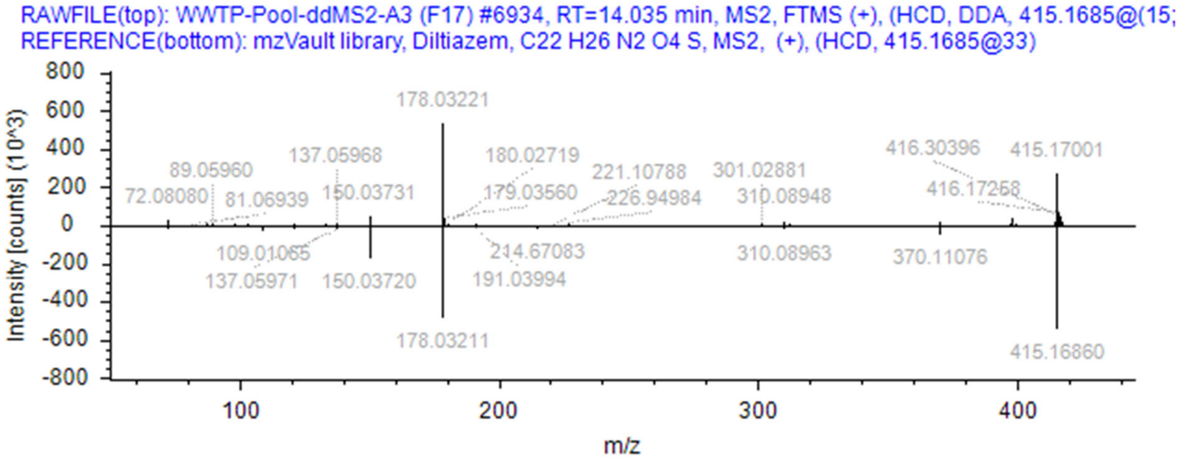

### Metabolites:

N-monodesmethyl diltilazem, C21H24N2O4S; deacetyl diltiazem , C20H24N2O3S; deacetyl N-monodesmethyl diltiazem, C19H22N2O3S; deacetyl diltiazem N-oxide, C20H24N2O4S; deacetyl O-desmethyl diltiazem, C19H22N2O3S; deacetyl N,O-didesmethyl diltiazem, C18H20N2O3S; O-desmethyl diltiazem, C21H25ClN2O4S.

Of these, two were identified in the dataset by in silico fragmentation.

N-monodesmethyl diltilazem, C21H24N2O4S. Pubchem ID 107891, Chemspider ID 97023, MetFrag hit 8/29 hits, database: NormanSusDat_20Nov2019. InChl key:YOMLDISQSWWYOT-UXHICEINSA-N

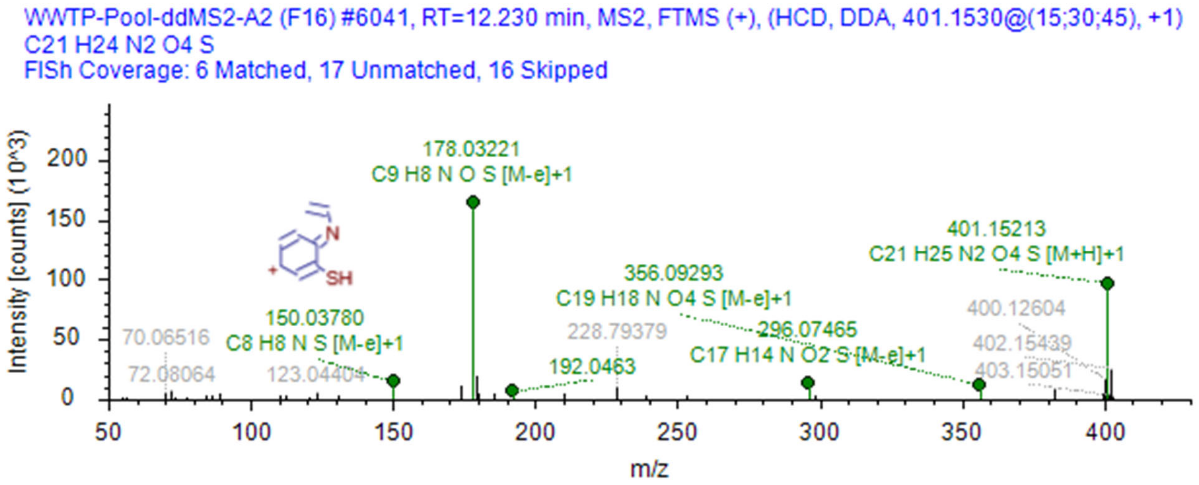


deacetyl diltiazem, C20H24N2O3S. Pubchem ID 91638, Chemspider ID 82743, MetFrag hit 5/53hits, database: NormanSusDat_20Nov2019. InChl key: NZHUXMZTSSZXSB-MOPGFXCFSA-N

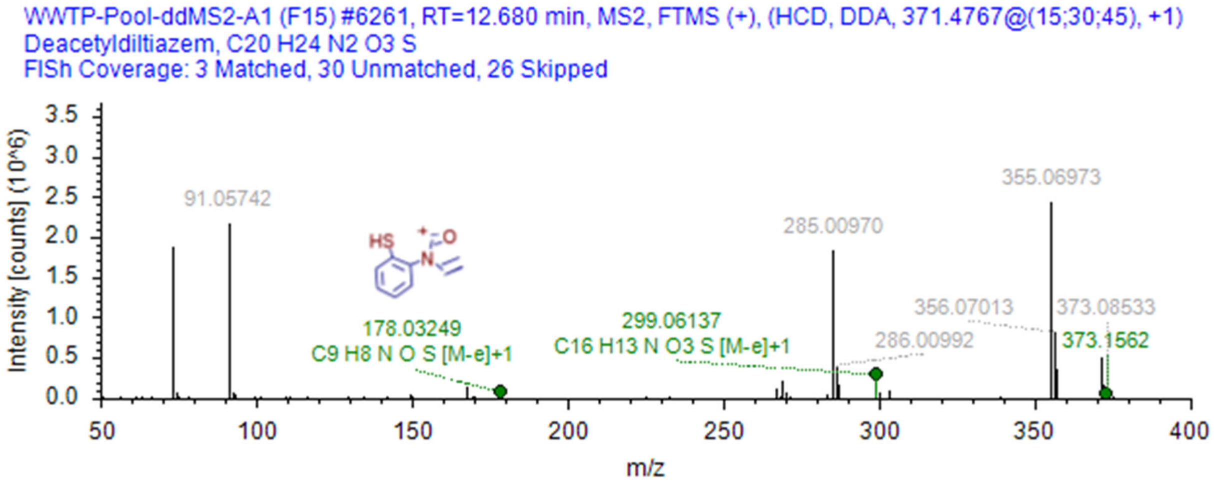


Refs: doi:10.1111/j.1365-2125.1987.tb03160.x; 10.1080/00498250110043517; *10.5281/zenodo.4056560*

### Fexofenadine

Chromatogram showing prevalence in samples

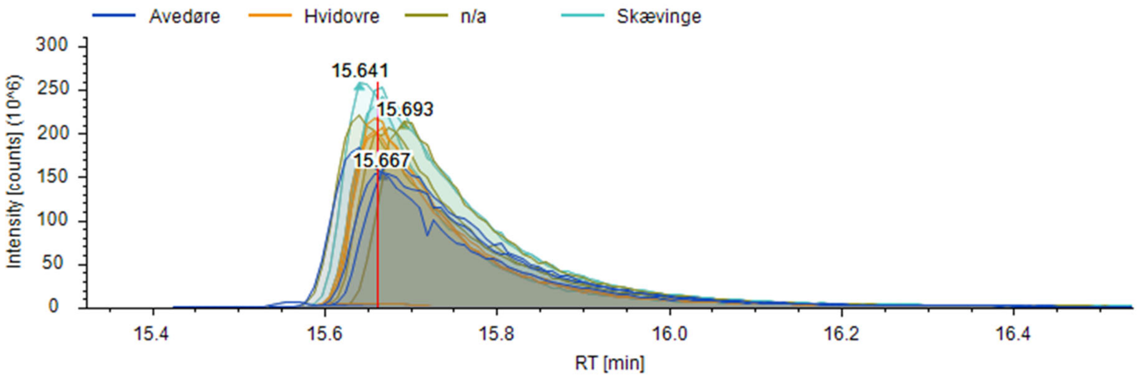


FISh coverage

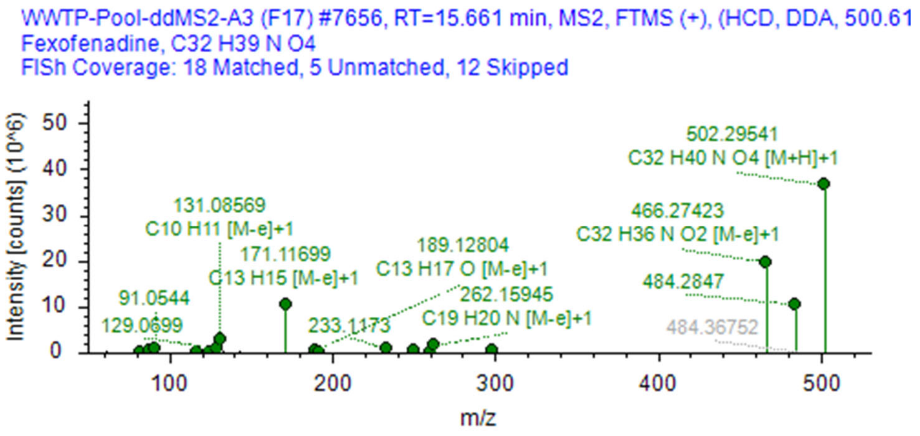


mzVault reference.

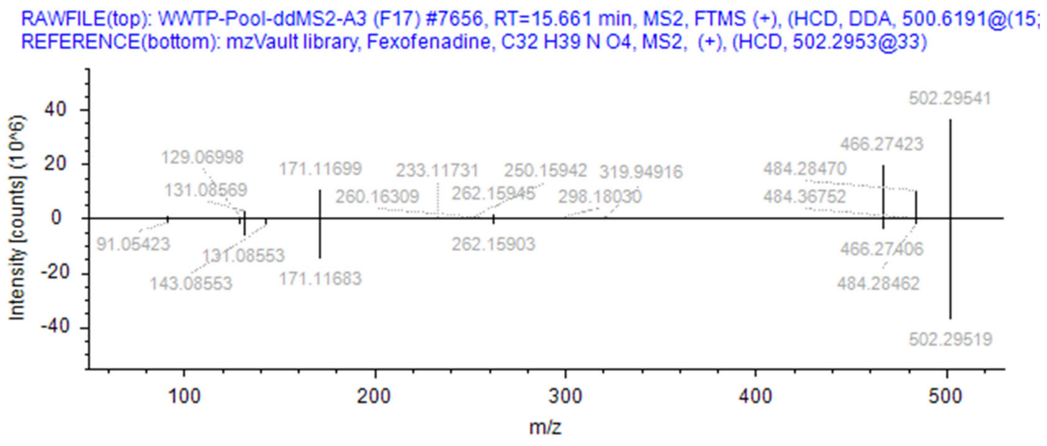

### Metabolites:

Identified via EnviPath. Identified in dataset by in silico fragmentation.

MDL 4829 (Azacyclonol), C18H21NO. Pubchem ID 15723, Chemspider ID 14952, MetFrag hit 13/64 hits, database: NormanSusDat_20Nov2019. InChl key: ZMISODWVFHHWNR-UHFFFAOYSA-N

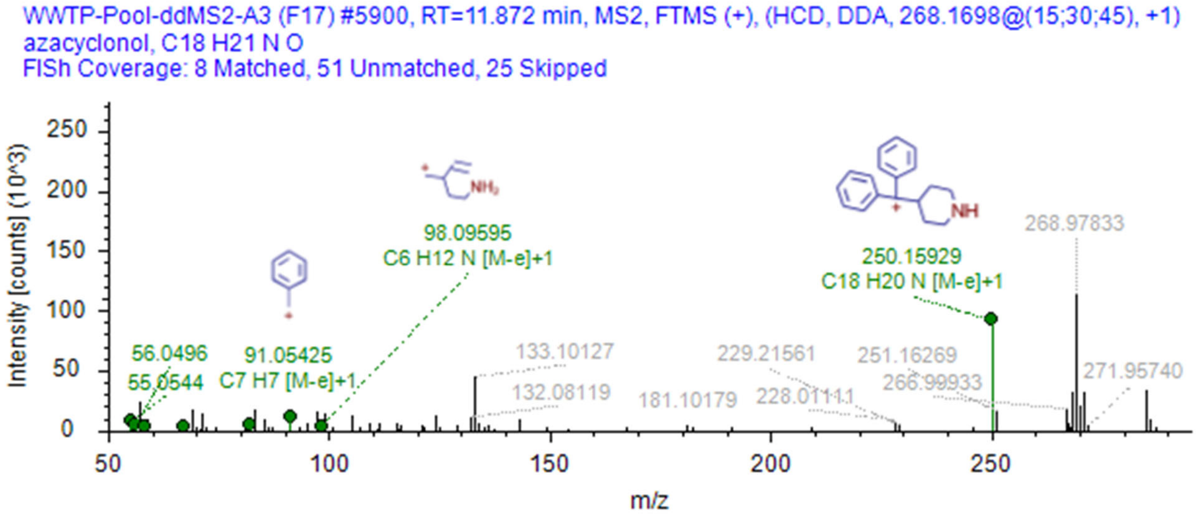


Fexofenadine Methyl Ester, C33H41NO4. Pubchem ID 9936344 , Chemspider ID 8111972, MetFrag hit 75/128 hits, database: NormanSusDat_20Nov2019. InChl key: GOUQSHOAAGQXNJ-UHFFFAOYSA-N

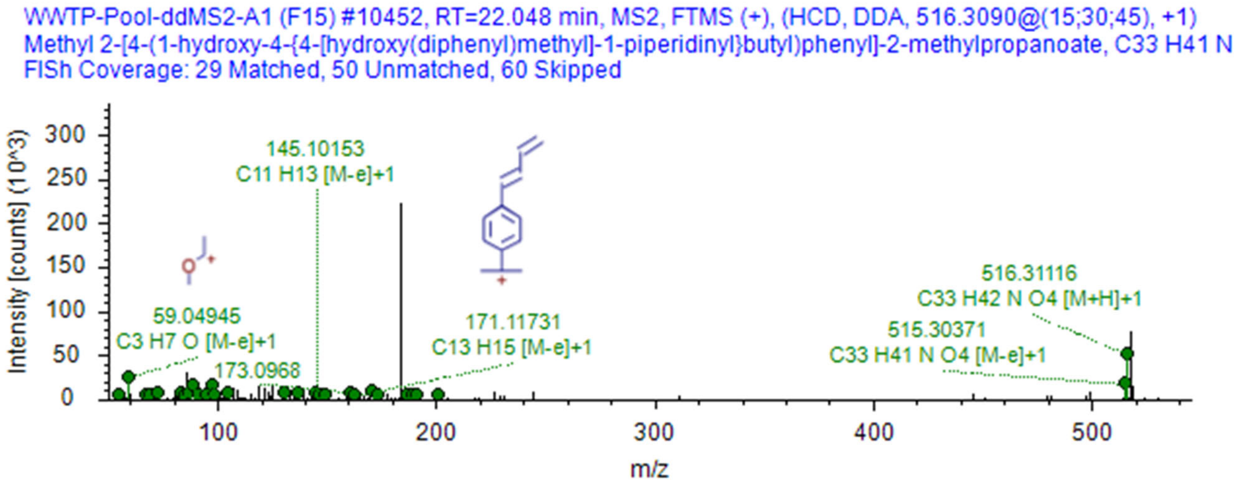

### Indomethacin

Chromatogram showing prevalence in samples

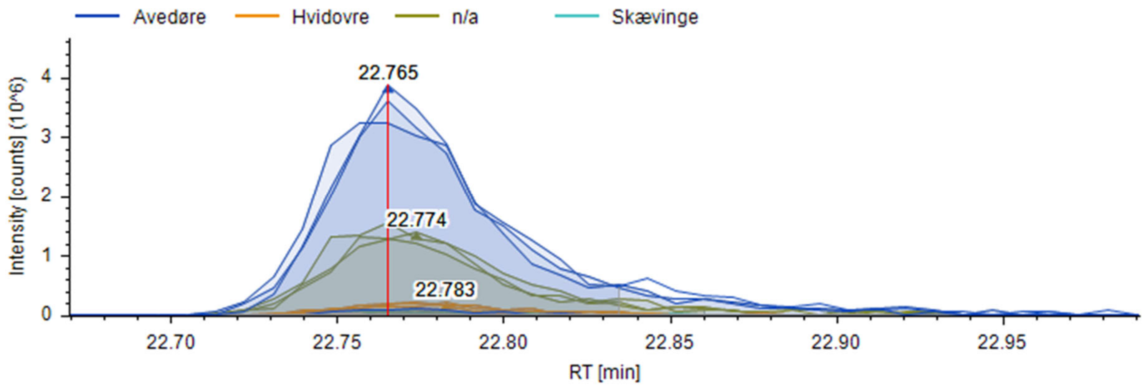


FISh coverage

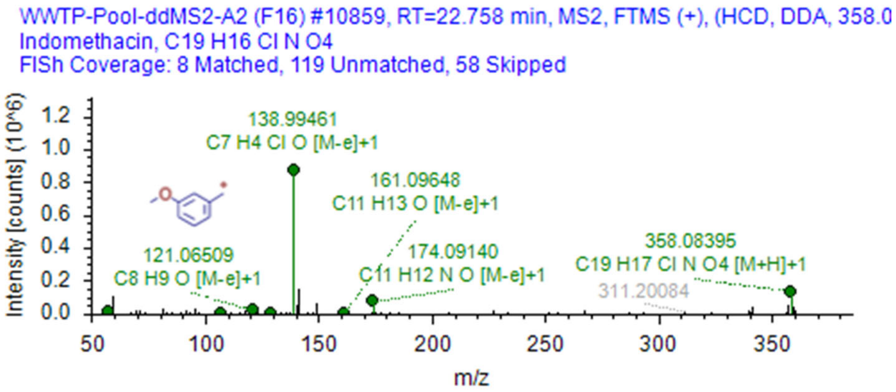


mzVault reference.

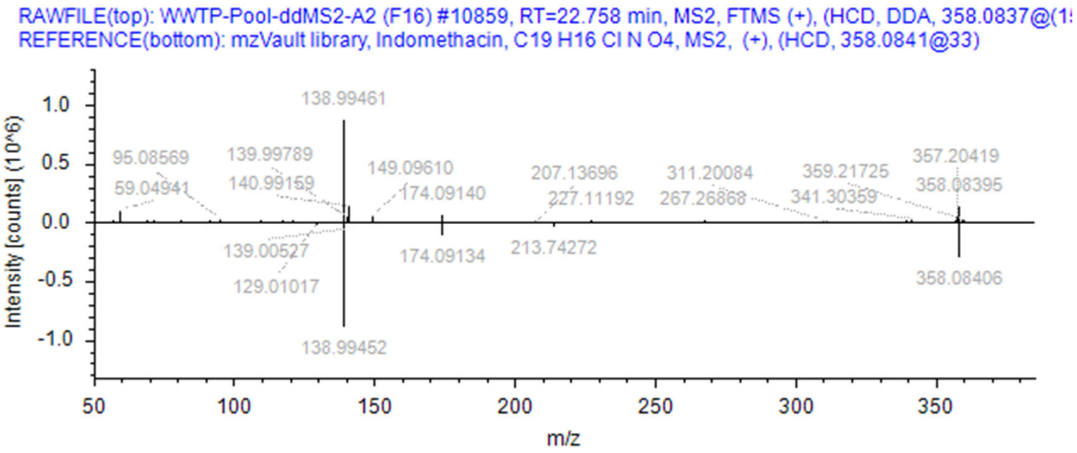

### Metabolites:

Identified via EnviPath.

O-desmethyl-indomethacin, $C_{18}H_{14}ClNO_4$; N-deschlorobenzoyl-indomethacin, $C_{12}H_{13}NO_3$; O-desmethyl-N-deschlorobenzoyl-indomethacin, $C_{11}H_{11}NO_3$; (2S,3S,4S,5R)-6-[2-[1-(4-chlorobenzoyl)-5-methoxy-2-methylindol-3-yl]acetyl]oxy-3,4,5-trihydroxyoxane-2-carboxylic acid, $C_{25}H_{24}ClNO_{10}$

None identified in dataset OR by in silico fragmentation.

### Oxcarbazepine

Chromatogram showing prevalence in samples

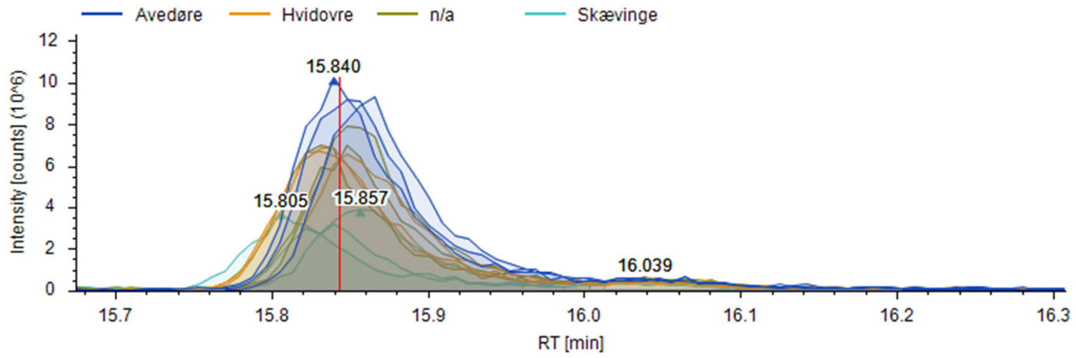


FISh coverage

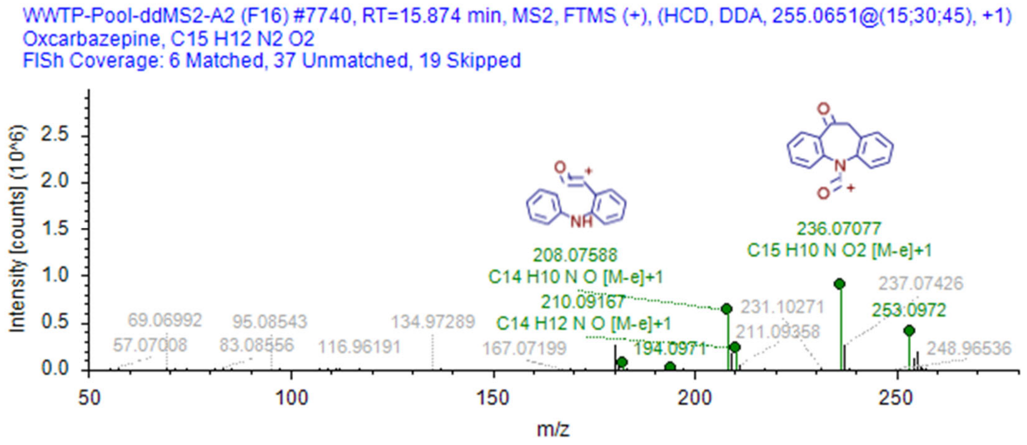


mzVault reference.

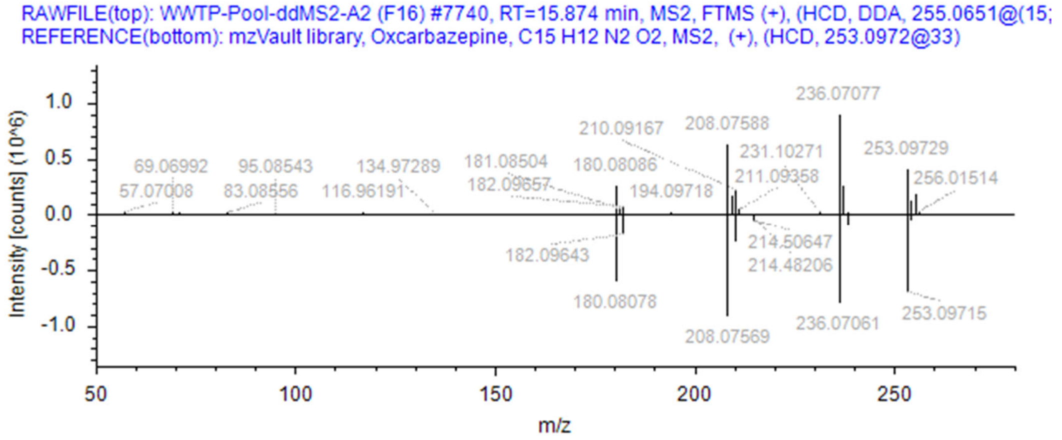

### Metabolites:

Identified via EnviPath. Identified in dataset by in silico fragmentation.

Licarbazepine, C15H14N2O2. Pubchem ID 114709, Chemspider ID 102704, MetFrag hit 22/99 hits, database: NormanSusDat_20Nov2019. InChl key: BMPDWHIDQYTSHX-UHFFFAOYSA-N

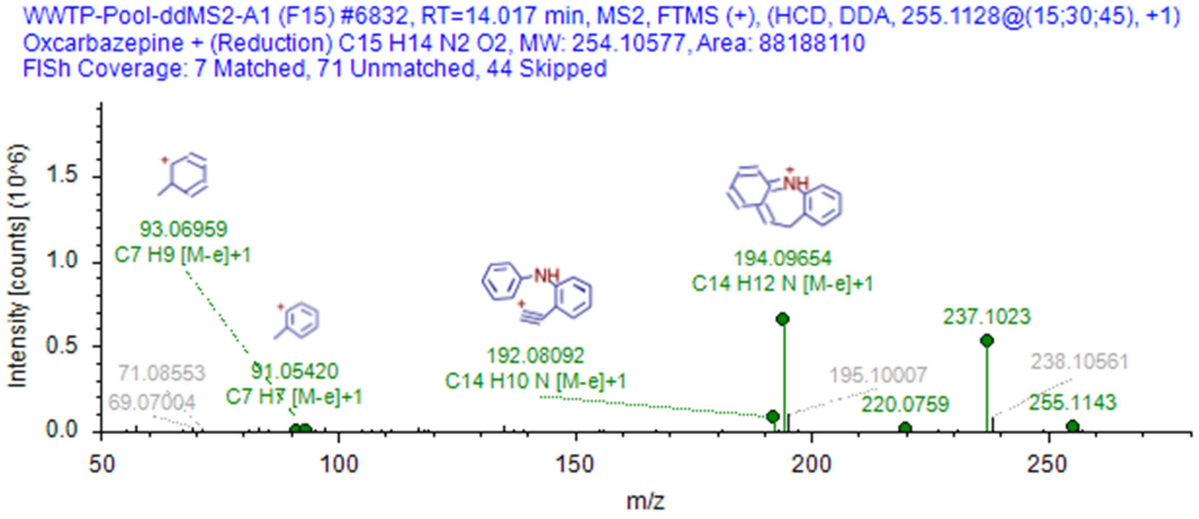


Dihydroxycarbazepine, C15H14N2O3. Pubchem ID 83852, Chemspider ID 102714, MetFrag hit 14/86 hits, database: NormanSusDat_20Nov2019. InChl key: PRGQOPPDPVELEG-UHFFFAOYSA-N

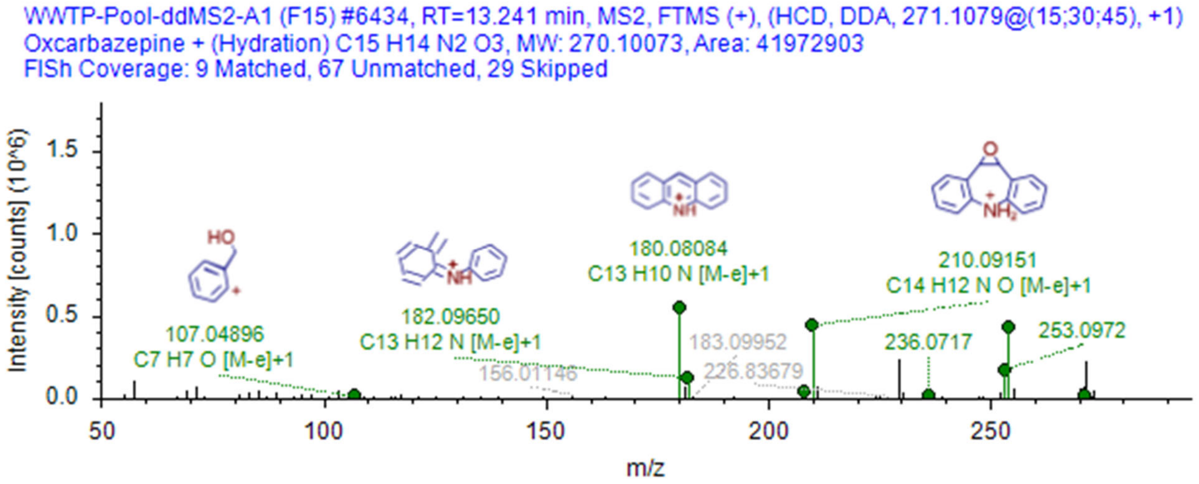

### Prednisone

Chromatogram showing prevalence in samples

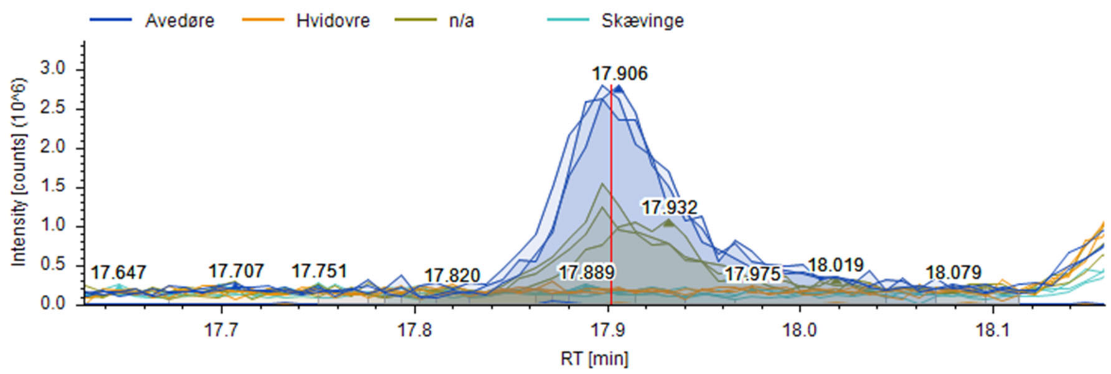


FISh coverage

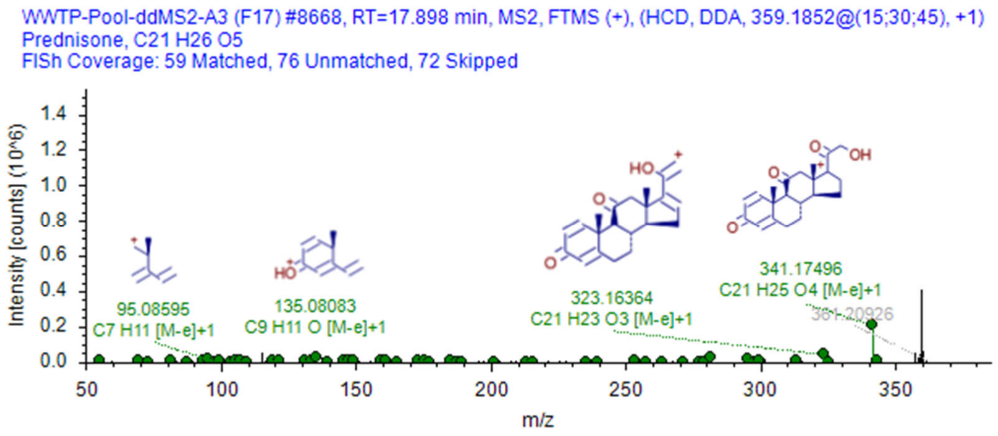


mzVault reference.

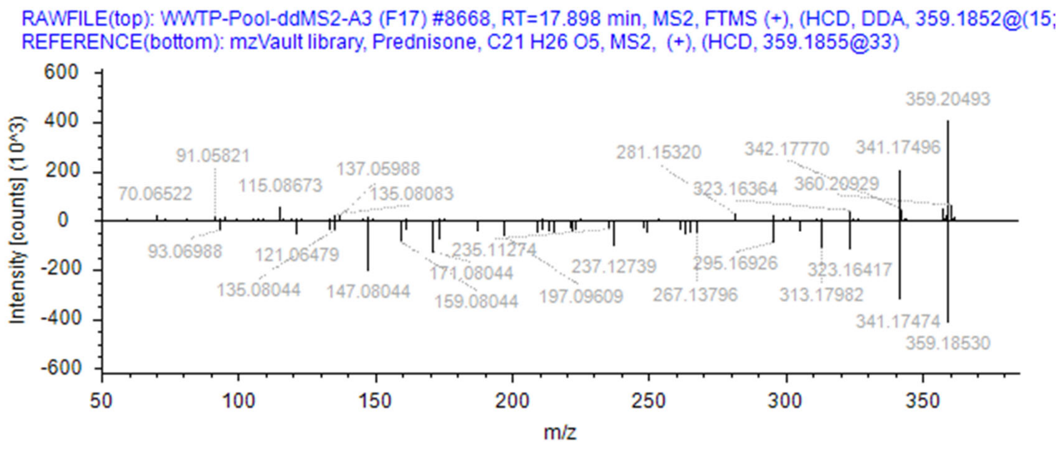

### Metabolites:

Identified via EnviPath.

17α,21-dihydroxy-pregnan-1,4,6-trien-3,11,30-trione, $C_{21}H_{24}O_5$; 20α-dihydro-prednisone, $C_{21}H_{28}O_5$; 6βhydroxy-prednisone, $C_{21}H_{26}O_6$; 6α-hydroxy-prednisone, $C_{21}H_{26}O_6$; 20β-dihydro-prednisone, $C_{21}H_{28}O_5$; 17α,20ξ,21-trihydroxy-5ξ-pregn-1-en-3,11-dione, $C_{21}H_{30}O_5$; Δ6-prednisolone, $C_{21}H_{26}O_5$; 20α-dihydro-prednisolone, $C_{21}H_{30}O_5$; 20β-dihydro-prednisolone, $C_{21}H_{30}O_5$; 6αhydroxy-prednisolone, $C_{21}H_{28}O_6$; 6βhydroxy-prednisolone, $C_{21}H_{28}O_6$; 6α,11β,17α,20β,21-pentahydroxypregnan-1,4-diene-3-one, $C_{21}H_{30}O_6$; 6β,11β,17α,20β,21-pentahydroxypregnan-1,4-diene-3-one , $C_{21}H_{30}O_6$; 6β,11β,17α,20α,21-pentahydroxypregnan-1,4-diene-3-one, $C_{21}H_{30}O_6$; 6β,11β,17α,21-tetrahydroxy-5ξ-pregn-1-en-3,20-dione, $C_{21}H_{30}O_6$; 6β,11β,17α,20β,21-pentahydroxy-5ξ-pregn-1-en-3-one, $C_{21}H_{32}O_6$; 6β,11β,17α,20α,21-pentahydroxy-5ξ-pregn-1-en-3-one, $C_{21}H_{32}O_6$.

Of these, only one (1) was identified in dataset by in silico fragmentation.

20α-dihydro-prednisolone, $C_{21}H_{30}O_5$. Pubchem ID 13962391, Chemspider ID 27524367, MetFrag hit 97/18086 hits, database: NormanSusDat_20Nov2019. InChl key: LCOVYWIXMAJCDS-LCGKLAOVSA-N

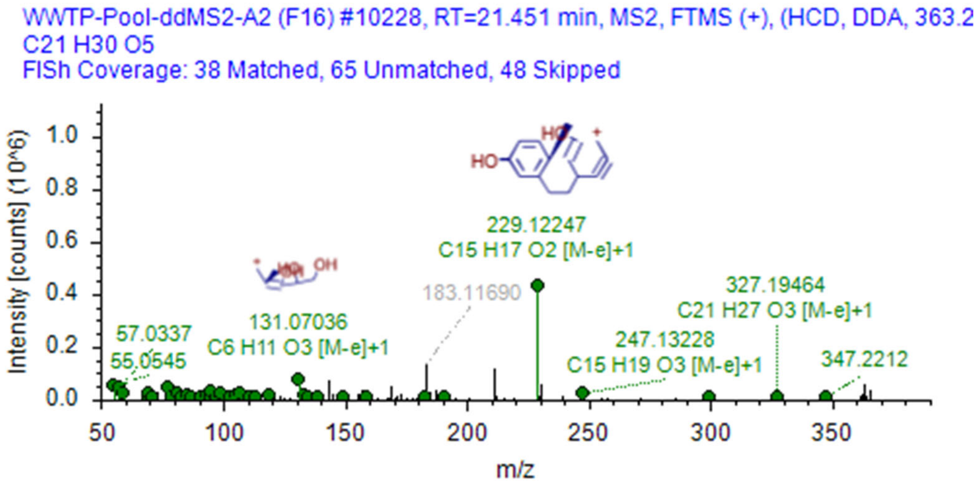

### Pregnenolone

Chromatogram showing prevalence in samples

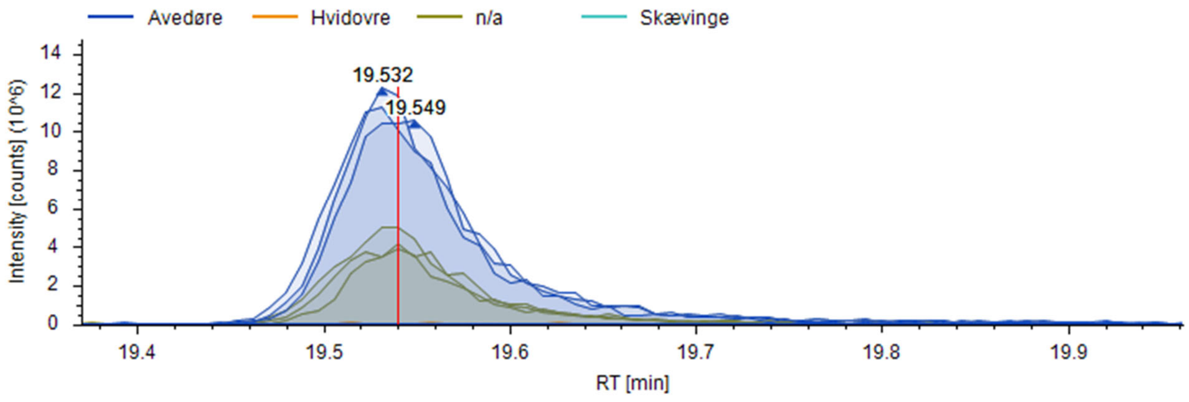


FISh coverage

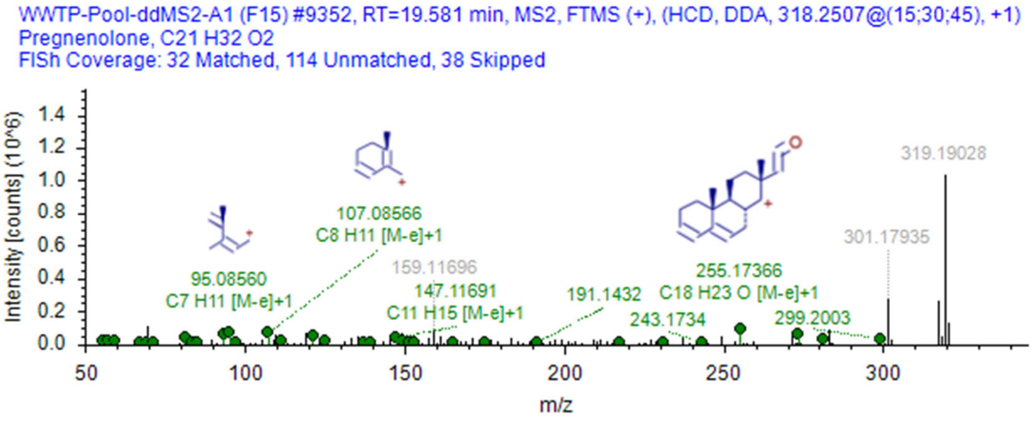


mzVault reference.

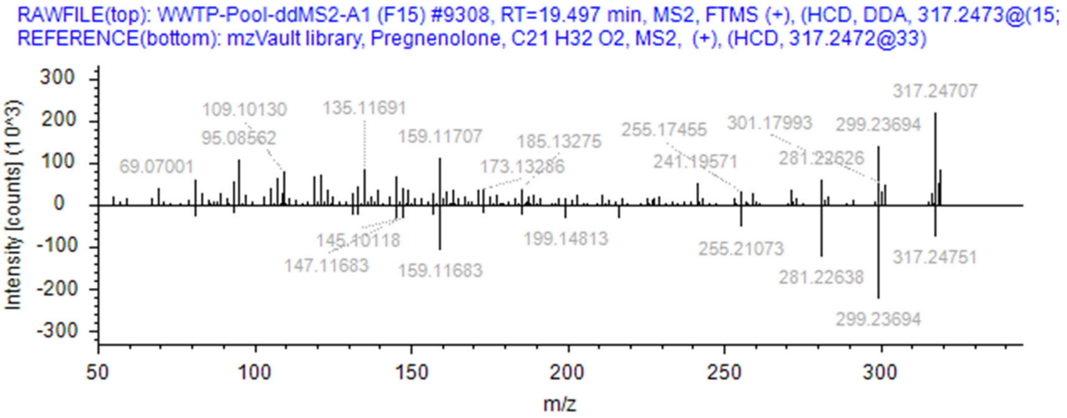

### Metabolites:

Identified via EnviPath.

1-(3,16-Dihydroxy-10,13-dimethyl-2,3,4,7,8,9,11,12,14,15,16,17-dodecahydro-1H-cyclopenta[a]phenanthren-17-yl)ethanone, $C_{21}H_{32}O_3$; 2-Hydroxy-1-(3-hydroxy-10,13-dimethyl-2,3,4,7,8,9,11,12,14,15,16,17-dodecahydro-1H-cyclopenta[a]phenanthren-17-yl)ethanone, $C_{21}H_{32}O_3$; Pregn-5-en-20-on-3b-yl sulfurate, $C_{21}H_{32}O_5S$.

One of these was identified in the dataset by in silico fragmentation.

Pregn-5-en-20-on-3b-yl sulfurate, $C_{21}H_{32}O_5S$. Pubchem ID 20845972, Chemspider ID 94802, MetFrag hit 64/173 hits, database: NormanSusDat_20Nov2019. InChl key:DIJBBUIOWGGQOP-OZIWPBGVSA-N

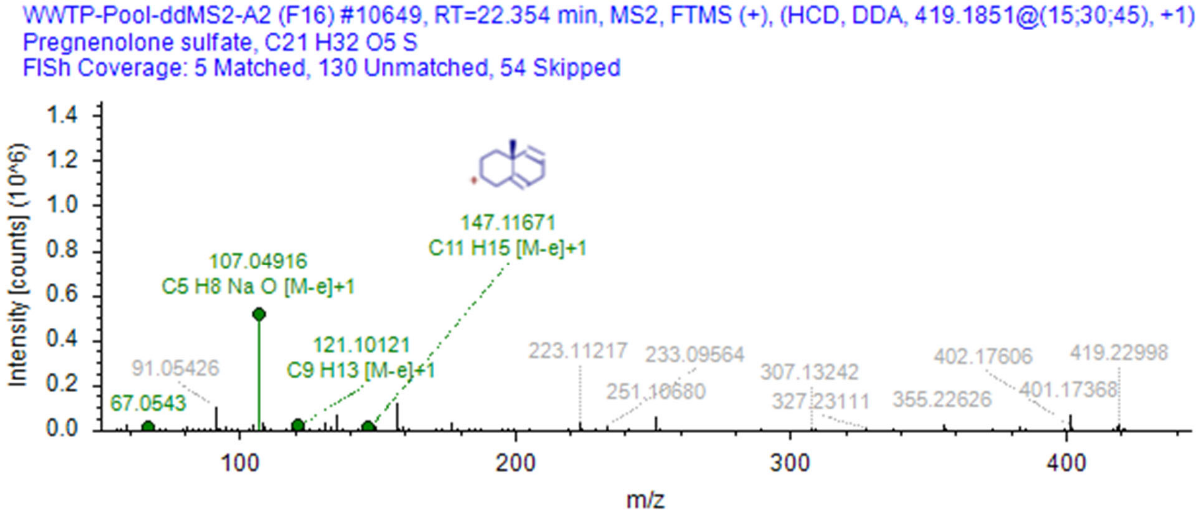

### Prometryn

Chromatogram showing prevalence in samples

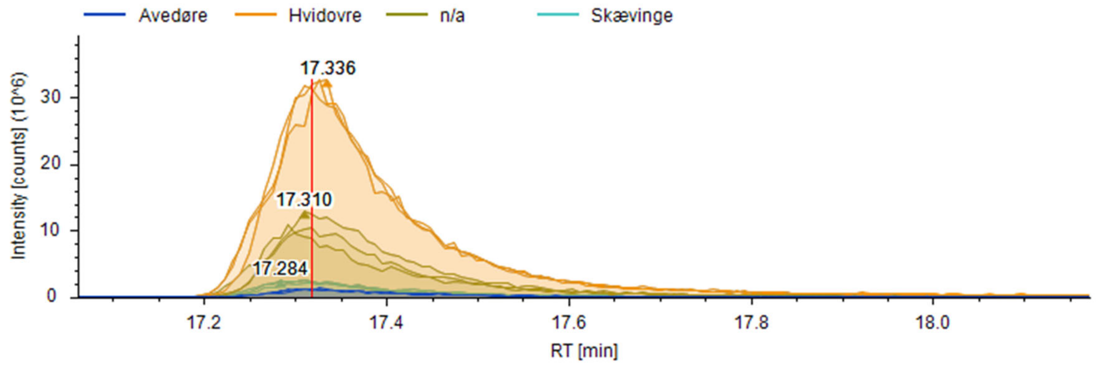


FISh coverage

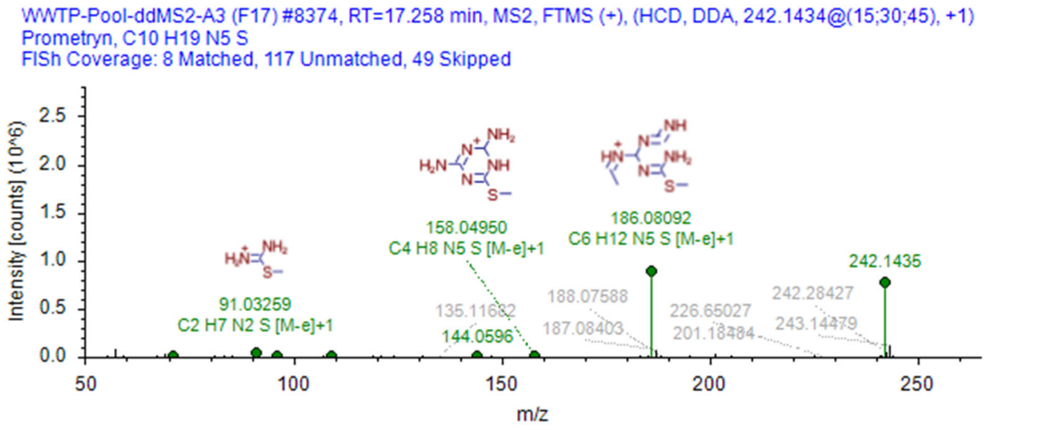


mzVault reference.

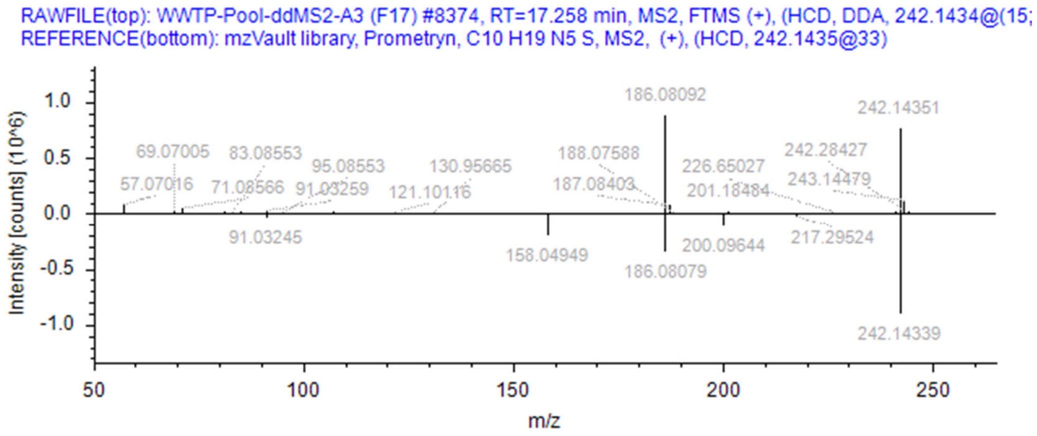

#### Metabolites:

2-mercapto-4-amino-6-isopropylamino-s-triazine; bis(2-amino-4-isopropylamino-s-triazinyl-6-yl)disulfide; 2-hydroxy-4-amino-6-isopropylamino-s-triazine

Ref: https://pubs.er.usgs.gov/publication/5230037

None identified in dataset OR by in silico fragmentation.

### Propiconazole

Chromatogram showing prevalence in samples

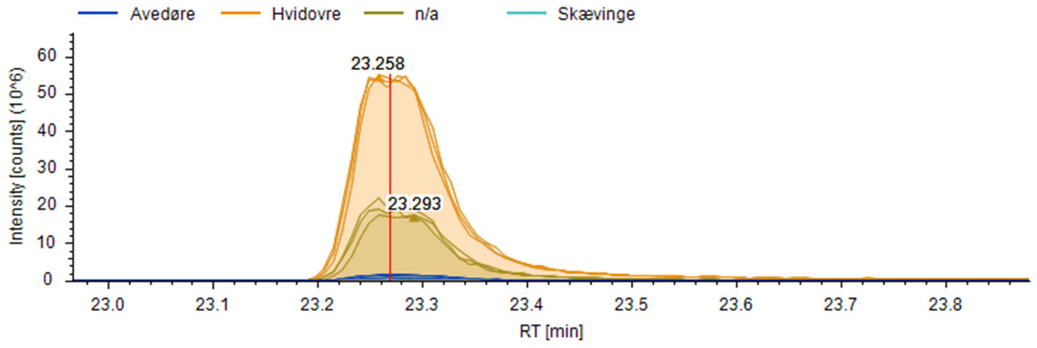


FISh coverage

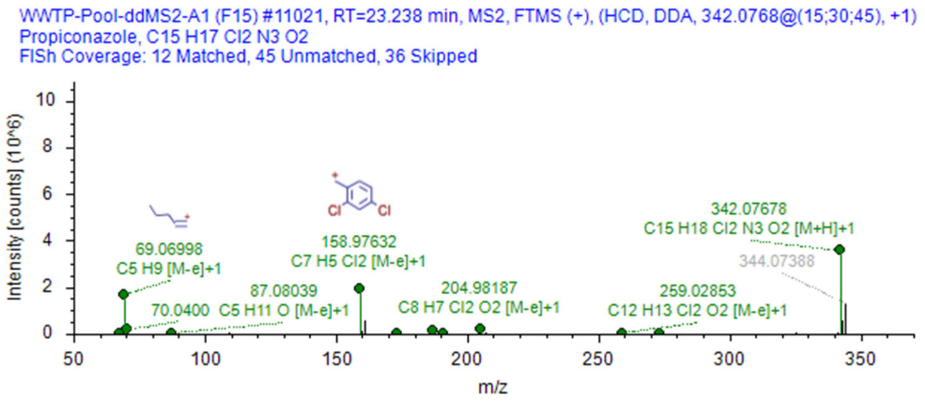


mzVault reference.

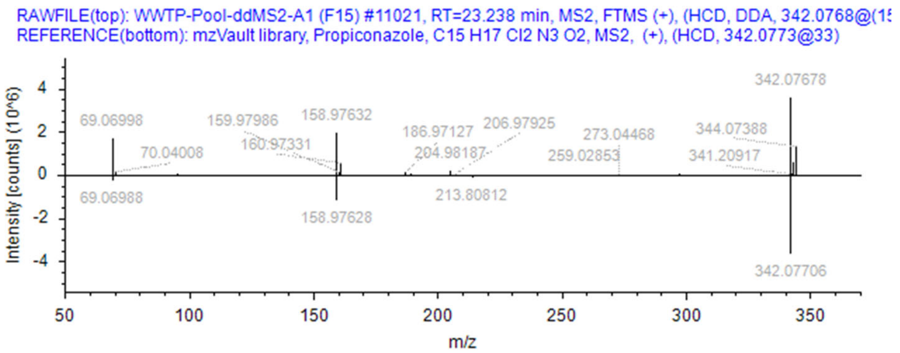


#### Metabolites:

None identified in dataset OR by in silico fragmentation.

### Propranolol

Chromatogram showing prevalence in samples

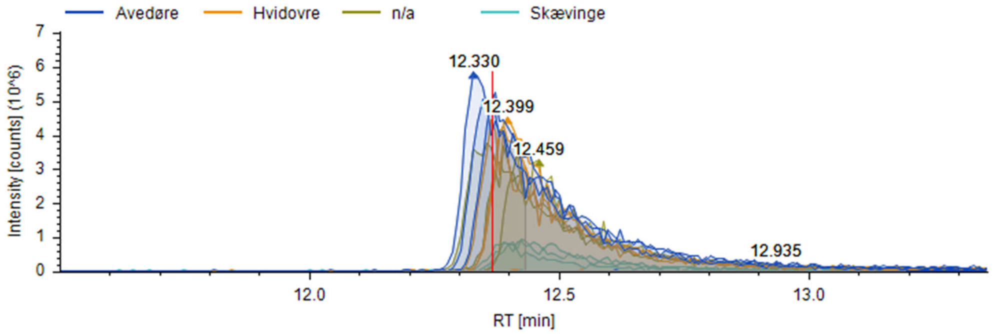


FISh coverage

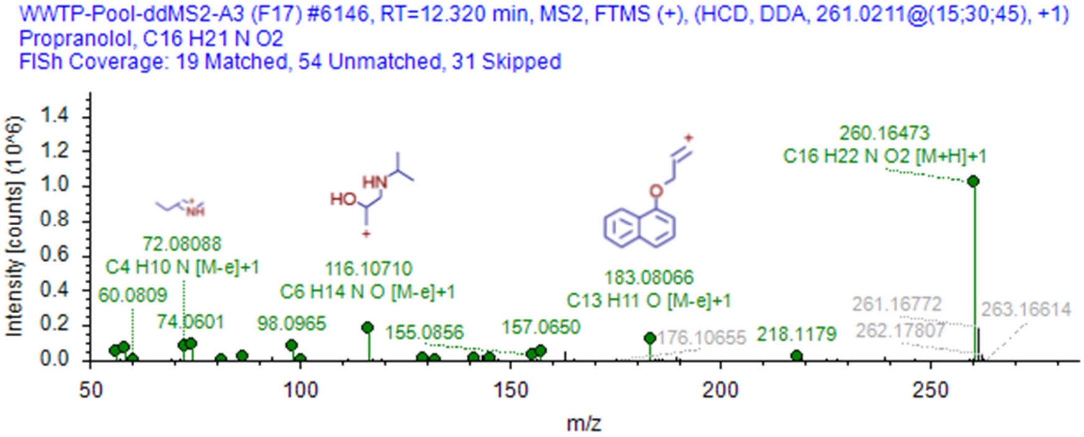


mzVault reference.

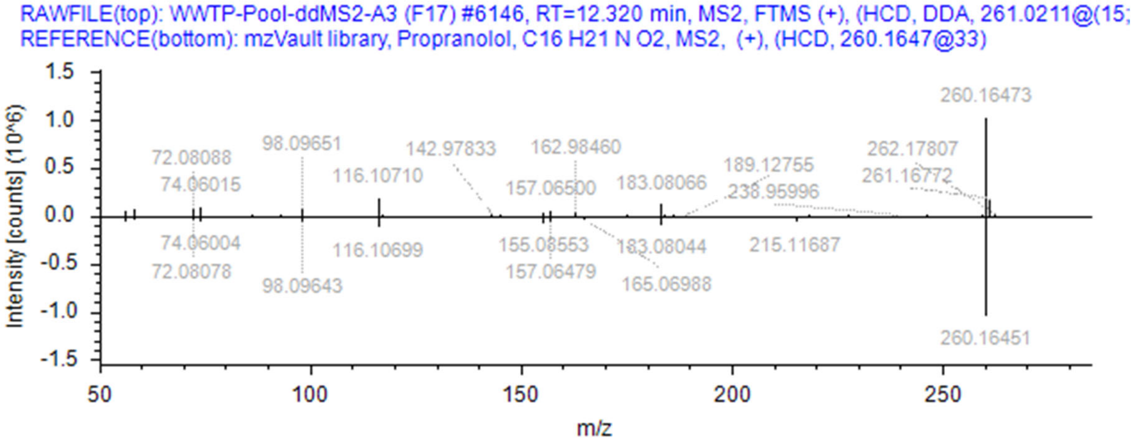

#### Metabolites:

α-naphthoxylactic acid, $C_{13}H_{12}O_4$; 4’-hydroxypropranolol, $C_{16}H_{21}NO_3$; propranolol glucuronide, $C_{22}H_{29}NO_8$; N-desisopropyl propranolol, $C_{13}H_{15}NO_2$; (2S,3S,4S,5R)-3,4,5-Trihydroxy-6-[1-naphthalen-1-yloxy-3-(propan-2-ylamino)propan-2-yl]oxyoxane-2-carboxylic acid, $C_{22}H_{29}NO_8$.

Ref: DOI:10.5281/zenodo.4056560; Drugbank acc.nr. DB00571

None identified in dataset OR by in silico fragmentation.

### Prosulfocarb

Chromatogram showing prevalence in samples

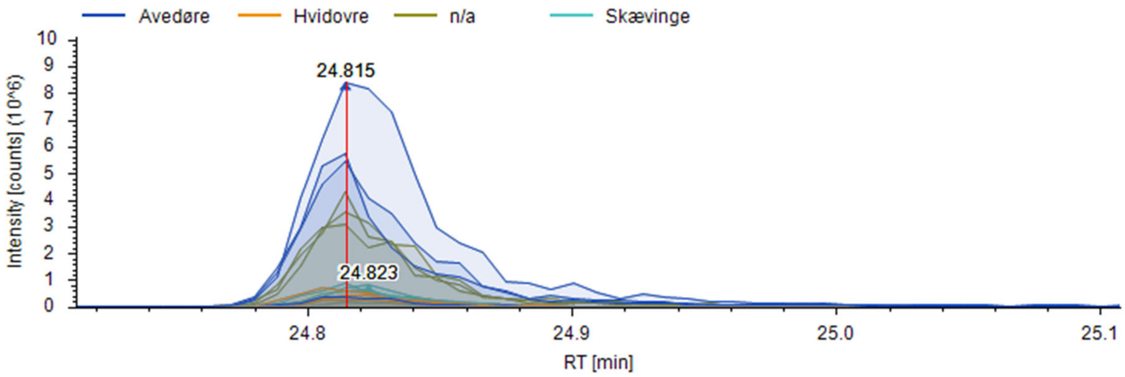


FISh coverage

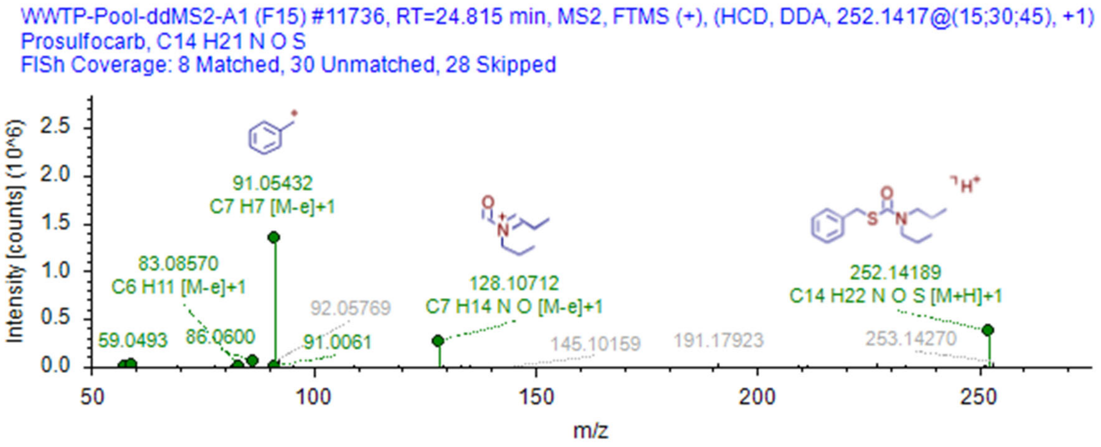


mzVault reference.

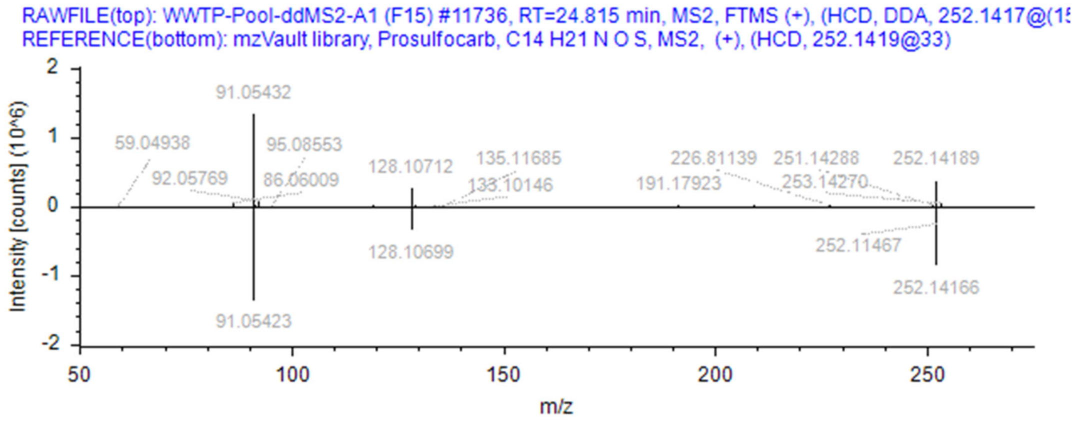

### Metabolites:

Identified via EnviPath. Identified in the dataset by in silico fragmentation.

Prosulfocarb sulfoxide, C14H21NO2S. Pubchem ID 57019542, Chemspider ID 71047475, MetFrag hit 10/202 hits, database: NormanSusDat_20Nov2019. InChl key:SRUUWJFBIOVZLU-UHFFFAOYSA-N

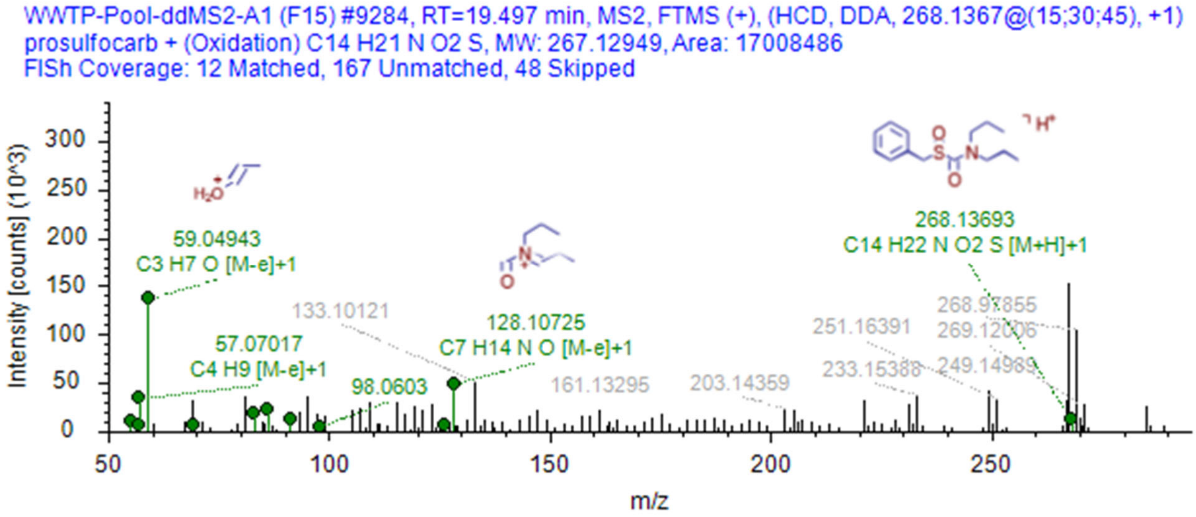

### Tributyl phosphate

Chromatogram showing prevalence in samples

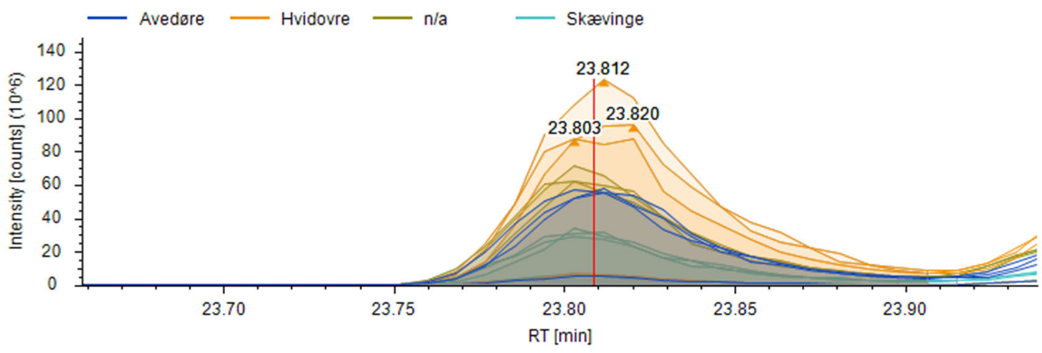


FISh coverage

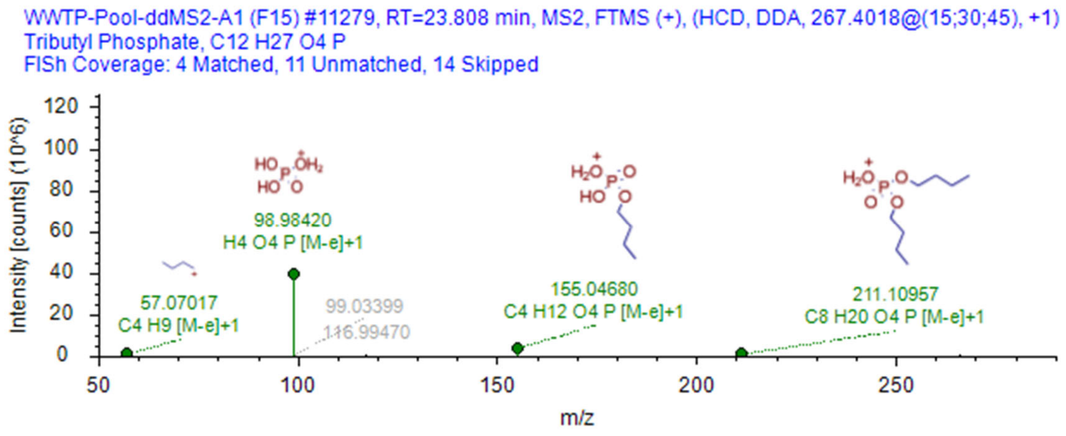


mzVault reference.

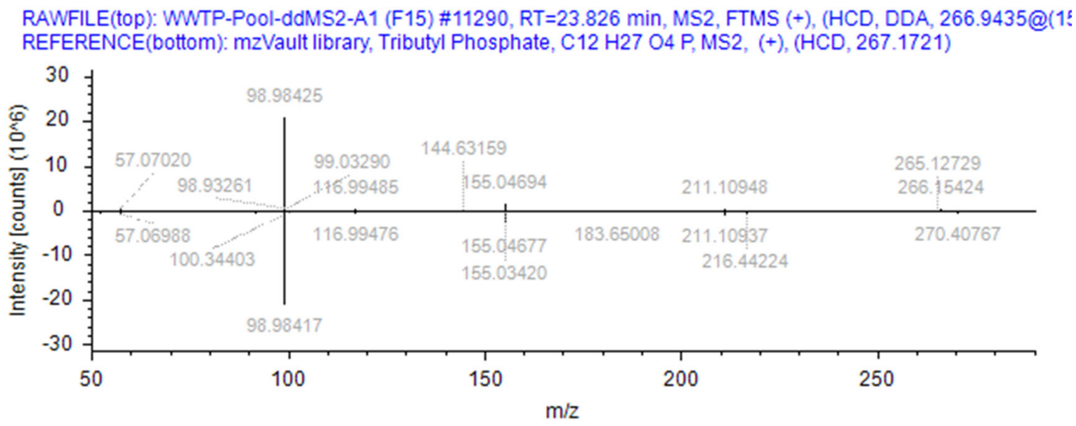

### Metabolites:

Identified via EnviPath.

Dibutyl phosphate; N-Acetyl-5-(3-hydroxybutyl)-L-cysteine; N-Acetyl-S-(3-oxobutyl)-L-cysteine

One of these was identified in the dataset by in silico fragmentation.

Dibutyl phosphate, C8H19O4P. Pubchem ID 7881, Chemspider ID 7593, MetFrag hit 6/114 hits, database: NormanSusDat_20Nov2019. InChl key:JYFHYPJRHGVZDY-UHFFFAOYSA-N

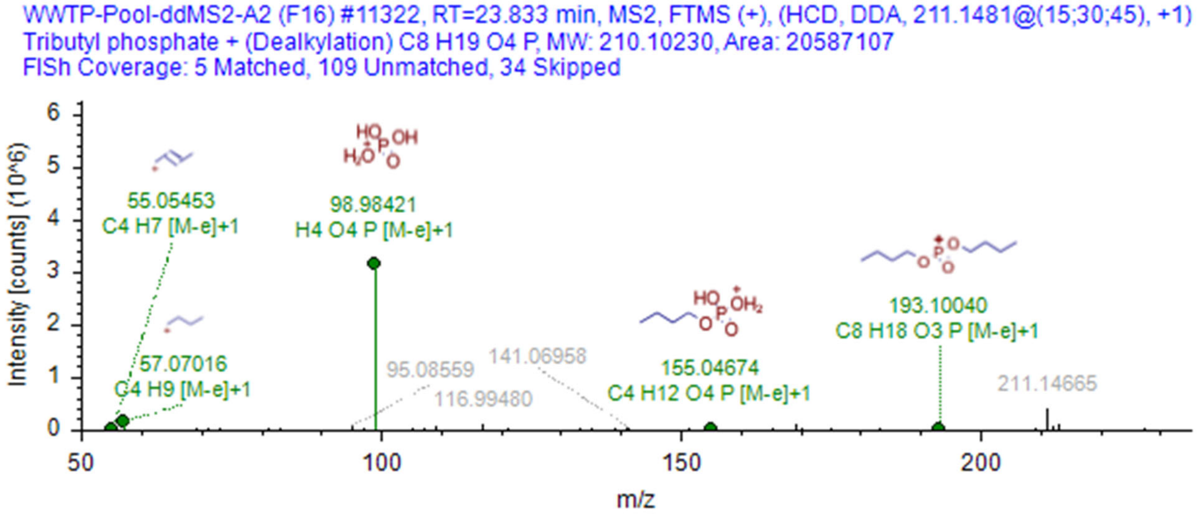

### Valsartan

Chromatogram showing Valsartan prevalence in samples

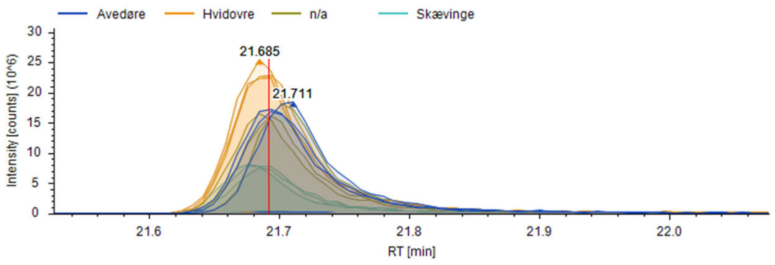


FISh coverage

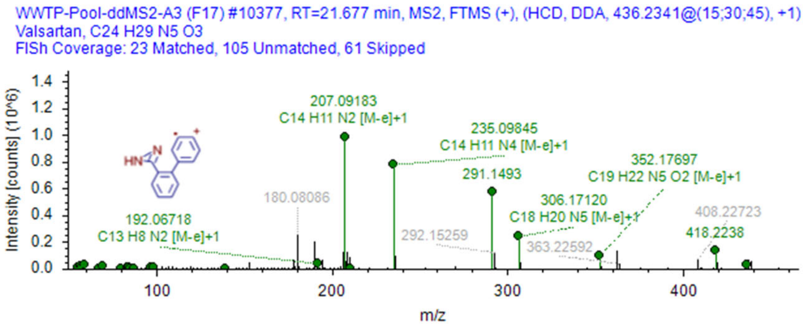


mzVault reference.

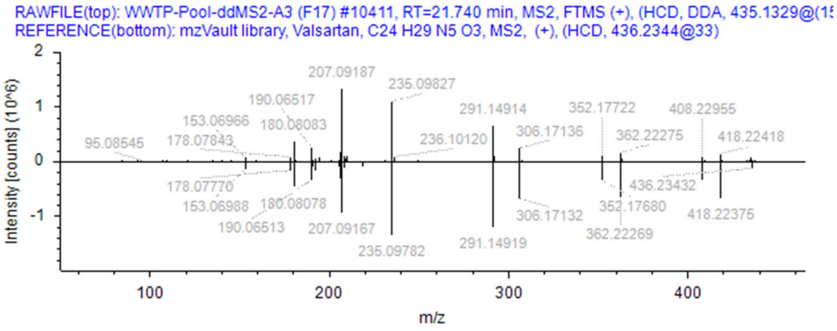


### Metabolites:

valeryl 4-hydroxy valsartan, $C_{29}H_{37}N_5O_5$.

Not identified in dataset OR by in silico fragmentation.
